# Charting the cognitive development of children using adult ‘polygenic g scores’

**DOI:** 10.64898/2025.12.19.695378

**Authors:** Yujing Lin, Robert Plomin

## Abstract

The most highly predictive polygenic scores in the behavioural sciences are for cognitive traits, especially general cognitive ability (g) and educational attainment. We combined polygenic scores derived from genome-wide association studies of adult g and educational attainment to create adult ‘polygenic g scores’ which we used to chart the course of cognitive development of 10,000 white British children from toddlerhood through early adulthood.

We integrated cross-sectional regression, latent growth curve, and confirmatory factor analysis to systematically characterise cognitive development. Polygenic g score showed minimal prediction in toddlerhood, modest prediction in childhood, and substantial prediction by early adulthood accounting for 12% of the variance. Higher polygenic g scores were associated with faster cognitive growth in latent growth models. Prediction was strongest for a cross-time latent cognitive factor (15%) capturing cognitive ability across development. By integrating polygenic prediction directly into a structural equation model framework, we provided a theoretical upper bound of genetic influences on g under minimal measurement error. We also examined the polygenic g score’s prediction of educational achievement, behaviour problems, and anthropometric outcomes and found similar developmental increases in prediction for educational achievement.

Together, our findings demonstrate that adult polygenic g scores can be a useful tool for charting the development of cognitive traits.

---

One of the most important outcomes of the DNA revolution for research on intelligence and cognitive abilities is the polygenic score (Plomin, 2019). Genome-wide association (GWA) studies correlate DNA differences, typically common genetic variants called single nucleotide polymorphisms (SNPs), with cognitive abilities. Many such genetic variants contribute to the substantial heritability of cognitive abilities, and the largest effect sizes are minuscule, accounting for less than 0.05% of the phenotypic variance (Plomin & von Stumm, 2018; Visscher et al., 2021). The small effect sizes make it difficult to trace pathways from genes to brain to behaviour. In contrast, polygenic scores sidestep the need to map these pathways. By aggregating thousands of these small but additive effects, polygenic scores translate GWA discoveries directly into a genetic predictor of individual differences among unrelated individuals in the population.

Despite the availability of many polygenic scores for cognitive outcomes to date, two scores consistently drive the strongest predictions (Procopio et al., 2025). The first is a polygenic score for intelligence. One of the largest GWA analyses of adult intelligence (N = ∼270,000) produced the IQ, or general cognitive ability (g), polygenic score (g-PGS), which predicts about 5% of the variance of adult intelligence in independent samples (Savage et al., 2018). GWA analyses require large sample sizes to detect the small effects of DNA variants. Conducting a GWA study (GWAS) of intelligence requires not only obtaining and genotyping DNA but also testing for intelligence, which is difficult with such large samples. For this reason, years of schooling (educational attainment) has been used as a proxy for intelligence (Rietveld et al., 2014). Educational attainment correlates strongly with intelligence, about 0.50 phenotypically (Deary, 2012) and 0.75 genetically (Hill et al., 2019). It is assessed with a single self-reported item about the highest level of education and, crucially, is routinely collected in most GWA studies as a demographic descriptor, making it possible to assemble huge samples for meta-analysis. The largest GWA analysis of educational attainment had a sample size of three million that yielded a polygenic score (EA-PGS) predicting up to 9% of the variance of intelligence, with the strongest prediction observed for verbal ability (Okbay et al., 2022). EA-PGS predicts more of the variance of intelligence than g-PGS, even though educational attainment is a coarse proxy for intelligence, because the EA-PGS sample size is more than ten times greater than the g-PGS sample size. Here, we combined g-PGS and EA-PGS to increase the prediction of adult g and refer to this composite as an adult ‘polygenic g score’.

Because inherited DNA differences are fixed at conception, polygenic scores remain constant during development, which is unique in developmental research. Although the genetic architecture of cognitive abilities exhibits some age-dependent effects, substantial genetic continuity across the lifespan has been well-documented (Hill et al., 2016; Savage et al., 2018). Since an adult-derived polygenic g score can theoretically predict adult g from birth just as well as in adulthood, we can leverage this genetic continuity in the opposite direction: using the adult polygenic g score as a tool to chart the emergence, growth, and changes of cognitive abilities across development and to pinpoint the earliest cognitive traits that are captured by adult-derived genetic predictors. To achieve this, we need longitudinal cognitive data that follow children from toddlerhood to adulthood.

Such longitudinal research takes three decades, and as a result, few data exist. Meta-analytic evidence indicates that while cognitive ability tends to fluctuate in toddlerhood, its stability increases rapidly across childhood, becoming highly stable by adolescence (Breit et al., 2024). Whereas these longitudinal phenotypic correlations represent a ceiling for predicting adult g from infant measures, adult polygenic g scores can be used as a sharper scalpel for dissecting facets of early cognitive development associated *genetically* with adult g.

A recent study with an average sample size of about 500 reported that adult-derived PGS predicted cognitive development with increasing strength across development (Gustavson et al., 2025). The highest prediction reported was for the EA-PGS, predicting a peak of 18% of the variance in cognitive ability at age 16, which is twice as much as the usual result of 9% for adult samples (Okbay et al., 2022). The g-PGS also showed a parallel pattern but with lower predictive strength. In addition, longitudinal analyses suggested that the increased prediction is attributable to the strengthening influence of stable genetic effects shared across ages, rather than the emergence of new, age-specific genetic factors.

Our study builds on these findings using data from the Twins Early Development Study (TEDS; Lockhart et al., 2023), a population-based cohort with over 10,000 British white children. TEDS provides considerably greater statistical power in sample size compared to the Gustavson et al. (2025) report. TEDS children were assessed using diverse measures of cognitive development in toddlerhood and early childhood (2, 3, and 4 years), middle childhood (7, 9, and 10 years), adolescence (12, 14, and 16 years), and adulthood (25 and 26 years), providing repeated measures of g, verbal ability, and nonverbal ability.

We aim to extend prior cross-sectional and confirmatory factor findings while addressing several developmental aspects that were not covered in previous work. First, we create a polygenic g score as a composite predictor by combining EA-PGS and g-PGS (Maier et al., 2018). Second, we include latent growth curve modelling to chart developmental trajectories as predicted by the polygenic g score. Third, we examine the tails of the polygenic distribution to investigate if cognitive development diverges for the highest and lowest scorers. Finally, educational achievement, behaviour problems, and anthropometric outcomes were also assessed longitudinally, enabling a comprehensive examination of polygenic g score prediction across trait domains.

In our preregistration (https://osf.io/vhuyj/overview), we specified five hypotheses: 1) the polygenic g score significantly predicts most cognitive outcomes, with educational achievement yielding the strongest correlation, 2) prediction for cognitive abilities increases from toddlerhood to early adulthood; 3) the polygenic g score also predicts behaviour problems and anthropometric outcomes; 4) prediction is linear across the distribution, indicating that high and low extremes are quantitatively, not qualitatively, different from the rest of the distribution; and 5) there are no significant sex differences in polygenic g prediction.

## Methods

### Sample

We leveraged data from the Twins Early Development Study (TEDS), a longitudinal cohort of 13,759 families with twins born between 1994 and 1996 in England and Wales (Lockhart et al., 2023). Phenotypic data were collected across multiple waves, including assessments at approximately ages 2, 3, 4, 7, 8, 9, 10, 12, 14, 16, 18, 21, 25, and 26 years. Genotypic data were available for 10,346 participants. Ethical approval for TEDS was obtained from King’s College London Research Ethics Committee (References: PNM/09/10–104 and HR/DP-20/2122060), and informed consent was obtained prior to each wave of data collection.

For cross-sectional analyses, all participants with available DNA data and at least one phenotypic measure were included. To calculate a total score for a given measure, participants were required to have completed at least half of the scale. For measures composed of multiple subscales, the same rule applied to each subscale, and a participant was included only if at least half of the subscales were available.

For longitudinal analyses, we included participants with data available for at least two ages (N = ∼4500 to ∼8000).

### Measures

#### Phenotypic Measures

The present study focuses on outcomes selected for their consistent assessment across the ages. From ages 2 to 26, we assessed cognitive abilities, educational achievement, behaviour problems, and anthropometric measures using age-appropriate measures. Additional measures specific to one age or one developmental stage are provided in the supplementary text. Complete documentation of all measures across ages is available in the TEDS data dictionary (https://datadictionary.teds.ac.uk/home.htm).

##### Early Cognitive Abilities (Ages 2-4)

Phenotypic measures at early ages were collected via booklets sent to families. At ages 2 and 3, booklets were sent only to families of twins born in 1994 and 1995, as twins born in 1996 were not age-appropriate for the tests. At age 4, booklets were sent to all families.

Verbal ability at ages 2 to 4 included vocabulary (what children can say) and grammar (how children use words). Vocabulary was assessed via parent-reported checklists. At age 2, parents completed a 100-word checklist adapted from the MacArthur Communicative Development Index (MCDI; Fenson et al., 1993, 2000). At age 3, the checklist included 45 MCDI words and 55 new words from a literature review and pilot testing, along with two additional questions about whether the child was talking and combining words. At age 4, 48 words selected from the literature review and pilot testing were used. Grammar was also measured using questions derived from the MCDI. At ages 2 and 3, parents completed the 6-item word use and 12-item sentence complexity scales. At age 3, the word use scale was expanded to 12 items. At age 4, a single 6-point global rating scale assessed language complexity from ‘not yet talking’ to ‘talking in long and complicated sentences.’ The verbal ability composite for the MCDI was calculated as the standardised mean of vocabulary and grammar scores. More detailed descriptions of the TEDS verbal measures between ages 2 to 4 are available in the TEDS data dictionary and previous TEDS publications (Dionne et al., 2003; Hayiou-Thomas et al., 2012).

Nonverbal ability was assessed using the Parent Report of Children’s Abilities (PARCA; Saudino et al., 1998), including parent-administered tasks and parent- report questionnaires. At age 2, parent-administered tasks included matching (8 items), brick building (4 items), folding (1 item), and copying (7 items) from the Bayley Scales of Infant Development (Bayley, 1993) and a design drawing task (4 items) adapted from the McCarthy Scales (McCarthy, 1972). The parent-report component assessed conceptual knowledge (26 items). At age 3, parent-administered tasks included odd-one-out (16 items), design drawing (6 items), and matching (16 items), with conceptual knowledge assessed via 24 items. At age 4, parent-administered tasks comprised the age 3 odd-one-out and design drawing tasks, plus draw-a-man (1 item) and puzzles (12 items). Conceptual knowledge was assessed via 12 items. The nonverbal ability composite was calculated as the standardised mean of parent-administered and parent-report PARCA scores, following the TEDS data dictionary and established practice in previous TEDS publications (Asbury et al., 2005; Oliver et al., 2004; Petrill et al., 2001; Saudino et al., 1998).

General cognitive ability at each age was calculated as the standardised mean of verbal and nonverbal composites.

##### Cognitive Abilities at Later Ages (Ages 7-25)

Cognitive ability measures at ages 7, 9, 10, 12, 16, and 25 have been described in detail in a previous TEDS publication and are only briefly summarised here (see Lin et al., 2025 supplementary materials). The only age not included in Lin et al. (2025) was age 14, though the measures are similar to other ages.

At age 7, cognitive assessments were conducted via telephone interviews. Verbal ability was measured using the Wechsler Intelligence Scale for Children (WISC-III) Similarity and Vocabulary tests (Wechsler, 1992). Nonverbal ability was assessed using the Conceptual Grouping Test and WISC Picture Completion Test (McCarthy, 1972).

At ages 9 and 10, verbal ability was assessed using WISC-III as a Process Instrument (WISC-III-PI) Vocabulary and General Knowledge tests (Kaplan et al., 1999). Nonverbal ability was measured using the Cognitive Abilities Test 3 (CAT3) figure classification and figure analogy tests at age 9 and WISC-III-UK Picture Completion and Raven’s tests at age 10 (Raven et al., 1996; Smith et al., 2001). Assessments were administered via mailed booklets at age 9 and online at age 10 and all subsequent ages.

At age 12, verbal ability comprised language tests (syntax, semantics, pragmatics) and reading tests (comprehension and fluency) (GOAL, 2002; Hammill et al., 1994; Markwardt, 1997; Torgesen et al., 1999; Wiig et al., 1989; Woodcock et al., 2001). Nonverbal ability was assessed using mathematical ability tests from the National Foundation for Education Research (Smith et al., 2001). General cognitive ability was independently assessed using WISC-III-PI Vocabulary, General Knowledge, Picture Completion (Wechsler, 1992), and Raven’s Pattern test (Raven et al., 1996).

At age 14, verbal ability was assessed using a 27-item WISC-III-PI vocabulary multiple-choice test (Kaplan et al., 1999). Nonverbal ability was measured using the 30-item Raven’s Standard Progressive Matrices (Raven et al., 1996).

At age 16, verbal and nonverbal abilities were assessed using the Mill Hill Vocabulary test (Raven et al., 1998) and Raven’s Standard and Advanced Progressive Matrices (Raven et al., 1996).

Between ages 7 and 16, except for age 12, verbal and nonverbal composites were calculated as standardised means of their respective component tests, and general cognitive ability was calculated as the standardised mean of the verbal and nonverbal composites.

At age 18, assessment only included two spatial ability online measures developed by TEDS researchers (Malanchini et al., 2020; Rimfeld et al., 2017): a bricks test and a navigation study. The Bricks test measured spatial ability through mental rotation and visualisation using both 2D and 3D stimuli across six subtests (9 items each): 2D rotation, 2D rotation and visualisation, 2D visualisation, 3D rotation, 3D rotation and visualisation, and 3D visualisation. Both individual subtest scores and the overall Bricks total score (mean of all six subtests) were included in the present study. The Navigation test included 30 tasks across six types (5 items each): orientation-direction, orientation-landmarks, map reading without memory, map reading with memory, perspective, and scanning. Each task generated accuracy, speed, and total scores; only the overall total score (mean of the six task types) was used in analyses.

At age 25, cognitive abilities were assessed using Pathfinder, a gamified web-based measure developed by TEDS researchers (Malanchini et al., 2021). Verbal ability (20 items) included the Mill Hill vocabulary, missing letter, and verbal reasoning tests. Nonverbal ability (20 items) included Raven’s standard progressive matrices and three visual puzzle tests on analogies, grouping, and logical sequences. Unlike earlier ages, cognitive ability scores at age 25 were not standardised; general cognitive ability scores ranged from 0 to 40, while verbal and nonverbal scores each ranged from 0 to 20.

##### Educational Achievement

Educational outcomes were examined using educational achievement from primary school to university. Educational outcomes at ages 7, 9, 10, 12, 16, 18, 21, and 26 have been described previously (Lin et al., 2025). The present study extended these general outcomes by including subject-specific grades up to age 18 and adding assessment at age 14.

At ages 7, 9, 10, and 12, teachers rated achievement in English and mathematics (starting at age 7) and science (starting at age 9) based on the National Curriculum Levels (https://www.gov.uk/national-curriculum/overview). The ratings ranged from 0-4 at age 7 and 0-9 at later ages.

At age 14, parents reported grades in English, mathematics, and science and the grades were translated to the 0-9 National Curriculum Levels.

At age 16, the General Certificate of Secondary Education (GCSE) is a national-level exam taken at the end of compulsory education. GCSE exam grades were obtained for core subjects (English, mathematics, and science), humanities, and languages. The grades ranged from 4 (G) to 11 (A*).

At age 18, A-level and AS-level qualifications were assessed across English, mathematics, science, technology, humanities, languages, and vocational subjects. A-levels are two-year qualifications completed after compulsory education and required for university entry. AS-levels represent completion of the first year only.

When A-level grades were unavailable, AS-level grades were used. Grades ranged from 1 (E) to 6 (A*).

At age 21, university degree classification was self-reported on a scale from 1 (lowest pass) to 5 (first-class honours).

At age 26, most twins had completed their education. Therefore, educational attainment (i.e., years of schooling) was used to measure educational outcomes. For twins missing age 26 data, age 21 educational attainment was used (correlation between two ages: r = 0.86).

##### Behaviour Problems

Behaviour problems were primarily assessed using the Strengths and Difficulties Questionnaire (SDQ) (Goodman, 1997), which was administered consistently across ages from 2 to 26 and across multiple informants. The SDQ yields five subscales: conduct problems, emotional problems, hyperactivity, peer problems, and prosocial behaviour. The first four problem subscales were summed to create a total problems score at each age.

Between ages 2 to 4, the Preschool Behaviour Questionnaire (Behar scales) was used to measure parent-reported behaviour problems (Behar, 1977), with items converted to SDQ-comparable components by TEDS researchers (https://datadictionary.teds.ac.uk/pdfs/4yr/234yr_behaviour_items.pdf). From age 7 onward, the standard 25-item SDQ was administered with multiple informants: parent reports at ages 7, 9, 12, 16, and 21 (emotional and peer problem subscales were not available from parents at age 16); teacher reports at ages 7, 9, and 12; and self-reports at ages 12, 16, 21, and 26.

We also included measures of anxiety and ADHD symptoms collected at multiple ages. Additional behaviour problems assessed at one or two ages were examined as outcomes; these results are presented in the Supplementary Materials.

##### Socioeconomic Status (SES)

Family SES was assessed at birth and at ages 7, 16, and 21. Each SES composite was standardised as z-scores and calculated from parental employment status (coded according to the UK Standard Occupational Classification or SOC, https://www.ons.gov.uk/methodology/classificationsandstandards/standardoccupationalclassificationsoc/), parental highest educational qualifications, and household income.

#### Genetic Measures

Genotyping for the TEDS participants was conducted on one of two platforms: the Affymetrix Genome-Wide Human SNP Array 6.0 and the Illumina HumanOmniExpressExome-8v1.2. DNA was obtained from either buccal cheek swabs or saliva samples collected over several waves.

Following quality control, genotypes from both platforms were separately phased using EAGLE2 and then imputed to the Haplotype Reference Consortium (release 1.1) (Durbin, 2014; Loh et al., 2016; McCarthy, 2016). After imputation, harmonisation, and merging of the two datasets, a final set of 7,363,646 SNPs for 10,346 twins remained for analysis. More details of the genotyping and imputation processes are described in previous TEDS publications (Lin et al., 2025; Selzam et al., 2018).

We constructed polygenic scores using LDpred2-auto, a Bayesian method that adjusts GWAS summary statistics for linkage disequilibrium (LD) using the HapMap3+ reference panel (Privé et al., 2021). Approximately 1.1 million SNPs common between the TEDS sample and the HapMap3+ panel were included. The most recent GWAS summary statistics of educational attainment (N = 765,283) and intelligence (N = 266,453) were used (Okbay et al., 2022; Savage et al., 2018). Although the educational attainment GWAS includes approximately three million individuals, the publicly available summary statistics exclude the 23andMe cohort due to data-sharing restrictions. Additionally, we acknowledge the availability of alternative GWAS for cognitive abilities (e.g., Davies et al., 2018; de la Fuente et al., 2021; Lam et al., 2021; Williams et al., 2023). These datasets represent partially overlapping cohorts and employ varying methodologies for constructing the latent g factor. We used the Savage summary statistics because they remain one of the most widely used and validated benchmarks for cognitive abilities.

To maximise the prediction of general cognitive ability, we combined the EA-PGS and g-PGS polygenic scores using SMTPred (Maier et al., 2018). SMTPred calculates optimal multi-trait weights by scaling each input polygenic score to its expected variance using GWAS summary statistics (i.e., SNP heritability, genetic correlation, and sample size). Because our input scores were derived using a Best Linear Unbiased Prediction (BLUP) approach, which accounts for linkage disequilibrium structure and prevents overestimating highly correlated variants, SMTPred utilises the same BLUP weighting to further increase prediction accuracy.

In our sample, EA-PGS was weighted 0.91 and g-PGS 0.31. The resulting combined score, termed the *polygenic g score*, correlated with g-PGS at 0.62 and with EA-PGS at 0.97 (Figure 1). While the correlation with EA-PGS is high, the composite was derived empirically, providing the strongest prediction of cognitive abilities that could be achieved using two polygenic scores on SMTPred. This polygenic g score was used as the primary predictor for all cognitive abilities, educational achievement, educational attainment, behaviour problems, and anthropometric outcomes in the present study.

**Figure 1.**
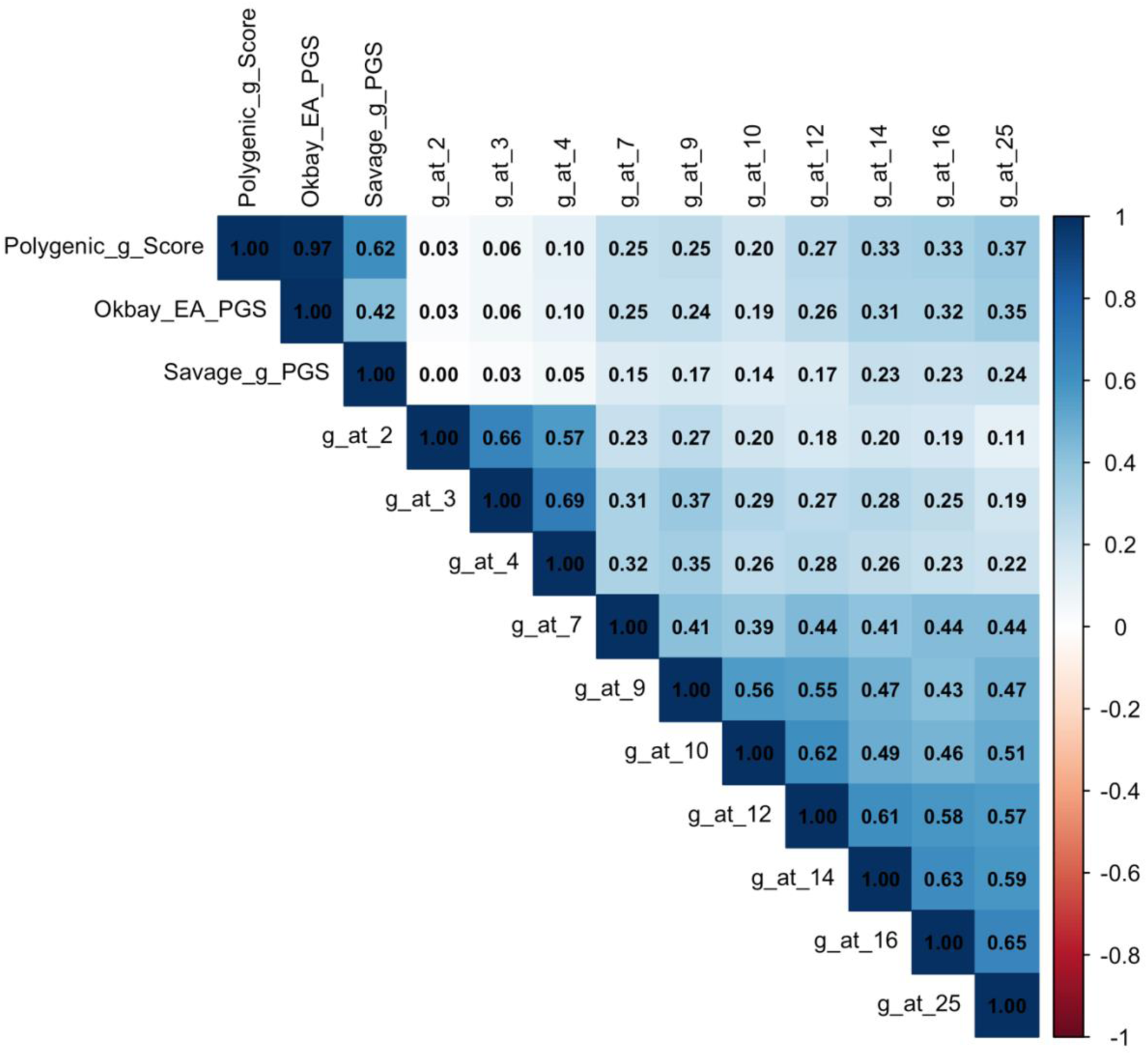
Correlations between polygenic g score, g-PGS, EA-PGS, and general cognitive ability (g) across ages.

### Statistical Analyses

The present study was pre-registered at https://osf.io/vhuyj/overview. Analyses were performed using RStudio 2023.09.1+494 with codes available on GitHub (https://github.com/YujingLinn/Cog-PGS).

Before carrying out the main analyses, we conducted sensitivity tests to examine potential effects of age, sex, zygosity, twin birth order, genotyping chip, and the first ten genomic principal components (PCs) on all phenotypes and family socioeconomic status measured across development. Sensitivity analyses of the polygenic g score were also conducted, except for age effects. Results are detailed in Supplementary Table S1.

Sensitivity analyses revealed significant associations between several covariates and the phenotypes. Specifically, age, sex, zygosity, genotyping chip, and the first ten genomic PCs showed significant effects on about half of the phenotypes. Twin birth order showed no significant effects. For the polygenic g score, significant effects were detected for zygosity and the ninth and tenth PCs. We adopted a conservative approach by including age, sex, genotyping chip, and the first ten genomic PCs as covariates in all cross-sectional analyses for consistency, even though not all phenotypes were significantly associated with every covariate. Age was excluded as a covariate in longitudinal analyses.

The apparent zygosity effect in the TEDS can arise from maternal characteristics associated with dizygotic twinning. DZ twinning is related to maternal fertility and increases with maternal age due to a higher probability of double ovulation (Hubers et al., 2024). Consequently, mothers of DZ twins tend to be older on average than mothers of MZ twins. Because maternal age is associated with SES, DZ twins in our sample had slightly higher SES at birth than MZ twins. After adjusting for SES at birth, the zygosity effect disappeared (variance explained = ∼0.2%). A minor contribution also arose from the genotyping design: the initial Affymetrix phase primarily included unrelated twins, whereas the later Illumina phase preferentially recruited DZ co-twins. Therefore, we performed additional analyses stratifying the sample into monozygotic and dizygotic twins to examine whether the polygenic g score predictions were robust across zygosity groups.

#### Polygenic Score Prediction

Our main analysis examined associations between our polygenic g score and outcomes at each age using a subsample of unrelated individuals (one randomly selected twin per pair). All phenotypes were analysed, including both composite scores and their constituent components or specific test scores.

All continuous outcomes were standardised prior to analysis. Models included age, sex, genotyping chip, and the first ten PCs as covariates to control for batch effects and population structure. We report standardised beta coefficients for the polygenic g predictor. Incremental variance explained was calculated as the difference in R² between the full model and a reduced model containing only covariates. Confidence intervals were estimated using percentile bootstrapping with 1000 iterations.

To ensure the robustness of our findings, we repeated all prediction analyses separately among females, males, monozygotic twins, and dizygotic twins.

#### Confirmatory Factor Analyses

Next, we examined the polygenic g score prediction of a cross-time confirmatory factor of the phenotypes. We extracted cross-time latent factors from repeated measures of the same construct, assuming these reflect stable underlying latent traits. Using multilevel structural equation modelling (SEM) to account for twin structure (i.e., family clustering), we conducted confirmatory factor analyses (CFA) for g, verbal ability, nonverbal ability, educational achievement, SDQ, anxiety, and ADHD. Whereas cross-sectional analyses used one randomly selected twin per family to ensure independence, SEM analyses included both twins from each pair to increase statistical power, retaining the full sample while appropriately adjusting for within-family non-independence.

We used three approaches to extract latent factors for total scales and subscales: a single cross-time latent factor across all ages, a higher-order latent factor derived from developmental stages, and a higher-order latent factor derived from shared measurement methods.

For cognitive and educational outcomes, informants and measurement methods varied by developmental stage. Early cognitive measures were parent- administered, while later tests were self-administered. Similarly, educational achievement was primarily teacher-evaluated, while national exam scores were parent-reported. Because different informants were used at different ages (e.g., parent-administered tests in early childhood, self-administered tests in adolescence), the confirmatory factors for cognitive abilities and educational achievement reflect developmental changes over time as well as potential method variance associated with different informants. To account for the resulting method variance and developmental changes, cognitive abilities were modelled by both developmental stage and assessment method. Developmentally, first-order latent factors were extracted for early childhood (ages 2, 3, and 4), middle childhood (ages 7, 9, and 10), and adolescence (ages 12, 14, and 16). Because age 25 constituted a single time point, no separate latent factor was estimated; instead, it was integrated directly alongside the first-order factors to establish a second-order latent factor. This developmental structure was replicated for verbal and nonverbal abilities.

Methodologically, cognitive measures were grouped into early measures (ages 2, 3, and 4), middle measures (ages 7, 9, 10, 12, and 14), and later measures (ages 16 and 25). This method structure was identical for verbal abilities and nonverbal abilities, with one exception for nonverbal tests: age 7 was modelled independently due to its unique use of the WISC, while ages 9 through 25 were grouped together based on their consistent use of the Raven’s matrices.

For educational achievement, we constructed a cross-time latent factor aggregating the teacher-rated primary school grades.

For behaviour outcomes, multiple informants were used at some ages. For example, SDQ was reported by parents at ages 2, 3, 4, 7, 9, 12, 16, and 21; by teachers at ages 7, 9, and 12; and by self-report at ages 12, 16, 21, and 26. This multi-informant approach reflects developmental appropriateness: parent reports are most suitable in early childhood when children cannot report reliably, teacher reports provide complementary school-based perspectives during school age, and self-reports become increasingly valid as adolescents develop greater self-awareness and autonomy. We therefore extracted both cross-rater latent factors (combining all informants) and within-rater latent factors when three or more assessments from the same informant were available for a given construct. Since the same assessment scales were used across ages for behaviour outcomes, we only constructed higher-order latent factors stratified by developmental stages.

We then used the polygenic g score to predict these latent factors and compared predictive validity against the age-specific cross-sectional predictions in the whole sample and the sex-stratified sample. To approximate real-world predictive utility, we extracted the latent factor scores for each individual and treated them as the dependent variables in our prediction models. Additionally, to establish the theoretical upper bound of the genetic effects, we incorporated the polygenic g score directly into our SEM as a predictor, generating an error-free prediction at the latent level. All models included age, sex, genotyping chip, and the first 10 genomic principal components as covariates, with standardised beta coefficients and incremental R² reported.

#### Latent Growth Curve Model

We conducted latent growth curve analyses to examine how the polygenic g score predicts both baseline levels (intercept) and developmental trajectories (slope) across time. The analyses were conducted for measures with repeated assessments across development, including general cognitive ability, verbal and nonverbal abilities, educational outcomes, the subscales of the SDQ, anxiety, ADHD, height, and BMI. Because all phenotypic measures were standardised (mean = 0, SD = 1) within each age group prior to analysis, our models capture changes in a participant’s relative ranking compared to their peers. Therefore, a positive association with the intercept indicates that higher polygenic g scores predict a higher initial relative ranking in the cohort, while a positive association with the slope indicates that higher polygenic g scores predict an upward shift in peer ranking over time. To account for varying developmental intervals between assessment waves, we used the exact age differences (in years) from baseline rather than treating the intervals between each assessment equally.

For measures with multiple potential informants, we prioritised consistency across developmental stages. For most phenotypes, we used parent reports before age 18 and self-reports from age 18 onwards. When parent reports were unavailable in childhood or adolescence, we prioritised teacher reports, followed by child reports; however, in most cases, child reports were used when parent reports were unavailable.

Missing data in the repeated measures were handled using Full Information Maximum Likelihood within the latent growth curve models (Enders & Bandalos, 2001). This approach uses all available data points for each participant, allowing inclusion of individuals with partially missing timepoints. Participants were included if they had data available for at least two timepoints. No additional imputation was performed for other variables included in the models.

Consistent with the confirmatory factor analyses, all latent growth models used the full twin sample with multilevel modelling to account for family clustering, maximising statistical power. Sex was included as a covariate in models for the whole sample. We also conducted multi-group analyses to compare intercepts and slopes between females and males, with family clustering accounted for within each sex-stratified sample.

#### Profile Analysis of Extremely High and Extremely Low Polygenic g Scores

We examined the developmental trajectories of participants with extreme polygenic g scores, defined as scores above or below three standard deviations from the population mean.

To maximise sample size, we leveraged the full sample and assigned polygenic g scores to the MZ co-twins of genotyped individuals, as only one twin per MZ pair was genotyped because MZ twins share identical genomes. This gave us 19 participants with scores three standard deviations above the mean and 12 participants with scores three standard deviations below the mean.

We described the observed values of cognitive, educational, and behaviour problem outcomes for these extreme groups across development (standardised for comparability).

To robustly compare differences in outcomes across development between high and low score groups, we divided the full sample into deciles based on polygenic g scores and conducted independent samples t-tests comparing the top and bottom deciles at each age.

#### Nonlinearity Tests

Finally, to test whether the relationship between polygenic g scores and outcomes is linear, we conducted regression analyses including a quadratic term for the polygenic g score. A significant quadratic term would indicate nonlinearity, representing either accelerating or decelerating effects at the extremes of the distribution. Conversely, a non-significant quadratic term would support linearity, suggesting that high and low extremes differ only quantitatively, not qualitatively, from the rest of the distribution.

#### Multiple Testing Correction

We applied the false discovery rate (FDR) correction to account for multiple testing across all regression analyses. All reported p-values are FDR-adjusted values.

## Results

We focus primarily on cognitive, educational, and behaviour problem phenotypes in this section, with other phenotypes discussed as relevant. Full results for all phenotypes, including anxiety, ADHD, anthropometric outcomes, and those assessed at just one or two ages, are reported in the Supplementary Materials.

A correlation matrix for all variables is presented in Supplementary Table S2, with domain-specific illustrations in Supplementary Figure S1. Descriptive statistics for all phenotypes and the polygenic g score are reported in Supplementary Table S3.

### Polygenic g Score Prediction from Toddlerhood to Early Adulthood

Polygenic g score significantly predicted most outcomes across phenotypes at most ages from childhood onwards. As shown in Figure 2a, for cognitive abilities, the prediction was weak or absent in toddlerhood (ages 2 to 4). From childhood, the prediction increased steadily and reached a peak in early adulthood (age 25) with standardised beta coefficients of 0.36 (95% CI [0.32, 0.40]) for both g and verbal ability, and 0.28 [0.24, 0.32] for nonverbal ability.

**Figure 2.**
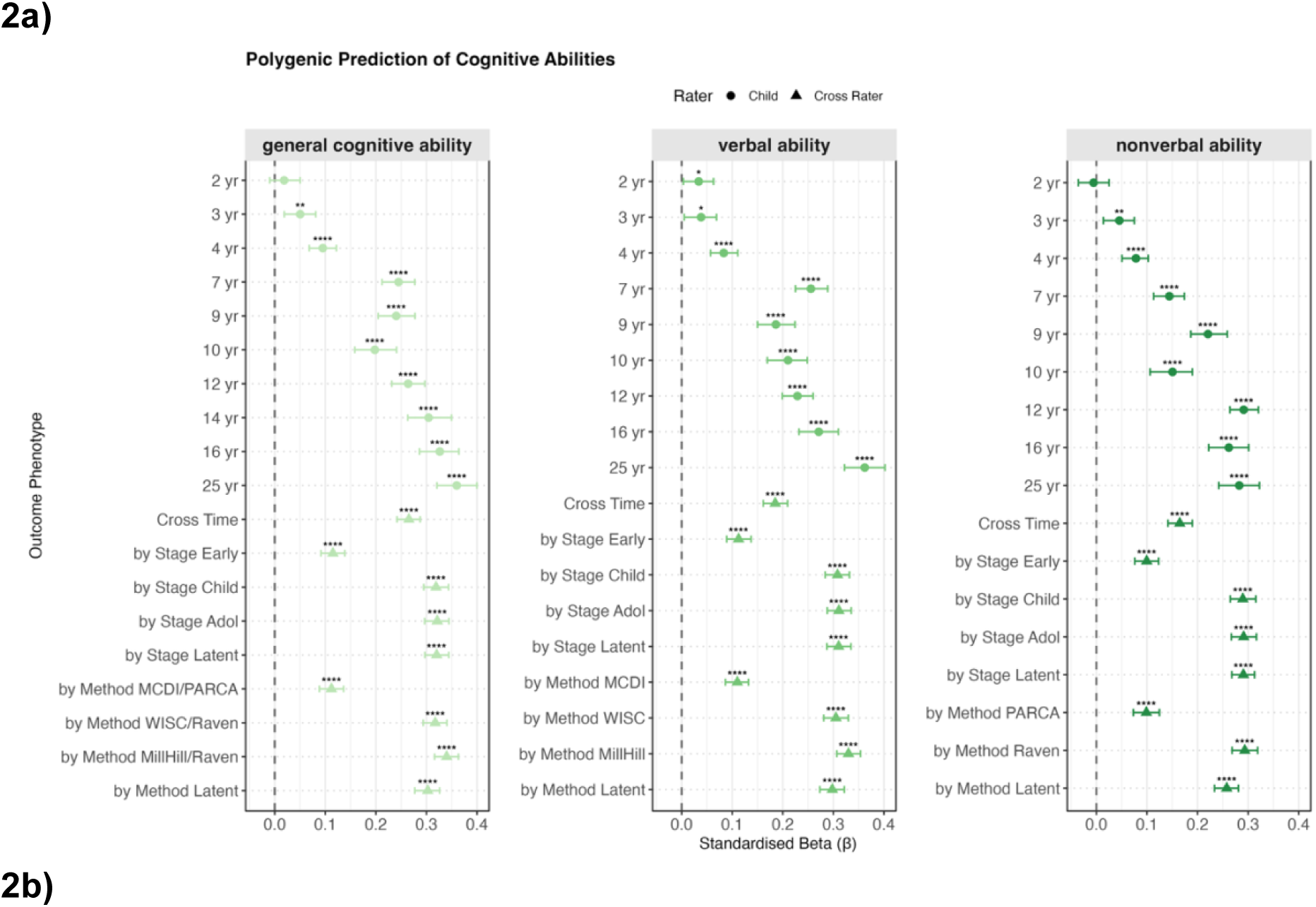
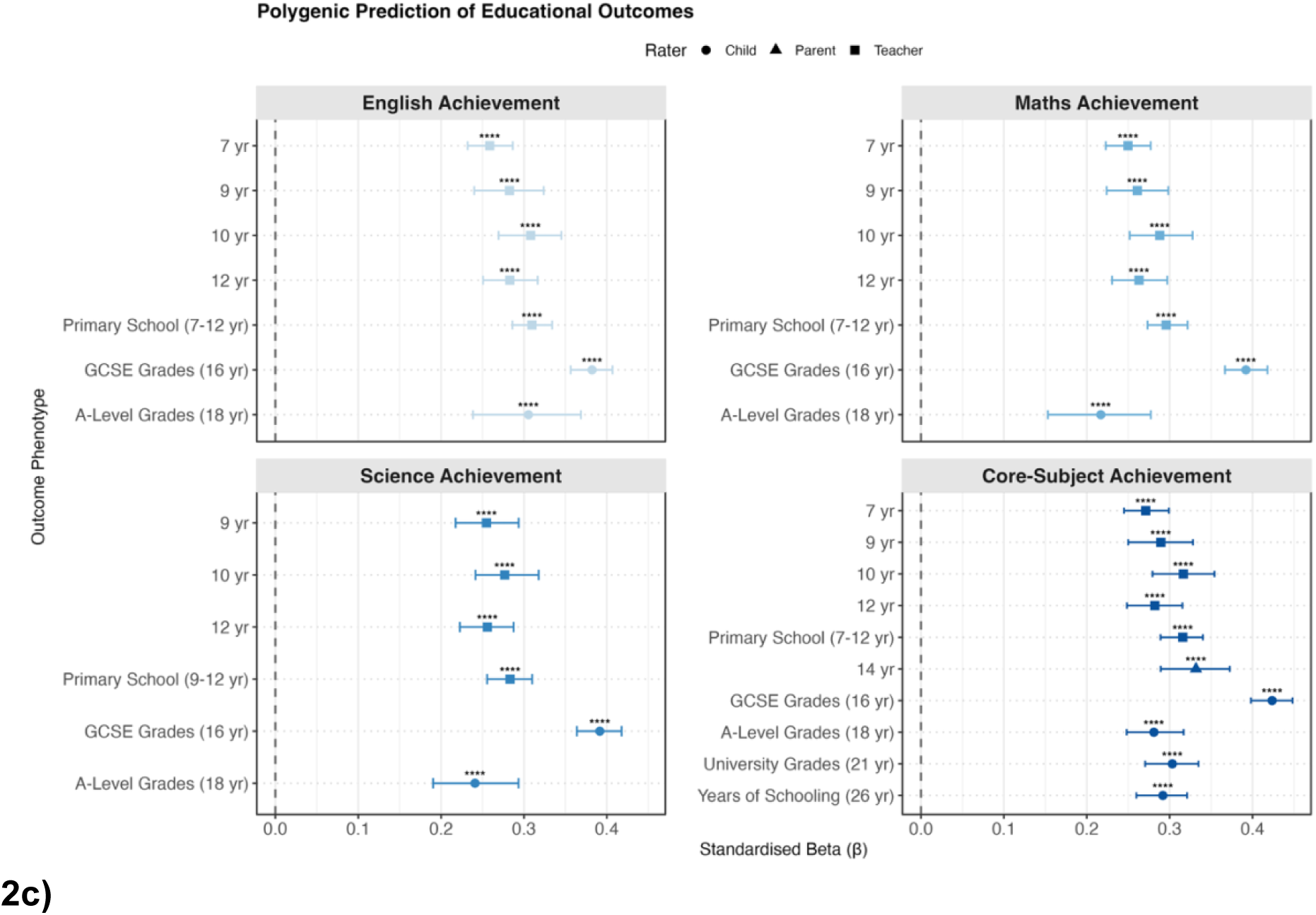
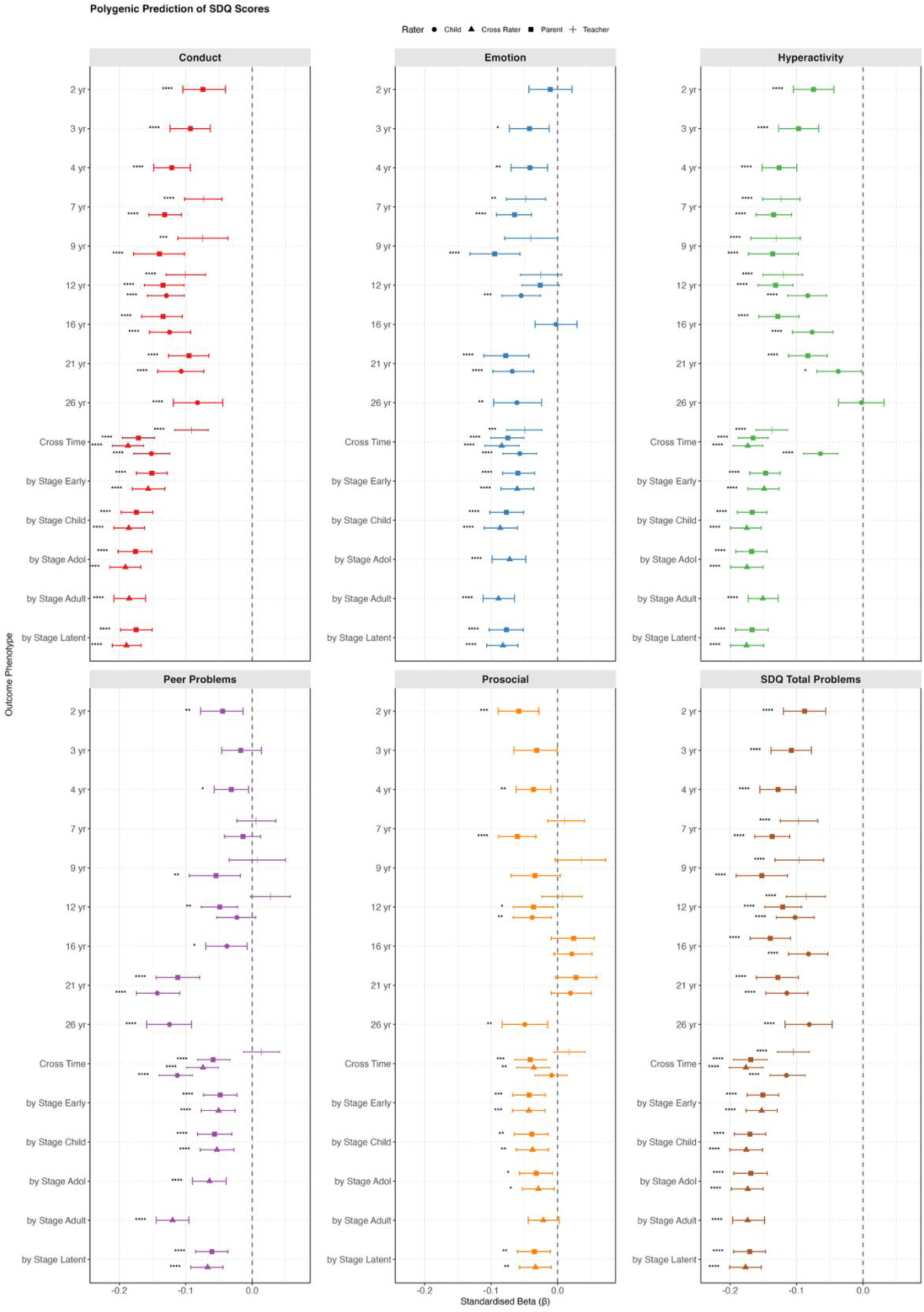
Polygenic g score prediction of cognitive abilities, educational achievement, and behaviour problems across development. Standardised beta (β) coefficients with 95% bootstrapped confidence intervals (1000 iterations) for the prediction of developmental outcomes by polygenic g score. **Panel a**: General cognitive ability (g) composite, and domain-specific verbal and nonverbal ability composites measured from ages 2 to 25. To account for variations in development and assessment, higher-order latent factors stratified by developmental stages and measurement methods, as well as cross-time common factors, were included. **Panel b**: Educational achievement in English, mathematics, and science, and a core-subject composite. Science was first measured at age 9, while English and mathematics were also assessed at age 7. The core-subject composite was derived from English, mathematics, and science (where available) from ages 7 to 18, with general university grades used at age 21 and years of schooling at age 26. For plotting purposes, ‘core subject’ serves as an overarching category representing educational achievement and attainment through age 26. A cross-time latent factor aggregating primary school grades was also included. **Panel c**: Strengths and Difficulties Questionnaire (SDQ) five subscales (conduct problems, emotion problems, hyperactivity, peer problems, and prosocial behaviour) and total problems score measured from ages 2 to 26. Cross-time latent factors and higher-order latent factors by developmental stages were also included. For SDQ, the rater is indicated by a symbol shape (parent, teacher, or child). For other outcomes, performance-based measures were used, with parent or teacher ratings at early ages and self-reports or test-based assessments at later ages. The vertical dashed line at β = 0 represents no association. Asterisks denote FDR-adjusted statistical significance: * p < 0.05, ** p < 0.01, *** p < 0.001, **** p < 0.0001. Complete numerical results, including sex-stratified and zygosity-stratified analyses, are reported in Supplementary Table S4.

Similar patterns were identified for the individual verbal and nonverbal tests: weak or absent prediction in early childhood that strengthened with age (see Supplementary Figures S2 and S3). Among verbal tests, the strongest prediction emerged for the Verbal Analogies from the age 25 Pathfinder battery (β = 0.33 [0.29, 0.37]). Among nonverbal tests, the strongest prediction was for the age 16 Understanding Number task (β = 0.31 [0.27, 0.35]).

For educational outcomes (Figure 2b), polygenic g score associations were moderate at age 7 (β = 0.25) and increased through mid-adolescence. Prediction strength peaked at age 16 for GCSE grades: English (β = 0.38 [0.36, 0.41]), mathematics (β = 0.39 [0.37, 0.42]), science (β = 0.39 [0.36, 0.42]), and the core-subject composite (β = 0.42 [0.40, 0.45]). Polygenic g score associations declined for A-level, university grades and years of schooling.

For behaviour problems, we focused on SDQ measures (Figure 2c). Negative associations were observed for most behaviour problem measures across ages and raters. Significant predictions emerged in toddlerhood for all SDQ subscales and peaked at a variance explained of 2% in late childhood and adolescence, then slightly weakened in early adulthood. For prosocial behaviour, the polygenic g score explained negligible variance (all <0.5%) after accounting for age and sex effects, despite slightly negative associations.

For other behaviour problems (anxiety in Supplementary Figure S4; ADHD in Supplementary Figure S5), weak negative associations were observed from early childhood onward. Interesting patterns emerged for height and BMI, with non-significant associations until early adulthood, then positive associations with height and negative associations with BMI (Supplementary Figure S6). For other miscellaneous outcomes, associations were largely positive for education- and cognitive-related phenotypes and showed mixed patterns for wellbeing-related phenotypes (Supplementary Figure S7).

As a sensitivity test, a full comparison confirmed that polygenic g score predictions outperform both the EA-only and g-only PGS predictions across phenotypes (Supplementary Table S4).

### Polygenic g Score Prediction of Cross-Time and Cross-Rater Latent Factor

We extracted cross-time latent factors to examine polygenic g score prediction across development and measurements. For behaviour outcomes, we additionally constructed both cross-rater and within-rater latent factors. All observed variables loaded significantly onto their respective latent constructs (Supplementary Table S5), with standardised loadings ranging from 0.15 to 0.95. Model fit indices are provided in Supplementary Table S6.

For models based on a single cross-time factor (and for cross-rater models of behaviour problems), overall model fit was modest, reflecting heterogeneity across developmental stages and measurements. However, when latent factors were first extracted within developmental stage or measurement and then combined into a higher-order latent factor, model fit improved substantially and was generally good (Supplementary Table S6). These hierarchical models better capture the developmental and methodological structure of the data while still isolating the shared variance representing the underlying trait.

To evaluate predictive validity, we first extracted individual-level factor scores and used them as outcomes in regression models predicted by polygenic g scores. In addition, to estimate the theoretical upper bound of prediction, we incorporated the polygenic g score directly into the SEM as a predictor of the latent factors. This approach estimates prediction at the latent level without measurement error in the outcome.

For cognitive abilities, polygenic prediction differed across latent modelling approaches. For general cognitive ability, prediction was strongest for the developmental-stage higher-order factor (β = 0.32), followed by the measurement-specific higher-order factor (β = 0.30) and the cross-time factor (β = 0.27). A similar pattern was observed for the latent factors representing verbal and nonverbal abilities. However, these latent-factor predictions did not exceed the strongest prediction observed at any individual age. One explanation is that cognitive measures in late adolescence or adulthood more closely capture the mature cognitive trait and may include age-specific genetic influences that are not shared with earlier developmental measures. In contrast, cross-time latent factors retain only the variance common across ages, which can attenuate prediction if the polygenic score also captures age-specific variance.

When the polygenic g score was incorporated directly into the SEM, the pattern of prediction across modelling approaches was similar, but effect estimates were stronger (Supplementary Table S7). Prediction was strongest for a cross-time latent cognitive factor capturing cognitive ability accounting for developmental stages within the SEM (β = 0.39). By integrating polygenic prediction directly into the SEM framework, this approach provides a theoretical upper bound of genetic influences on g under minimal measurement error.

For educational outcomes, the primary school stage latent-factor predictions were similar to and did not substantially exceed those obtained for individual measures. When the prediction was estimated directly within the SEM, the strongest effect was observed for the core-subject latent factor (β = 0.38), compared to β = 0.32 using extracted factor scores.

For behaviour problems, the cross-time cross-rater factor and the developmental-stage higher-order cross-rater factor showed similar predictions for SDQ total problems (β = 0.18), representing the strongest estimates across behaviour models and exceeding those observed at individual ages. When the prediction was estimated directly within the SEM, the corresponding effect increased to β = 0.20.

Within-rater models generally showed better model fit, with the developmental-stage higher-order within-rater models providing the best overall fit. However, cross-rater predictive estimates were higher, though not significantly higher, than the corresponding within-rater models. One possible explanation is that different informants capture partially distinct aspects of behaviour problems. Combining information across raters may therefore capture a broader representation of the underlying behaviour construct, which could be more strongly aligned with the stable liability indexed by the polygenic g score.

Standardised path diagrams for all 73 models appear in Supplementary Figures S8 and S9.

### Polygenic g Score Prediction of Developmental Trajectories

Beyond examining associations at individual ages and cross-time common factors, we used latent growth curve models to examine how the polygenic g score predicts both baseline levels (intercepts) and rates of change (slopes). Only longitudinal measures were included for cognitive abilities, educational achievement, behaviour problems, and anthropometric outcomes. The full results are presented in Supplementary Table S8 for the full sample and in Table S9 for the sex-stratified sample.

Overall, the latent growth curve models demonstrated acceptable fit to the data across phenotypes (average Comparative Fit Index [CFI] = 0.88; full fit indices for all models are detailed in Supplementary Tables S8 and S9).

At the baseline, the polygenic g score already displayed significant positive associations with cognitive, educational, and anthropometric outcomes (Figure 3a for g and Figure S10 for all phenotypes). The strongest baseline prediction was for core-subject achievement (β_standardised_ = 0.31), followed by English, maths, and science achievement (β = 0.26 to 0.27). The baseline predictions for cognitive abilities were between 0.06 and 0.07. In contrast, all behaviour problem intercepts showed negative associations ranging from −0.02 to −0.13, indicating that children with higher polygenic g scores generally exhibited higher initial cognitive and educational performance and fewer behaviour problems.

**Figure 3.**
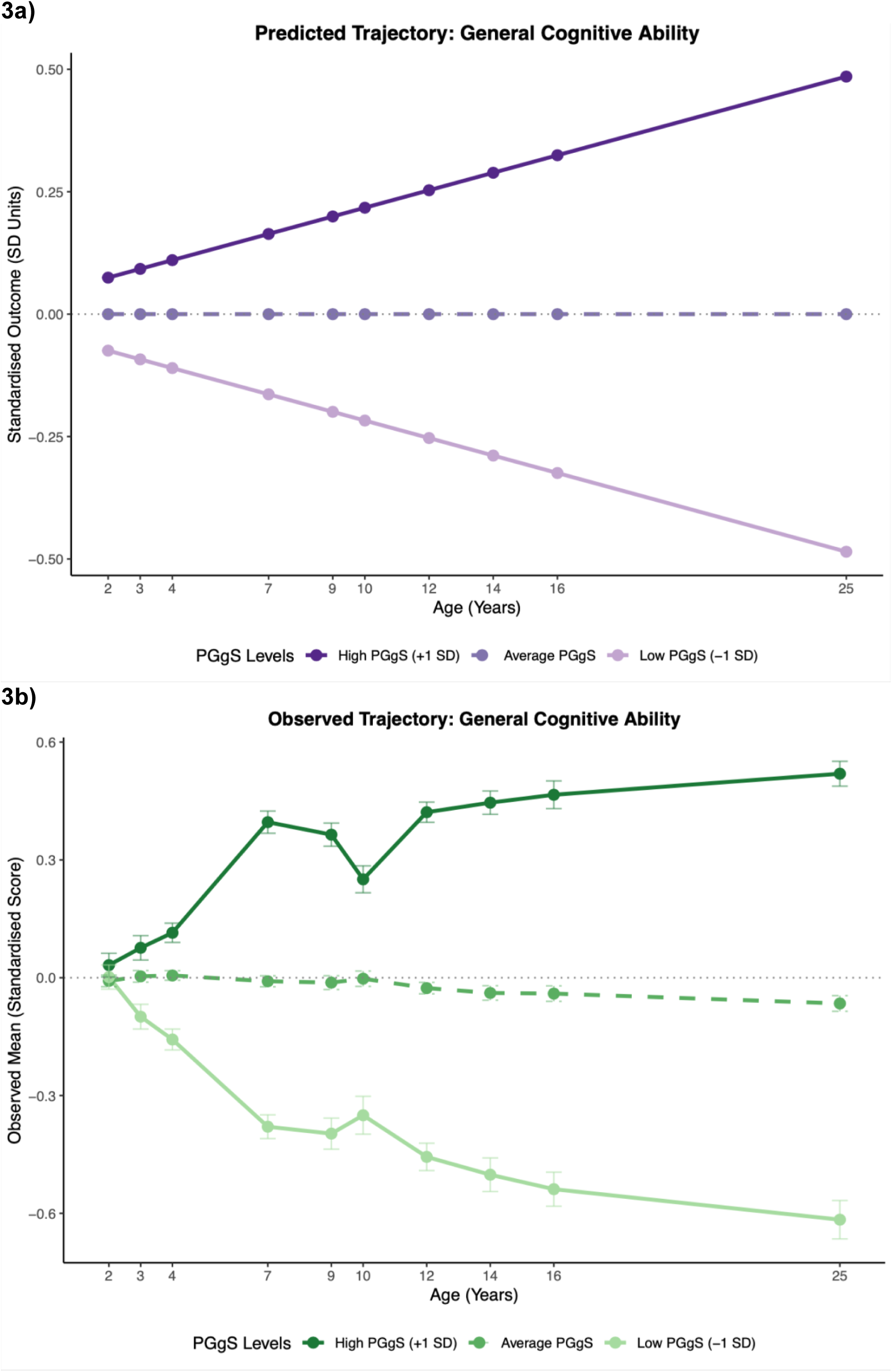
Predicted (3a) and empirically observed (3b) latent growth trajectories for general cognitive ability from ages 2 to 25. Trajectories are presented for individuals with high (+1 SD), average (Mean), and low (–1 SD) polygenic g scores. The predicted model was derived directly using the unstandardised estimates from the latent growth curve. The observed model represents the descriptive average scores at each developmental wave for the three groups. All observed scores were standardised within each age wave prior to plotting to reflect rank-order changes. Visualisations of the predicted and observed models for all longitudinal outcomes are available in Supplementary Figures S10 and S11, respectively. Corresponding sex-stratified models are provided in Supplementary Figures S12 and S13. PGgS = polygenic g score

Across development, the polygenic g score predicted steeper relative increases for most cognitive (β = 0.02) and educational outcomes (β = 0.01).

Together, the positive baseline and growth effects for cognitive and educational outcomes implied that children with higher polygenic g scores started with higher cognitive and educational performance and continued to gain a relative advantage at a steady rate across development.

For behaviour problems, while the baseline effects were mostly significant, the slope effects were not, suggesting that children with higher polygenic g scores begin with fewer behaviour problems but show slightly faster relative increases over time, gradually moving closer to the average developmental trajectories.

In contrast, we found the opposite for BMI, which yielded a combination of a positive intercept (0.05) and a negative slope (−0.12), indicating that children with higher polygenic g scores started with a higher BMI in relation to their peers, while gradually moving down to a BMI lower than their peers across development.

To complement the modelled estimates, observed trajectories are provided (Figure 3b for g and Figure S11 for all phenotypes). The empirical trajectories align with the predicted trajectories, particularly for cognitive and educational outcomes. The notable exception was height; the observed data revealed substantial developmental fluctuations, which correspond to the poor fit of the latent growth model for height.

### Polygenic g Score Prediction Across the Distribution

To examine whether polygenic g score prediction varies across the distribution, we first tested for nonlinear effects by including both linear and quadratic terms in the same model. Across all outcomes, no significant quadratic effects were observed in either the full sample or sex-stratified analyses (all incremental R² < 0.01, all FDR-adjusted p > 0.05; Supplementary Table S10). These results indicate that associations are linear throughout the distribution, with no evidence of stronger or weaker effects at the ends of the polygenic score distribution.

Second, we compared phenotypic outcomes between individuals at the distributional tails—those in the top versus bottom polygenic g score deciles (Supplementary Table S11). For cognitive abilities, individuals at the top of the distribution consistently scored significantly higher than those at the bottom from age 7 onwards, with occasional exceptions (e.g., spatial ability in adolescence and adulthood). Differences were non-significant in toddlerhood. For educational achievement, all comparisons between distributional extremes were statistically significant. For behaviour problems, approximately half of the phenotypes showed non-significant differences between the top and bottom deciles. For example, peer problems across most ages and raters, as well as height and BMI, showed no significant differences between top and bottom deciles. The magnitude of decile differences generally corresponded to population-level prediction strength.

Outcomes with stronger overall associations (such as cross-age and cross-rater composite scores) consistently showed significant differences between the top and the bottom polygenic g score deciles.

### Polygenic g Score Prediction for the Highest and Lowest Individuals

To examine outcomes at the distributional extremes in greater detail, we identified individuals with the highest (N = 19) and lowest (N = 12) polygenic g scores (Figure 4). Sample characteristics are presented in Supplementary Table S3 and group comparisons (t-tests) in Supplementary Table S12. Due to small sample sizes, most comparisons were not statistically significant. Notably, SES differed significantly between groups at all ages.

**Figure 4.**
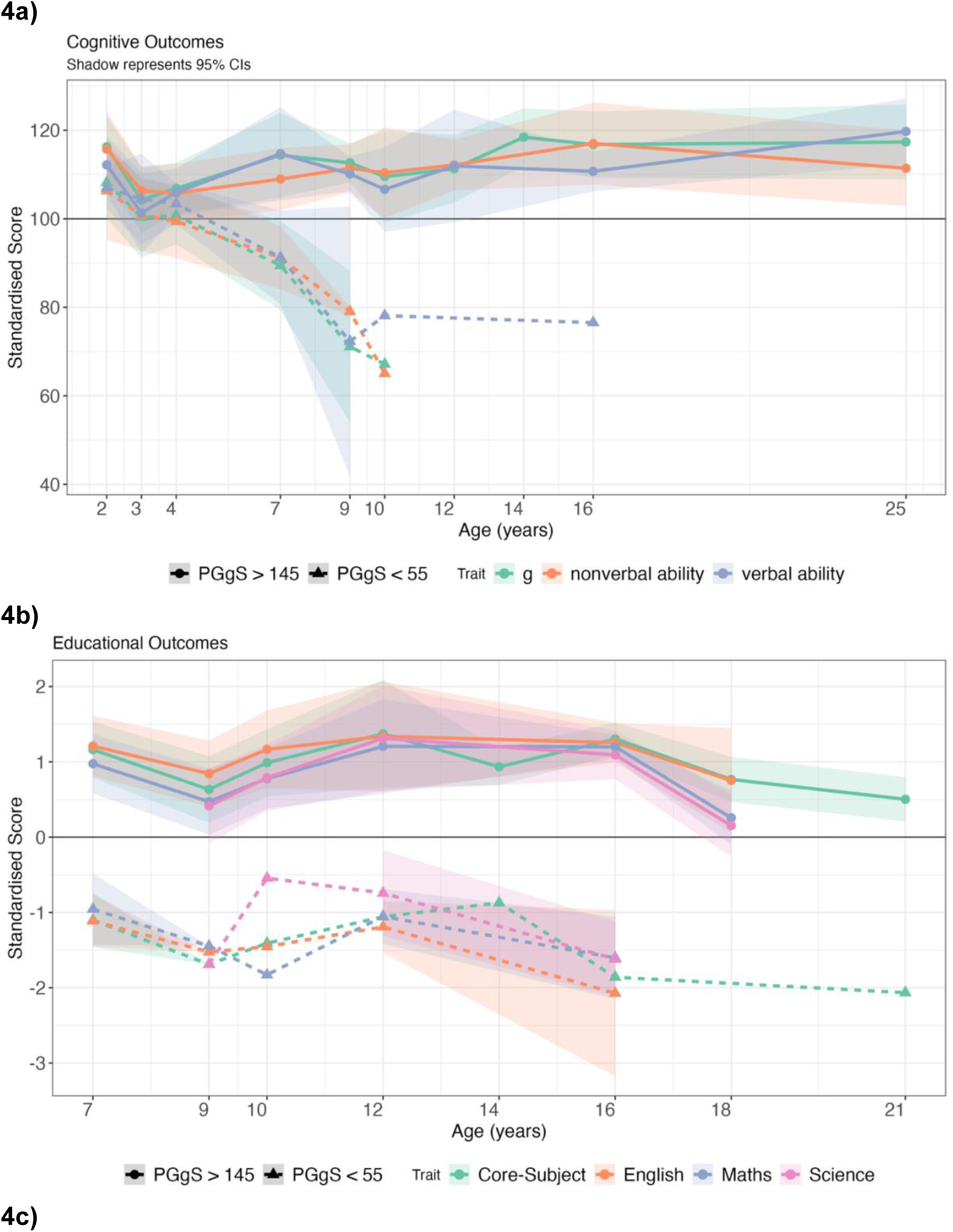
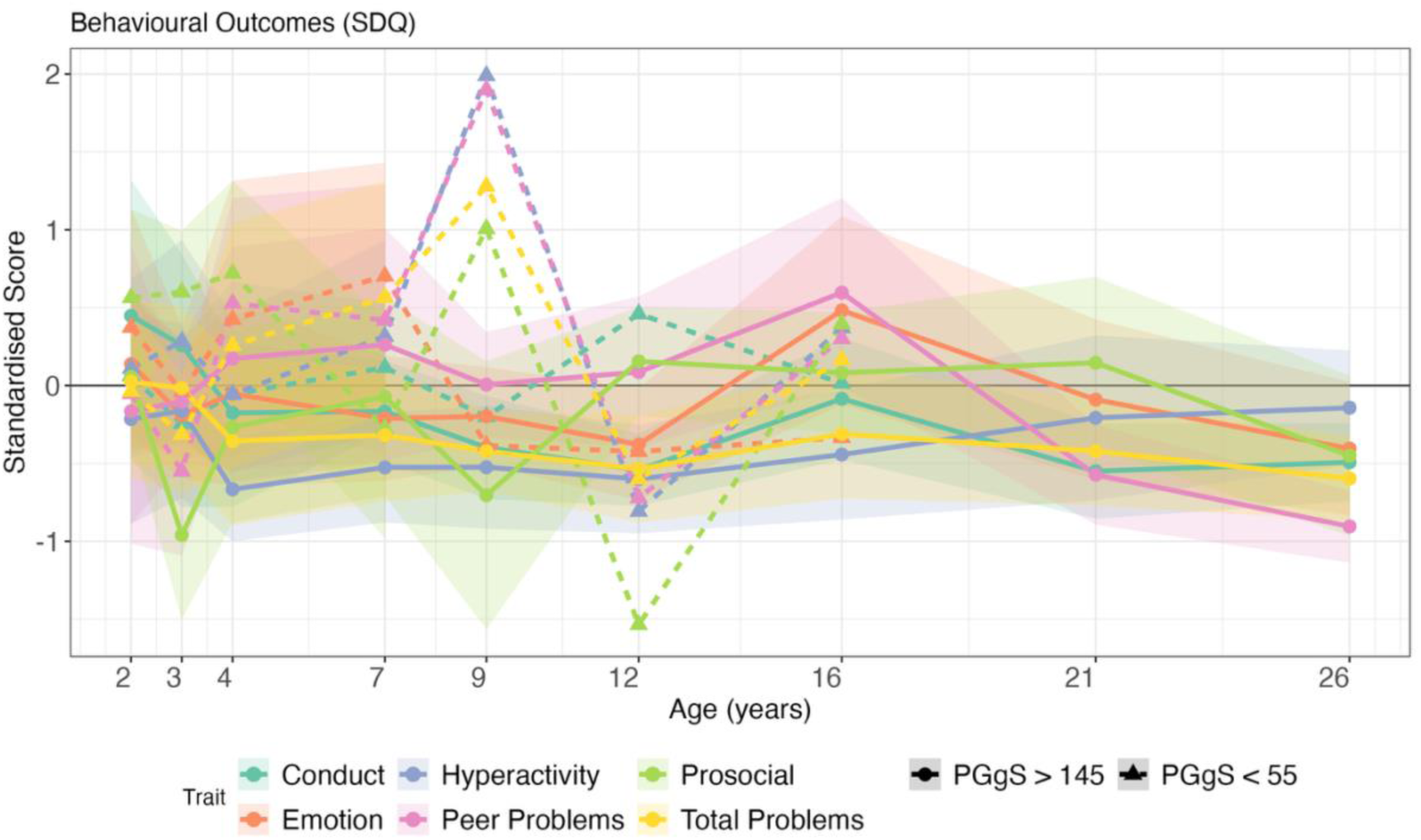
Mean developmental trajectories for individuals with the highest and lowest polygenic g scores. Mean trajectories across development for individuals with polygenic g scores >145 and <55. **Panel a:** General cognitive ability (g) composite and domain-specific composites (nonverbal and verbal ability). **Panel b:** educational achievement outcomes including English, mathematics, science, and core-subject composite (sum of English, mathematics, and science). **Panel c:** Strengths and Difficulties Questionnaire (SDQ) five subscales and total problem score. All measures are standardised to mean = 0, SD = 1, except for cognitive ability measures (mean = 100, SD = 15). Shaded areas represent 95% confidence intervals; absence of confidence intervals indicates that only one individual was measured at that age. For phenotypes with multiple raters, one rater per age is used: parent ratings before age 18 and child/self-ratings at age 18 and older. The black horizontal line at y = 0 represents the population average. PGgS = polygenic g score

For cognitive abilities, both groups initially performed within one standard deviation (SD) of the population mean in toddlerhood and early childhood (Figure 4a). By middle childhood, the high polygenic score group had IQ-equivalent scores of about 110. The low polygenic score group had IQ-equivalent scores between 80 and 90 in middle childhood but experienced substantial attrition; by adulthood, only one individual remained in the study.

Educational achievement yielded similar developmental trends. The high polygenic score group performed about one SD above the population mean, while the low polygenic score group performed about one SD below the mean (Figure 4b). For behaviour problems, neither group differed significantly from the population mean or from each other (Figure 4c). Mean trajectory comparisons for the additional phenotypic groups are presented in Supplementary Figure S14.

### Polygenic g Score Prediction Among Subsamples

Polygenic g score predictions showed high consistency across sexes at each age, across ages, across developmental trajectories, across distributions, and among the extreme scorers. Estimates were typically within one to two standard errors of each other, well within the range of overlapping 95% confidence intervals. Cognitive composites and individual cognitive tests showed nearly identical predictions for males and females. Educational outcomes showed similar patterns for both sexes, with predictions increasing through high school and declining thereafter. For behaviour problems, predictions were also comparable between sexes, with only occasional deviations. For example, the polygenic g score predicted life satisfaction at age 16 more strongly for females (β = 0.19) than for males (0.05). On average, the differences in standardised beta coefficients between sexes were smaller than 0.01.

Developmental trajectories were also largely similar between sexes, particularly for cognitive and educational outcomes. Occasional differences emerged for behaviour measures. For instance, for the ARBQ shyness subscale, females showed relatively stable differences across polygenic score groups across ages, whereas for males, these group differences gradually attenuated over time, with all groups converging toward the cohort average.

Zygosity comparisons likewise revealed minimal differences. Predictions were highly similar for monozygotic and dizygotic twins, with an average difference in standardised beta coefficients of only 0.01 across all phenotypes. The largest observed difference was for A-level technology grades at age 16 (0.02 for monozygotic twins vs 0.16 for dizygotic twins), though such differences were rare exceptions.

## Discussion

Polygenic scores derived from adult intelligence and educational attainment continue to be the strongest predictors of cognitive development. By combining g-PGS and EA-PGS, we created an adult polygenic g score to chart cognitive development from toddlerhood (age 2) to early adulthood (age 26). Our polygenic g score improved prediction from 9% variance explained using EA-PGS alone to 12% for g at age 25. Prediction was even higher when estimated using an error-free structural equation model with cross-time latent factors, reaching 15% for g. Additionally, for educational achievement, the polygenic g score predicted up to 18% of variance at age 16, and up to 3% for behaviour problems and anthropometric outcomes.

The polygenic g score predicts cognitive outcomes as early as age 2 (β_standardised_ = 0.03 to 0.05). Although the prediction was largely negligible in toddlerhood and early childhood, it increased steadily from childhood to early adulthood, peaking at β_standardised_ = 0.36 at age 25, supporting our first two hypotheses on the significant and increased polygenic score prediction with age for cognitive abilities. These age-related increases in PGS prediction are consistent with both twin and genomic studies. For example, twin heritability estimated in TEDS increased from 41% at age 9 to 66% at age 17 (Haworth et al., 2010). Another study extended the finding and showed that cognitive heritability remains stable from young adulthood into late midlife (Lyons et al., 2017). A similar pattern was identified in genomic studies using PGS (Allegrini et al., 2019; Gustavson et al., 2025). This pattern is known as the Wilson effect, which refers to the increasing heritability of IQ with age (Bouchard, 2013).

Several mechanisms help explain this phenomenon. First, discovery GWA studies of cognitive abilities are conducted in adults; thus, the SNP effect estimates in these GWA studies more closely match the genetic architecture of cognitive outcomes in later developmental stages than in toddlerhood or early childhood. Second, phenotypic measures are less stable and reliably assessed at younger ages, which limits how well the polygenic g score can predict cognitive abilities in early childhood, as the phenotypic correlation represents the upper bound of genetic prediction. For example, the phenotypic correlation of intelligence between ages 2 and 25 was modest (r = 0.11), but increased substantially by late adolescence, reaching r = 0.65 between ages 16 and 25. Nevertheless, substantial genetic continuity across the lifespan supports our approach of using an adult-derived polygenic g score to chart cognitive development, as noted in the Introduction. In addition, a genome-wide association study on general cognitive function reported a genetic correlation of 0.71 between childhood and older ages (Hill et al., 2016) and Savage et al. (2018) also found high genetic correlations (> 0.62) across childhood, young adulthood, and adulthood. Third, while developmental changes in genetic architecture play a role, increasing active and evocative gene–environment correlation continues contributing to the magnification of genetic effects over time, as individuals increasingly select, evoke, and create environments that align with their genetic propensities (Plomin, 1994).

Despite potential constraints in early-age measurement, the polygenic g score is still one of the best predictors of cognitive development. By combining g-PGS and EA-PGS, we introduced additional genetic signals relevant to cognitive abilities (Plomin & von Stumm, 2018). We achieved even higher predictive power within an error-free SEM framework of g, verbal ability, and nonverbal ability with 15%, 16%, and 13% variance explained, respectively.

However, in practice, extracting cross-time latent factor scores did not outperform the strongest predictions from later-age observed phenotypes. Although one might expect the cross-time factor to perform best by stripping away age-specific noise, this was not the case here. As our participants age, their genetic architecture increasingly aligns with the adult cohorts used in the original GWAS, which likely contributes more to the increase in prediction accuracy at later ages than the removal of age-specific noise by cross-time latent factors.

The increasing alignment between participants’ genetic architecture and the adult GWAS reference also helps contextualise the challenge of measurement invariance. Because cognitive assessments inevitably change as children grow, methodological variance and developmental progression are inextricably linked. However, the steady rise in predictive power as participants approach adulthood strongly suggests that genuine developmental shifts may be the primary driver, rather than a mere measurement artefact of using different tests.

In addition, our latent growth curve models captured both baseline differences and developmental changes. Children with higher polygenic g scores showed higher baseline cognitive performance and a modest but statistically significant increase in relative growth rate, consistent with increasing gene-environment correlation over time. These findings illustrate that the influence of polygenic g score is not limited to predicting outcomes at isolated ages but also characterises the overall developmental trajectory of cognitive abilities from early childhood to adulthood.

We next examined educational achievement, behaviour problems, and anthropometric measures. Polygenic g score predicted most trait domains, supporting the third hypothesis. The strongest prediction was for educational achievement, consistent with previous research (Wilding et al., 2024). Prediction of educational achievement increased until the end of secondary school, peaking at 18% for GCSE grades, and then declined for subsequent educational milestones, reaching approximately 7% for years of schooling (educational attainment) at age 26. Similar developmental patterns were reported in independent research (Kvalvik et al., 2025). While a polygenic g score captures the variance in standardised, fine-grained cognitive batteries and school test scores, educational milestones are increasingly influenced by non-cognitive abilities and environmental variance (Malanchini et al., 2024), which explains the attenuated predictive power of polygenic g score after age 16.

Moreover, as with cognitive traits, higher polygenic g scores are associated with higher baseline educational achievement and steeper relative gain. In contrast, predictions for behaviour problems and anthropometric outcomes were negligible to weak and showed less consistency. Behaviour problems showed negative baseline associations and small positive slope estimates, indicating that children with higher polygenic g scores began with fewer behaviour problems relative to their peers, but exhibited a slight upward shift in relative ranking over time. This pattern might reflect regression to the cohort mean and artefacts from standardisation across varying instruments, rather than substantive developmental change. No consistent sex differences were observed.

In line with our fourth hypothesis, prediction across the distribution of polygenic g scores was linear. The absence of significant quadratic effects indicates that individuals at the end of the distribution differ quantitatively rather than qualitatively from the rest of the sample. At the distributional extremes of the polygenic g scores (+/–three standard deviations), individuals who were three standard deviations above the mean showed, on average, around one to two standard deviations advantage in cognitive and educational outcomes in adulthood. Behaviour problems and anthropometric outcomes showed almost no meaningful differences between extremes.

Finally, consistent with our fifth hypothesis, we found no evidence for sex differences in the prediction of the polygenic g score across outcomes.

Our study has several limitations. First, we applied European-derived GWAS weights to a British white cohort. While this shared genetic background reduced confounding and made our estimates robust, it limits the generalisability of our findings to other populations with differing allele frequencies and environmental contexts.

Second, the weaker prediction for nonverbal relative to verbal ability reflects the composition of our polygenic g score. By incorporating EA-PGS, our composite captures a substantial amount of education-specific genetic variance, which is more strongly associated with language and reading skills than with mathematical or spatial outcomes (Procopio et al., 2025). Although the UK Biobank verbal numerical reasoning test contains slightly more nonverbal/numerical items, these questions are predominantly text-based word problems (e.g., applied arithmetic and age comparisons). The tasks categorised as numerical reasoning rely heavily on reading comprehension (Genç et al., 2021). Both components of our composite therefore share a verbal bias. We nevertheless term the composite the ‘polygenic g score’ because its weights were derived empirically to maximise prediction of g, and because educational attainment is highly genetically correlated with g, suggesting that much of the predictive power of EA-PGS reflects shared genetic influences on cognitive abilities. Moreover, we acknowledge that the EA-PGS also derives predictive power from noncognitive skills (Demange et al., 2021), and caution that our composite should be interpreted as a genetic predictor of adult g rather than a genetically specific measure of g.

Third, our g composite was constructed by averaging verbal and nonverbal abilities. Although this composite performed well, hierarchical phenotypic models would allow us to estimate g more precisely from the underlying tests and could improve prediction. Fourth, discovery GWA studies rely on common variants. Recent evidence suggests that rare genetic variants contribute substantially to cognitive differences and could improve prediction with whole-genome sequencing (Wainschtein et al., 2025).

Fifth, caution is needed when applying polygenic g scores for genetic explanation beyond prediction (Plomin & von Stumm, 2022). Recent debates about environmental confounding in polygenic prediction reflect the distinction between causal explanation and prediction. Whereas causal models aim to isolate biological mechanisms, our predictive framework intentionally captures all correlated pathways to maximise explained variance. Accordingly, our beta estimates represent an upper bound of predictive accuracy rather than so-called ‘direct genetic effects’ (Lin et al., 2025).

As twin analyses suggest increased heritability of intelligence and cognitive abilities from childhood to young adulthood, molecular genetic studies have made use of this finding by focusing on the genetics of adults. While twin designs rely on phenotypic measurements at each developmental stage to estimate heritability, GWAS-based and PGS-based approaches can quantify genetic propensities from conception onward. After decades of research, we are now able to leverage adult molecular genetic findings in the reverse direction to chart child cognitive development as it is genetically associated with adult cognitive abilities. Our results show that prediction begins as early as age 2, increases steadily through development, and is especially powerful in predicting growth trajectories. Importantly, this predictive utility does not depend on a causal explanation of individual SNP effects, but rather on the aggregation of genetic signals associated with the phenotype (Lin et al., 2025; Plomin & von Stumm, 2022).

Together, we show that combining two highly predictive polygenic scores yields substantial insights into cognitive developmental trajectories from toddlerhood and early childhood to early adulthood. Alongside other early-life predictors, polygenic scores fixed at conception continue to be one of the strongest predictors available for long-term cognitive development.

## Acknowledgments

We gratefully acknowledge the ongoing contribution of the participants in the Twins Early Development Study (TEDS) and their families. TEDS is supported by the UK Medical Research Council (MR/V012878/1 and previously MR/M021475/1). Y.L. is supported by a KCL-CSC joint scholarship awarded by King’s College London and the China Scholarship Council.

For the purposes of open access, the author has applied a Creative Commons Attribution (CC BY) license to any Accepted Author Manuscript version arising from this submission.

