## Supplementary Figures for "Charting the cognitive development of children using adult ‘polygenic g scores’"

**Figure S1_1 Correlation Matrix for Cognitive Ability Composites**

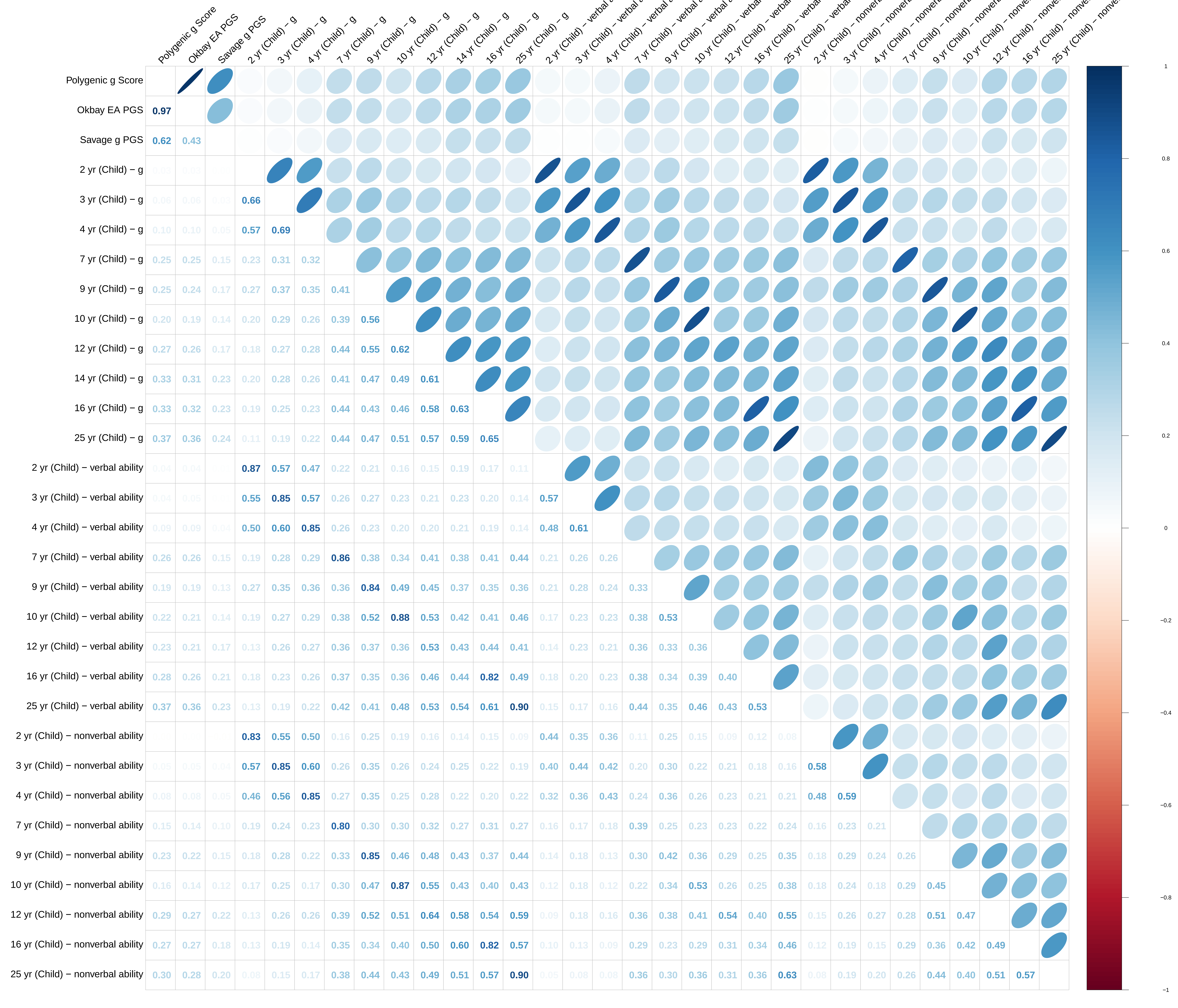

**Figure S1_2 Correlation Matrix for General Cognitive Ability (g)**

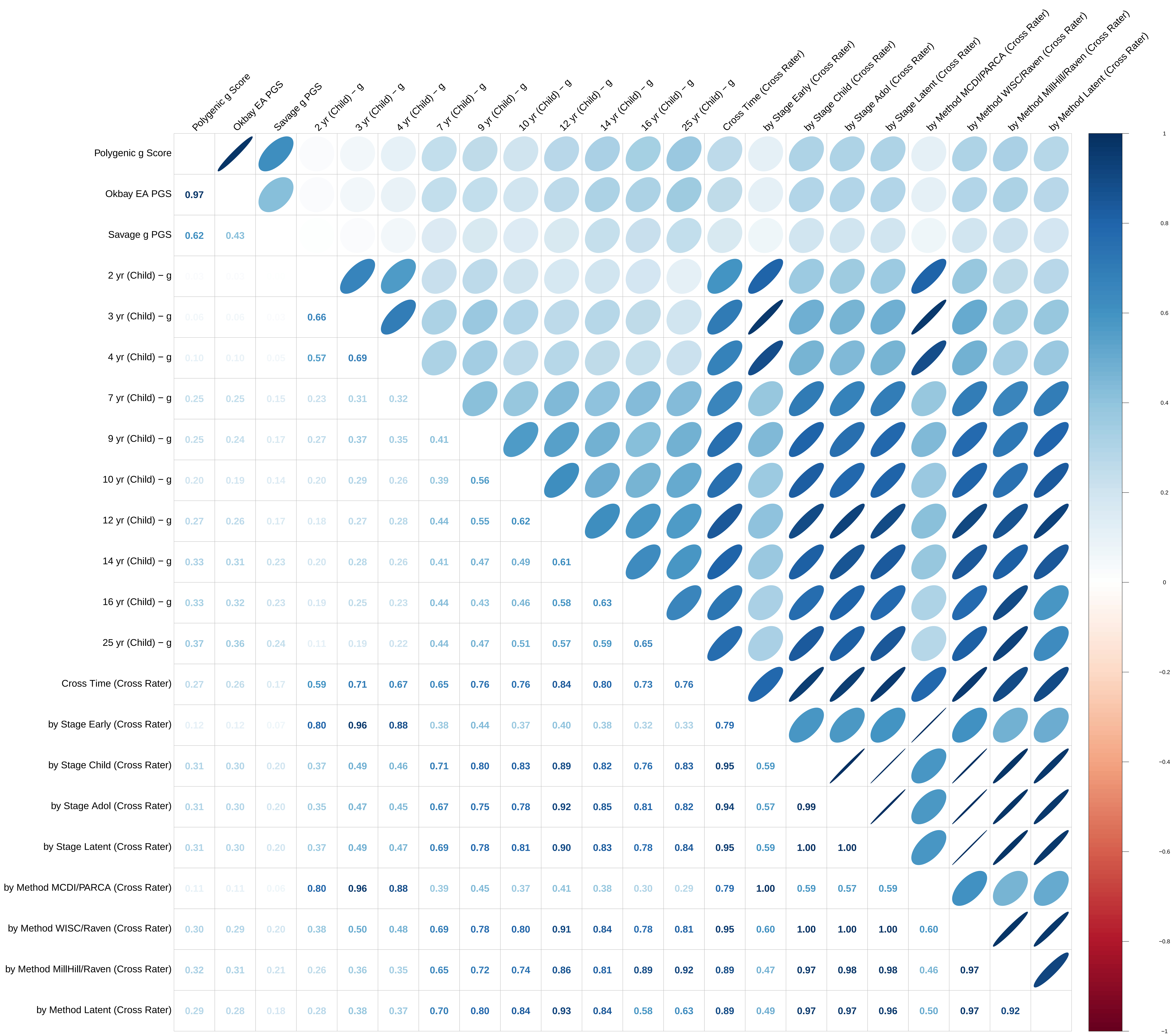

**Figure S1_3 Correlation Matrix for Verbal Tests, Composites, and Latent Factors**

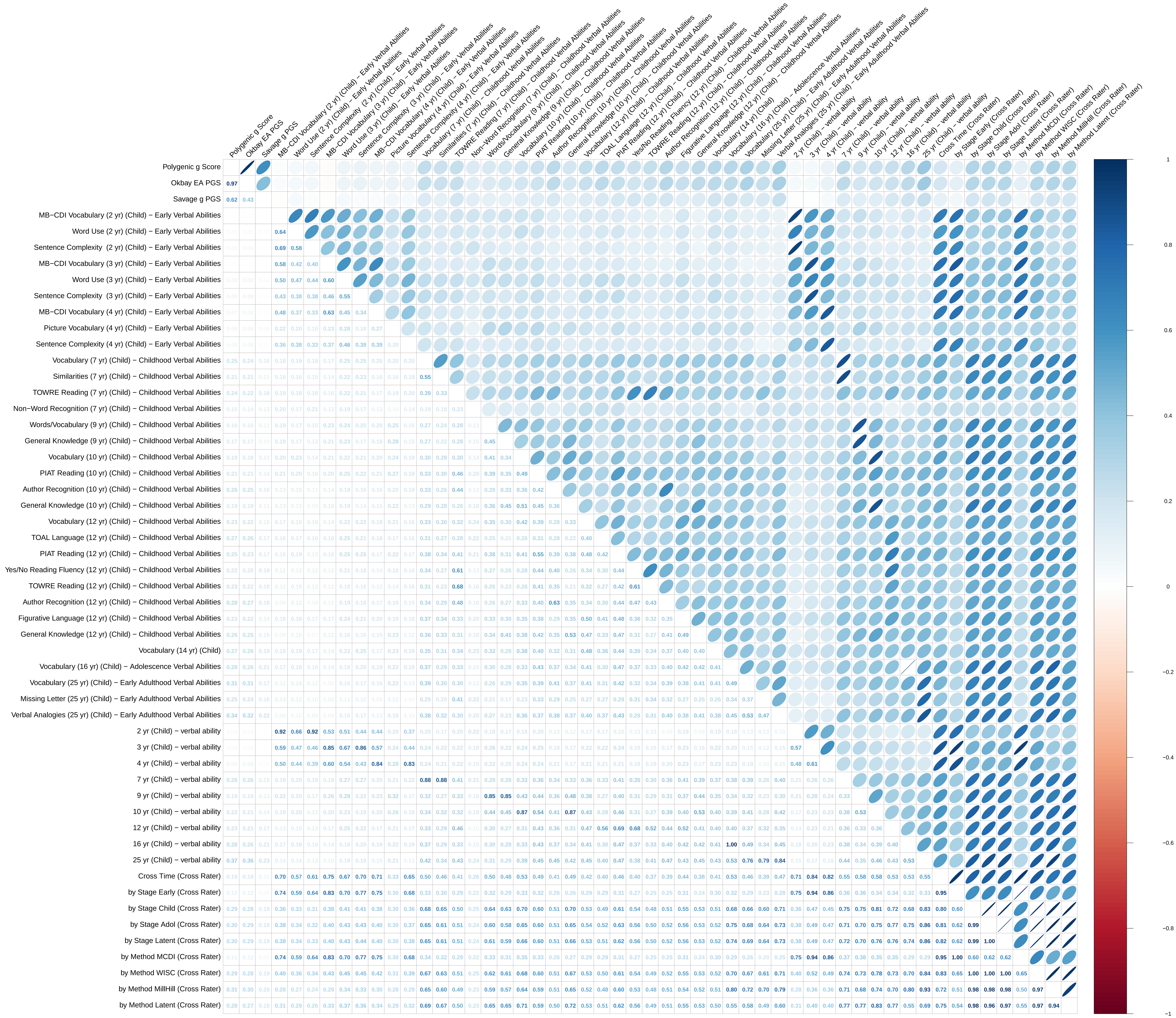

**Figure S1_4 Correlation Matrix for Nonverbal Tests, Composites, and Latent Factors**

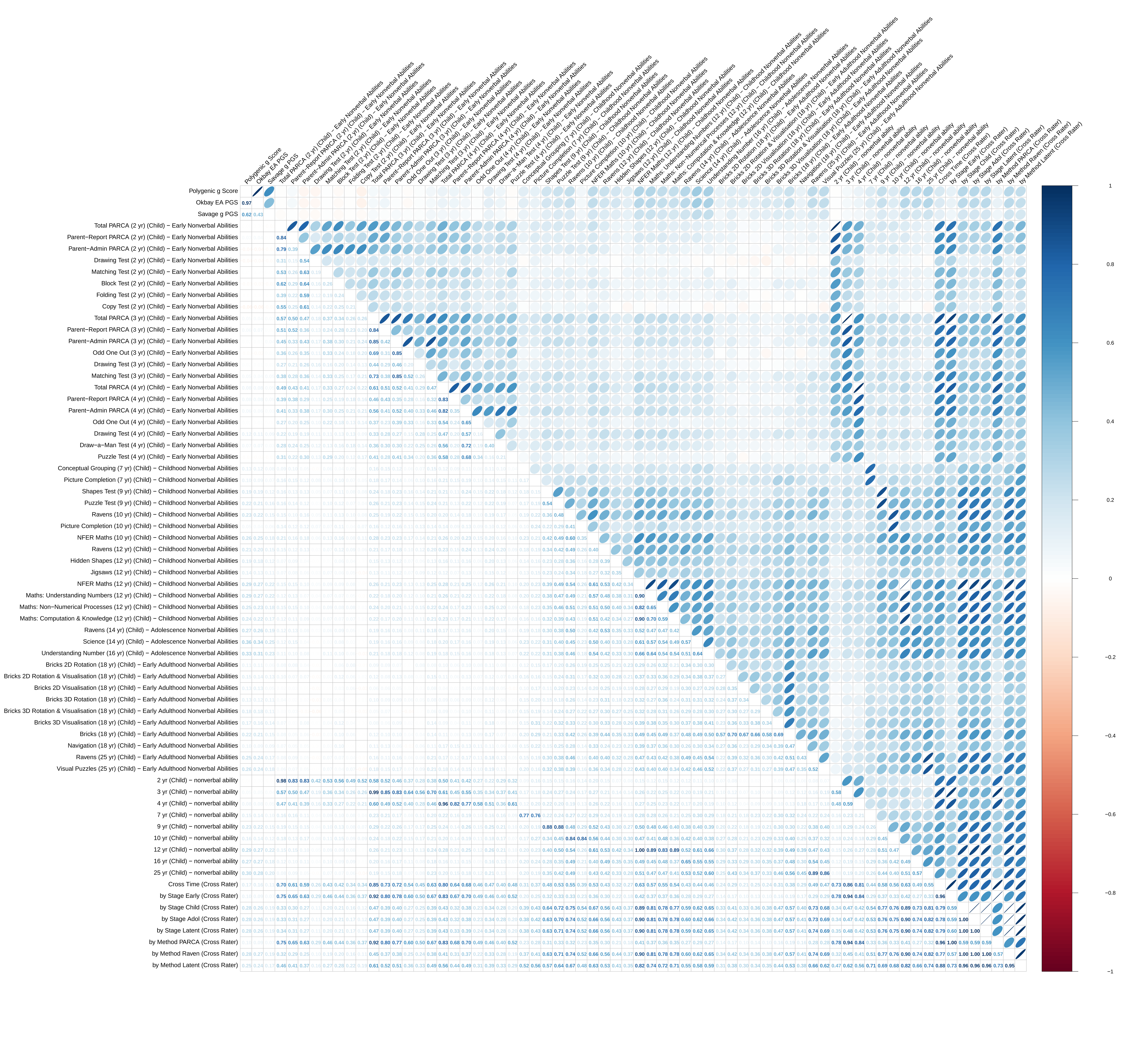

**Figure S1_5 Correlation Matrix for Educational Achievement and Attainment (with Latent Factors)**

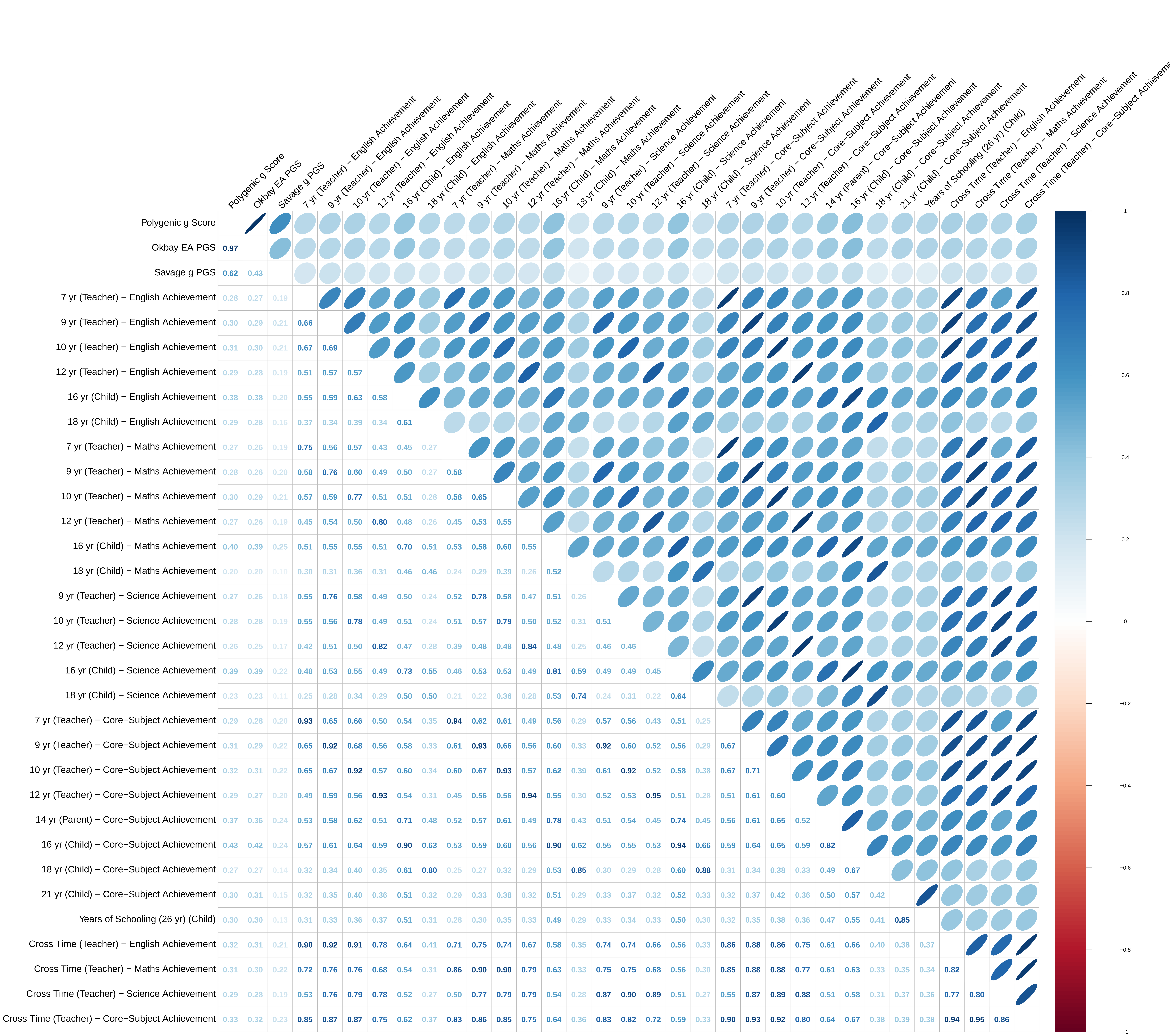

**Figure S1_6 Correlation Matrix for SDQ (with Latent Factors)**

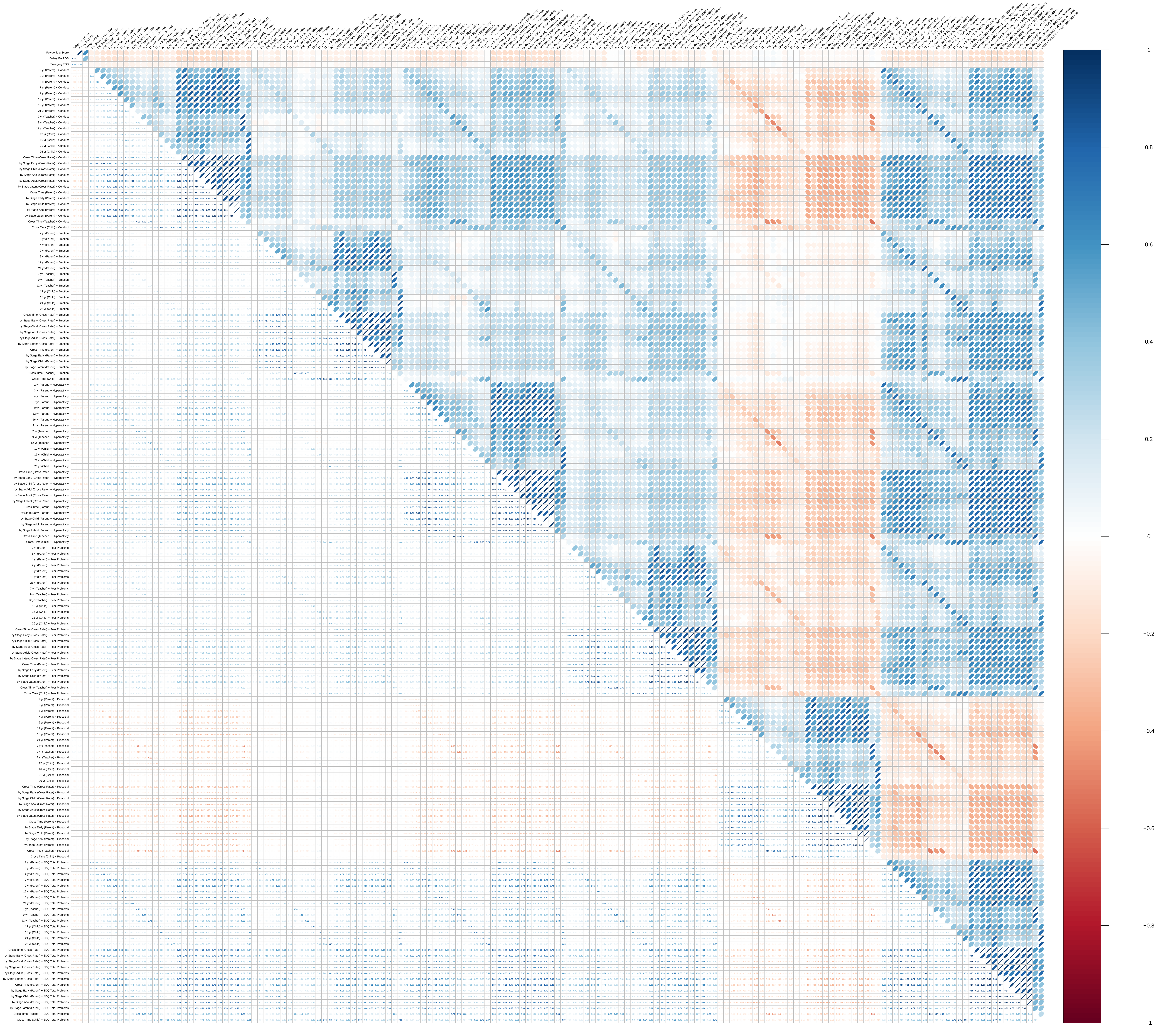

**Figure S1_7 Correlation Matrix for Anxiety Measures (with Latent Factors)**

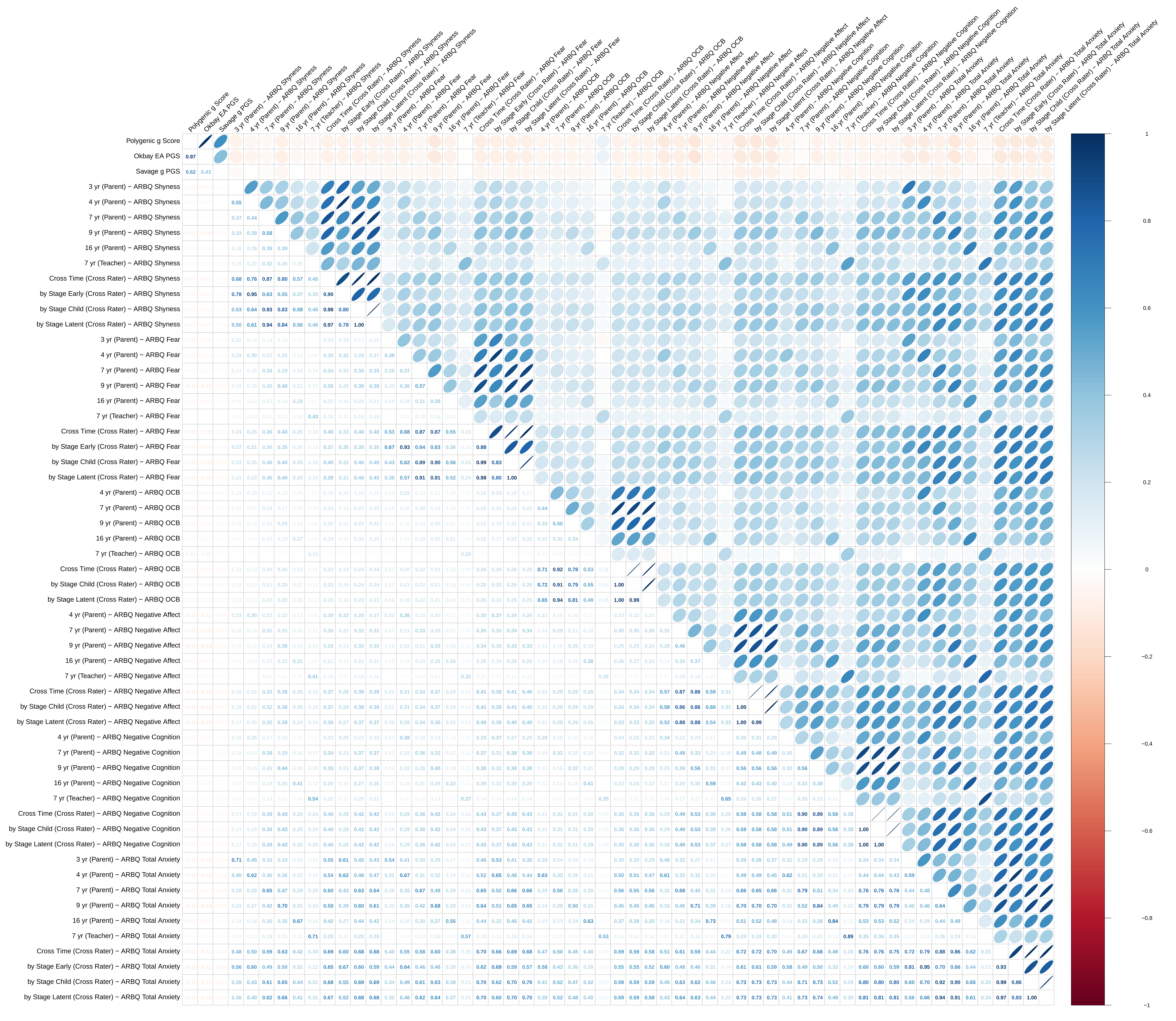

**Figure S1_8 Correlation Matrix for ADHD Measures (with Latent Factors)**

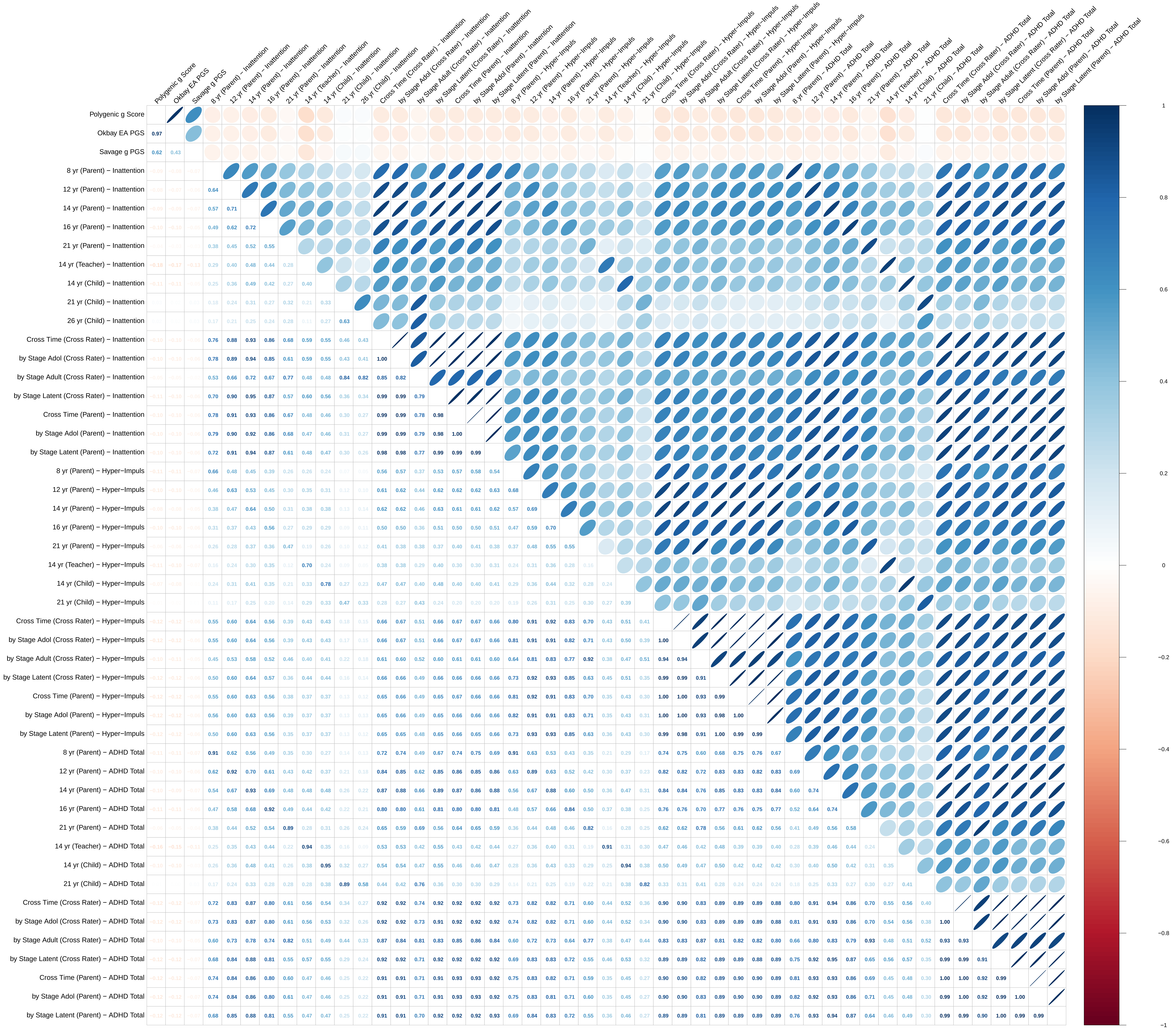

Figure S1_9 Correlation Matrix for Anthropometric Measures

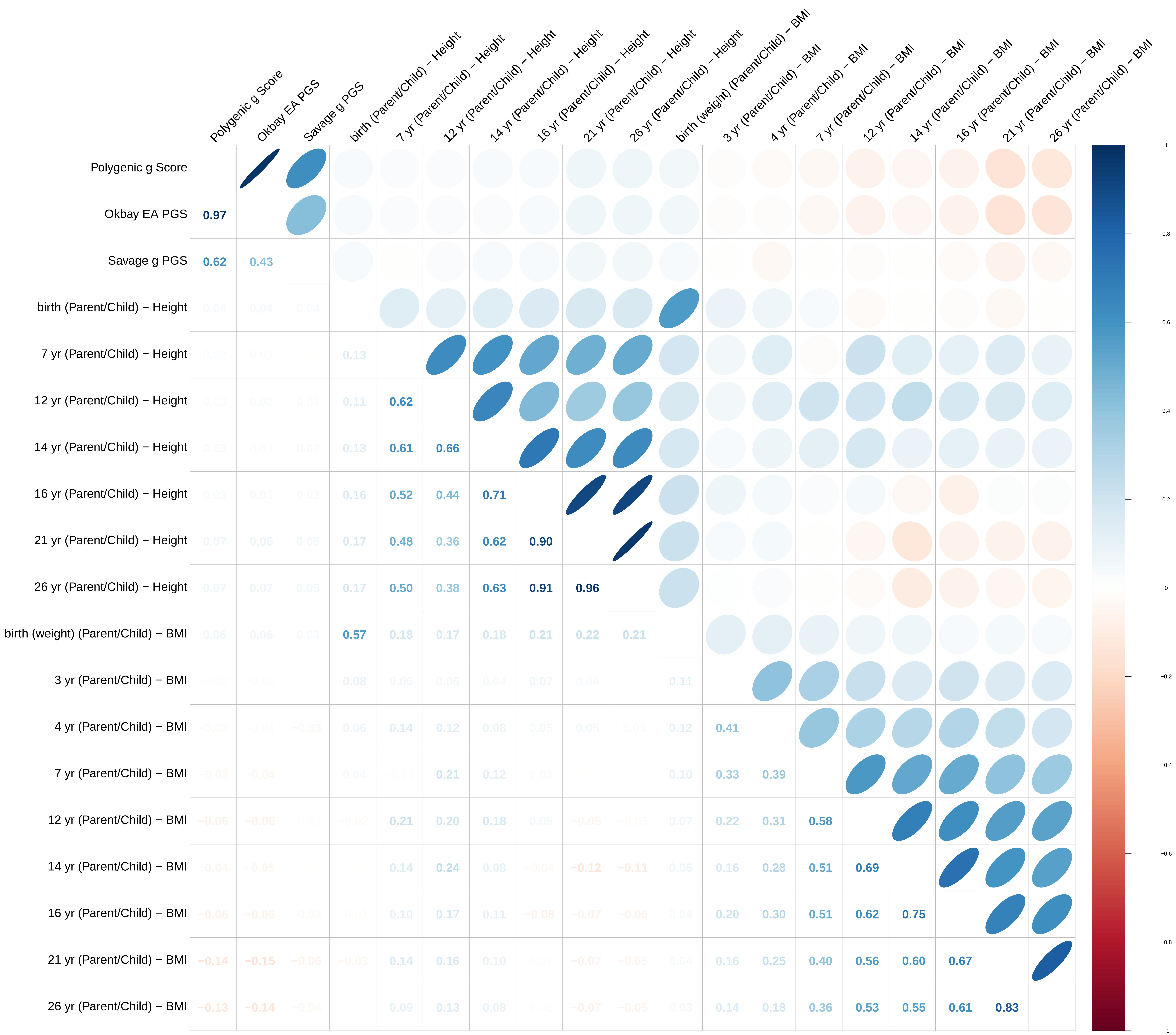

**Figure S8_1: General Cognitive Ability (g) − Overall**

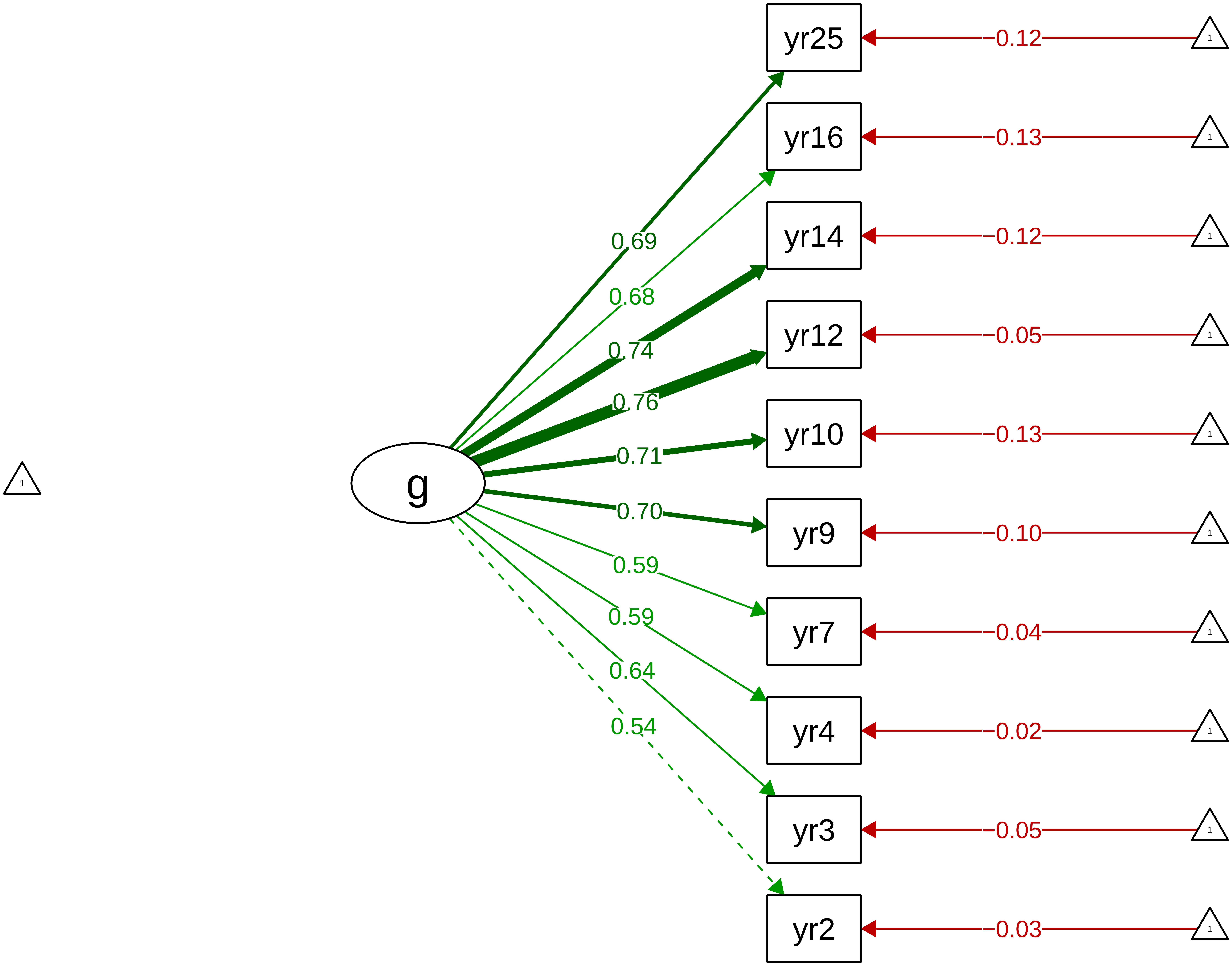

**Figure S8_2: General Cognitive Ability (g) − Stage**

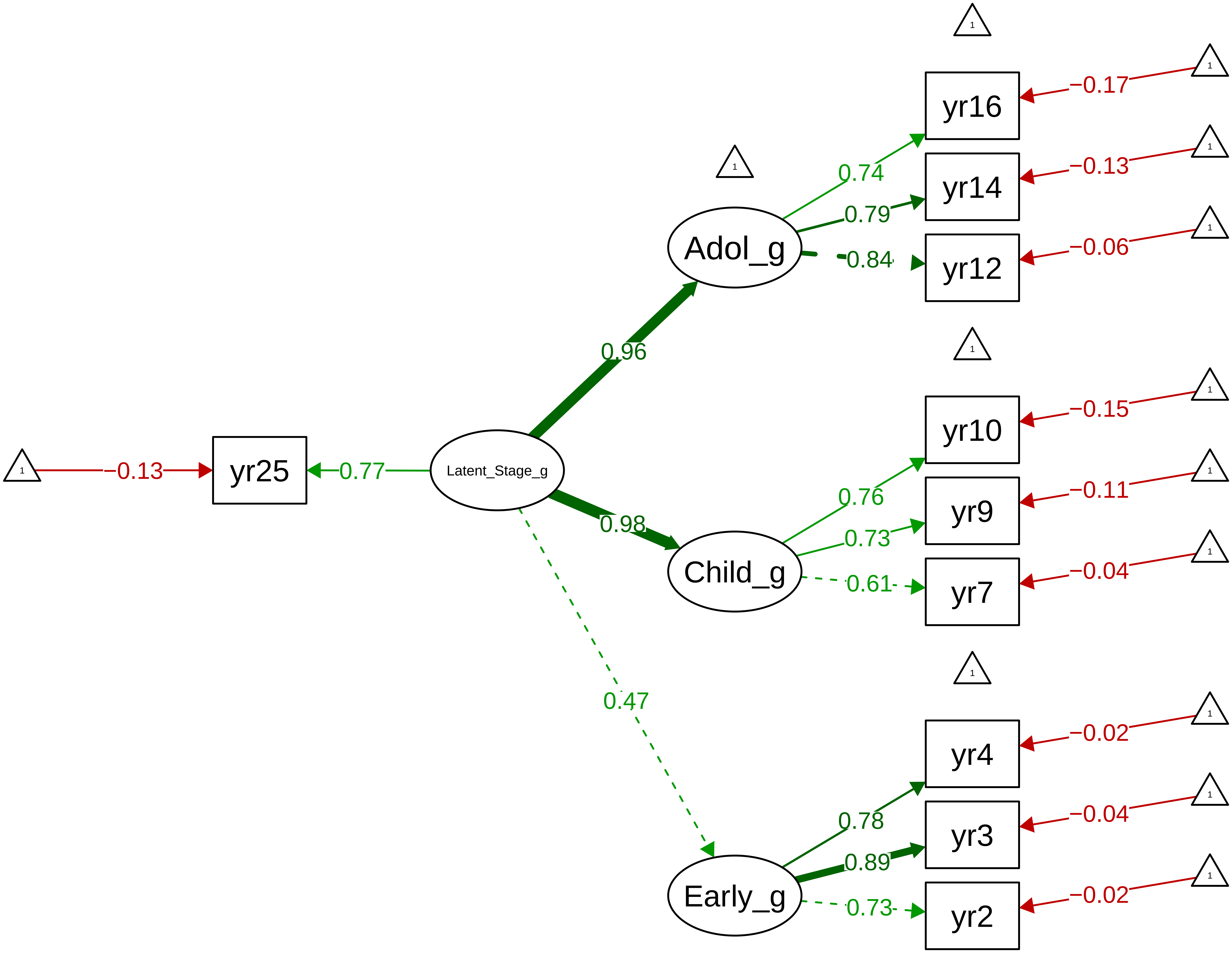

### Figure S8_3: General Cognitive Ability (g) − Method

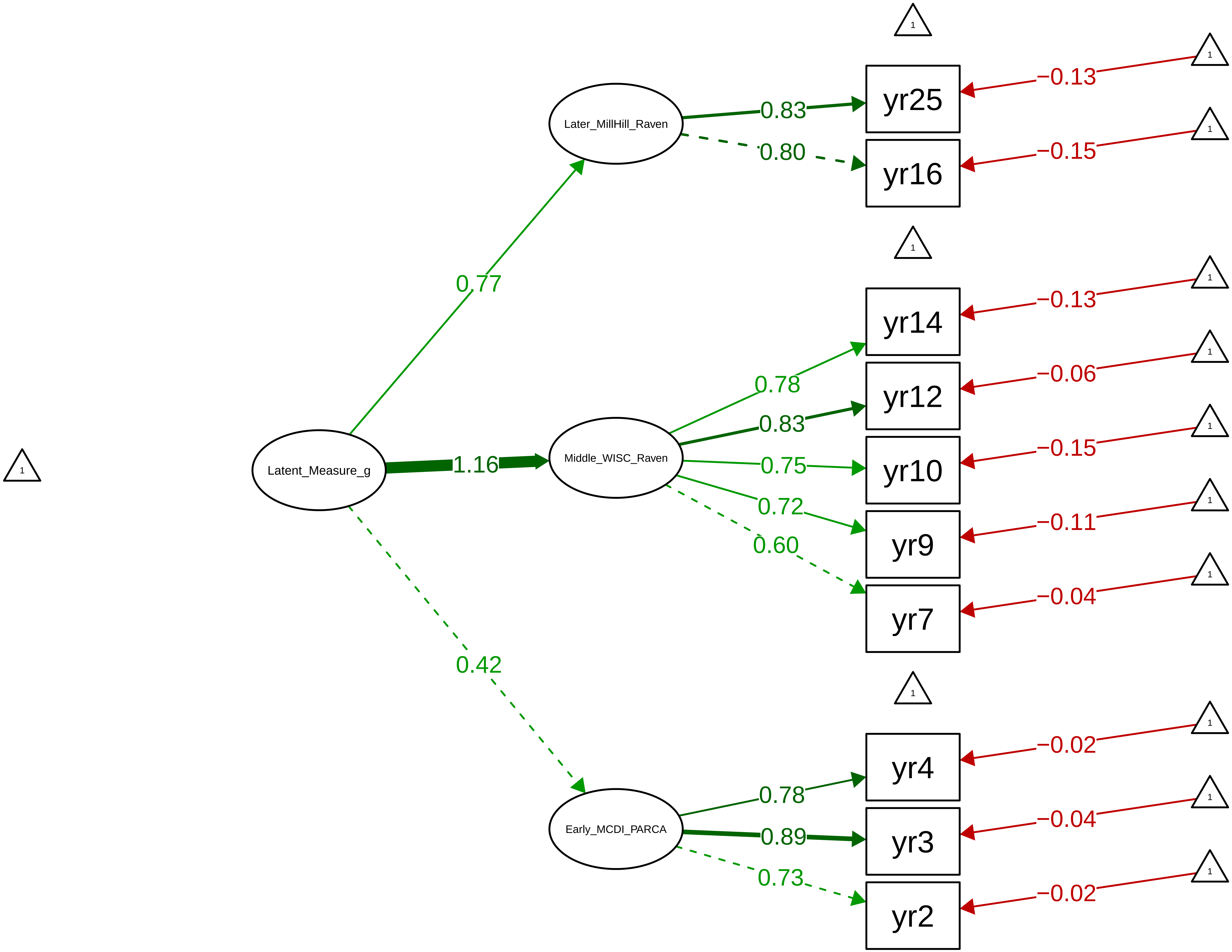

Figure S8_4: Verbal Ability − Overall

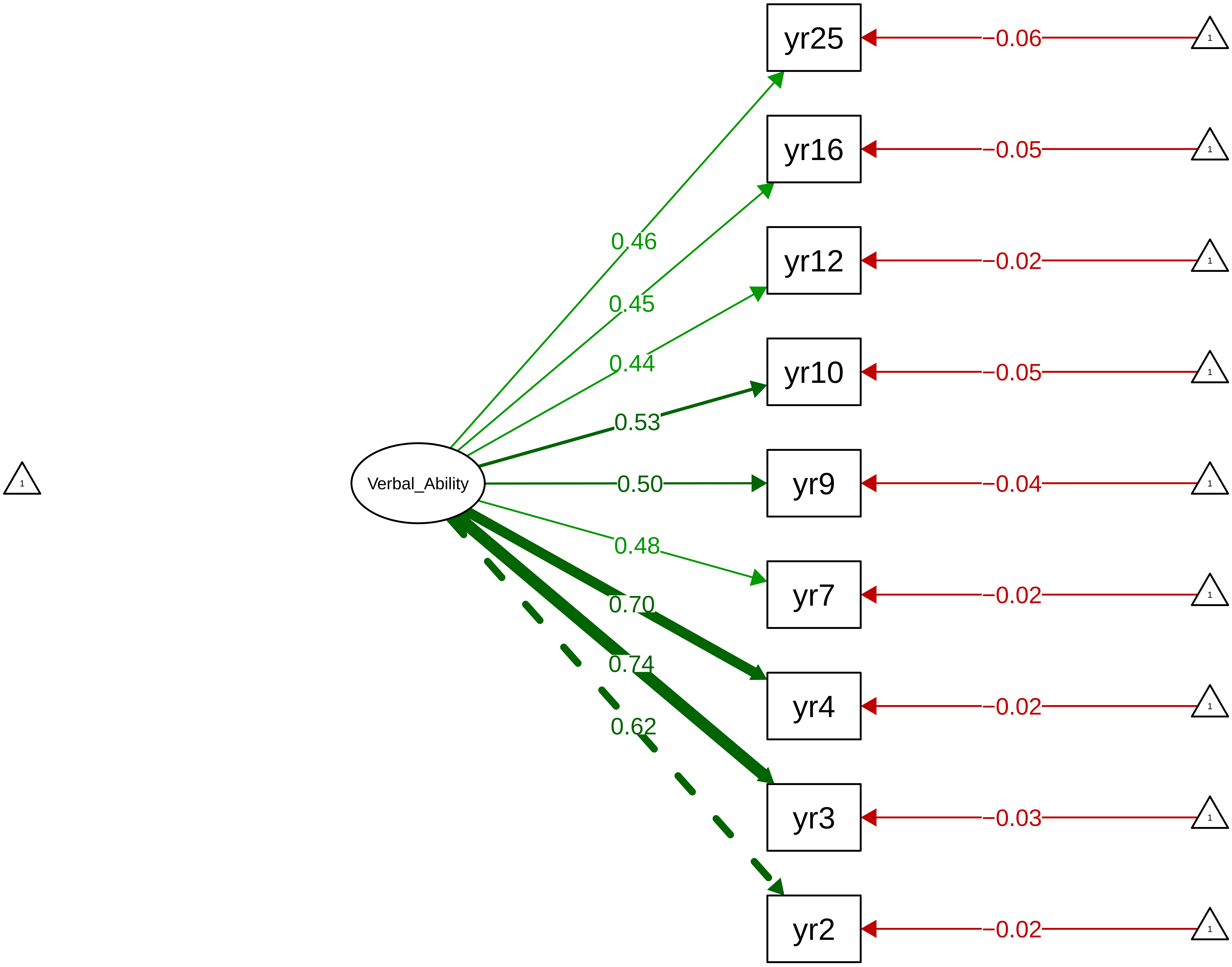

**Figure S8_5: Verbal Ability − Stage**

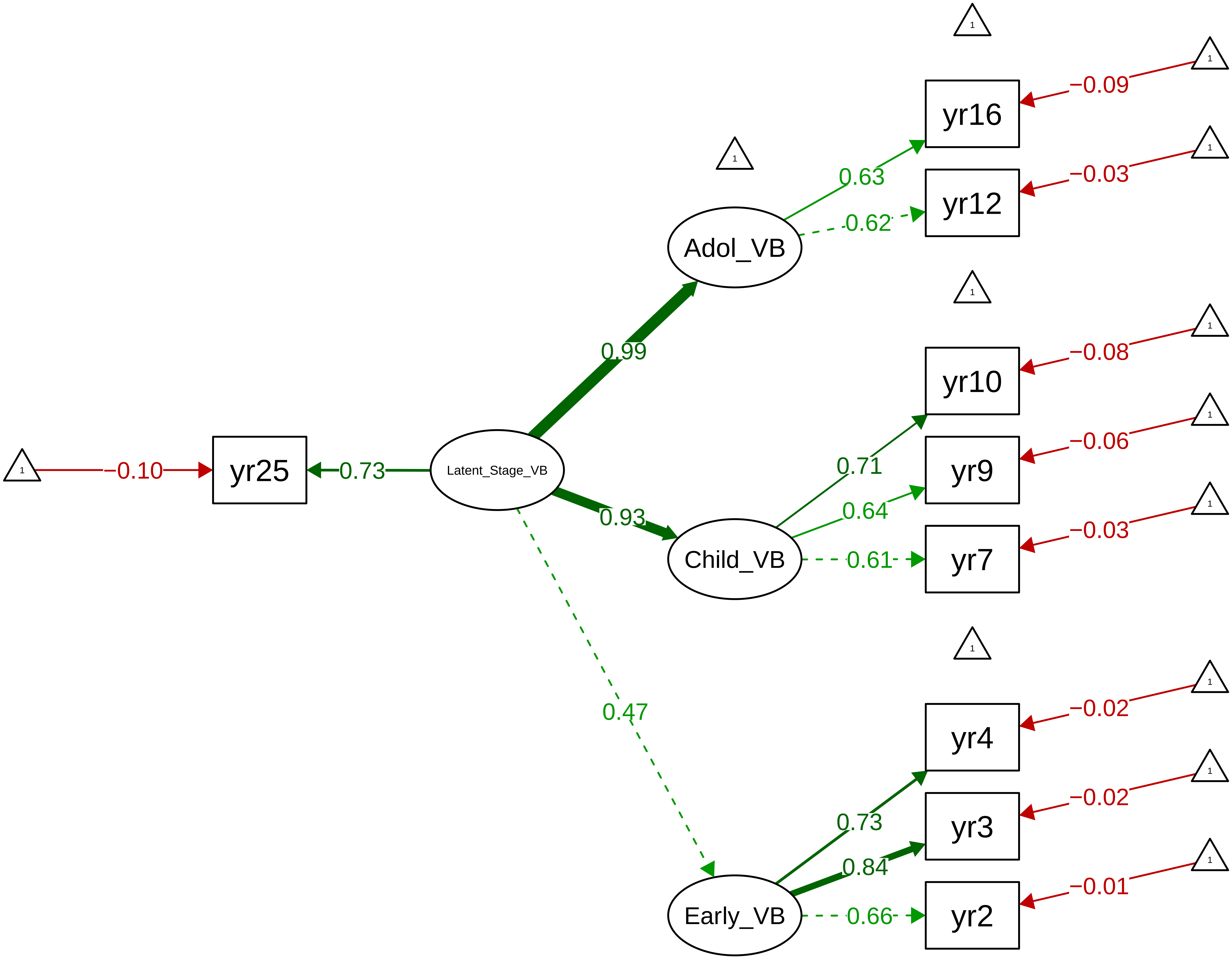

**Figure S8_6: Verbal Ability − Method**

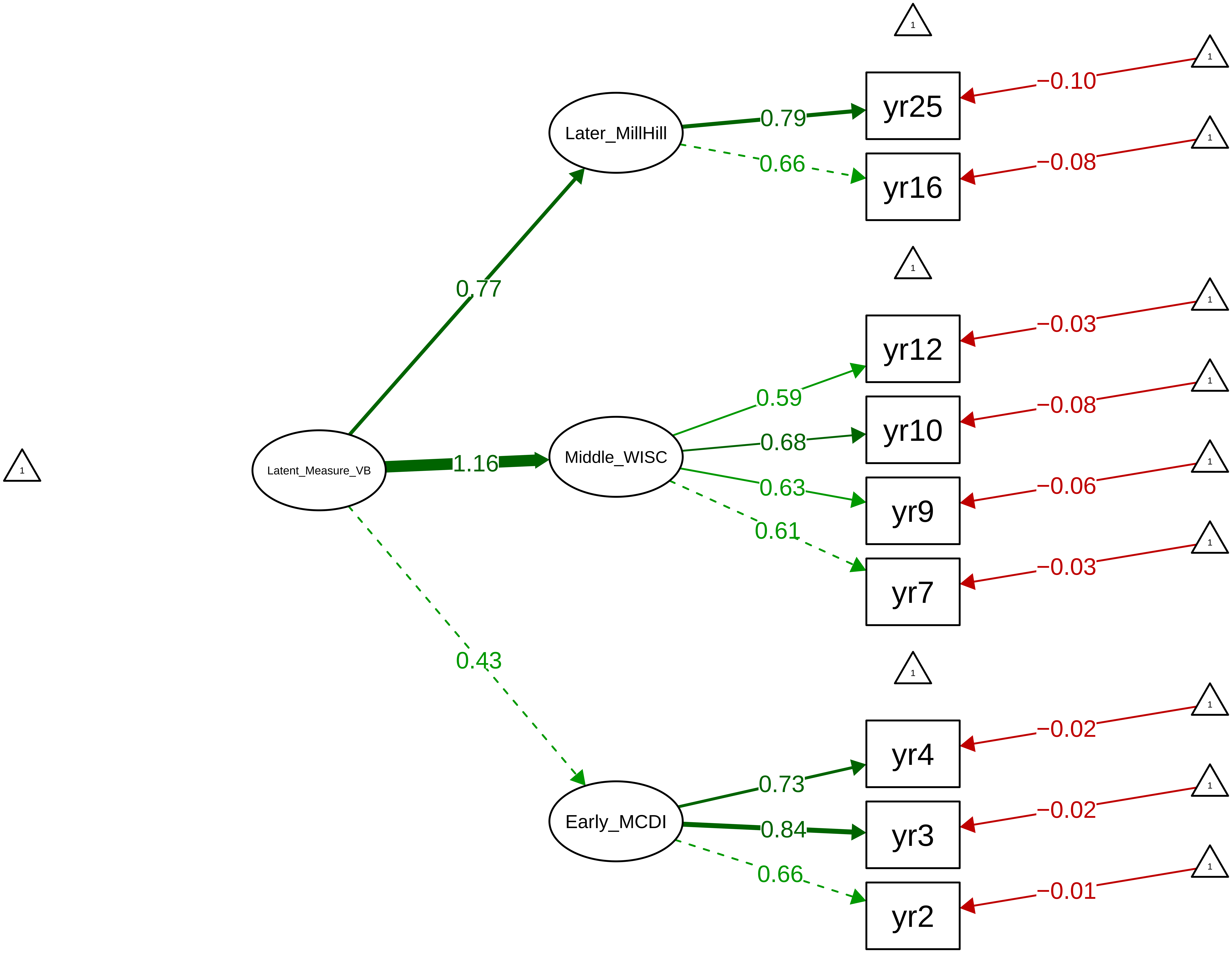

**Figure S8_7: Nonverbal Ability − Overall**

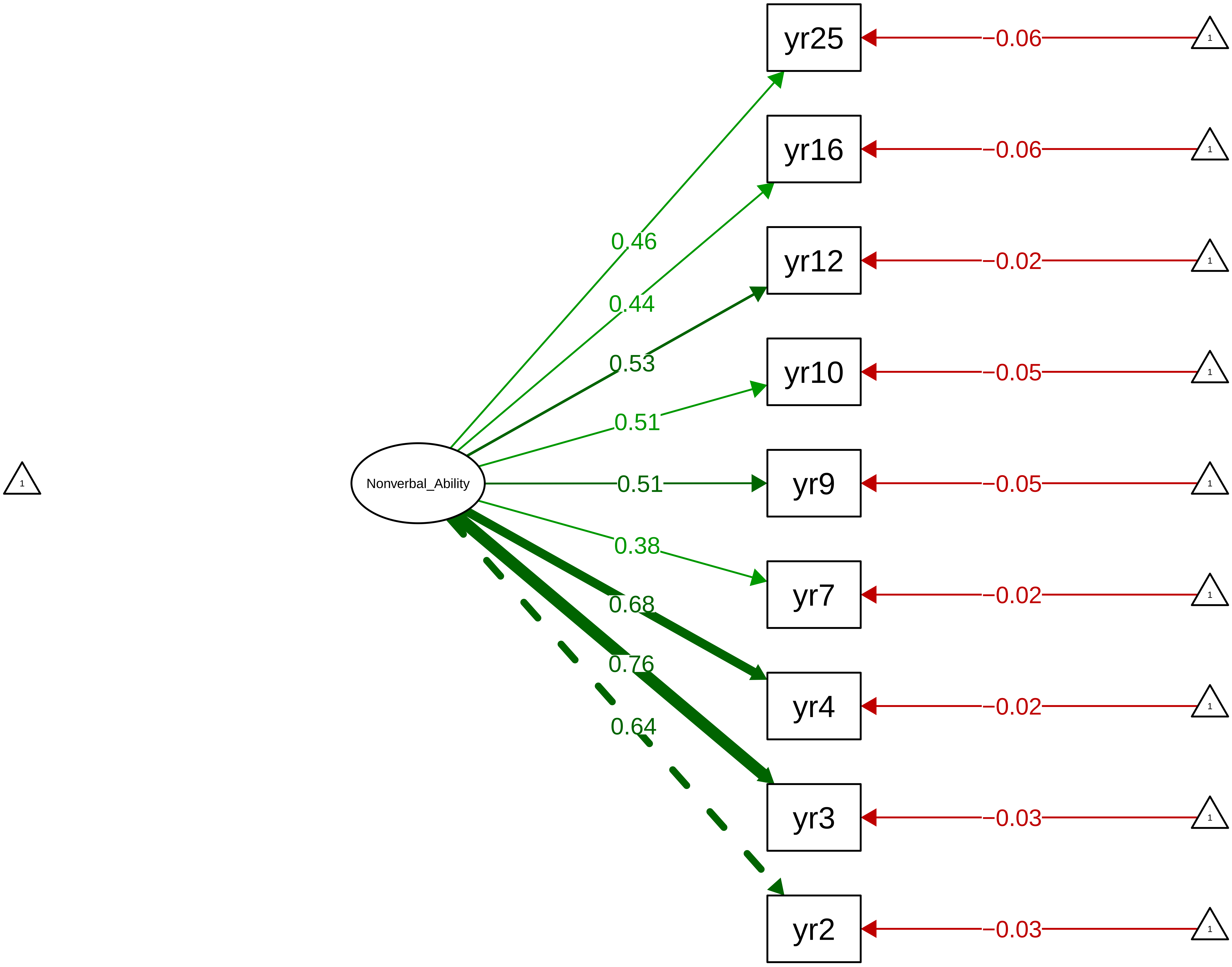

**Figure S8_8: Nonverbal Ability − Stage**

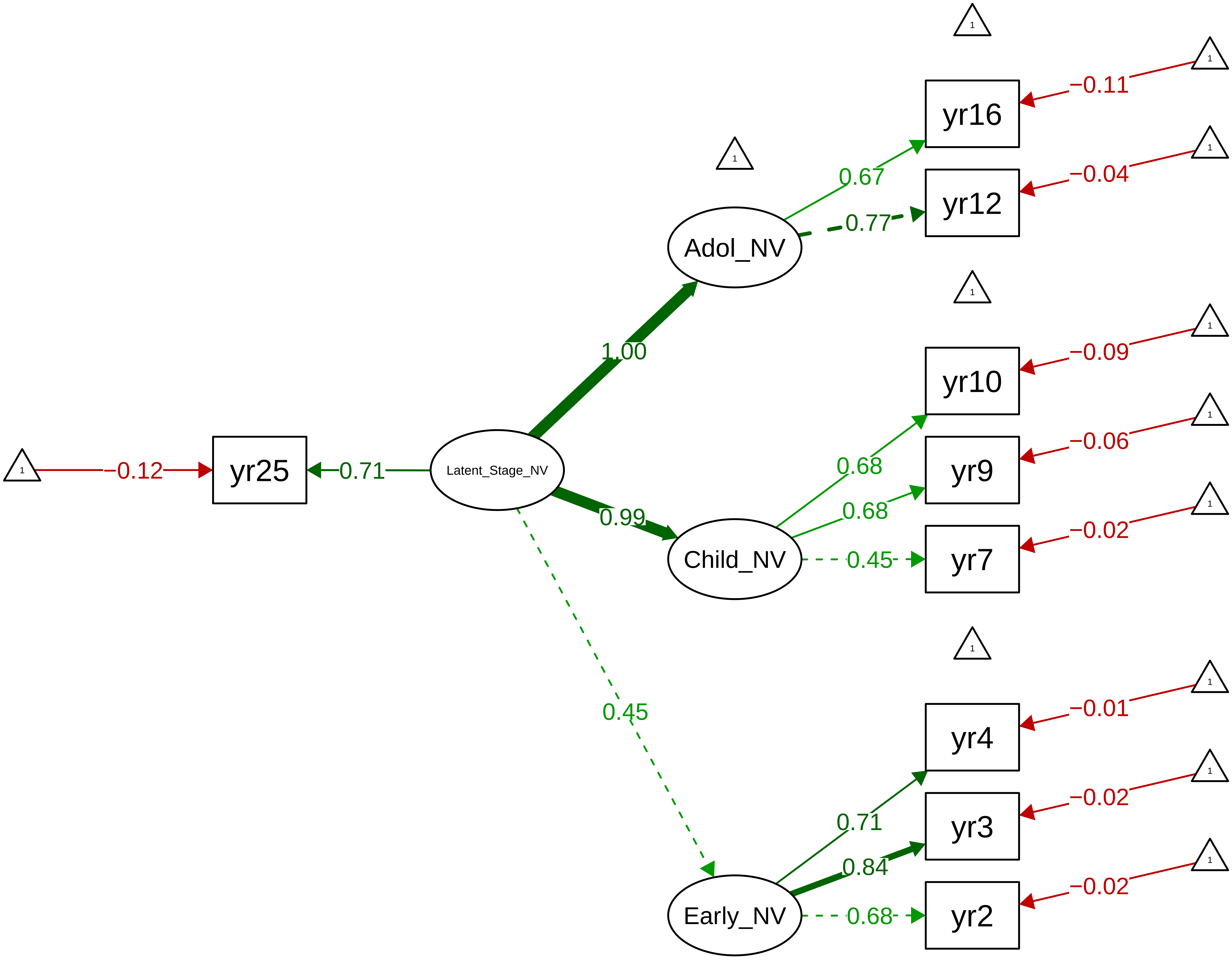

**Figure S8_9: Nonverbal Ability − Method**

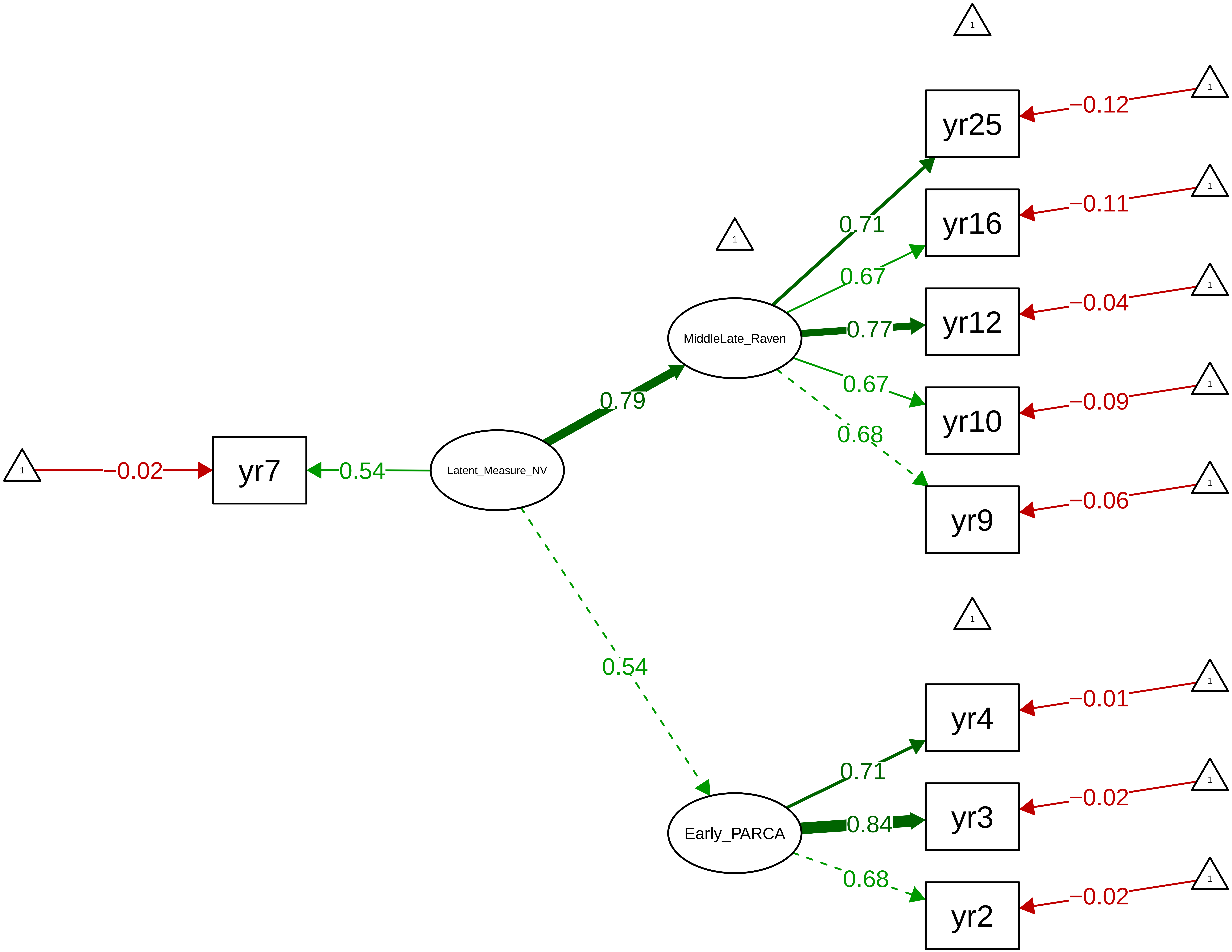

Figure S8_10: English Achievement Latent (Teacher 7−12)

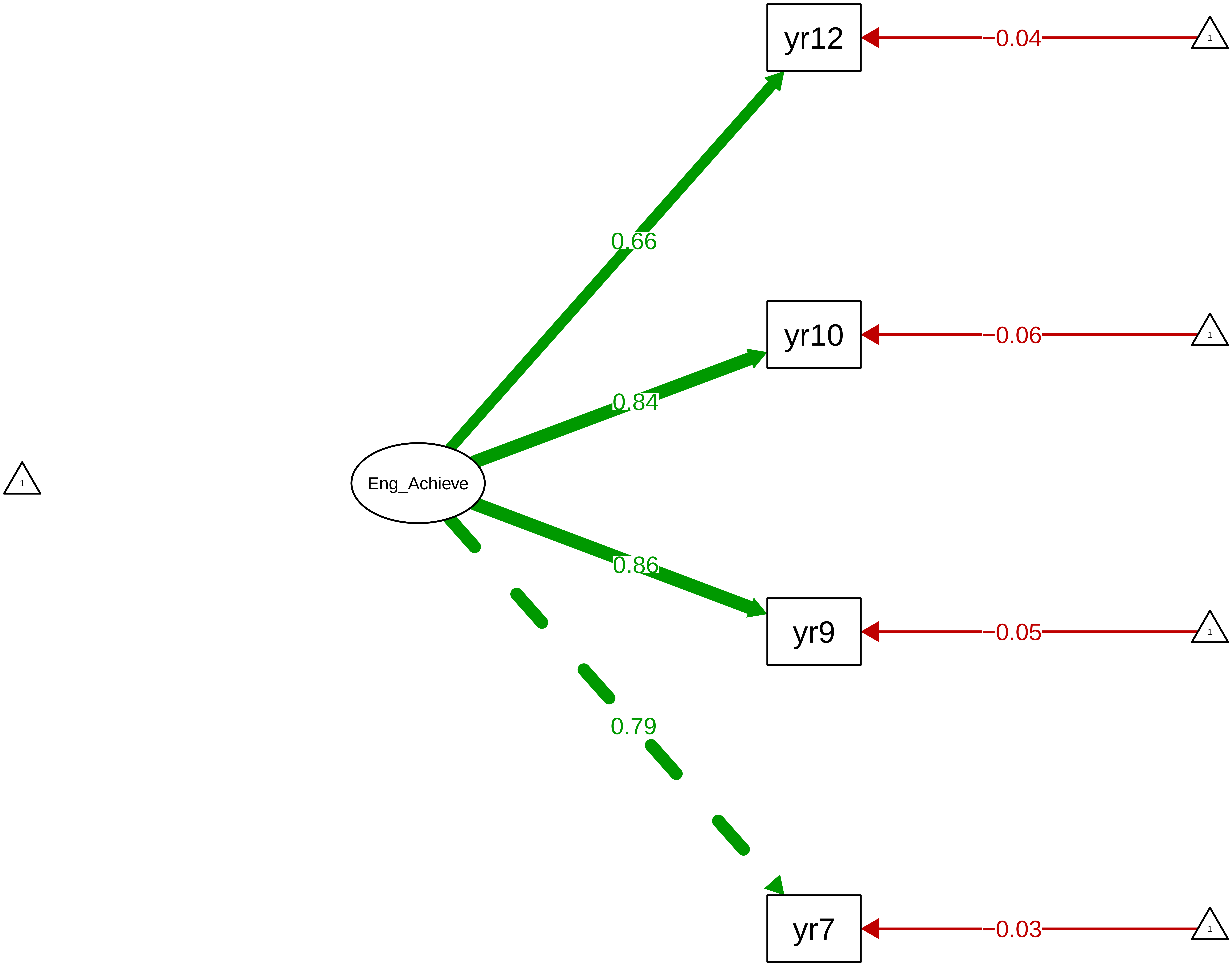

Figure S8_11: Maths Achievement Latent (Teacher 7−12)

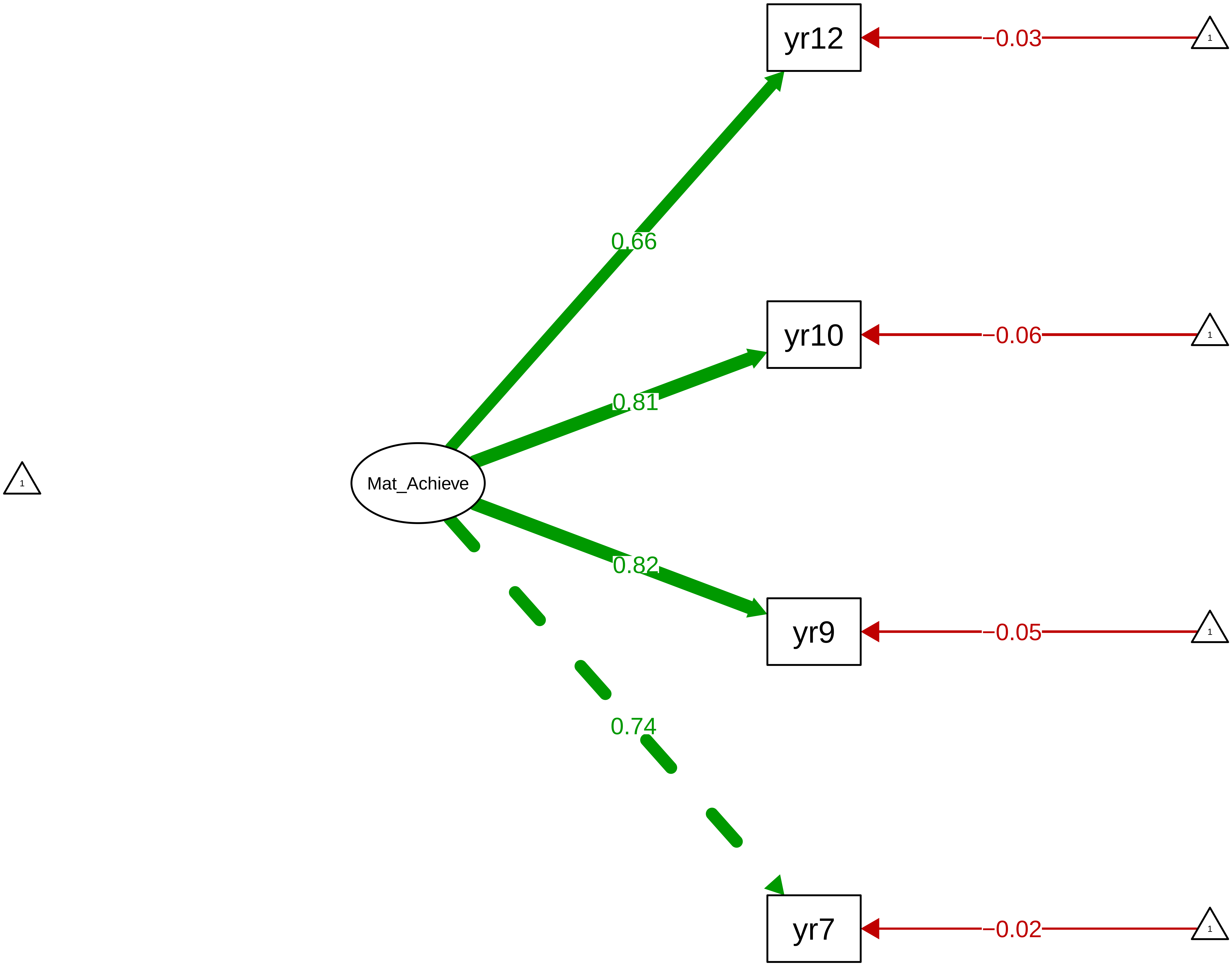

Figure S8_12: Science Achievement Latent (Teacher 9−12)

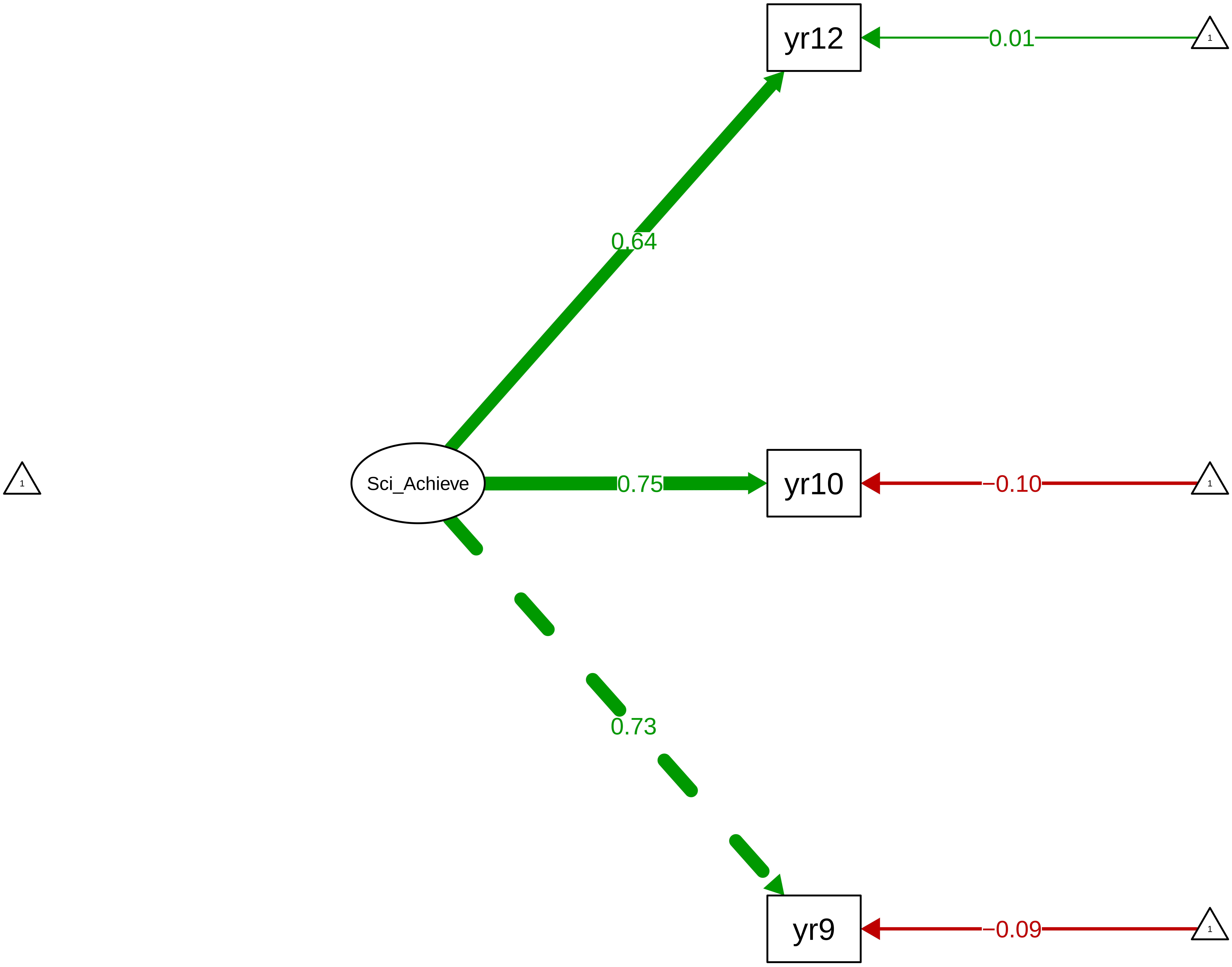

**Figure S8_13: Core−Subject Achievement Latent (Teacher 7−12)**

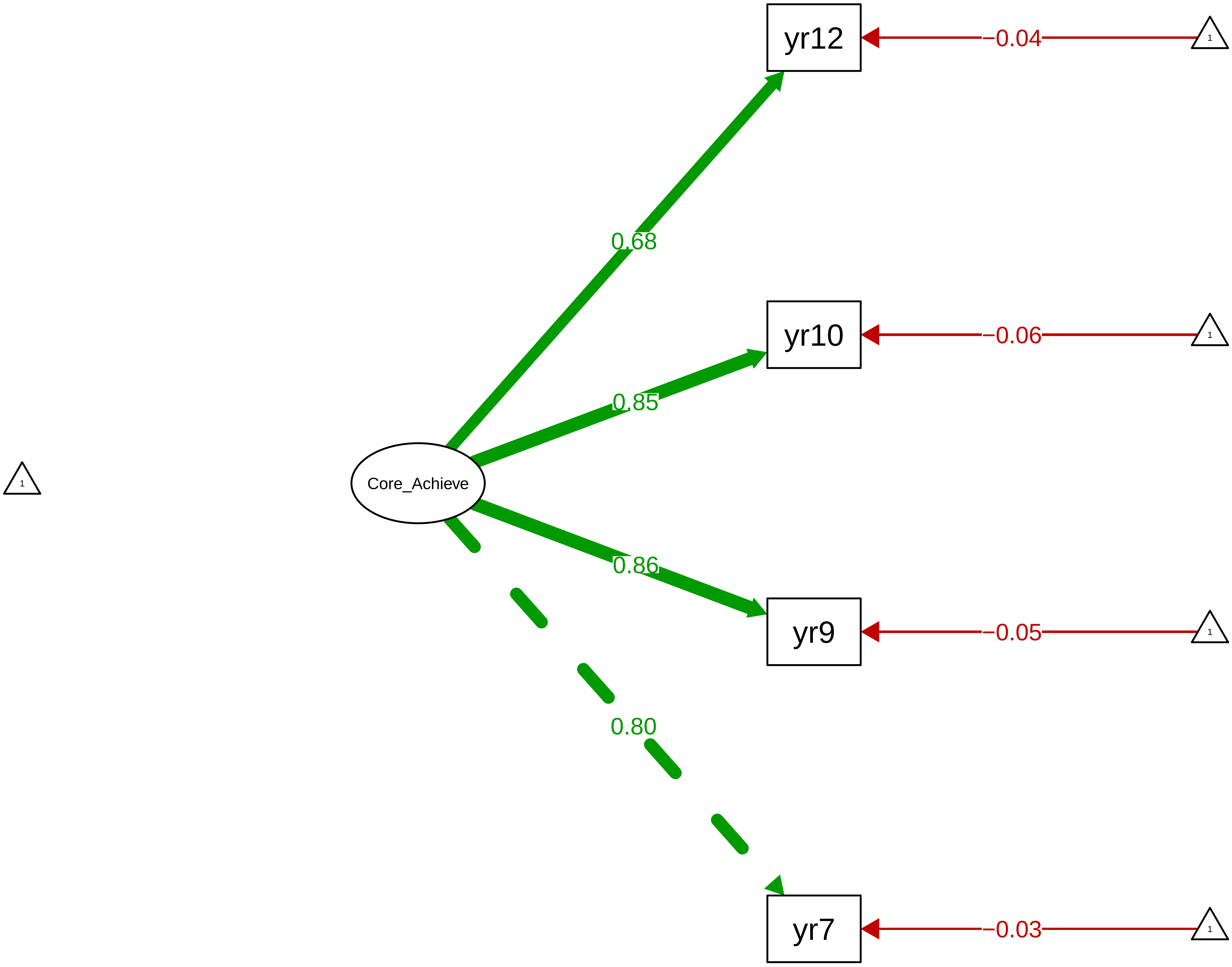

Figure S8_14: ARBQ Shyness − CTCR Overall

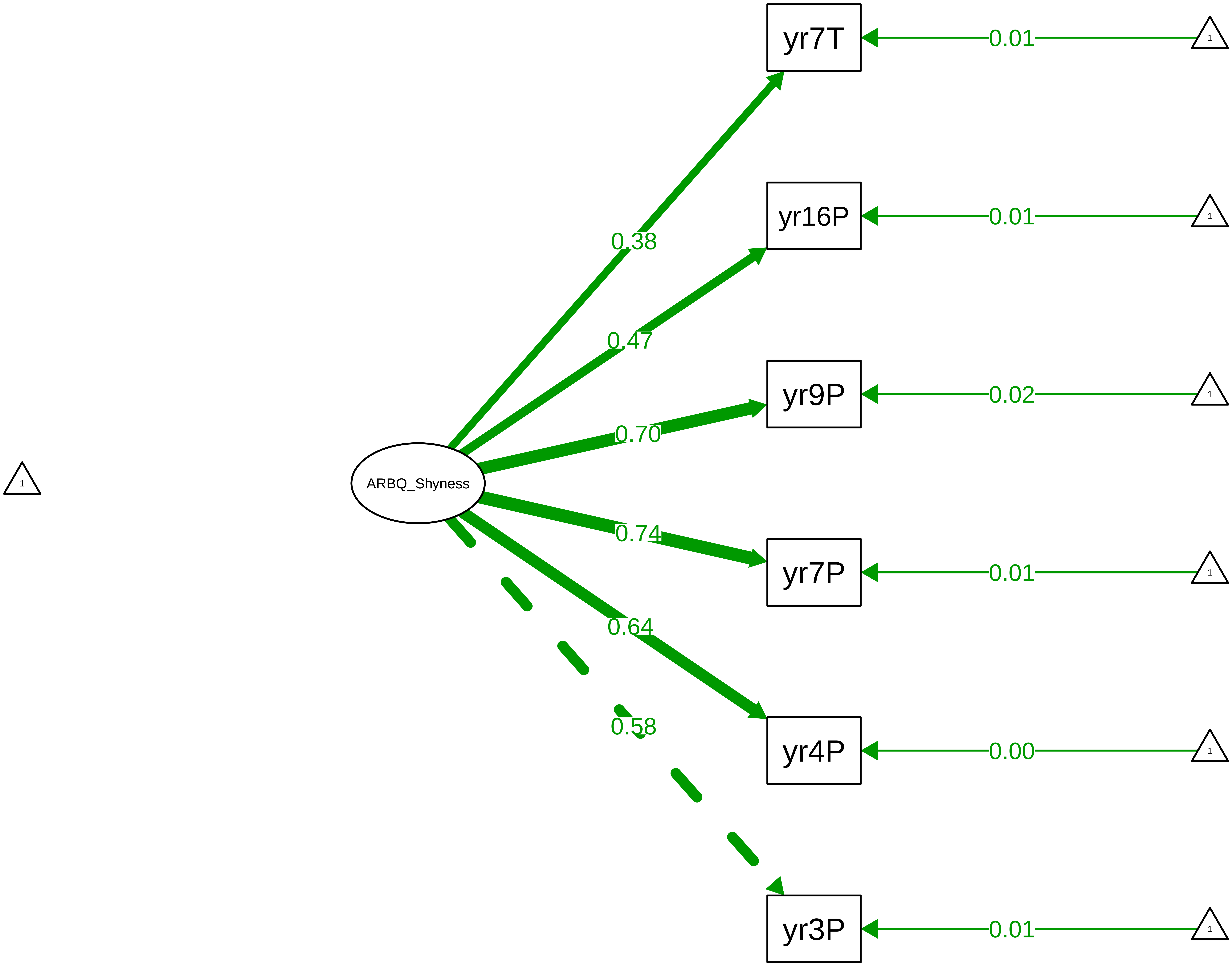

**Figure S8_15: ARBQ Shyness − CTCR Stage**

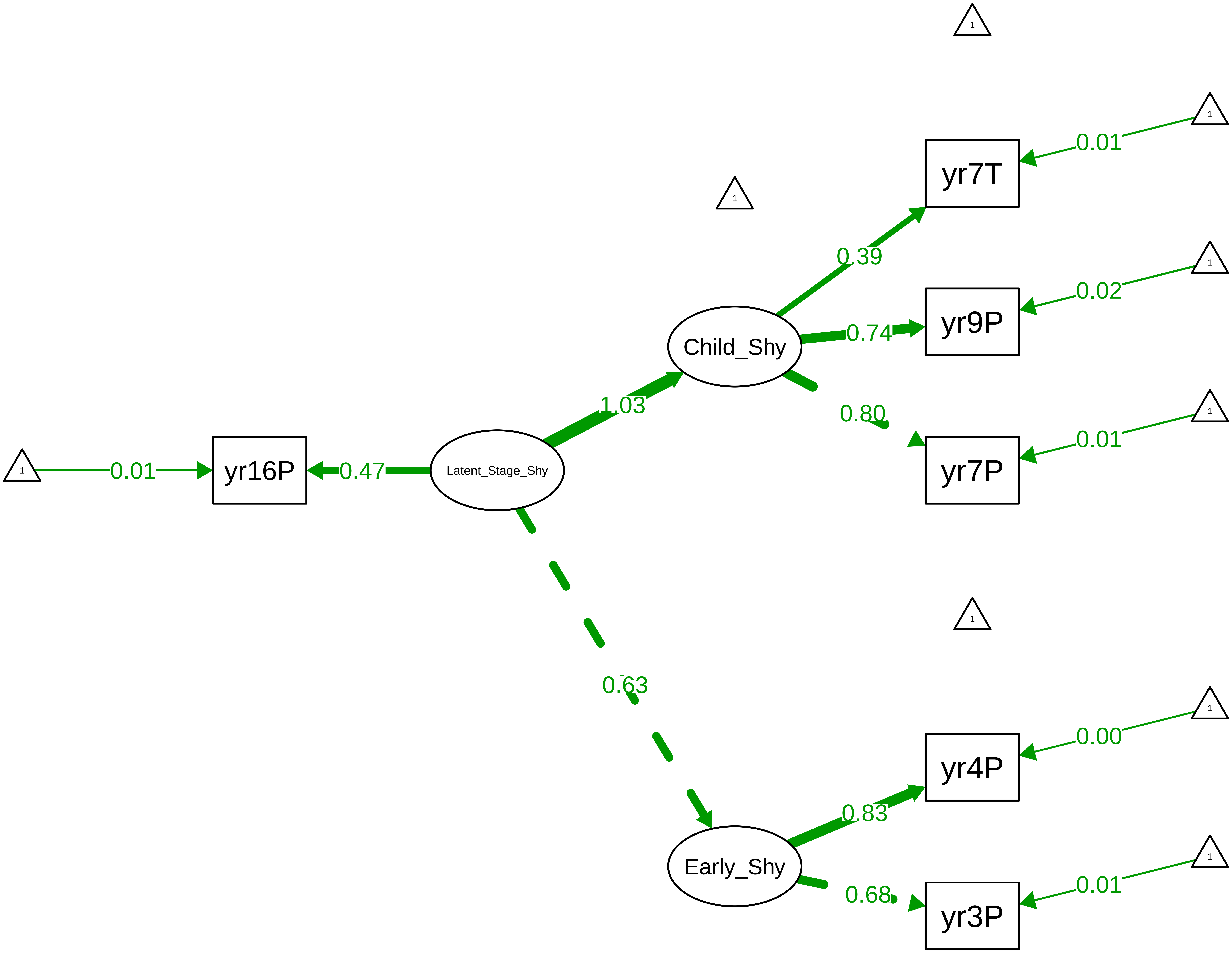

**Figure S8_16: ARBQ Fear − CTCR Overall**

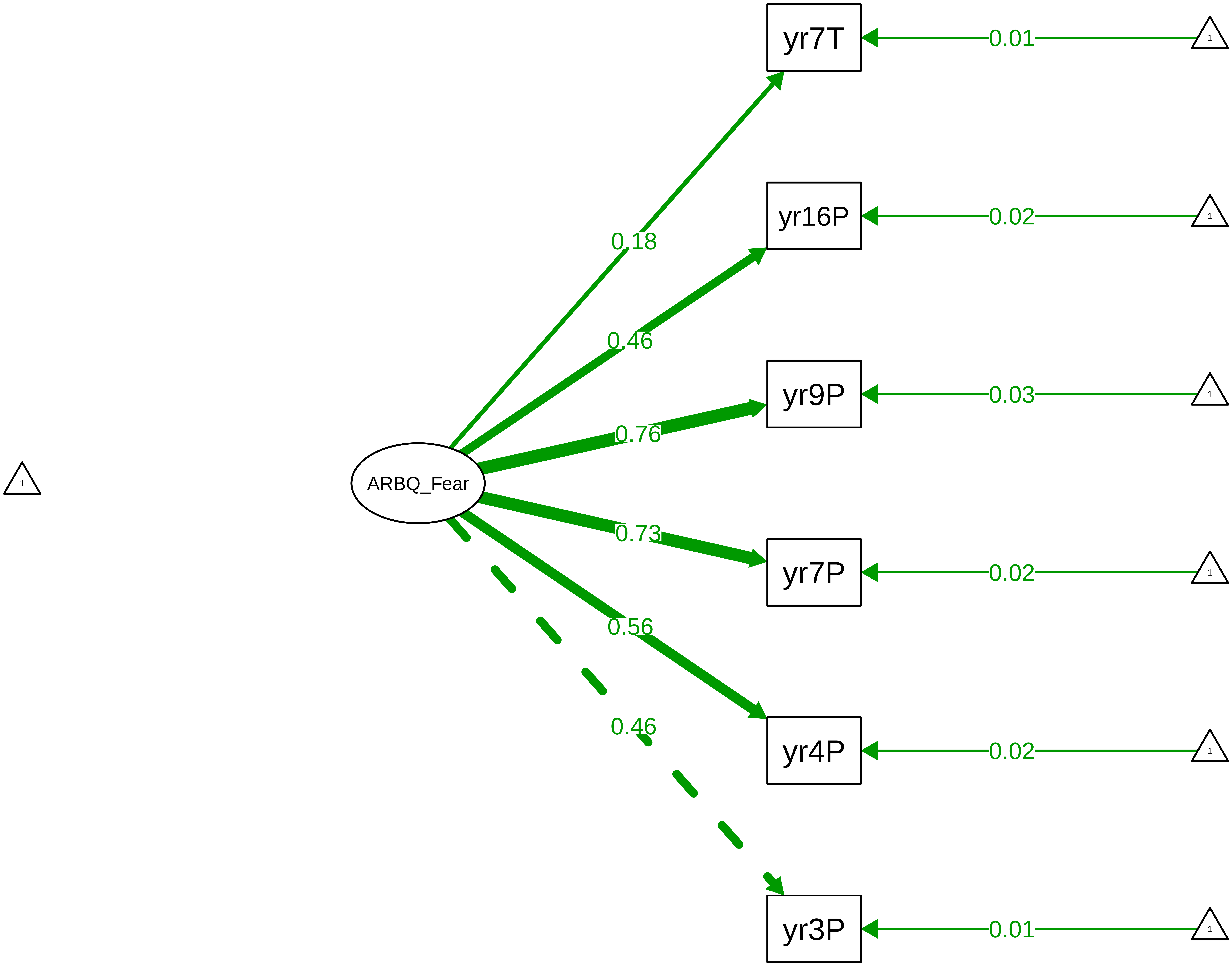

Figure S8_17: ARBQ Fear – CTCR Stage

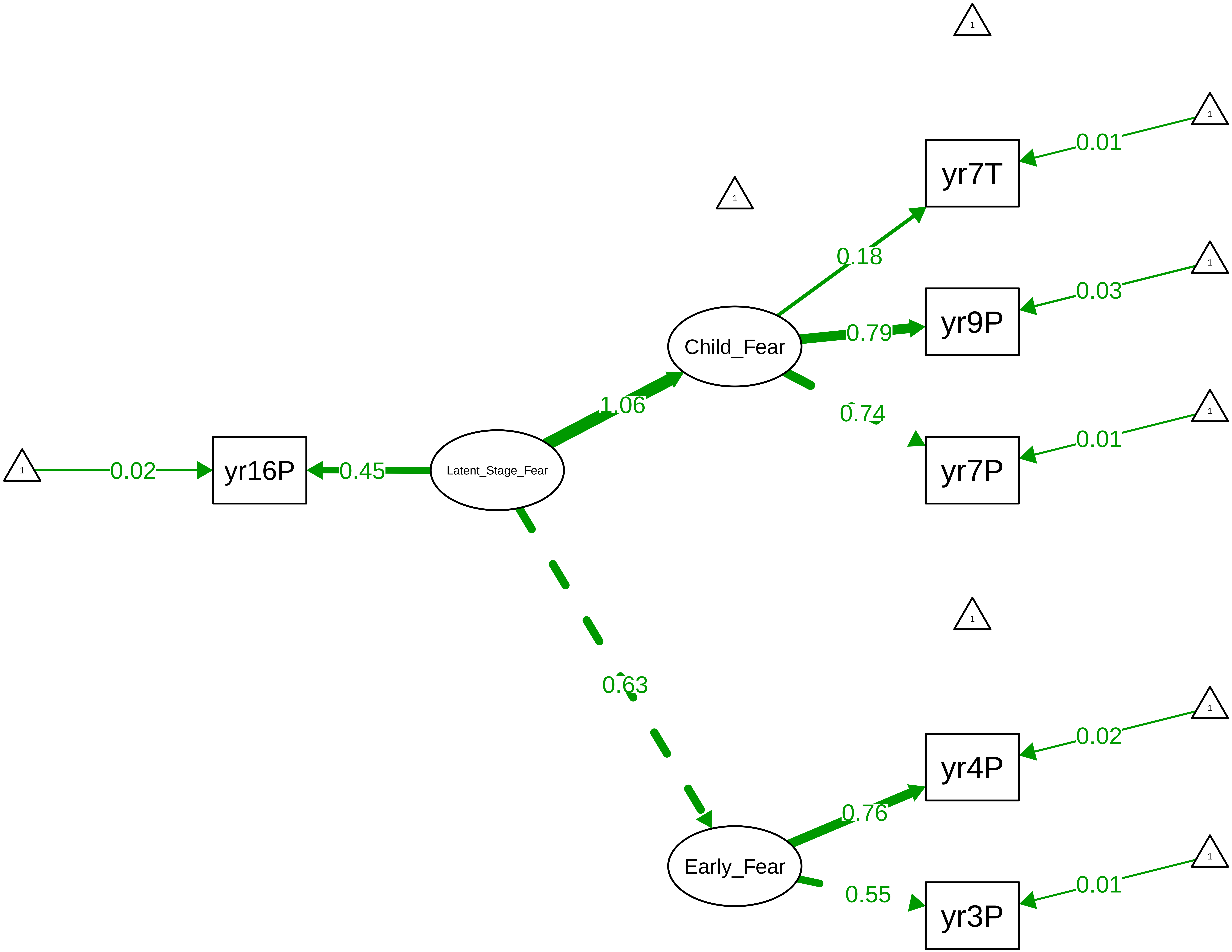

Figure S8_18: ARBQ Obsessive−Compulsive − CTCR Overall

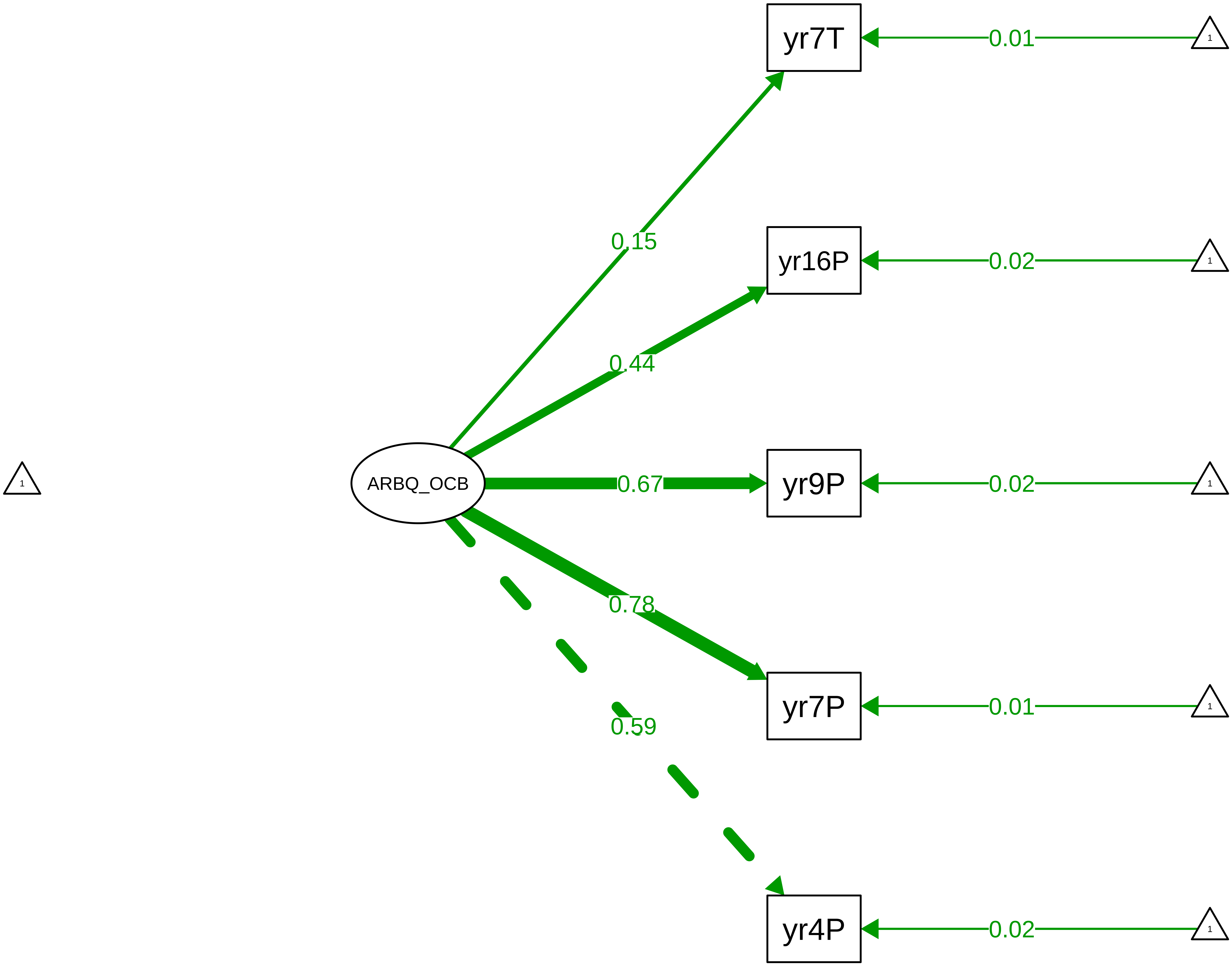

**Figure S8_19: ARBQ Obsessive−Compulsive − CTCR Stage**

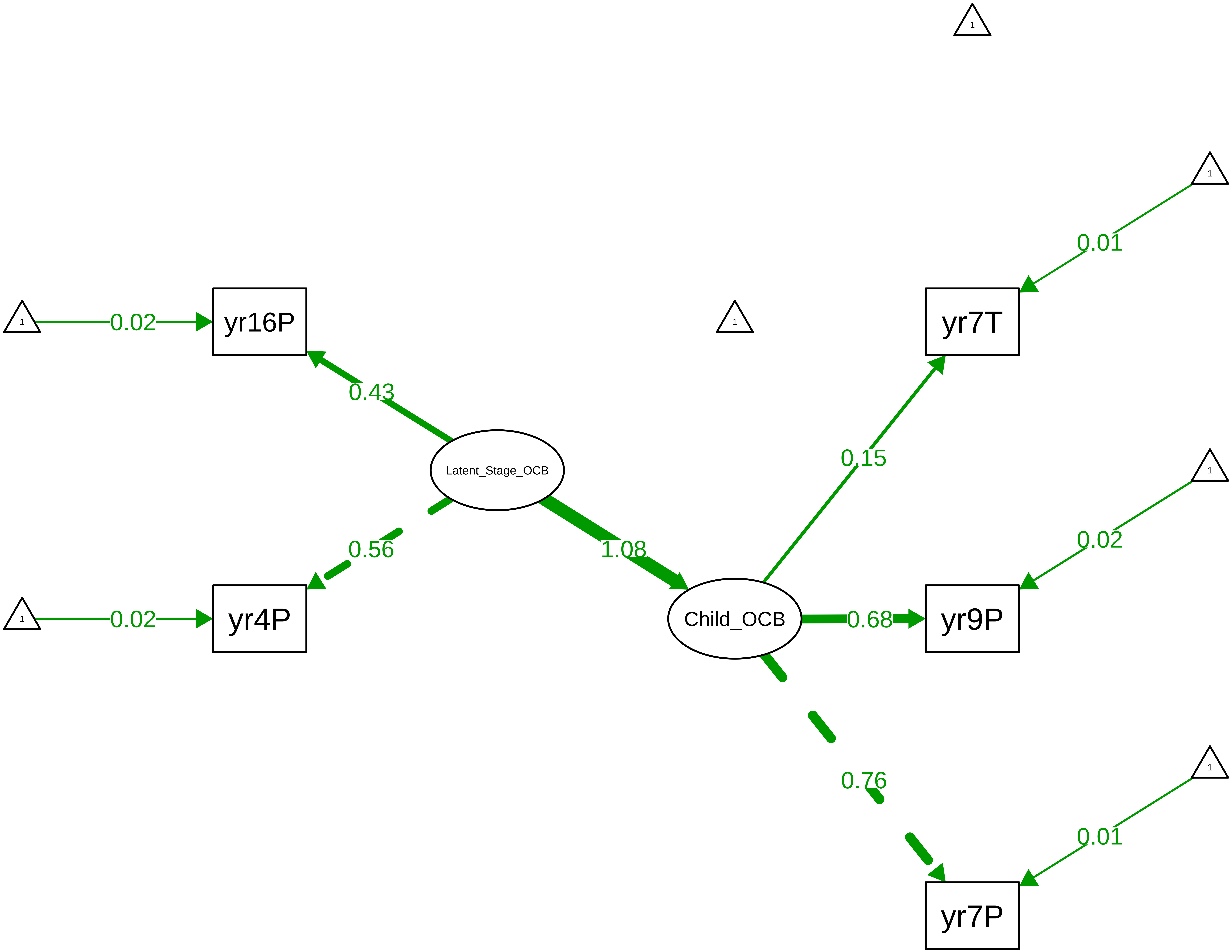

**Figure S8_20: ARBQ Negative Affect − CTCR Overall**

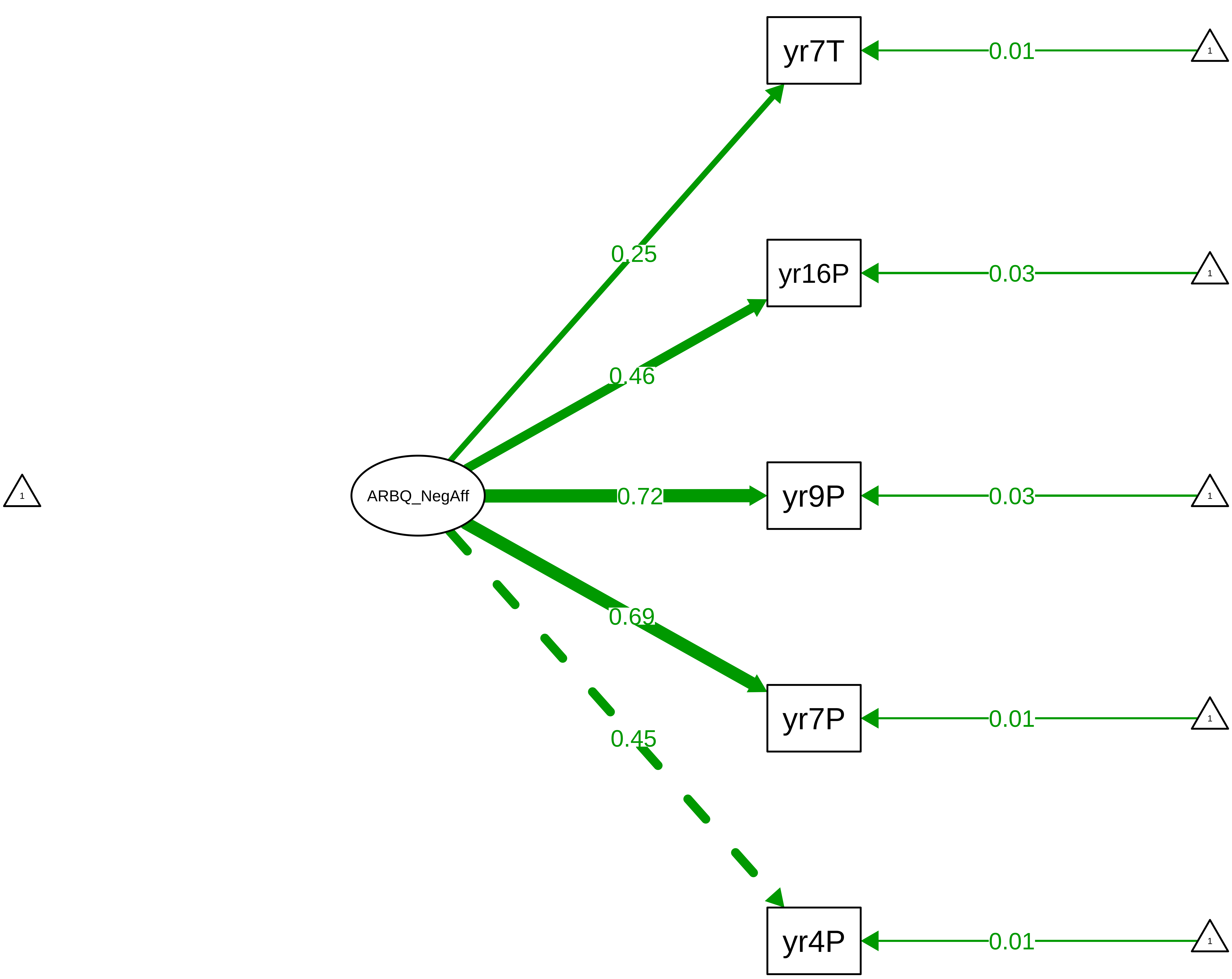

Figure S8_21: ARBQ Negative Affect − CTCR Stage

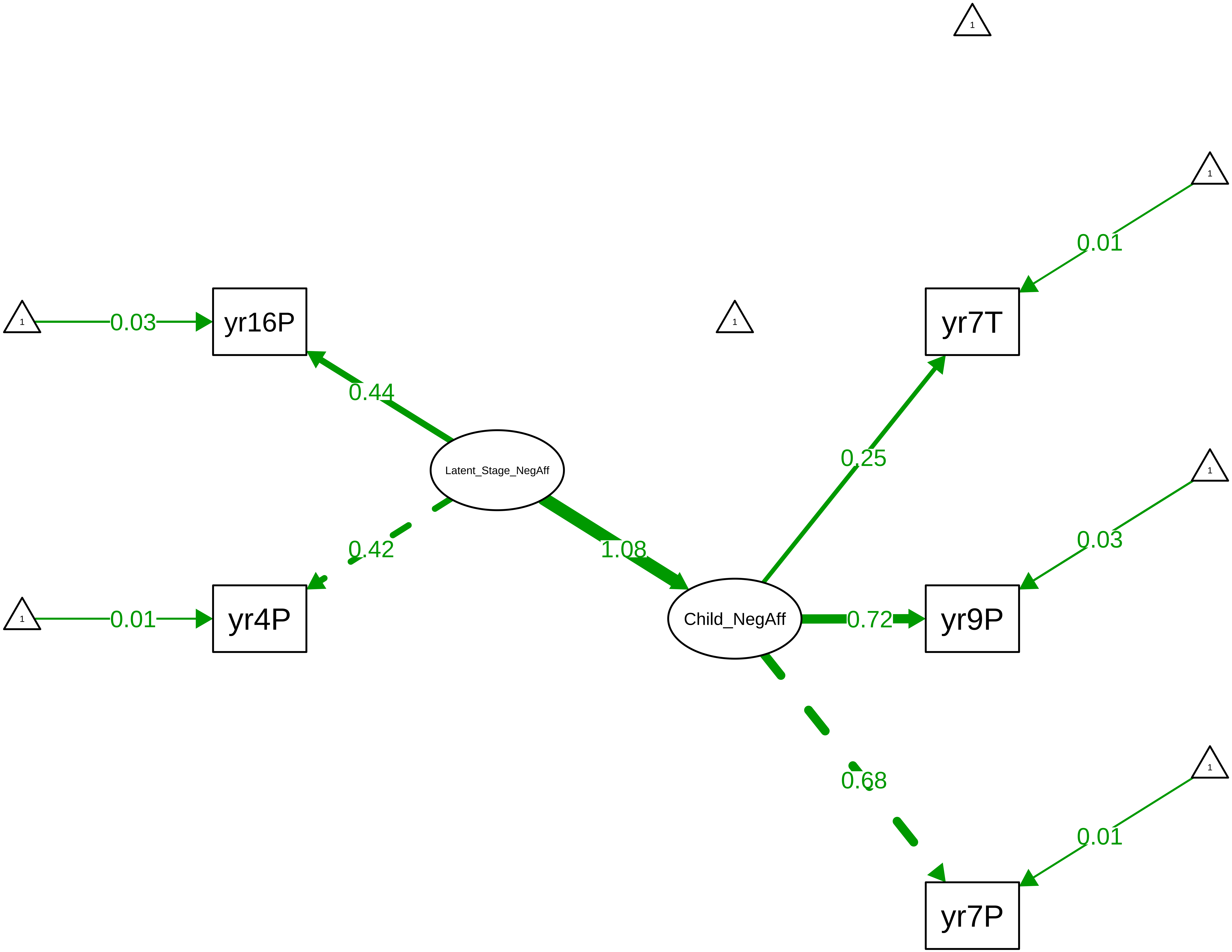

**Figure S8_22: ARBQ Negative Cognition − CTCR Overall**

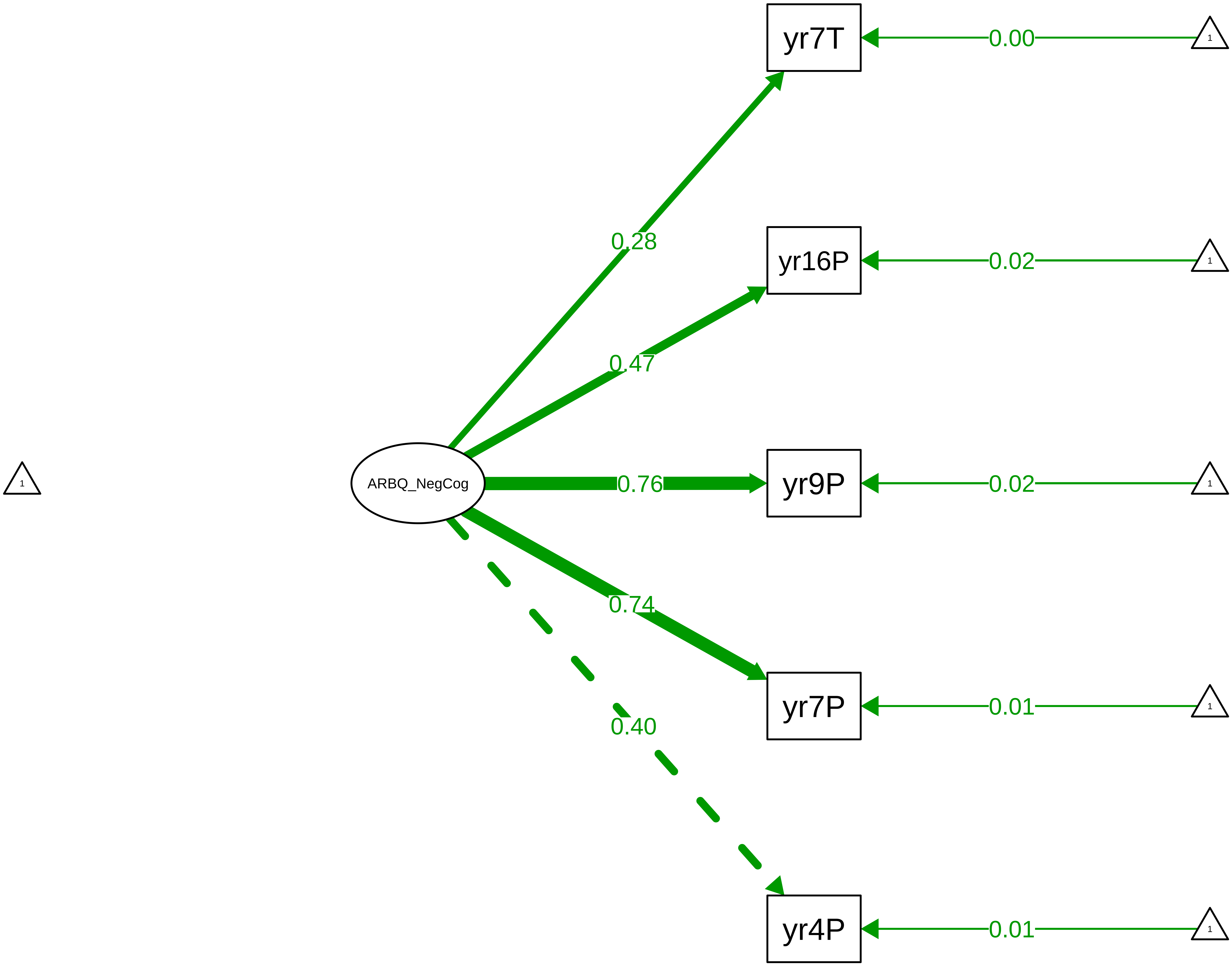

**Figure S8_23: ARBQ Negative Cognition − CTCR Stage**

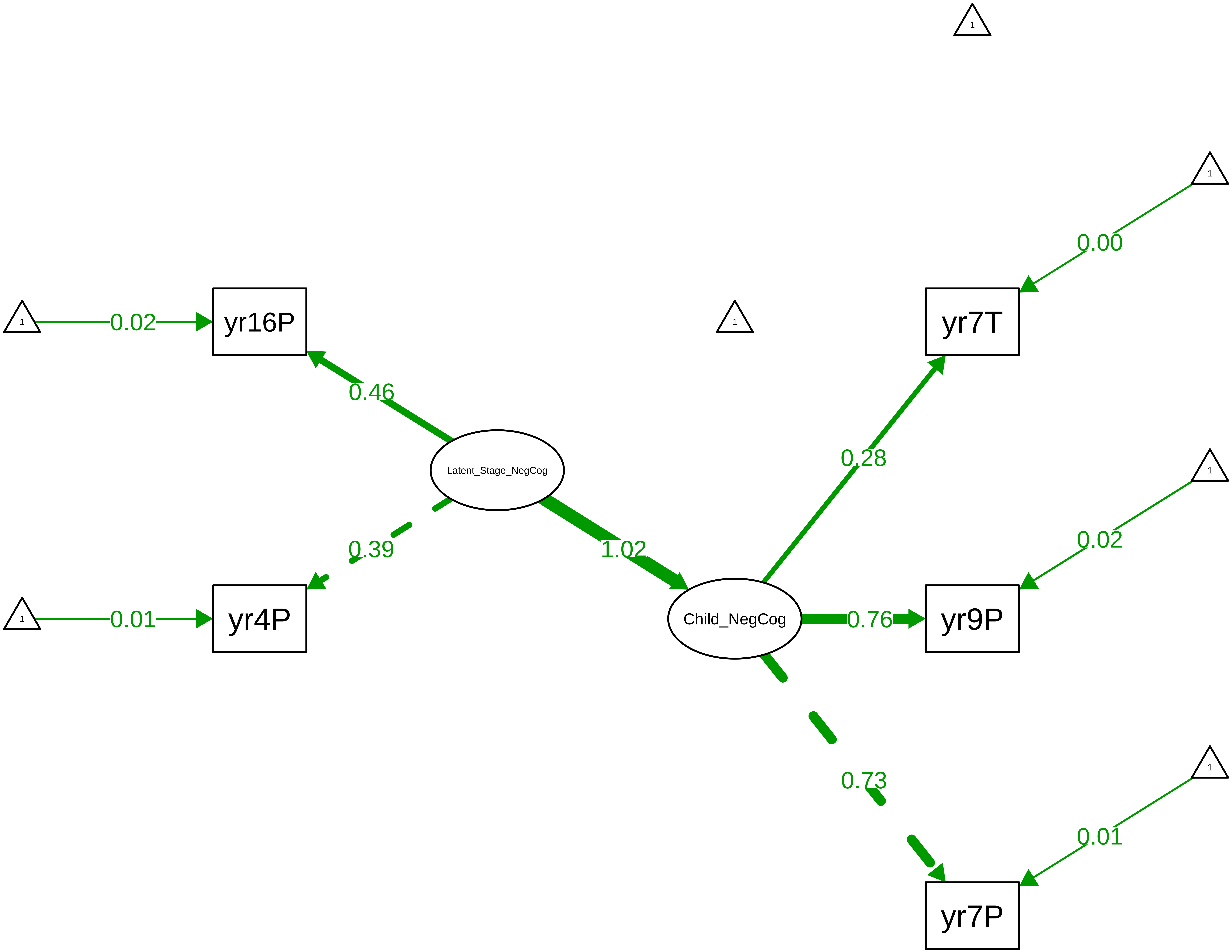

**Figure S8_24: ARBQ Anxiety Total − CTCR Overall**

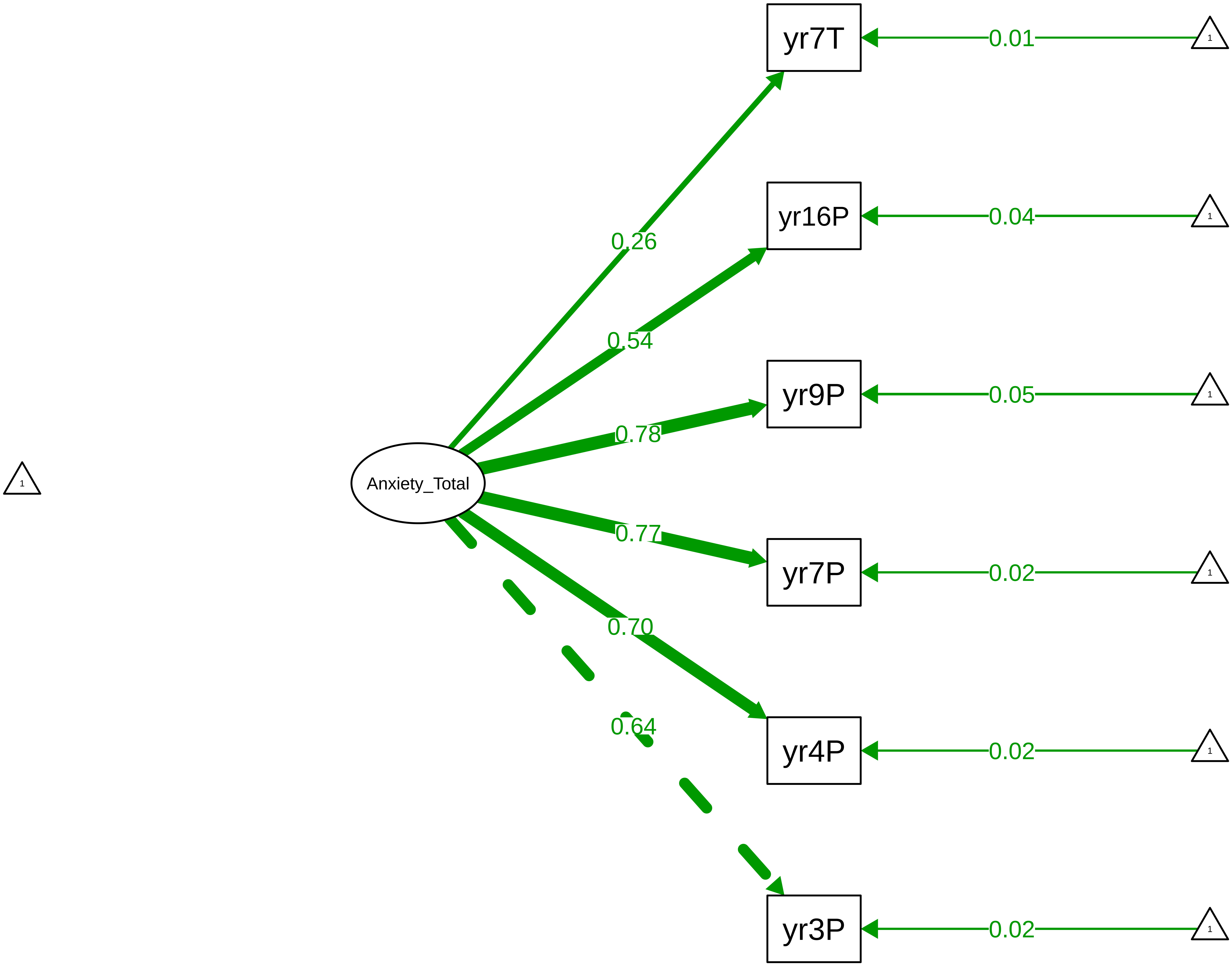

Figure S8_25: ARBQ Anxiety Total – CTCR Stage

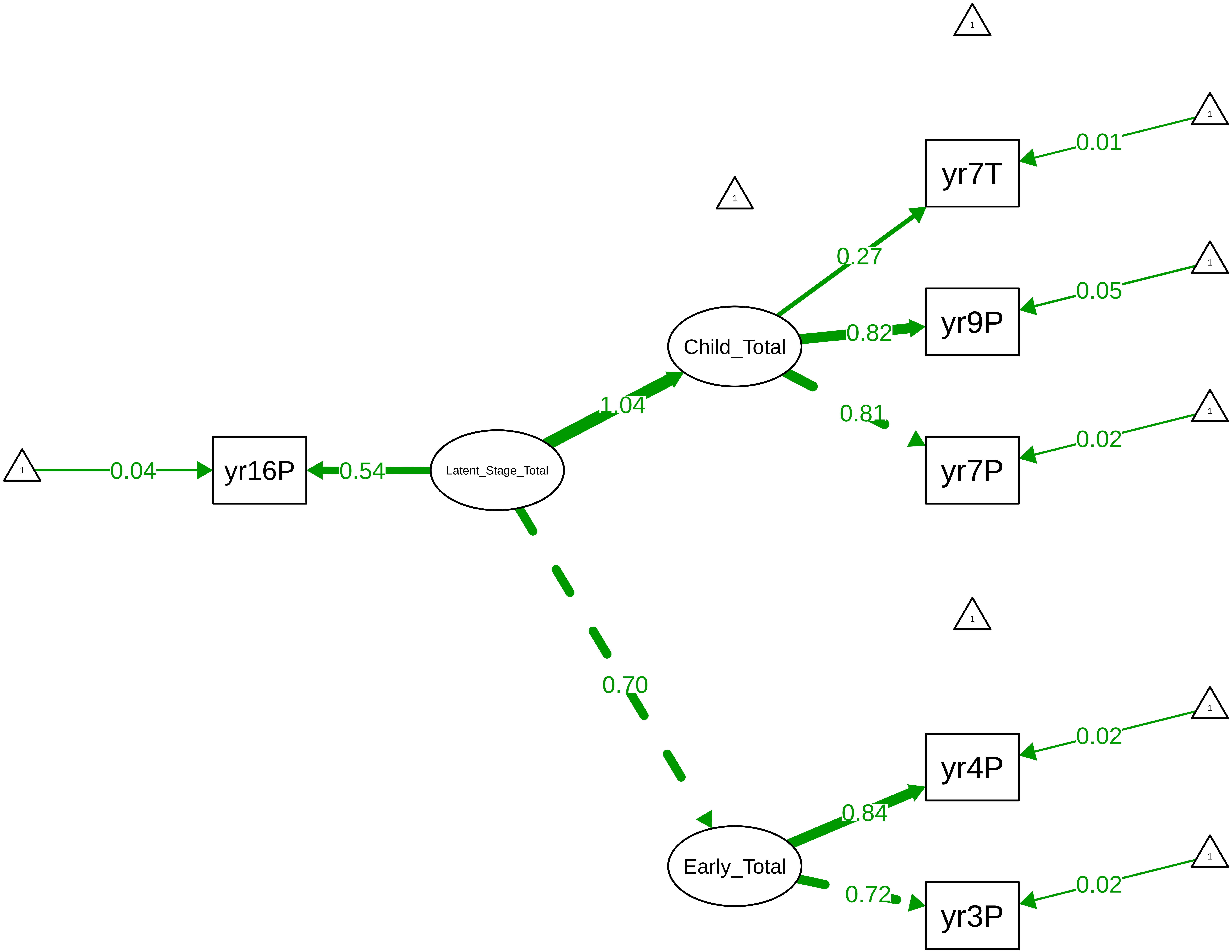

Figure S8_26: Conners Inattention − CTCR Overall

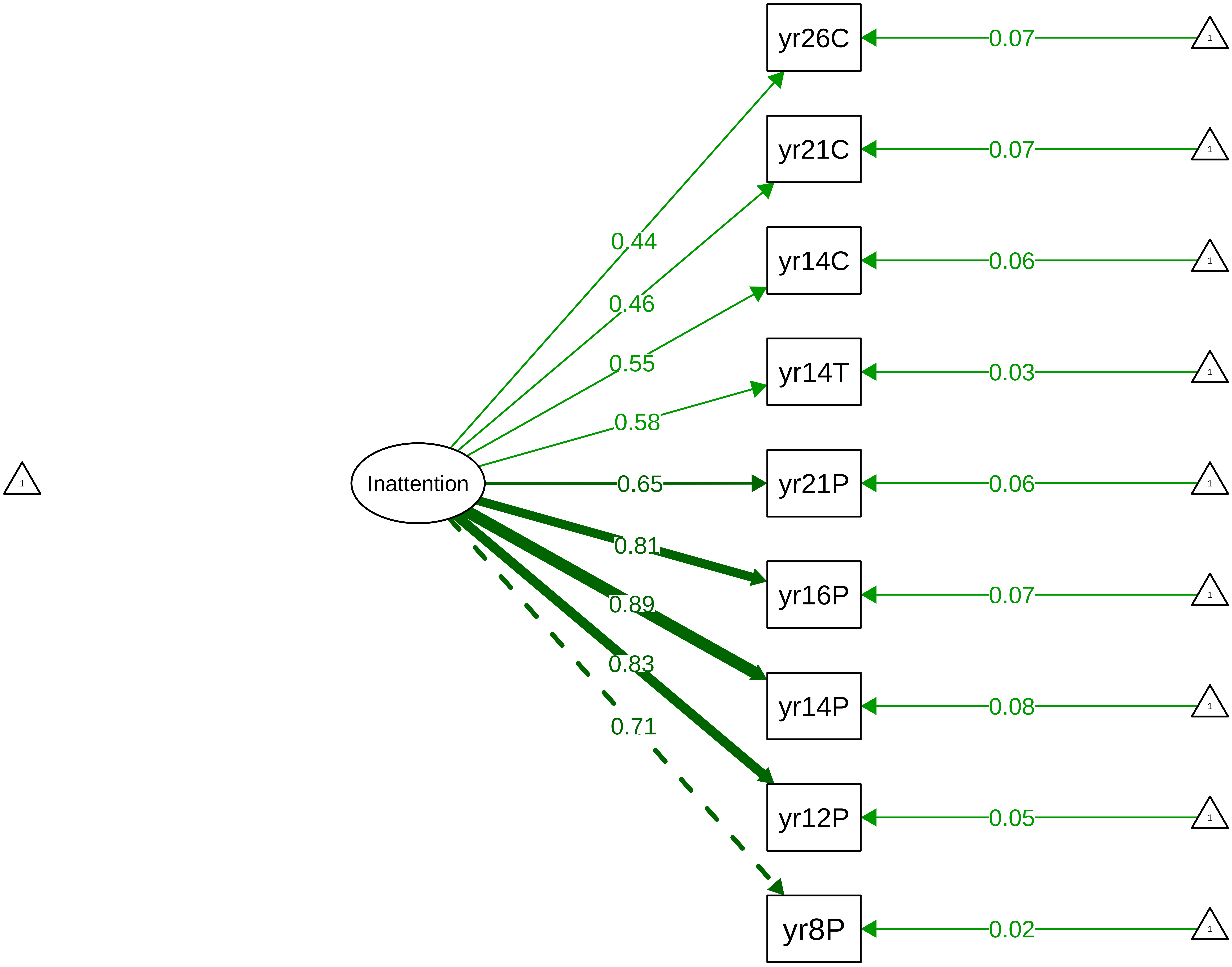

Figure S8_27: Conners Inattention − CTCR Stage

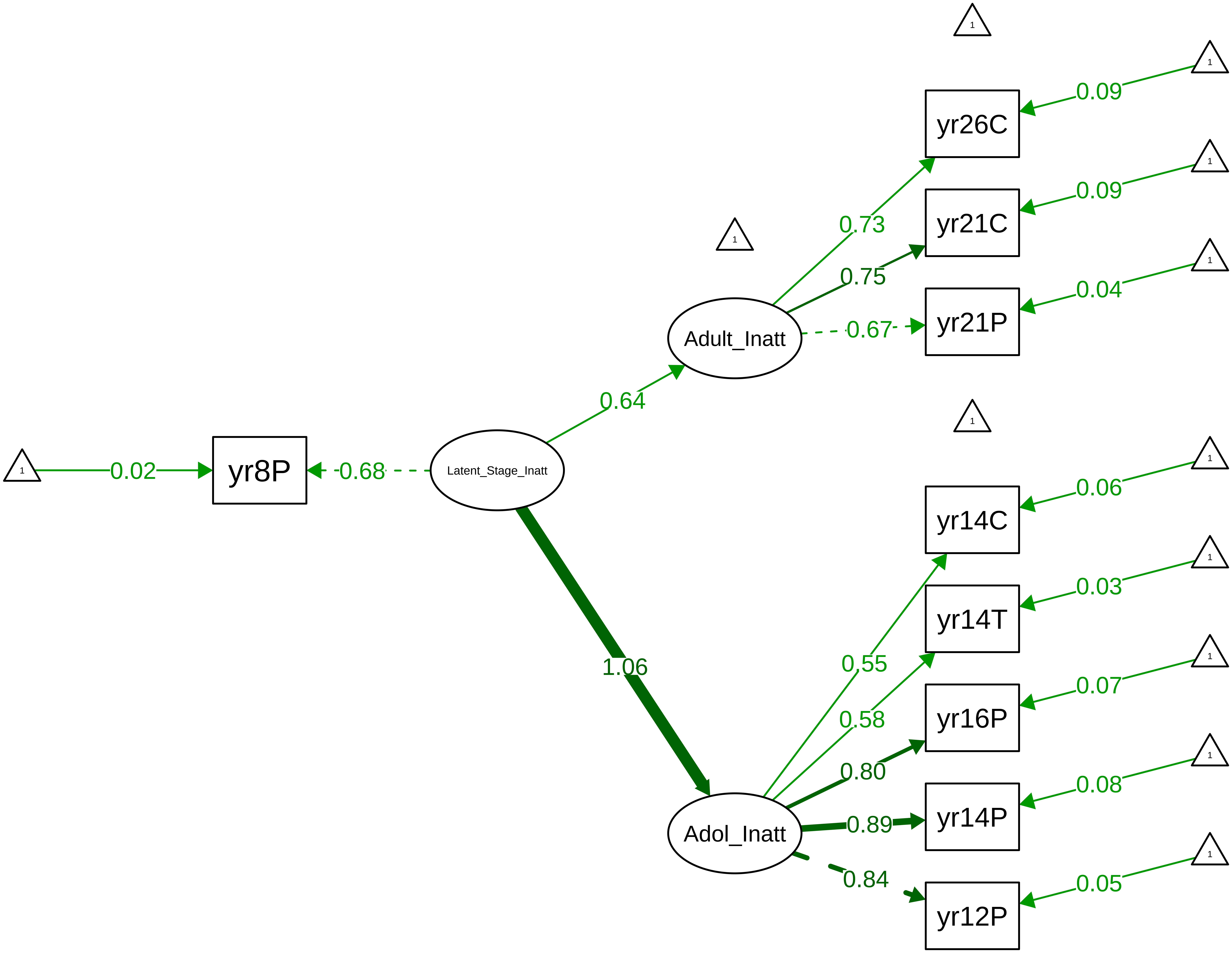

**Figure S8_28: Conners Inattention − Parent Overall**

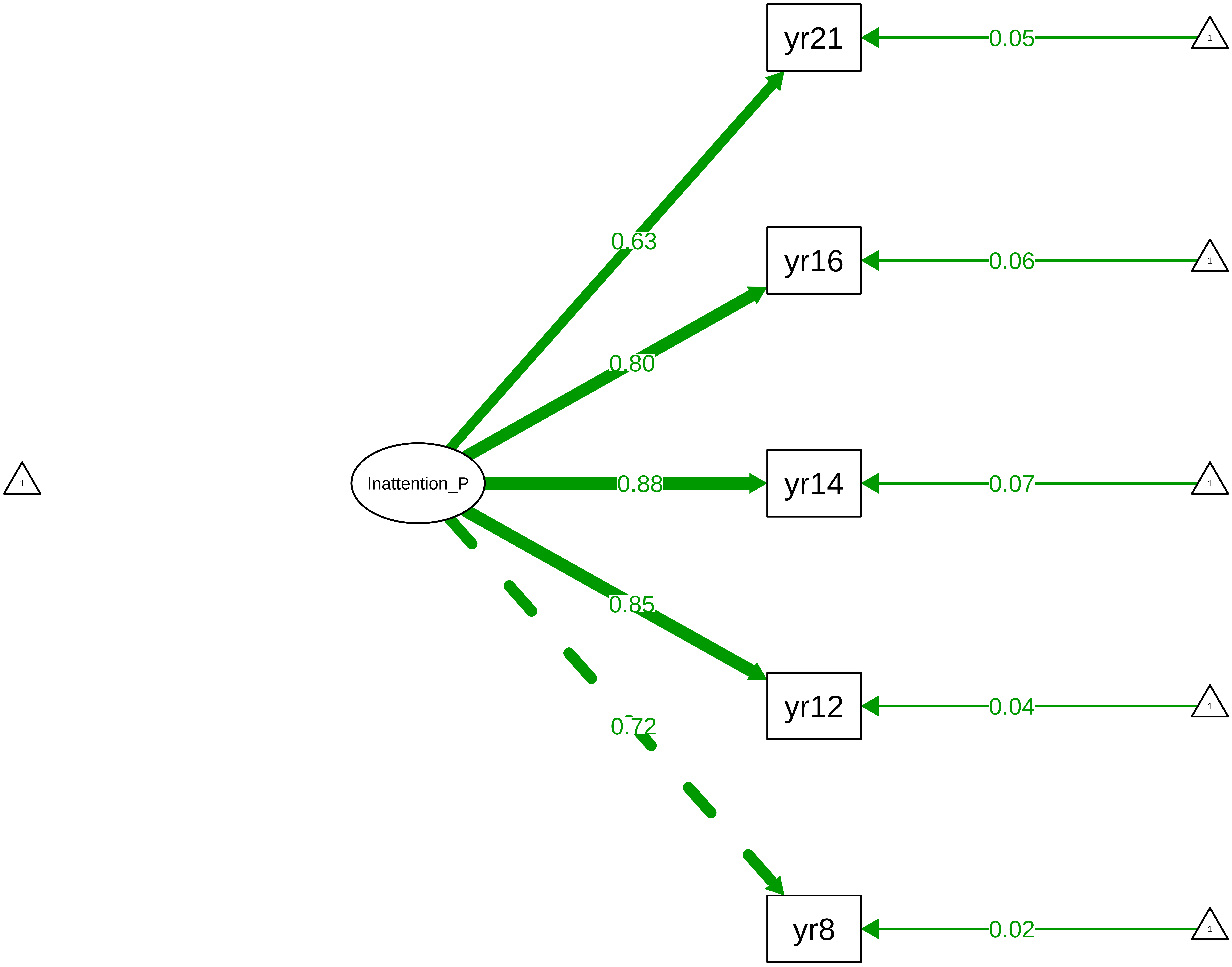

**Figure S8_29: Conners Inattention − Parent Stage**

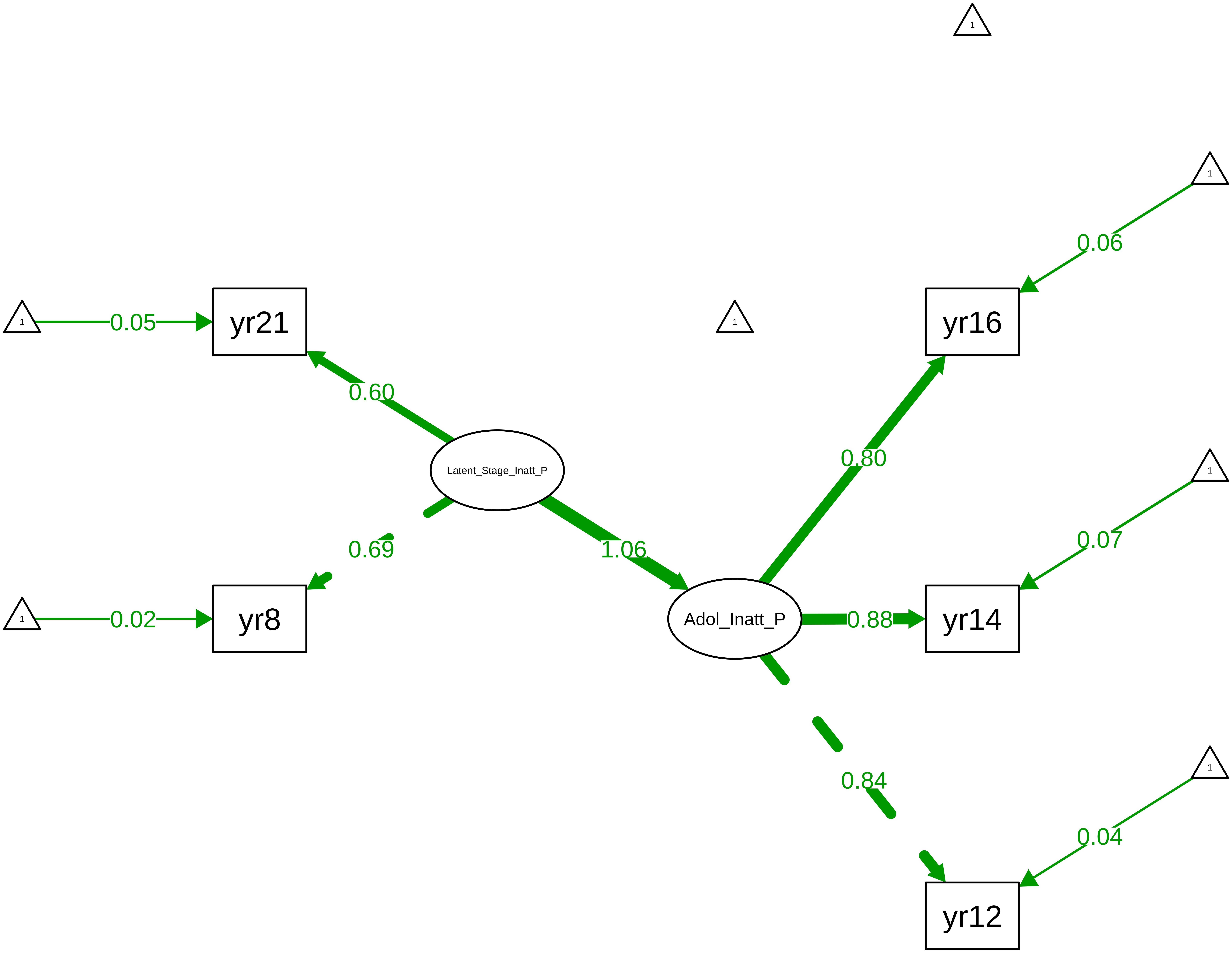

**Figure S8_30: Conners Hyperactivity−Impulsivity − CTCR Overall**

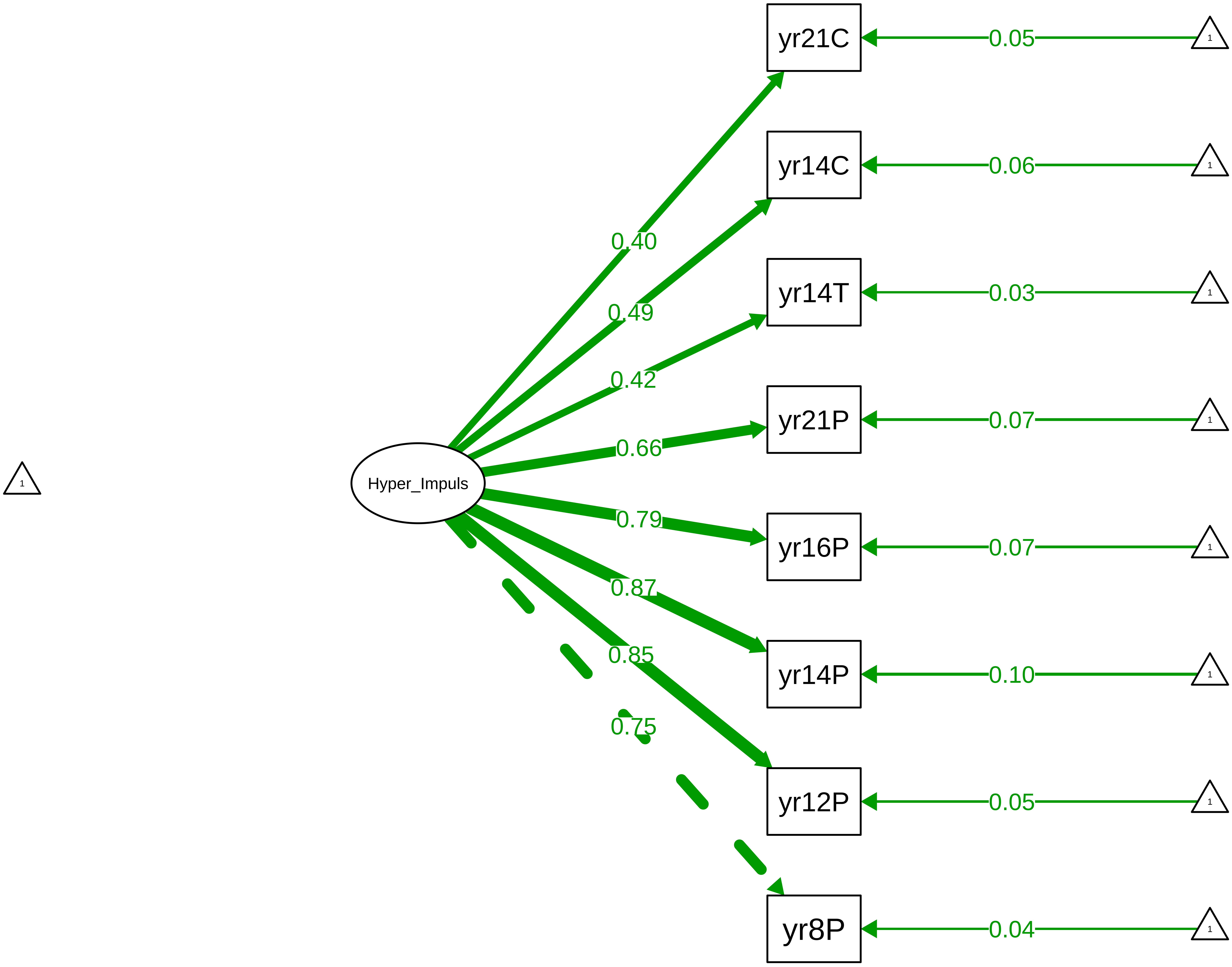

Figure S8_31: Conners Hyperactivity−Impulsivity − CTCR Stage

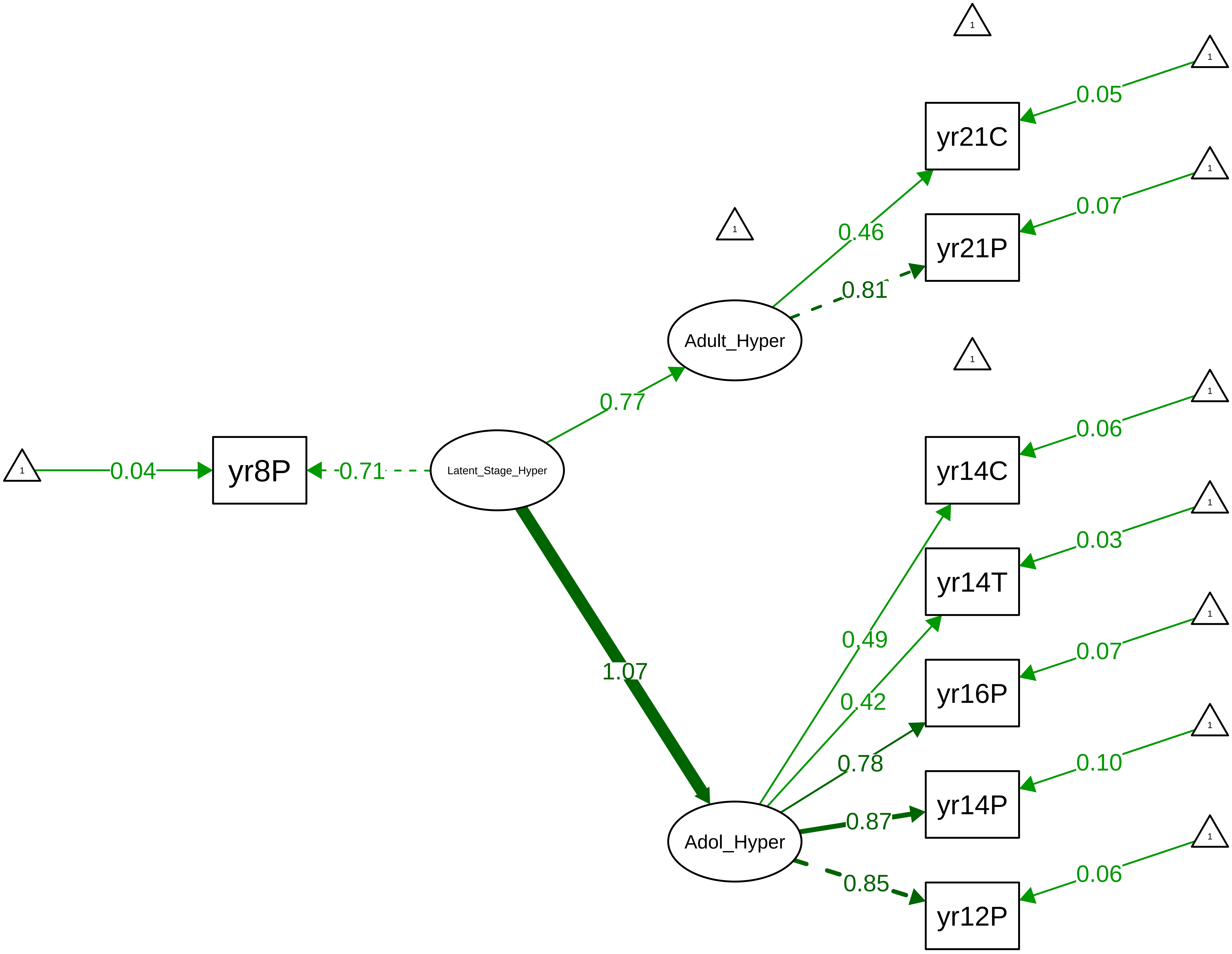

Figure S8_32: Conners Hyperactivity−Impulsivity − Parent Overall

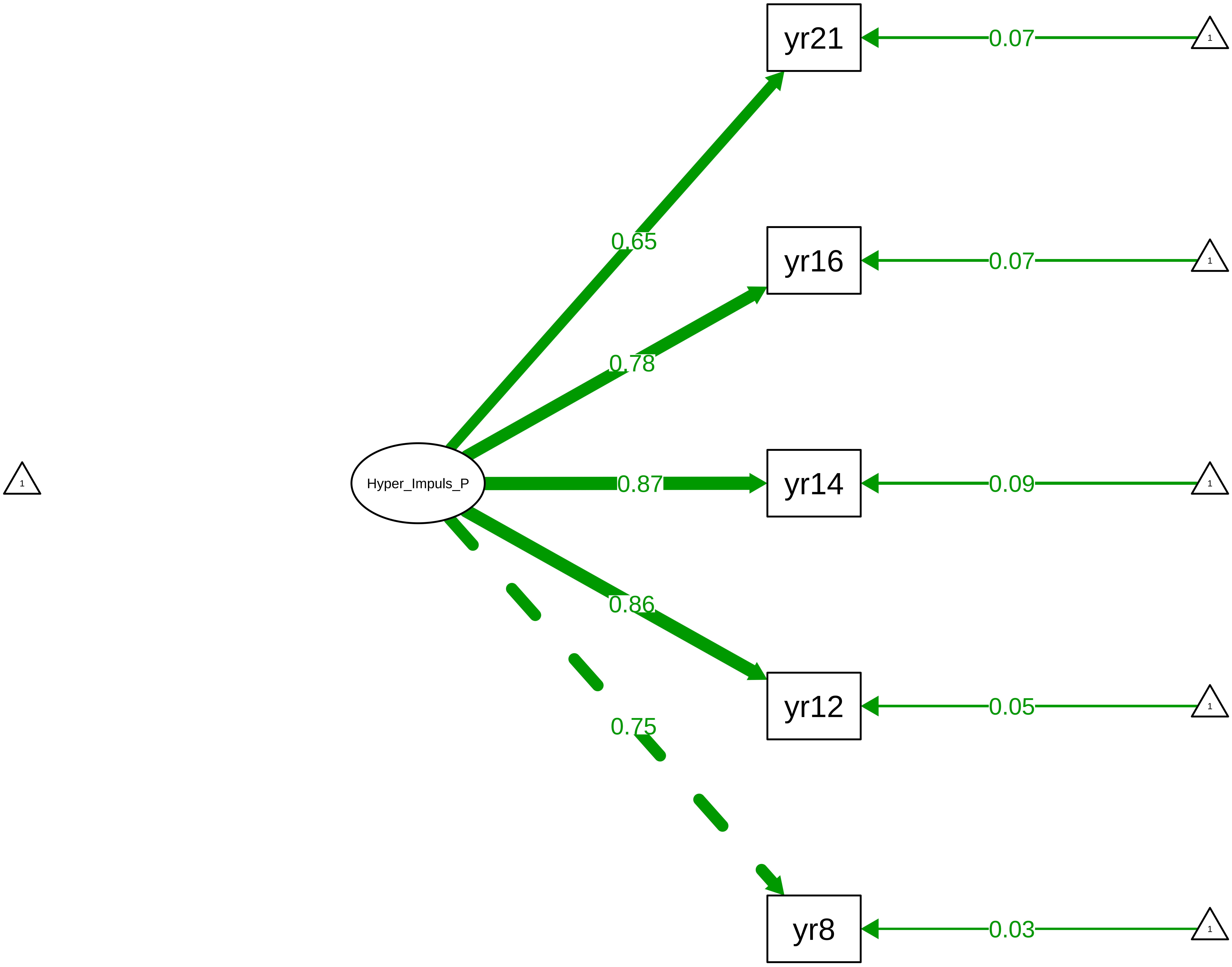

#### Figure S8_33: Conners Hyperactivity−Impulsivity − Parent Stage

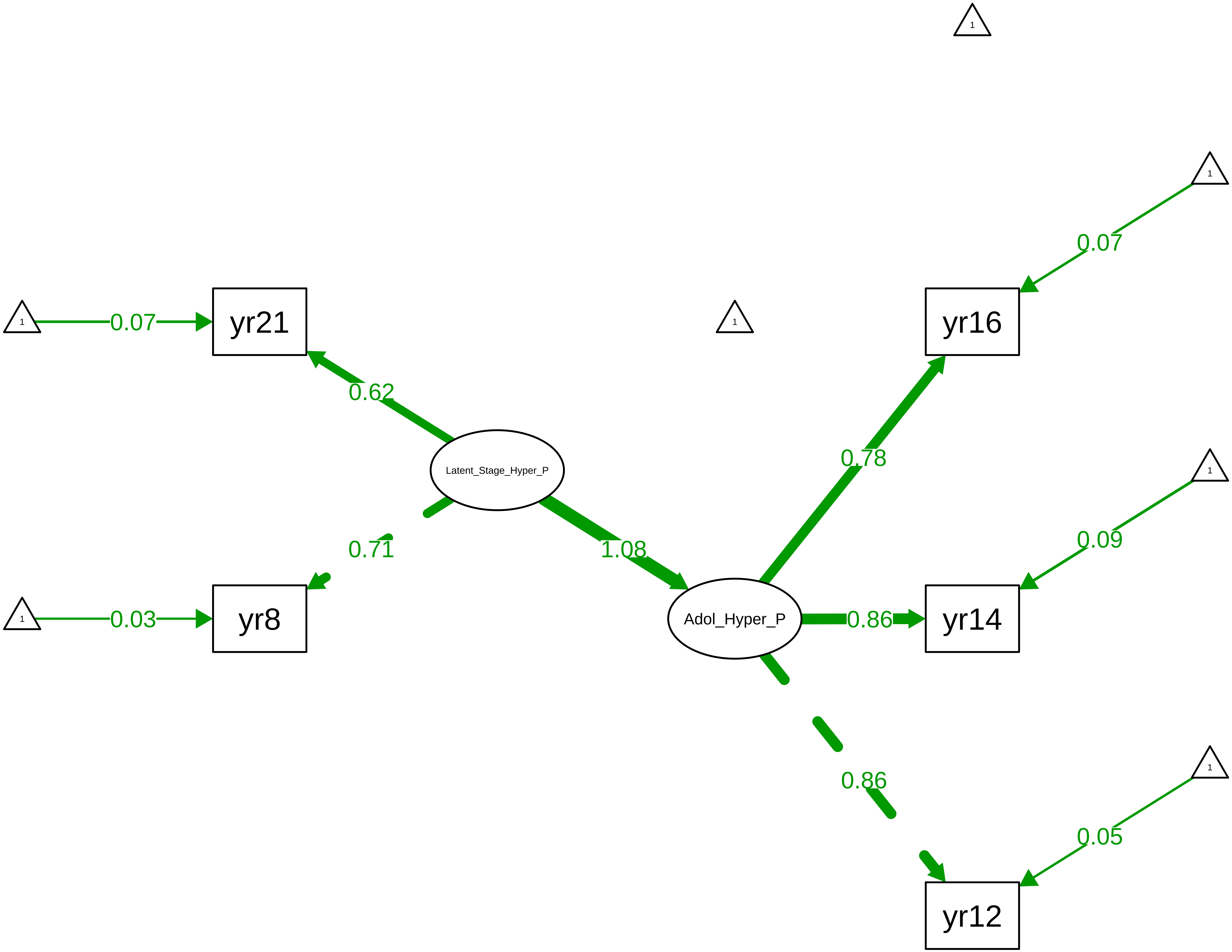

**Figure S8_34: Conners Total − CTCR Overall**

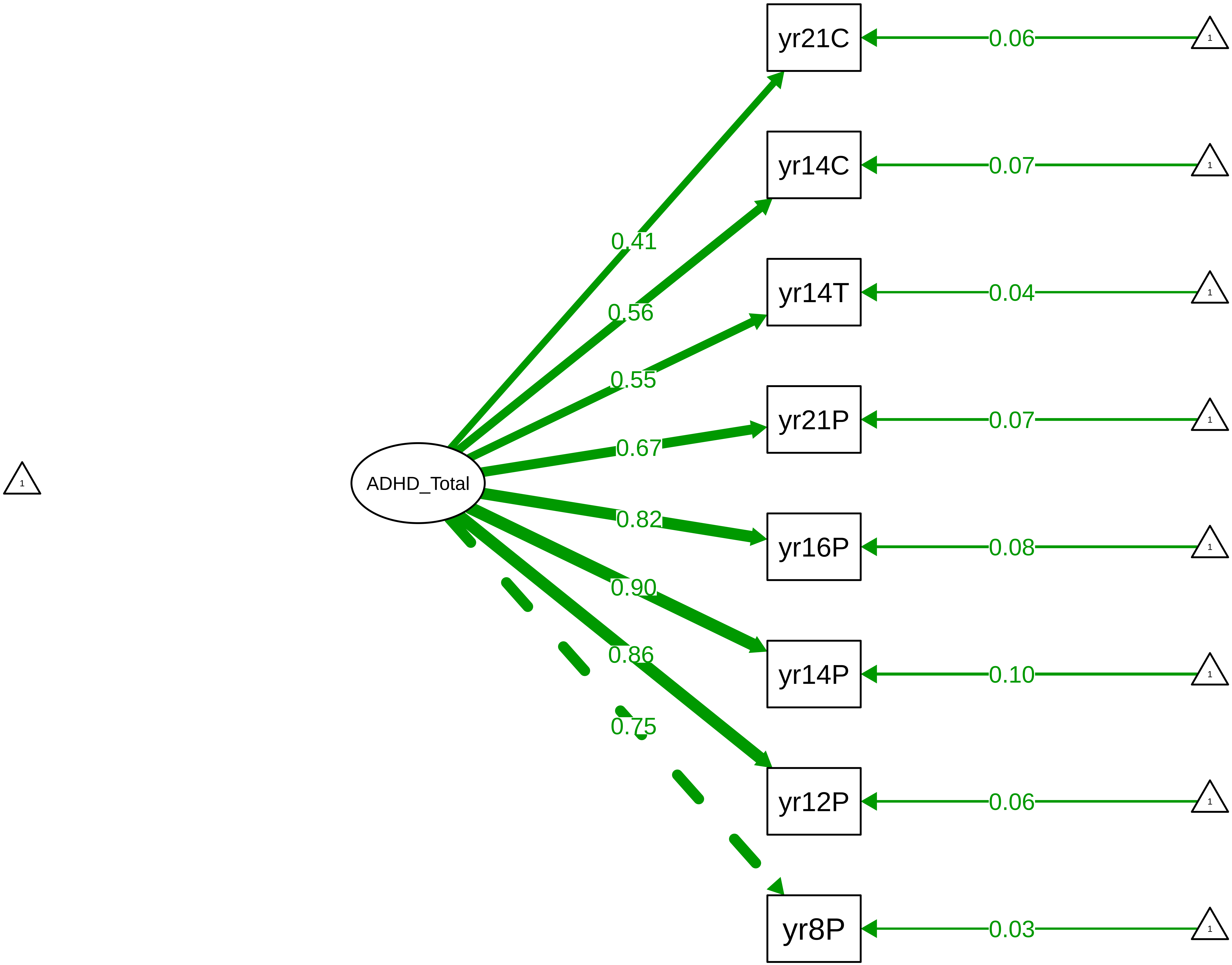

Figure S8_35: Conners Total − CTCR Stage

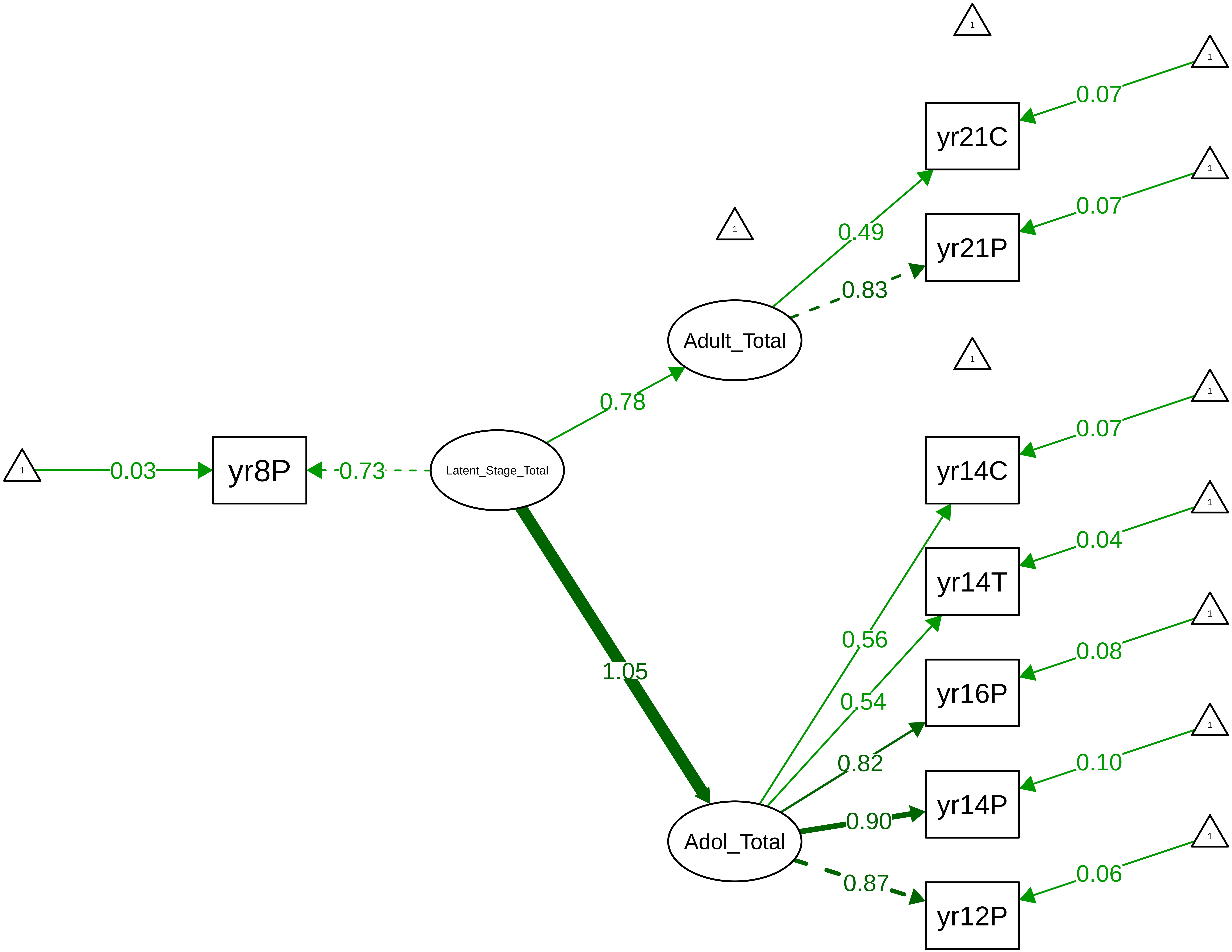

Figure S8_36: Conners Total – Parent Overall

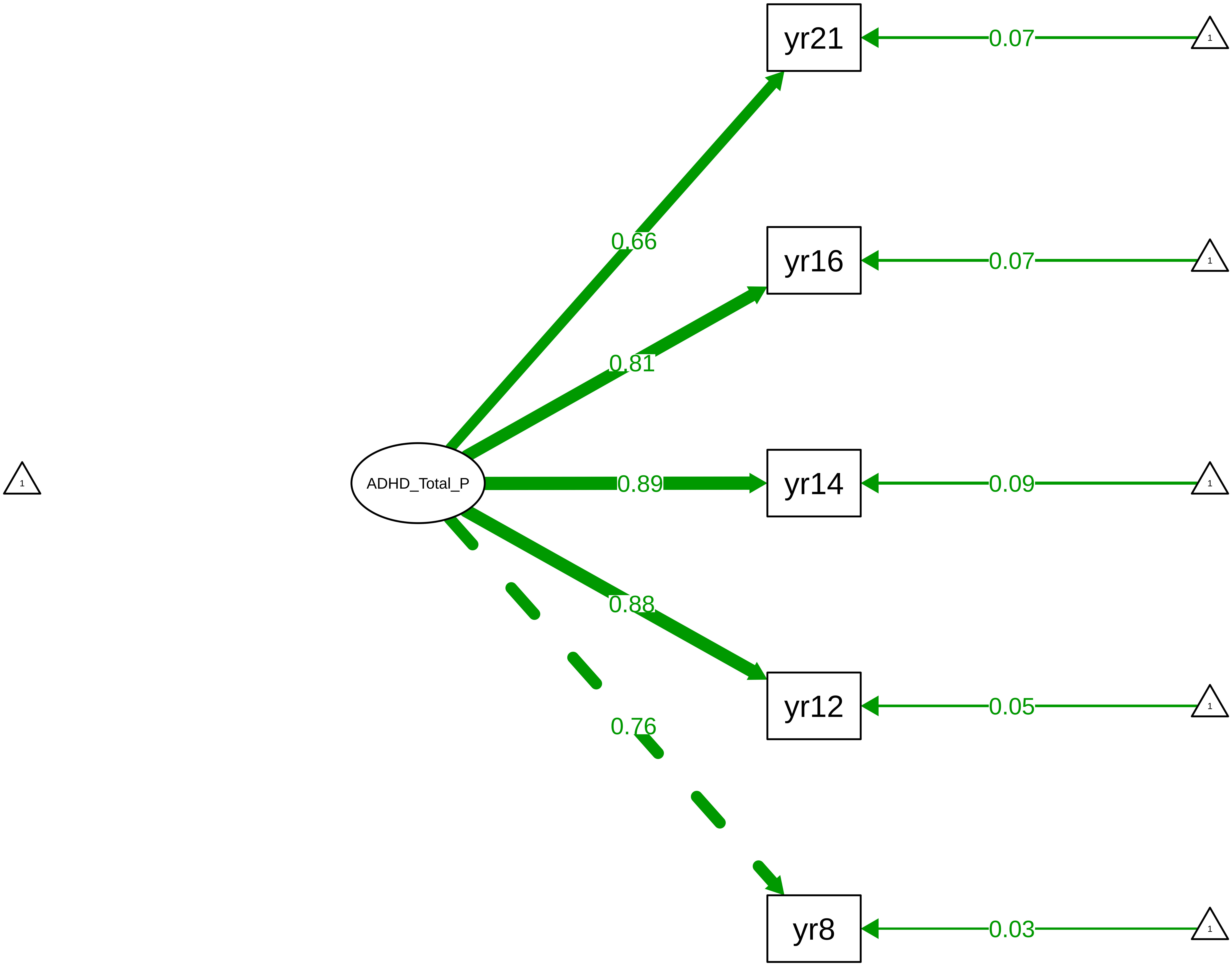

**Figure S8_37: Conners Total − Parent Stage**

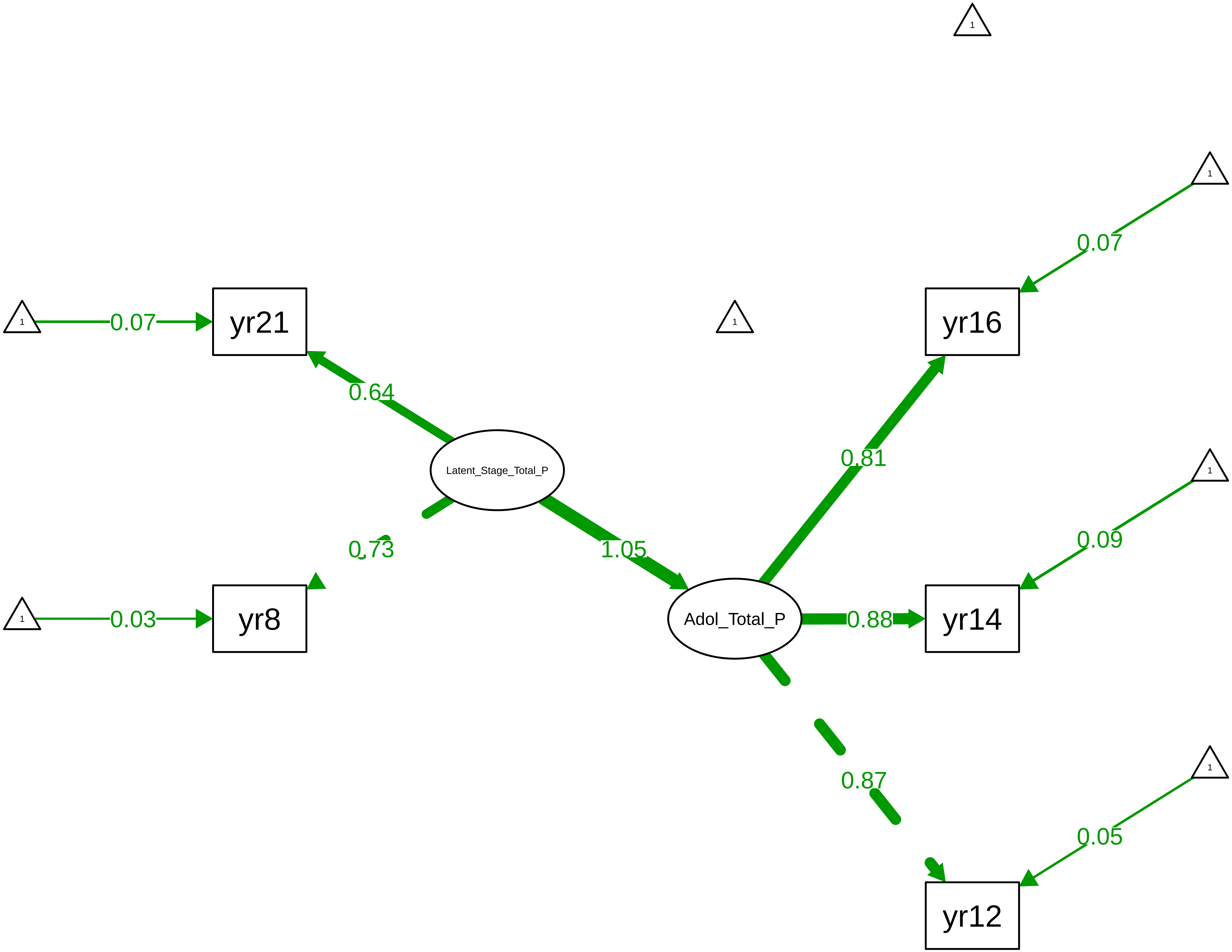

**Figure S8_38: SDQ Conduct − CTCR Overall**

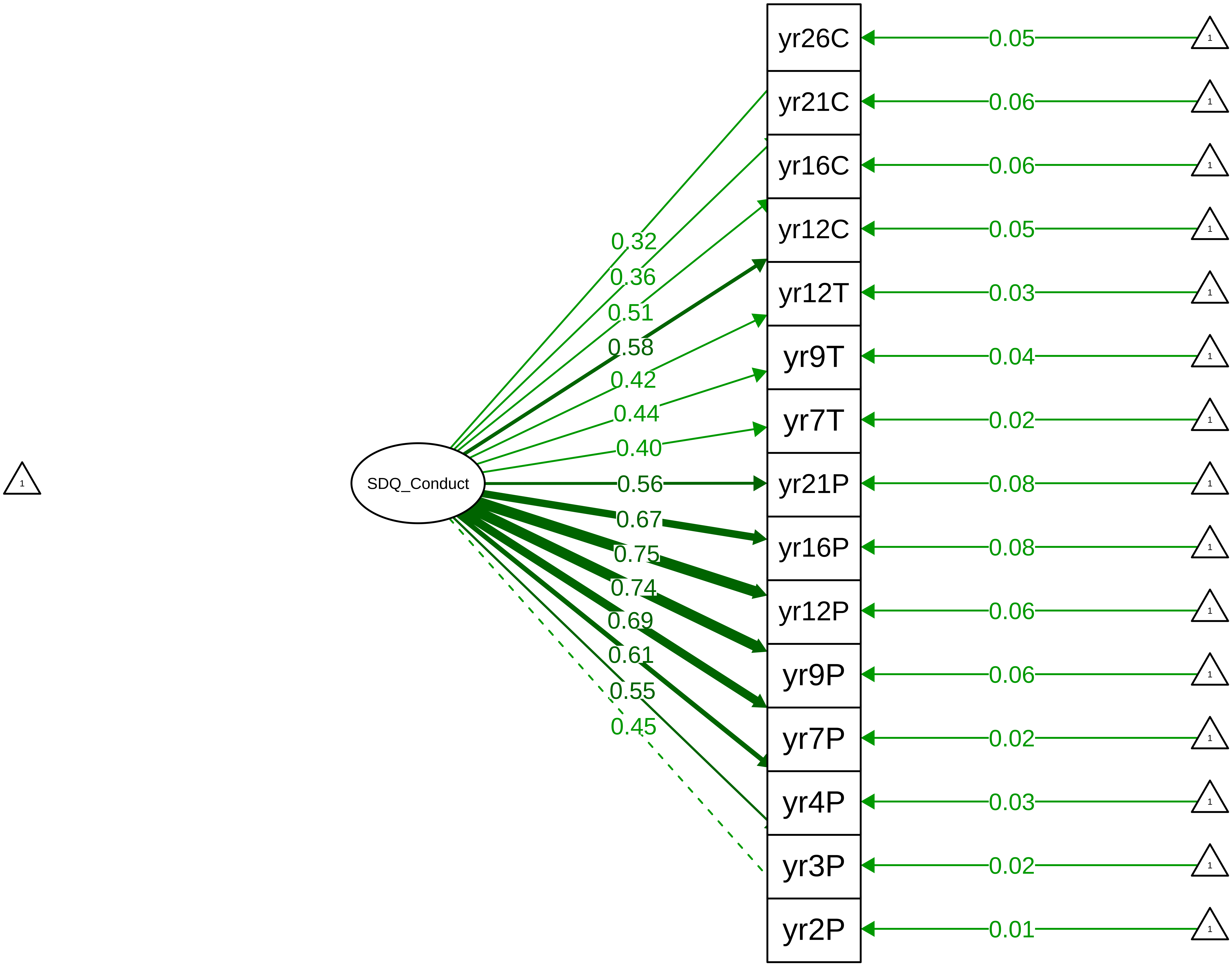

**Figure S8_39: SDQ Conduct − CTCR Stage**

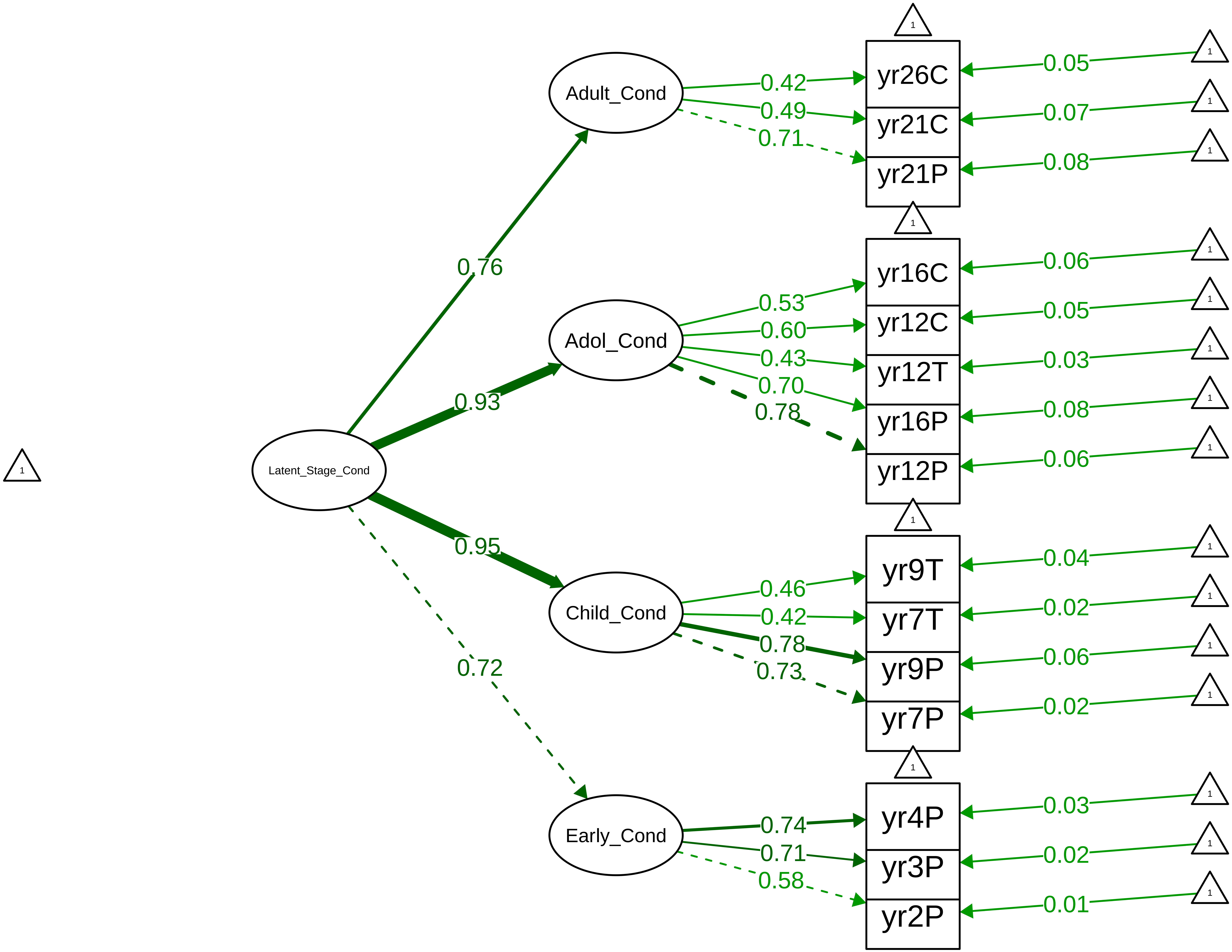

Figure S8_40: SDQ Conduct − Parent Overall

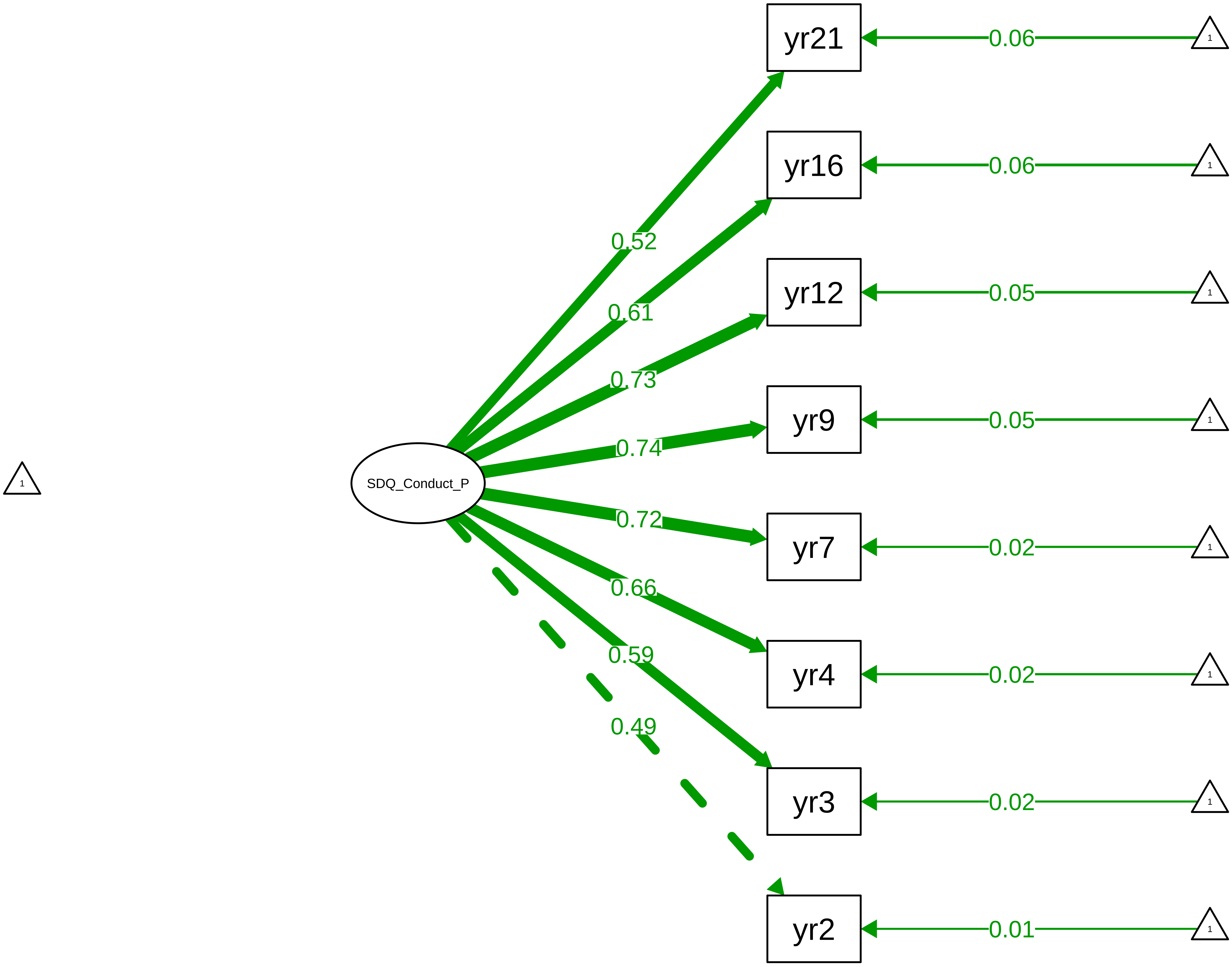

Figure S8_41: SDQ Conduct – Parent Stage

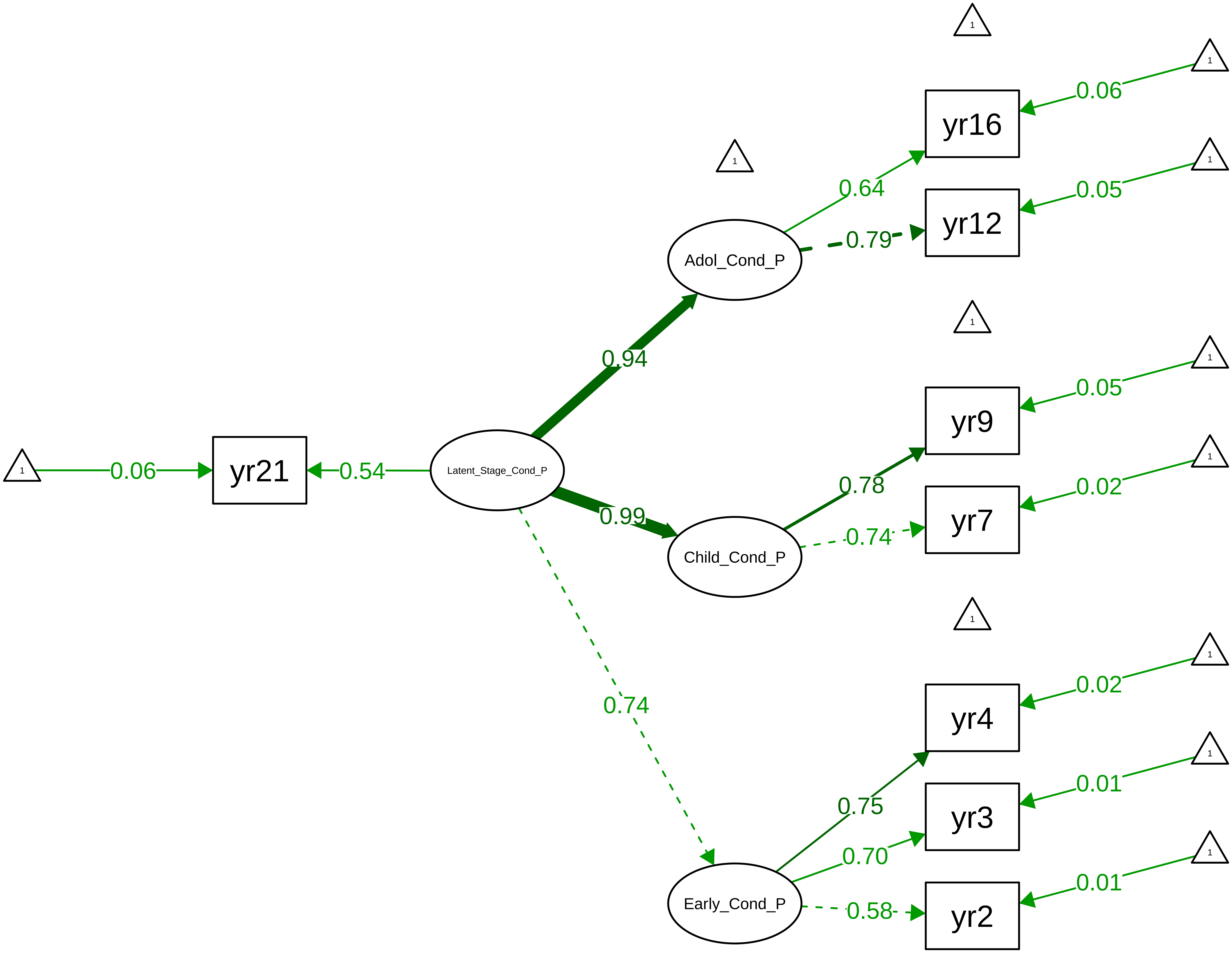

Figure S8_42: SDQ Conduct − Teacher Overall

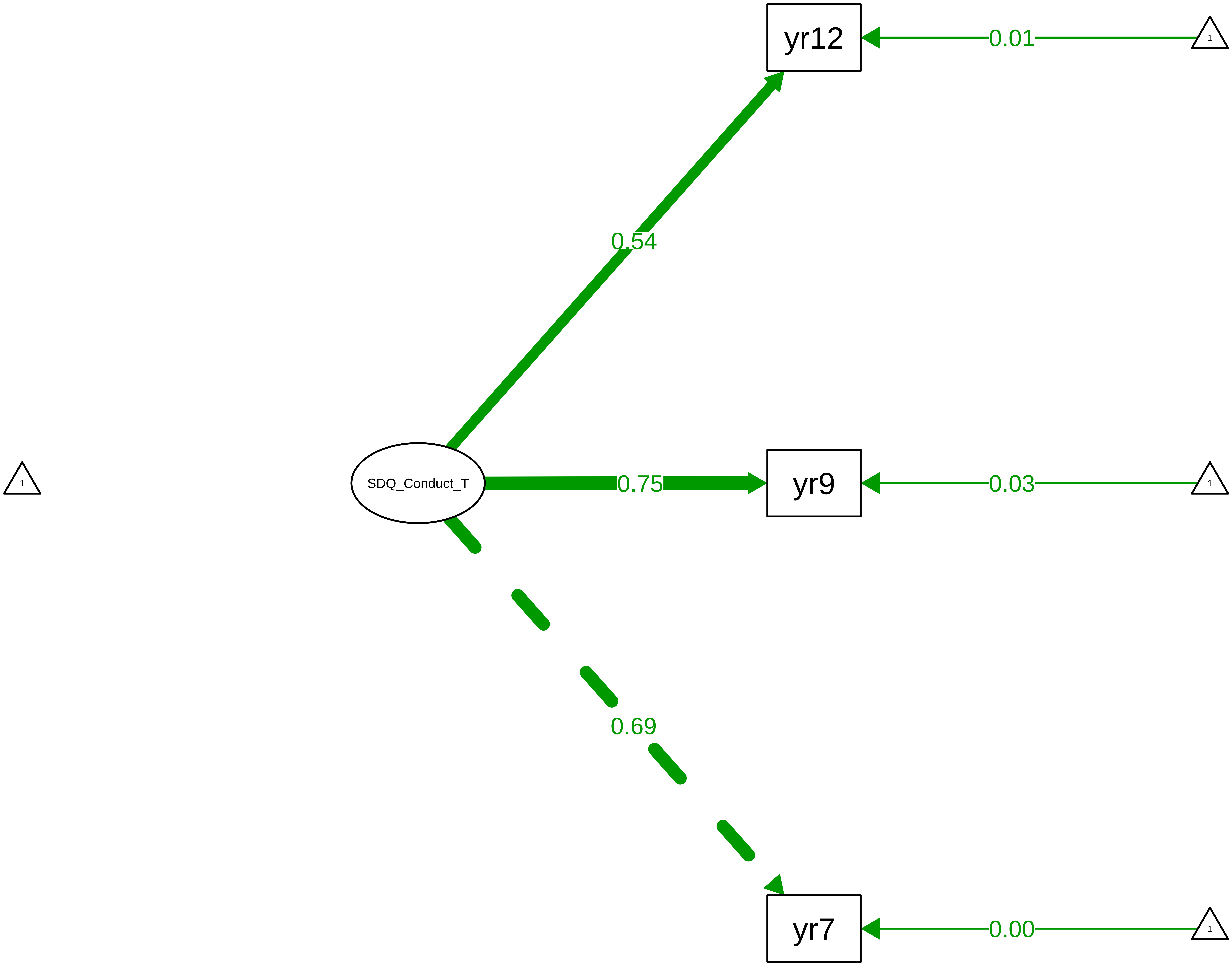

Figure S8_43: SDQ Conduct – Child Overall

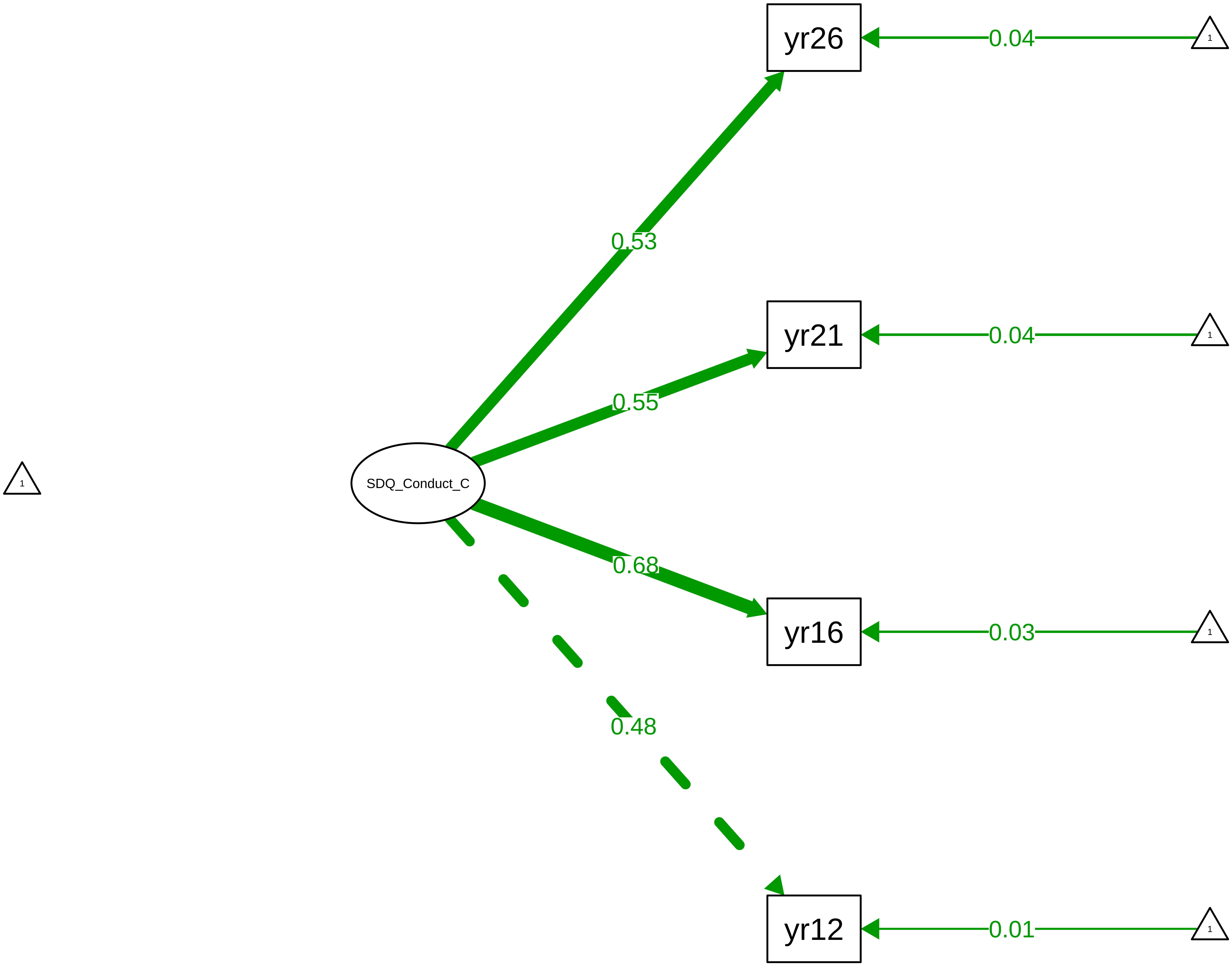

**Figure S8_44: SDQ Emotion − CTCR Overall**

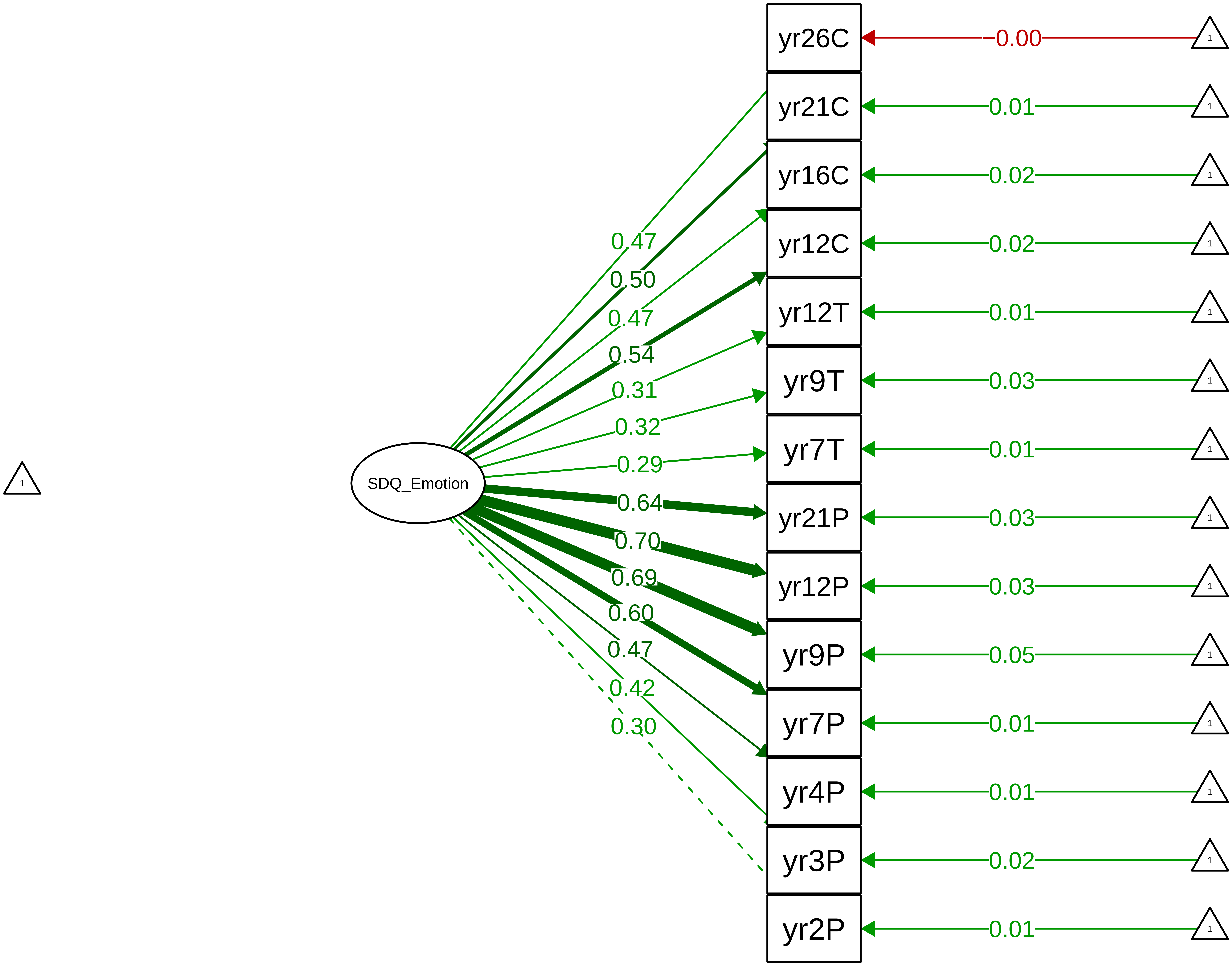

#### Figure S8_45: SDQ Emotion − CTCR Stage

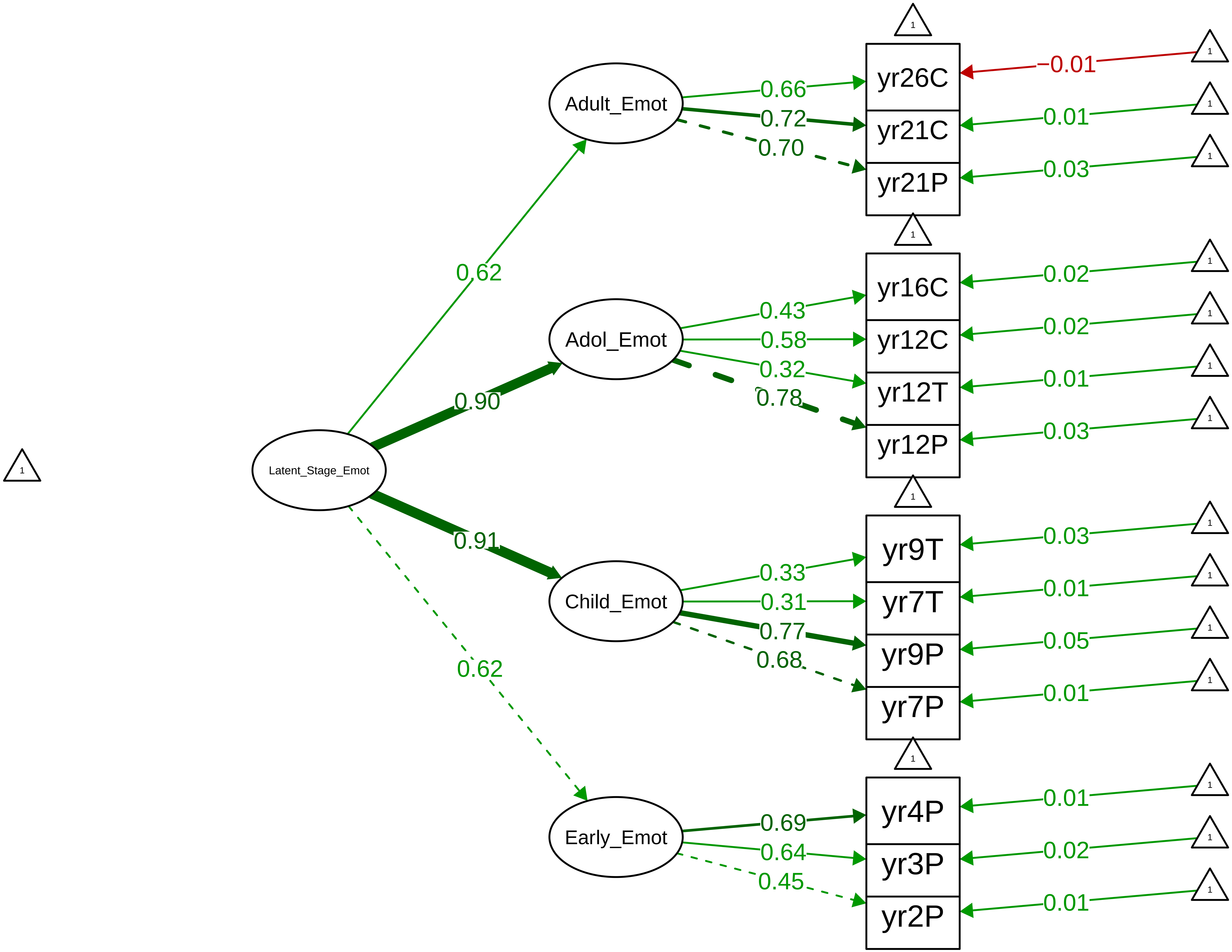

Figure S8_46: SDQ Emotion − Parent Overall

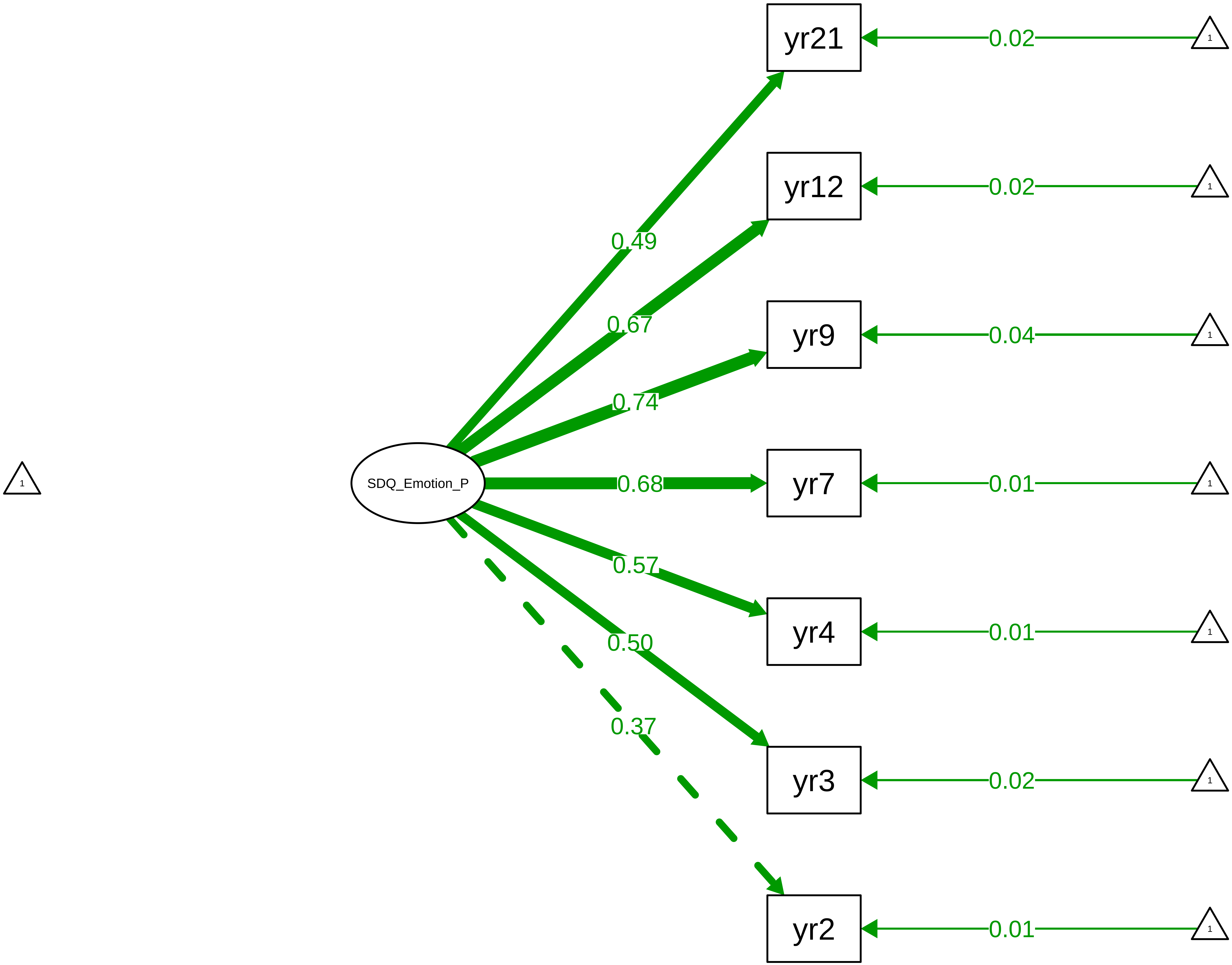

**Figure S8_47: SDQ Emotion − Parent Stage**

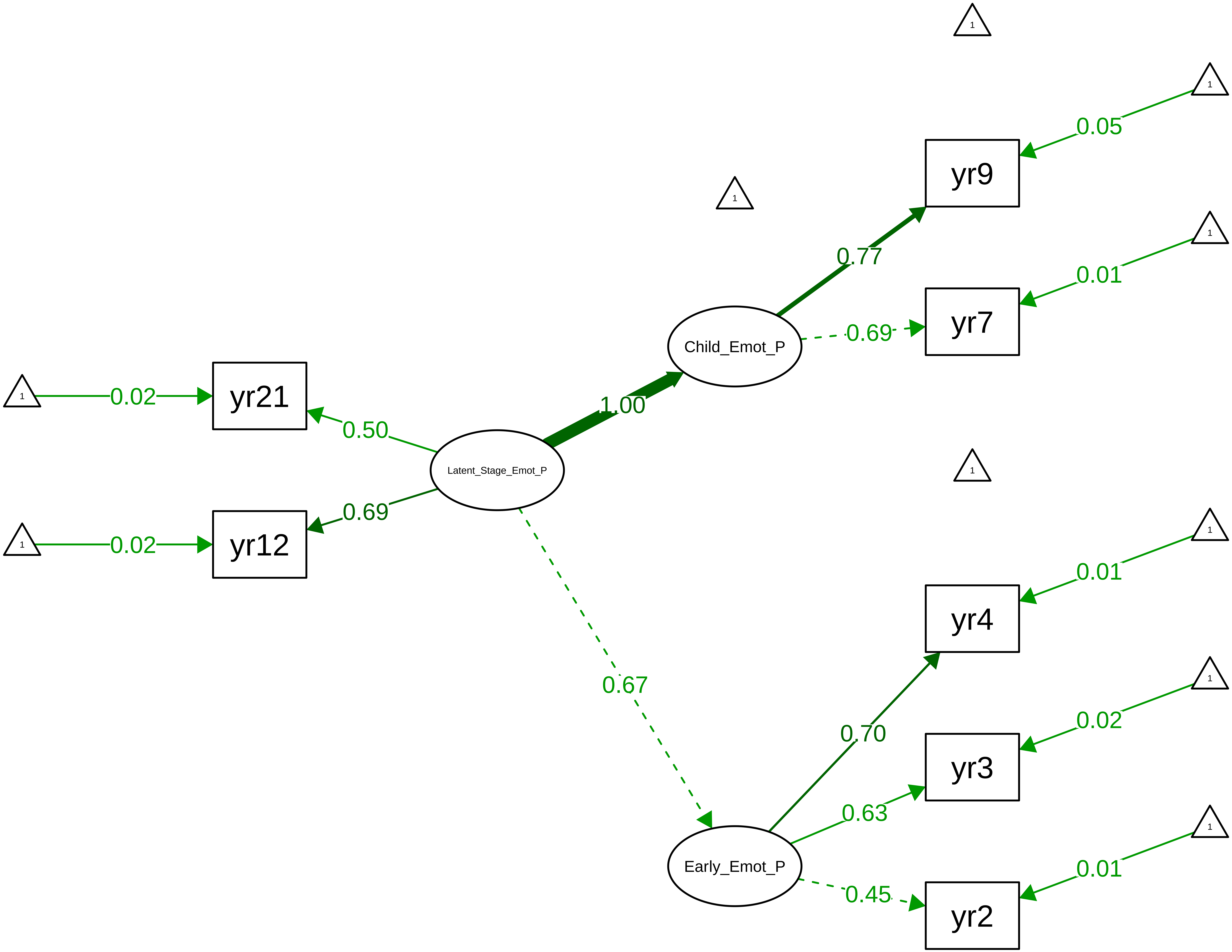

**Figure S8_48: SDQ Emotion − Teacher Overall**

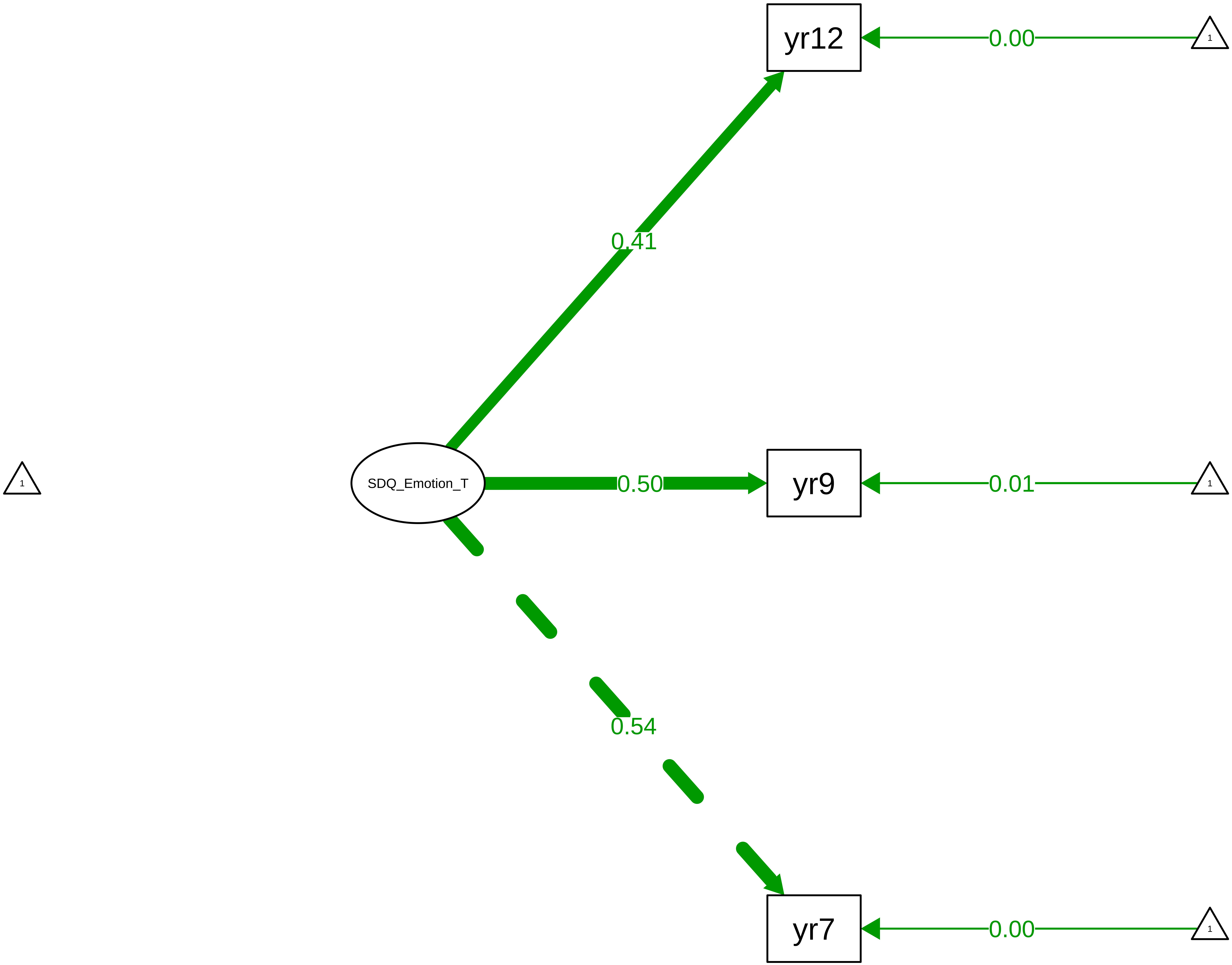

**Figure S8_49: SDQ Emotion − Child Overall**

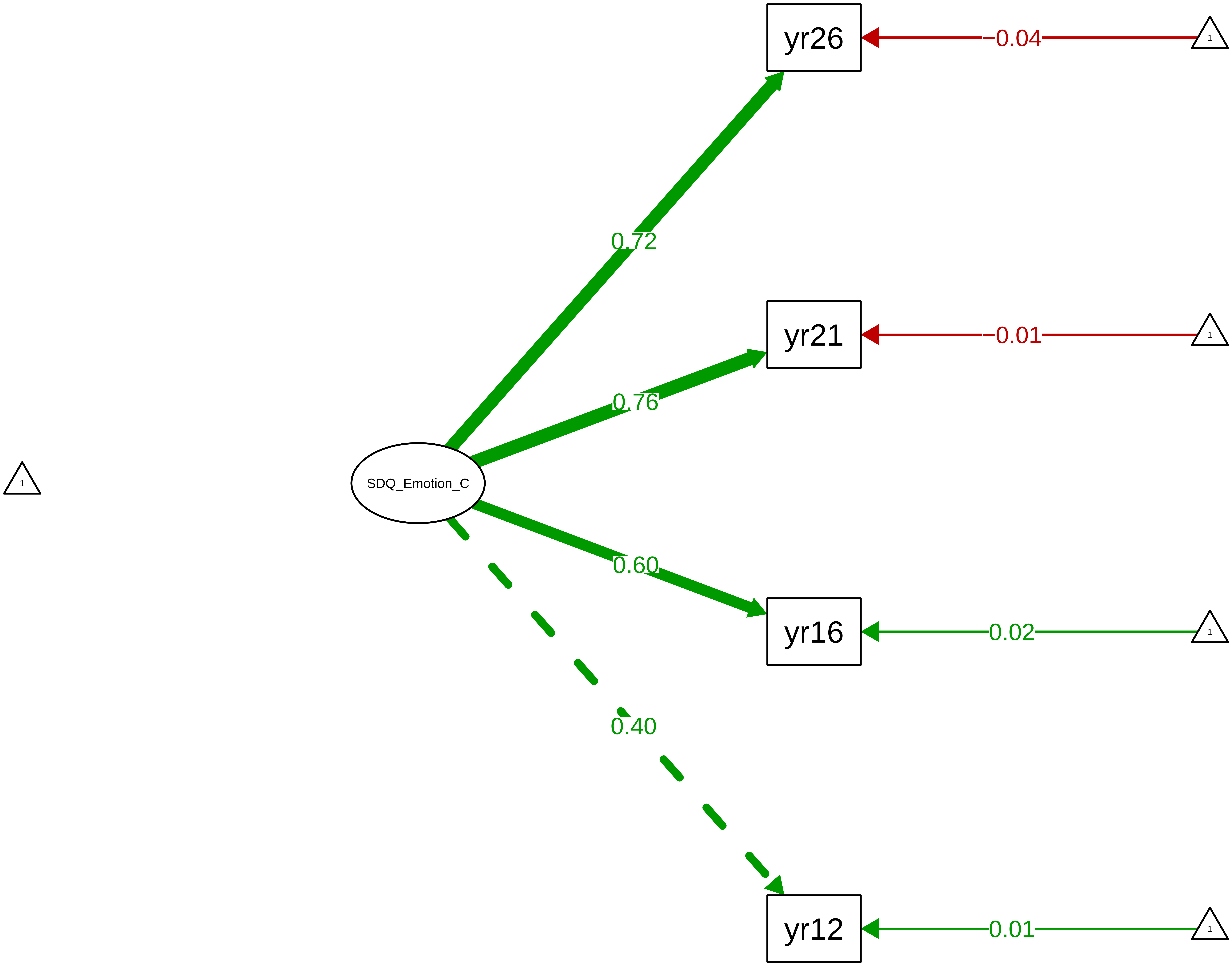

**Figure S8_50: SDQ Hyperactivity − CTCR Overall**

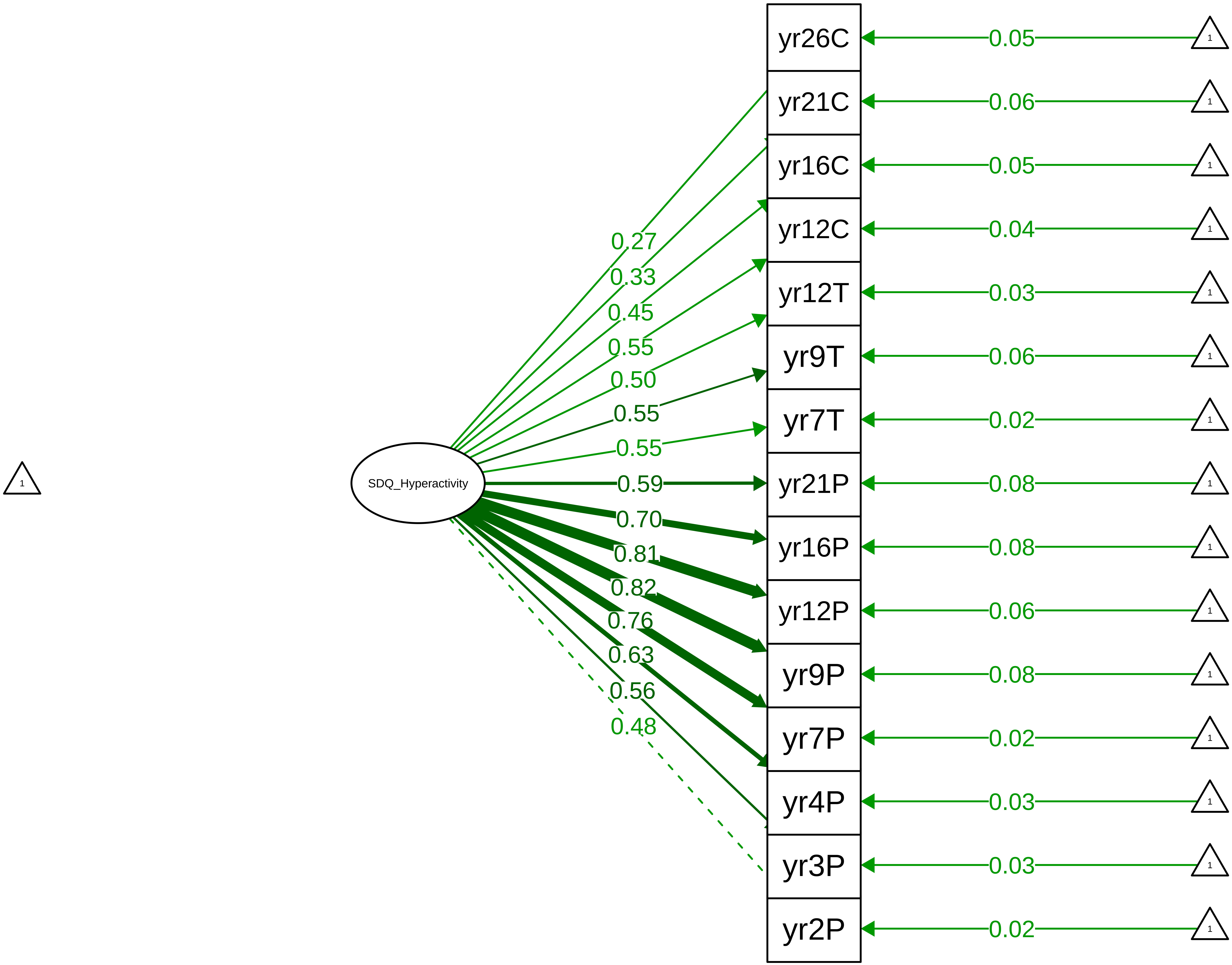

#### Figure S8_51: SDQ Hyperactivity – CTCR Stage

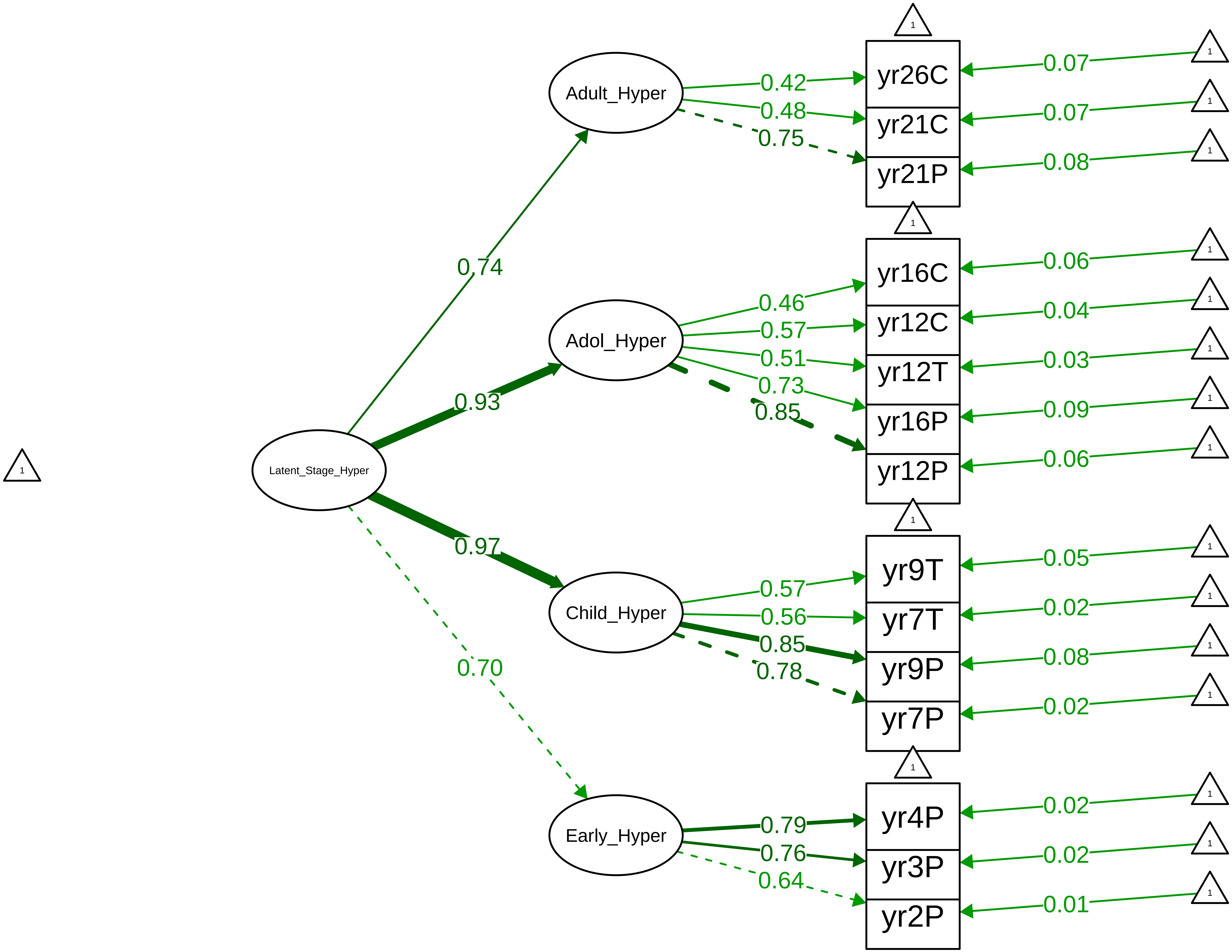

**Figure S8_52: SDQ Hyperactivity – Parent Overall**

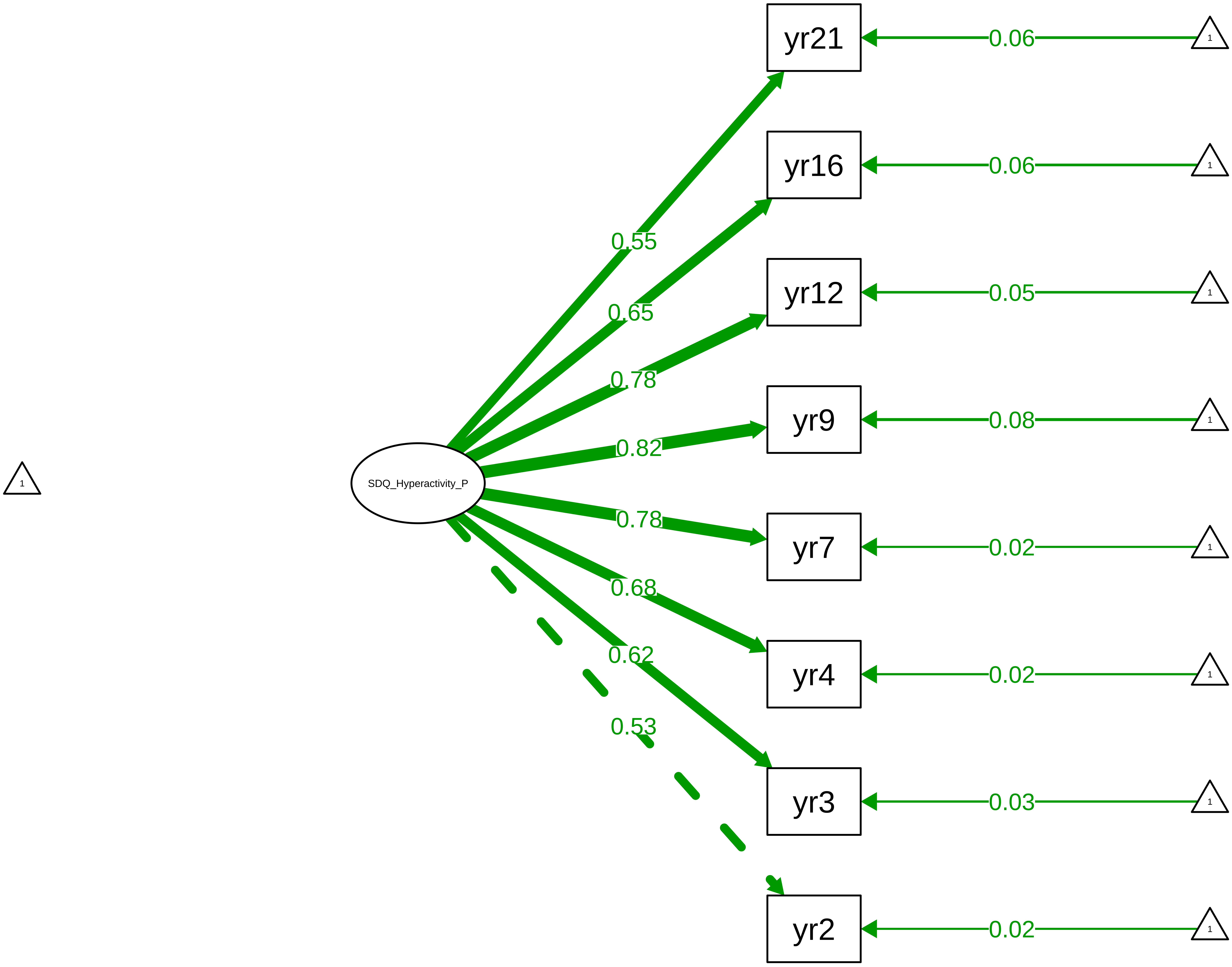

**Figure S8_53: SDQ Hyperactivity – Parent Stage**

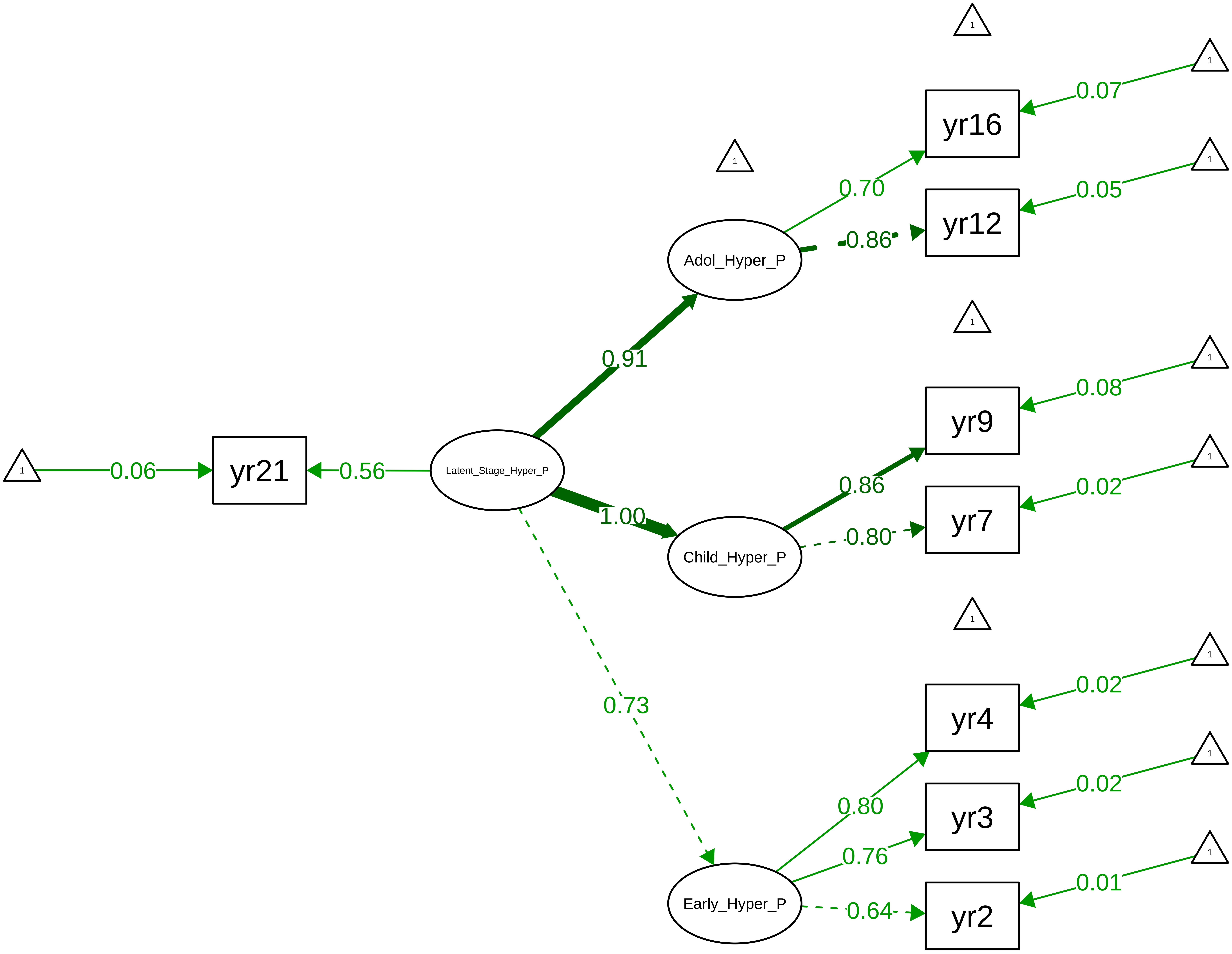

**Figure S8_54: SDQ Hyperactivity − Teacher Overall**

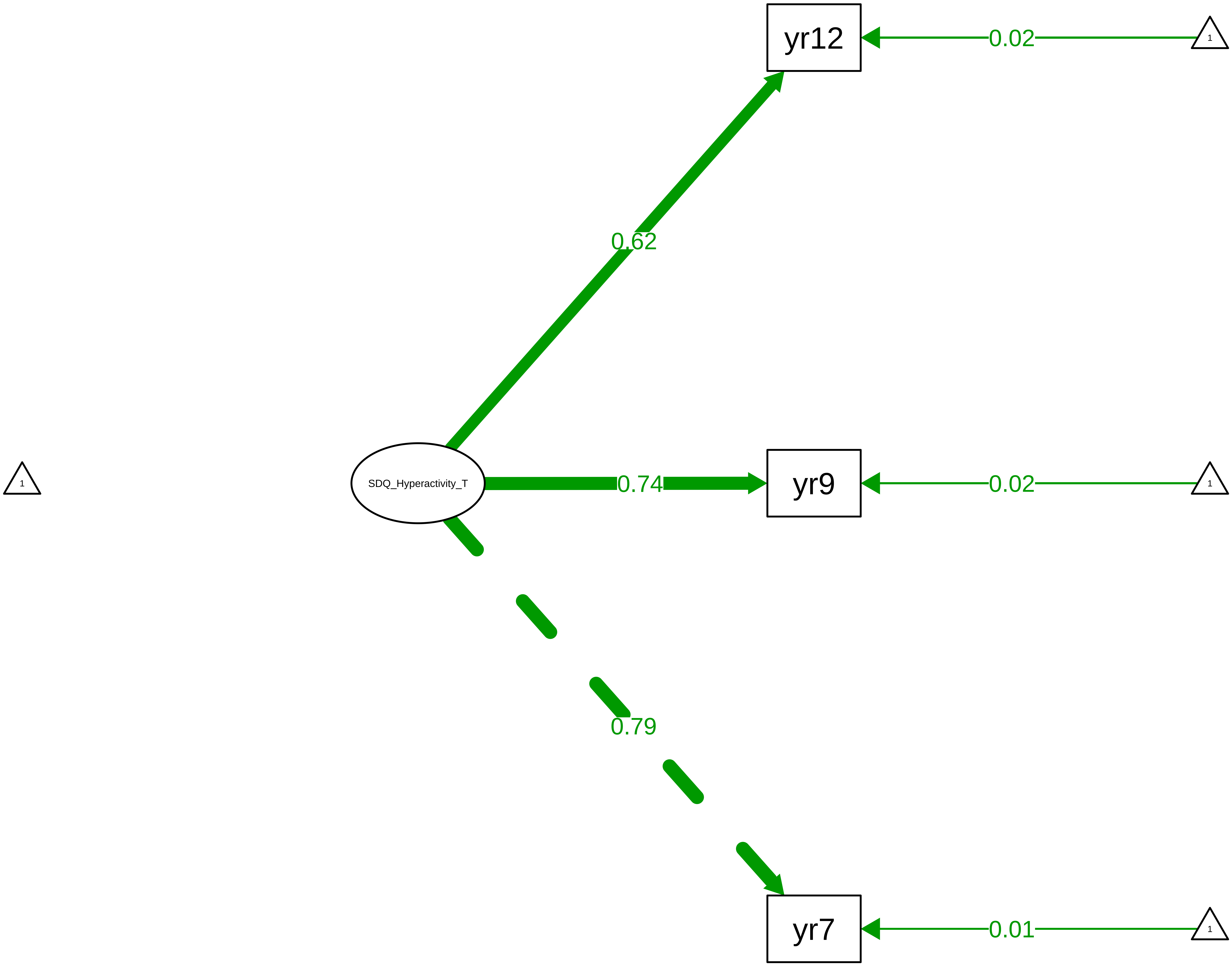

**Figure S8_55: SDQ Hyperactivity − Child Overall**

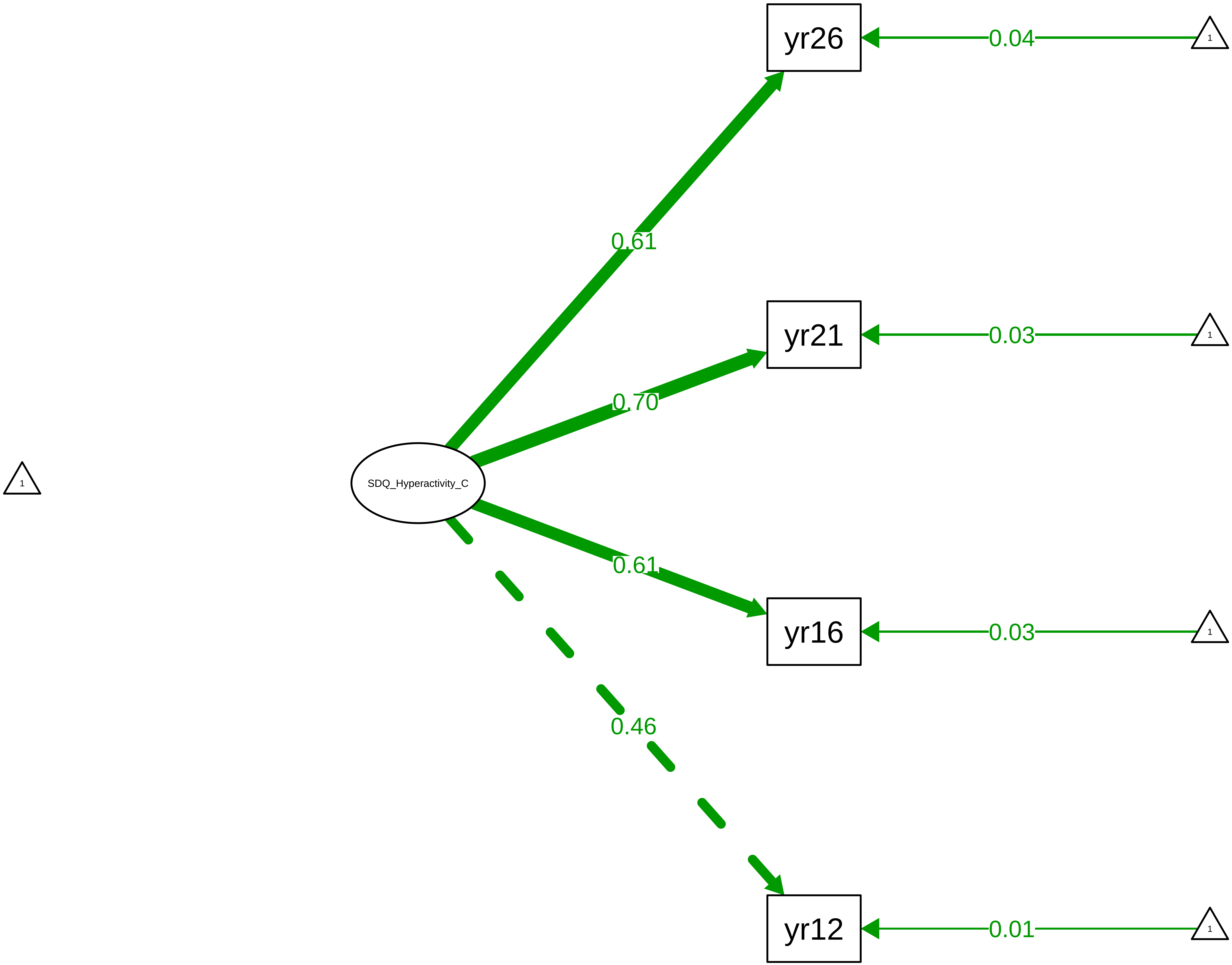

Figure S8_56: SDQ Peer Problems − CTCR Overall

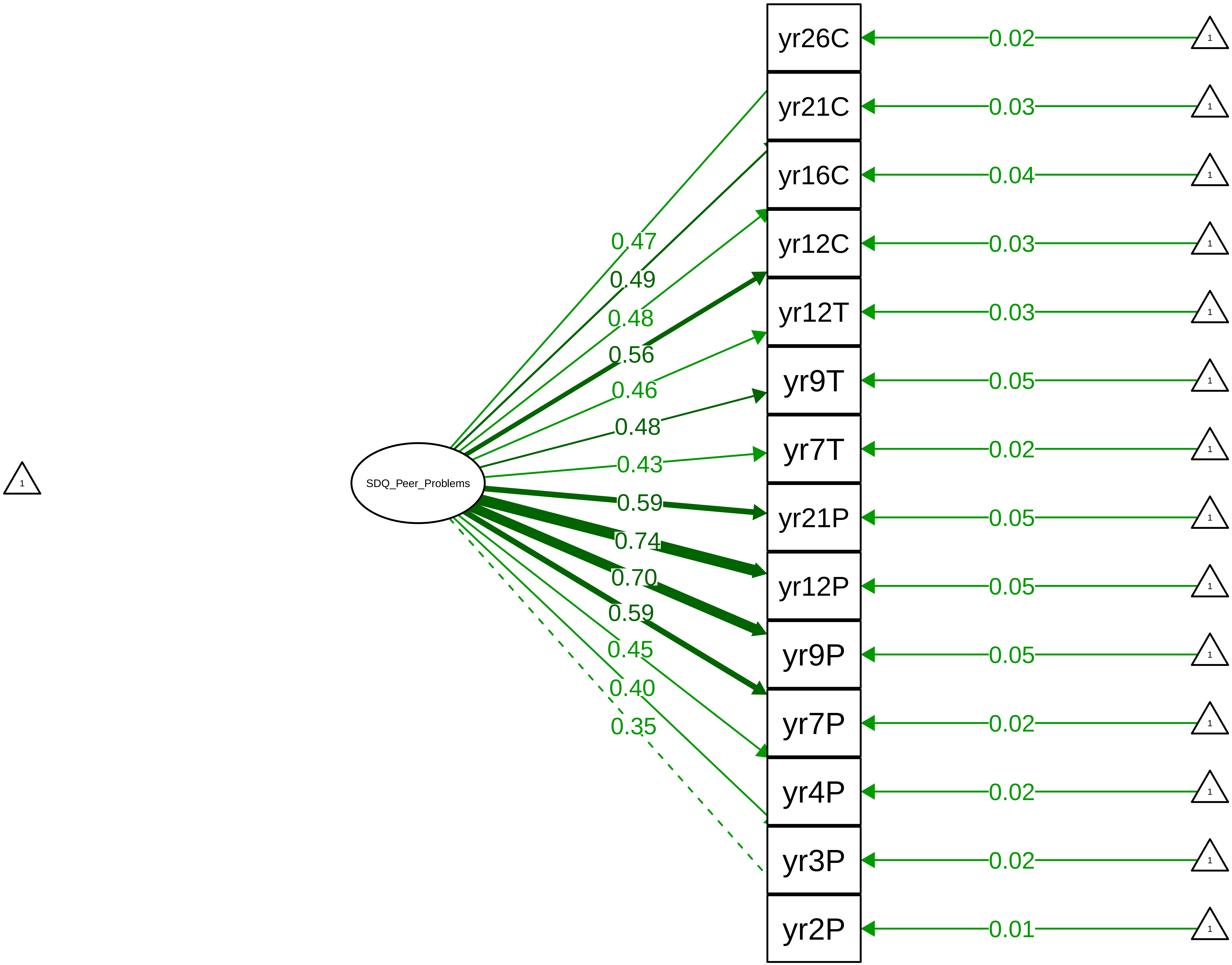

**Figure S8_57: SDQ Peer Problems − CTCR Stage**

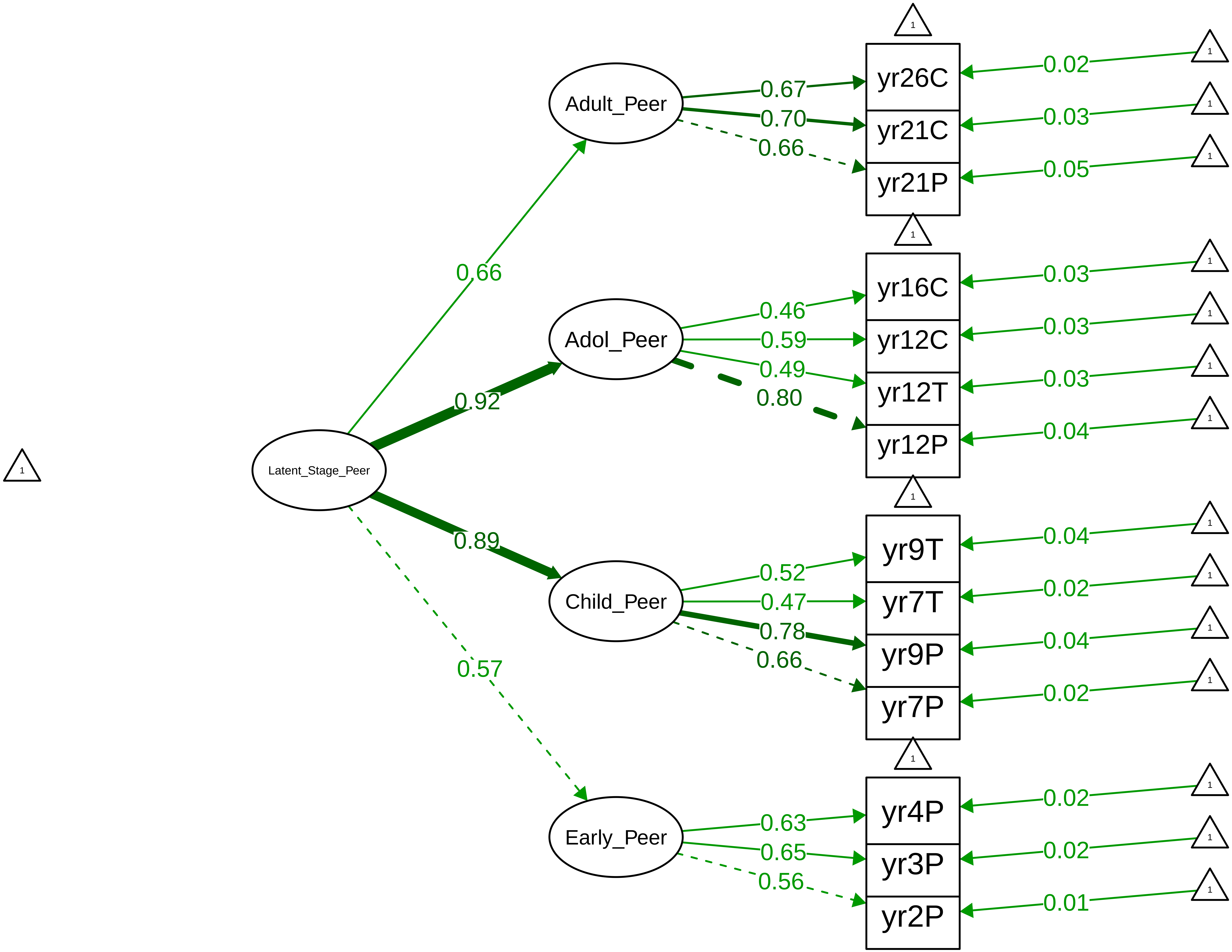

Figure S8_58: SDQ Peer Problems − Parent Overall

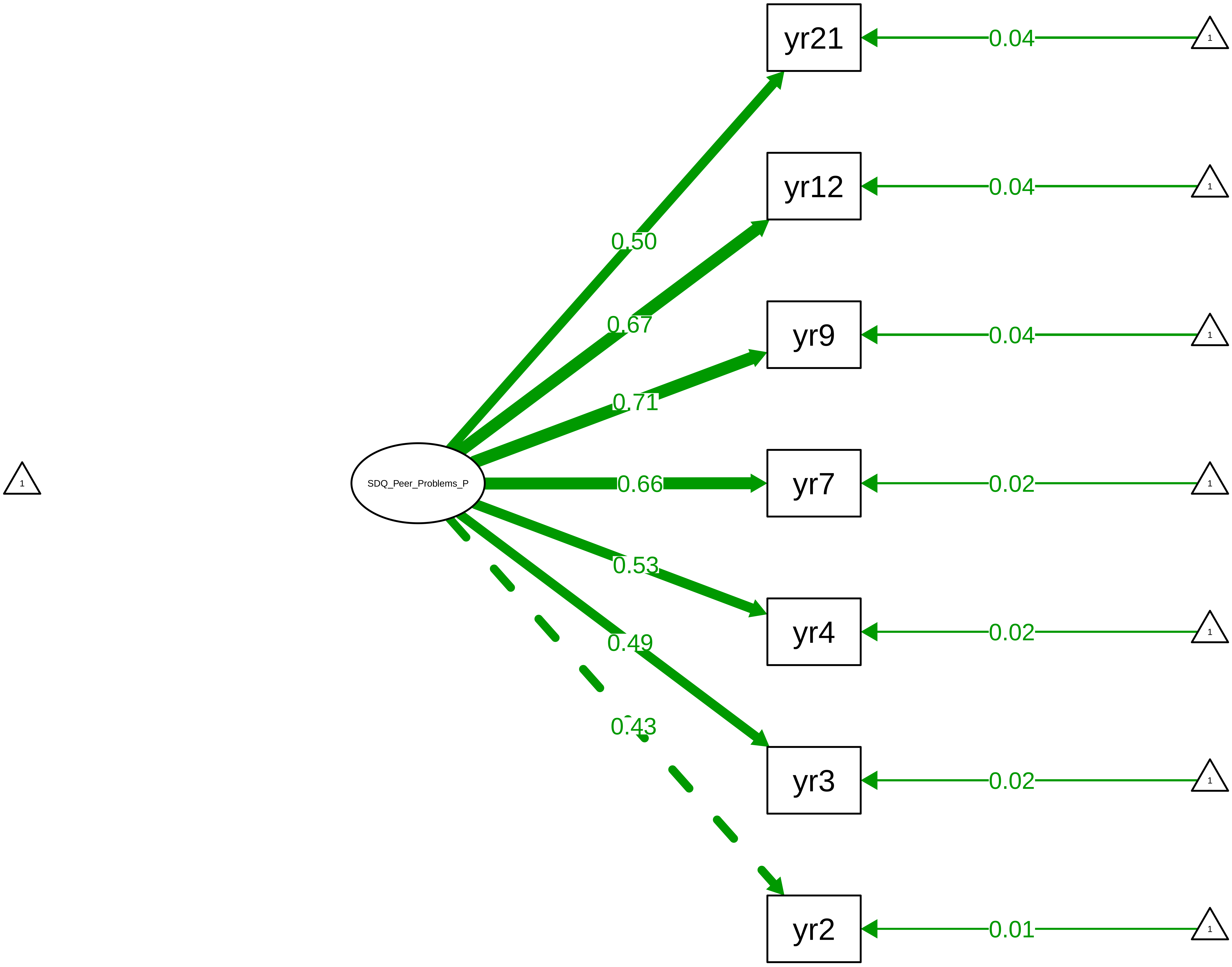

#### Figure S8_59: SDQ Peer Problems – Parent Stage

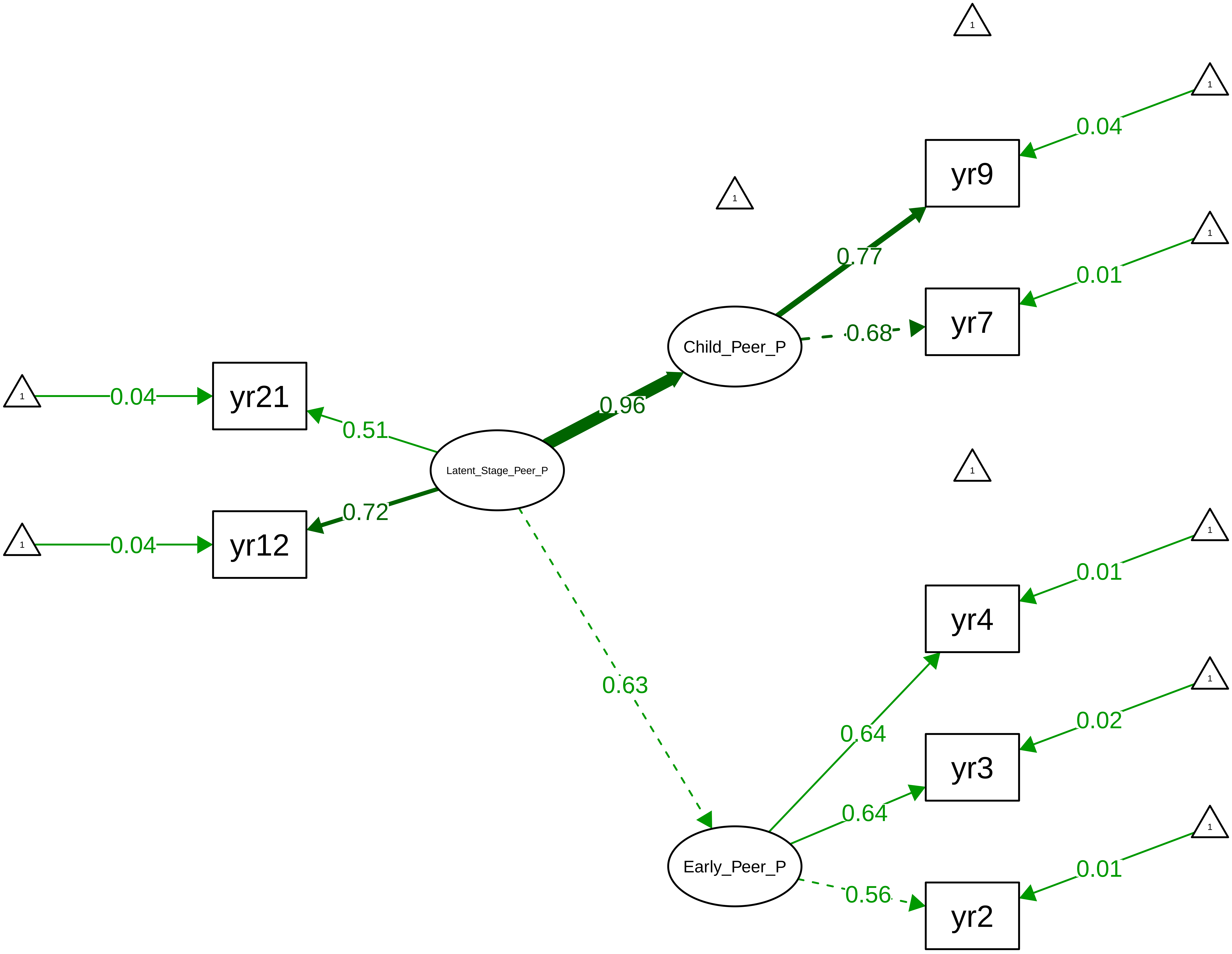

Figure S8_60: SDQ Peer Problems − Teacher Overall

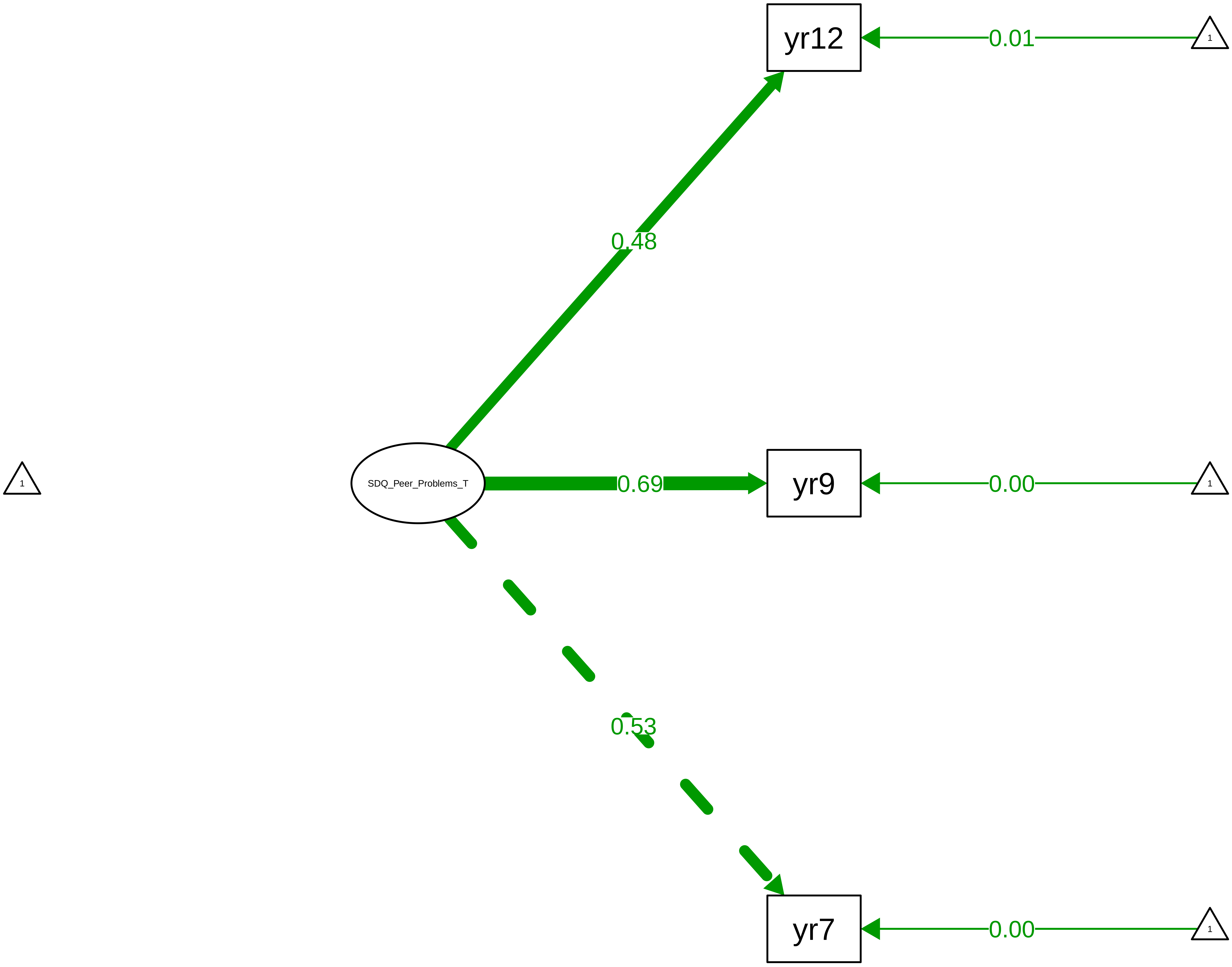

Figure S8_61: SDQ Peer Problems − Child Overall

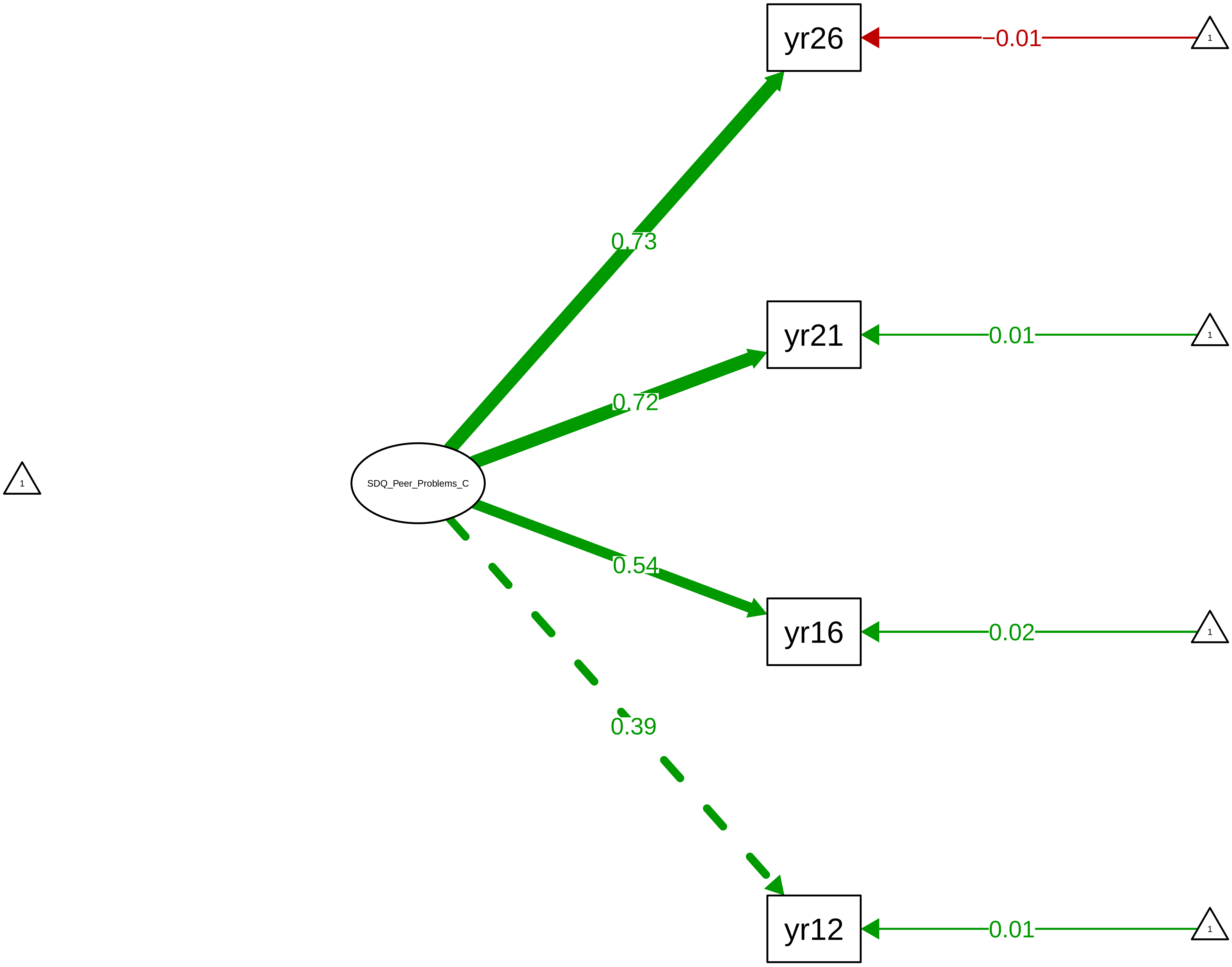

#### Figure S8_62: SDQ Prosocial – CTCR Overall

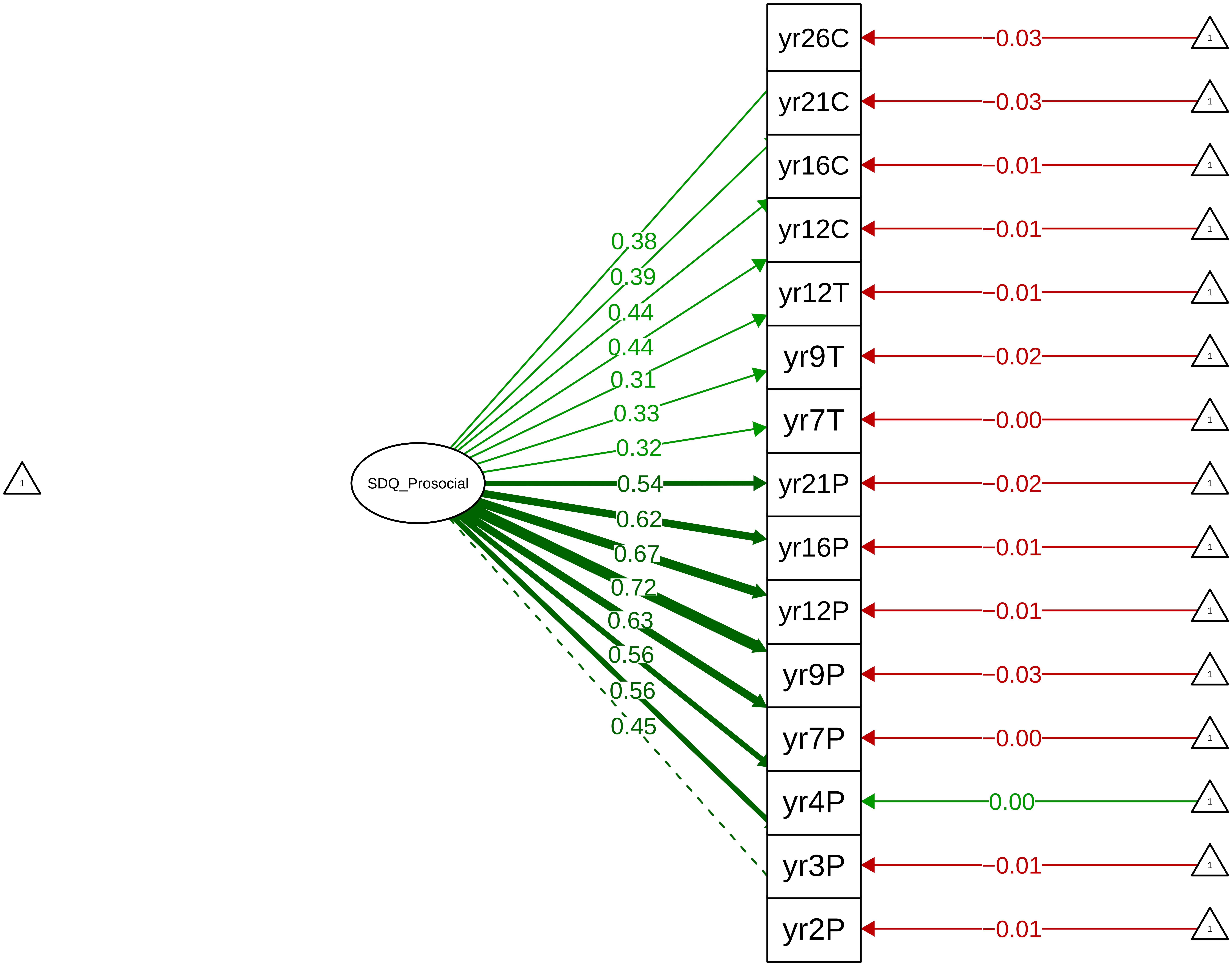

**Figure S8_63: SDQ Prosocial − CTCR Stage**

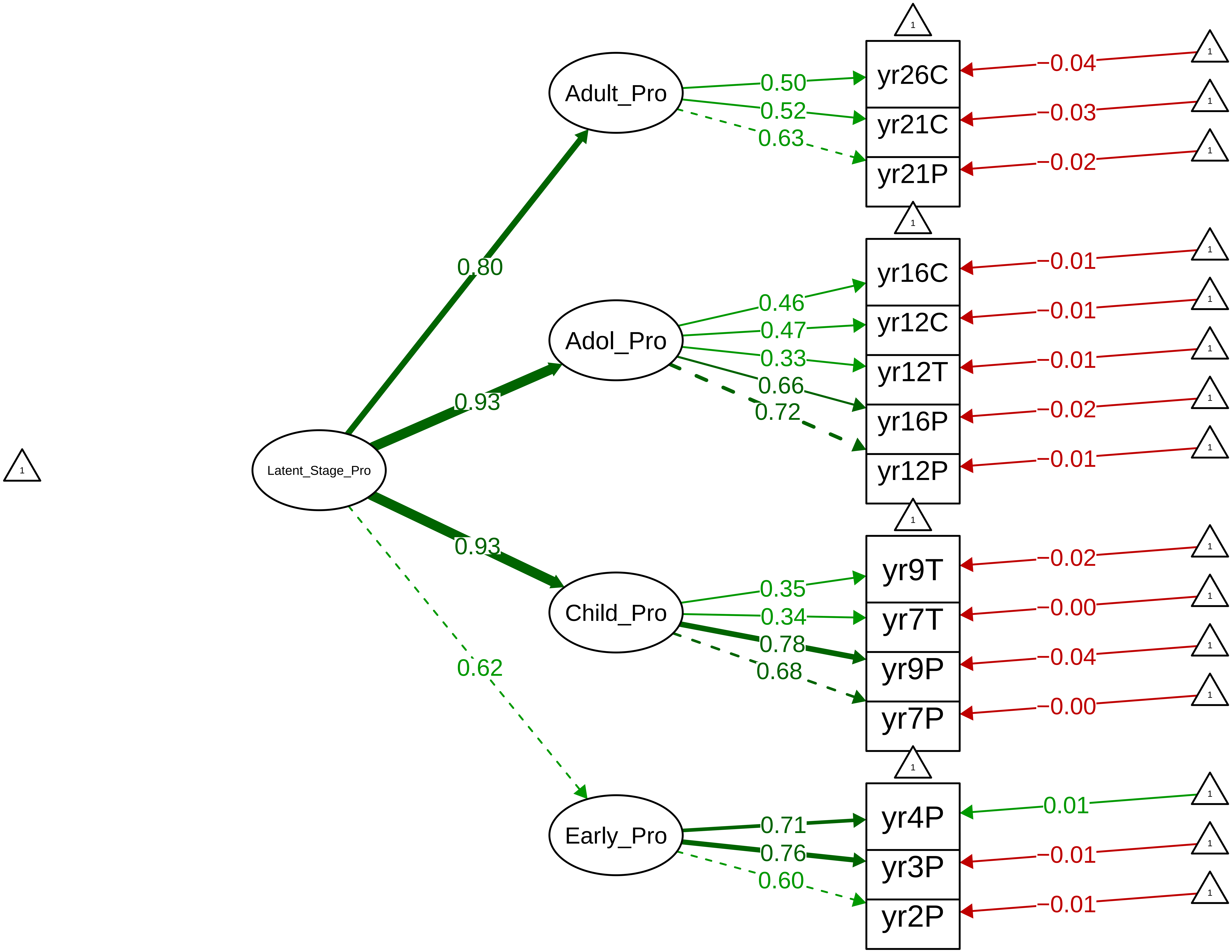

Figure S8_64: SDQ Prosocial − Parent Overall

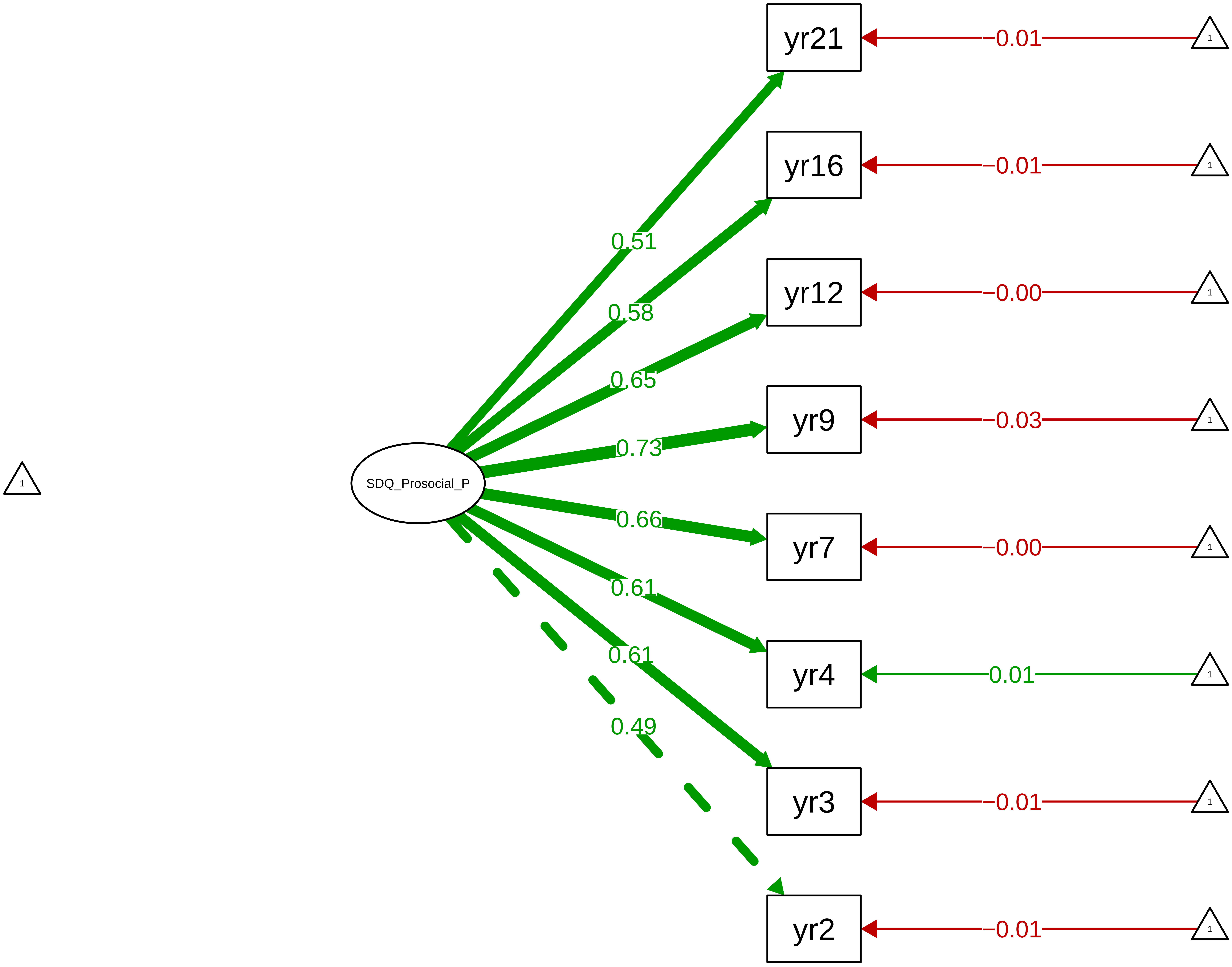

Figure S8_65: SDQ Prosocial – Parent Stage

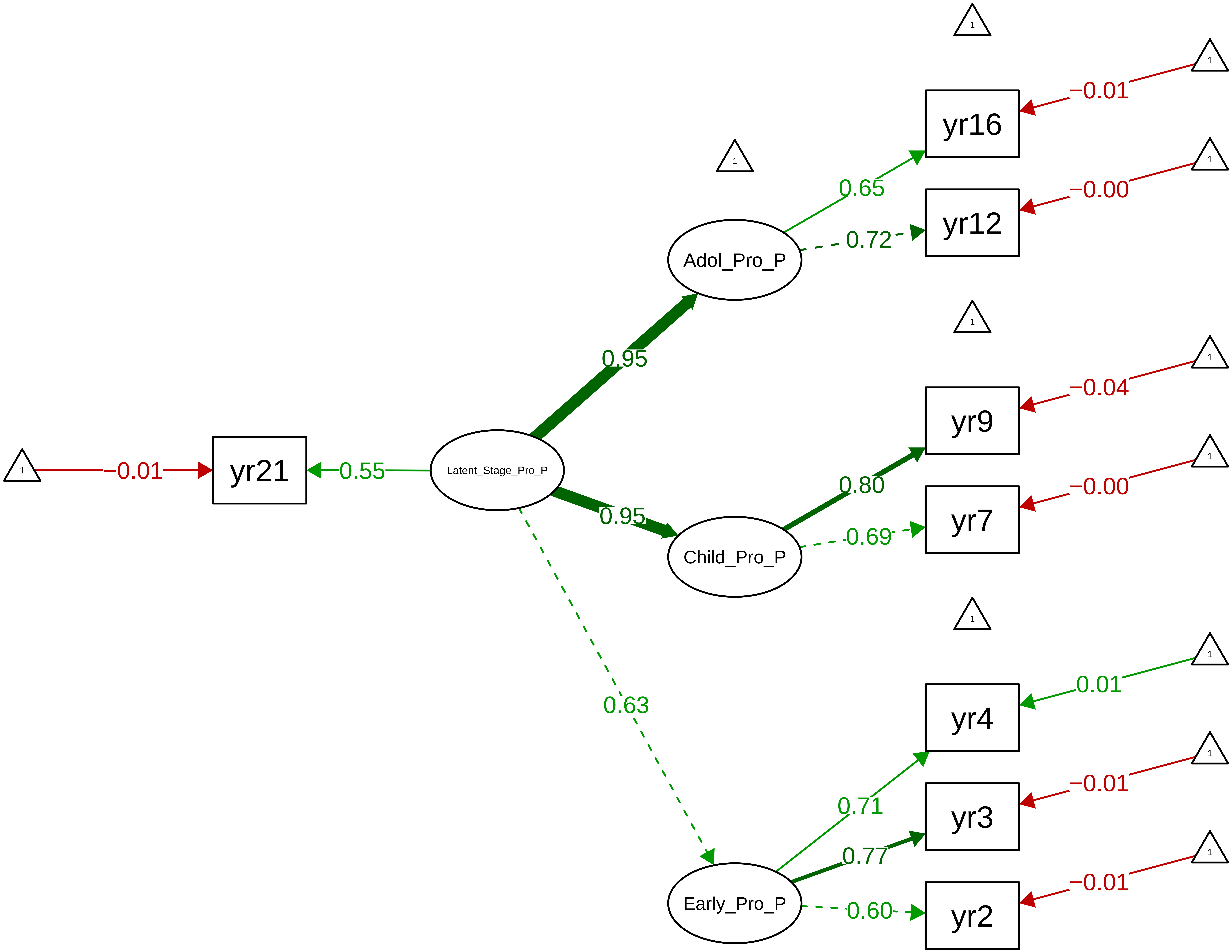

Figure S8_66: SDQ Prosocial − Teacher Overall

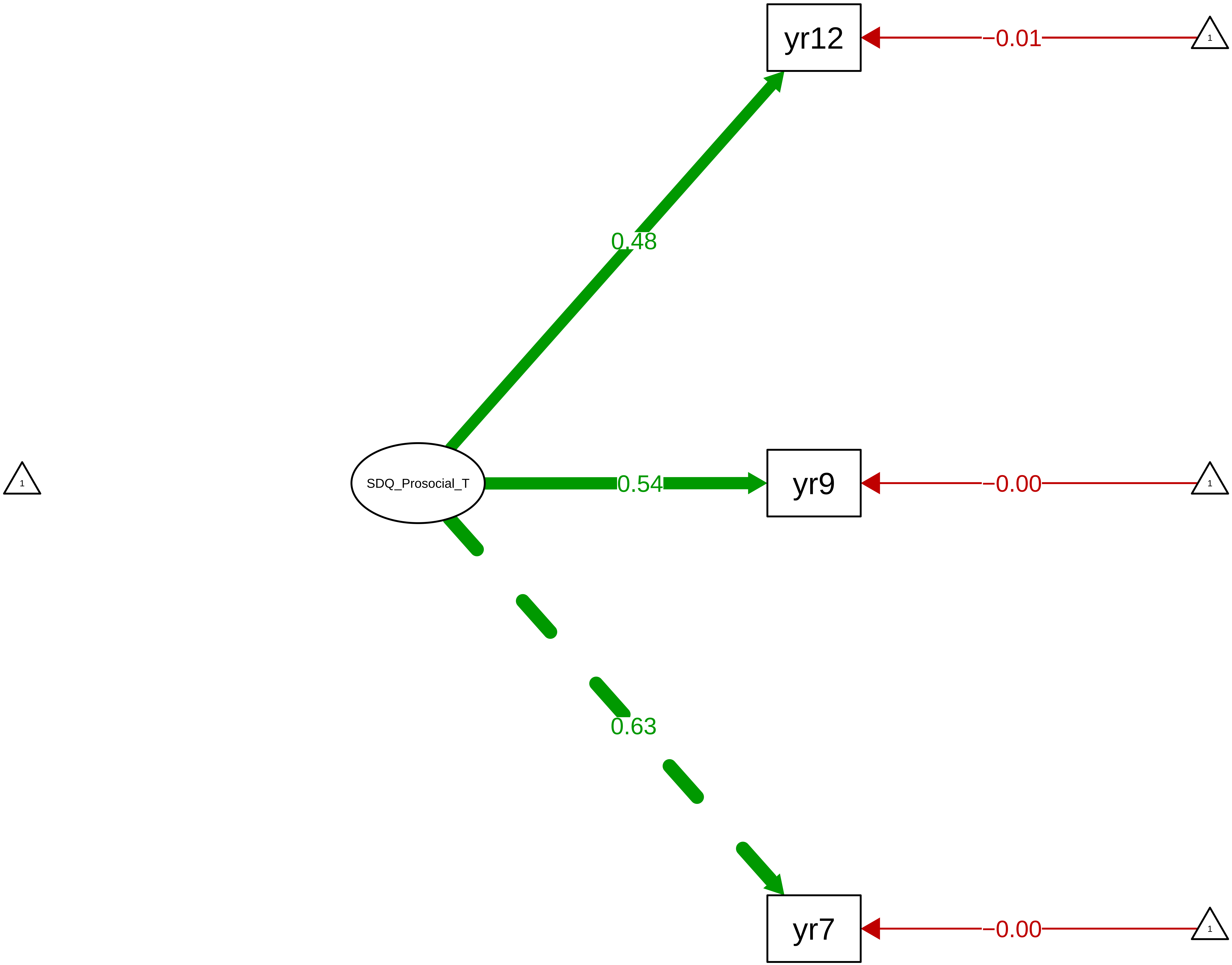

**Figure S8_67: SDQ Prosocial − Child Overall**

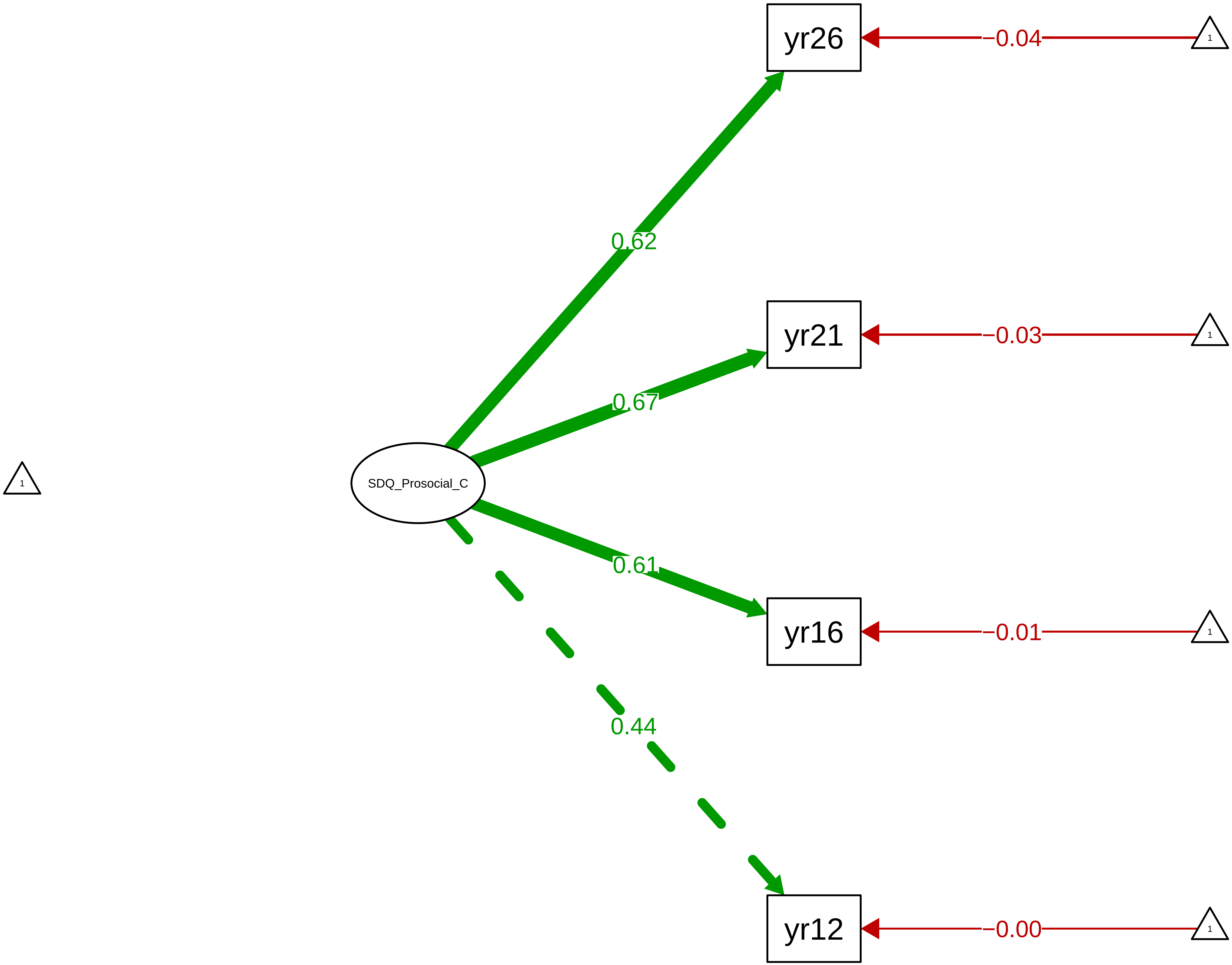

Figure S8_68: SDQ Total Problems − CTCR Overall

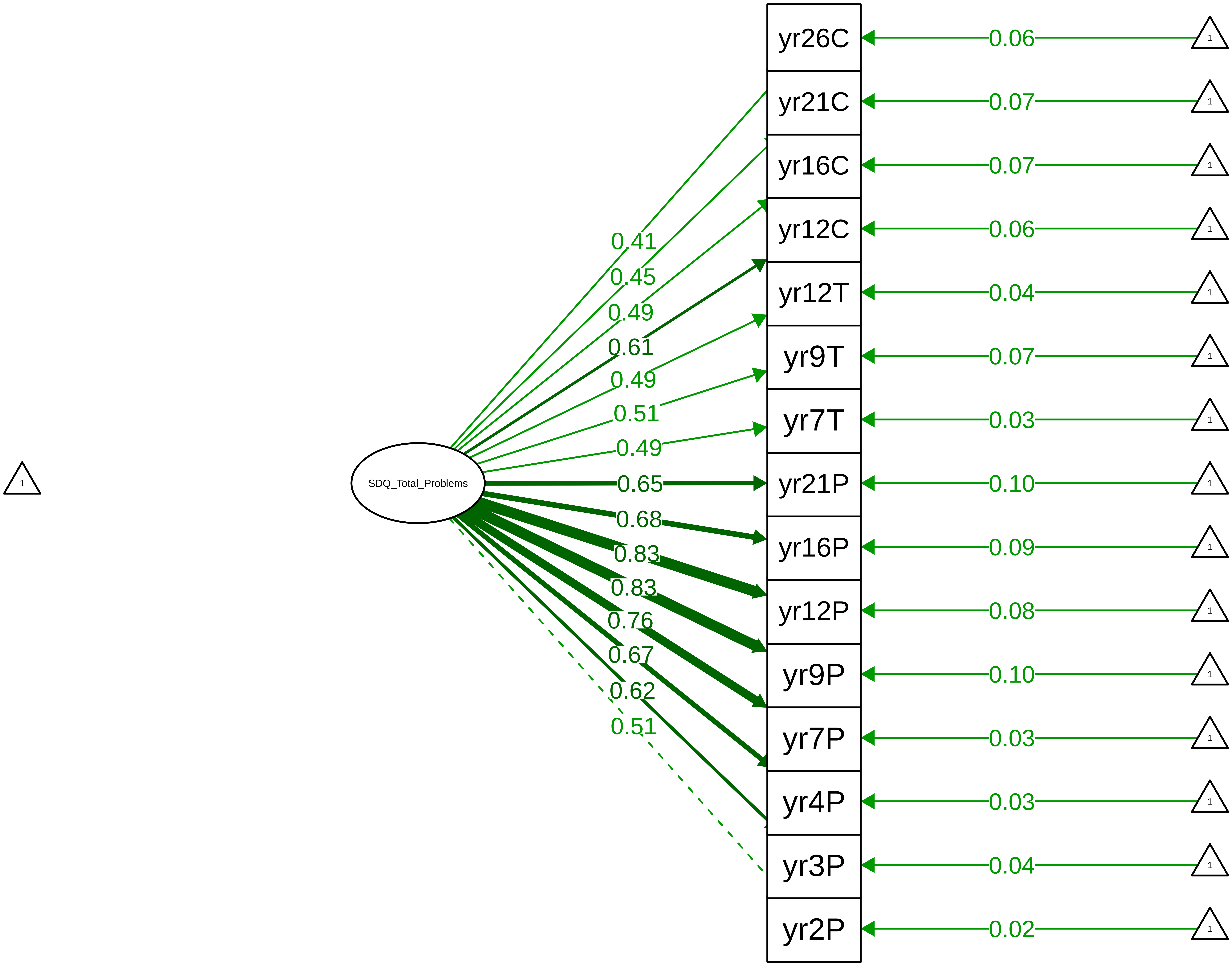

Figure S8_69: SDQ Total Problems – CTCR Stage

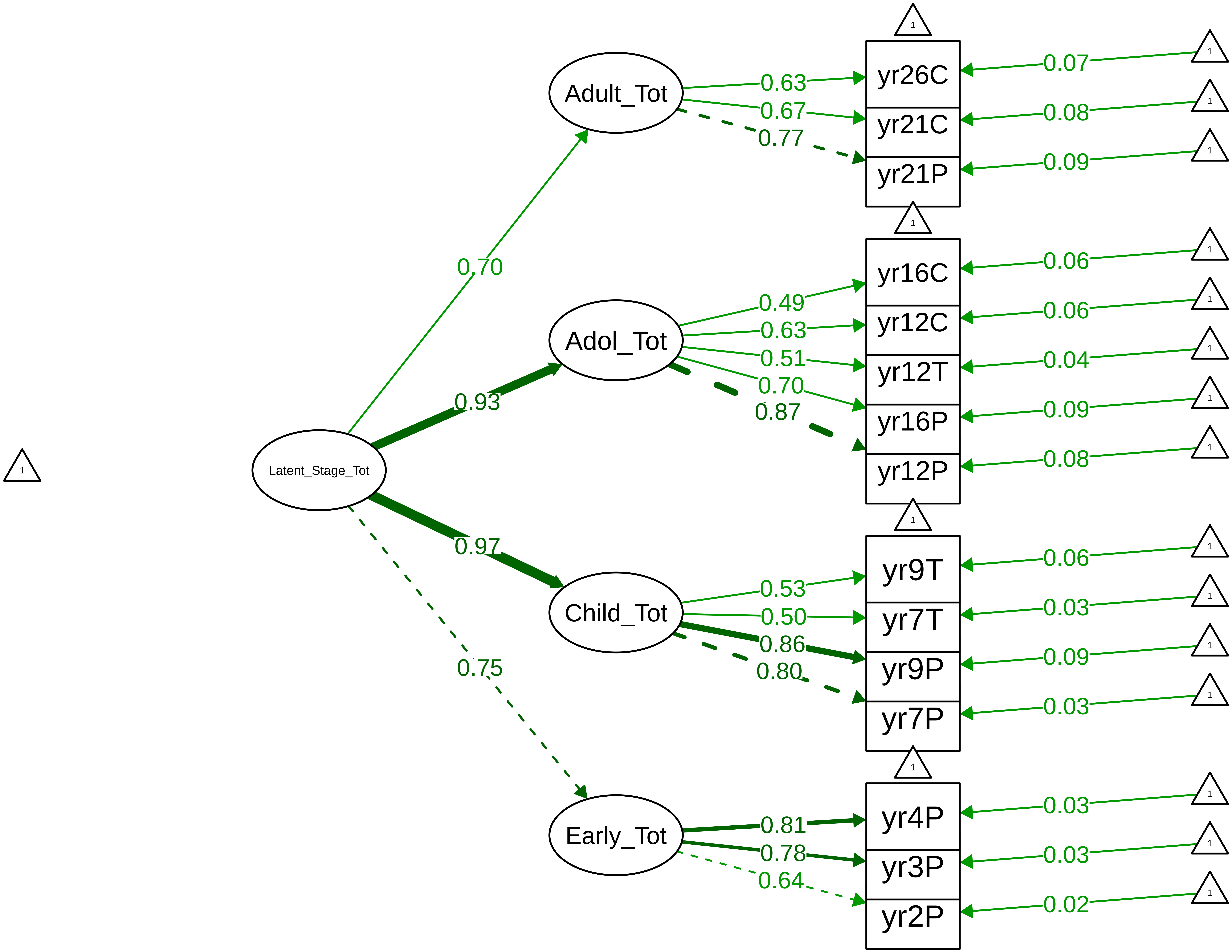

**Figure S8_70: SDQ Total Problems − Parent Overall**

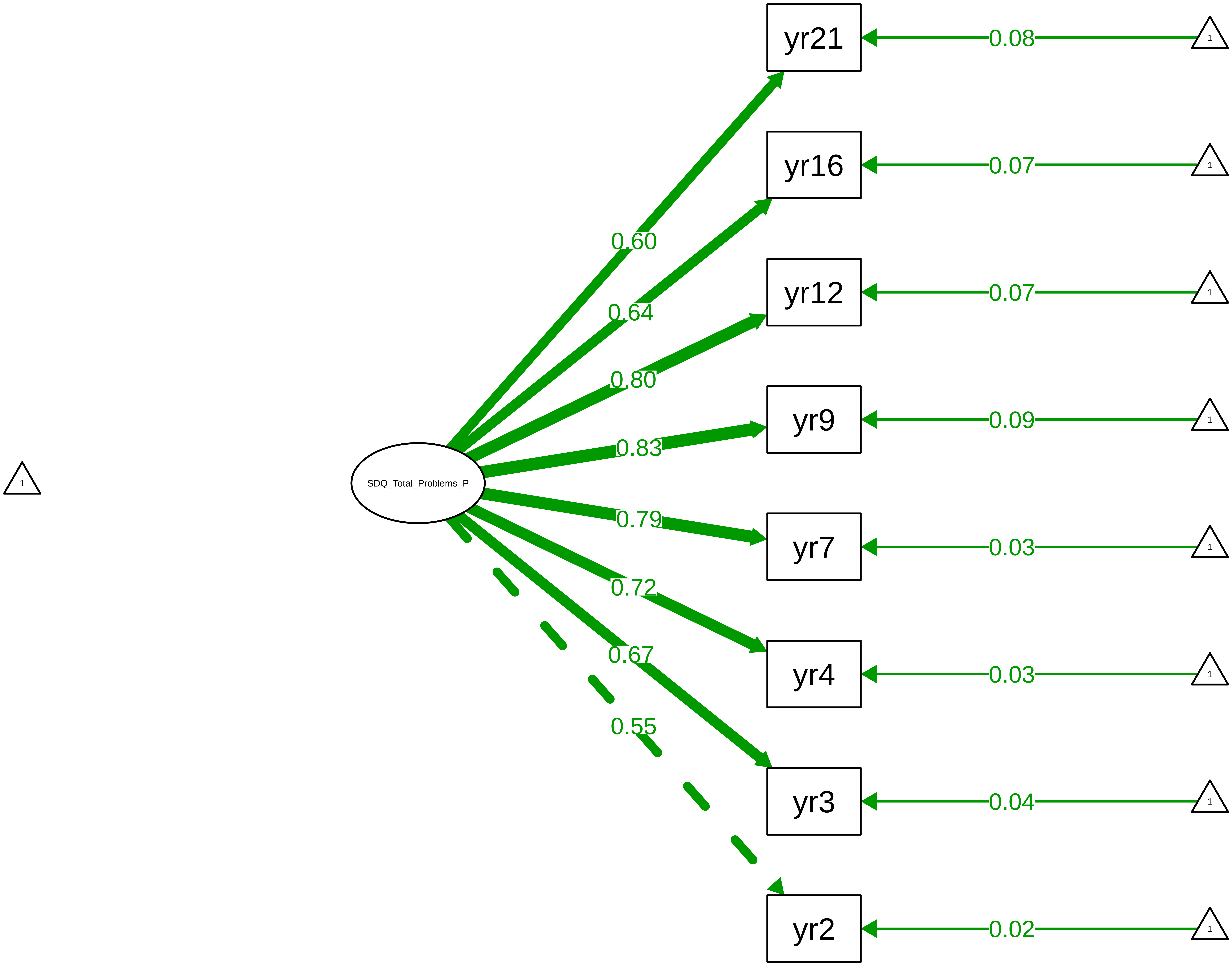

**Figure S8_71: SDQ Total Problems – Parent Stage**

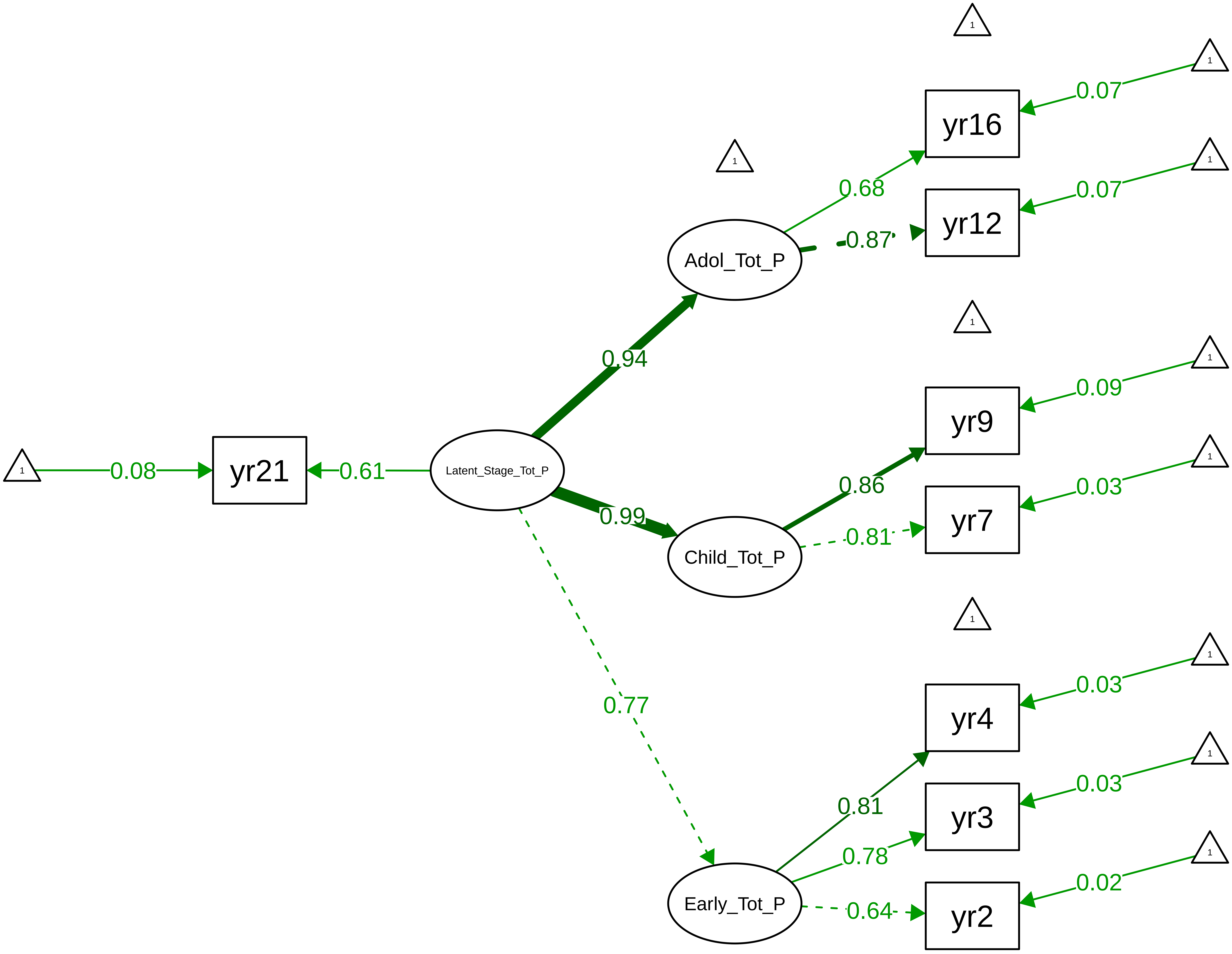

Figure S8_72: SDQ Total Problems − Teacher Overall

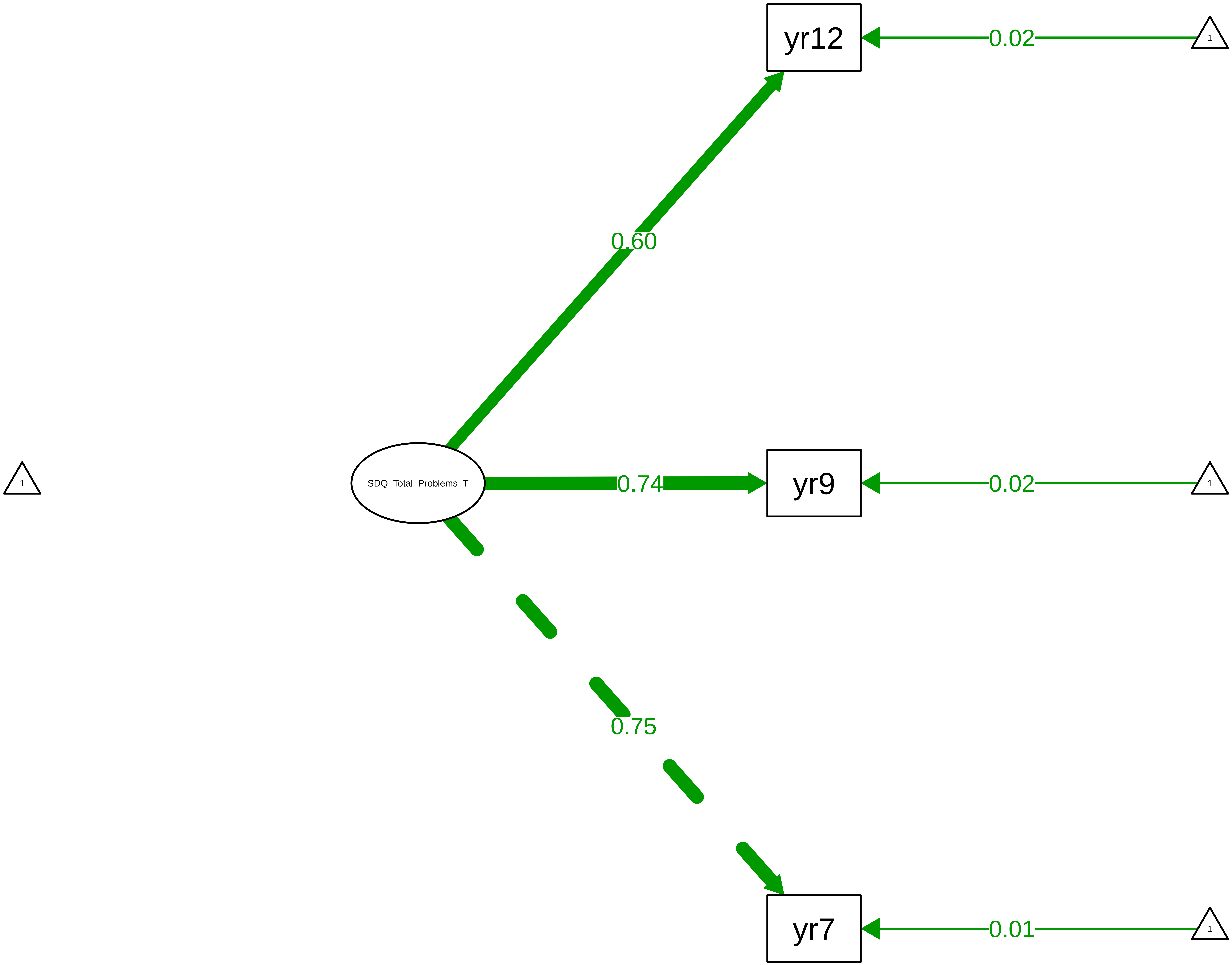

Figure S8_73: SDQ Total Problems − Child Overall

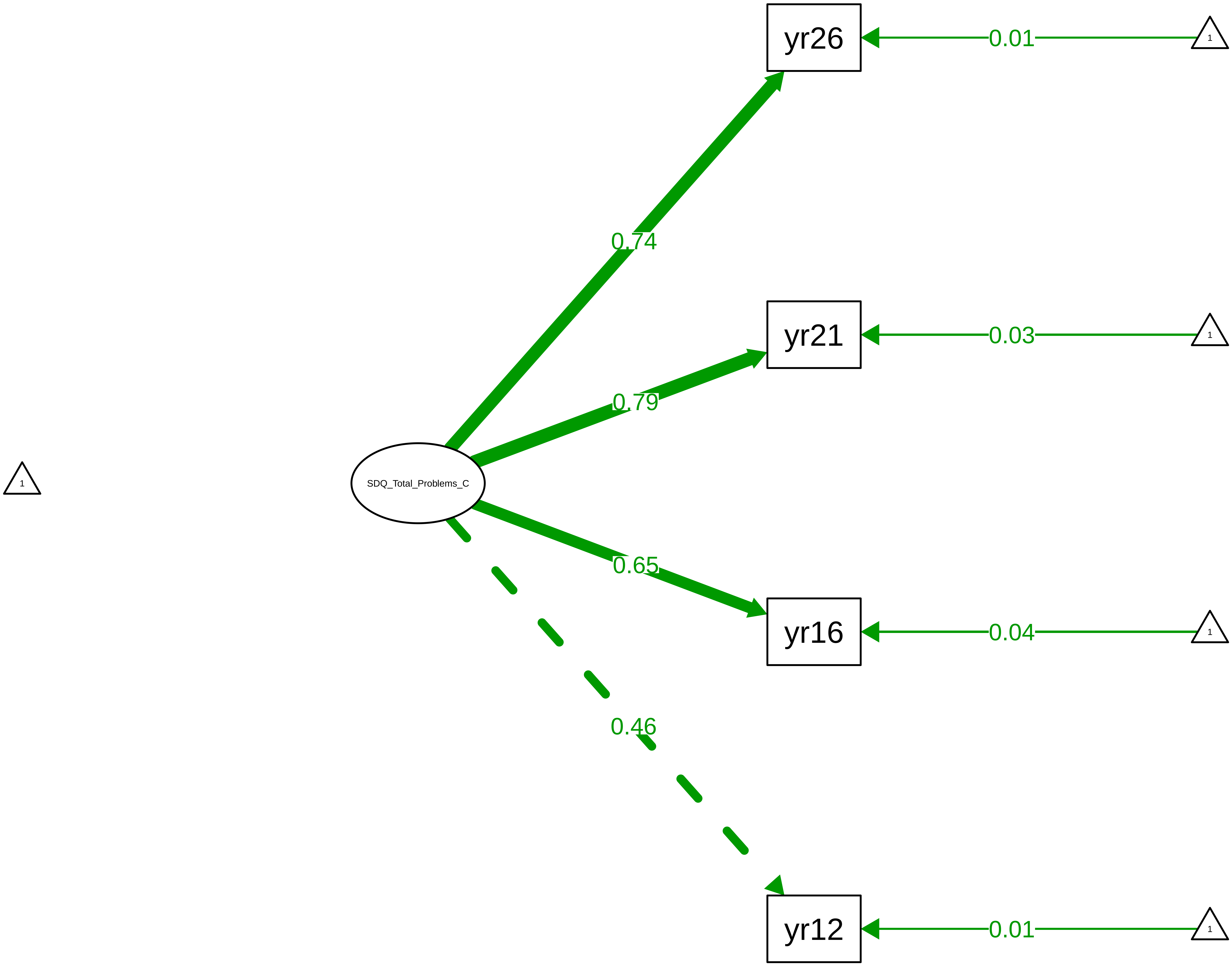

Figure S9_1: General Cognitive Ability (g) − Overall

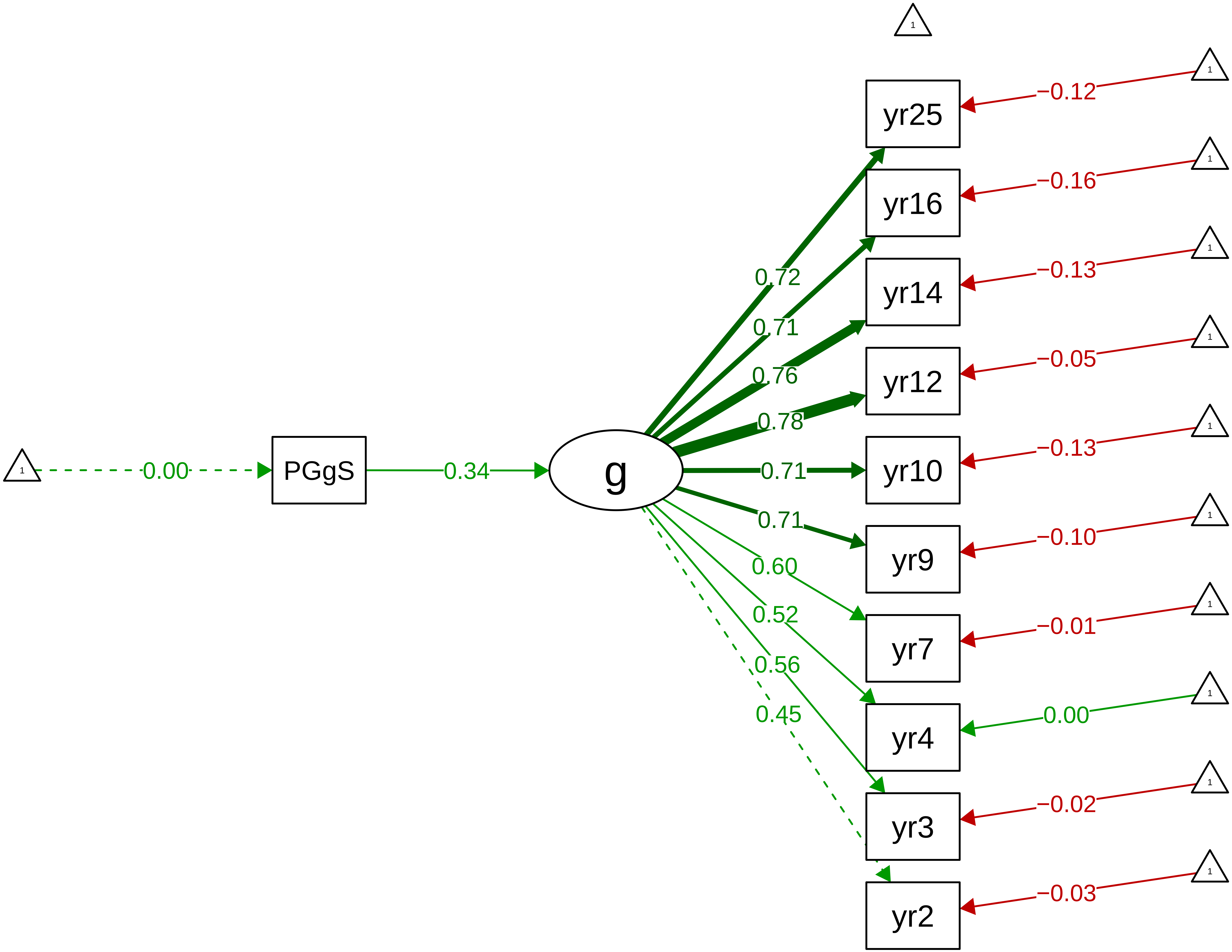

Figure S9_2: General Cognitive Ability (g) − Stage

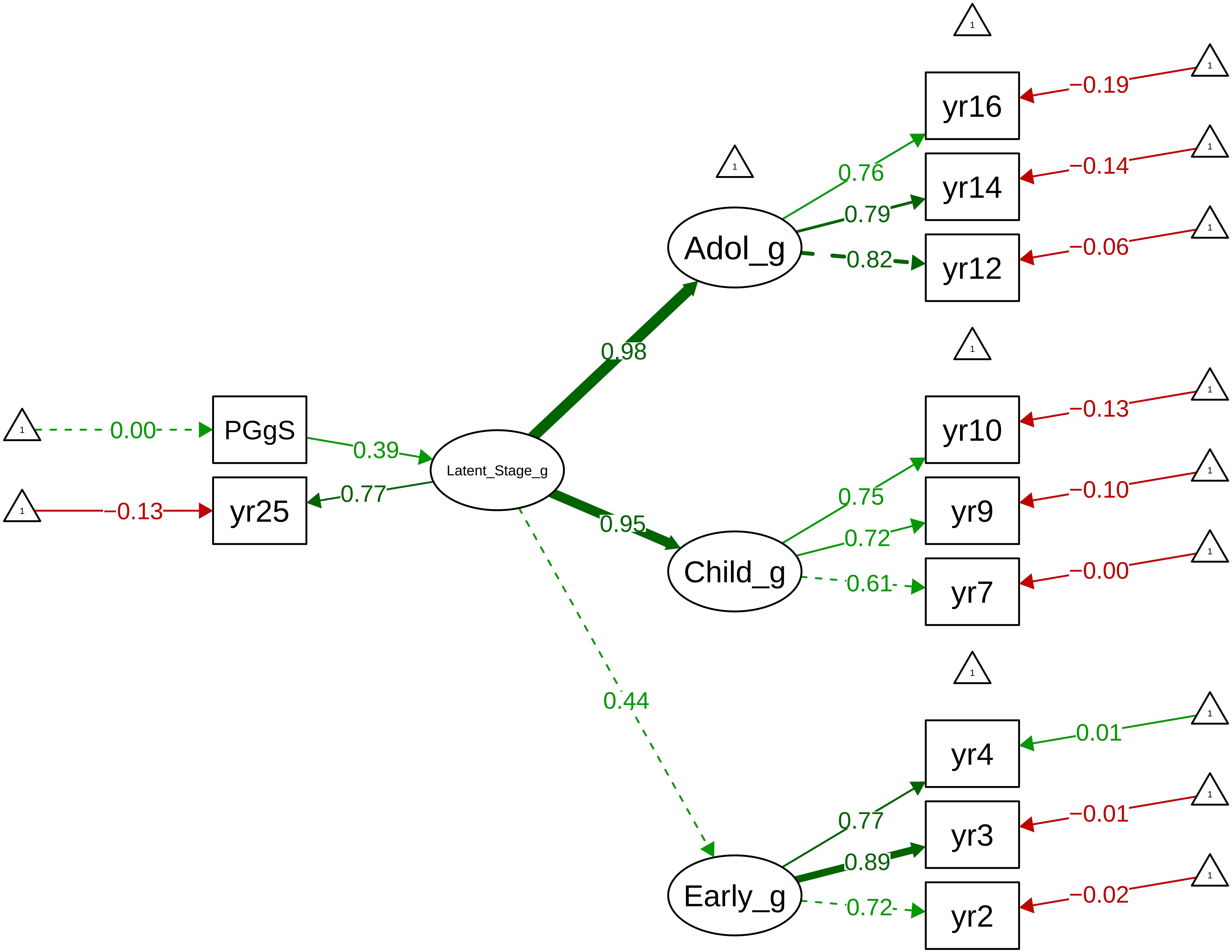

**Figure S9_3: General Cognitive Ability (g) − Method**

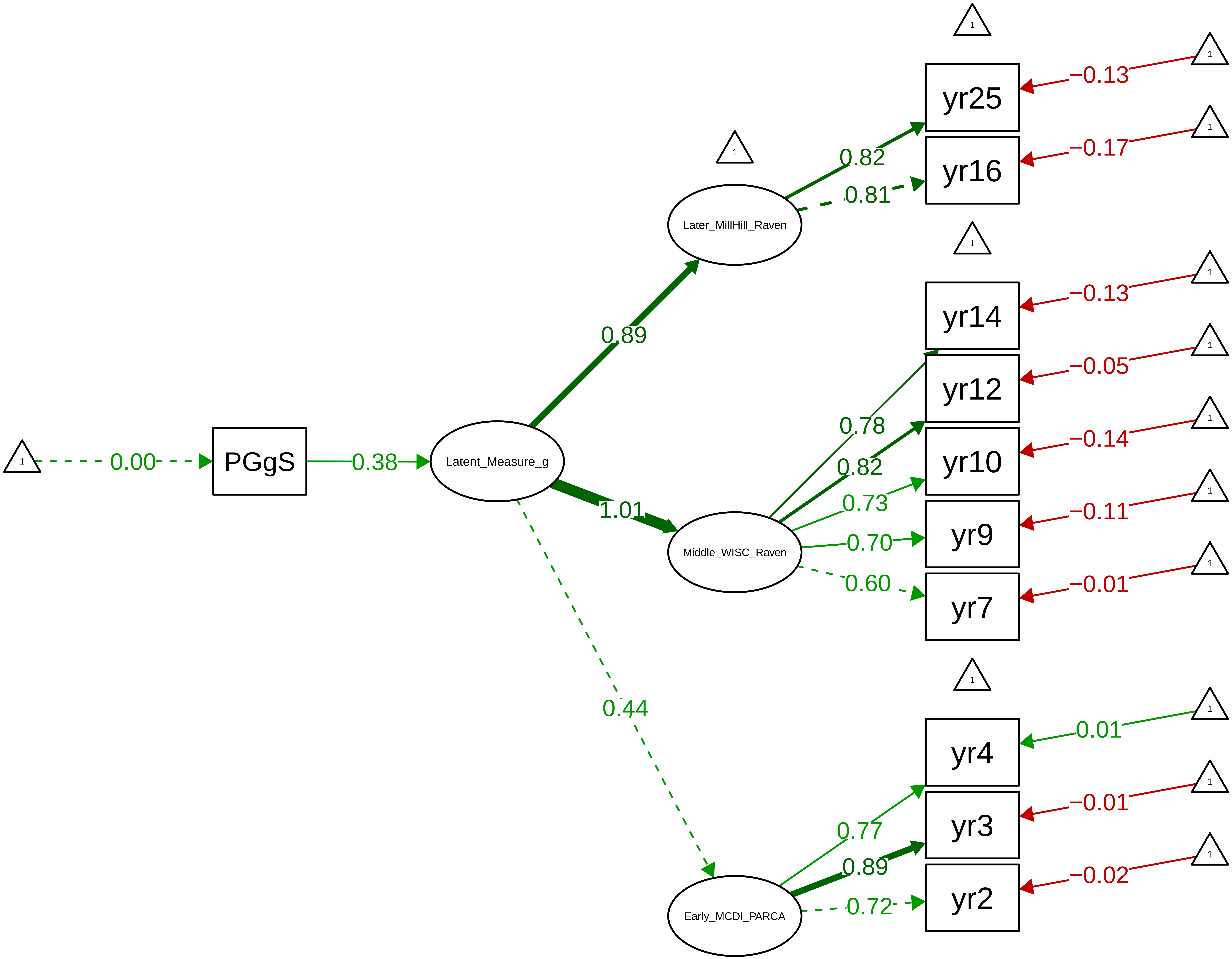

Figure S9_4: Verbal Ability − Overall

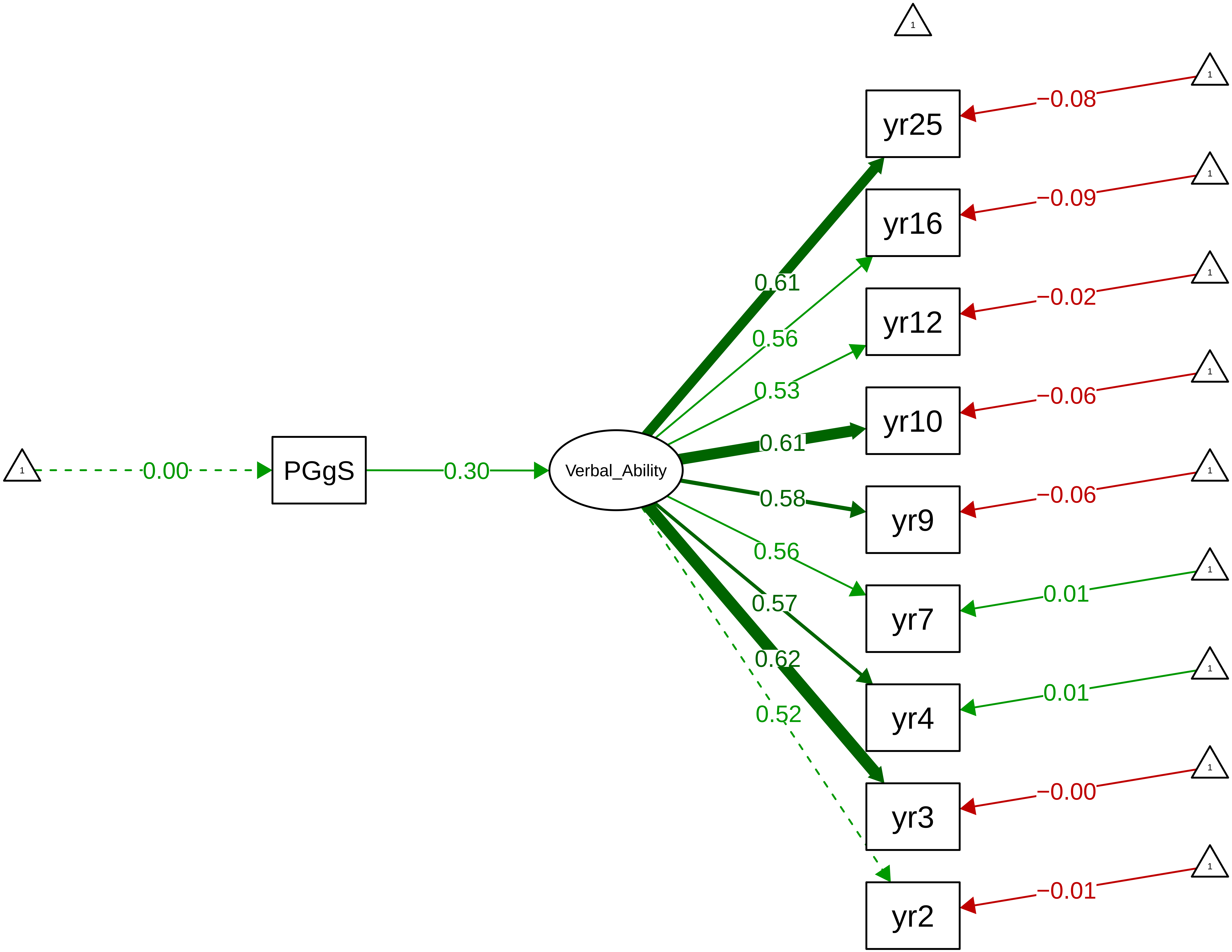

Figure S9_5: Verbal Ability − Stage

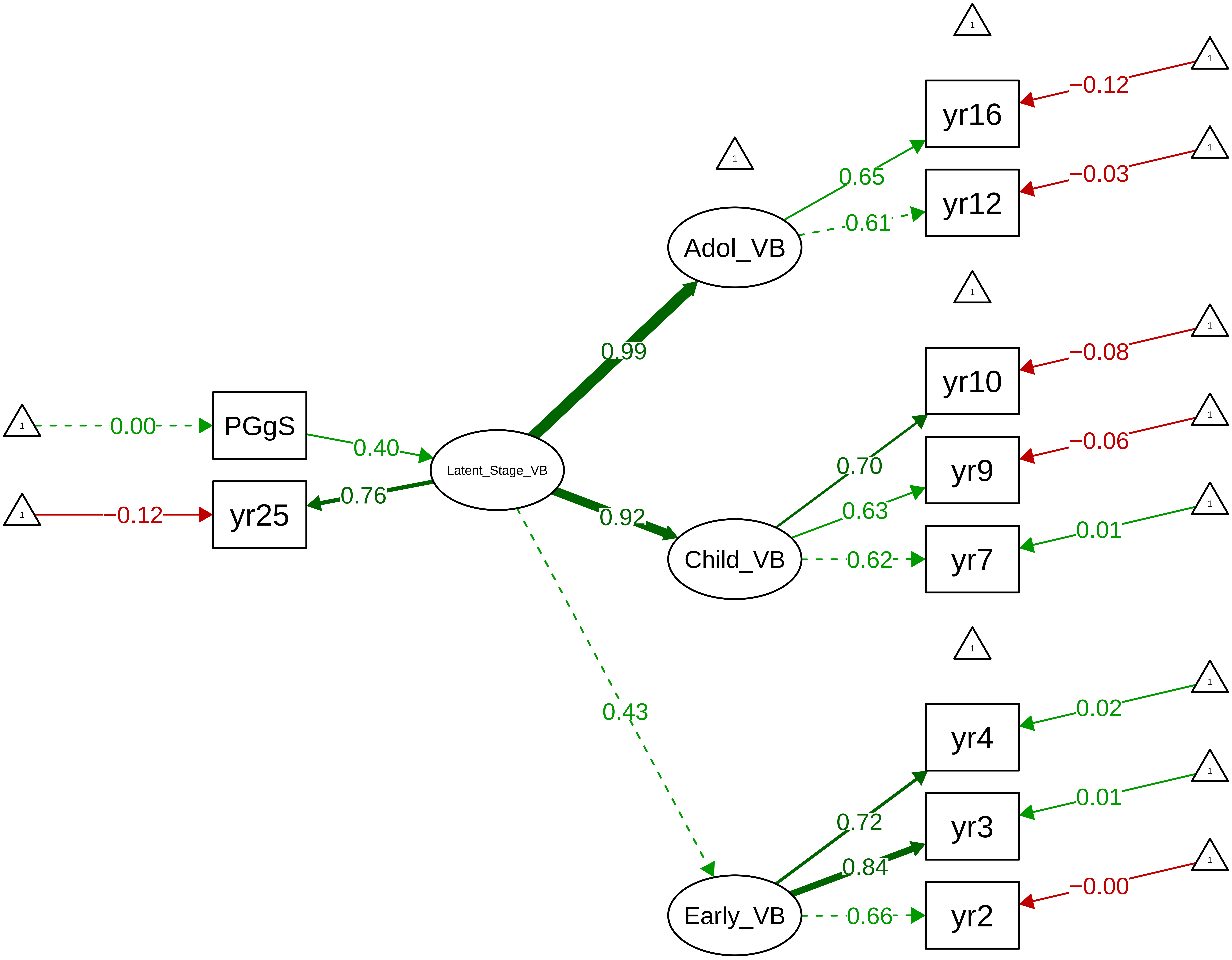

**Figure S9_6: Verbal Ability − Method**

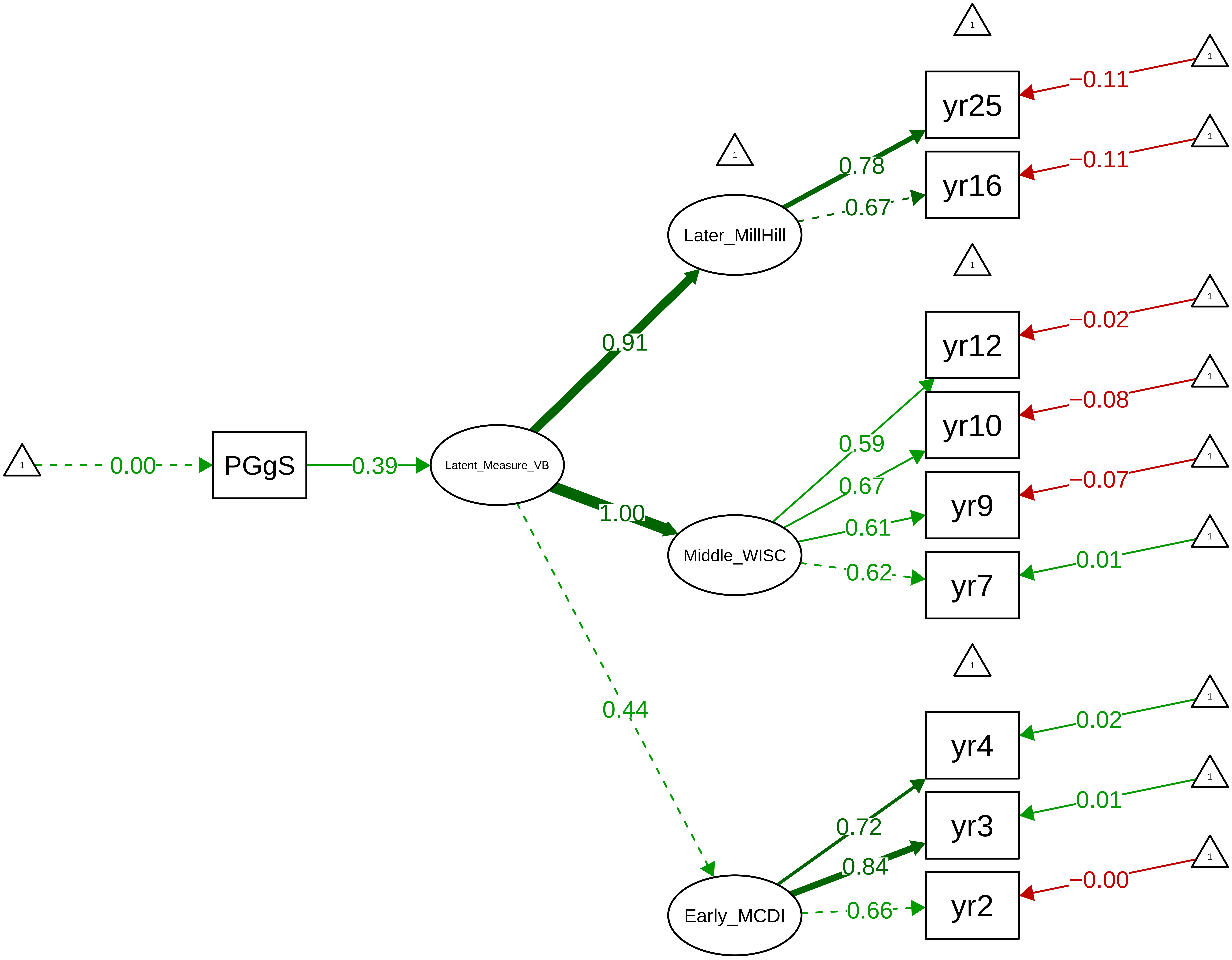

**Figure S9_7: Nonverbal Ability − Overall**

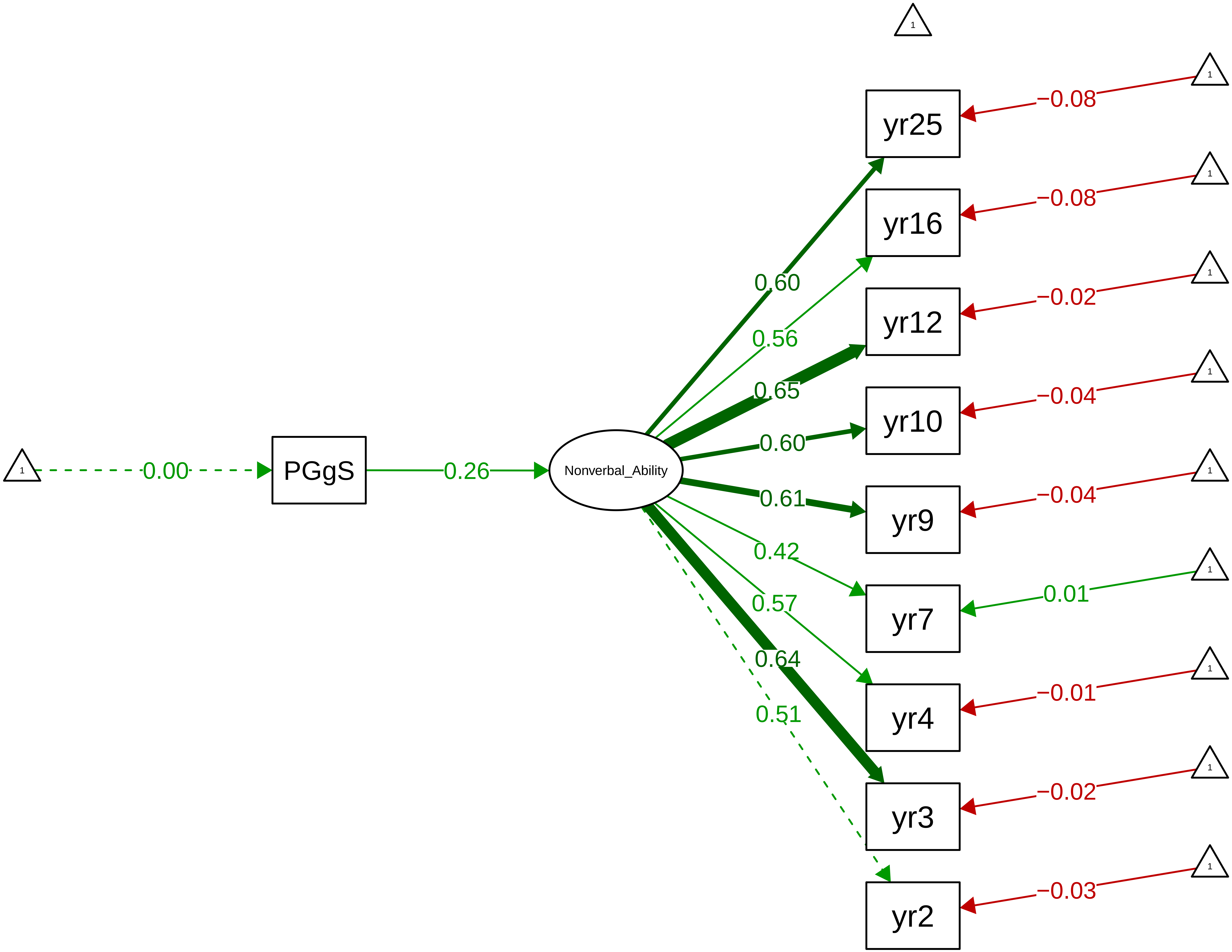

**Figure S9_8: Nonverbal Ability − Stage**

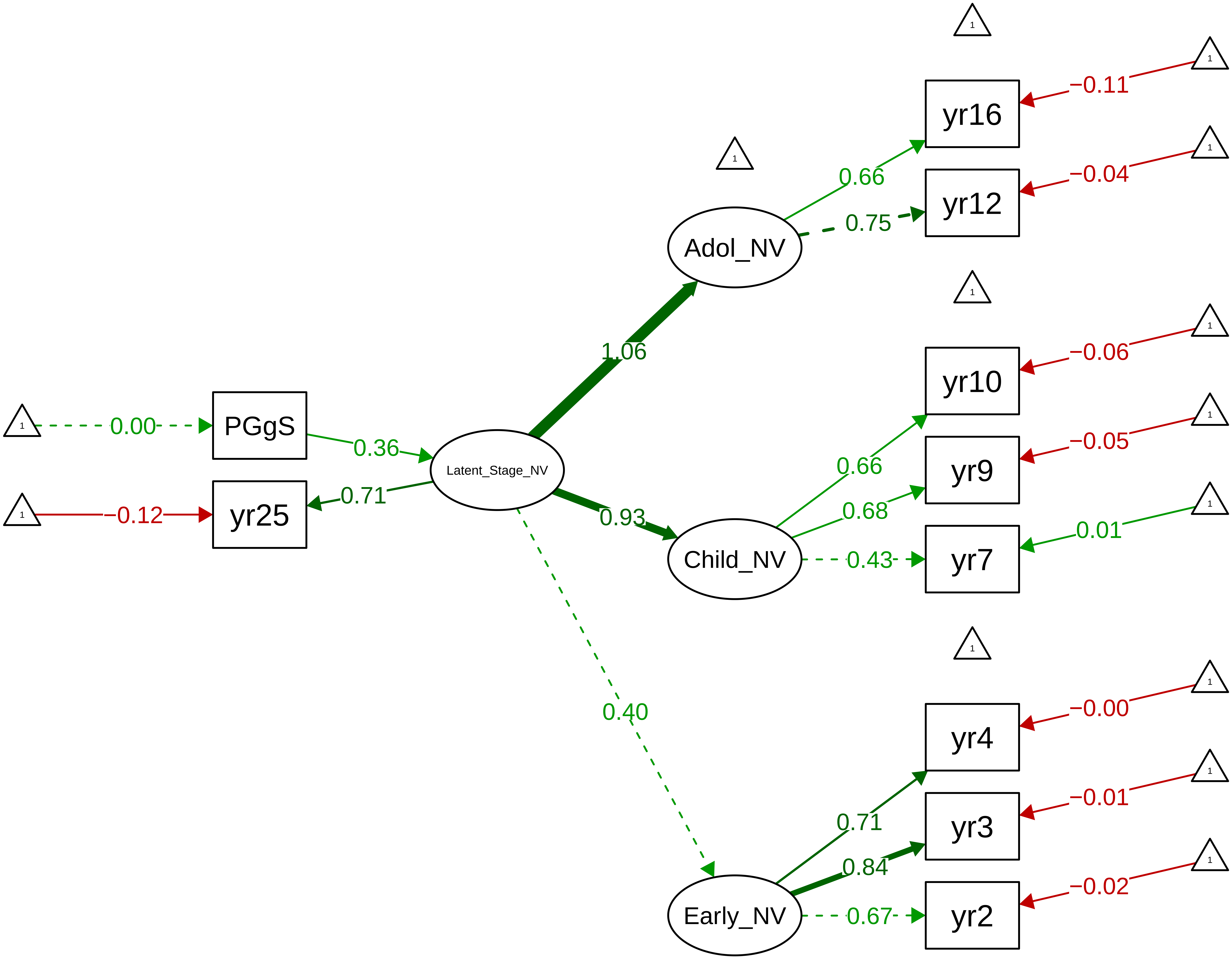

**Figure S9_9: Nonverbal Ability − Method**

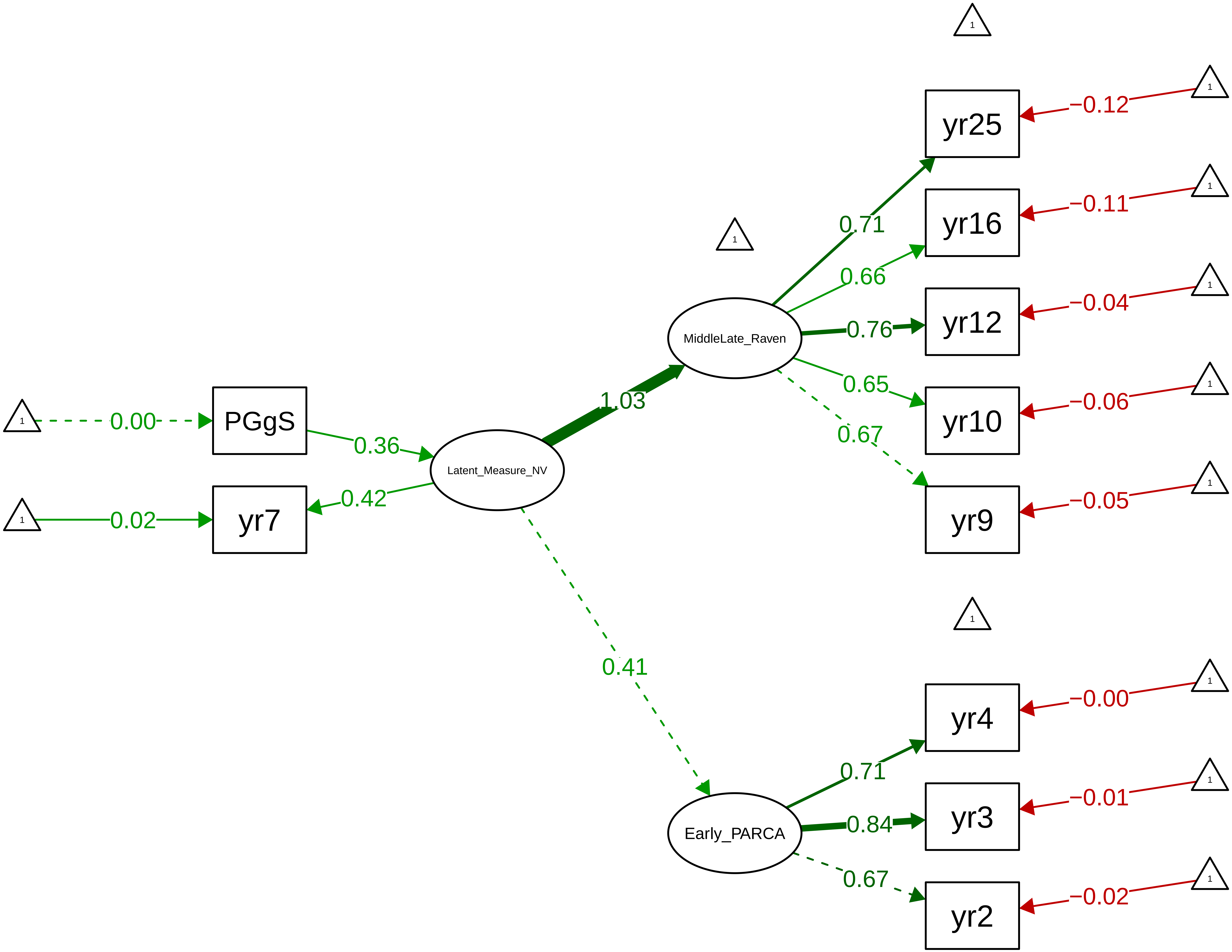

Figure S9_10: English Achievement Latent (Teacher 7−12)

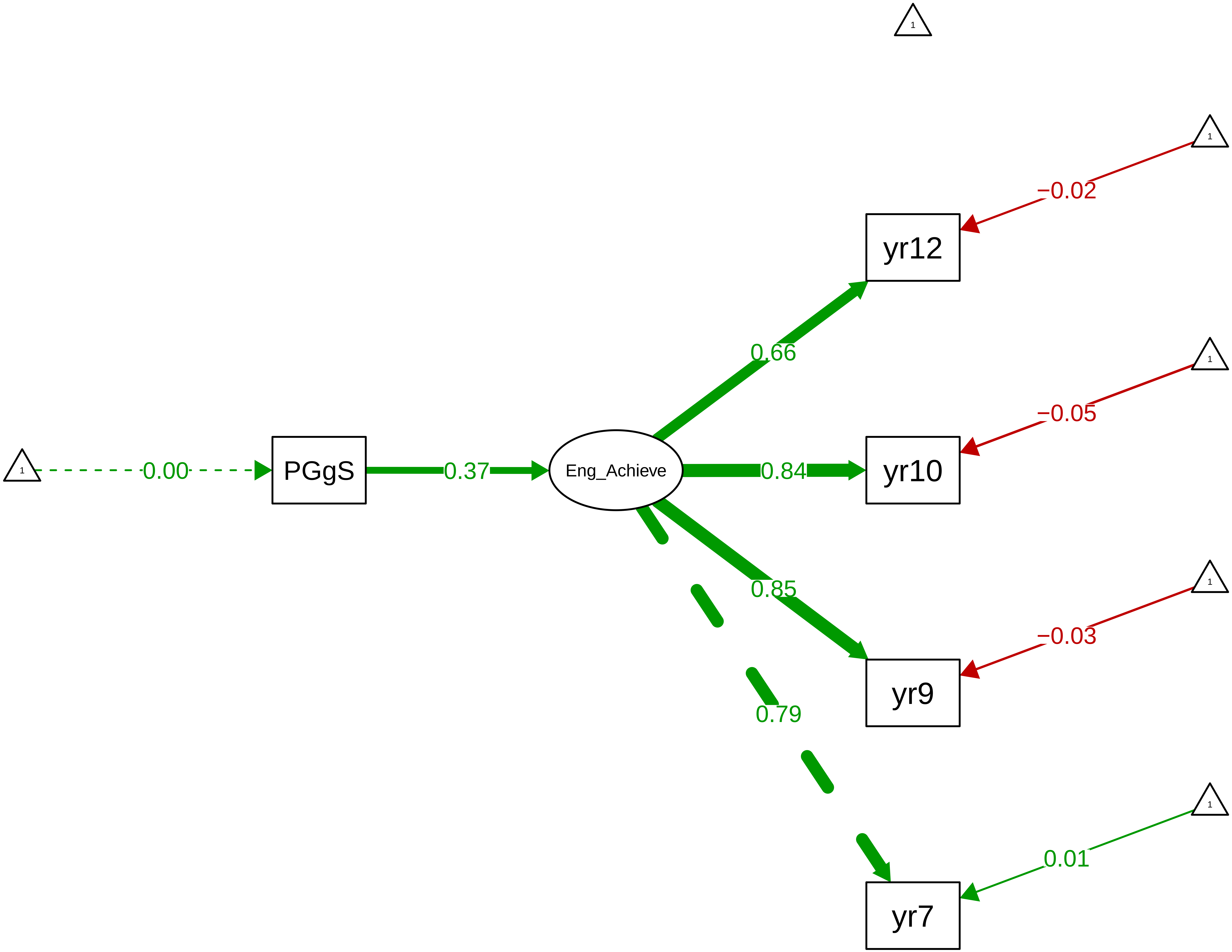

**Figure S9_11: Maths Achievement Latent (Teacher 7−12)**

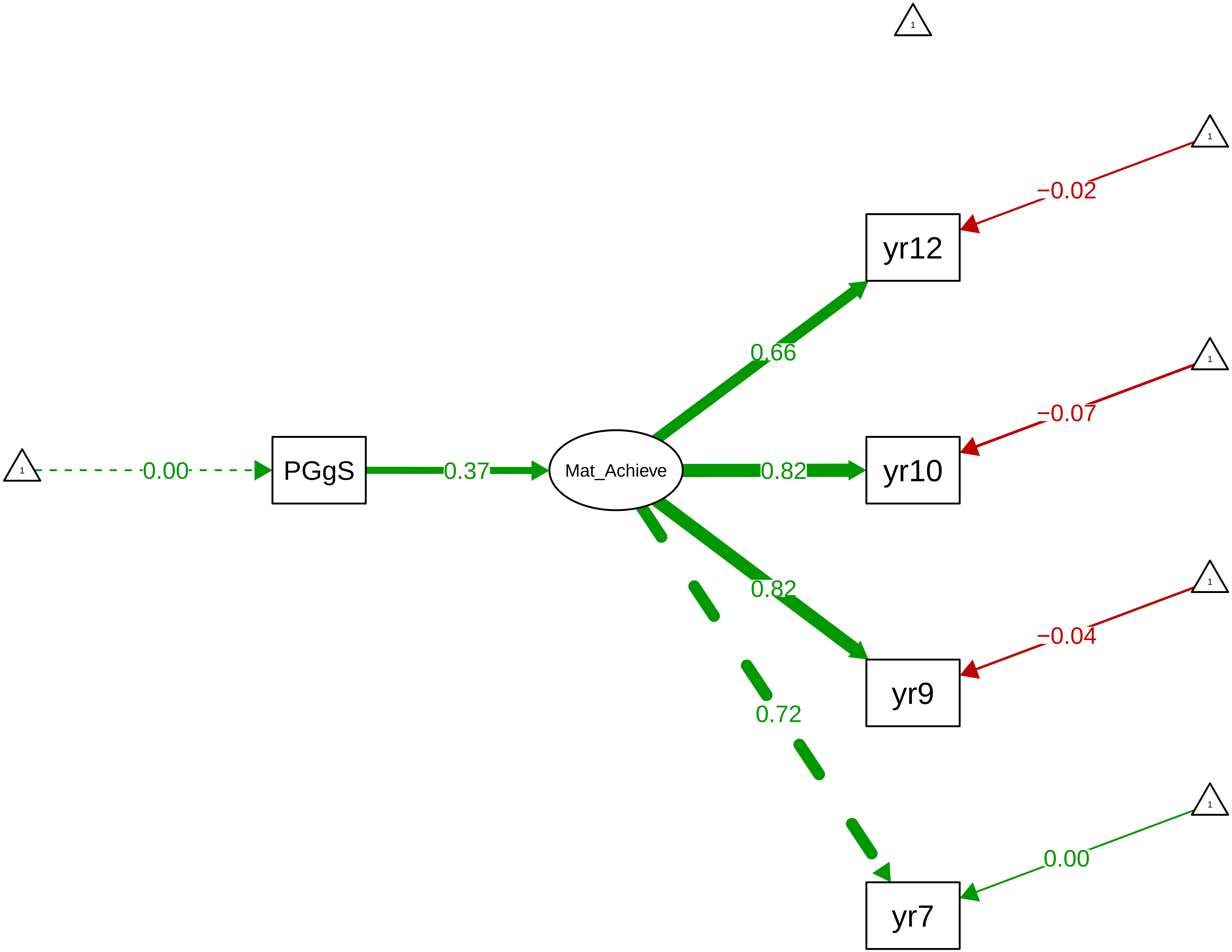

**Figure S9_12: Science Achievement Latent (Teacher 9–12)**

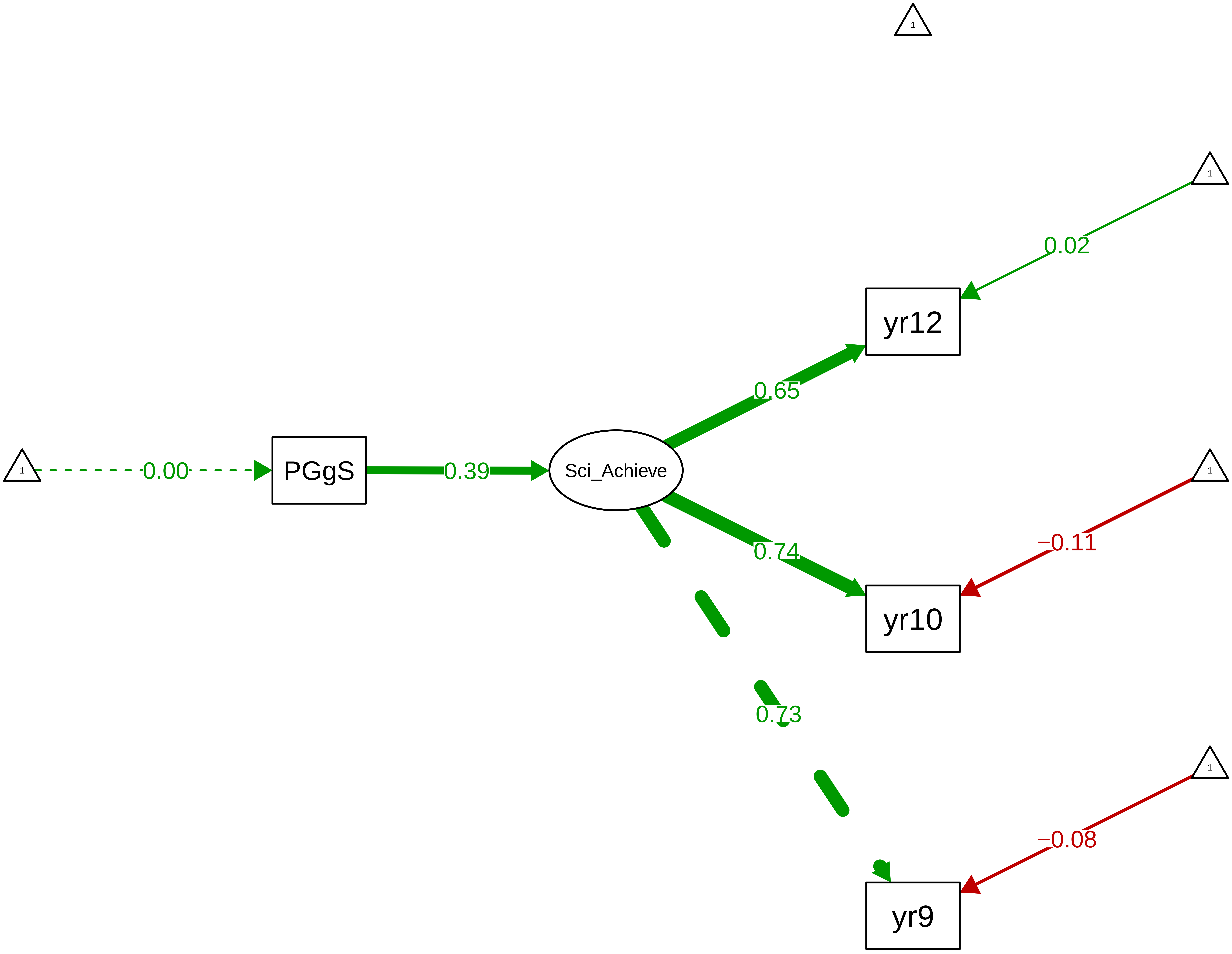

**Figure S9_13: Core−Subject Achievement Latent (Teacher 7−12)**

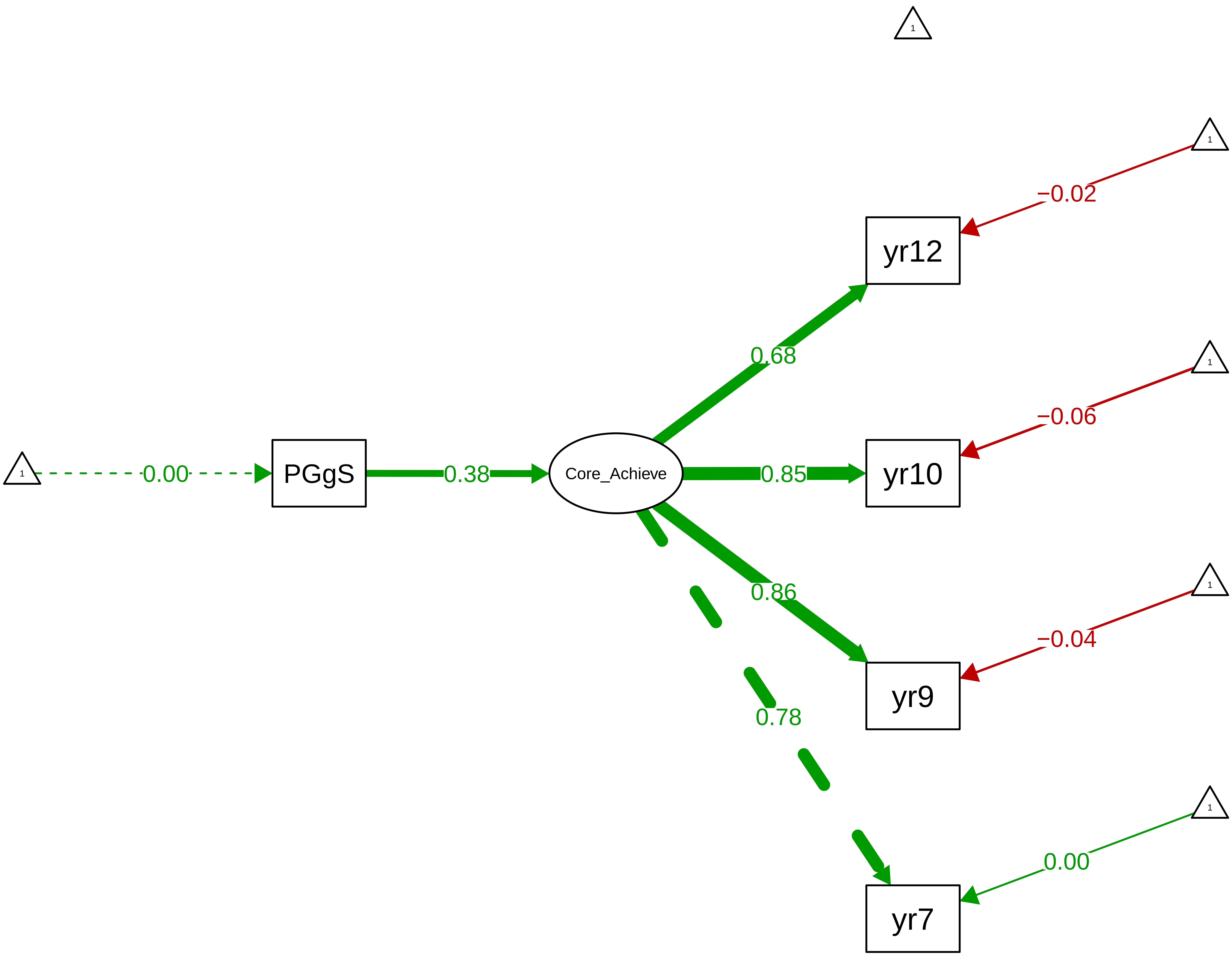

Figure S9_14: ARBQ Shyness − CTCR Overall

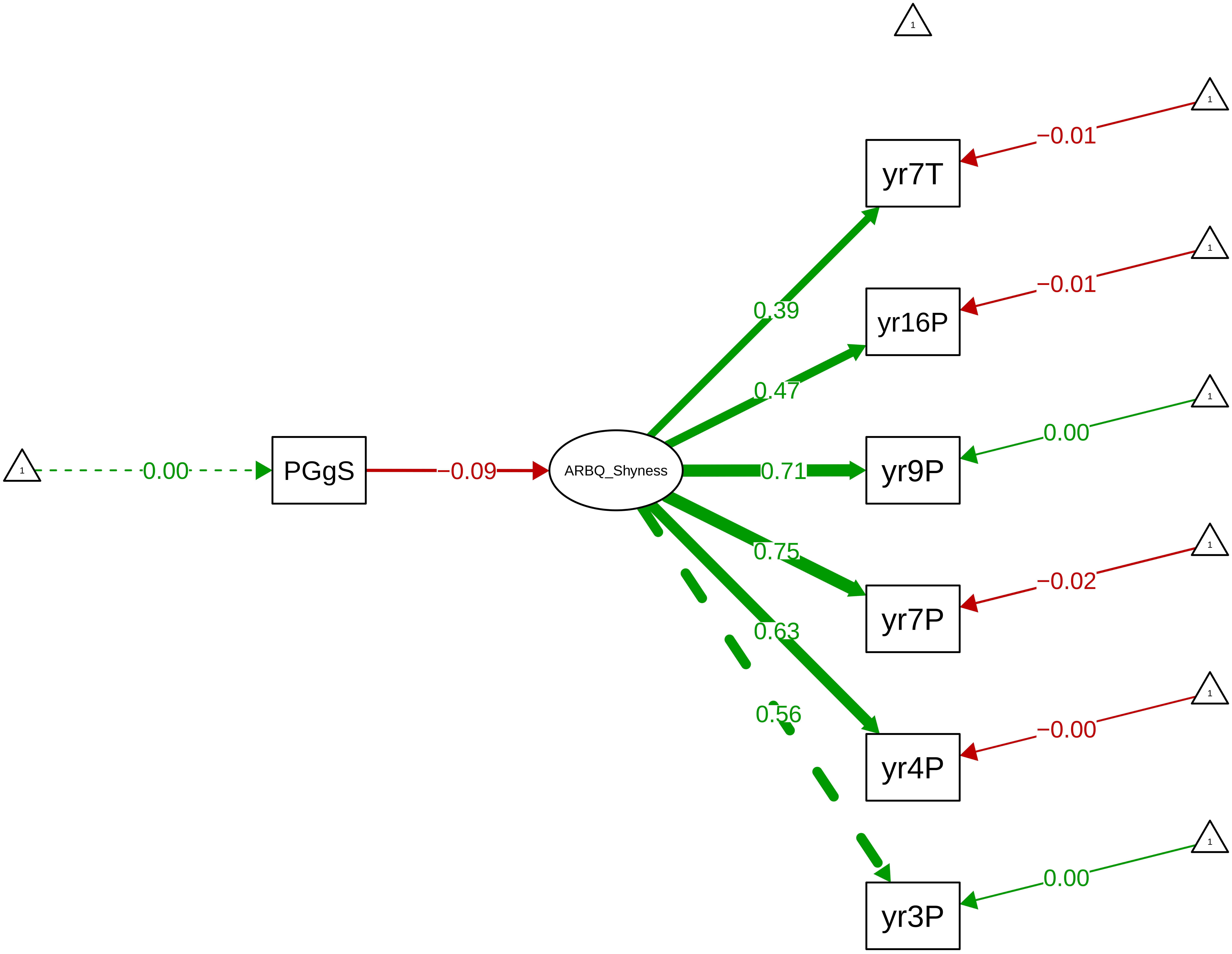

Figure S9_15: ARBQ Shyness − CTCR Stage

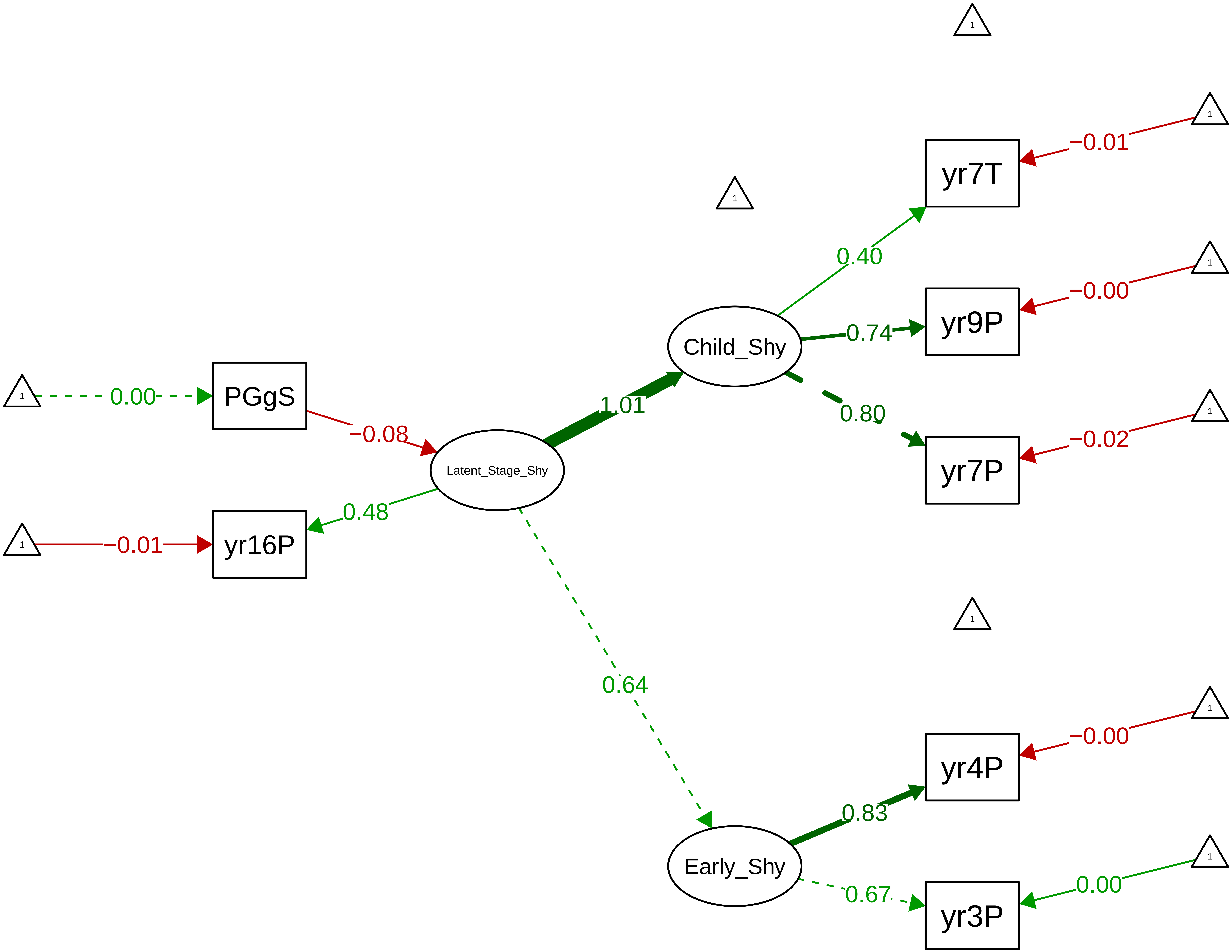

Figure S9_16: ARBQ Fear − CTCR Overall

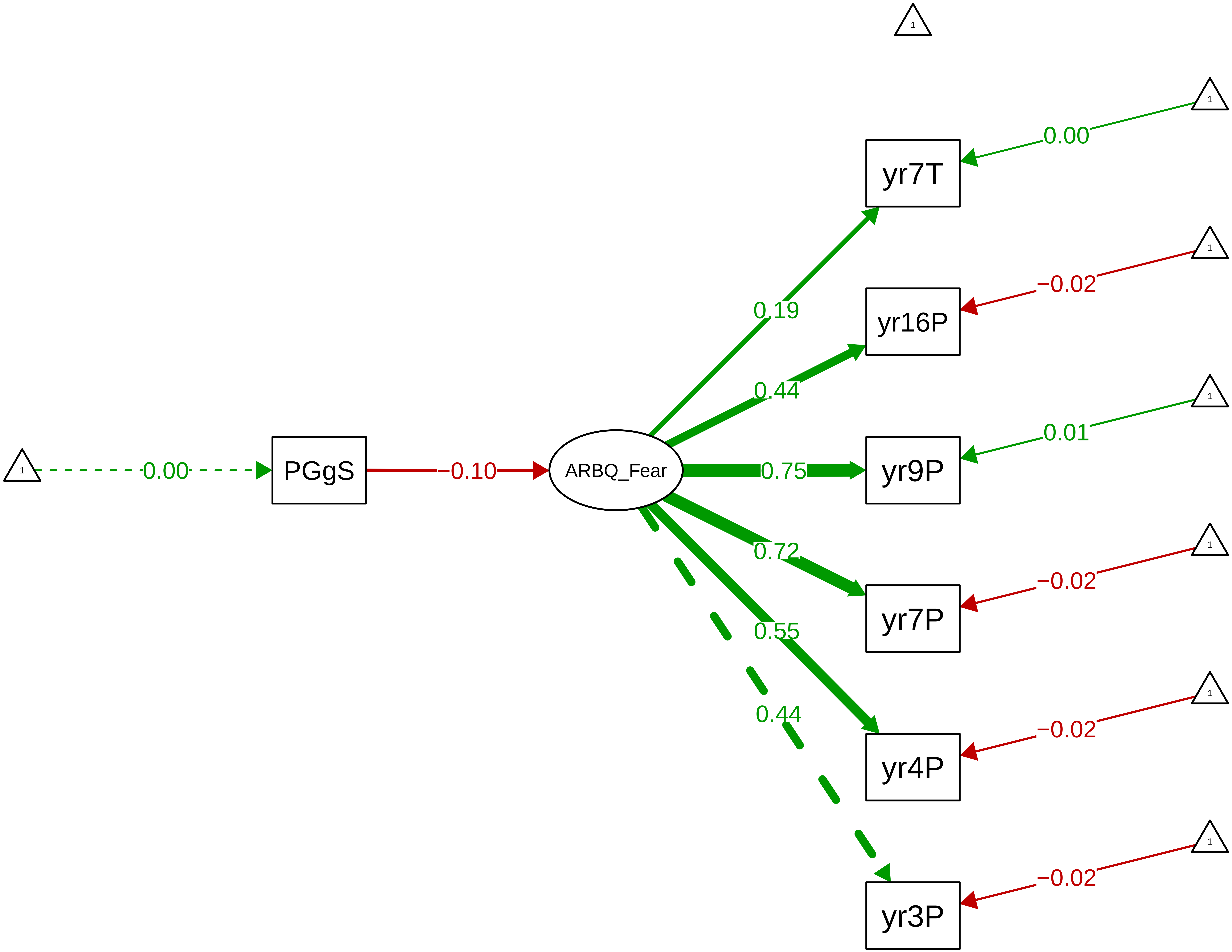

**Figure S9_17: ARBQ Fear – CTCR Stage**

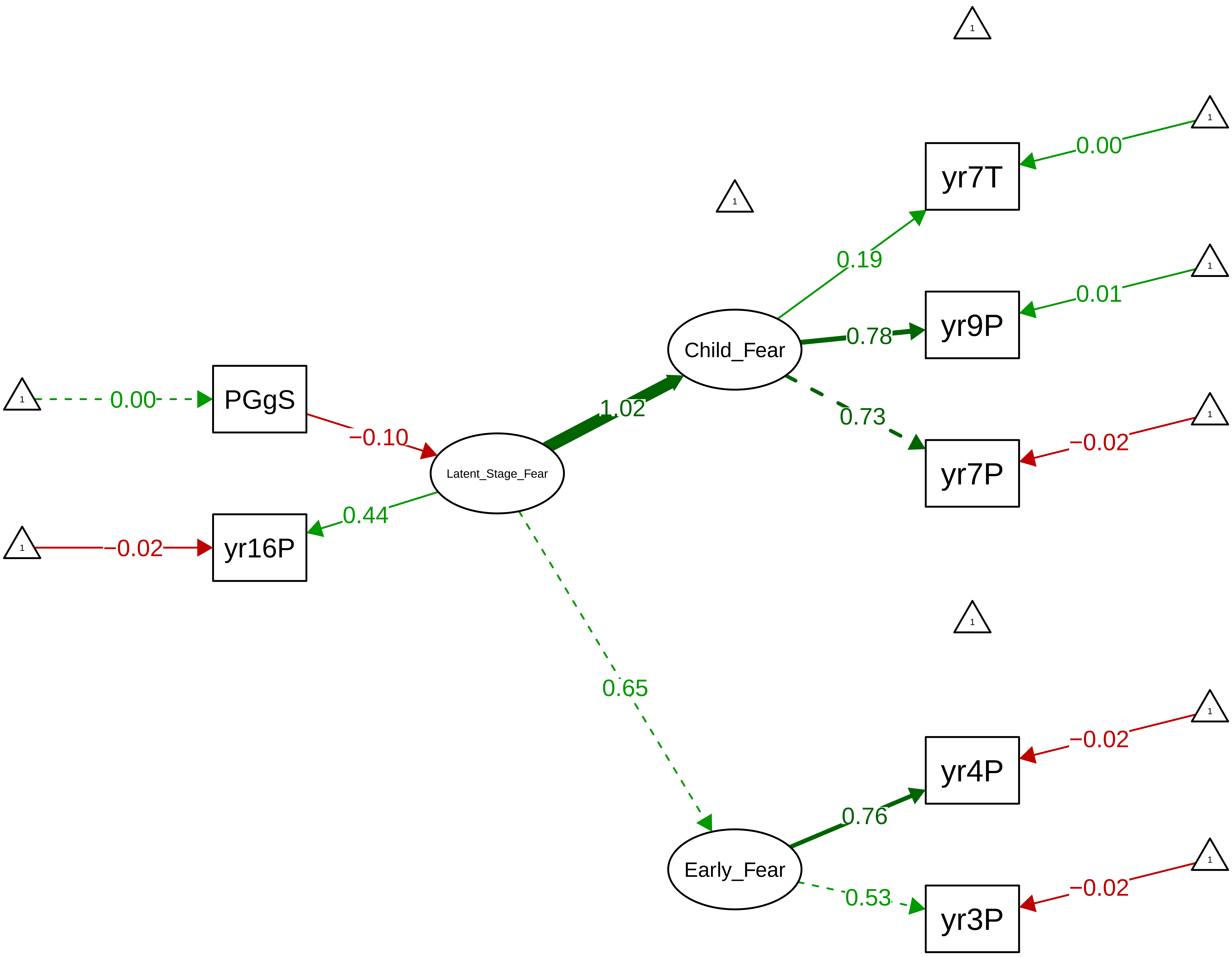

Figure S9_18: ARBQ Obsessive−Compulsive − CTCR Overall

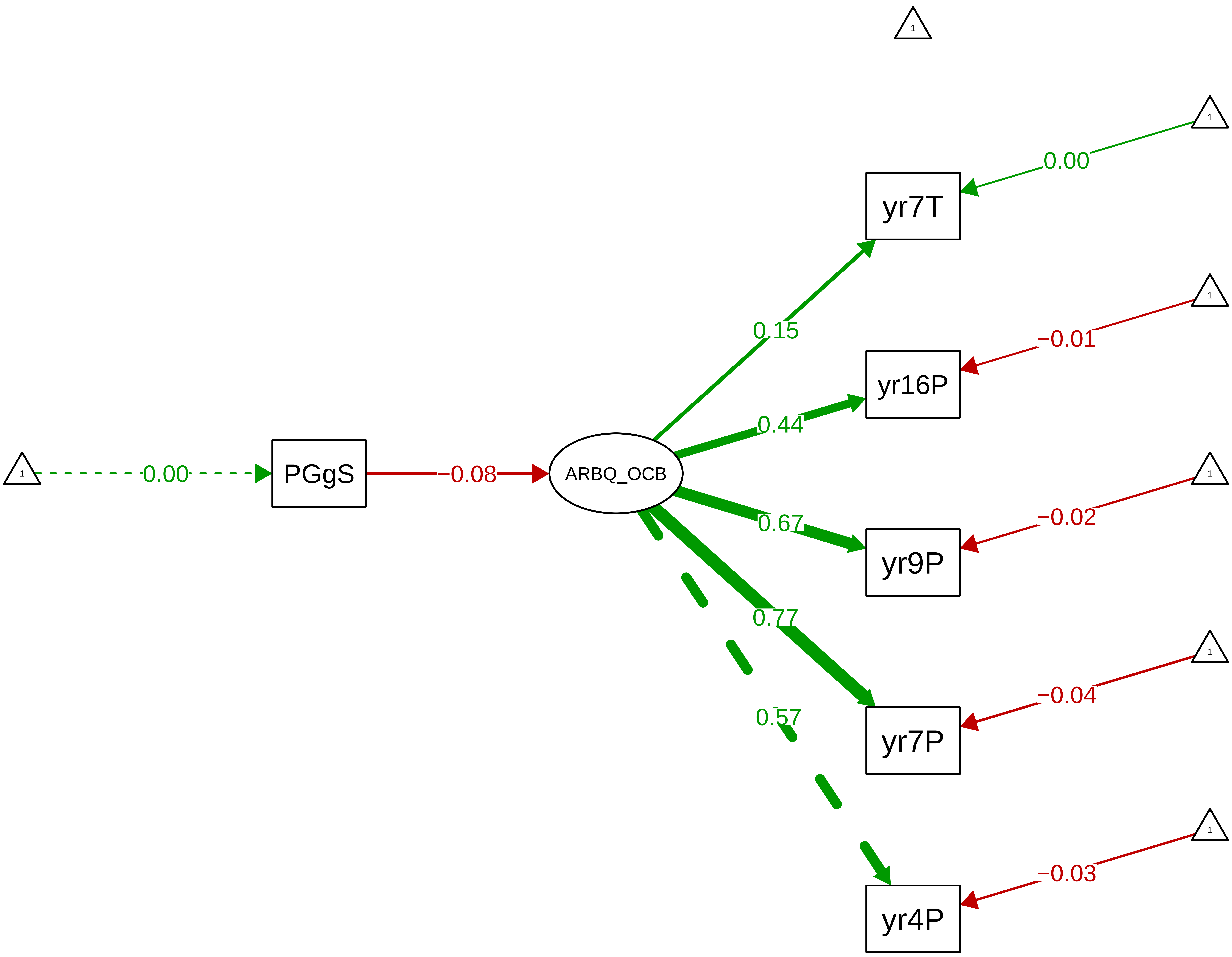

#### Figure S9_19: ARBQ Obsessive−Compulsive − CTCR Stage

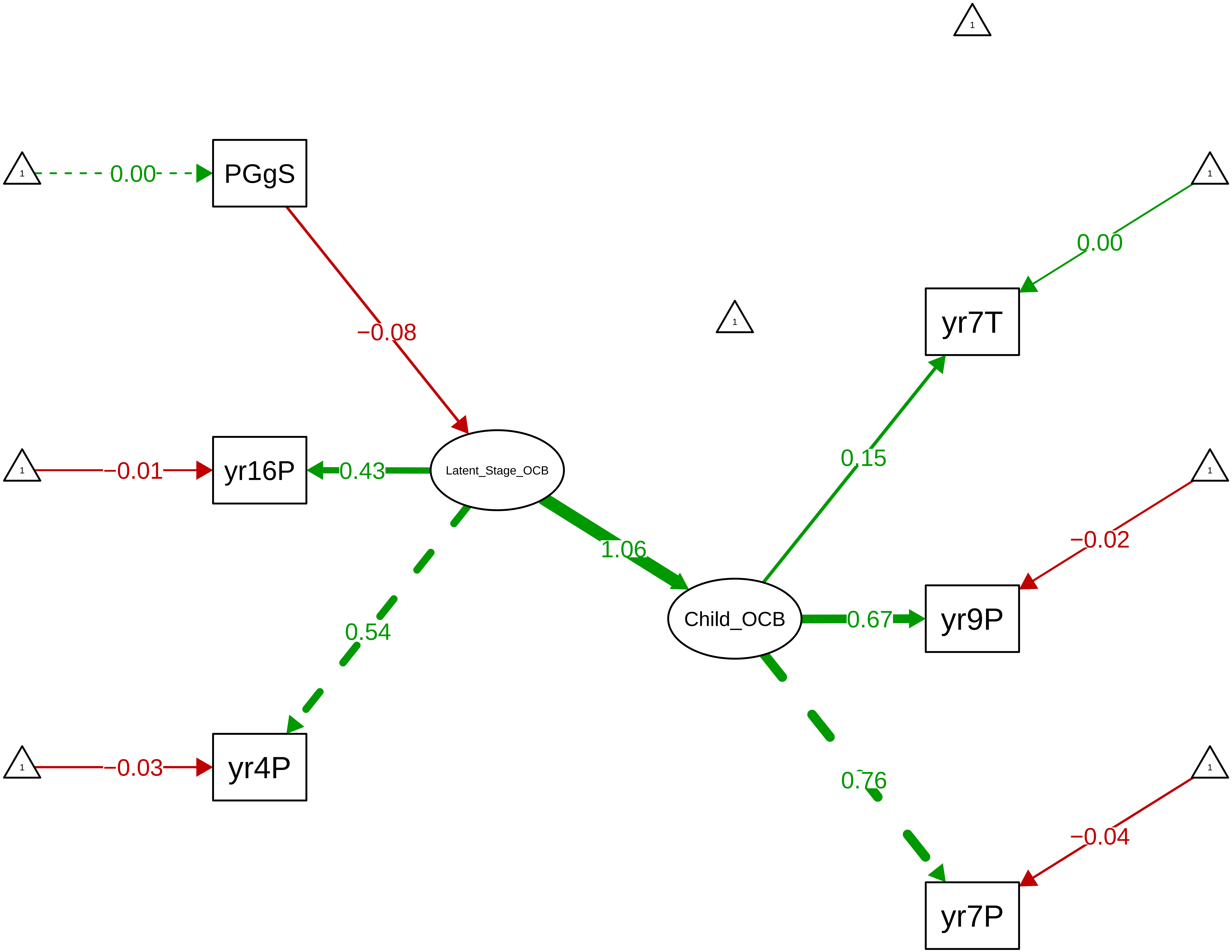

Figure S9_20: ARBQ Negative Affect − CTCR Overall

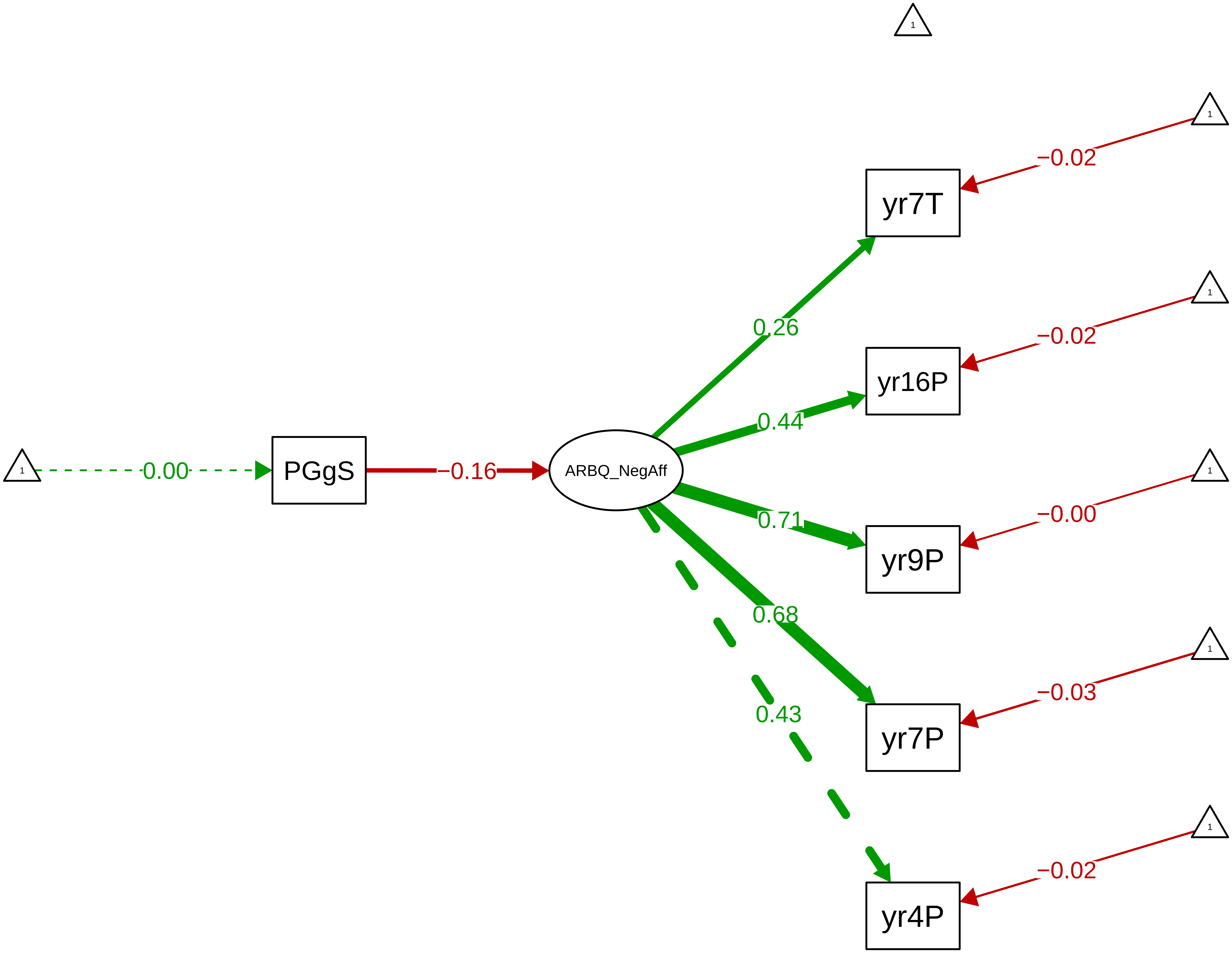

**Figure S9_21: ARBQ Negative Affect − CTCR Stage**

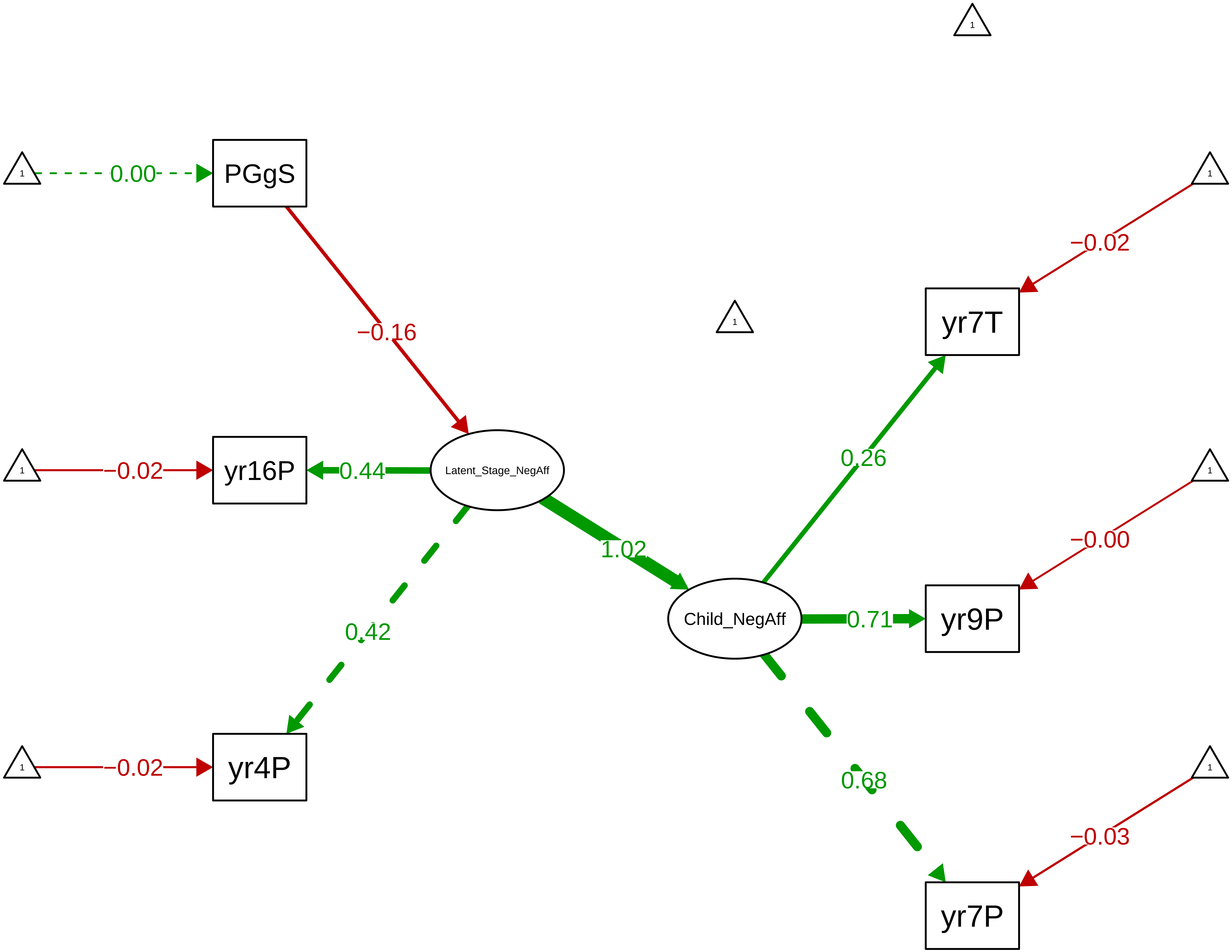

Figure S9_22: ARBQ Negative Cognition − CTCR Overall

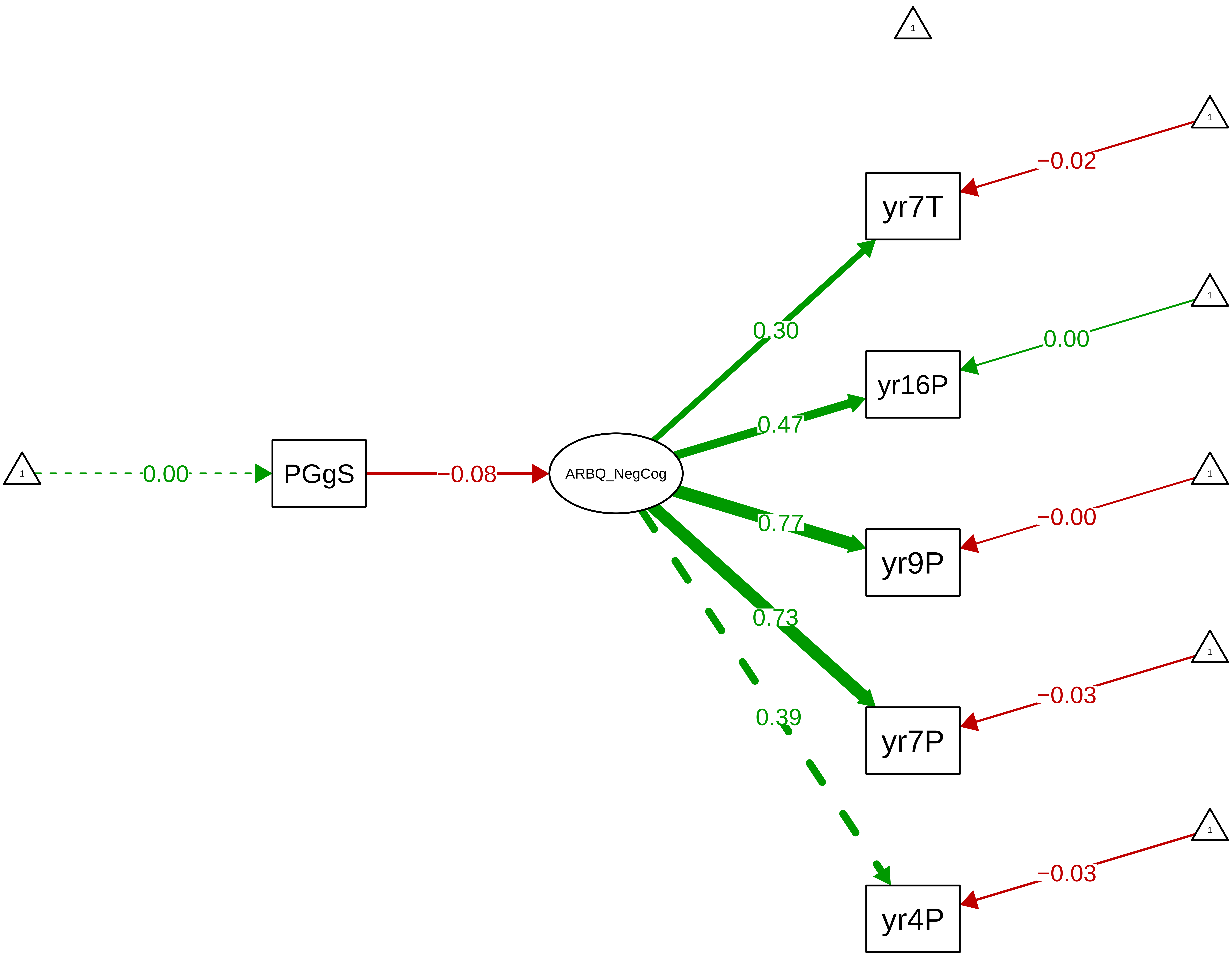

**Figure S9_23: ARBQ Negative Cognition − CTCR Stage**

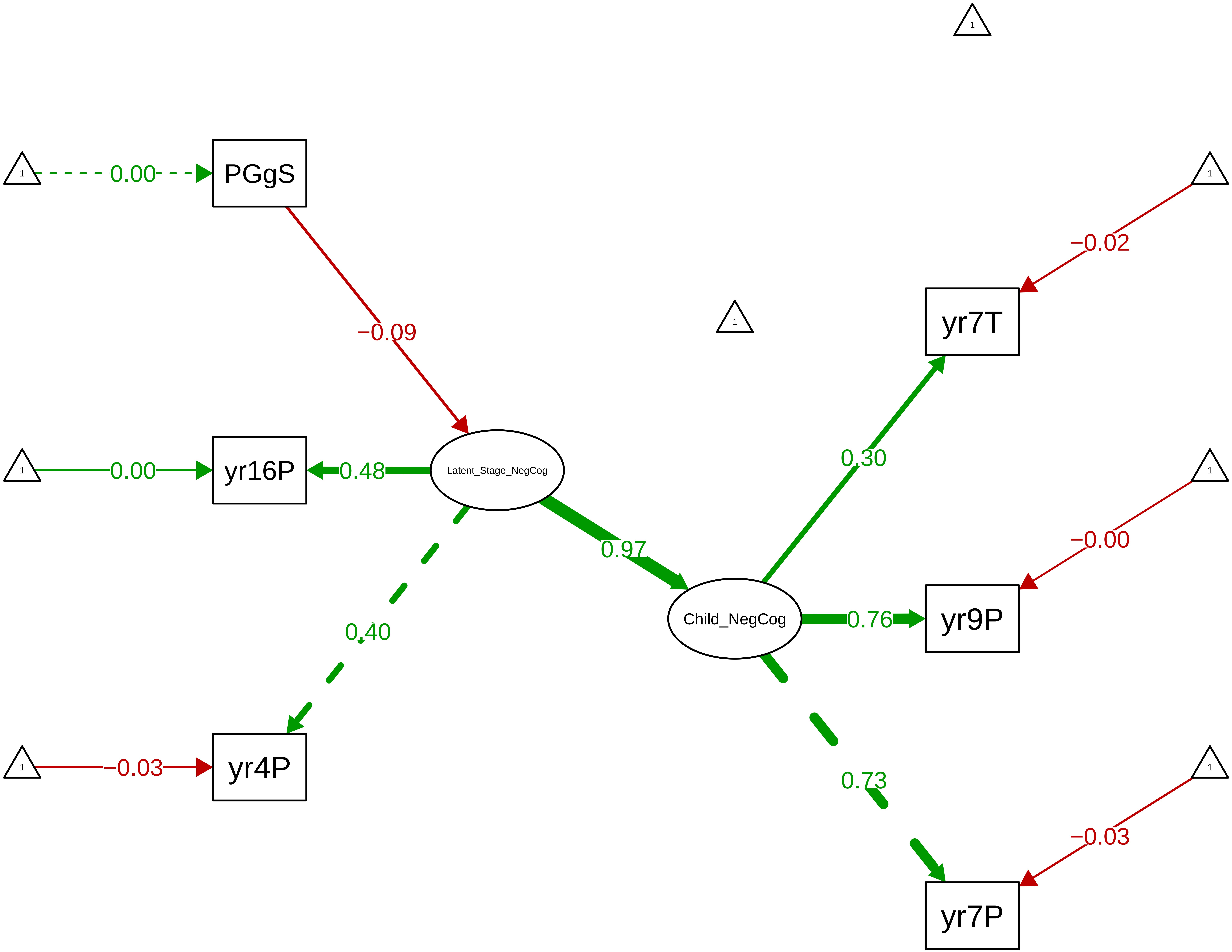

Figure S9_24: ARBQ Anxiety Total – CTCR Overall

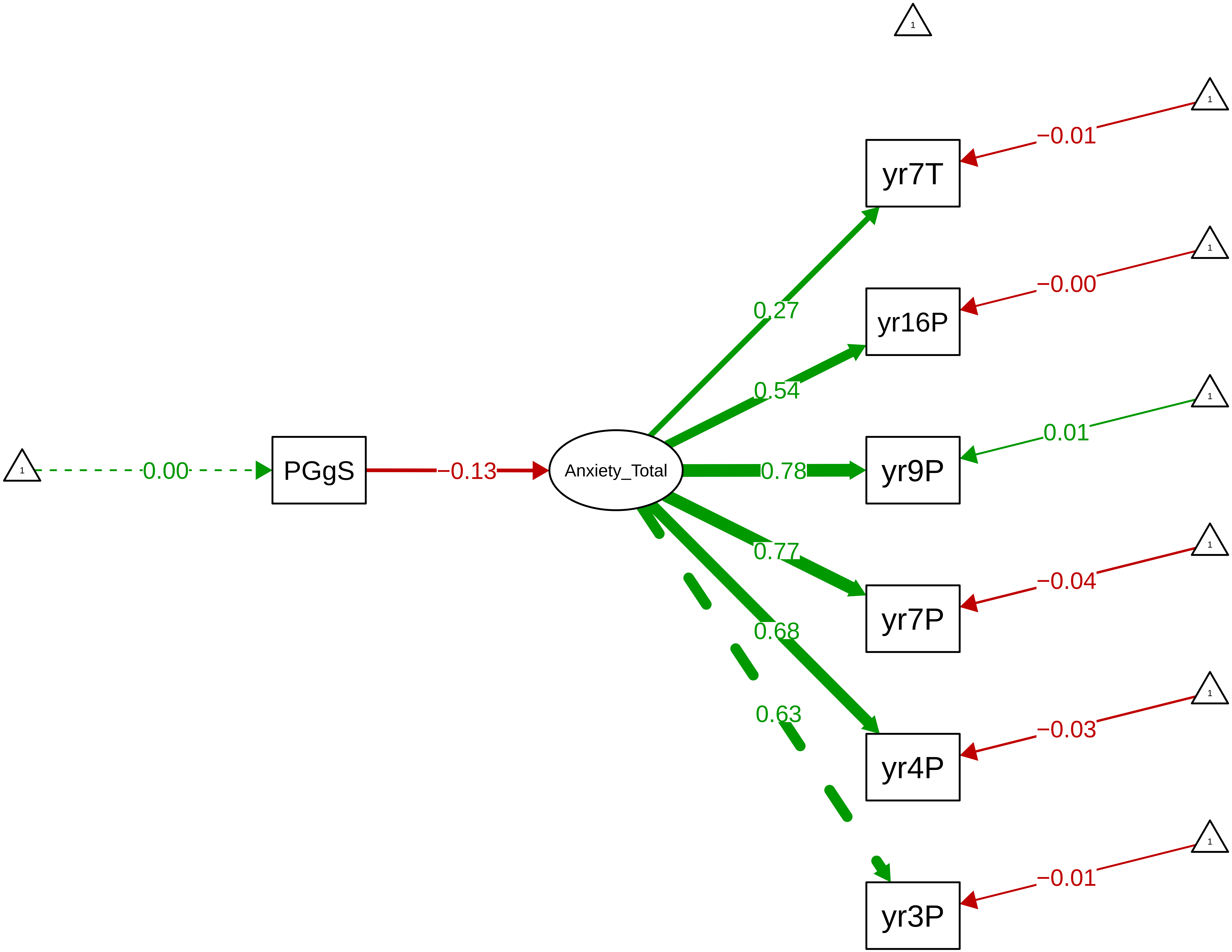

Figure S9_25: ARBQ Anxiety Total − CTCR Stage

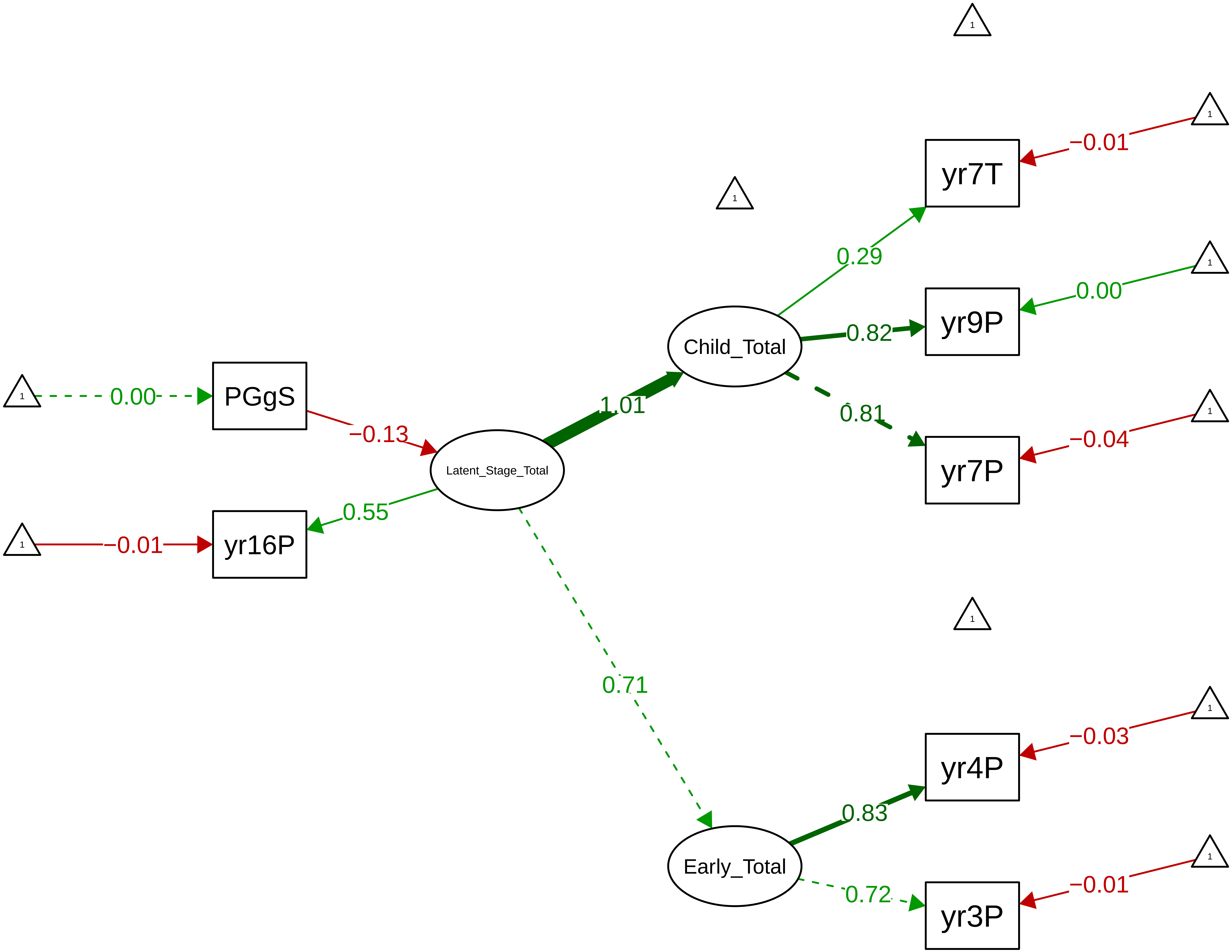

Figure S9_26: Conners Inattention − CTCR Overall

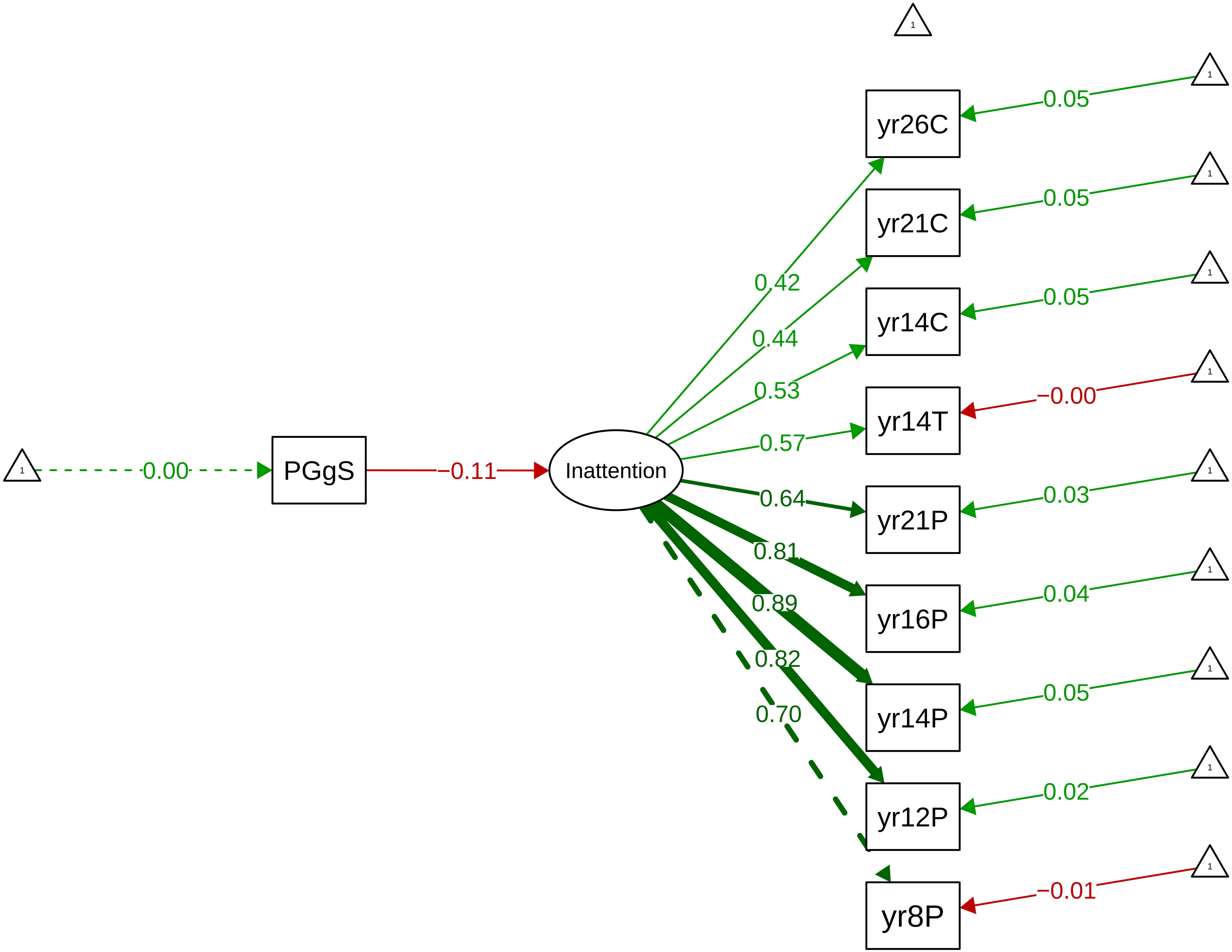

Figure S9_27: Conners Inattention − CTCR Stage

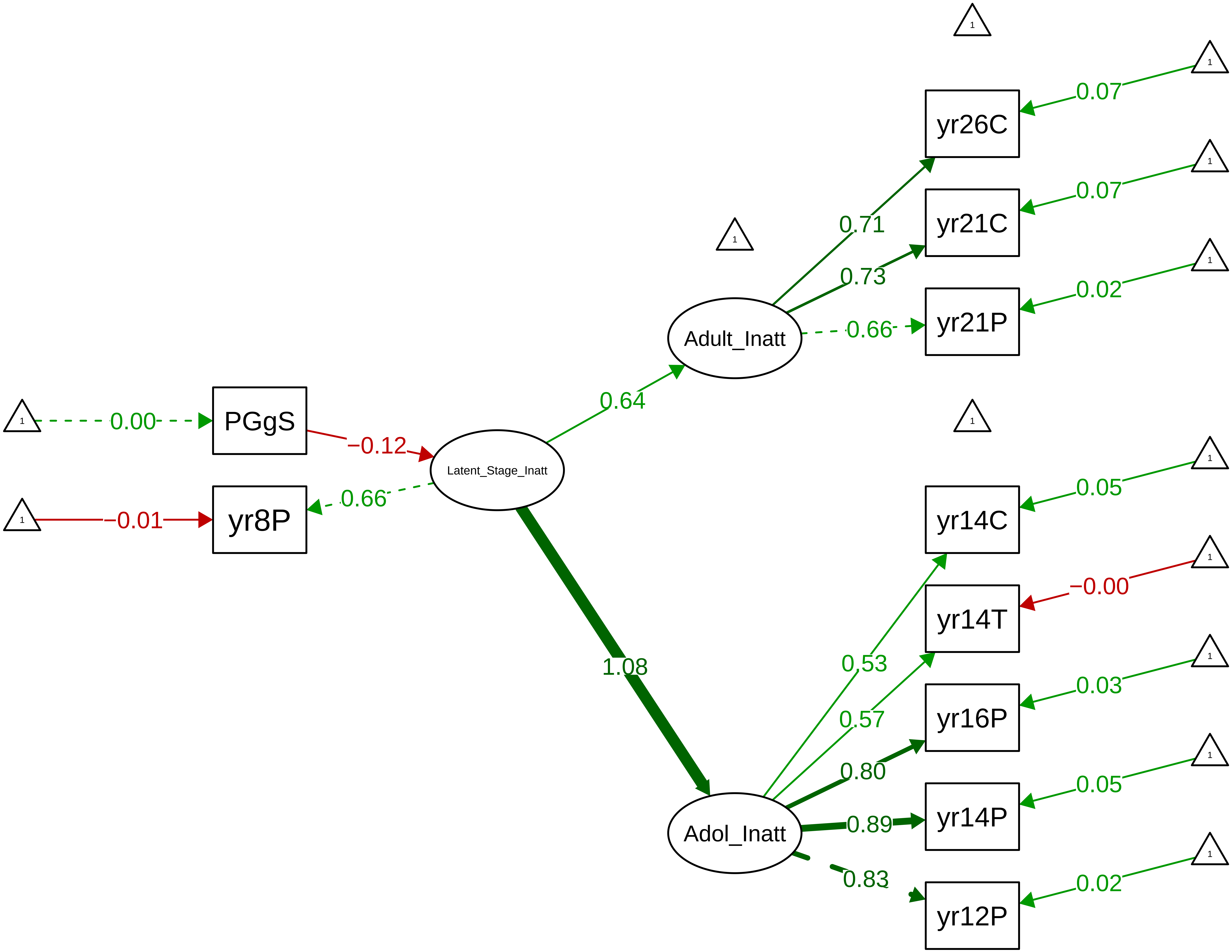

Figure S9_28: Conners Inattention – Parent Overall

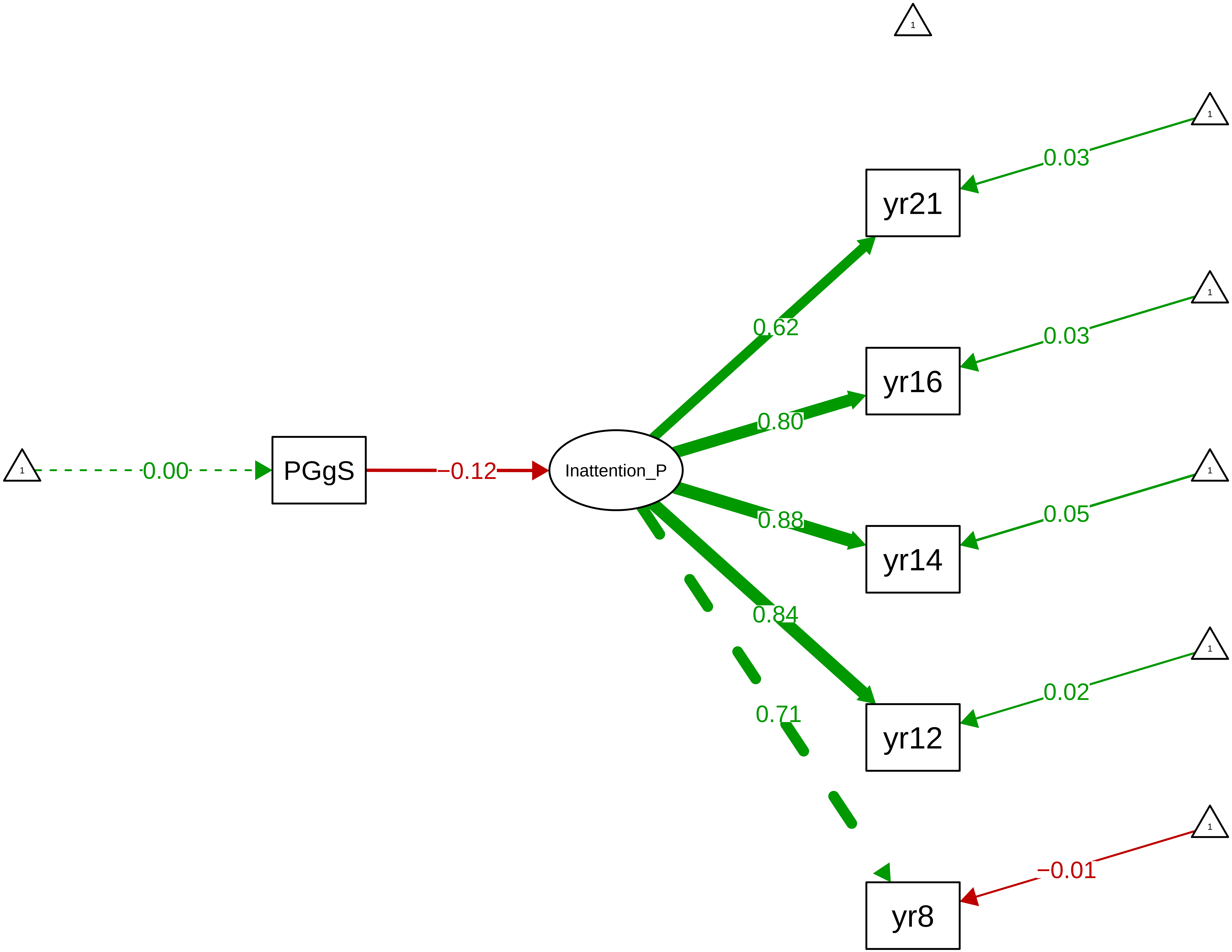

**Figure S9_29: Conners Inattention − Parent Stage**

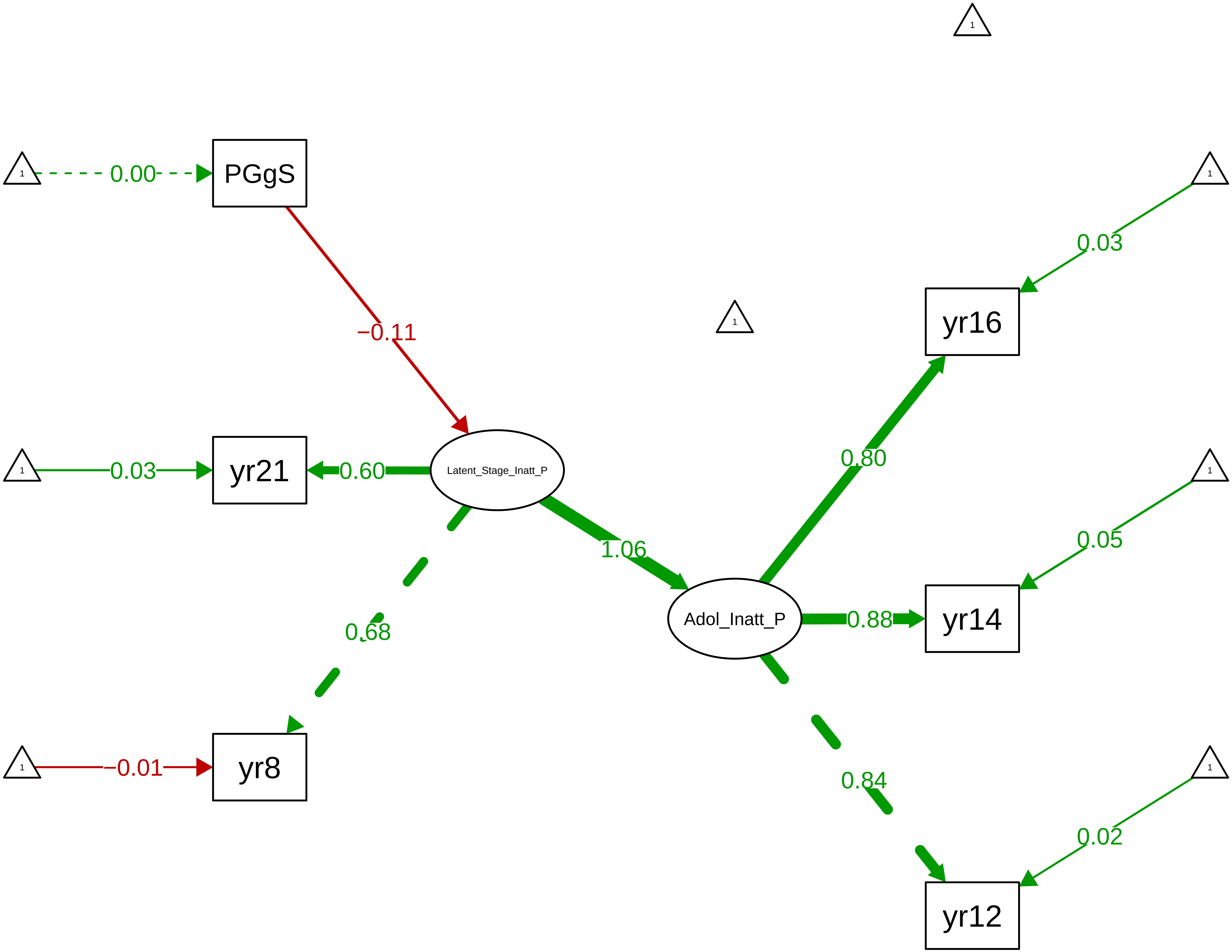

Figure S9_30: Conners Hyperactivity−Impulsivity − CTCR Overall

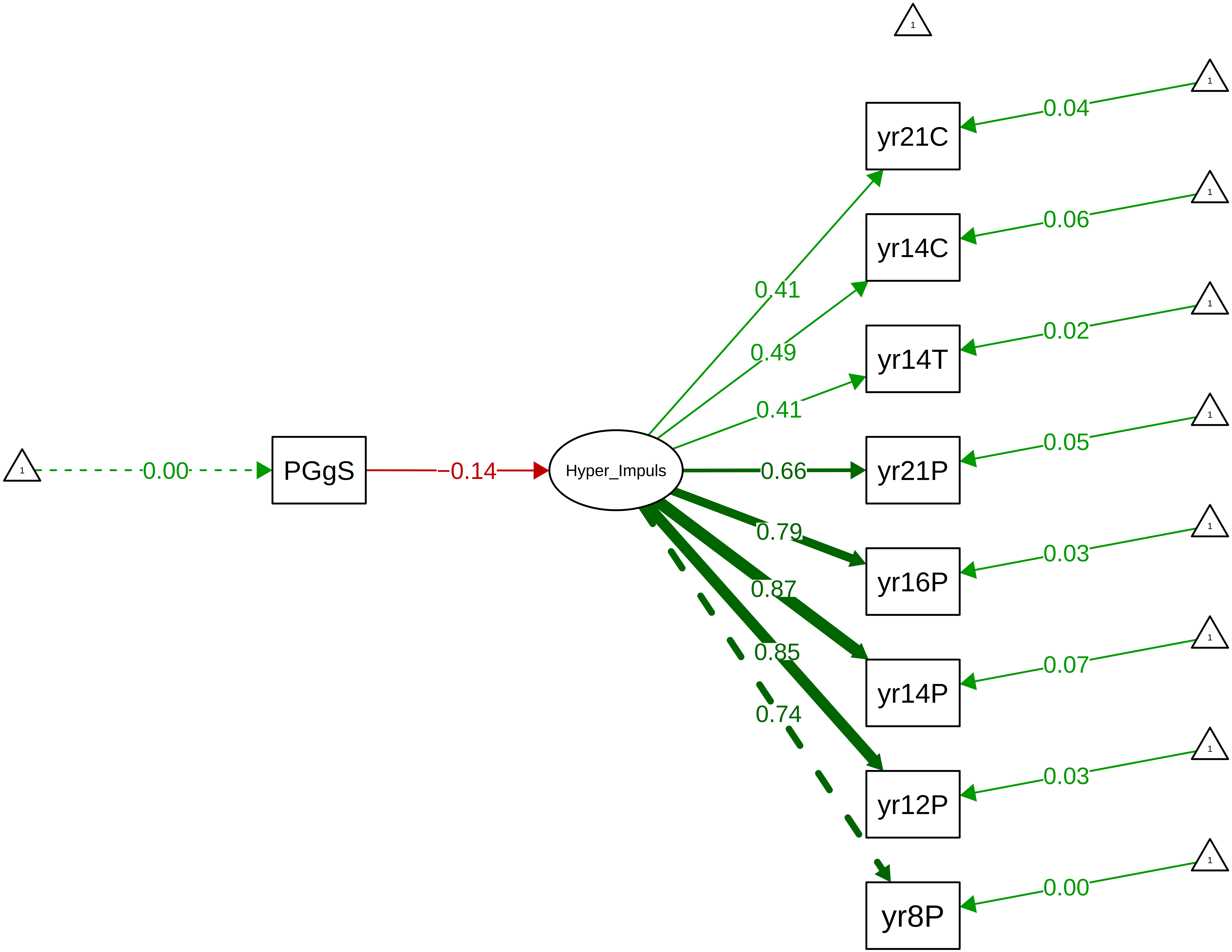

**Figure S9_31: Conners Hyperactivity−Impulsivity − CTCR Stage**

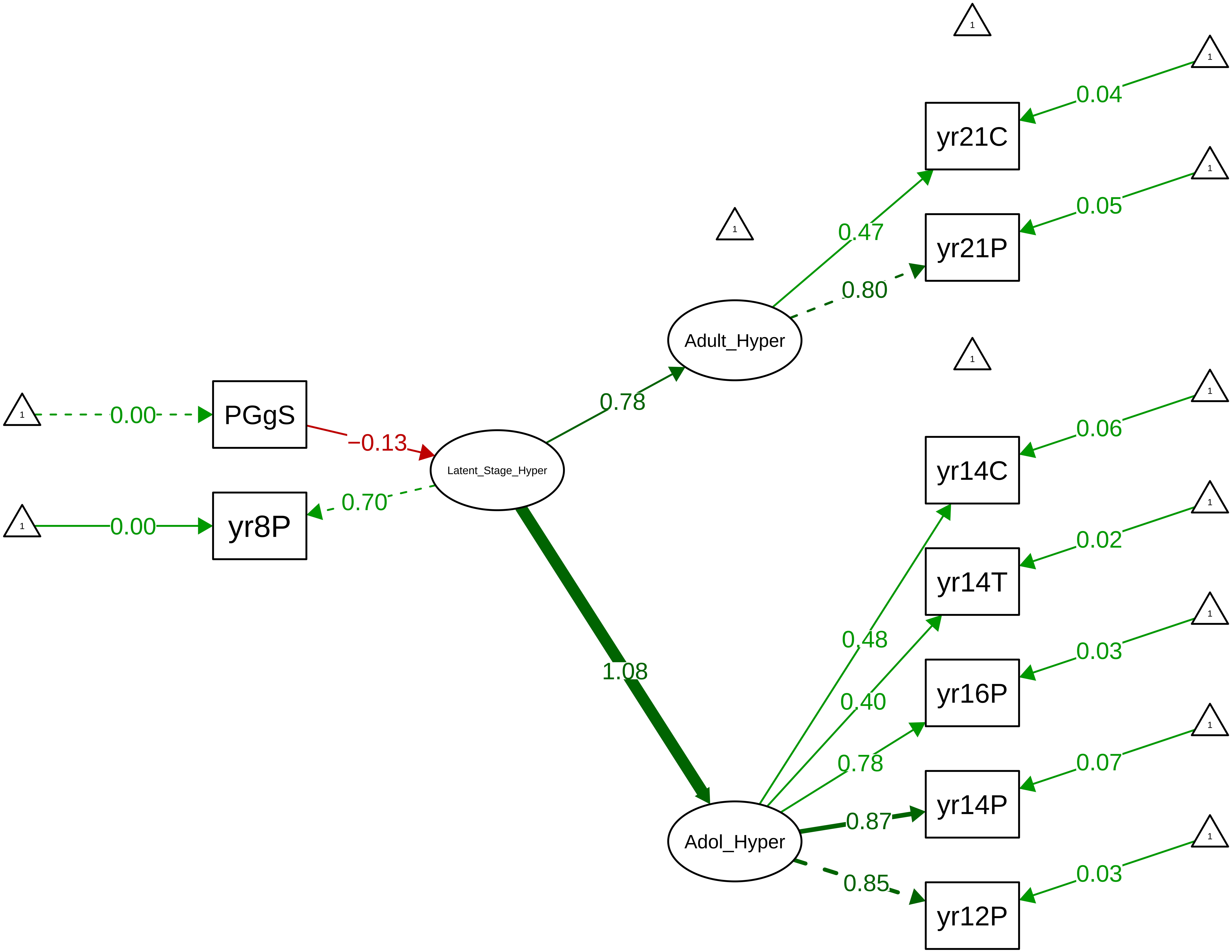

Figure S9_32: Conners Hyperactivity−Impulsivity − Parent Overall

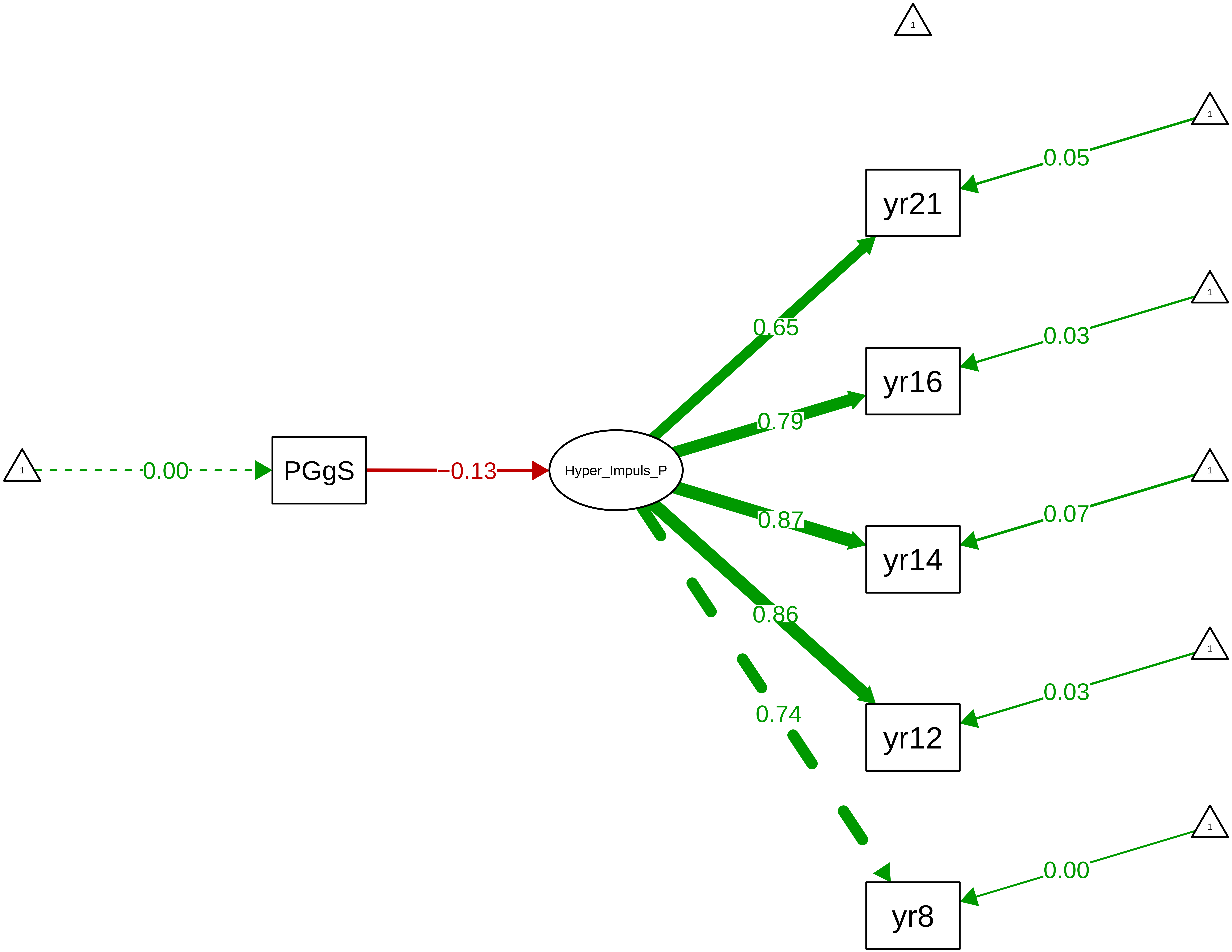

Figure S9_33: Conners Hyperactivity−Impulsivity − Parent Stage

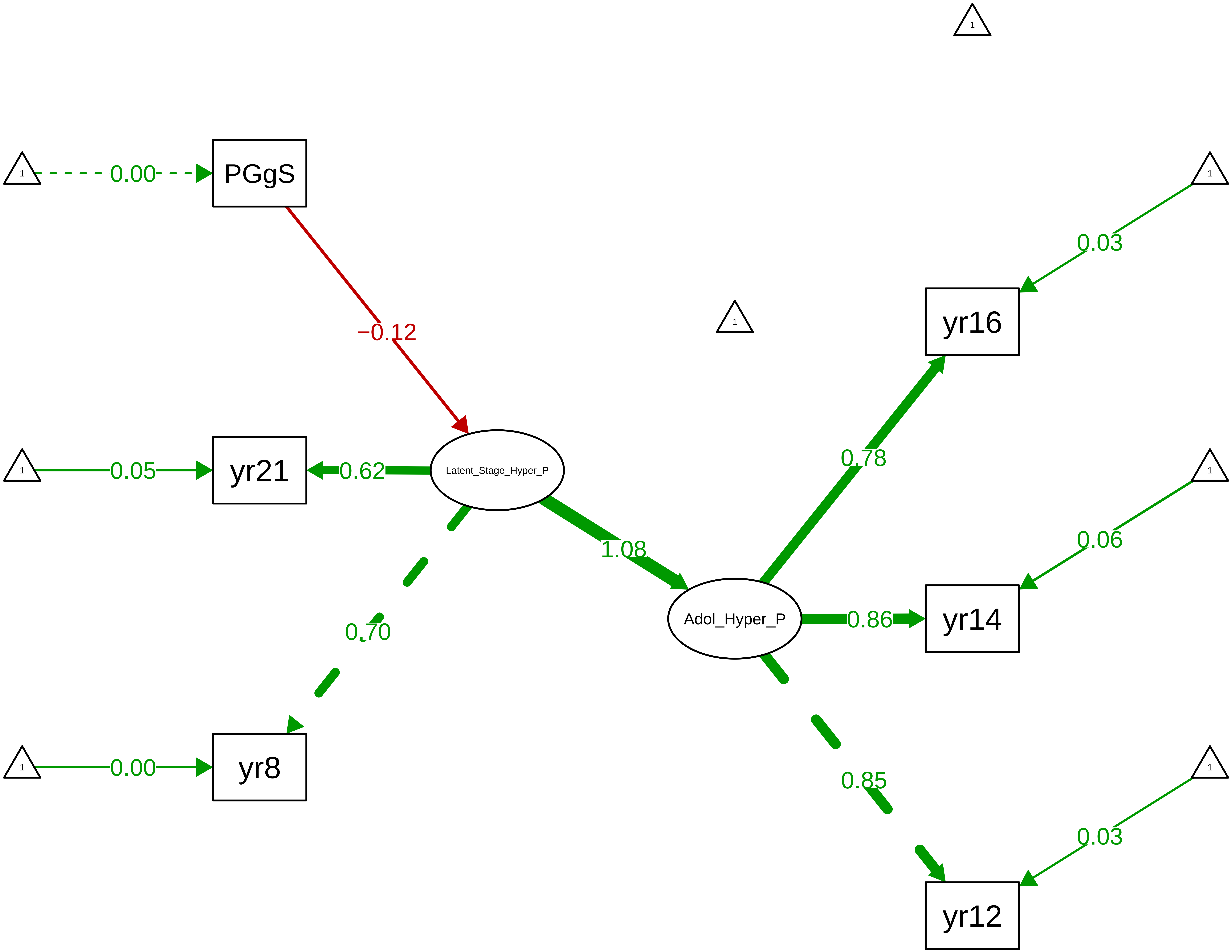

**Figure S9_34: Conners Total − CTCR Overall**

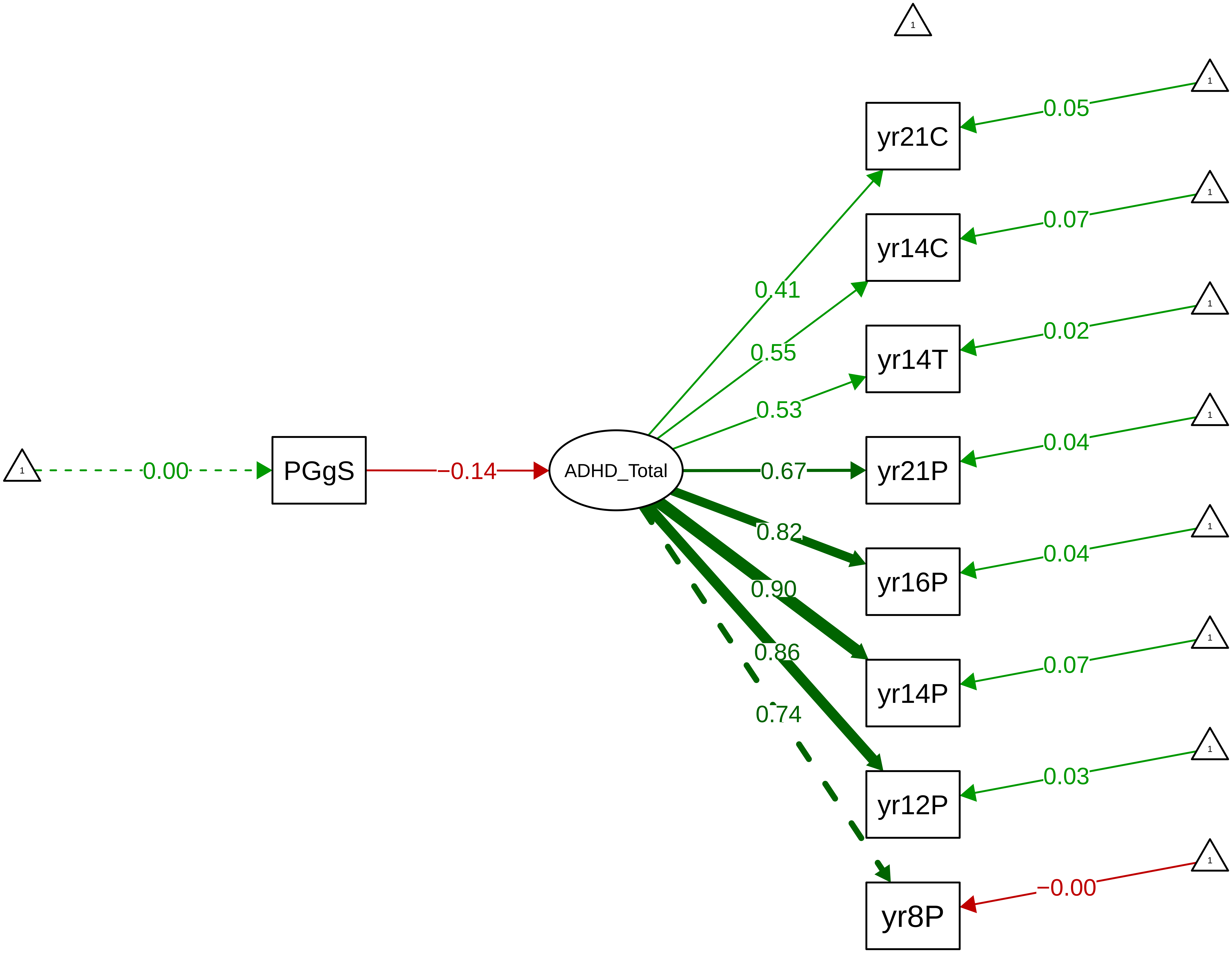

**Figure S9_35: Conners Total − CTCR Stage**

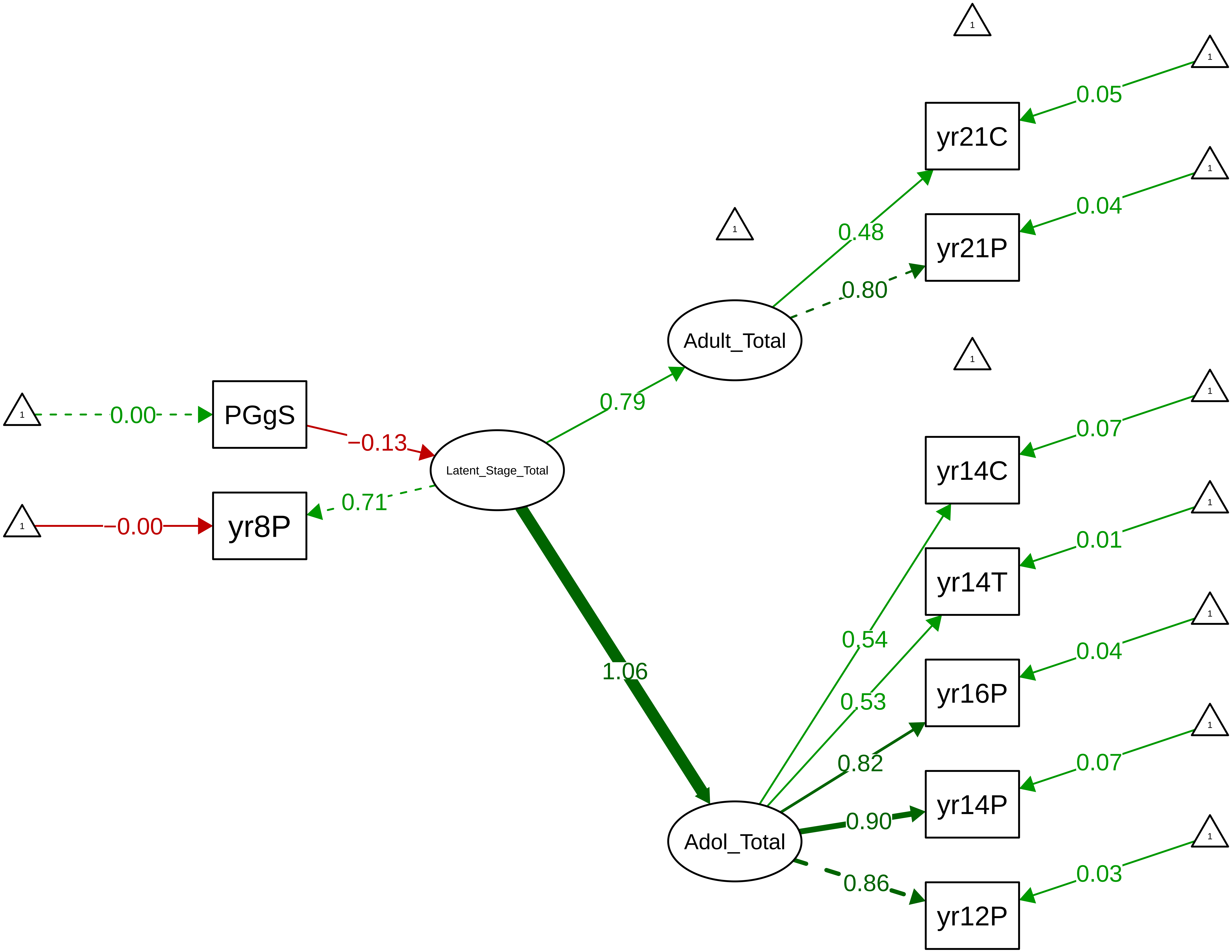

#### Figure S9_36: Conners Total − Parent Overall

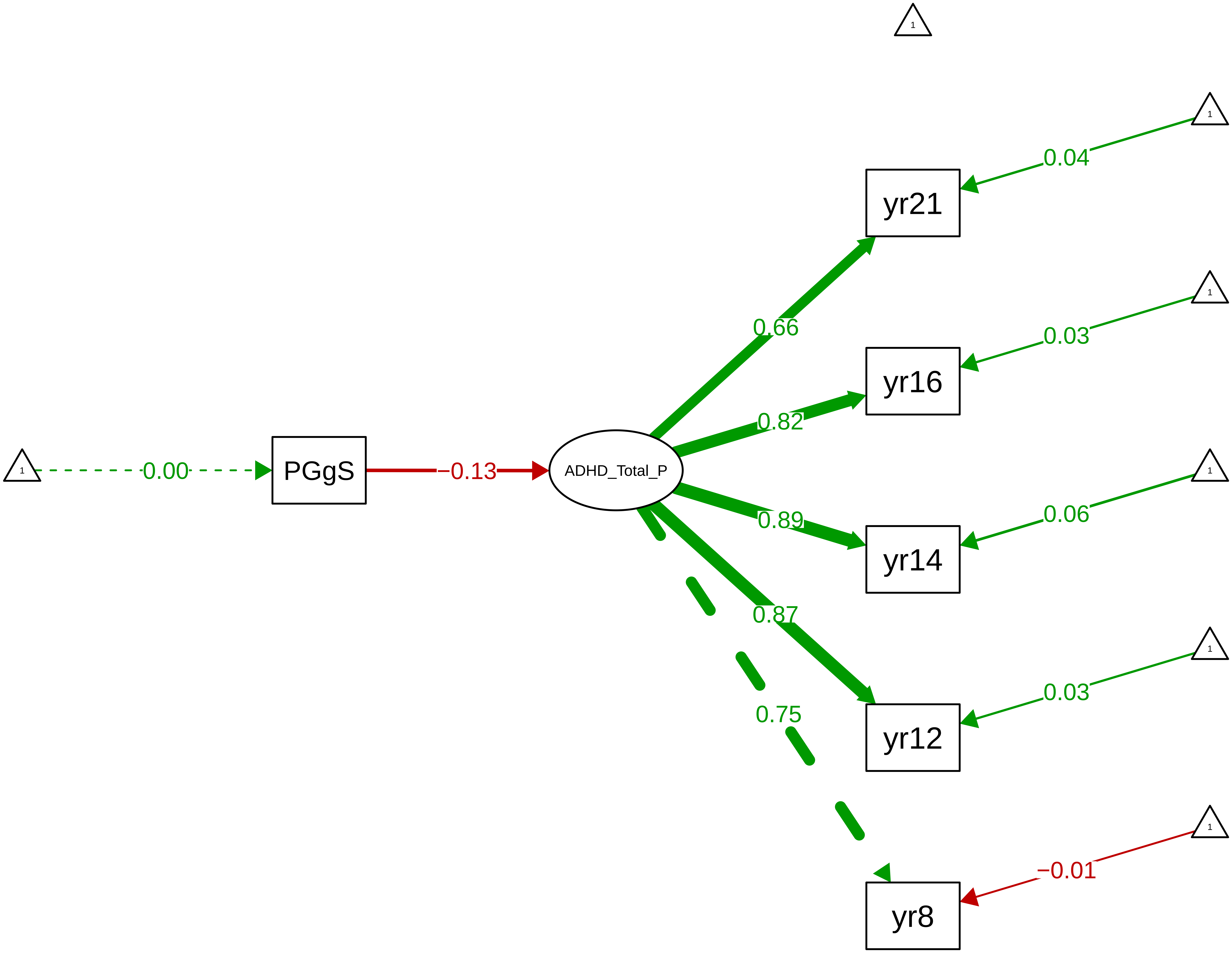

**Figure S9_37: Conners Total − Parent Stage**

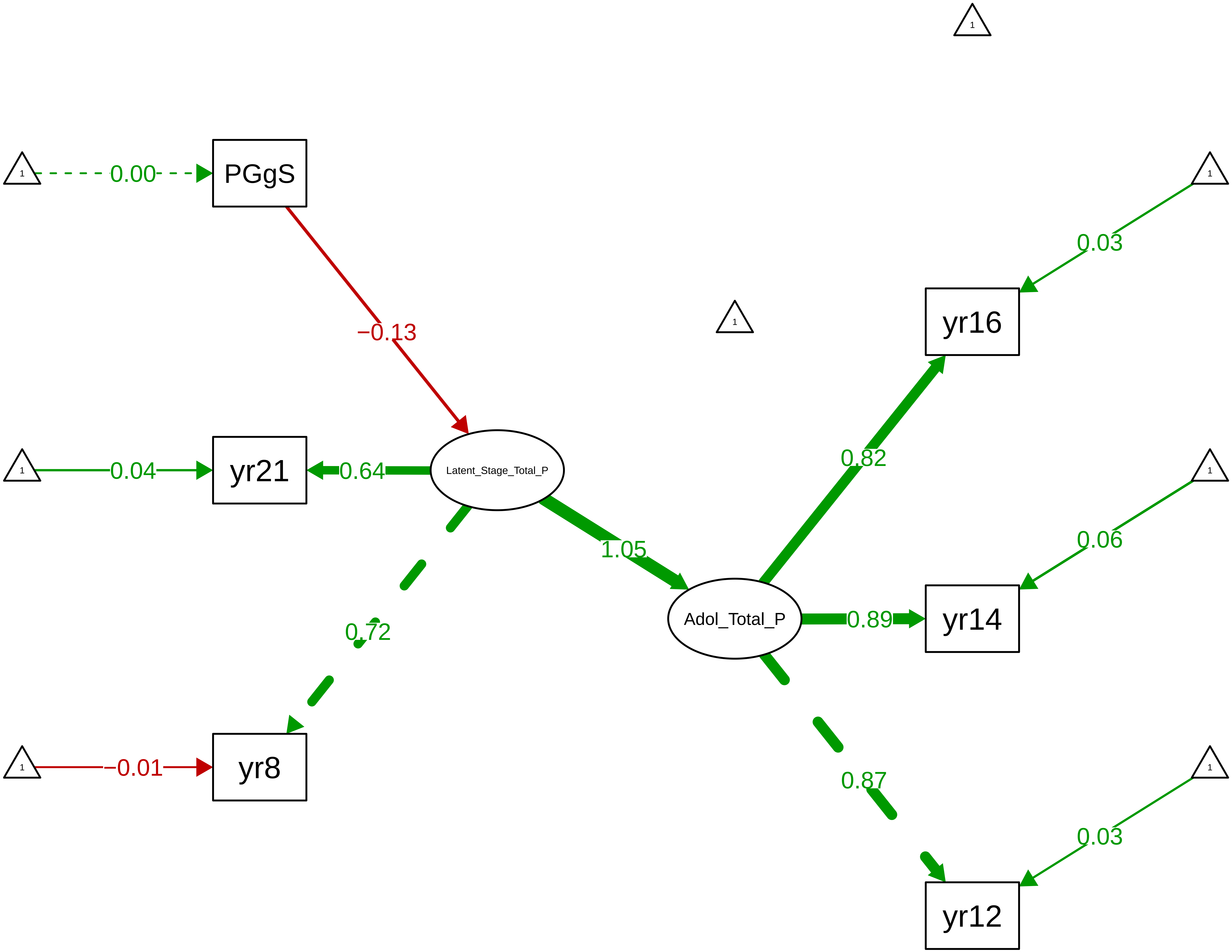

**Figure S9_38: SDQ Conduct − CTCR Overall**

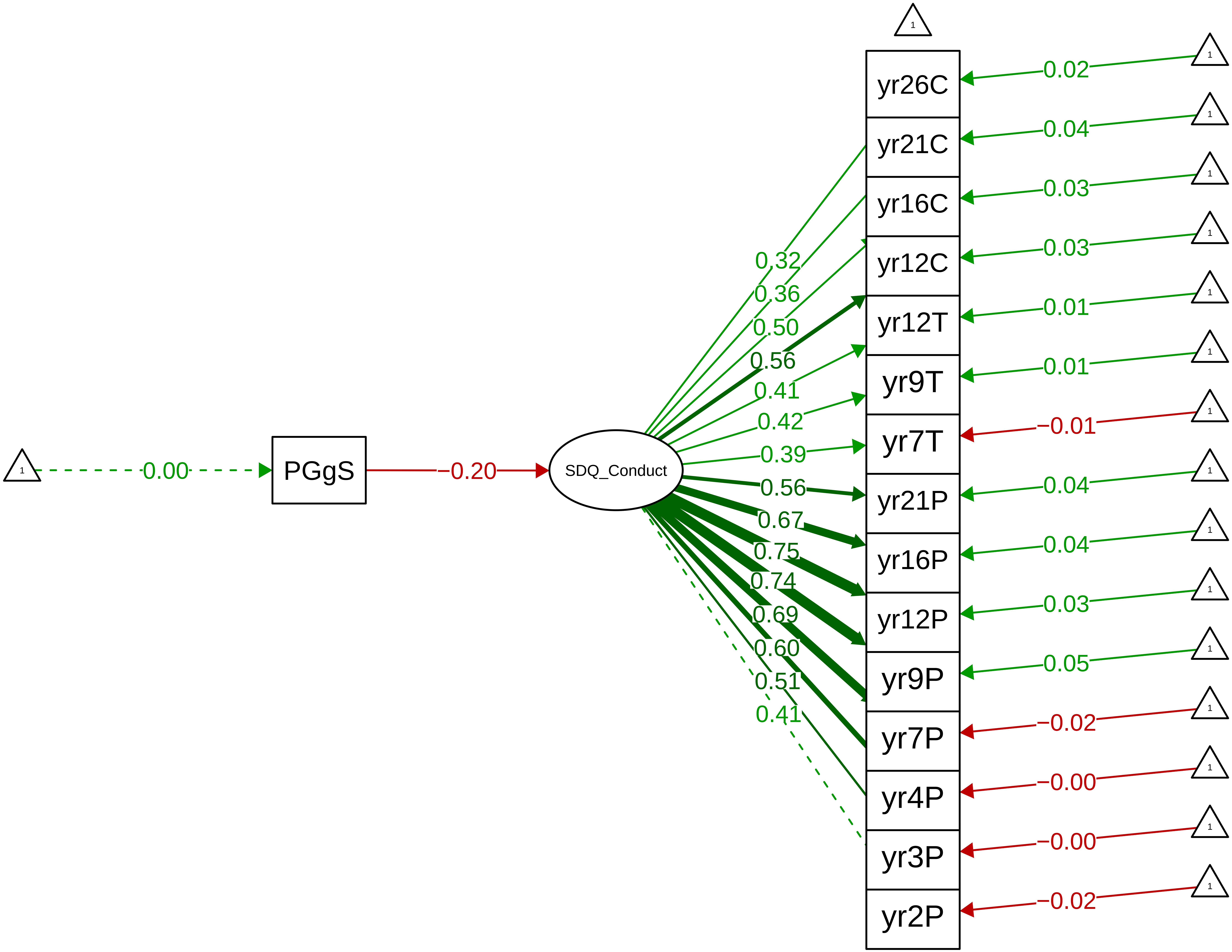

**Figure S9_39: SDQ Conduct − CTCR Stage**

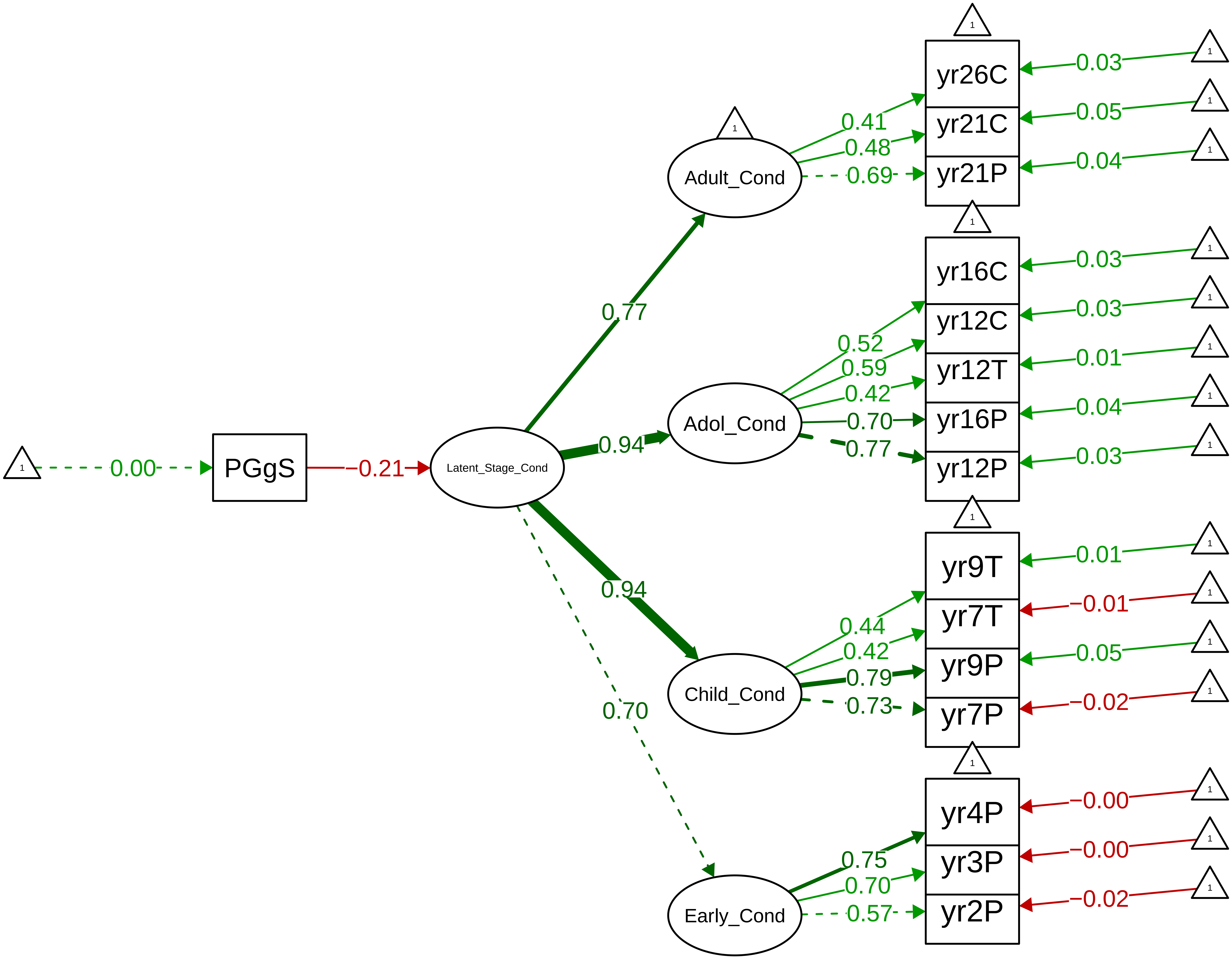

**Figure S9_40: SDQ Conduct − Parent Overall**

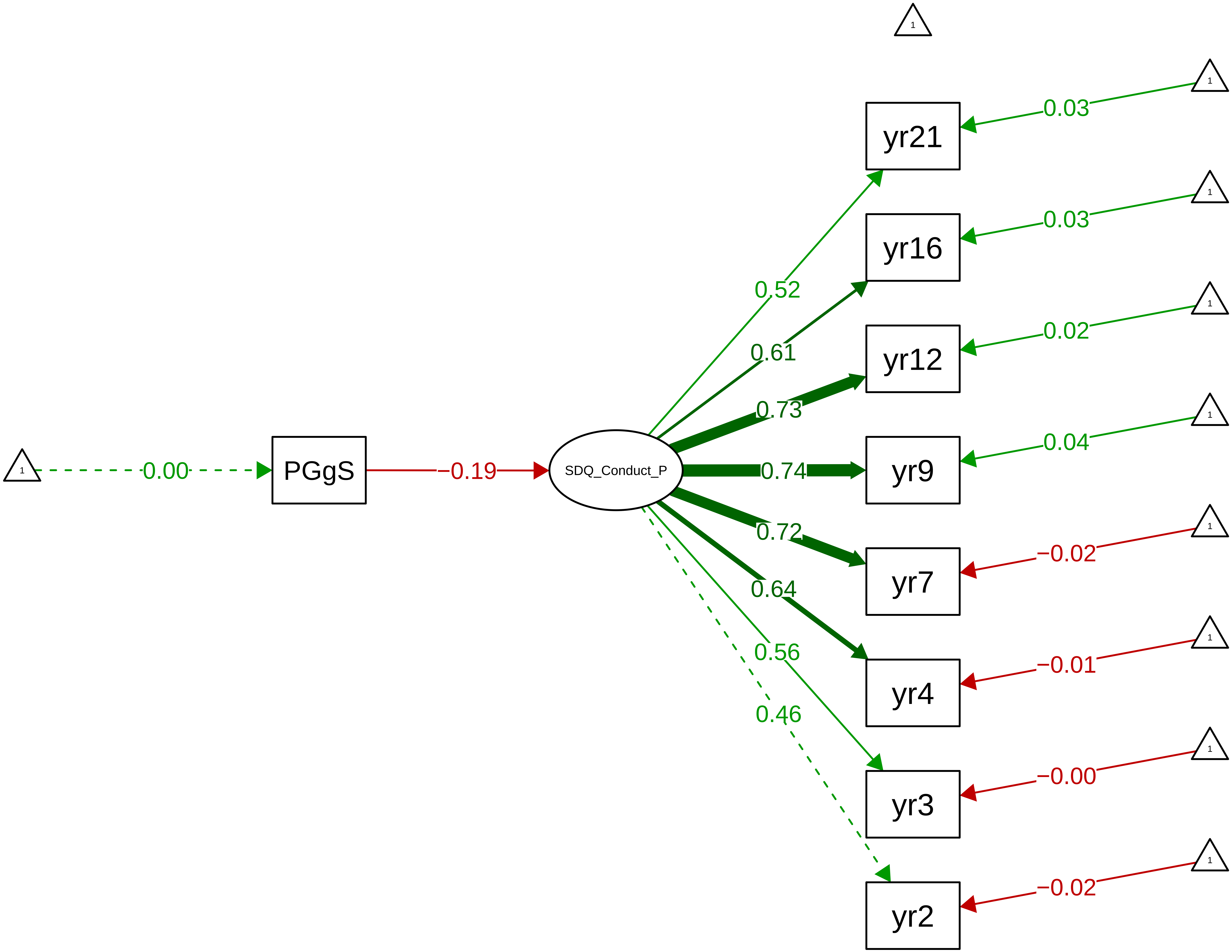

**Figure S9_41: SDQ Conduct − Parent Stage**

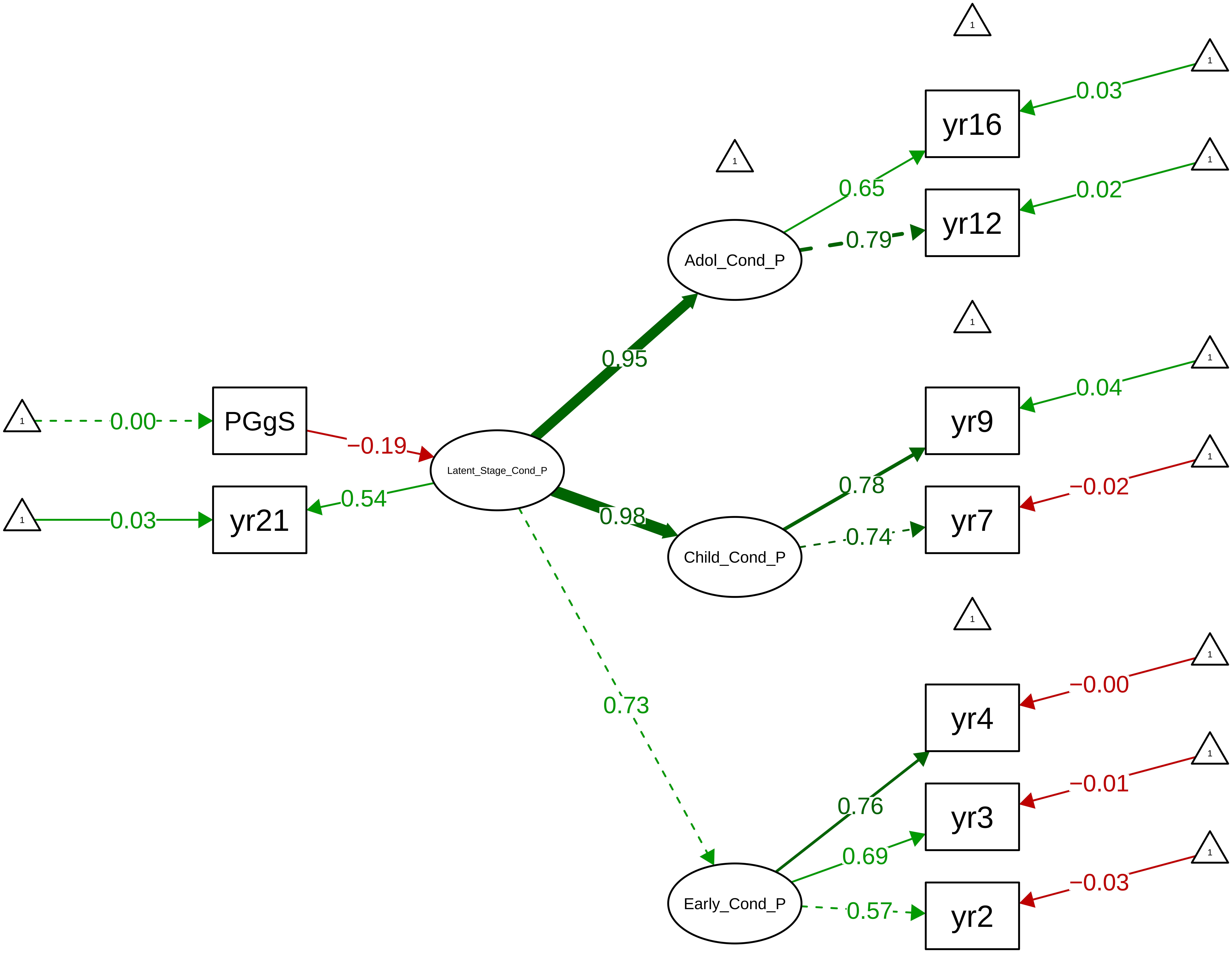

Figure S9_42: SDQ Conduct − Teacher Overall

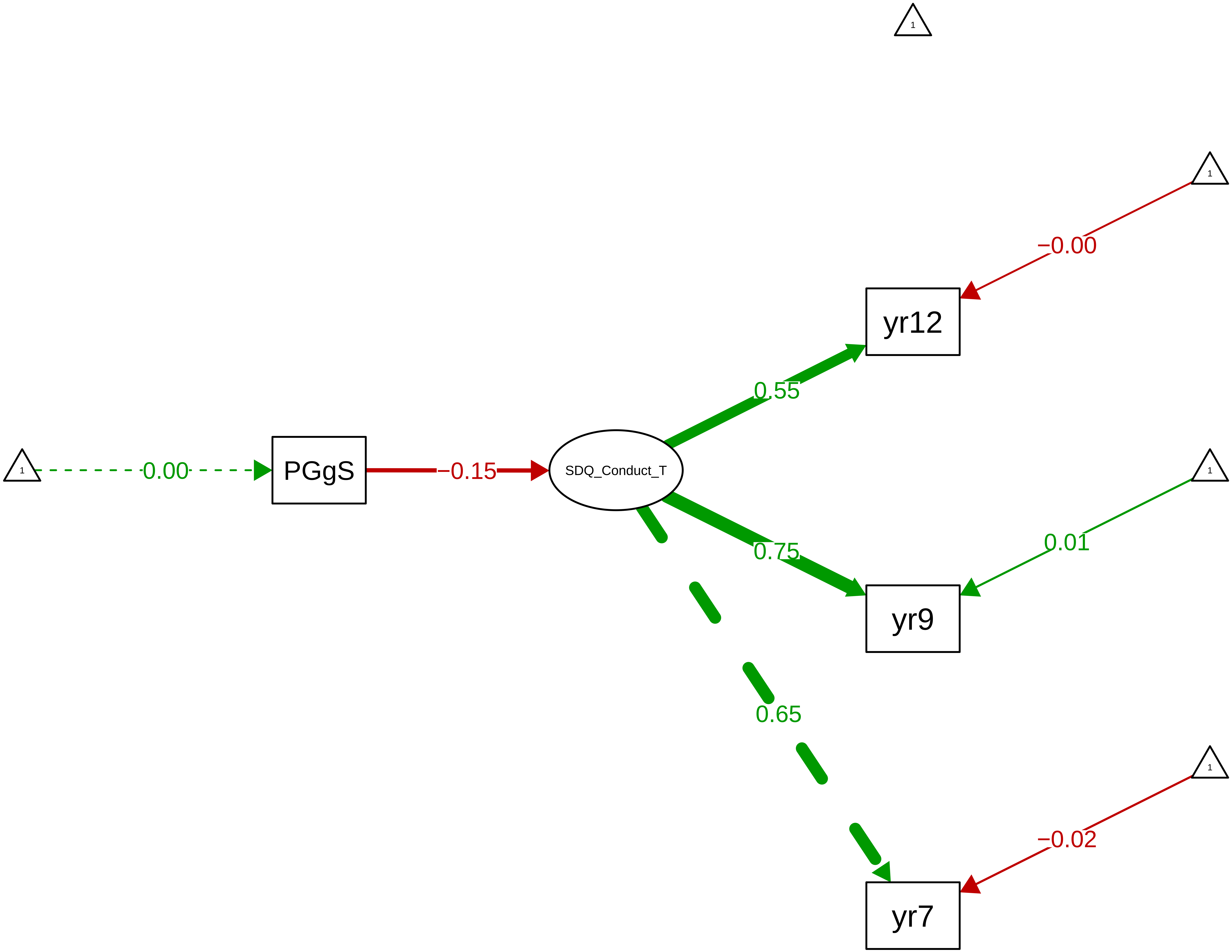

**Figure S9_43: SDQ Conduct − Child Overall**

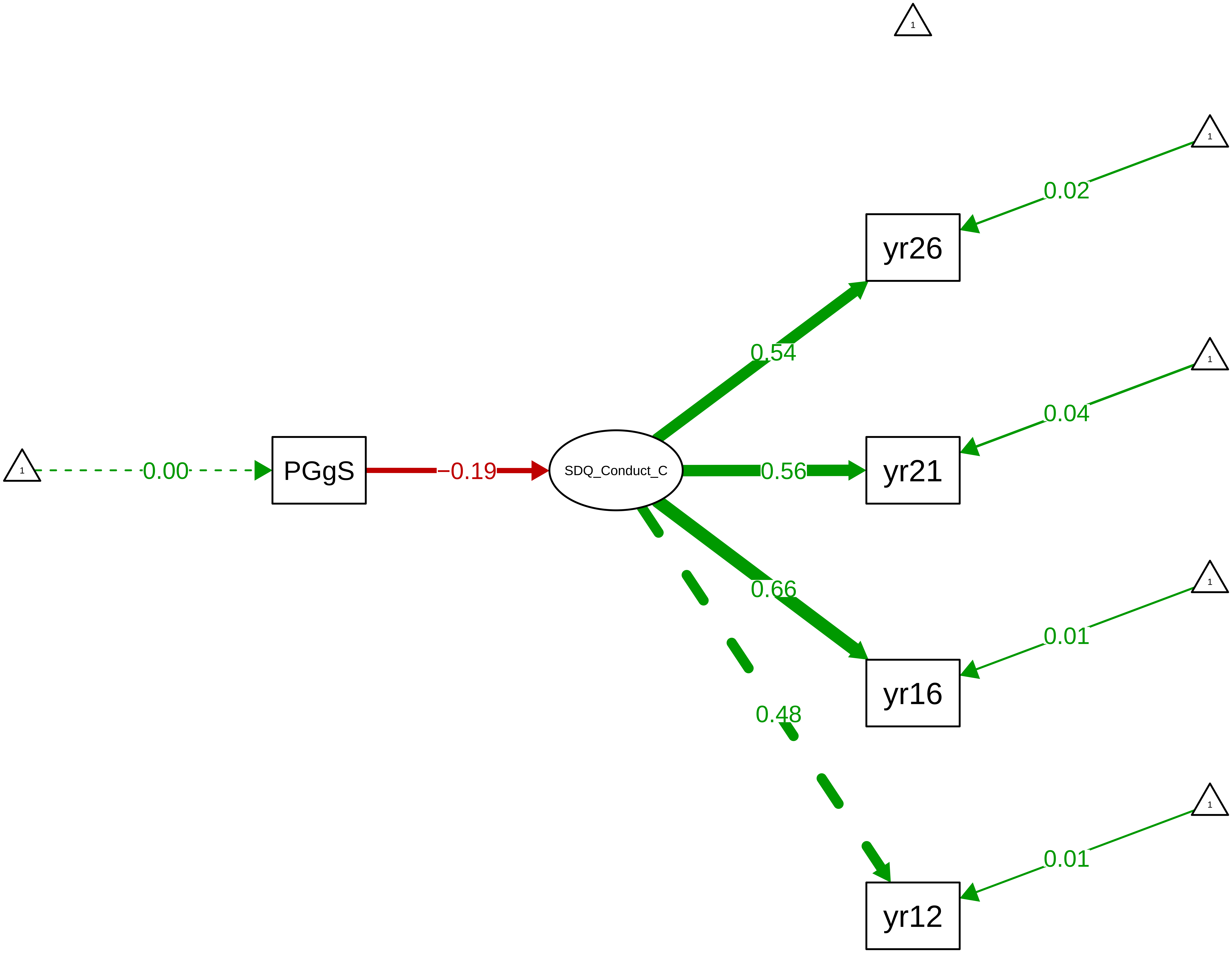

**Figure S9_44: SDQ Emotion – CTCR Overall**

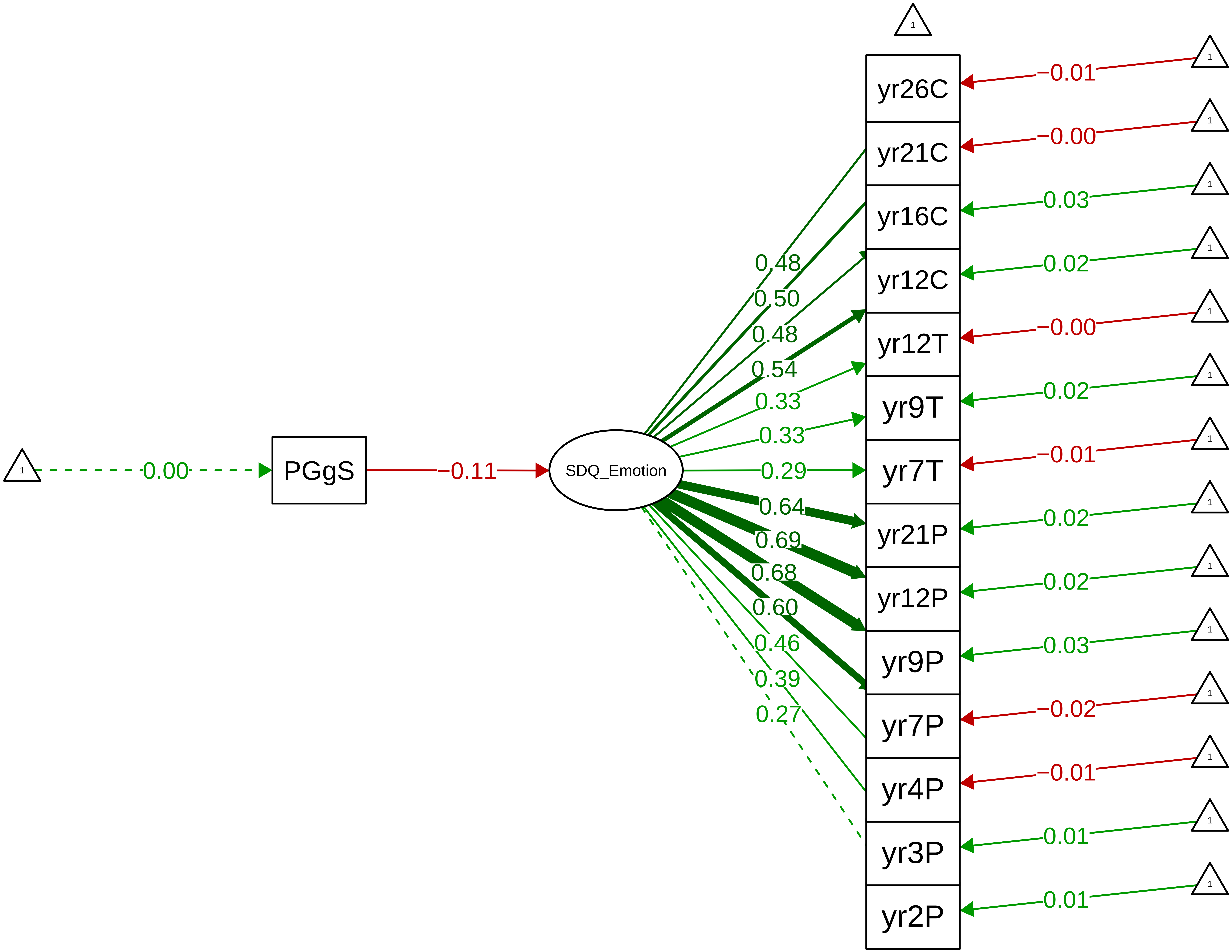

**Figure S9_45: SDQ Emotion − CTCR Stage**

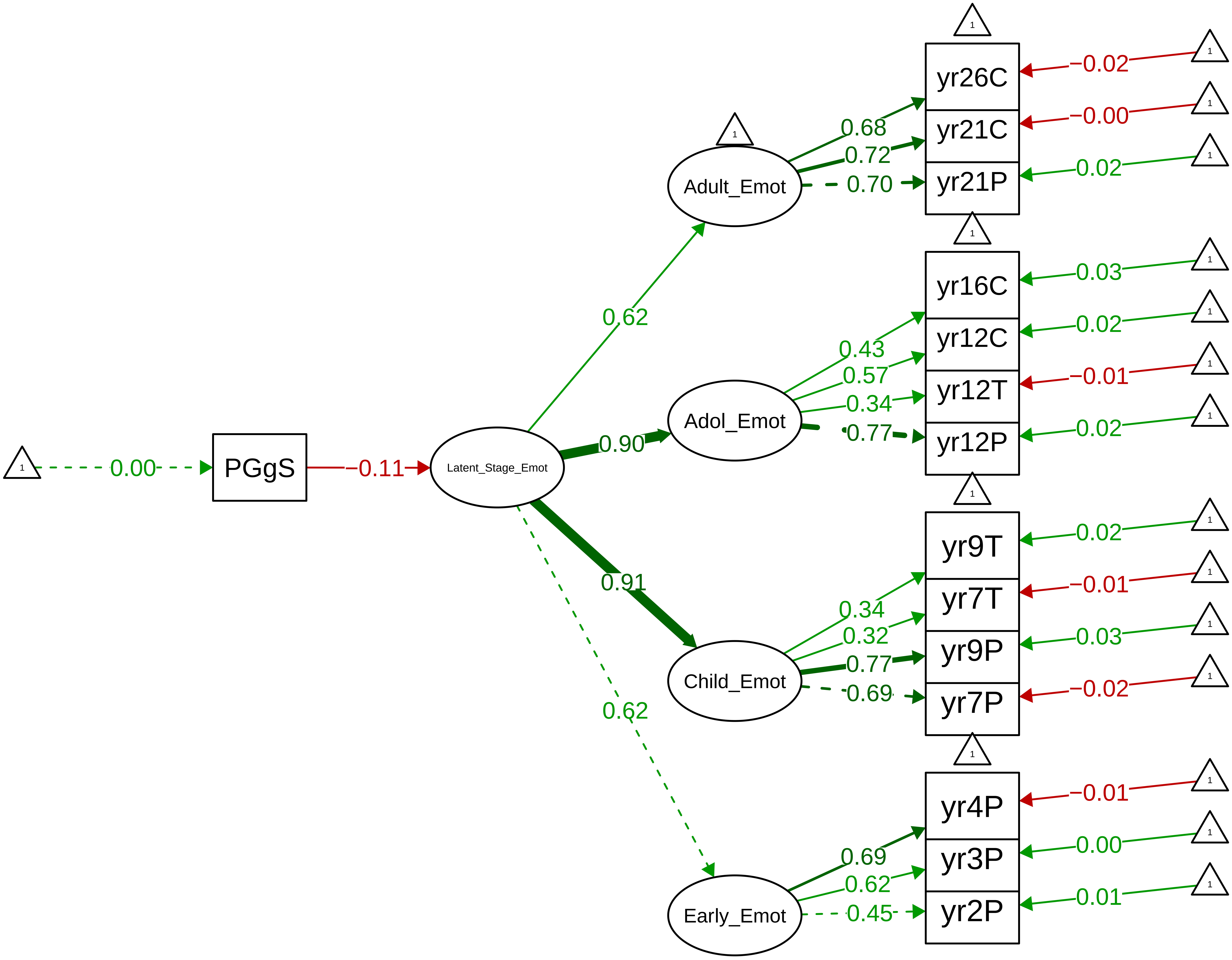

**Figure S9_46: SDQ Emotion − Parent Overall**

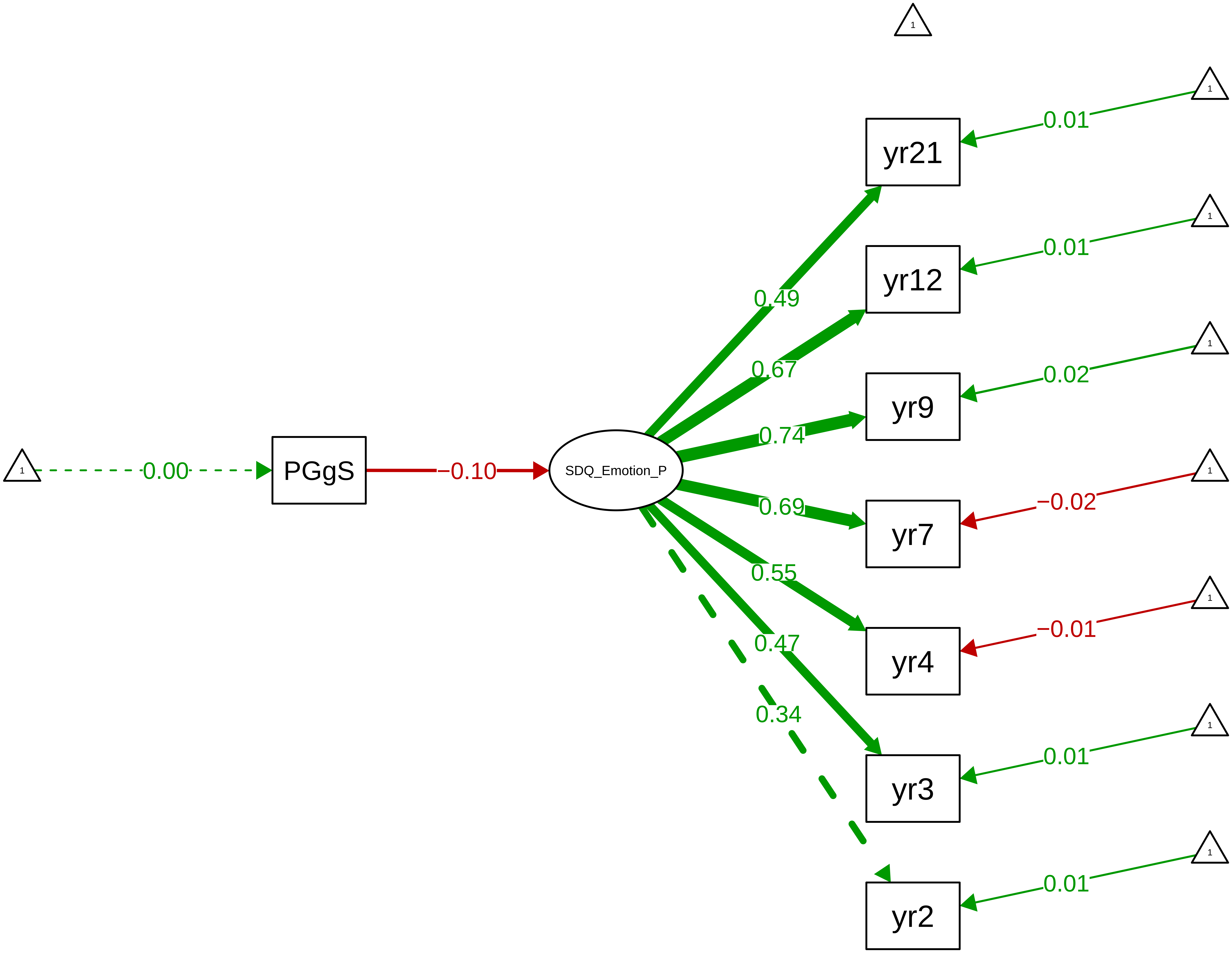

#### Figure S9_47: SDQ Emotion − Parent Stage

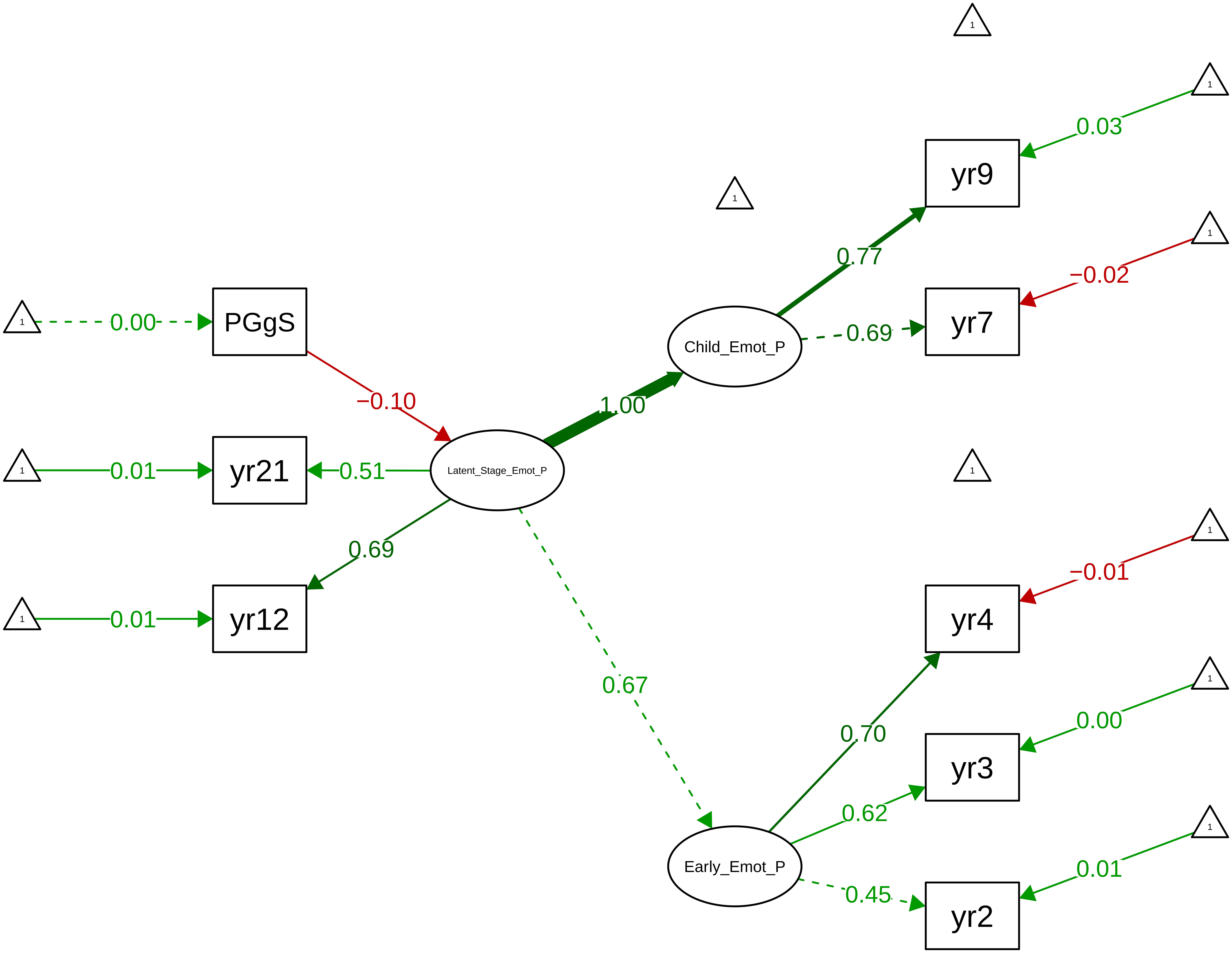

Figure S9_48: SDQ Emotion − Teacher Overall

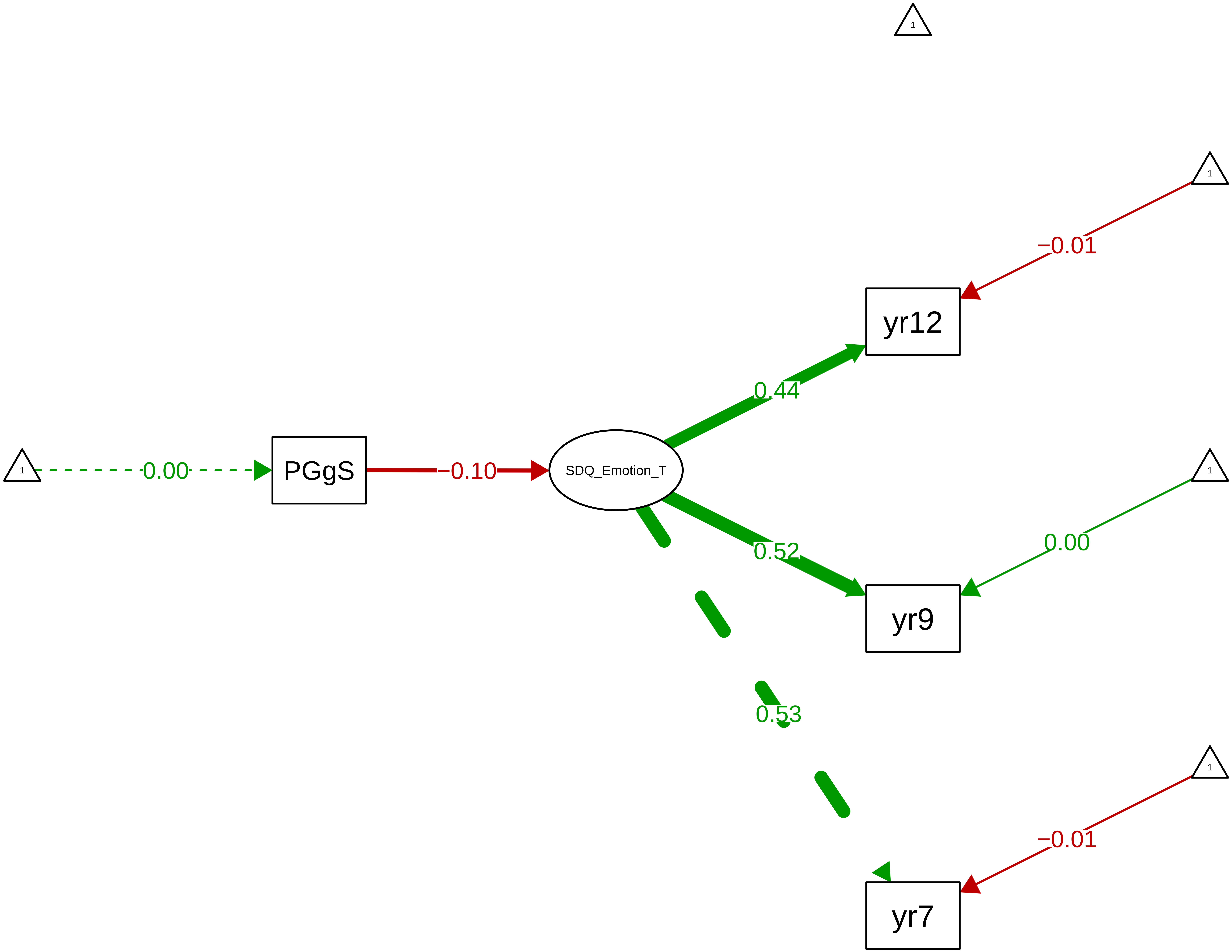

#### Figure S9_49: SDQ Emotion − Child Overall

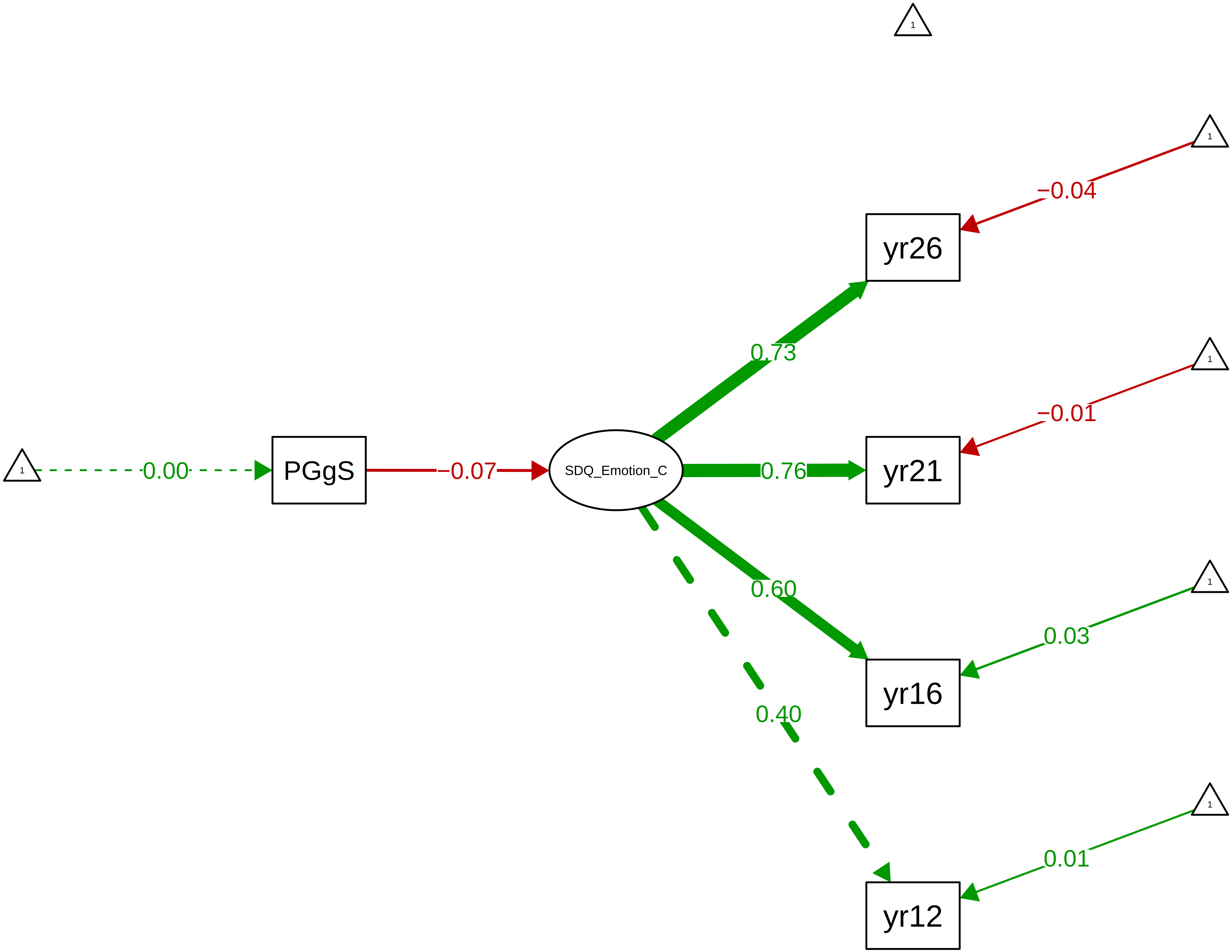

#### Figure S9_50: SDQ Hyperactivity − CTCR Overall

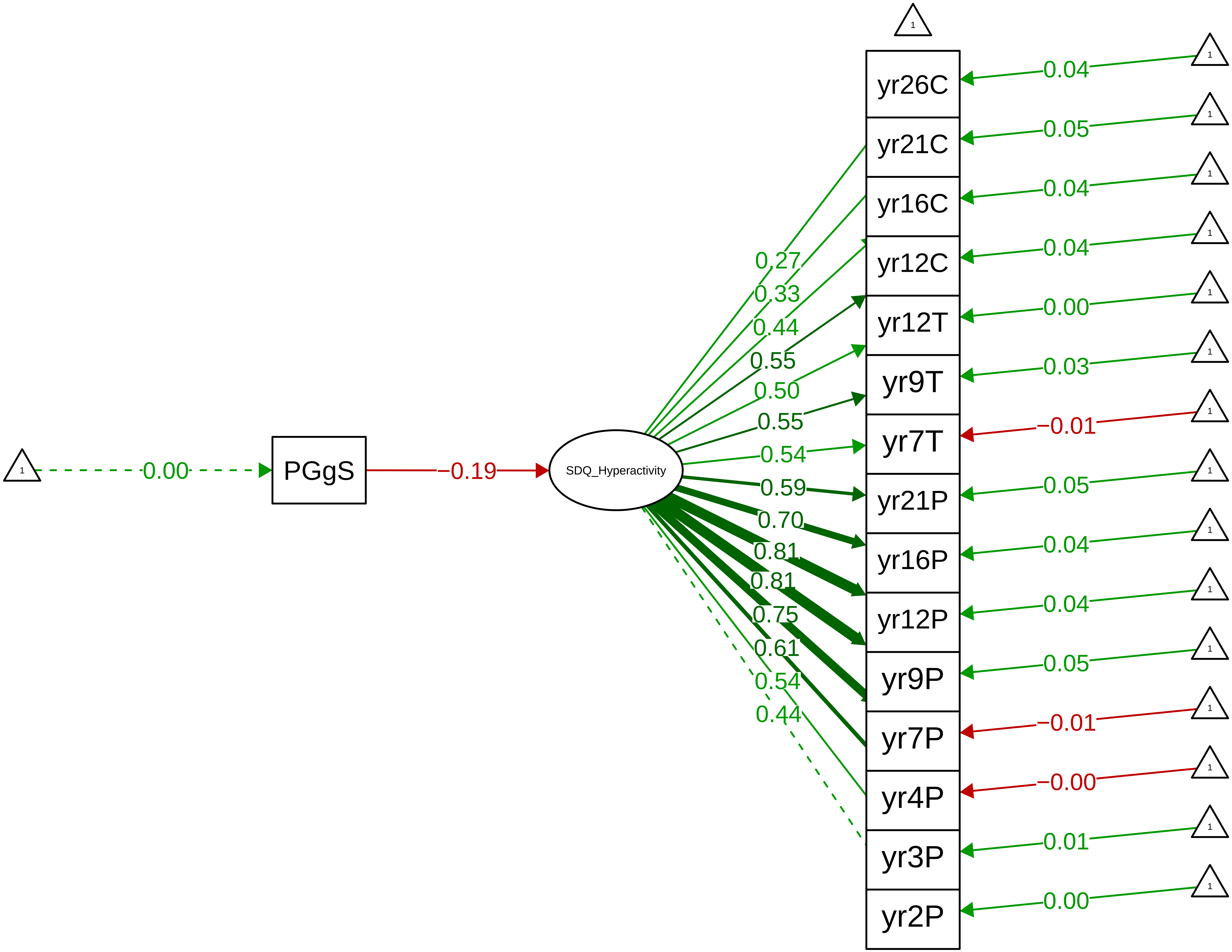

**Figure S9_51: SDQ Hyperactivity − CTCR Stage**

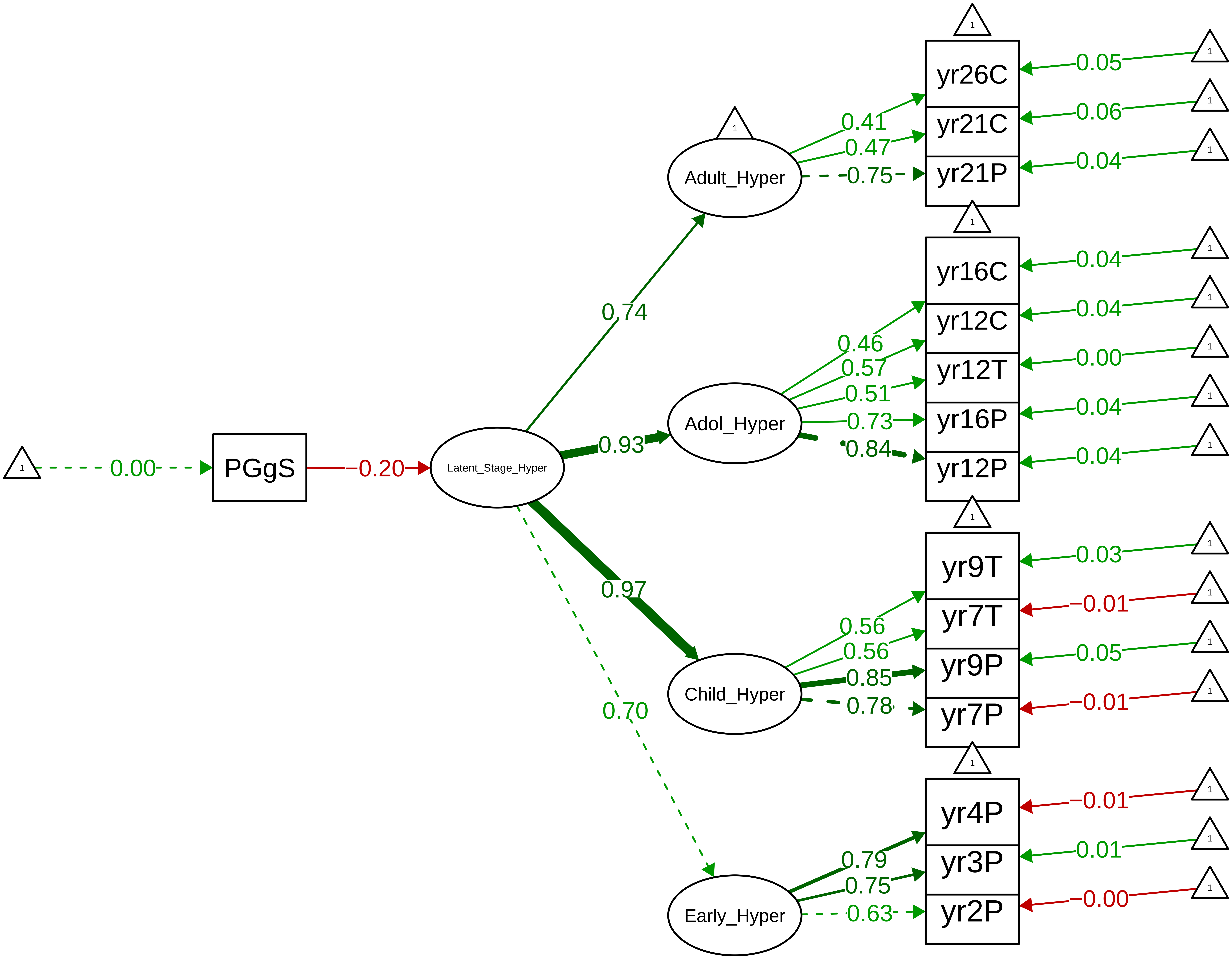

Figure S9_52: SDQ Hyperactivity − Parent Overall

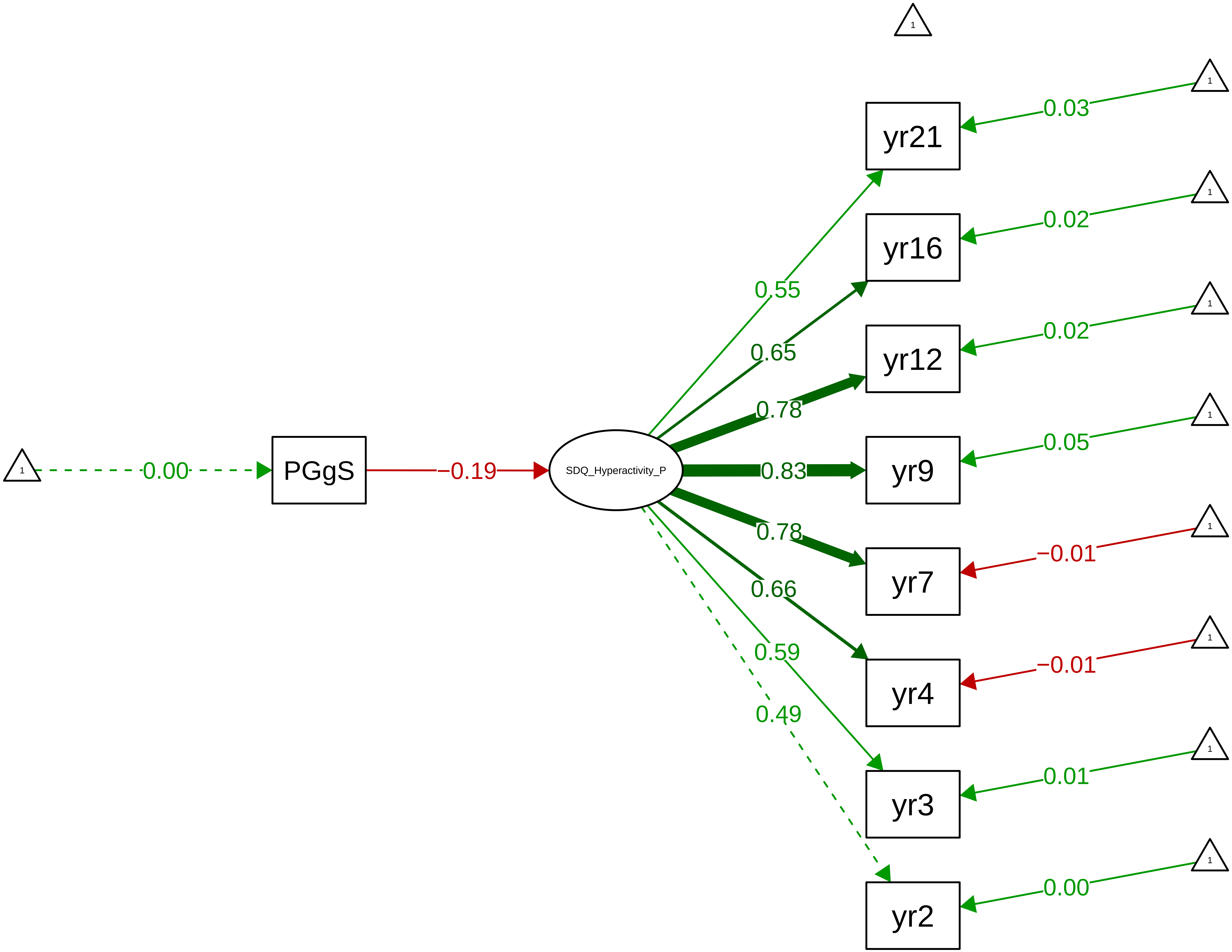

**Figure S9_53: SDQ Hyperactivity − Parent Stage**

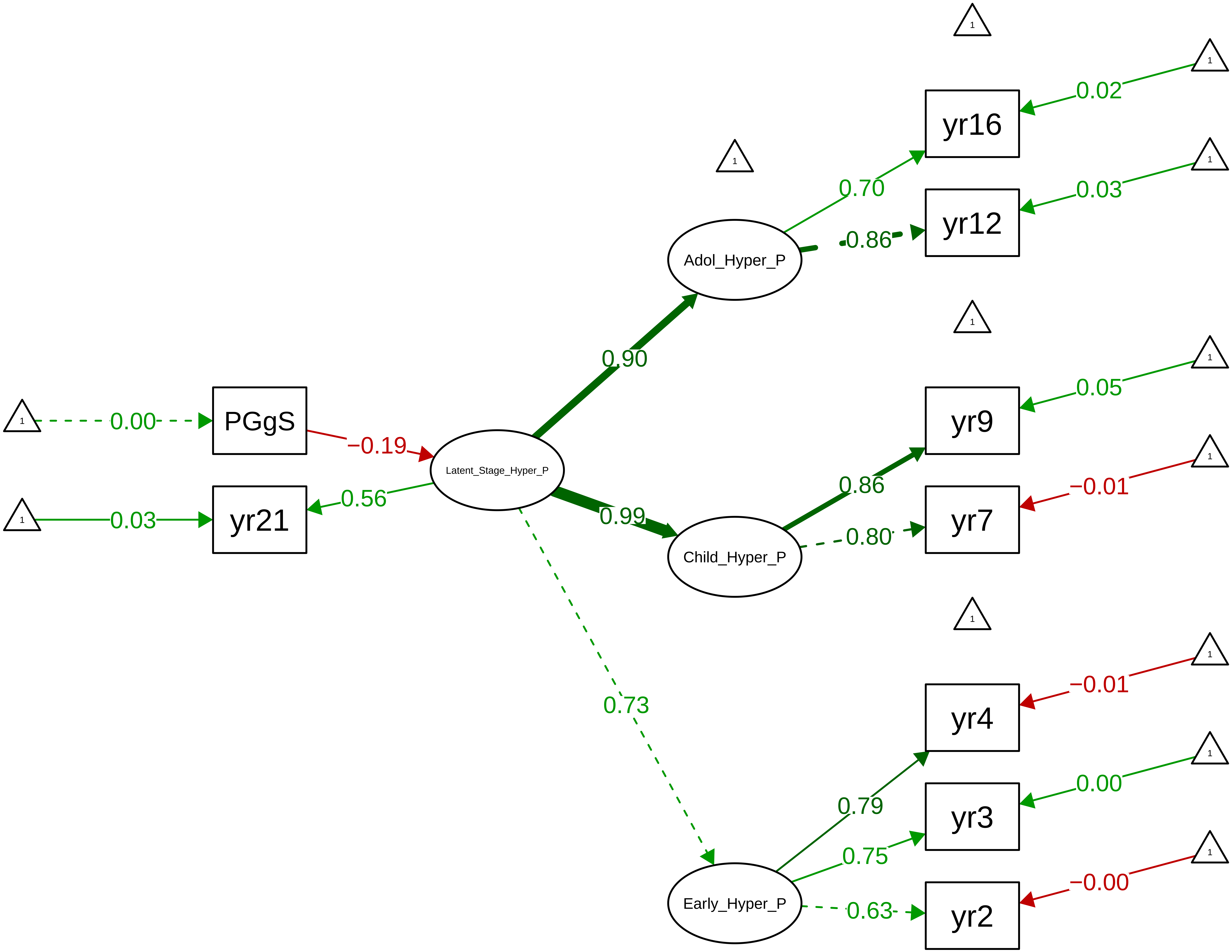

Figure S9_54: SDQ Hyperactivity − Teacher Overall

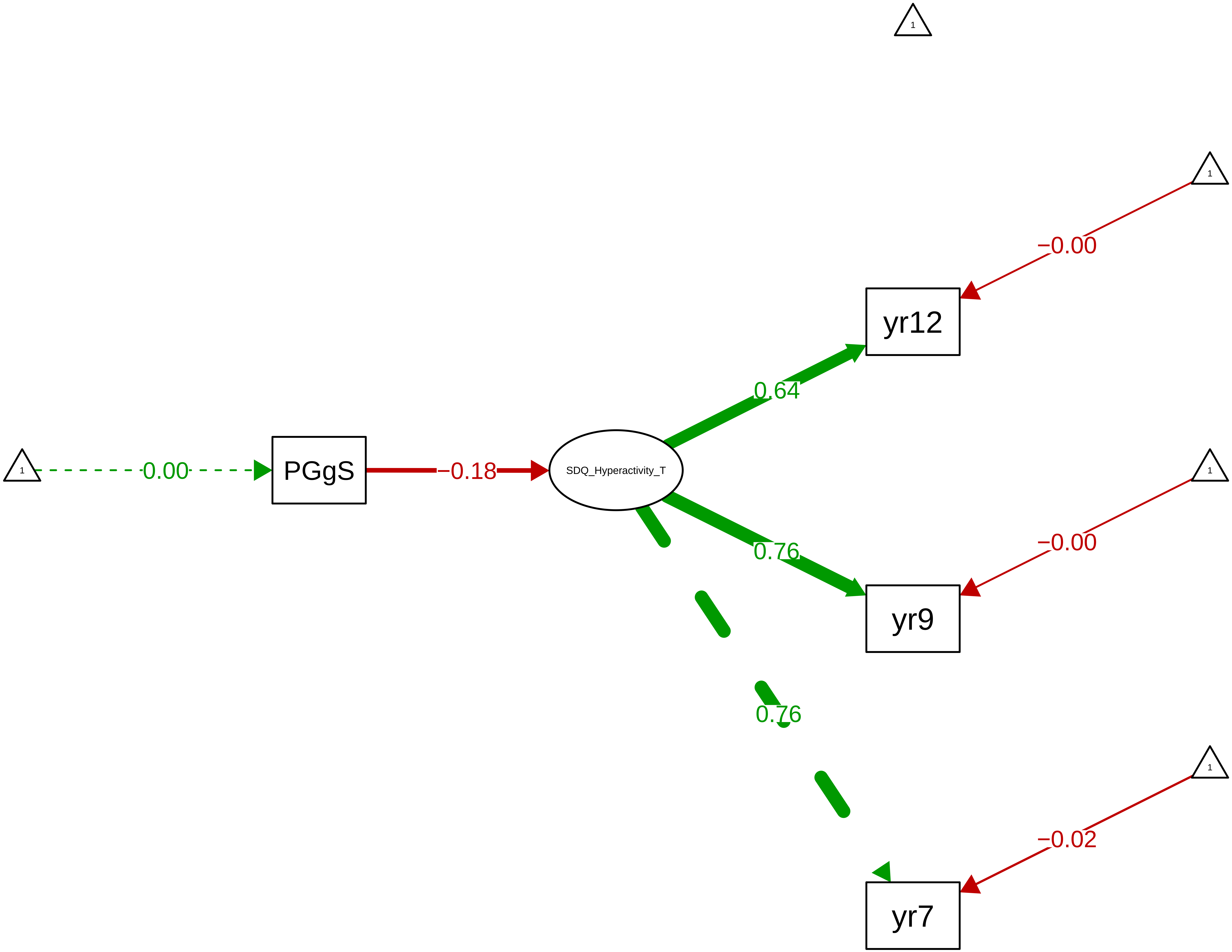

Figure S9_55: SDQ Hyperactivity – Child Overall

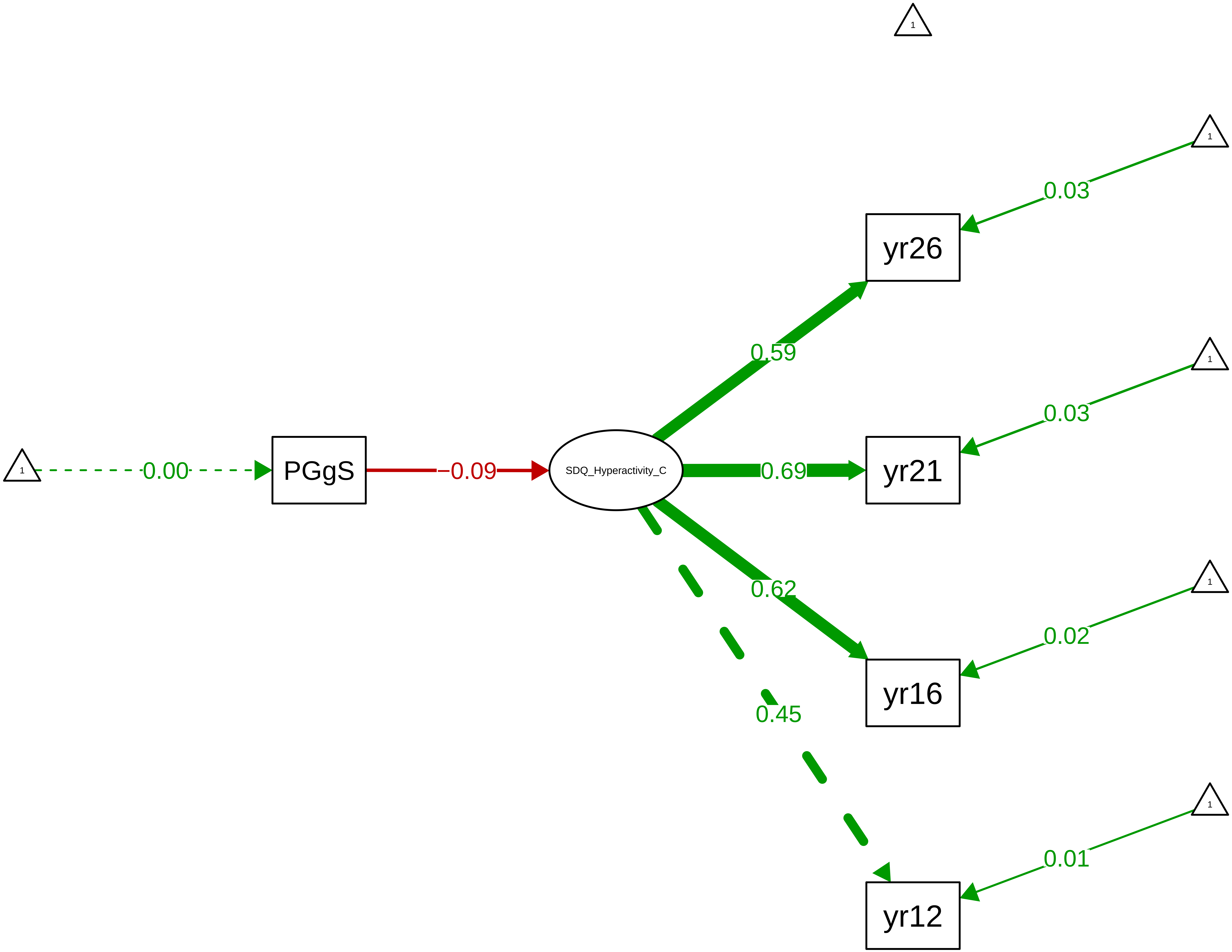

**Figure S9_56: SDQ Peer Problems − CTCR Overall**

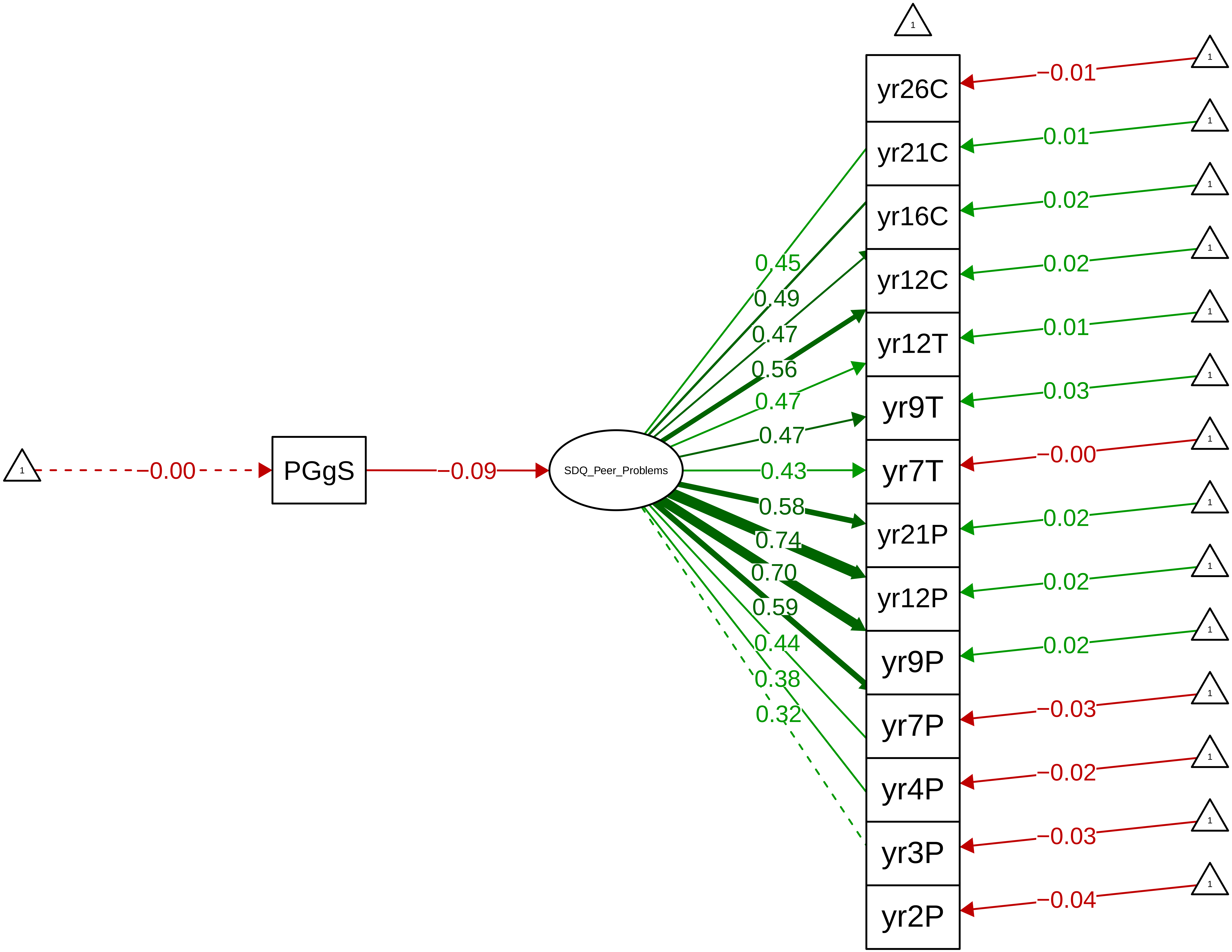

**Figure S9_57: SDQ Peer Problems − CTCR Stage**

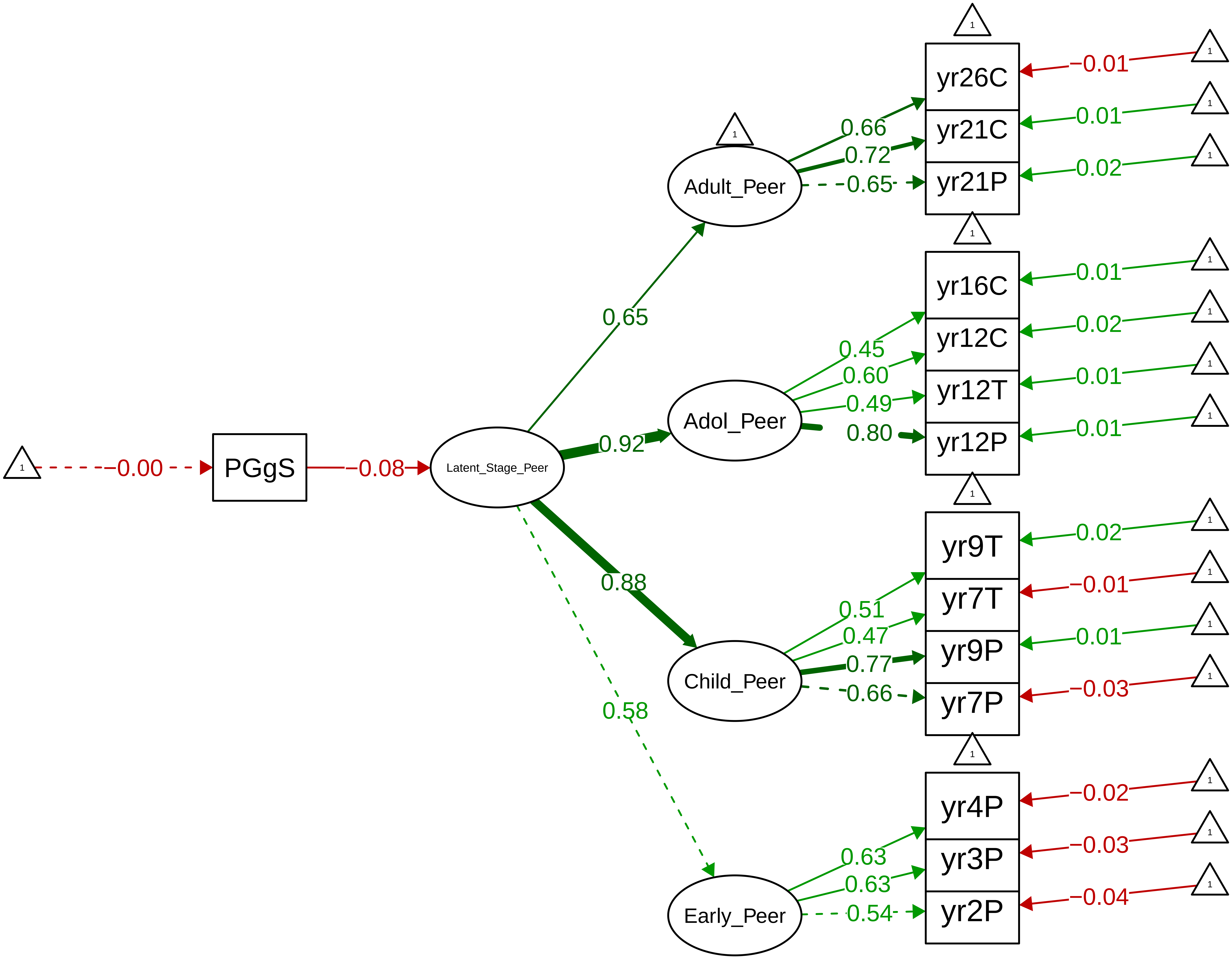

Figure S9_58: SDQ Peer Problems – Parent Overall

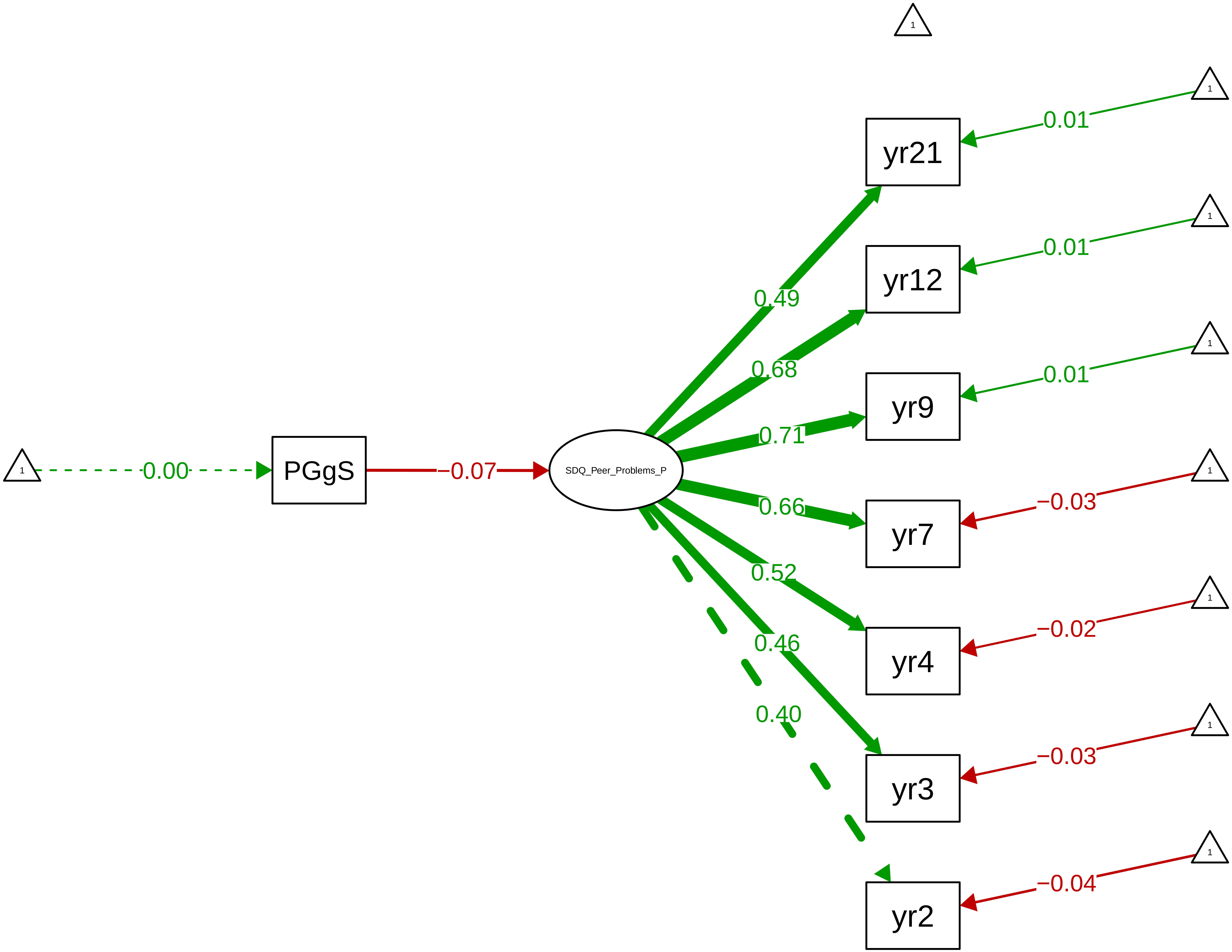

**Figure S9_59: SDQ Peer Problems − Parent Stage**

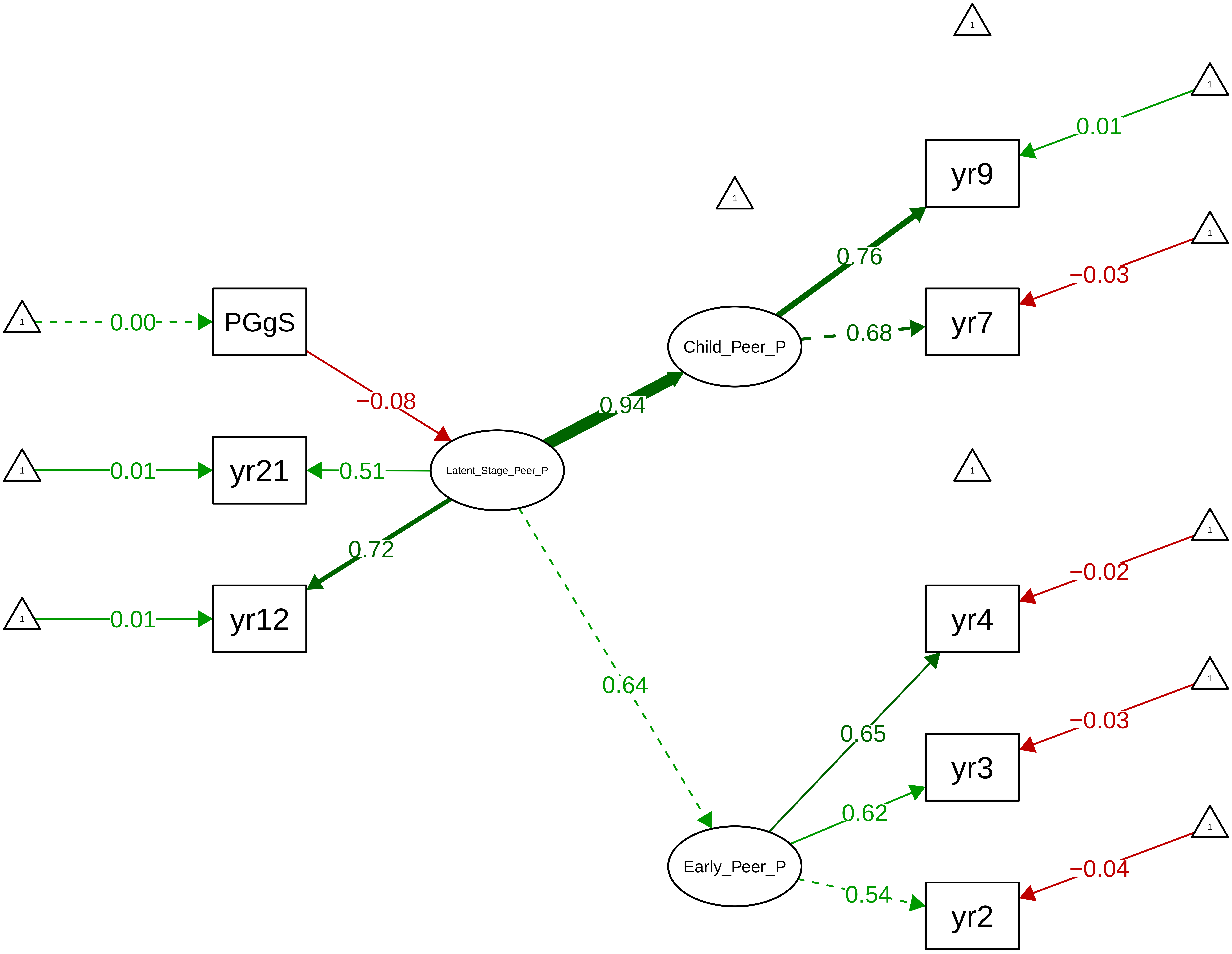

#### Figure S9_60: SDQ Peer Problems − Teacher Overall

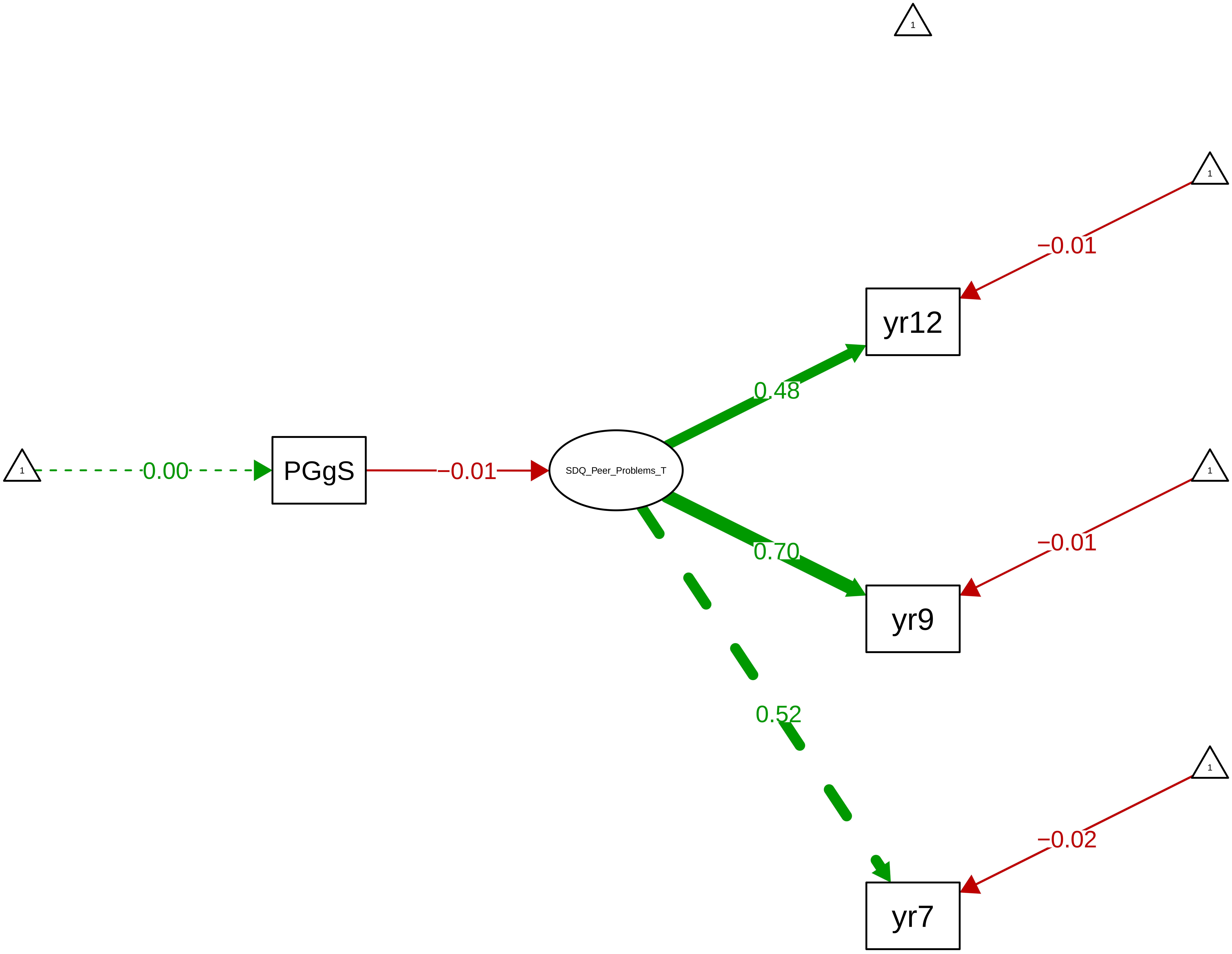

Figure S9_61: SDQ Peer Problems − Child Overall

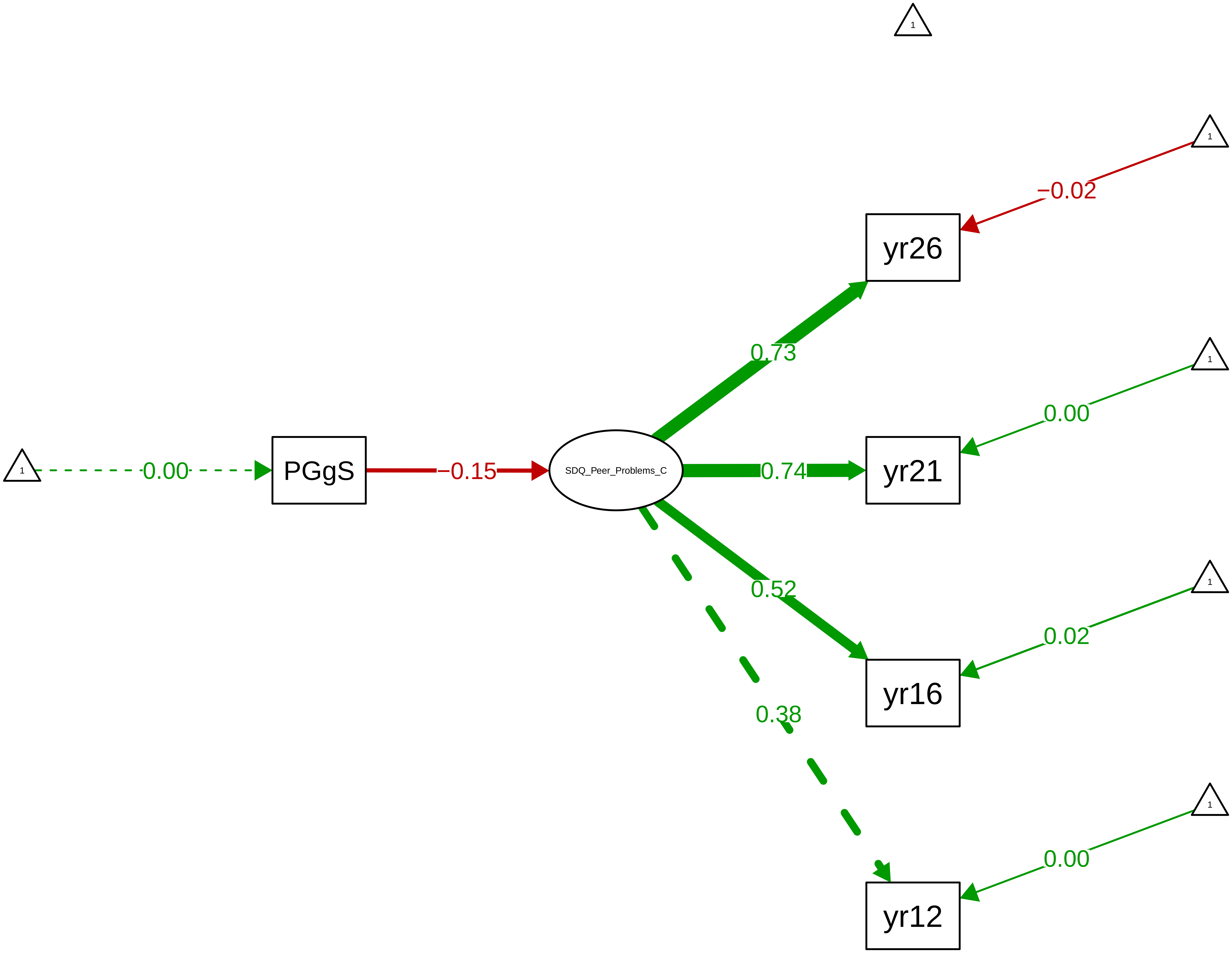

**Figure S9_62: SDQ Prosocial − CTCR Overall**

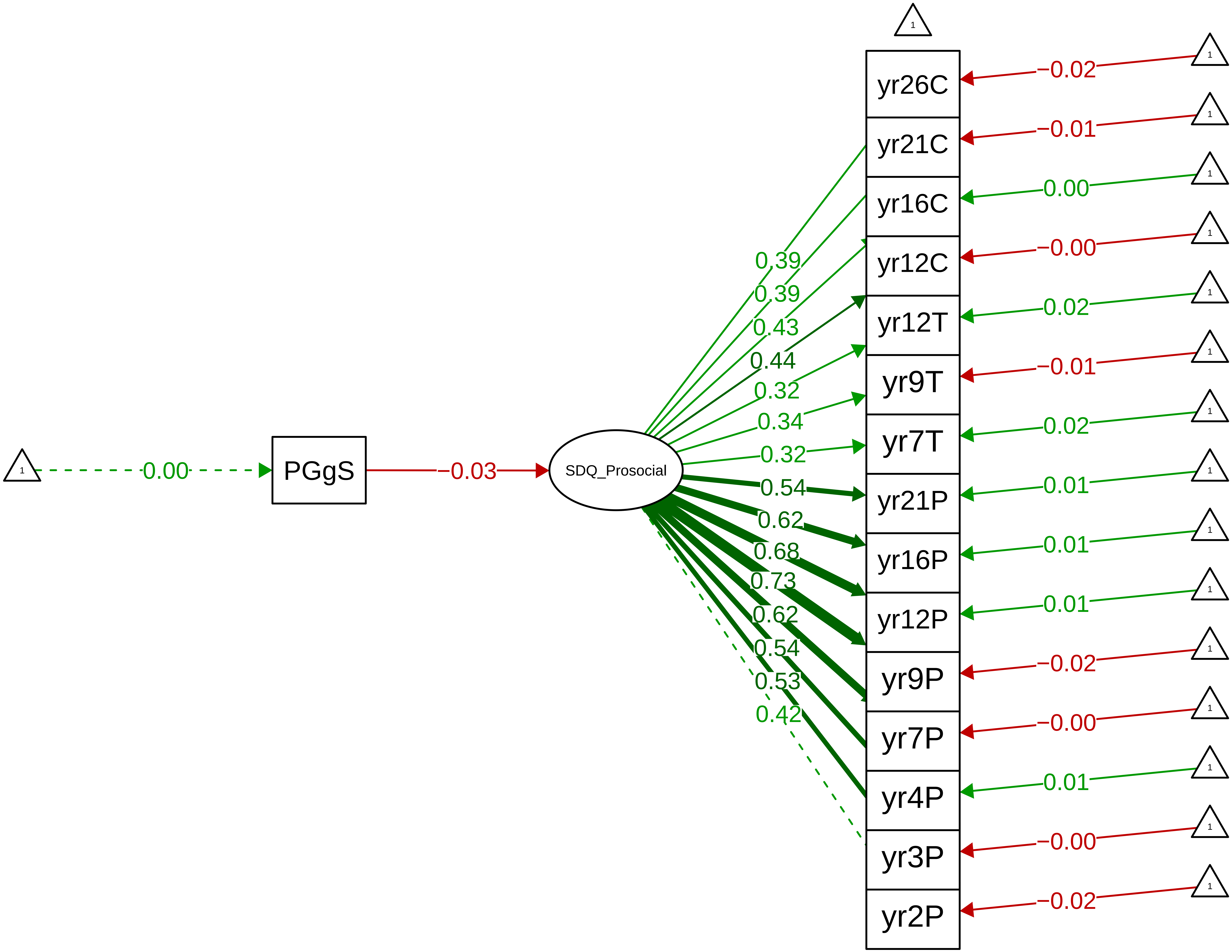

**Figure S9_63: SDQ Prosocial − CTCR Stage**

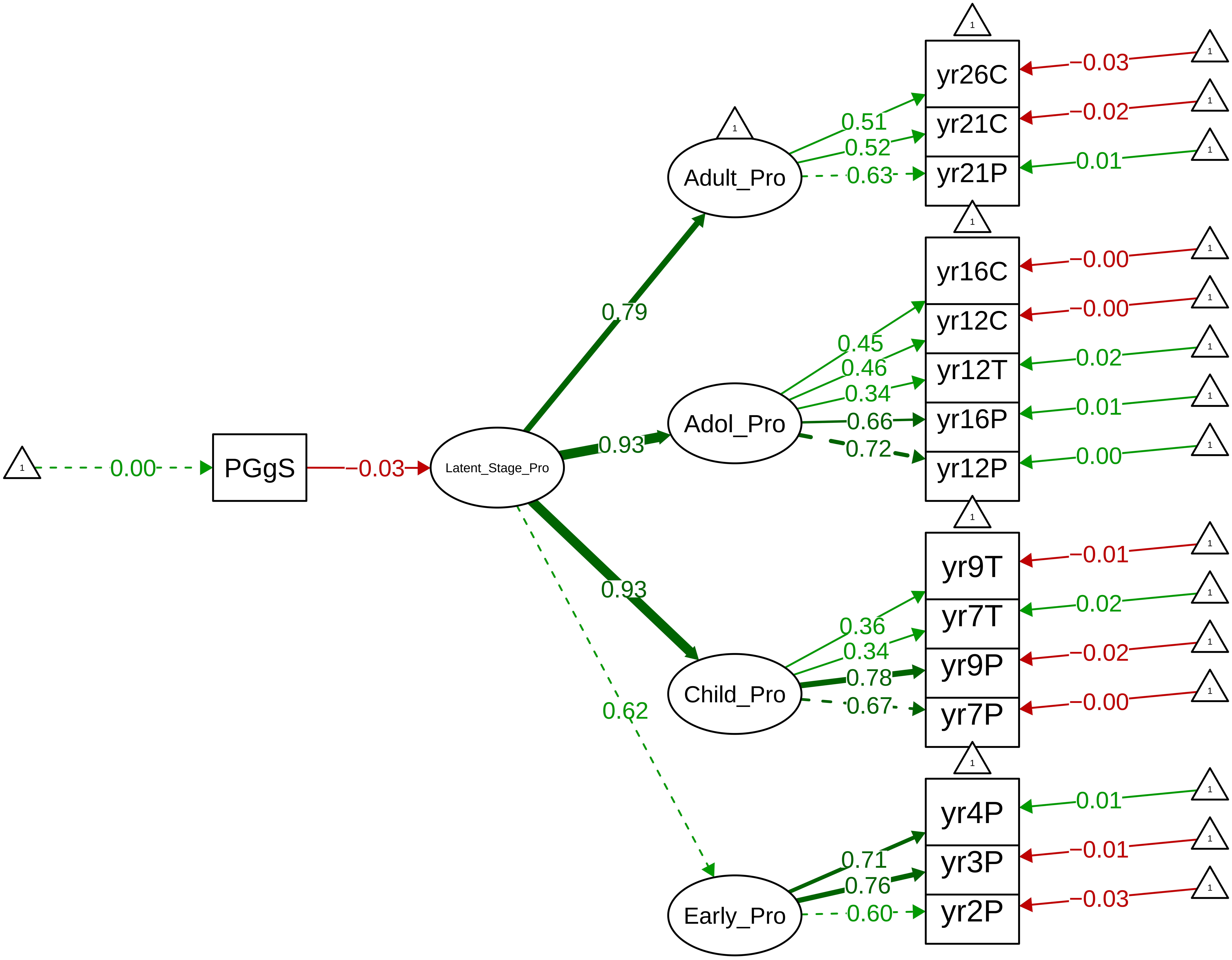

Figure S9_64: SDQ Prosocial − Parent Overall

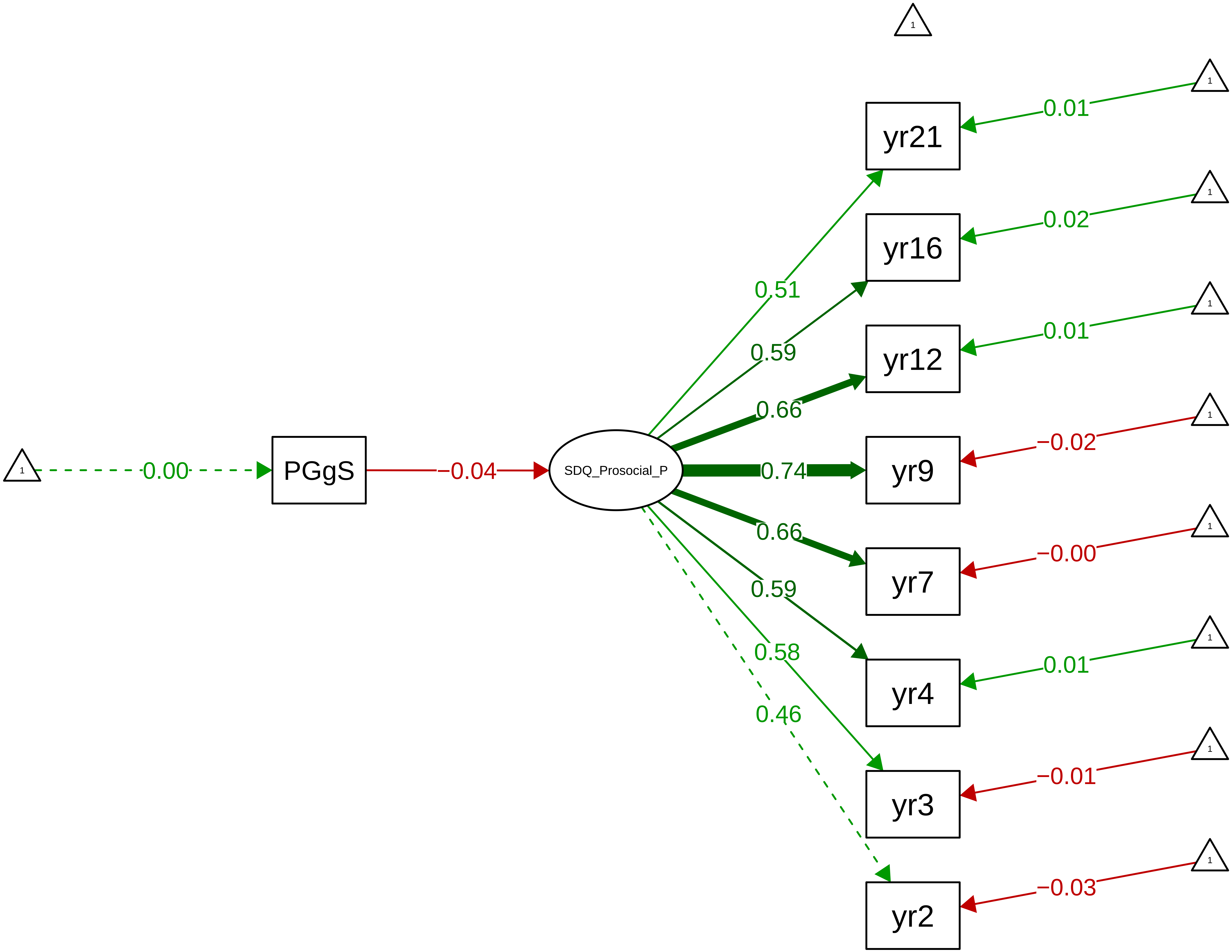

**Figure S9_65: SDQ Prosocial − Parent Stage**

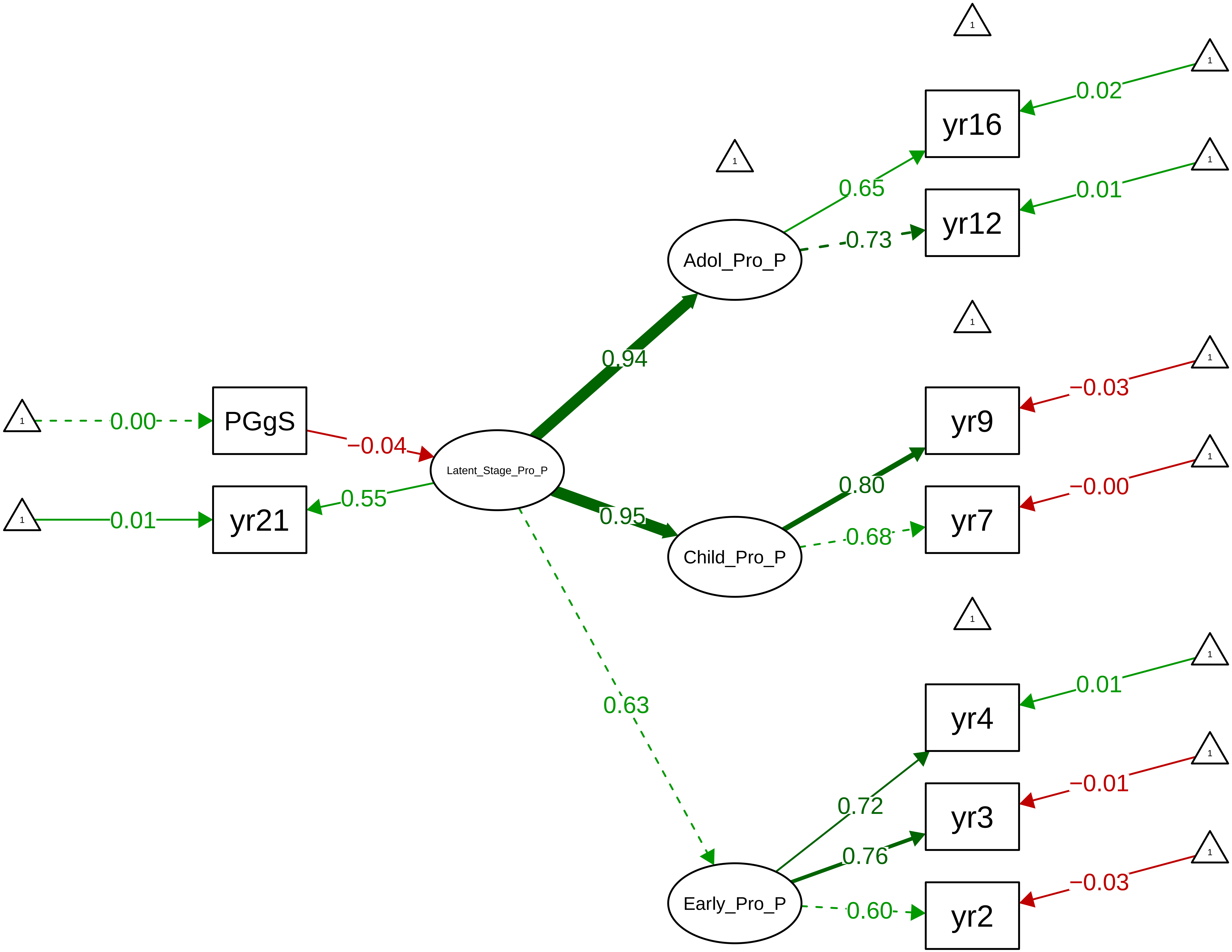

**Figure S9_66: SDQ Prosocial − Teacher Overall**

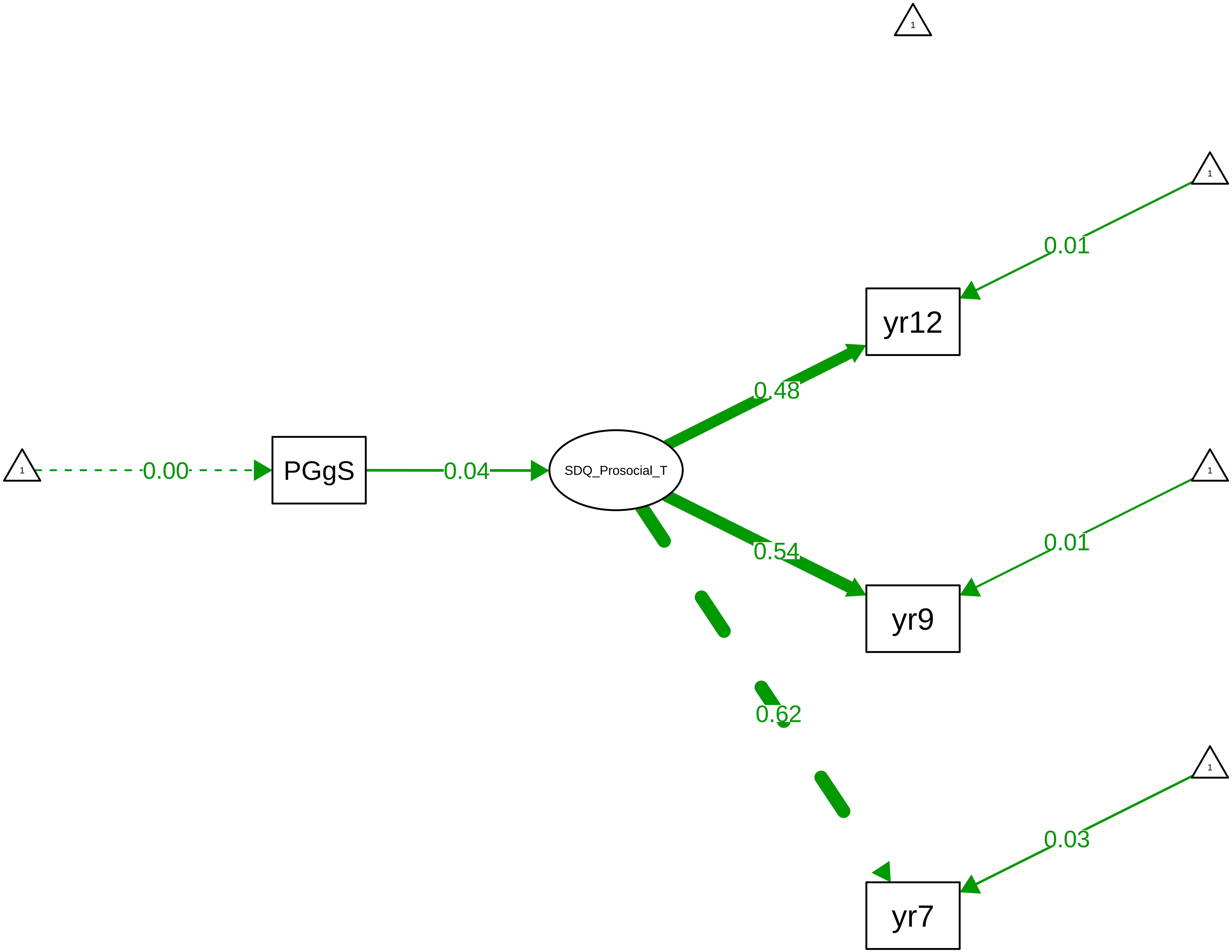

Figure S9_67: SDQ Prosocial − Child Overall

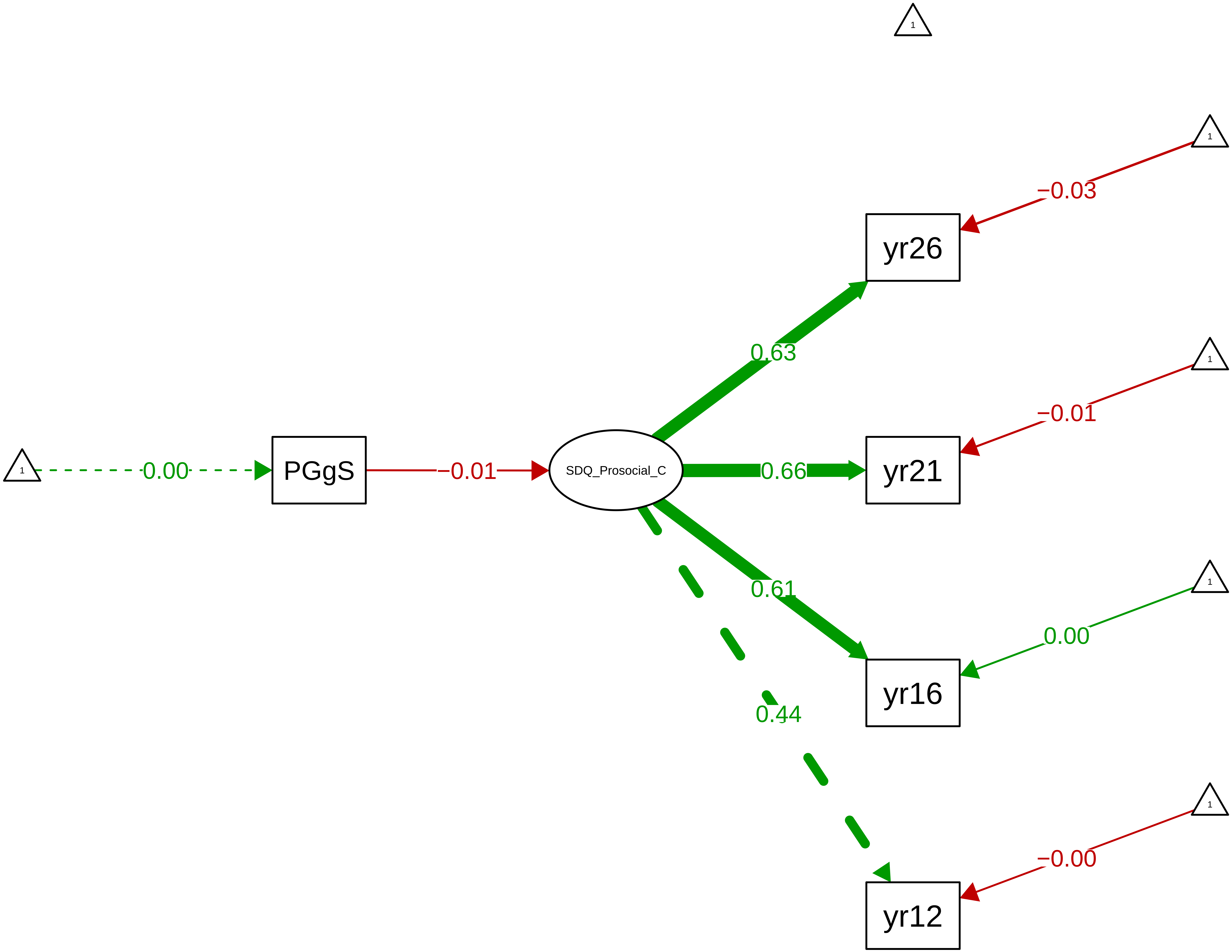

Figure S9_68: SDQ Total Problems − CTCR Overall

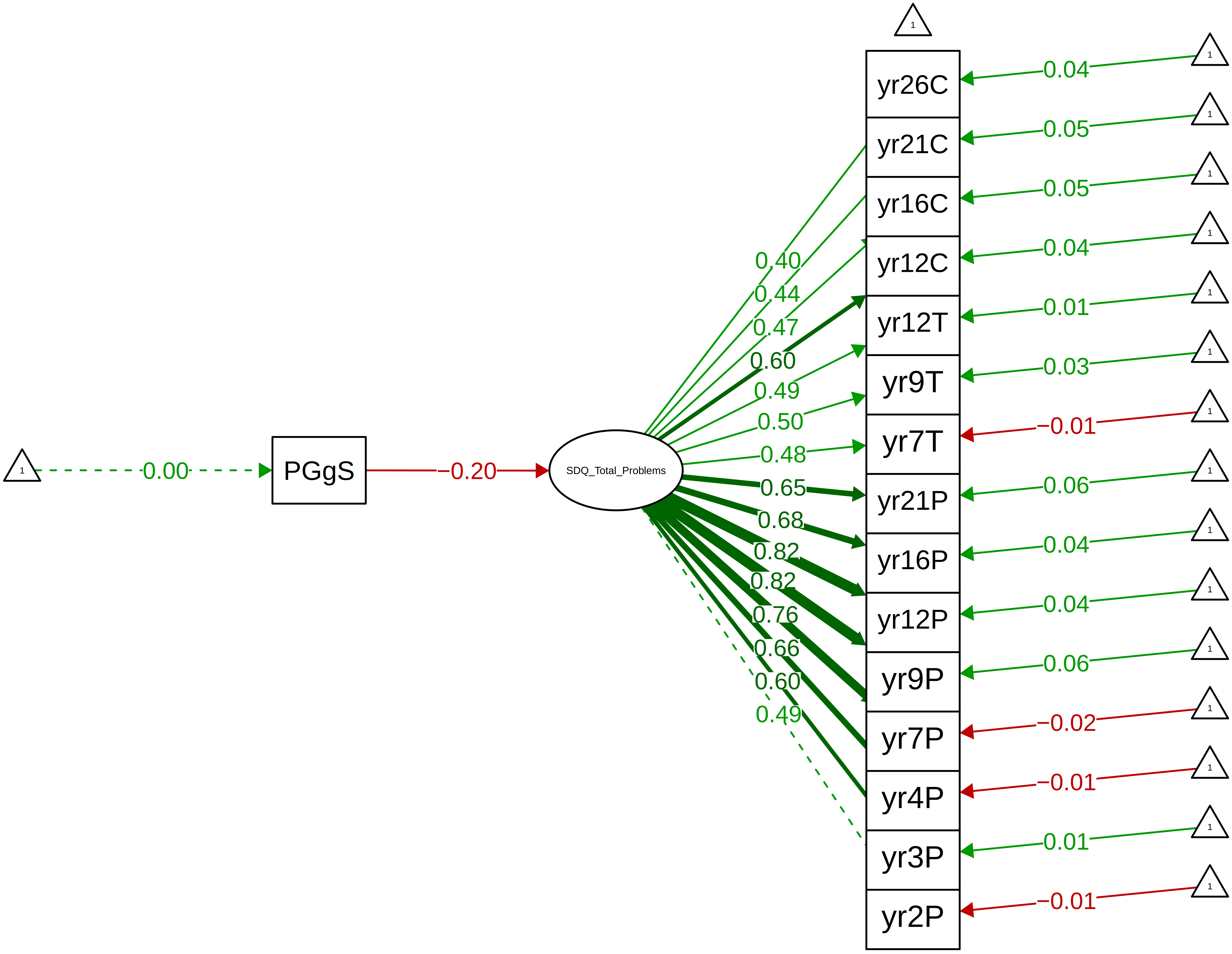

**Figure S9_69: SDQ Total Problems − CTCR Stage**

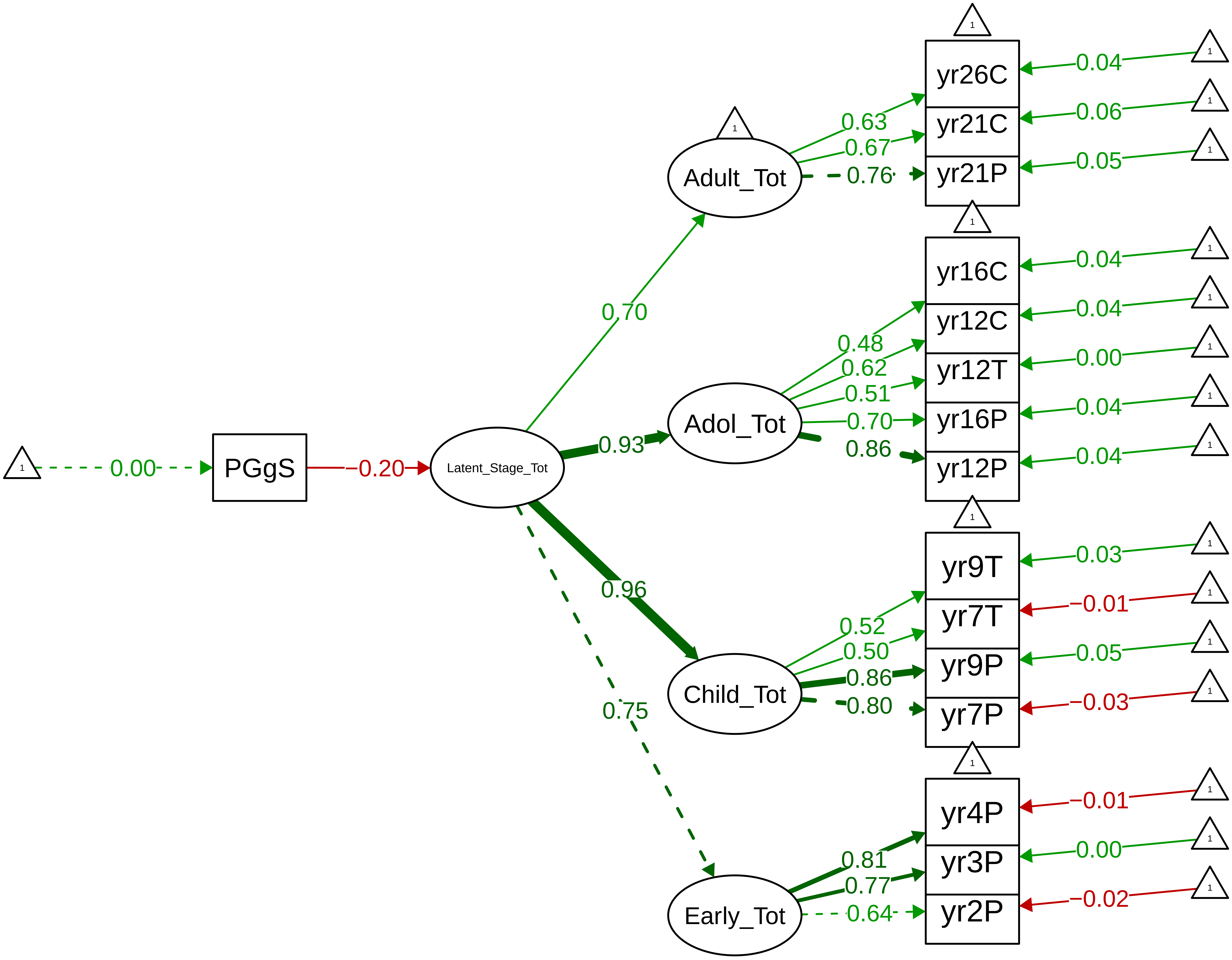

#### Figure S9_70: SDQ Total Problems − Parent Overall

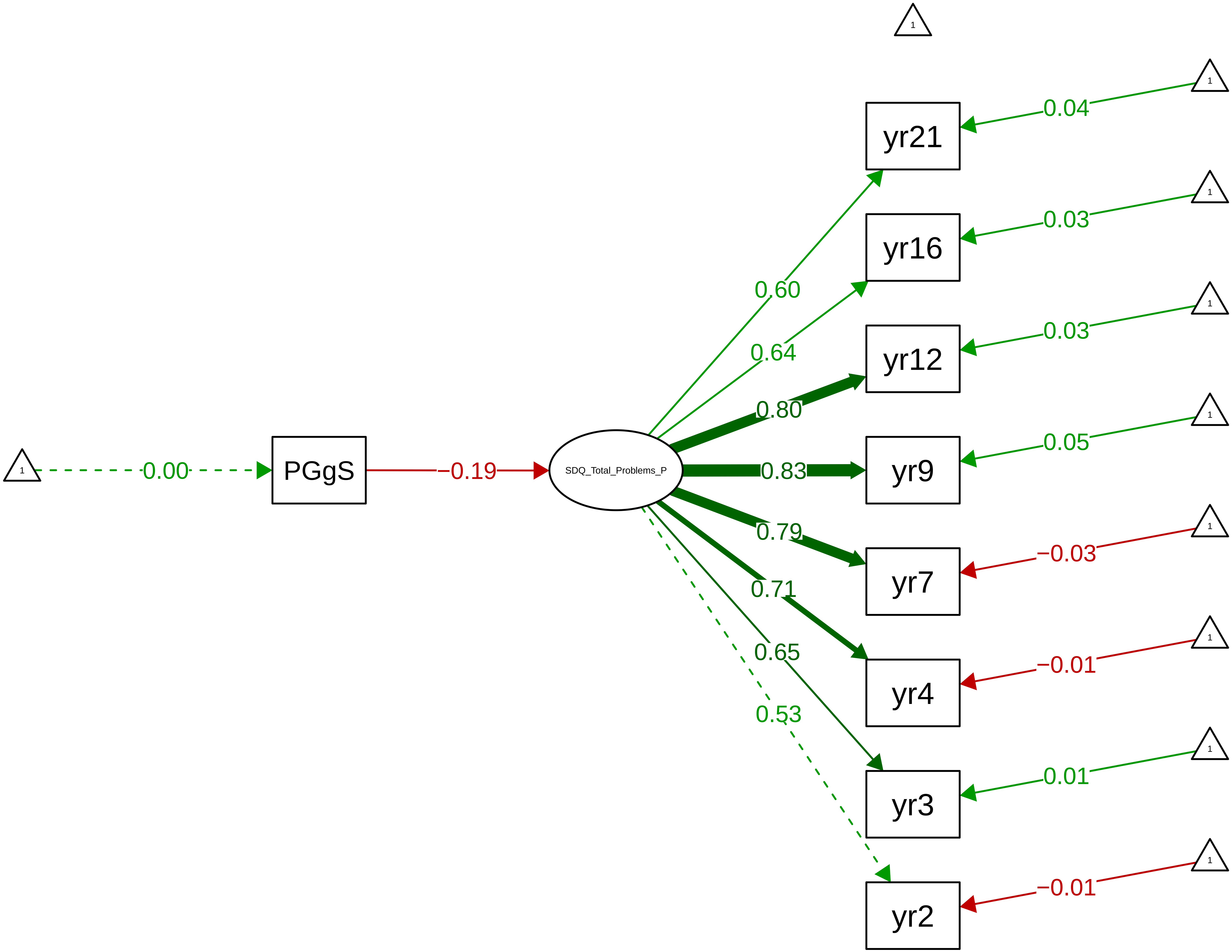

**Figure S9_71: SDQ Total Problems − Parent Stage**

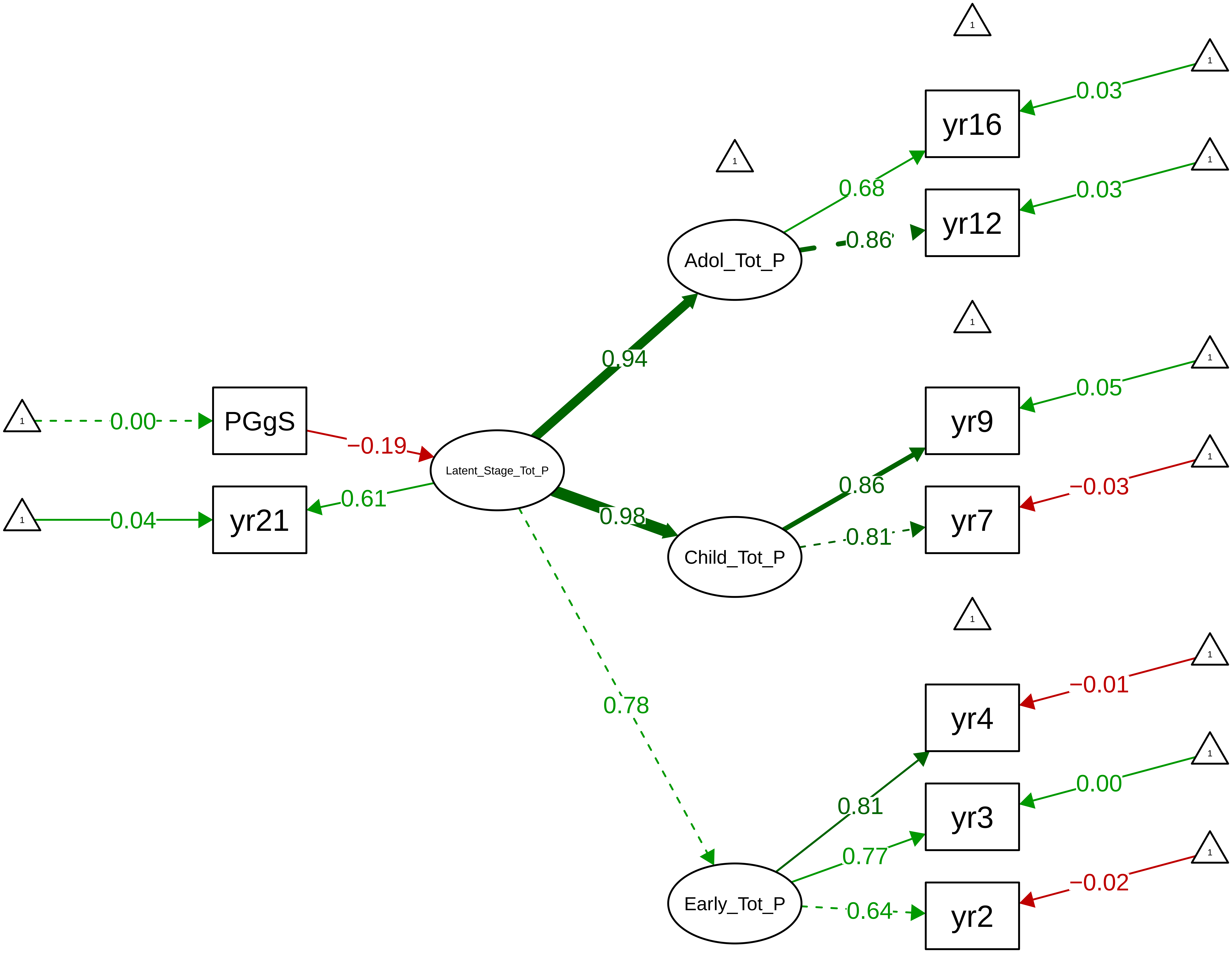

**Figure S9_72: SDQ Total Problems − Teacher Overall**

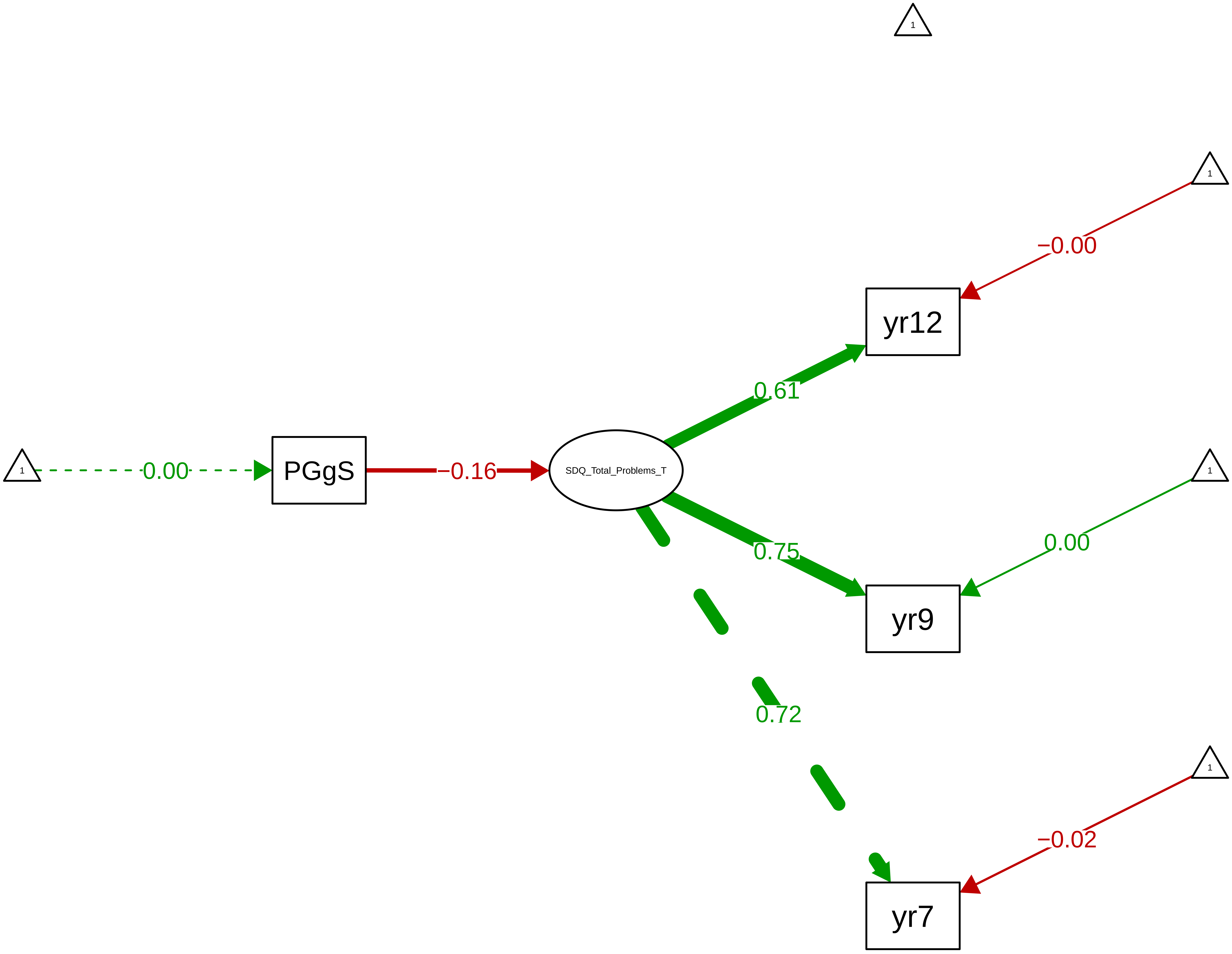

**Figure S9_73: SDQ Total Problems – Child Overall**

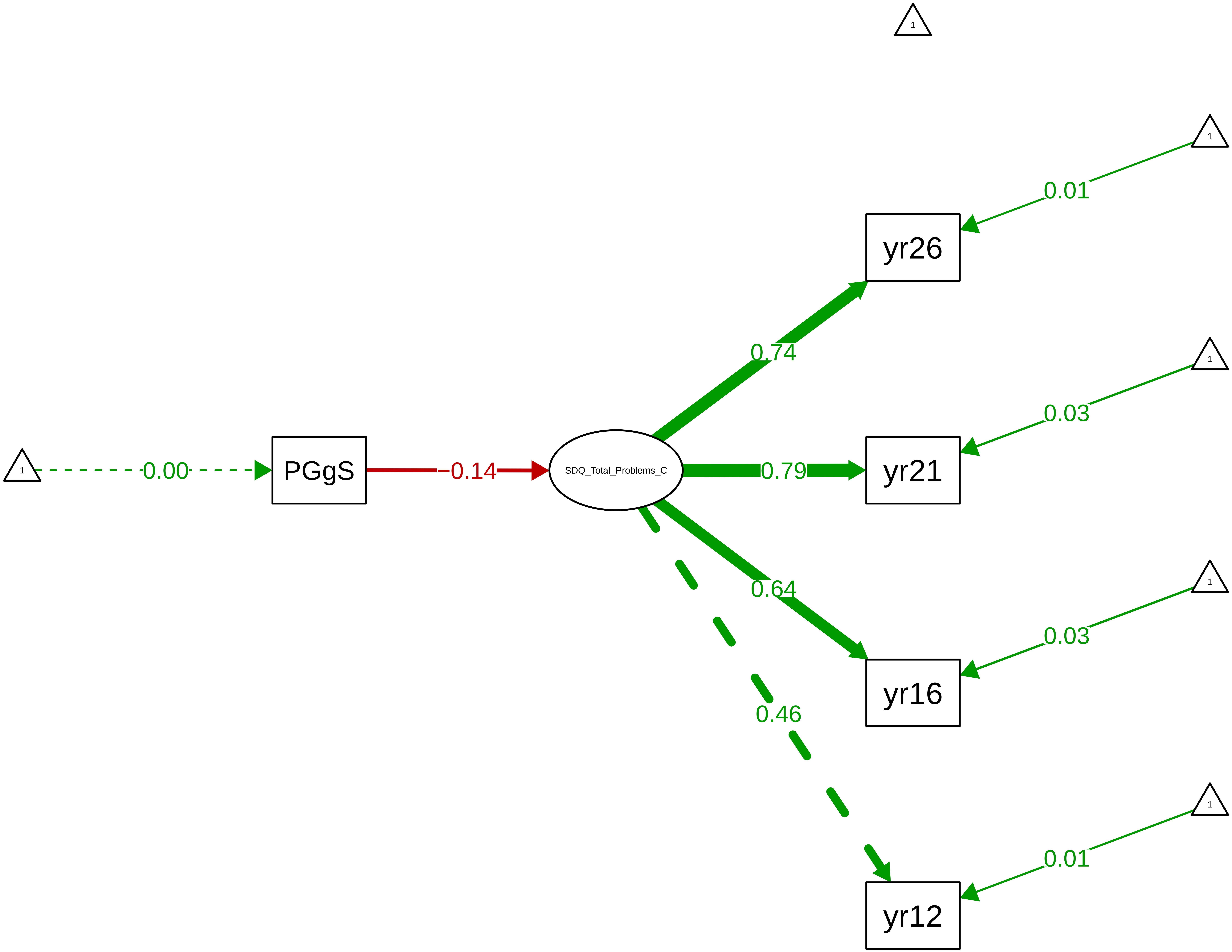

### Figure S10_1 Predicted Trajectory: General Cognitive Ability

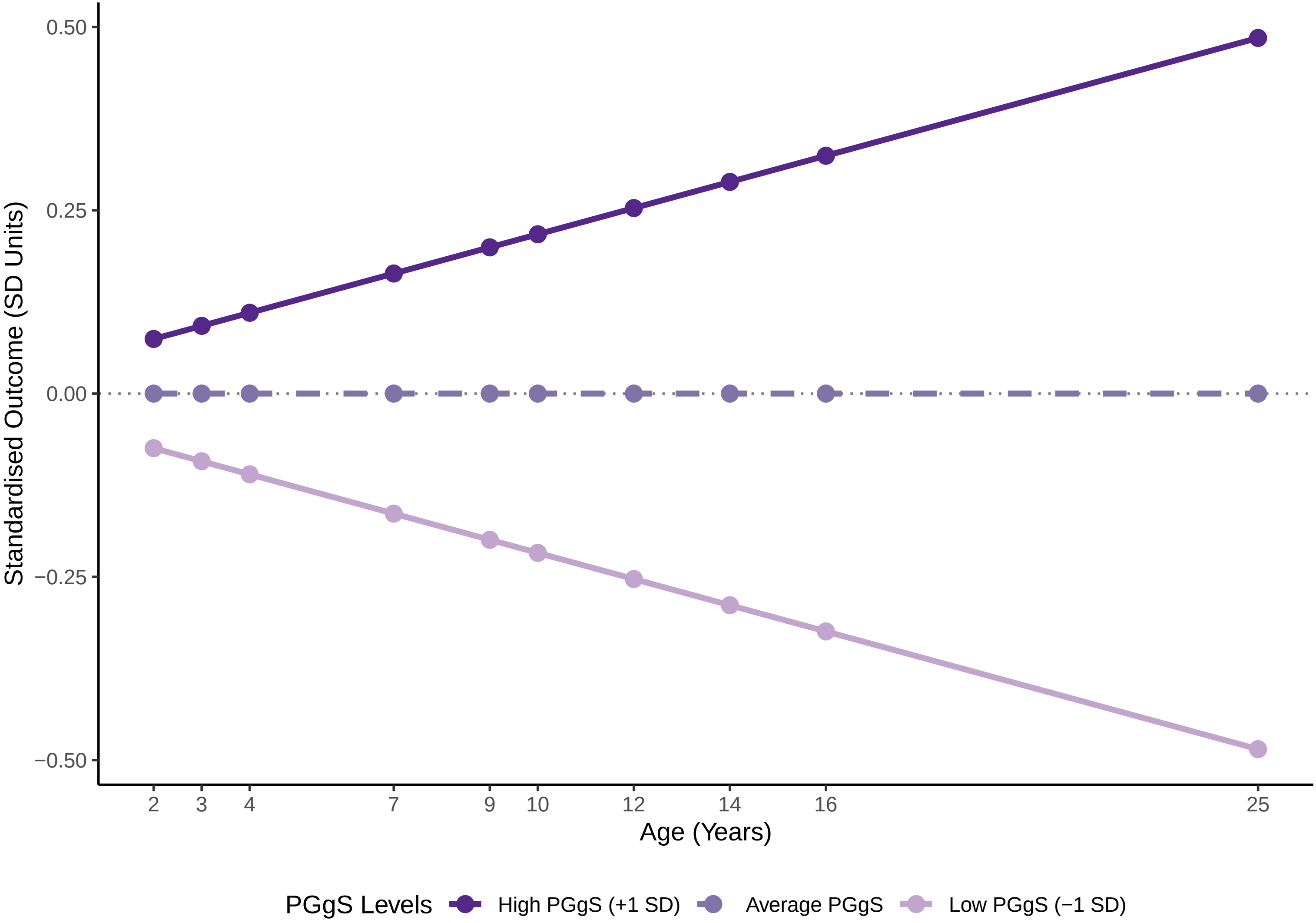

### Figure S10_2 Predicted Trajectory: Verbal Abilities

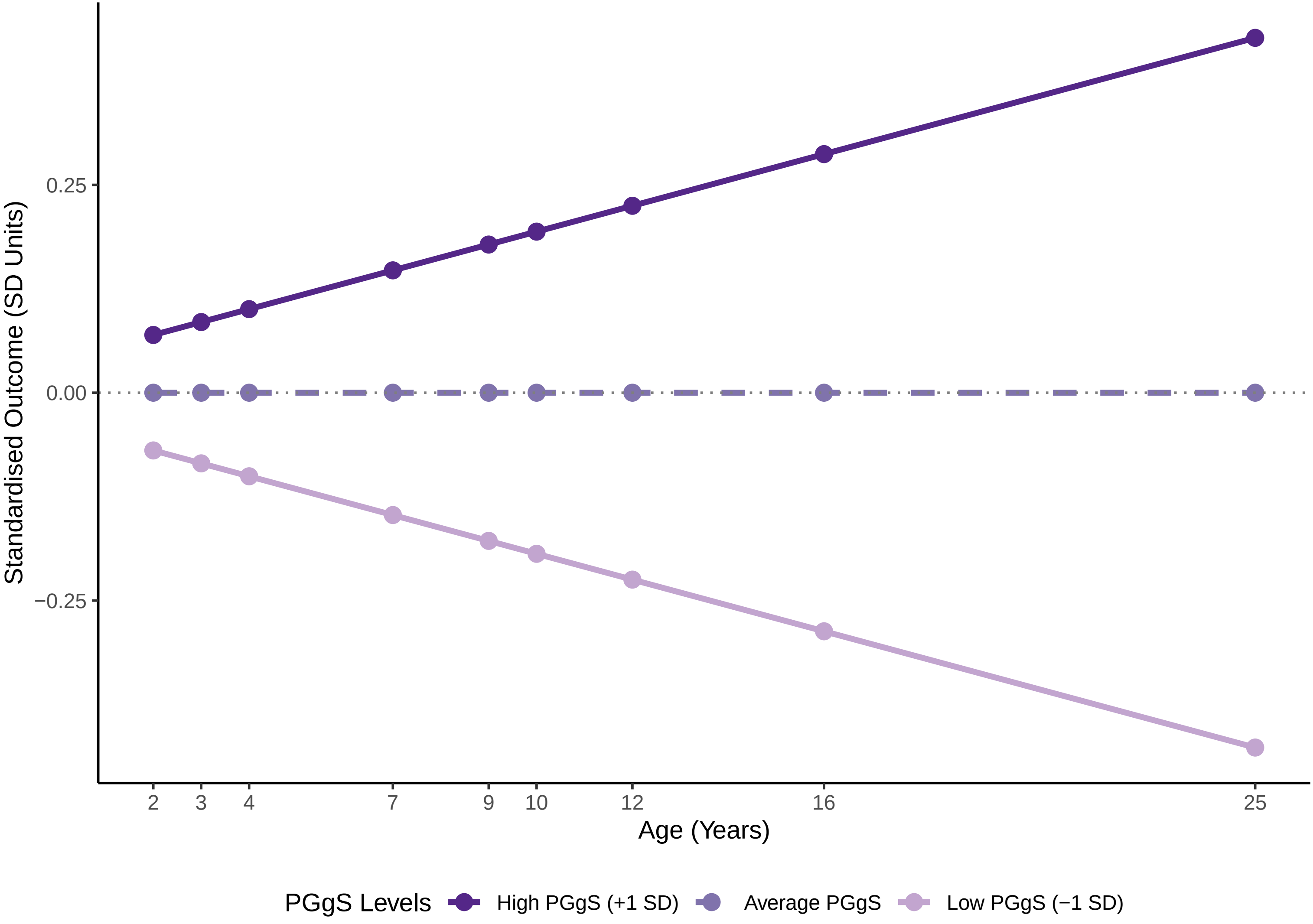

Figure S10_3 Predicted Trajectory: Nonverbal Abilities

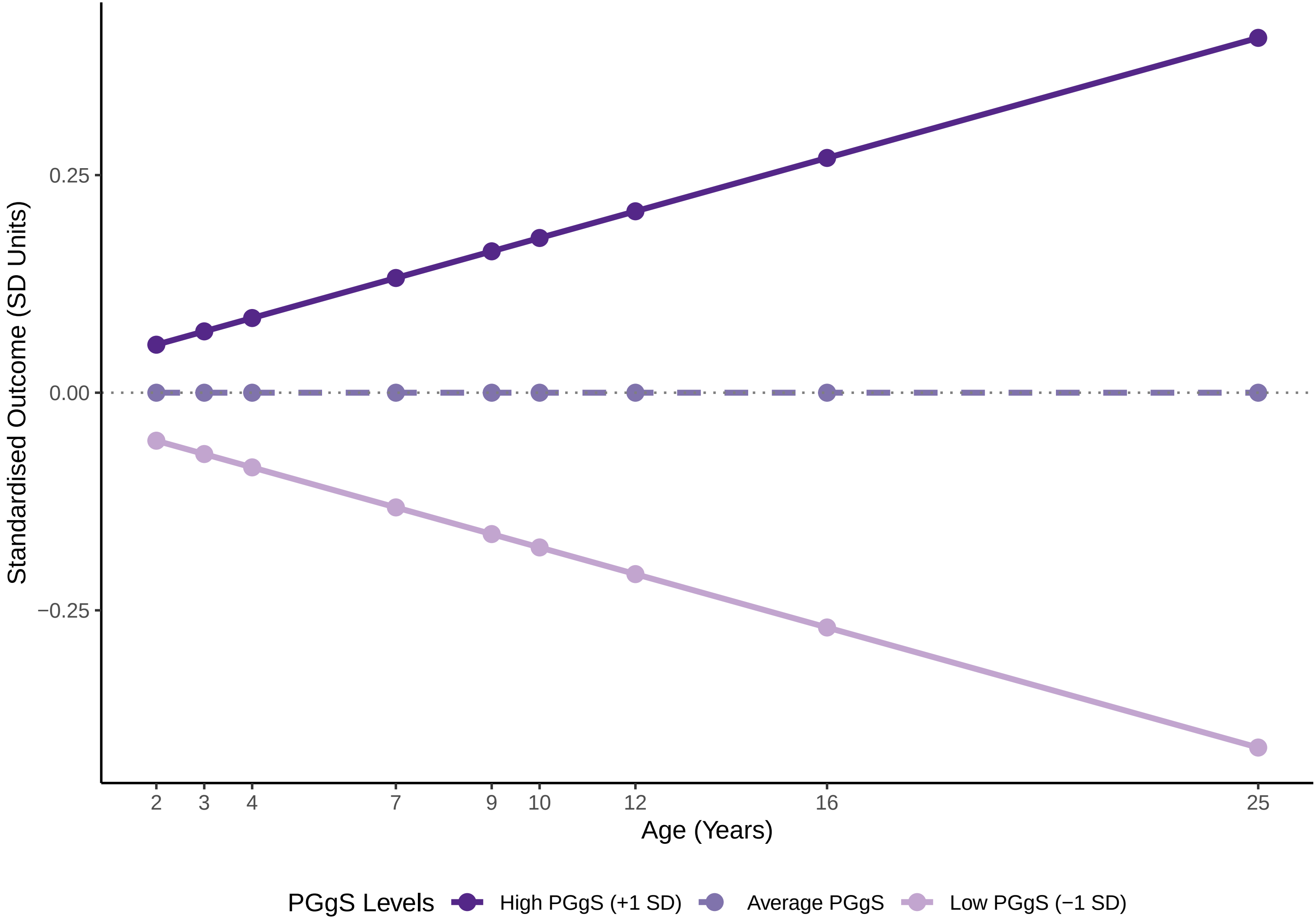

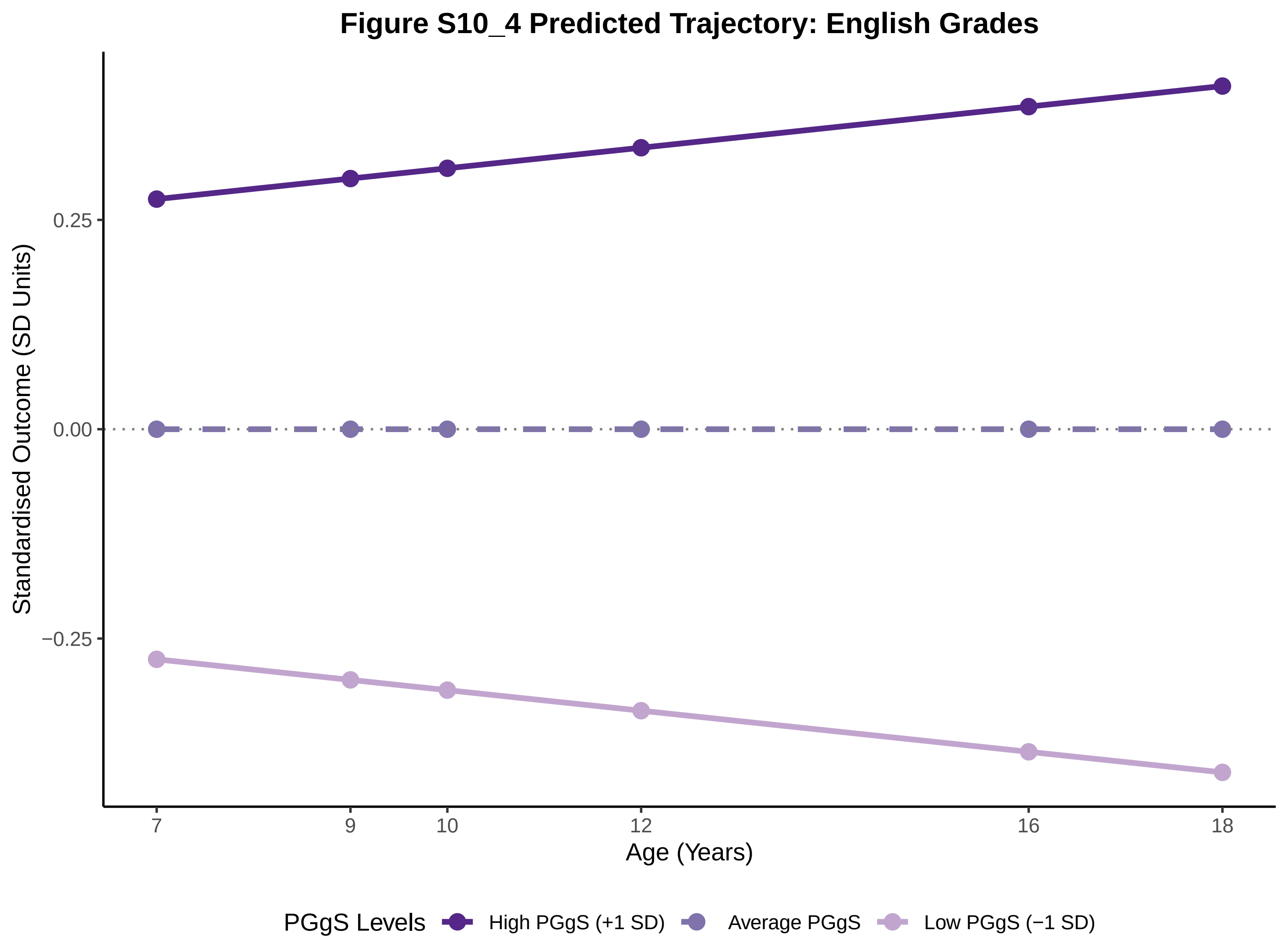

Figure S10_4 Predicted Trajectory: English Grades
Standardised Outcome (SD Units)
0.25
0.00
−0.25
7
9
10
12
16
18
Age (Years)
PGgS Levels
High PGgS (+1 SD)
Average PGgS
Low PGgS (−1 SD)

### Figure S10_5 Predicted Trajectory: Maths Grades

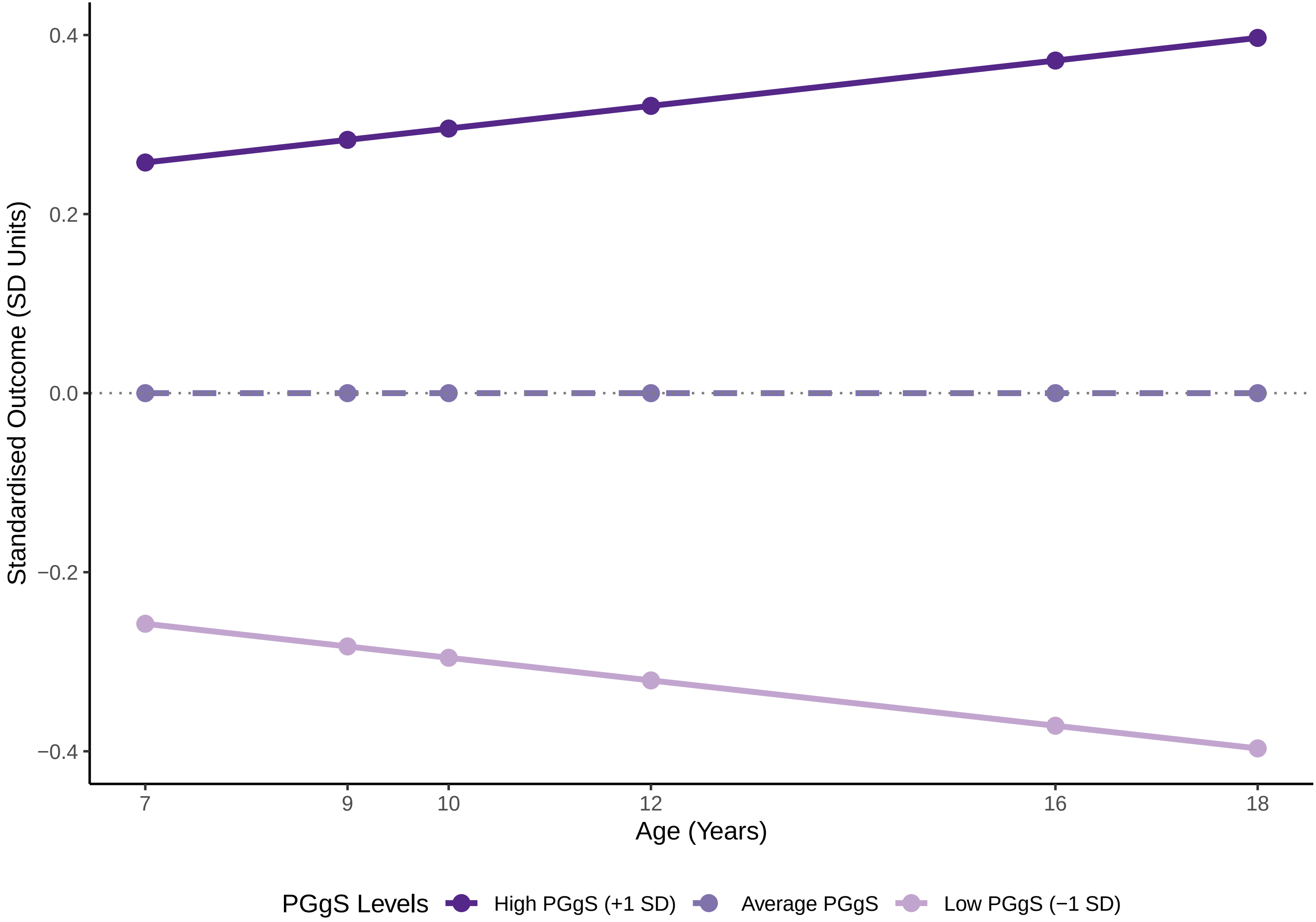

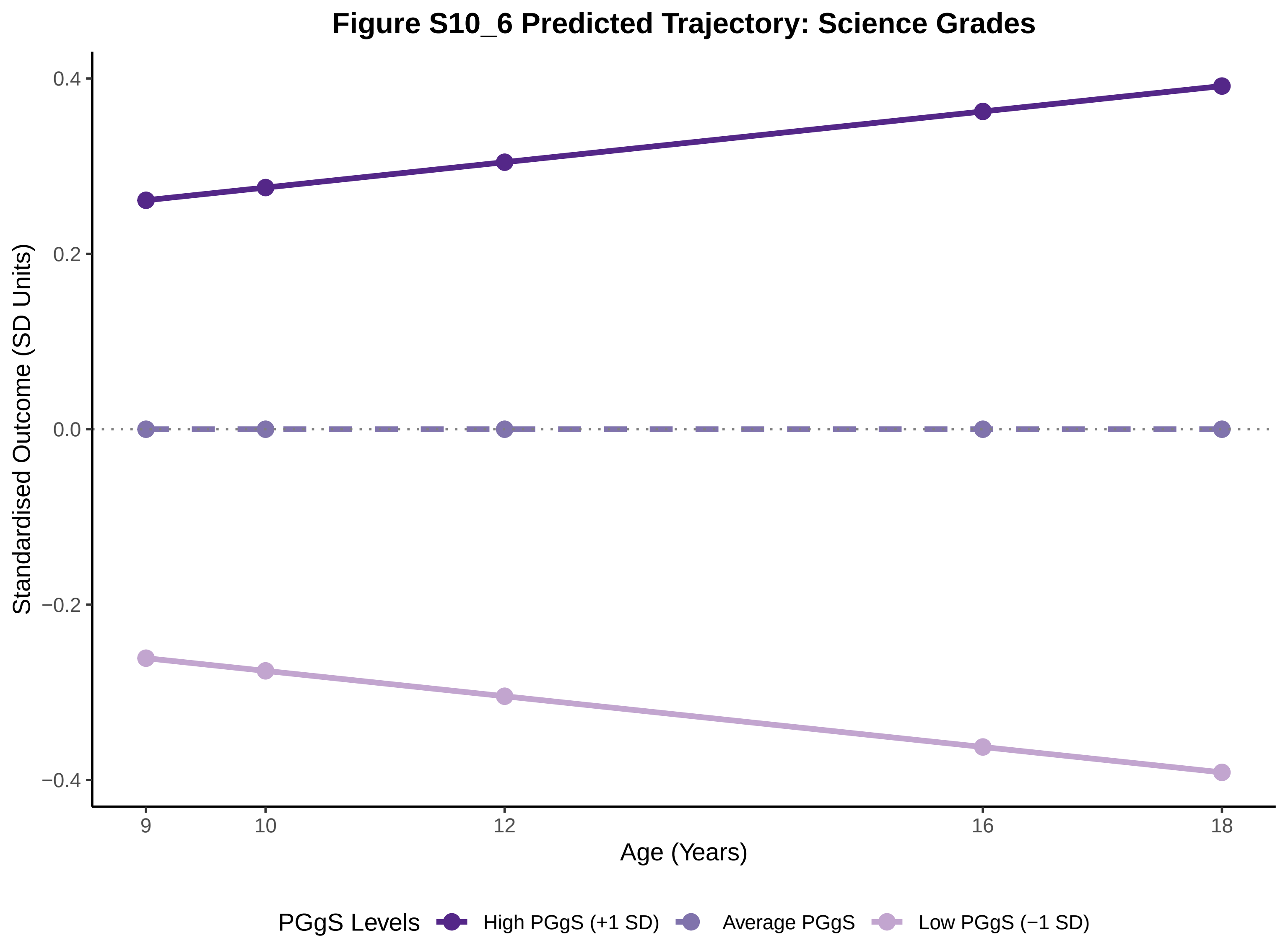
Figure S10_6 Predicted Trajectory: Science Grades
Standardised Outcome (SD Units)
0.4
0.2
0.0
−0.2
−0.4
9
10
12
16
18
Age (Years)
PGgS Levels
High PGgS (+1 SD)
Average PGgS
Low PGgS (−1 SD)

Figure S10_7 Predicted Trajectory: Core Subject Grades

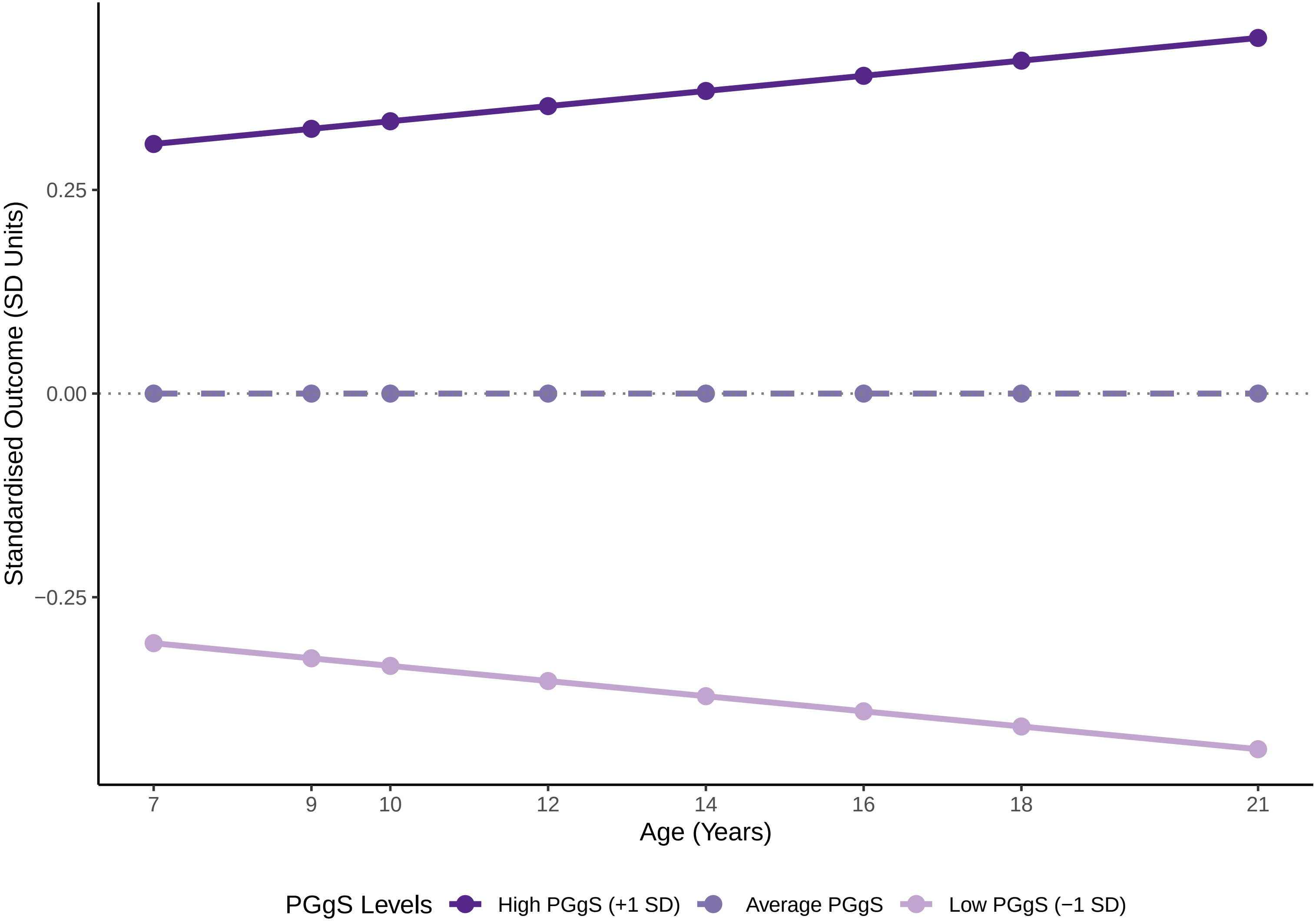

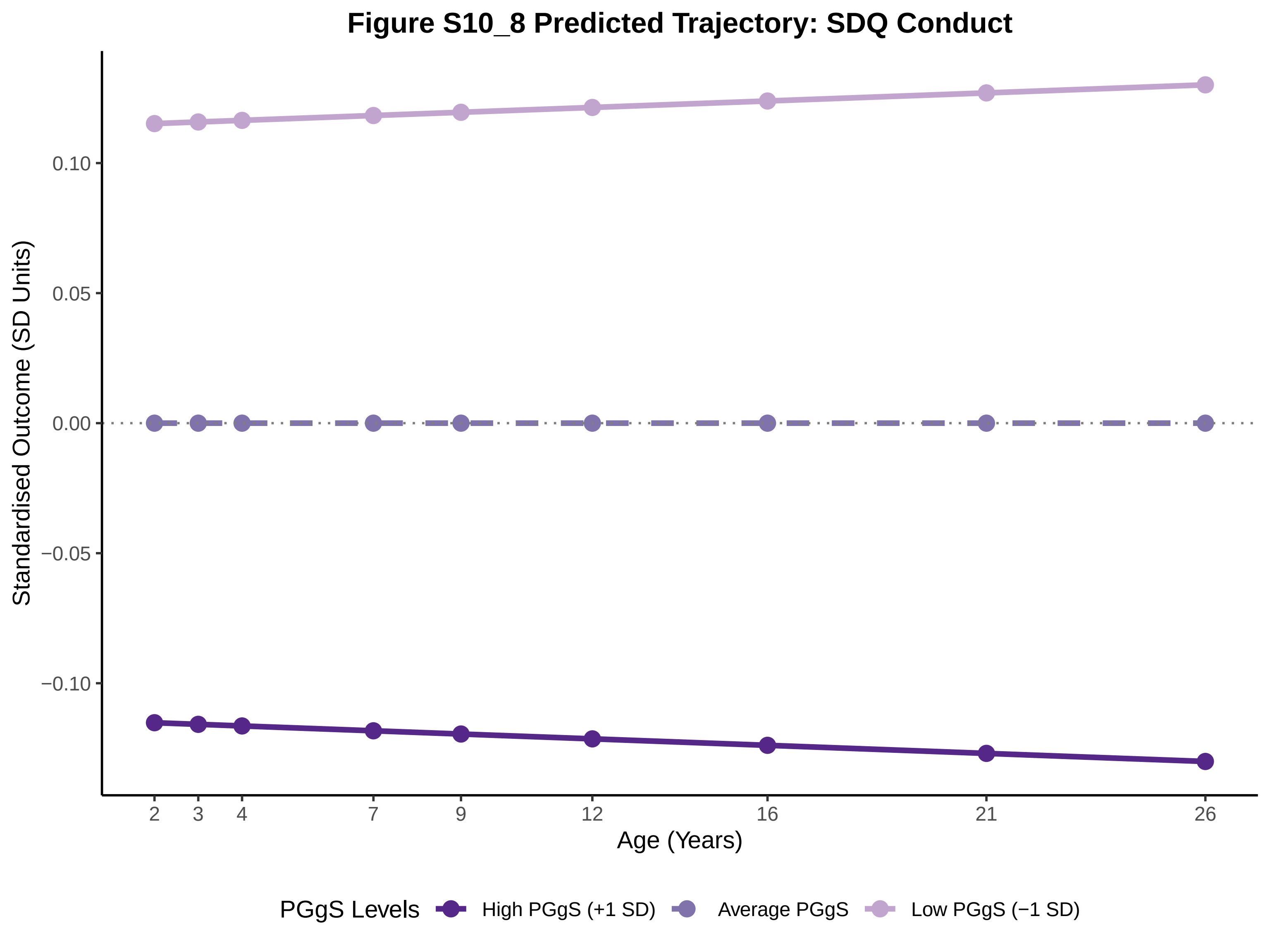
Figure S10_8 Predicted Trajectory: SDQ Conduct
Standardised Outcome (SD Units)
0.10
0.05
0.00
−0.05
−0.10
2
3
4
7
9
12
16
21
26
Age (Years)
PGgS Levels
High PGgS (+1 SD)
Average PGgS
Low PGgS (−1 SD)

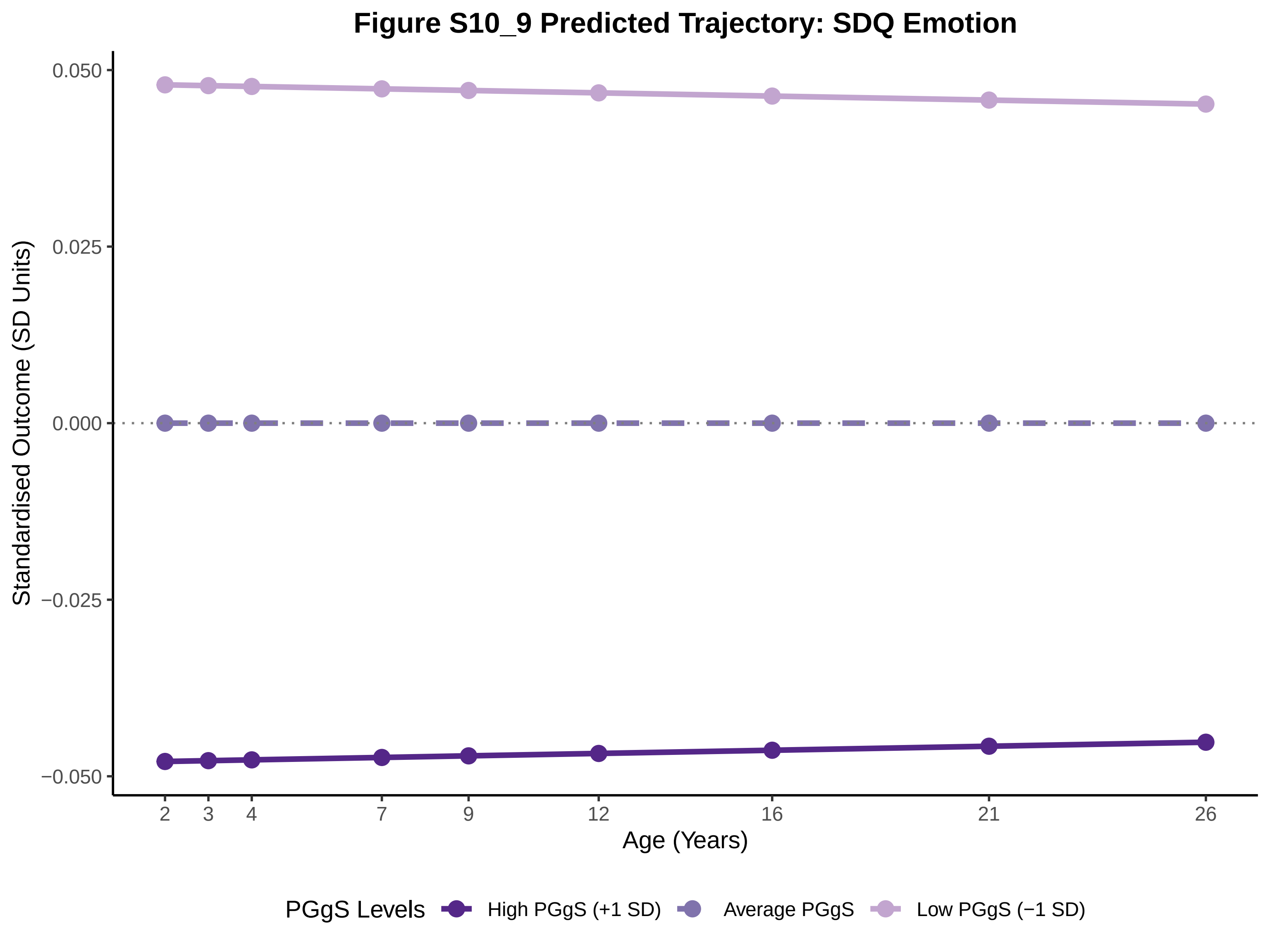
Figure S10_9 Predicted Trajectory: SDQ Emotion
Standardised Outcome (SD Units)
0.050
0.025
0.000
−0.025
−0.050
Age (Years)
2
3
4
7
9
12
16
21
26
PGgS Levels
High PGgS (+1 SD)
Average PGgS
Low PGgS (−1 SD)

**Figure S10_10 Predicted Trajectory: SDQ Hyperactivity_SDQ**

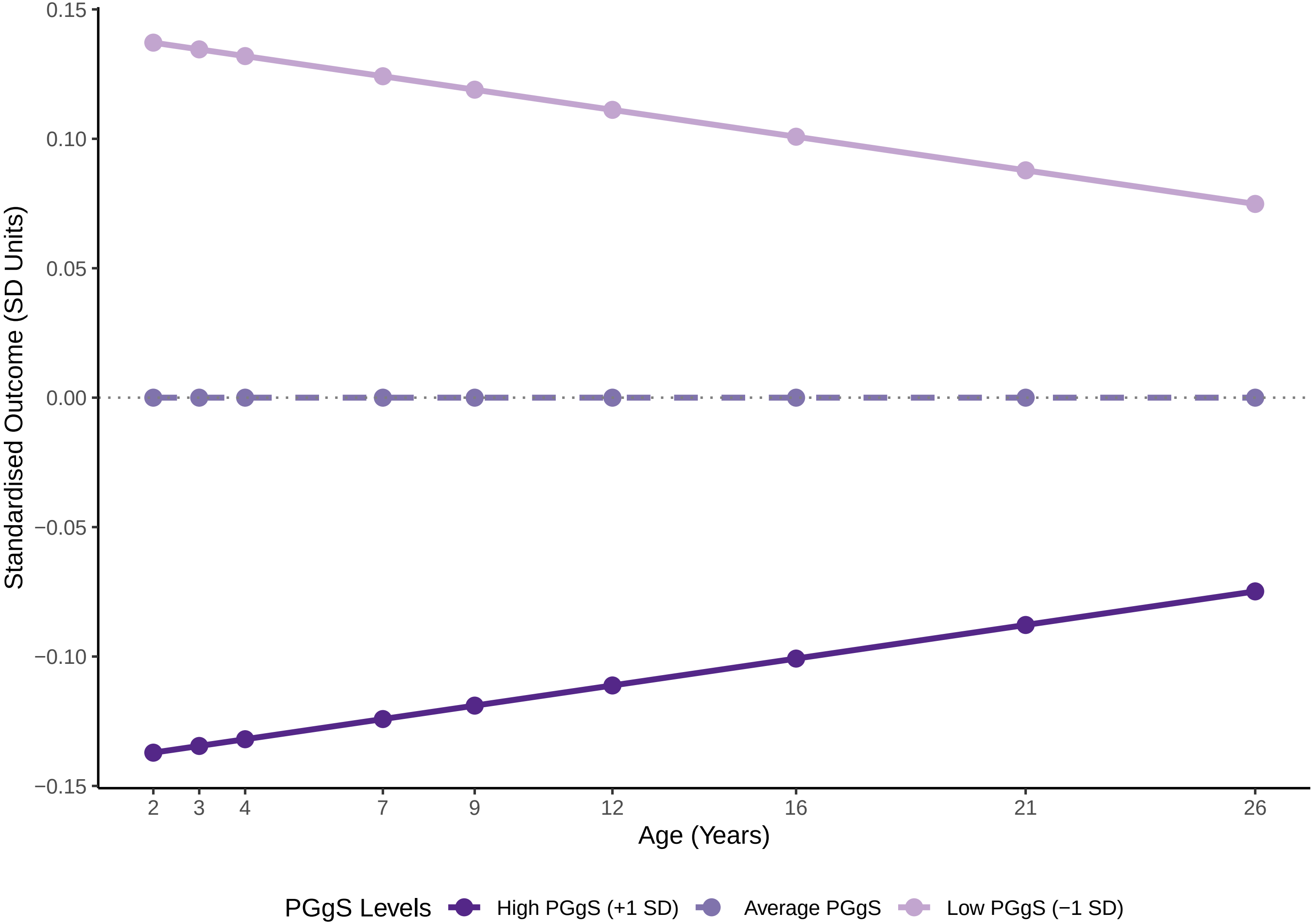

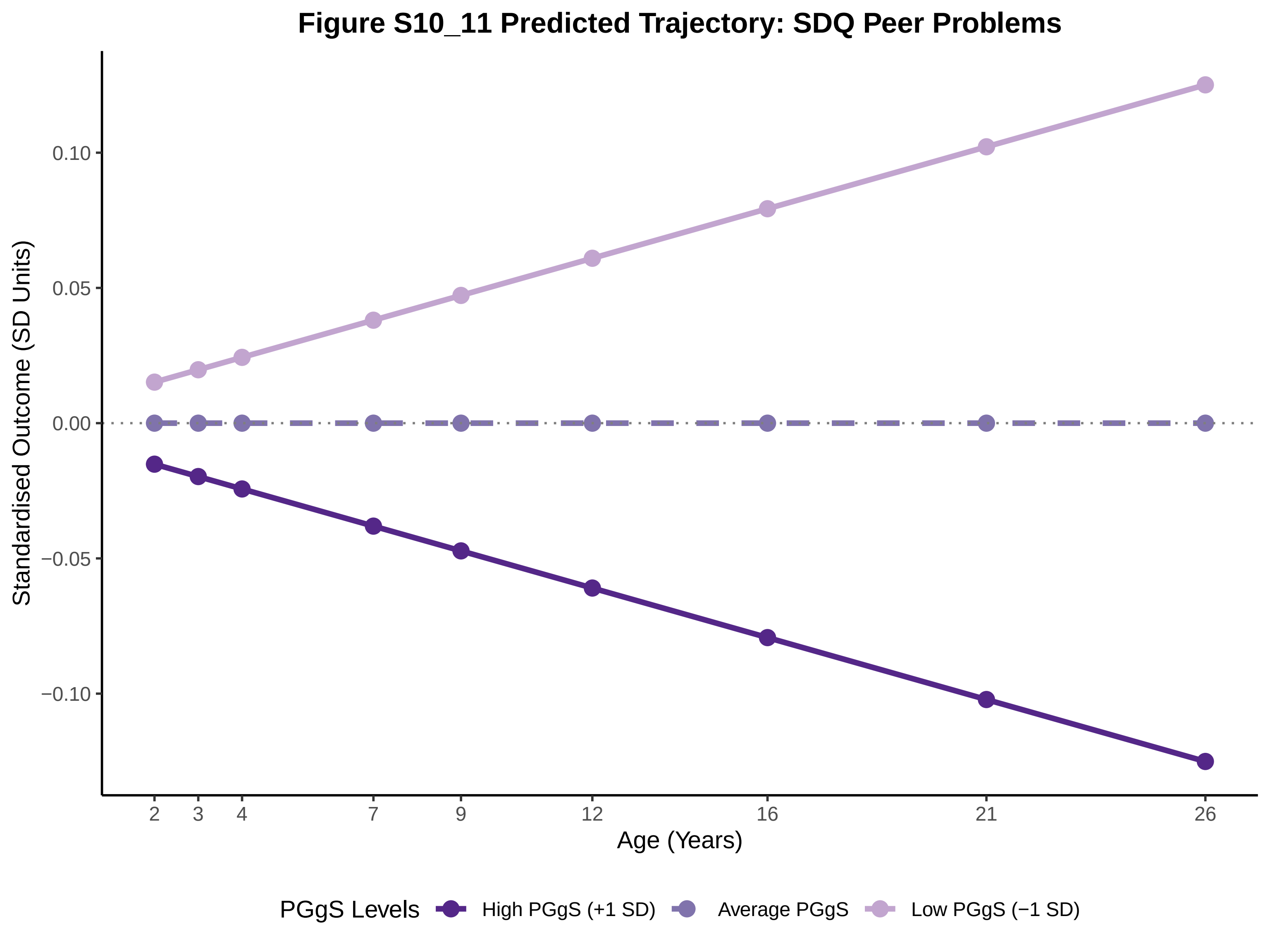
Figure S10_11 Predicted Trajectory: SDQ Peer Problems
Standardised Outcome (SD Units)
0.10
0.05
0.00
−0.05
−0.10
2
3
4
7
9
12
16
21
26
Age (Years)
PGgS Levels
High PGgS (+1 SD)
Average PGgS
Low PGgS (−1 SD)

### Figure S10_12 Predicted Trajectory: SDQ Prosocial

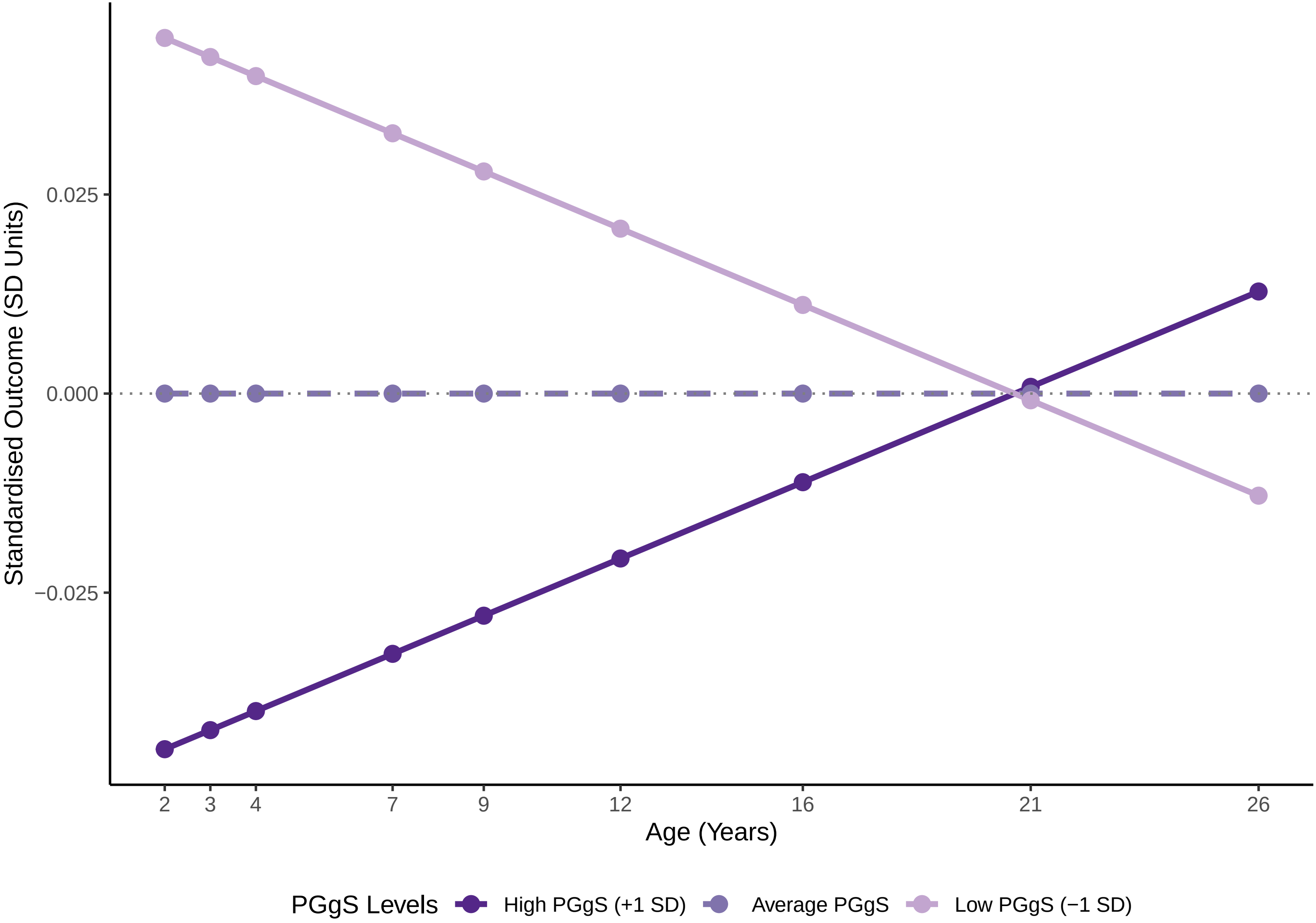

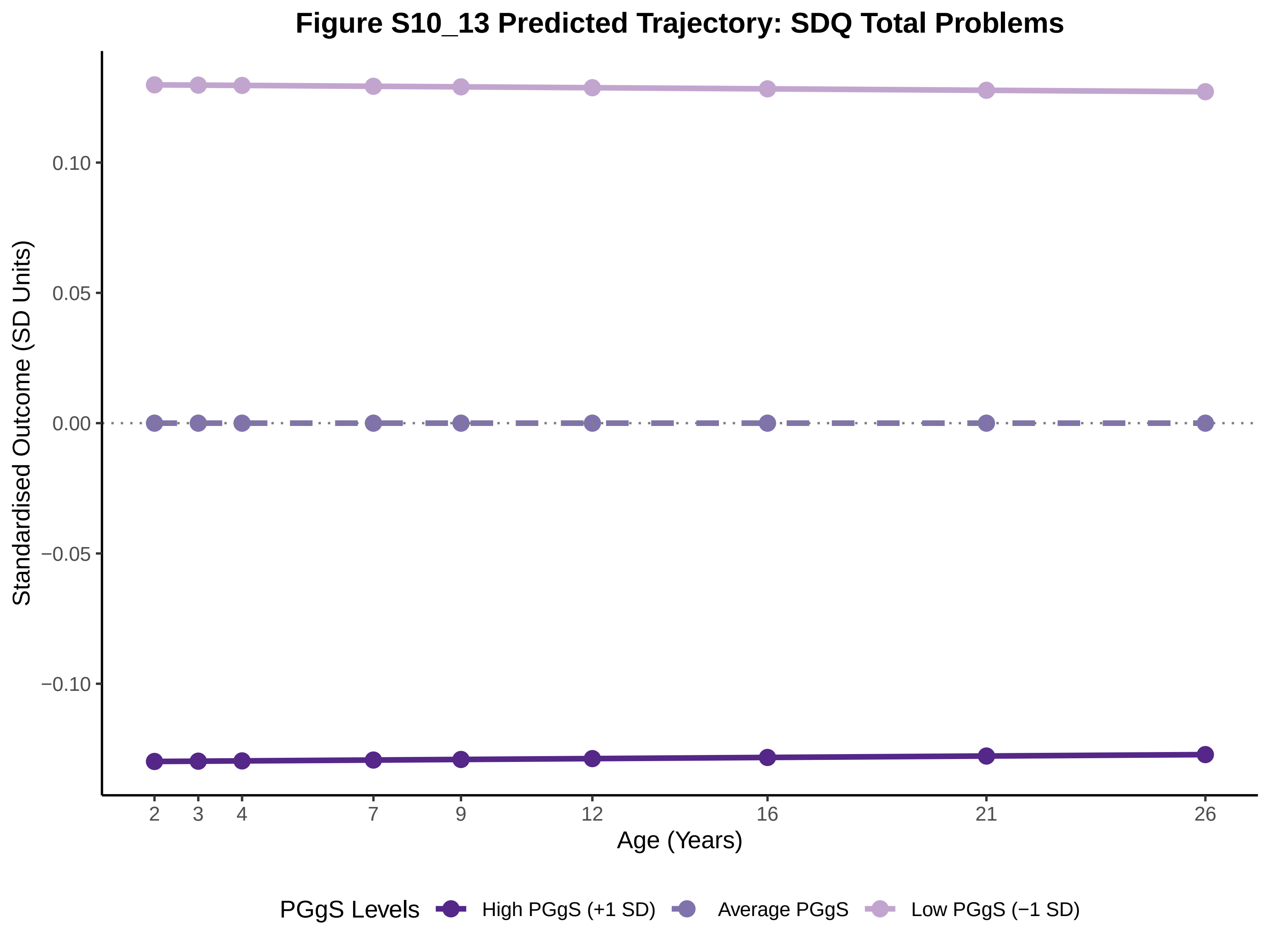
Figure S10_13 Predicted Trajectory: SDQ Total Problems
Standardised Outcome (SD Units)
0.10
0.05
0.00
−0.05
−0.10
2
3
4
7
9
12
16
21
26
Age (Years)
PGgS Levels
High PGgS (+1 SD)
Average PGgS
Low PGgS (−1 SD)

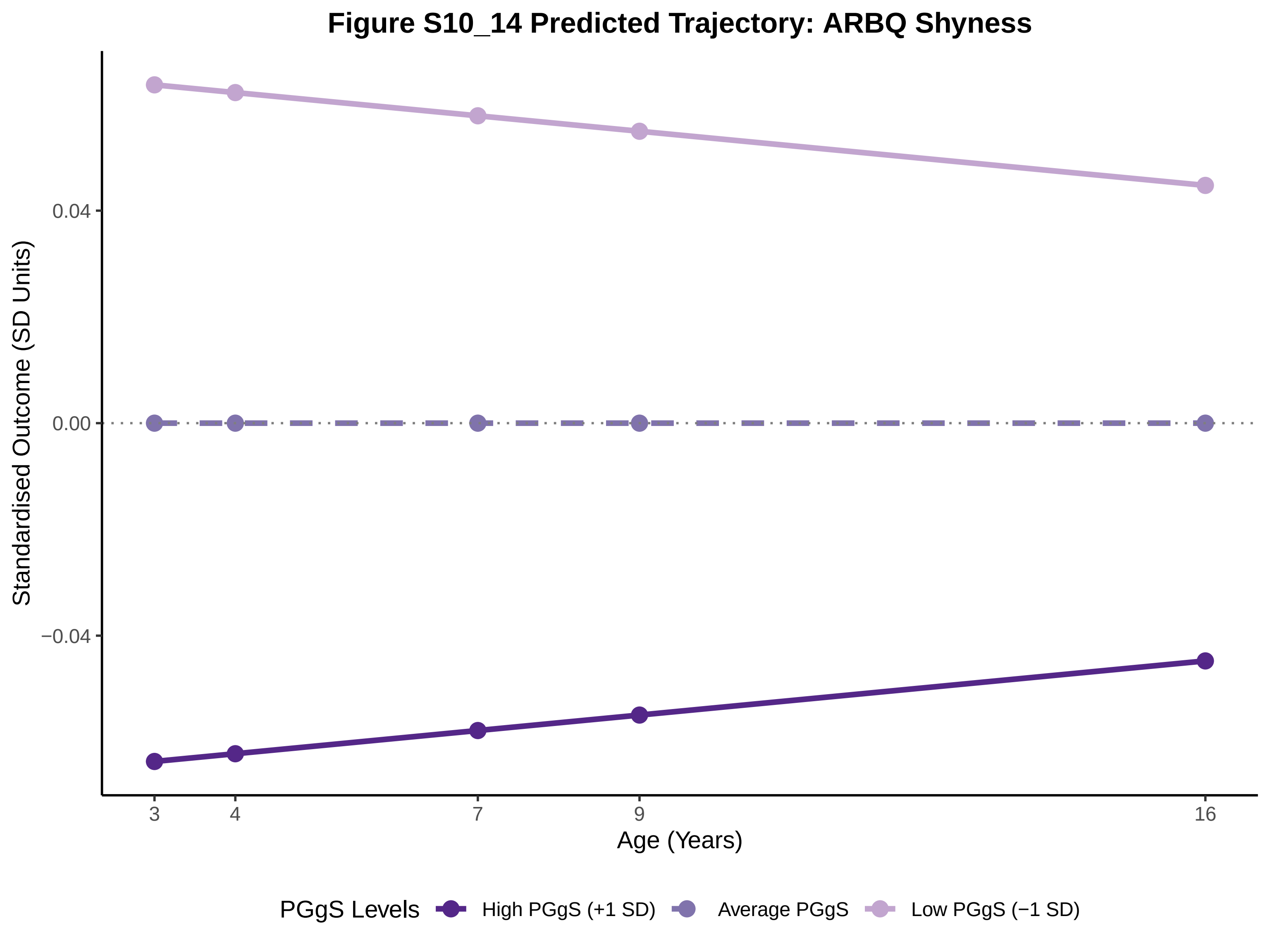
Figure S10_14 Predicted Trajectory: ARBQ Shyness
Standardised Outcome (SD Units)
0.04
0.00
−0.04
3
4
7
9
16
Age (Years)
PGgS Levels
High PGgS (+1 SD)
Average PGgS
Low PGgS (−1 SD)

### Figure S10_15 Predicted Trajectory: ARBQ Fear

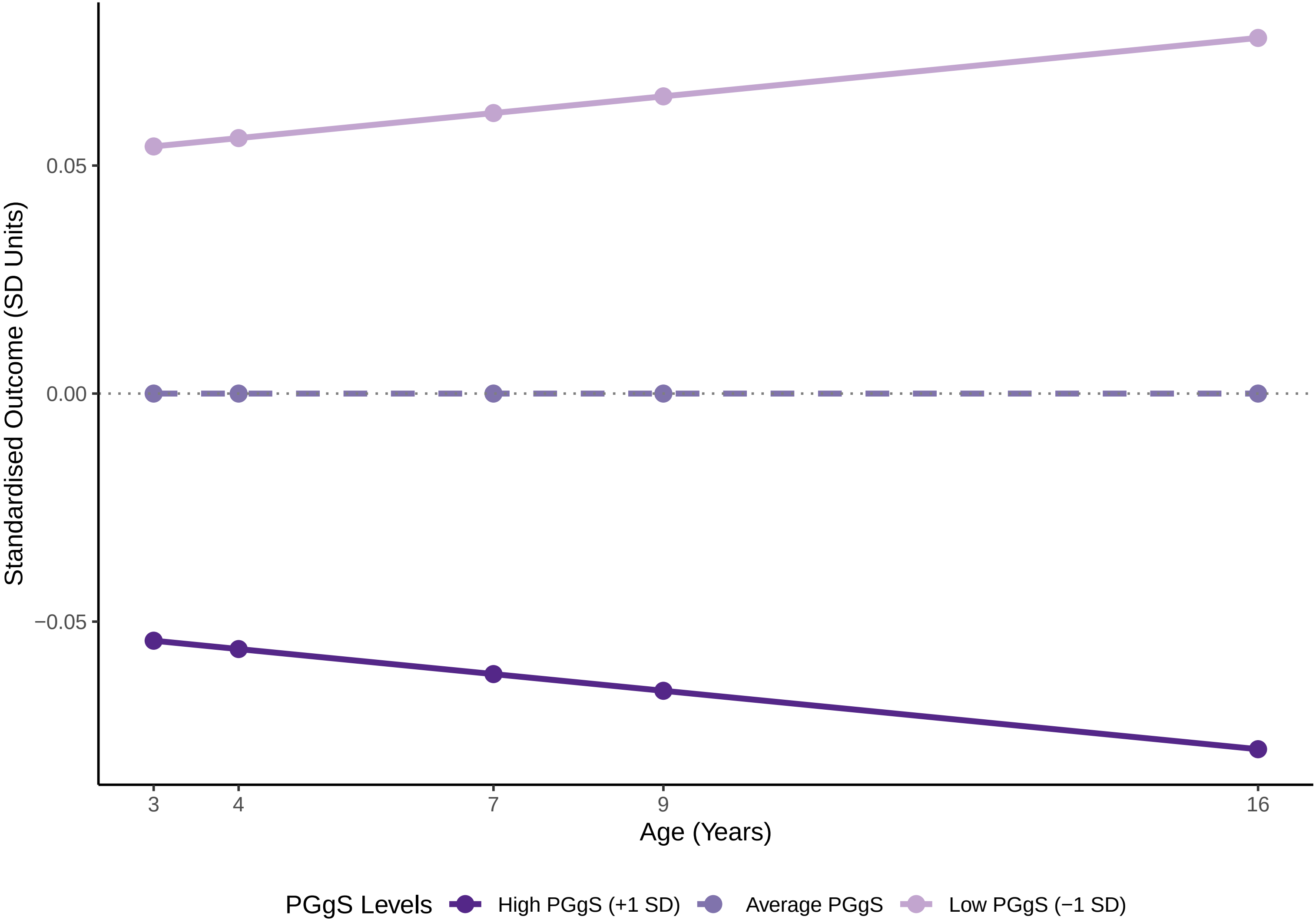

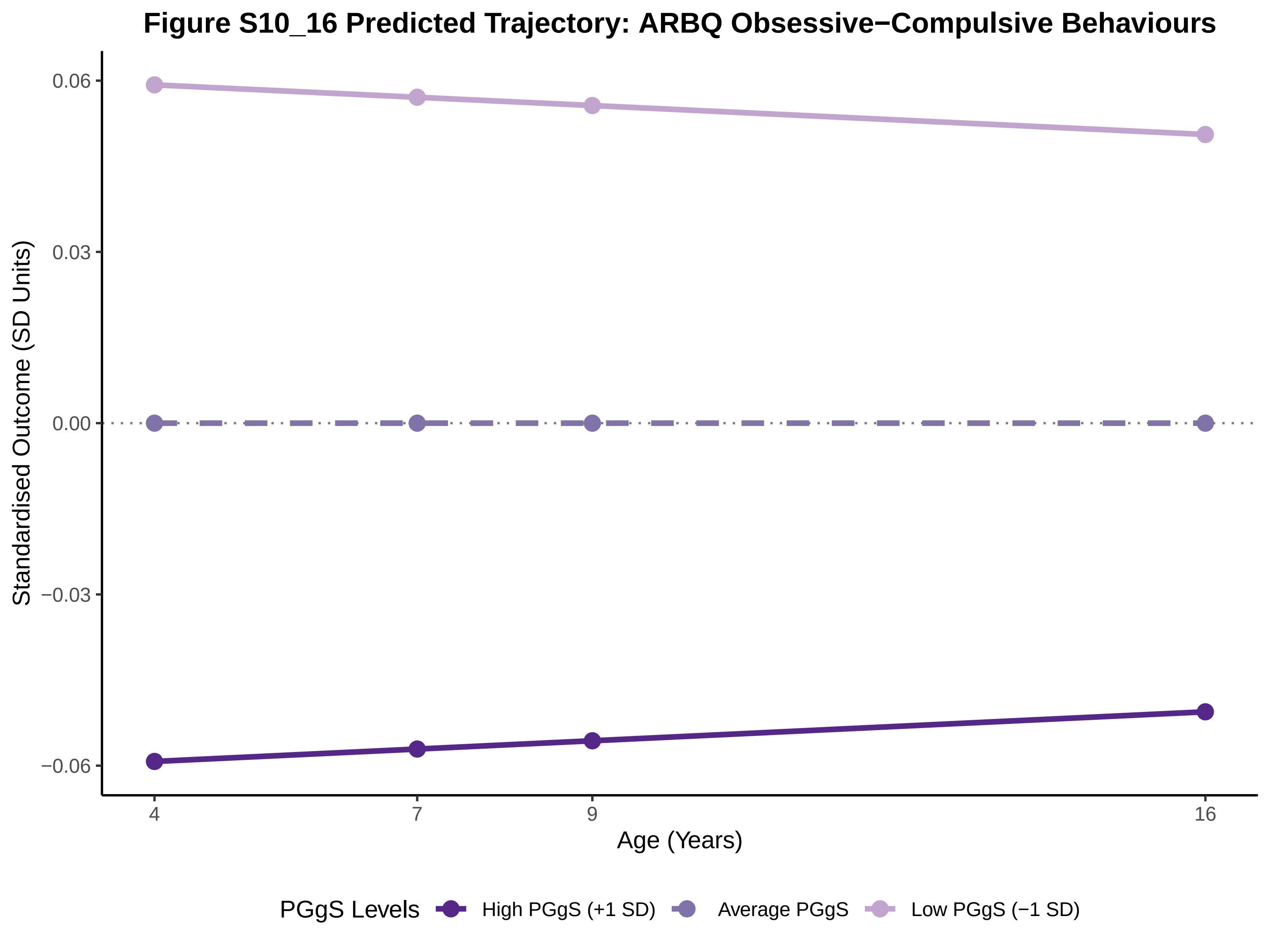

Figure S10_16 Predicted Trajectory: ARBQ Obsessive−Compulsive Behaviours
Standardised Outcome (SD Units)
0.06
0.03
0.00
−0.03
−0.06
4
7
9
16
Age (Years)
PGgS Levels
High PGgS (+1 SD)
Average PGgS
Low PGgS (−1 SD)

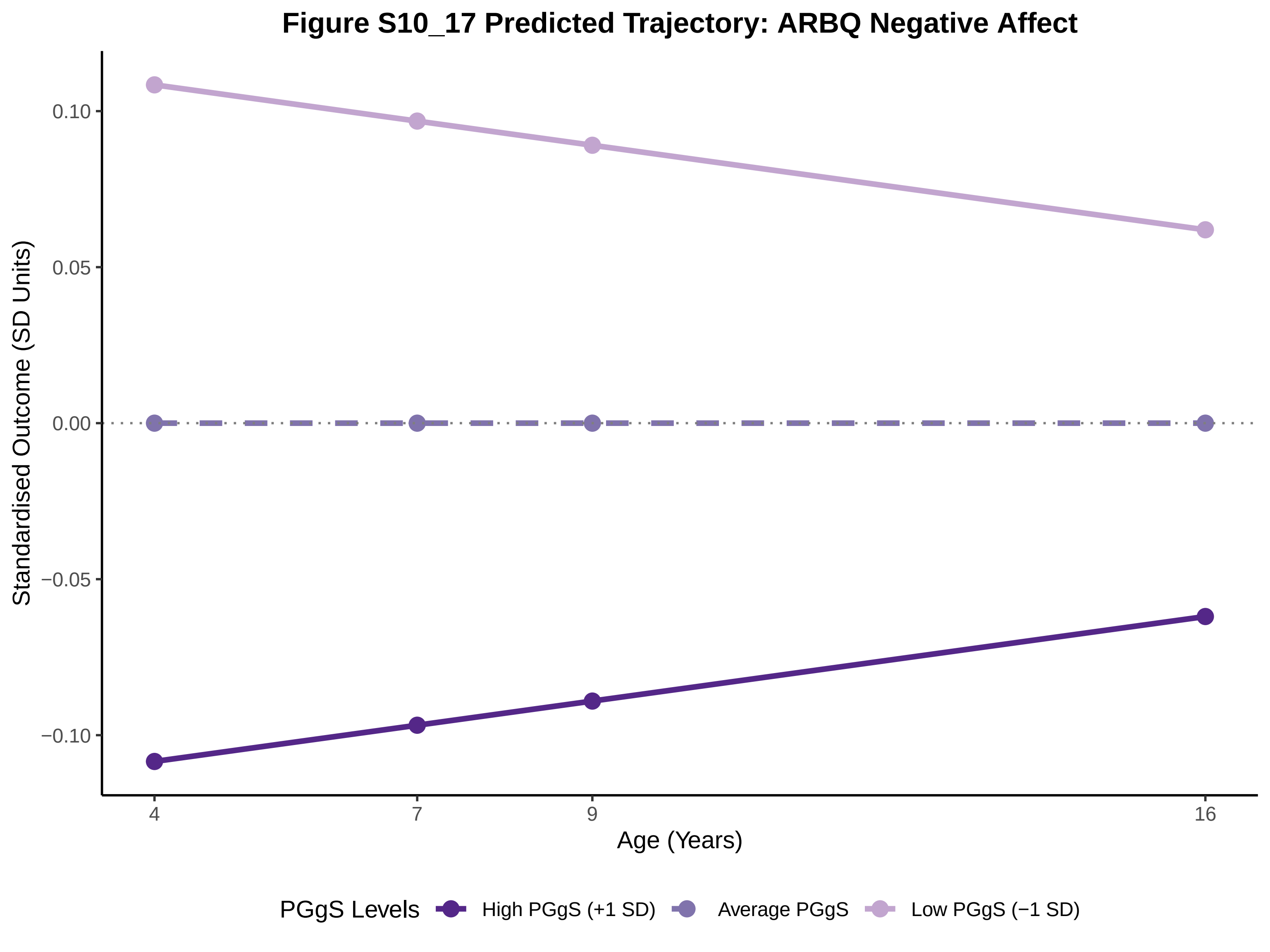
Figure S10_17 Predicted Trajectory: ARBQ Negative Affect
Standardised Outcome (SD Units)
0.10
0.05
0.00
−0.05
−0.10
4
7
9
16
Age (Years)
PGgS Levels
High PGgS (+1 SD)
Average PGgS
Low PGgS (−1 SD)

### Figure S10_18 Predicted Trajectory: ARBQ Negative Cognition

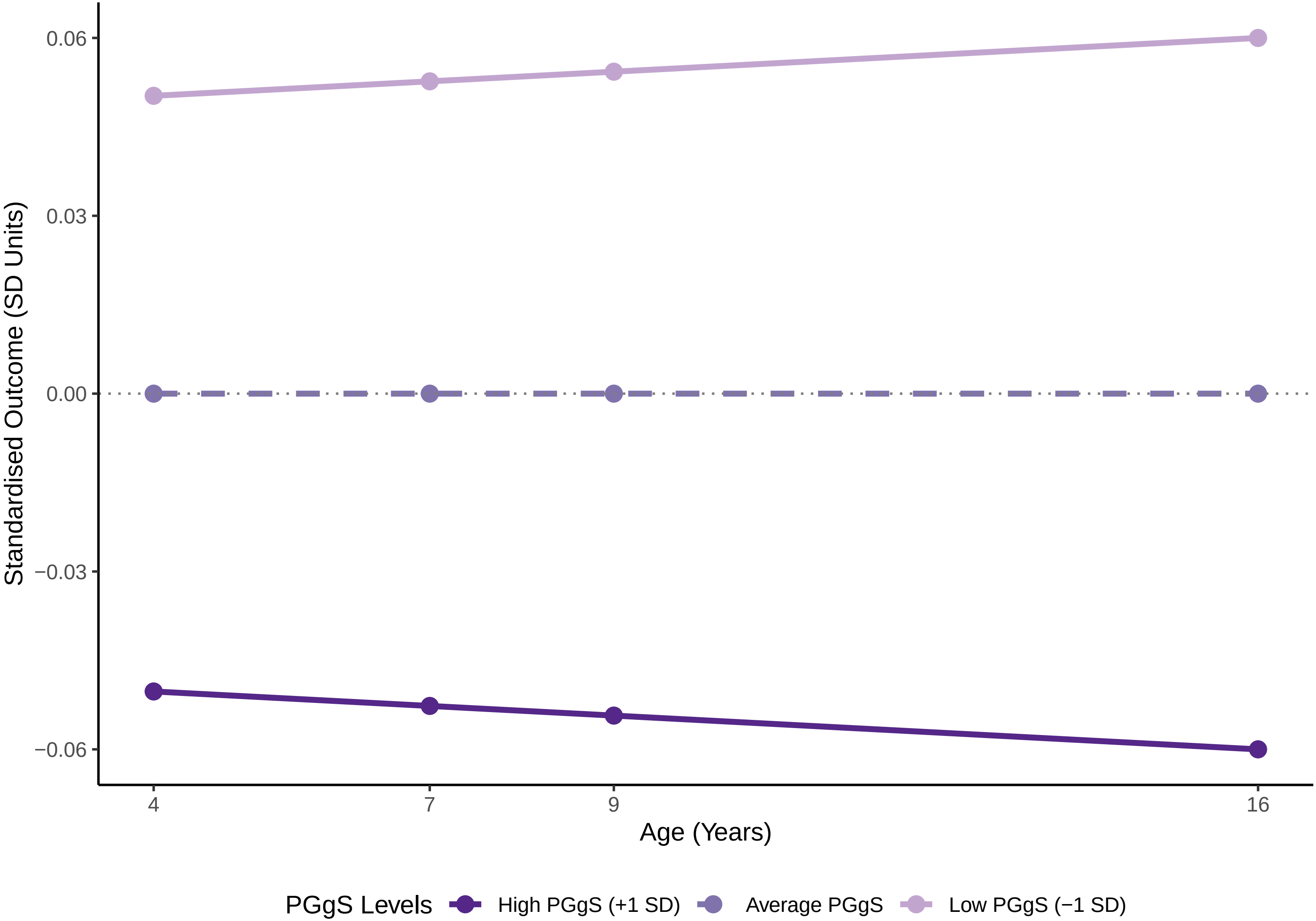

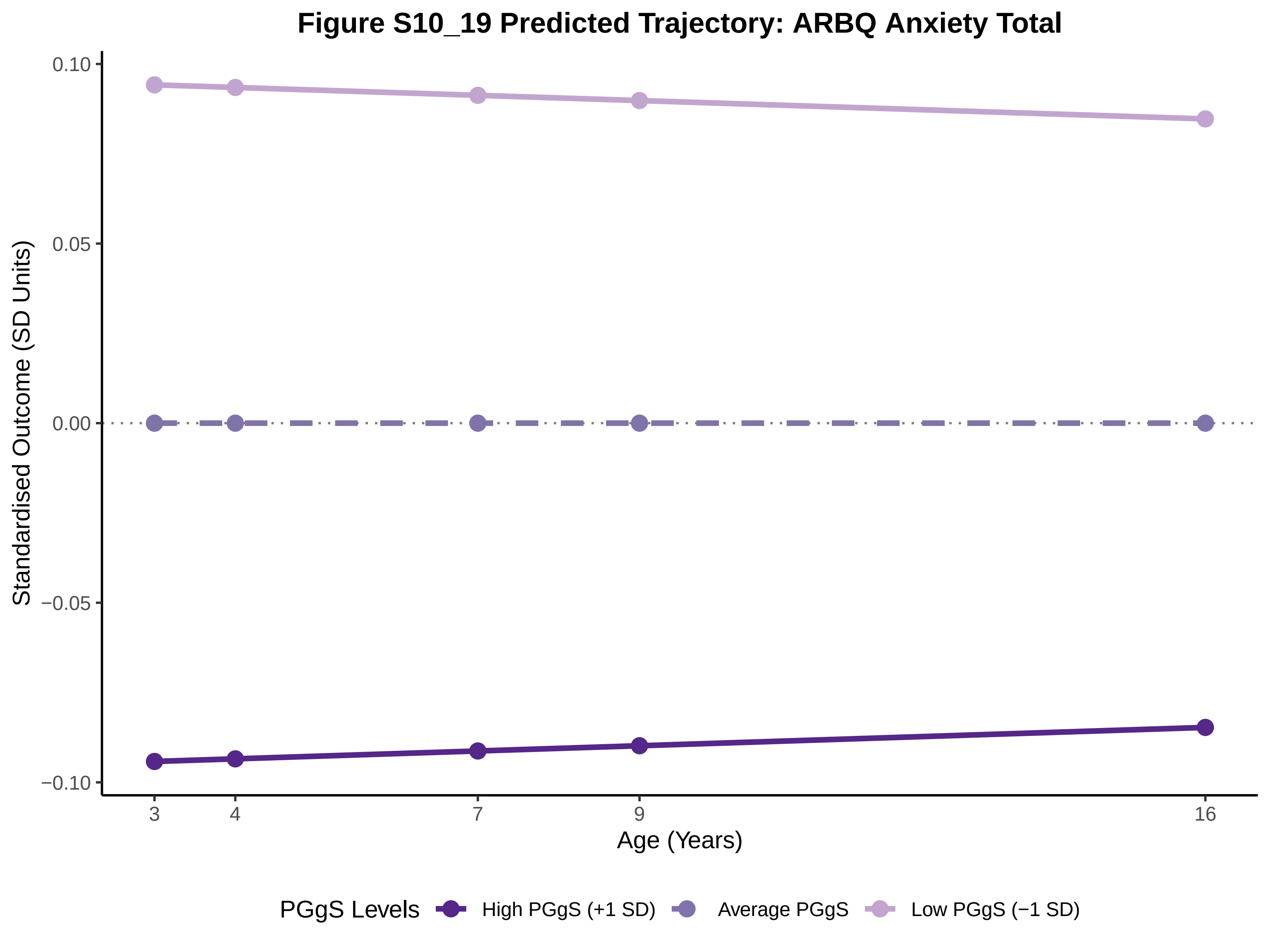
Figure S10_19 Predicted Trajectory: ARBQ Anxiety Total
0.10
0.05
0.00
−0.05
−0.10
Standardised Outcome (SD Units)
3
4
7
9
16
Age (Years)
PGgS Levels
High PGgS (+1 SD)
Average PGgS
Low PGgS (−1 SD)

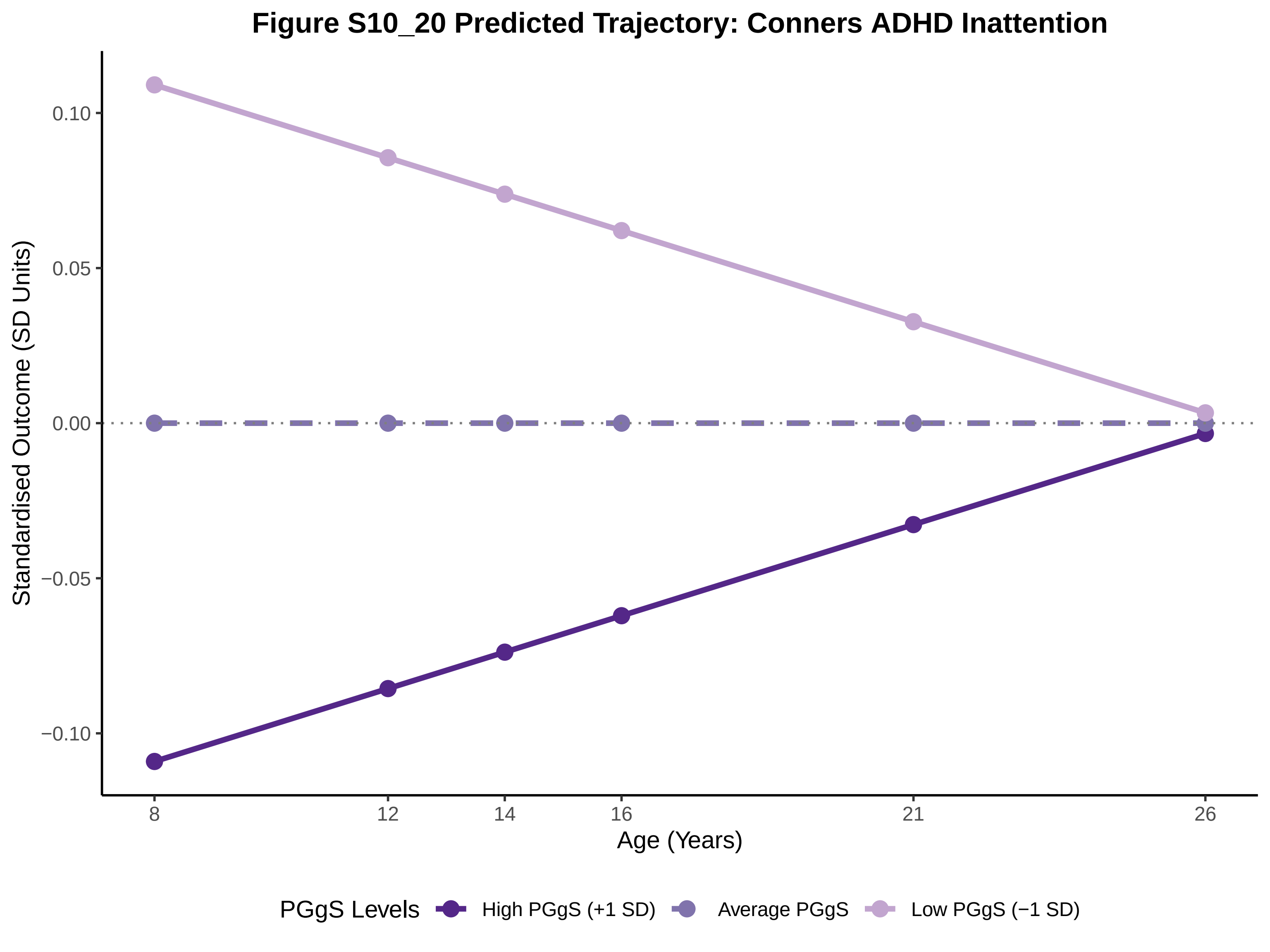

Figure S10_20 Predicted Trajectory: Conners ADHD Inattention
Standardised Outcome (SD Units)
0.10
0.05
0.00
−0.05
−0.10
8
12
14
16
21
26
Age (Years)
PGgS Levels
High PGgS (+1 SD)
Average PGgS
Low PGgS (−1 SD)

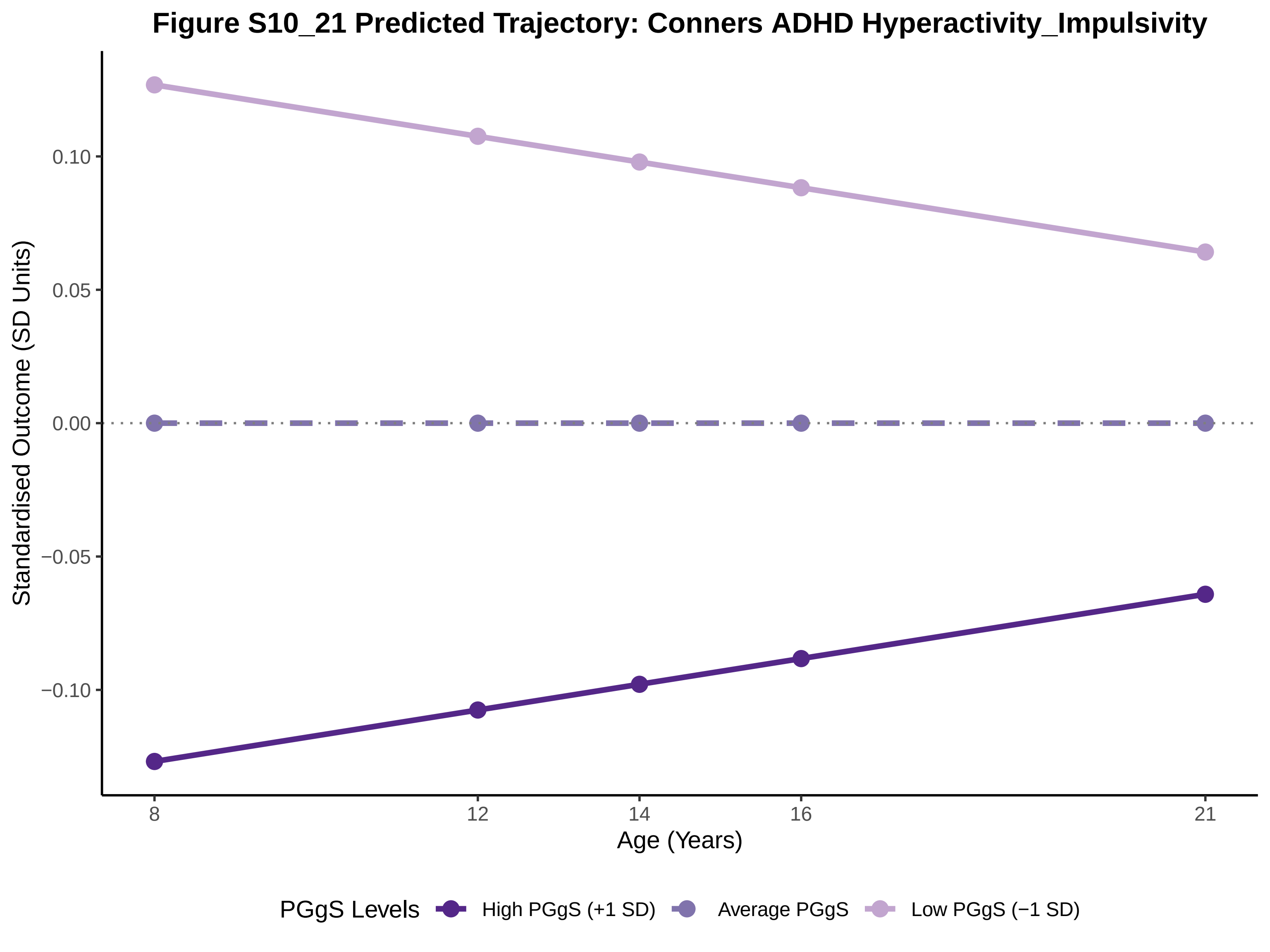

Figure S10_21 Predicted Trajectory: Conners ADHD Hyperactivity_Impulsivity
Standardised Outcome (SD Units)
0.10
0.05
0.00
−0.05
−0.10
8
12
14
16
21
Age (Years)
PGgS Levels
High PGgS (+1 SD)
Average PGgS
Low PGgS (−1 SD)

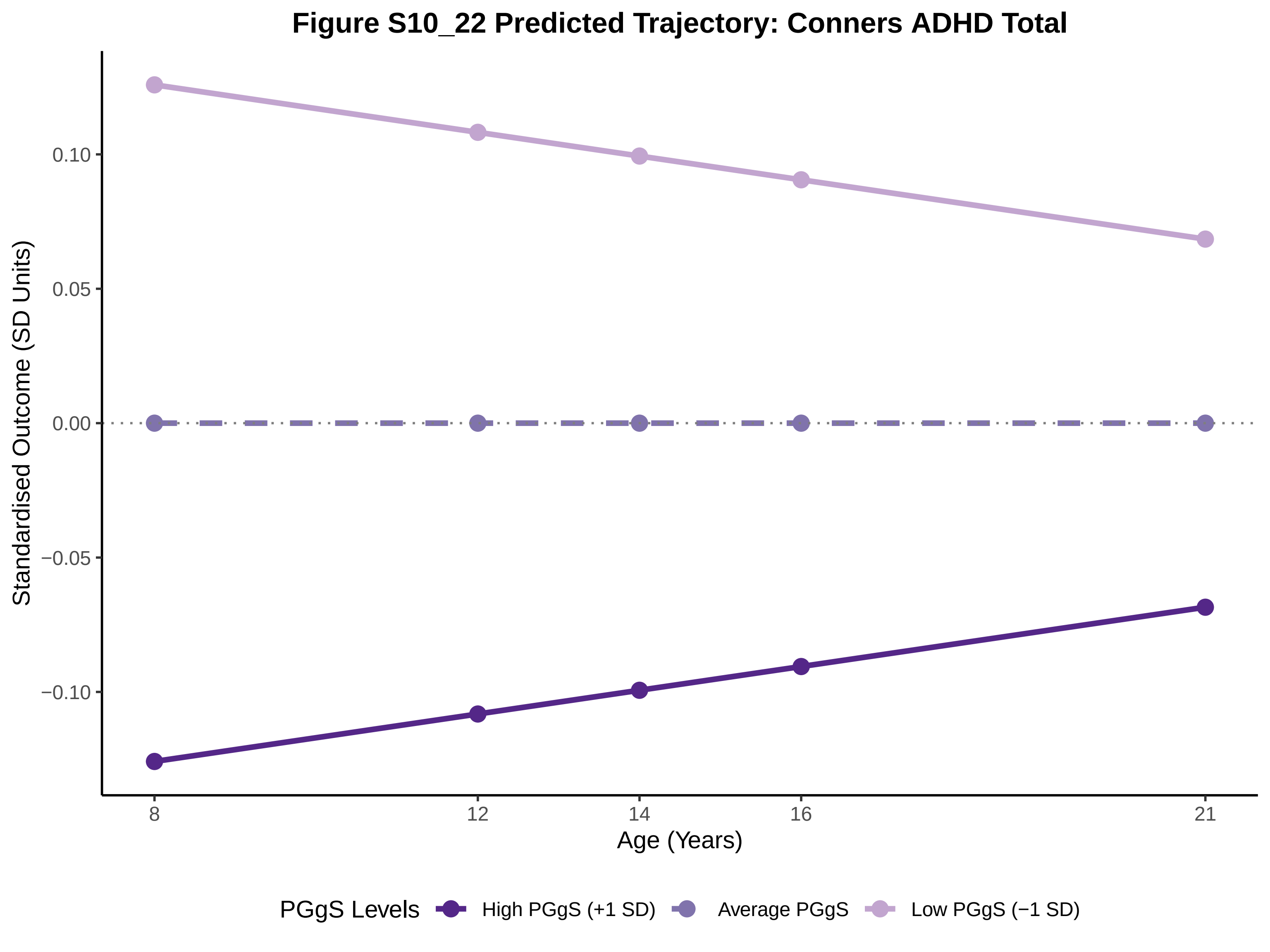
Figure S10_22 Predicted Trajectory: Conners ADHD Total
Standardised Outcome (SD Units)
0.10
0.05
0.00
−0.05
−0.10
8
12
14
16
21
Age (Years)
PGgS Levels
High PGgS (+1 SD)
Average PGgS
Low PGgS (−1 SD)

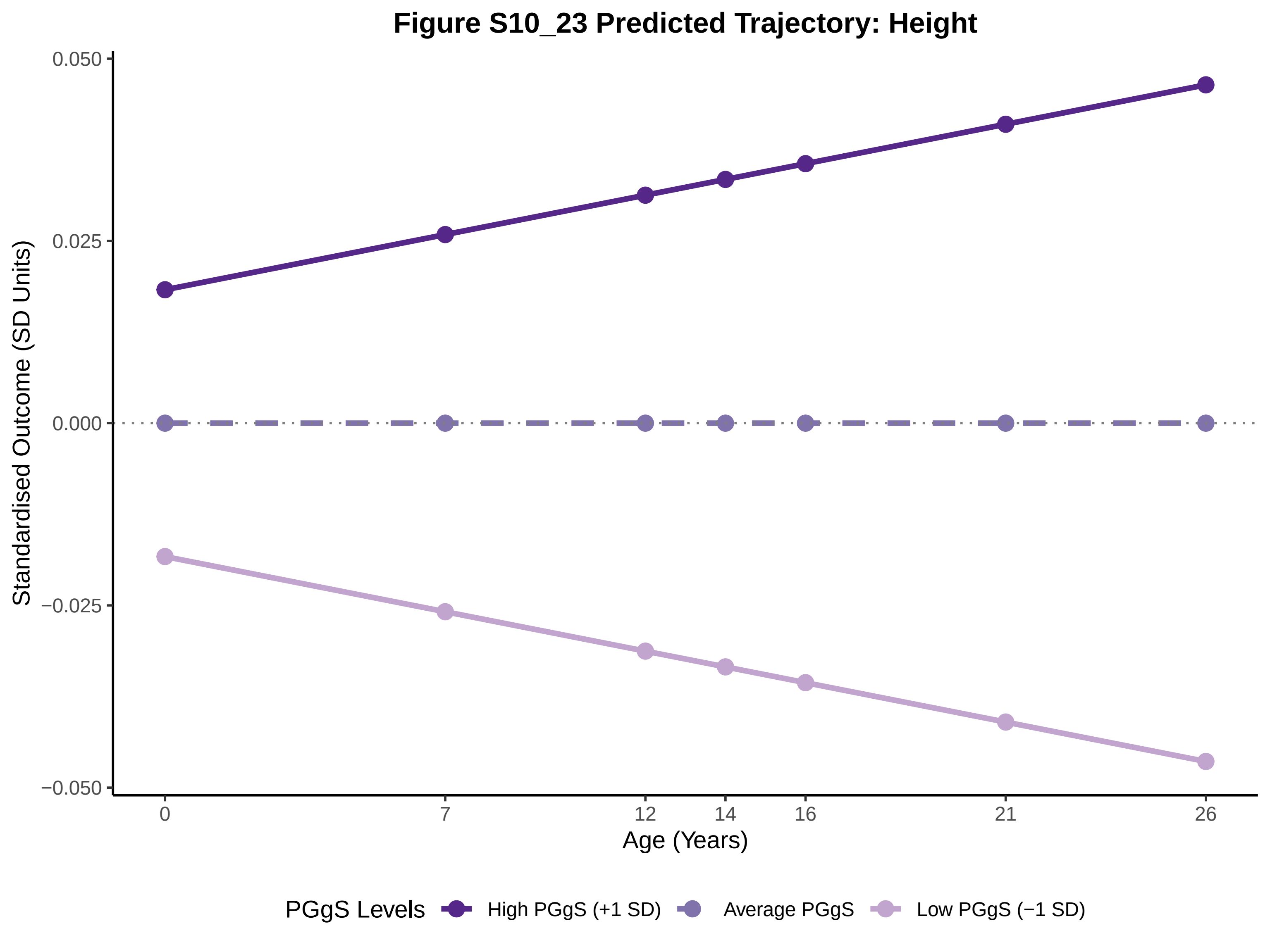

Figure S10_23 Predicted Trajectory: Height
Standardised Outcome (SD Units)
0.050
0.025
0.000
−0.025
−0.050
0
7
12
14
16
21
26
Age (Years)
PGgS Levels
High PGgS (+1 SD)
Average PGgS
Low PGgS (−1 SD)

### Figure S10_24 Predicted Trajectory: BMI (as weight at birth)

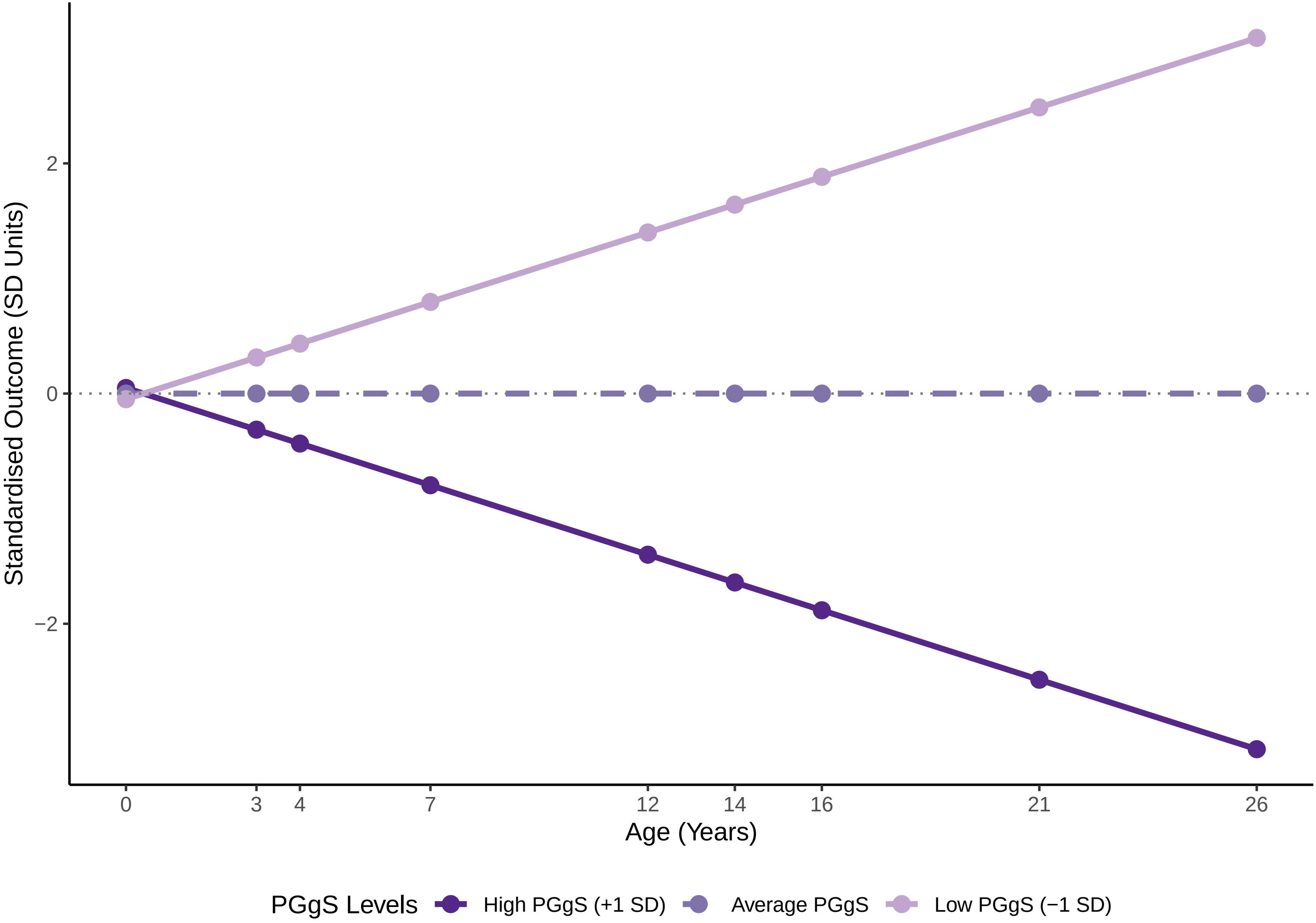

### Figure S12_1 Predicted Trajectory: General Cognitive Ability

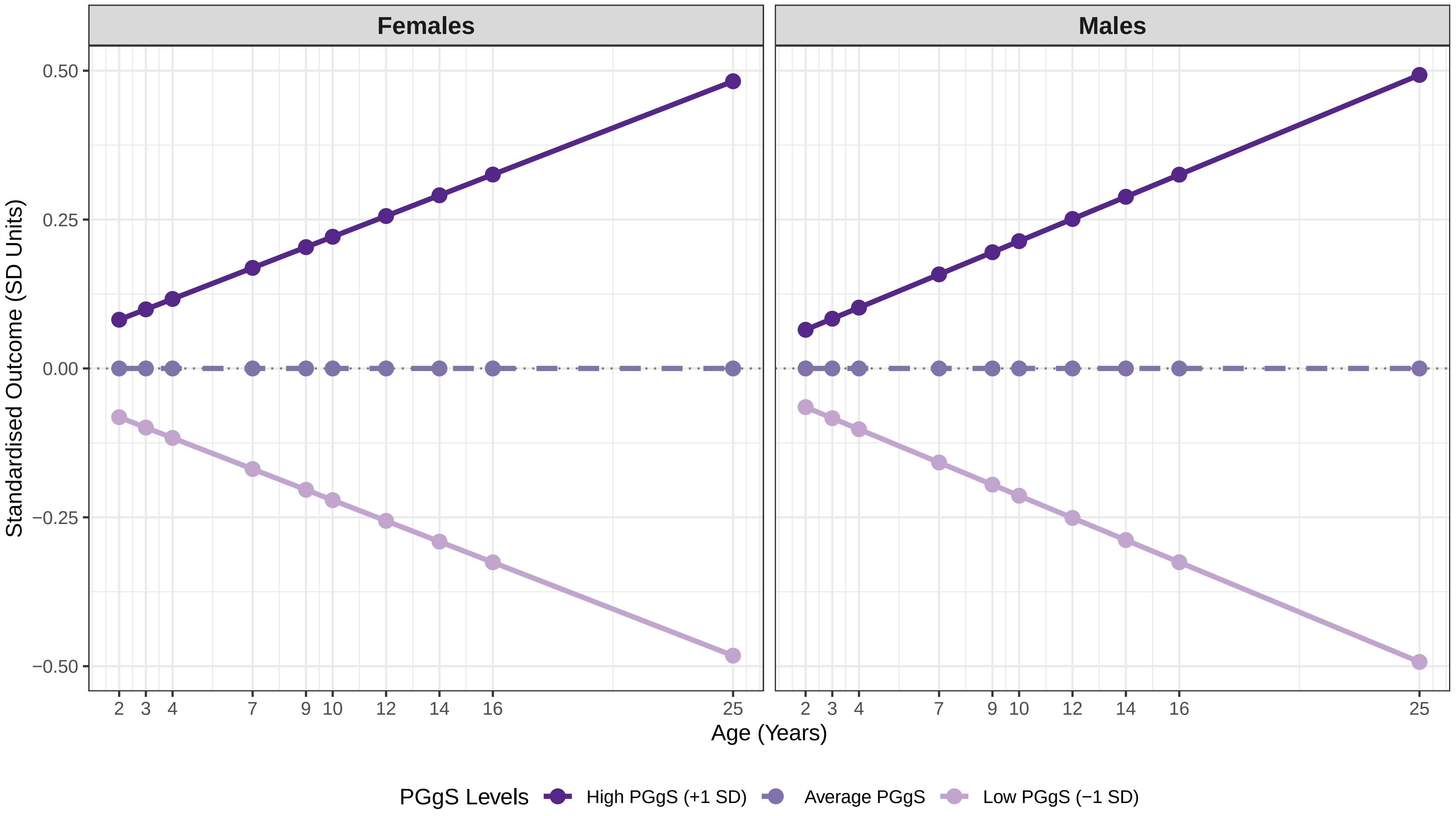

#### Figure S12_2 Predicted Trajectory: Verbal Abilities

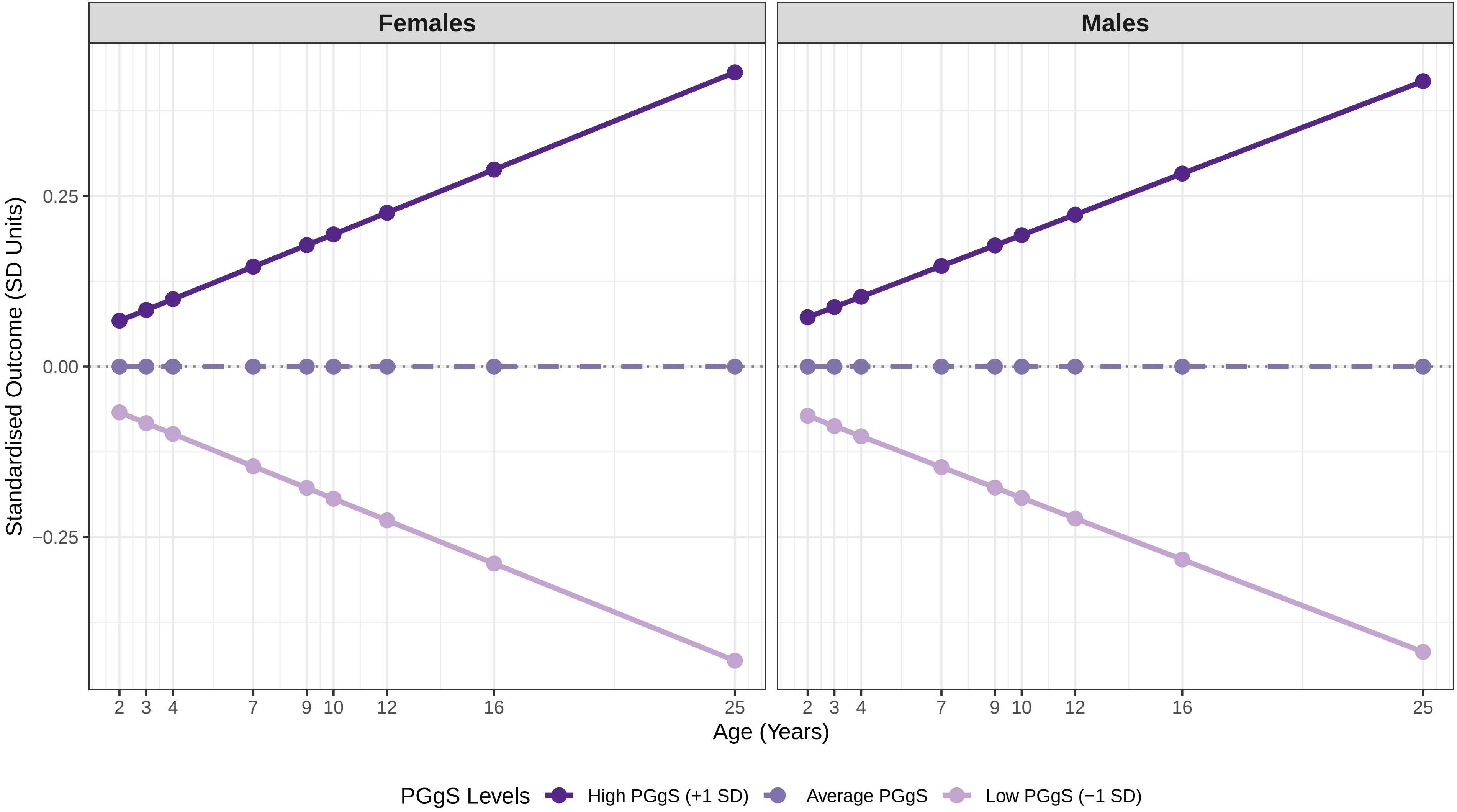

### Figure S12_3 Predicted Trajectory: Nonverbal Abilities

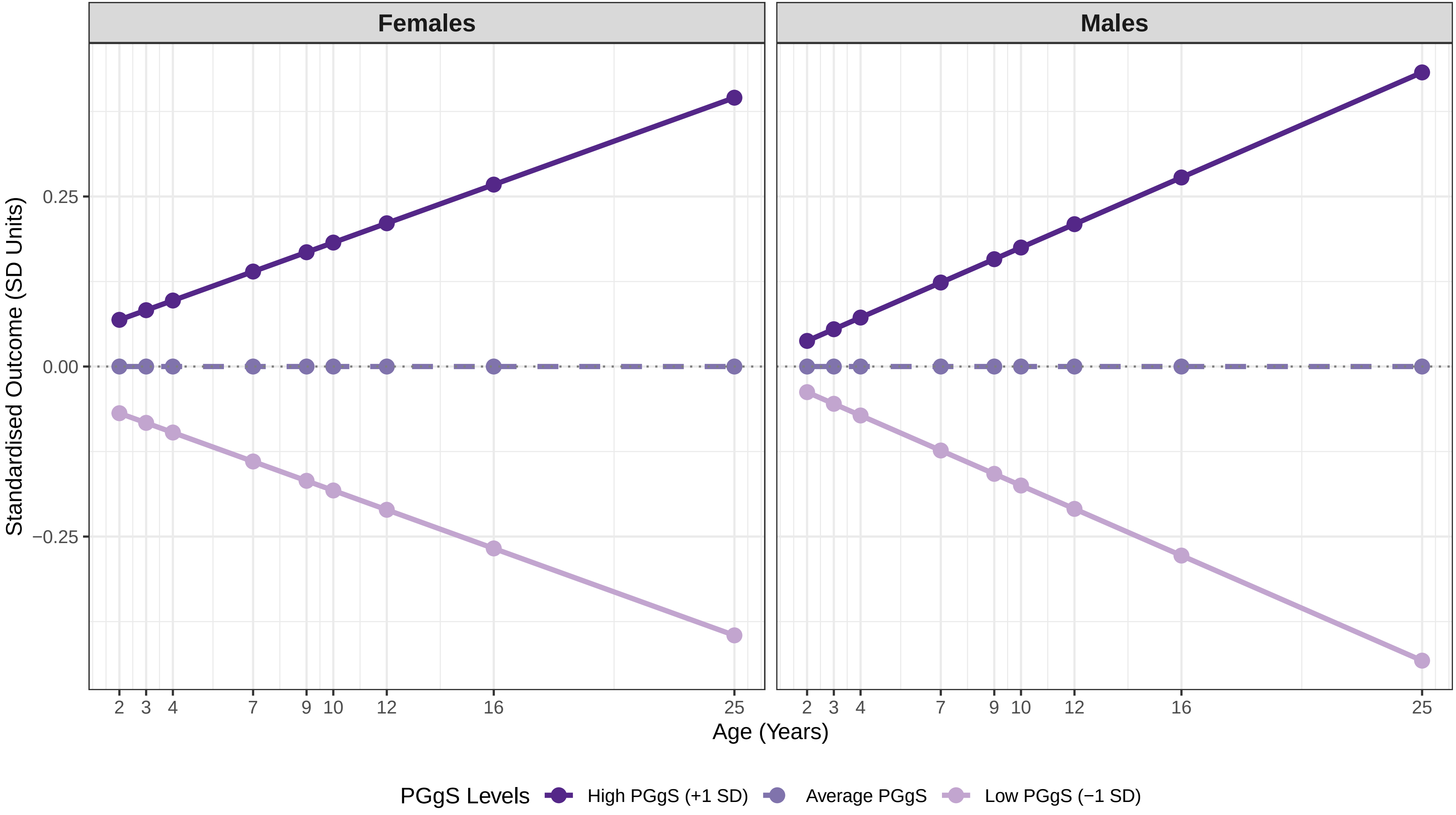

#### Figure S12_4 Predicted Trajectory: English Grades

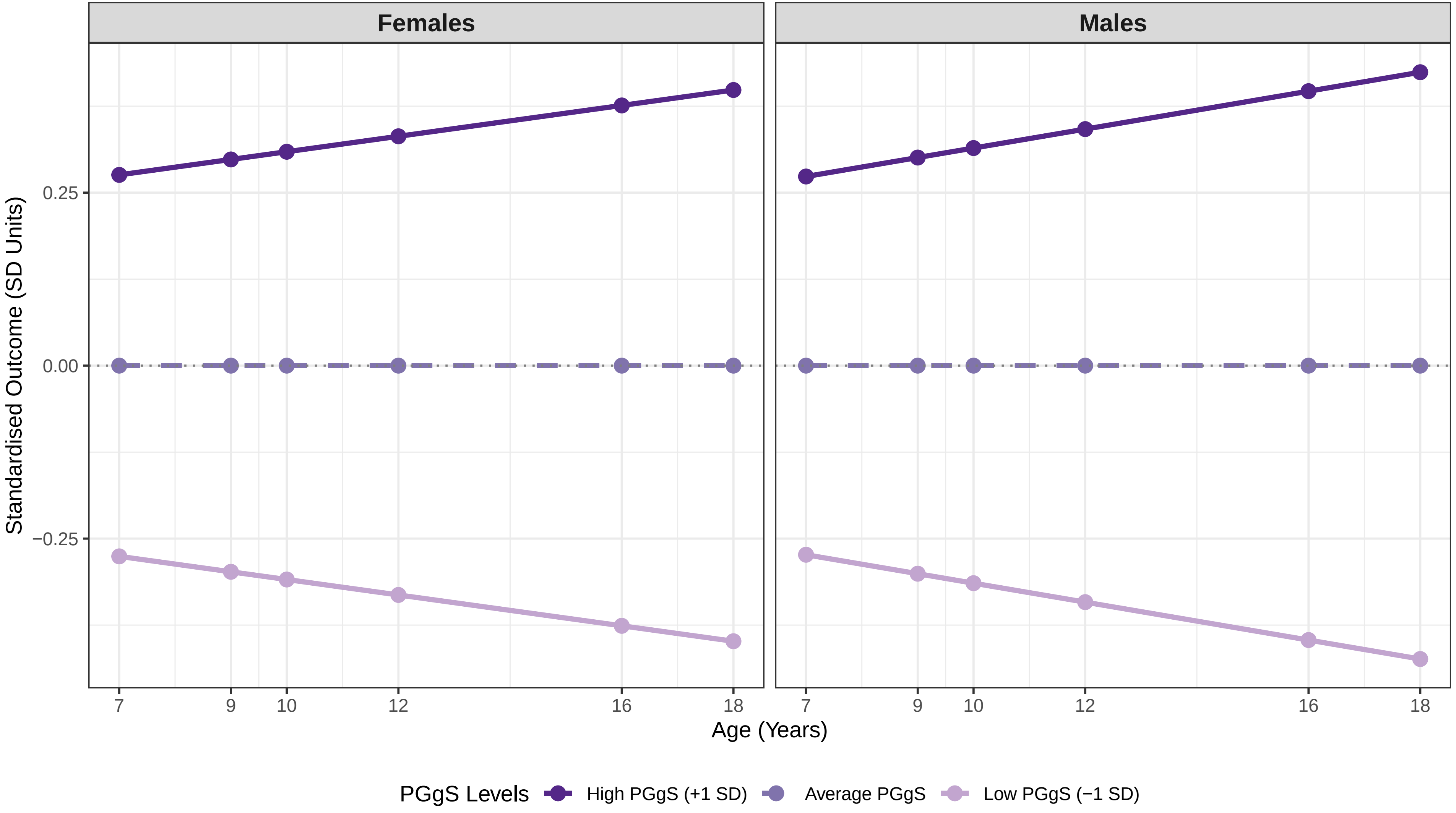

### Figure S12_5 Predicted Trajectory: Maths Grades

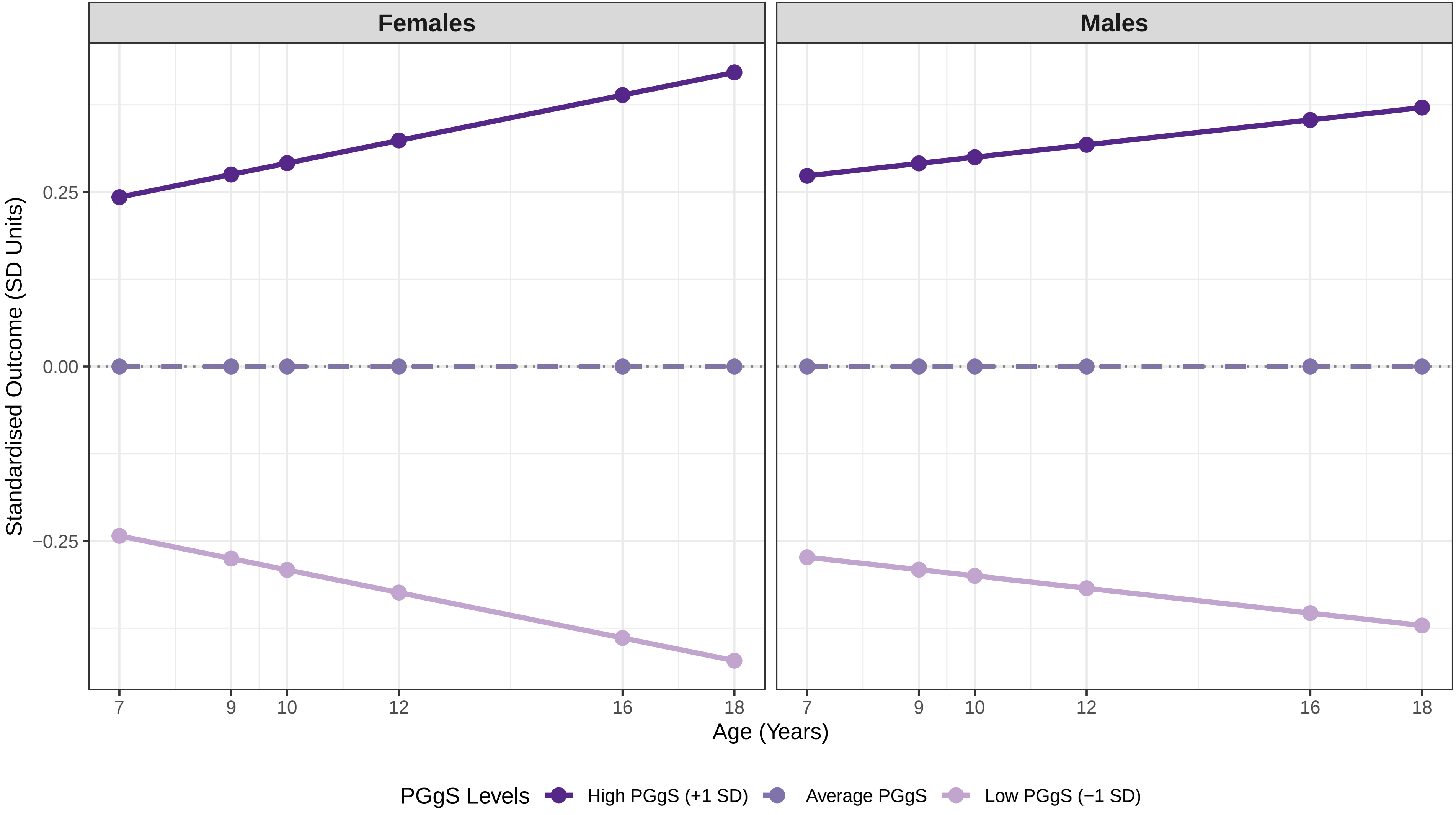

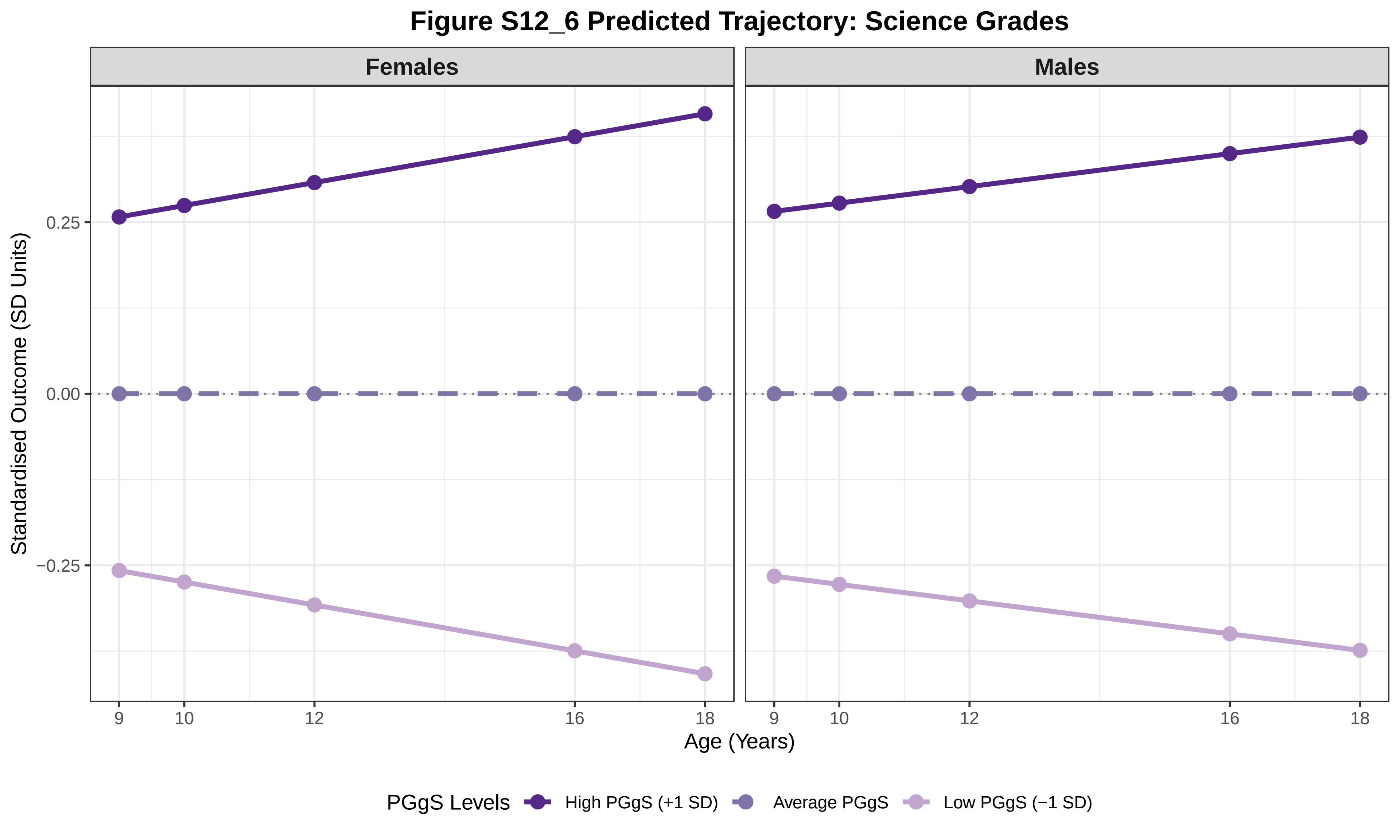

Figure S12_6 Predicted Trajectory: Science Grades
Females
Males
Standardised Outcome (SD Units)
0.25
0.00
−0.25
9
10
12
16
18
Age (Years)
PGgS Levels
High PGgS (+1 SD)
Average PGgS
Low PGgS (−1 SD)

**Figure S12_7 Predicted Trajectory: Core Subject Grades**

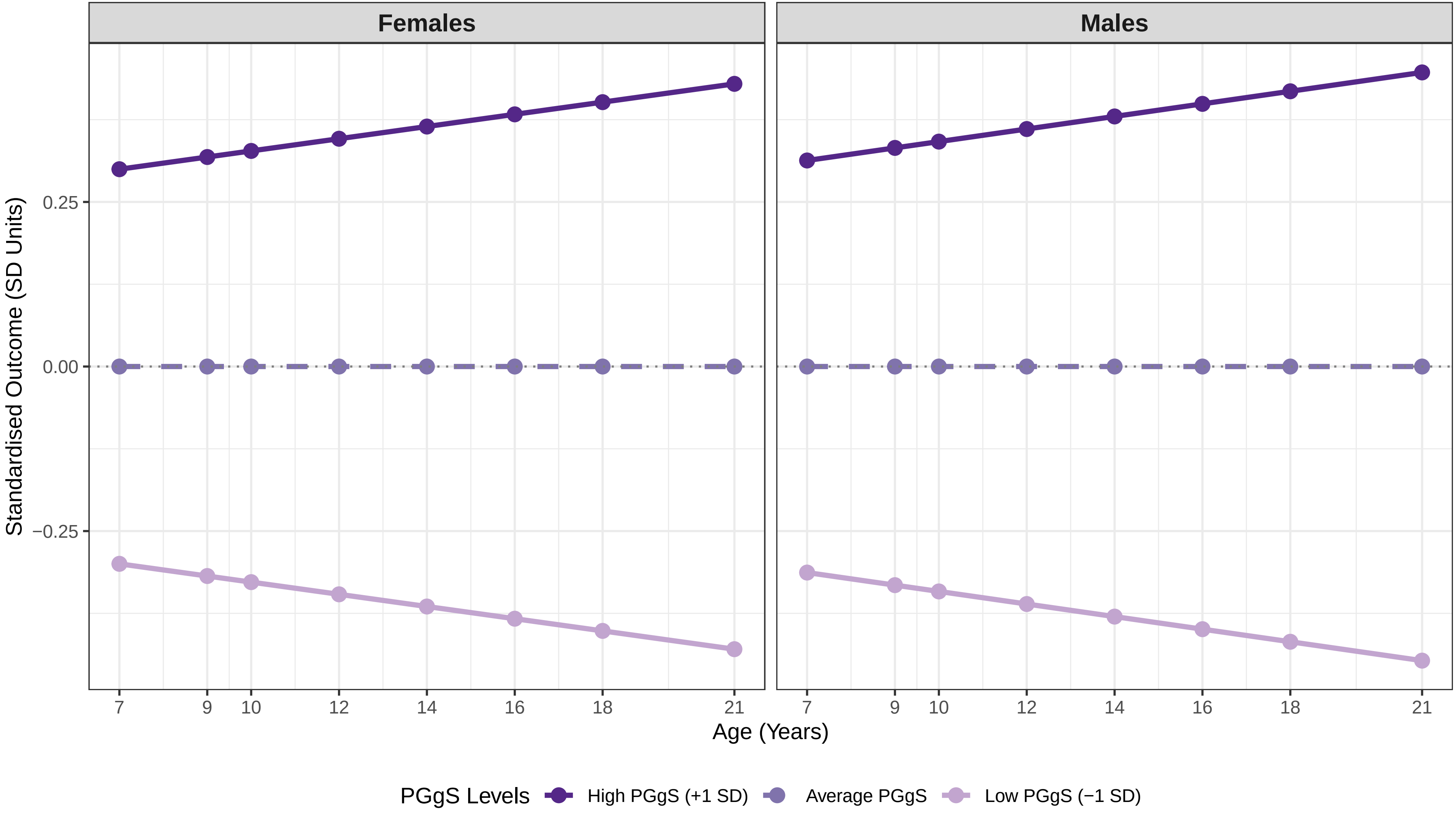

### Figure S12_8 Predicted Trajectory: SDQ Conduct

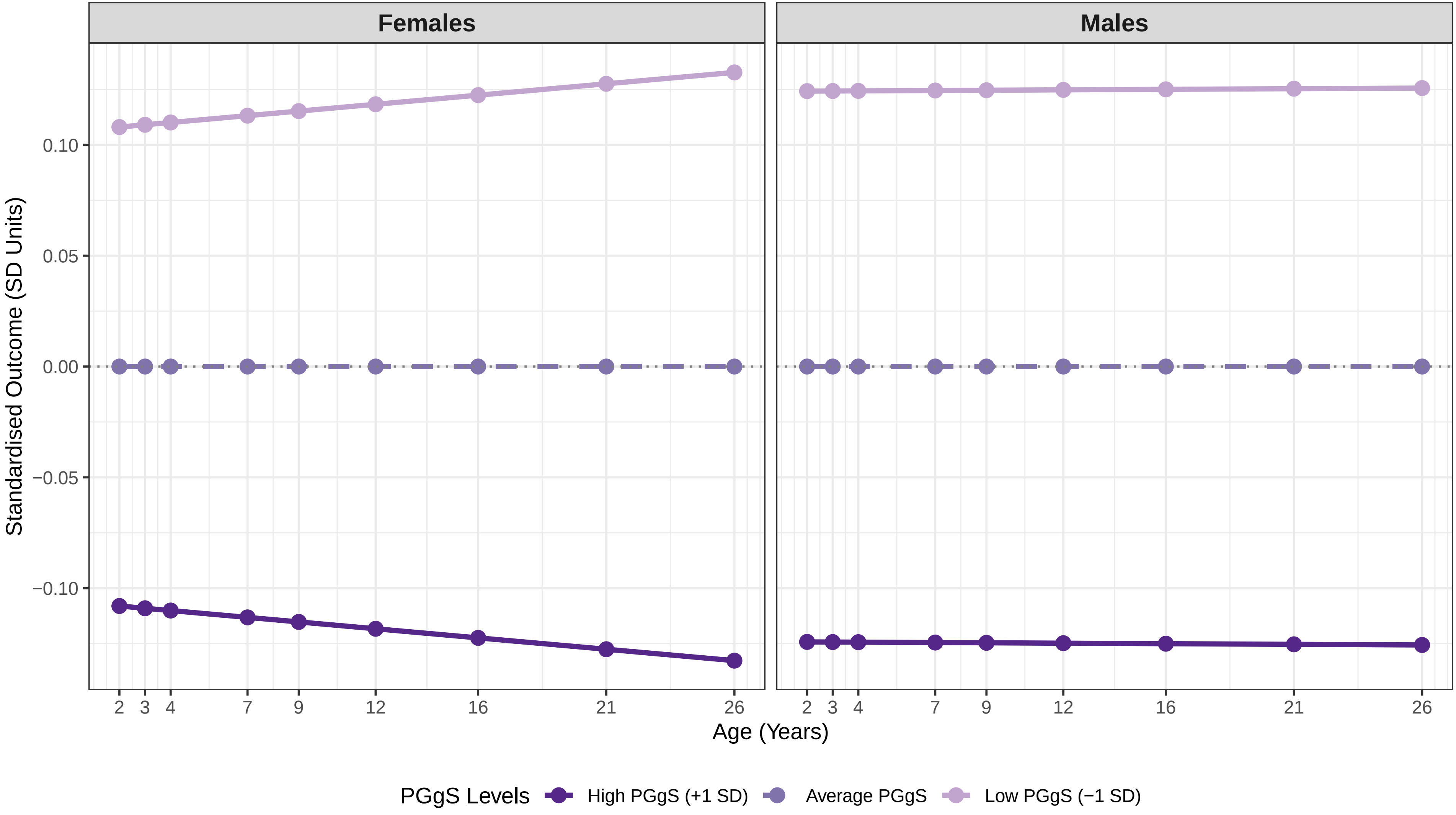

**Figure S12_9 Predicted Trajectory: SDQ Emotion**

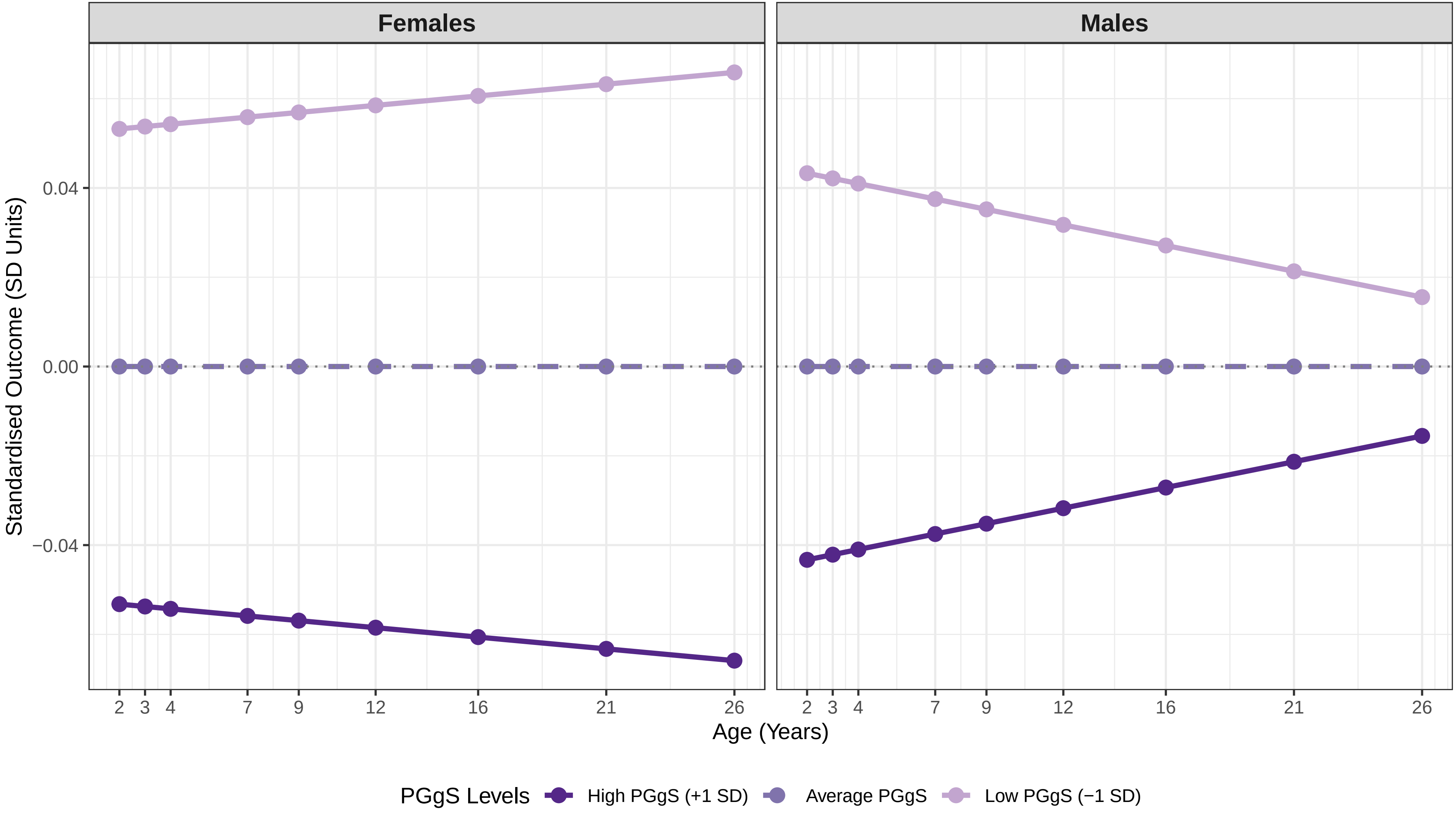

### Figure S12_10 Predicted Trajectory: SDQ Hyperactivity_SDQ

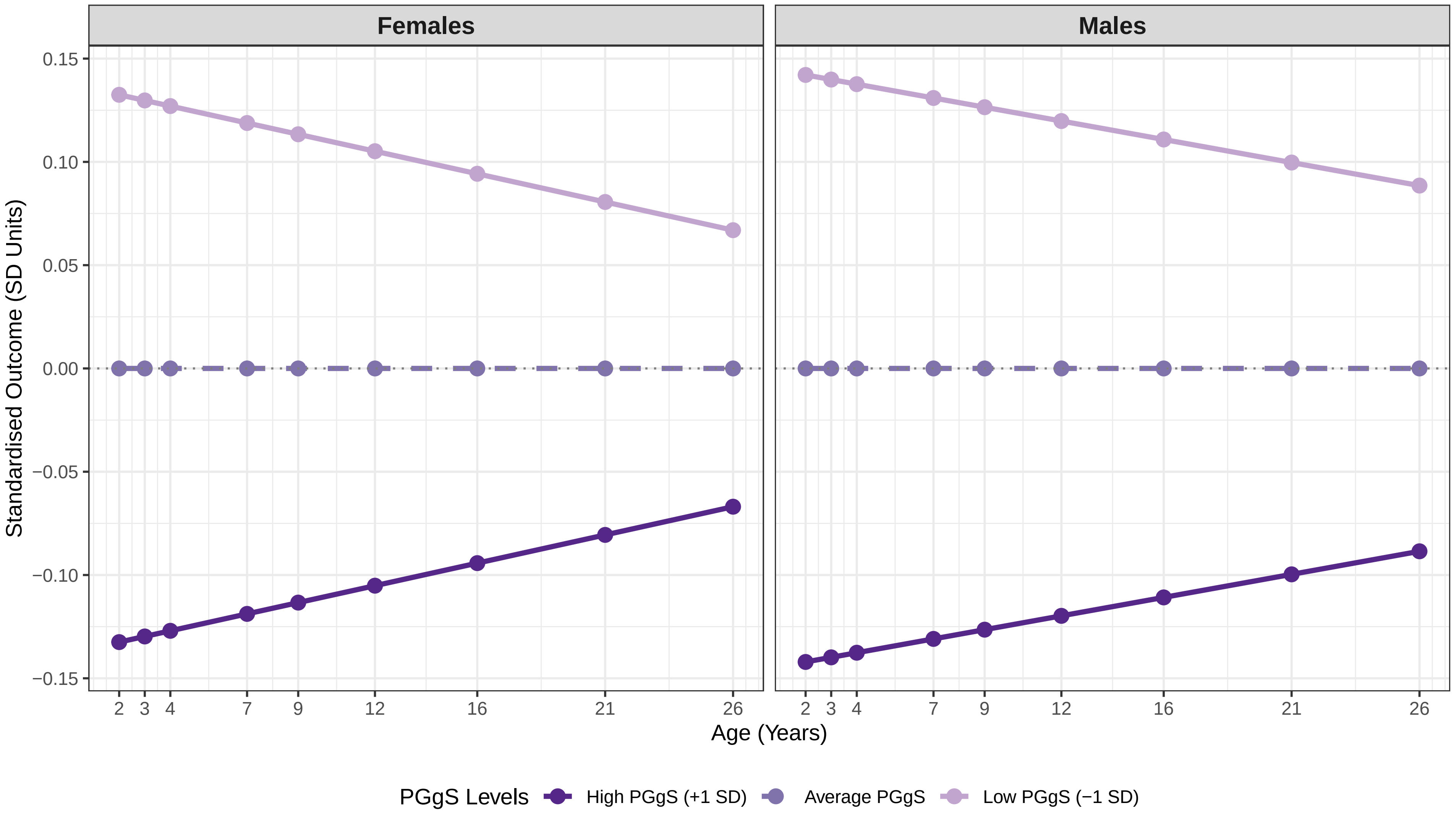

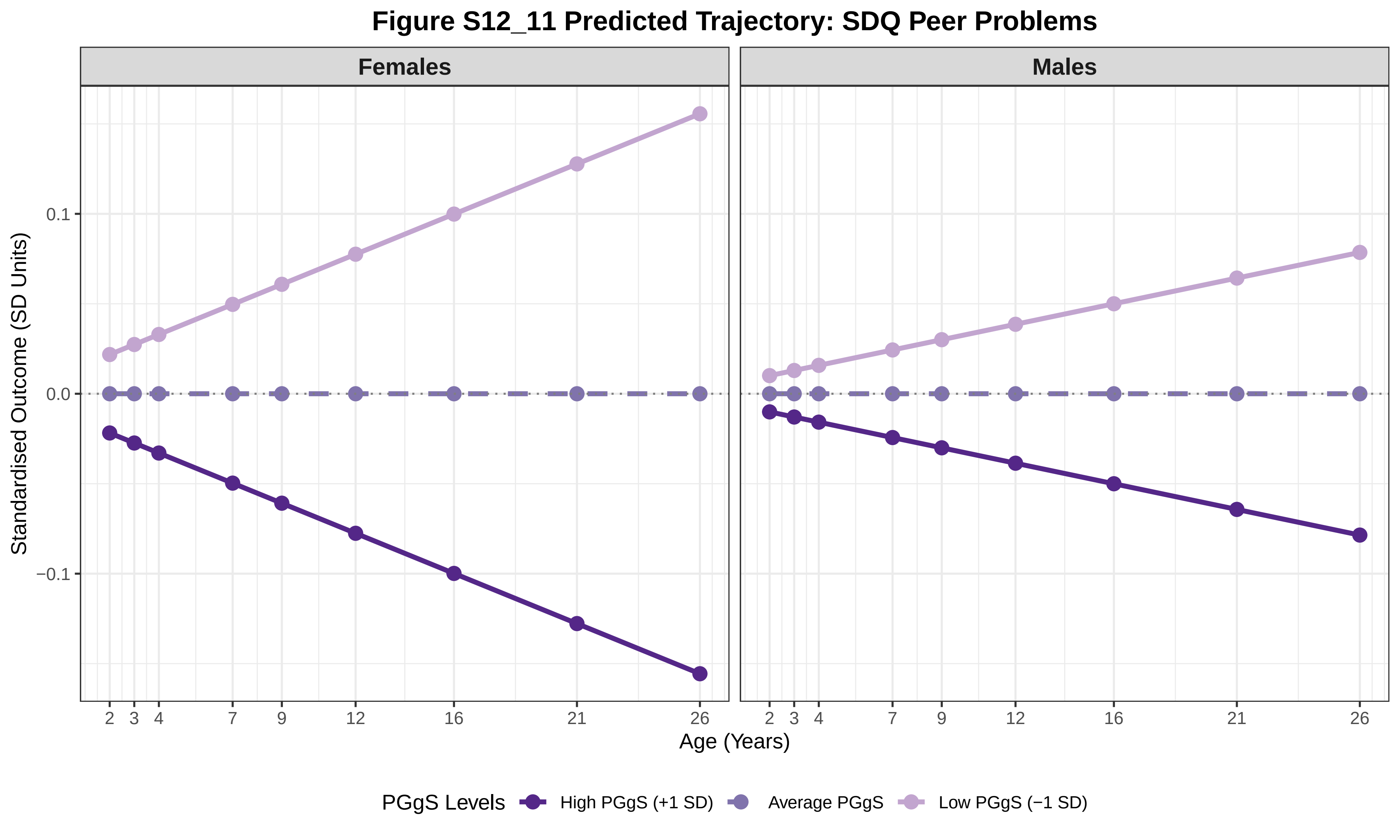

Figure S12_11 Predicted Trajectory: SDQ Peer Problems
Females
Males
Standardised Outcome (SD Units)
0.1
0.0
−0.1
2
3
4
7
9
12
16
21
26
Age (Years)
PGgS Levels
High PGgS (+1 SD)
Average PGgS
Low PGgS (−1 SD)

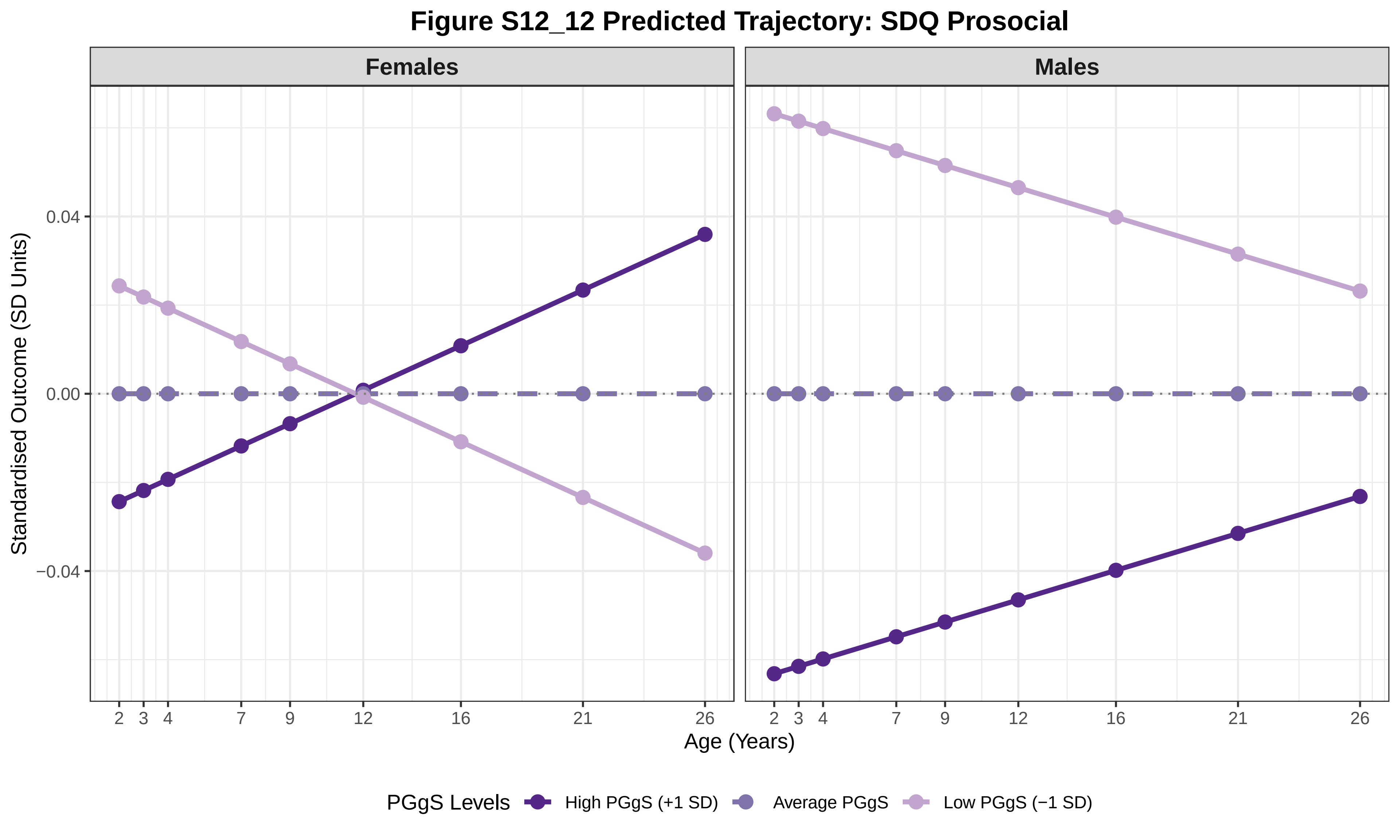

Figure S12_12 Predicted Trajectory: SDQ Prosocial
Females
Males
Standardised Outcome (SD Units)
0.04
0.00
−0.04
2
3
4
7
9
12
16
21
26
Age (Years)
PGgS Levels
High PGgS (+1 SD)
Average PGgS
Low PGgS (−1 SD)

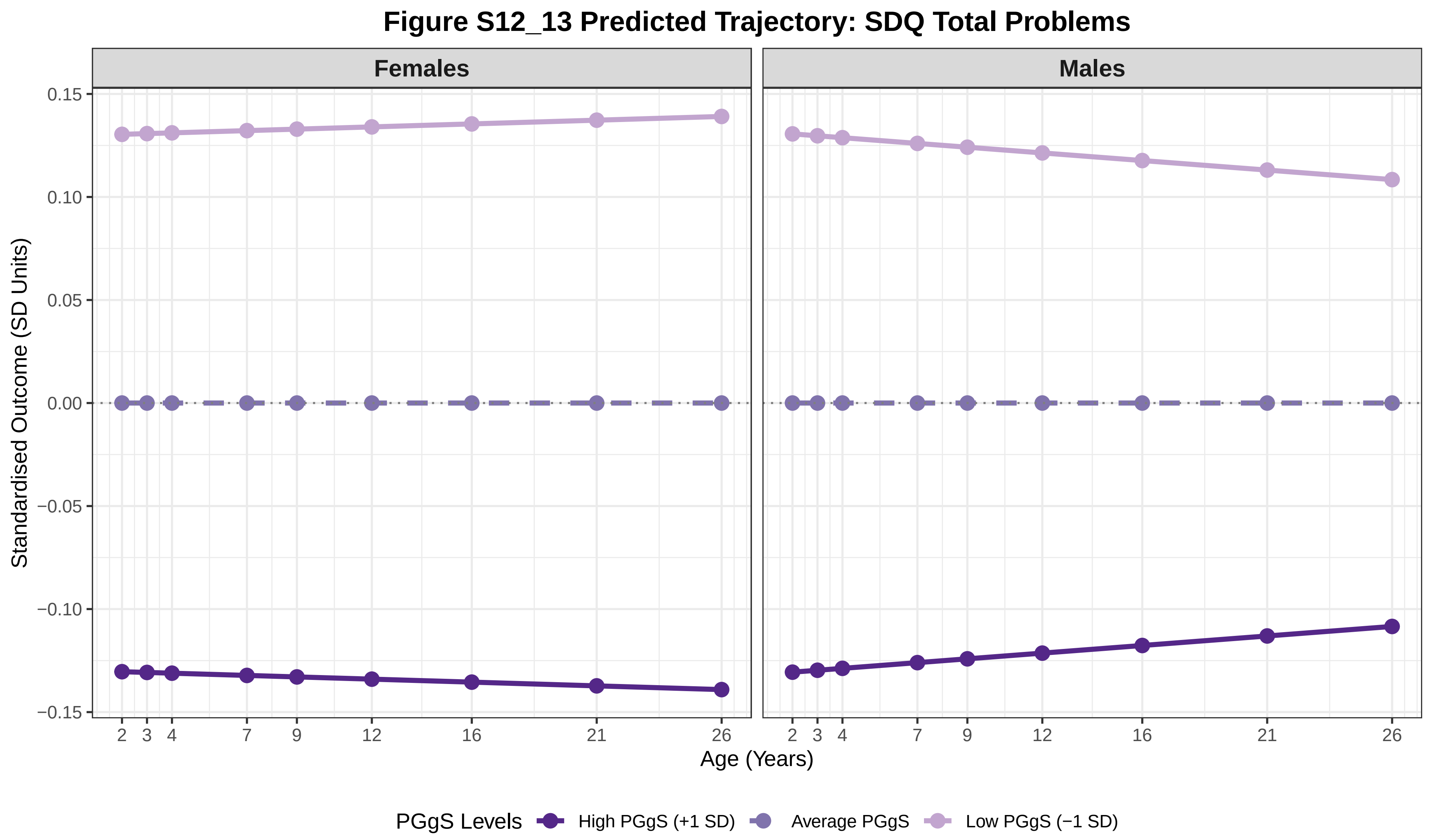

Figure S12_13 Predicted Trajectory: SDQ Total Problems
Females
Males
0.15
0.10
0.05
0.00
−0.05
−0.10
−0.15
Standardised Outcome (SD Units)
2 3 4 7 9 12 16 21 26
Age (Years)
PGgS Levels
High PGgS (+1 SD)
Average PGgS
Low PGgS (−1 SD)

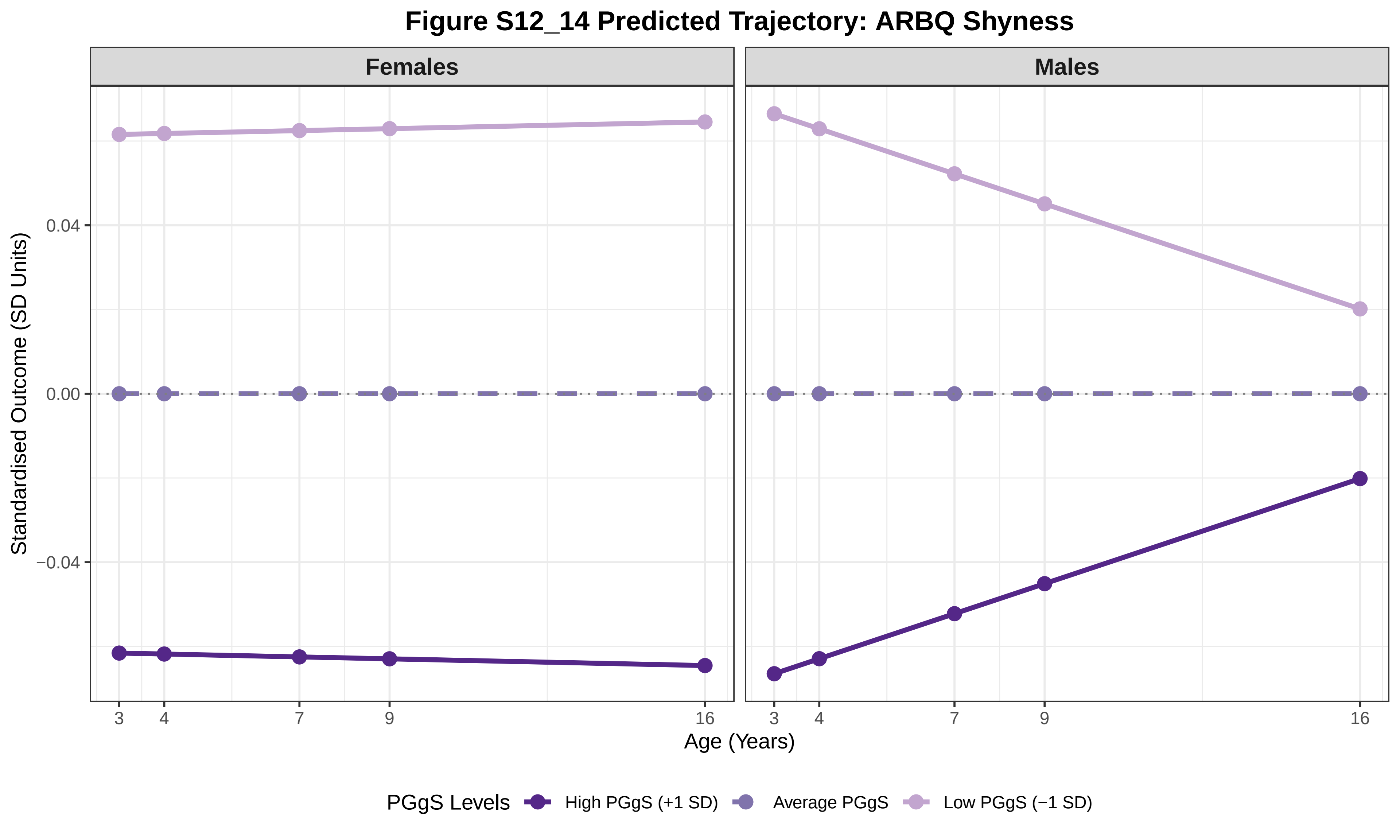
Figure S12_14 Predicted Trajectory: ARBQ Shyness
Females
Males
Standardised Outcome (SD Units)
0.04
0.00
−0.04
3
4
7
9
16
Age (Years)
PGgS Levels
High PGgS (+1 SD)
Average PGgS
Low PGgS (−1 SD)

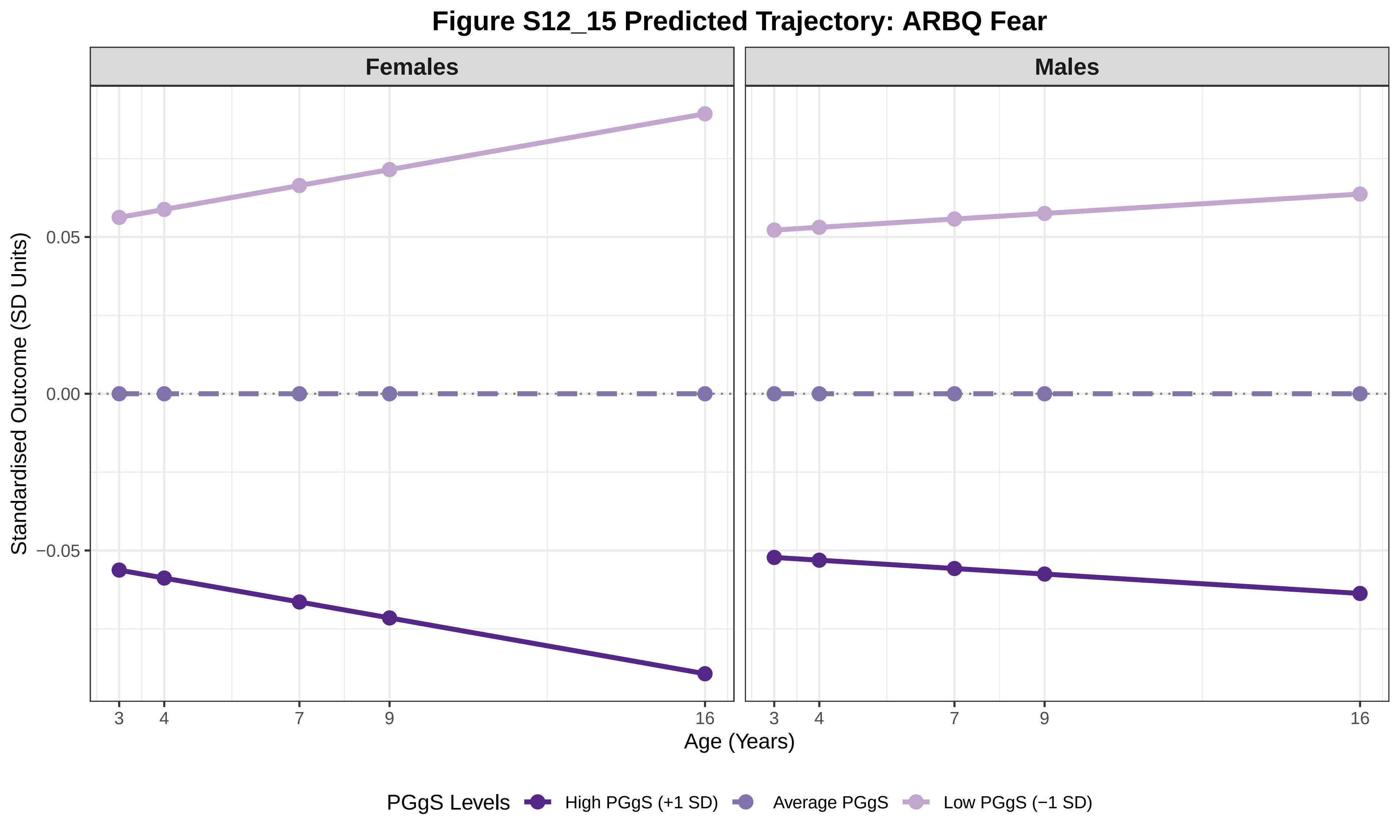

Figure S12_15 Predicted Trajectory: ARBQ Fear
Females
Males
Standardised Outcome (SD Units)
0.05
0.00
−0.05
3
4
7
9
16
Age (Years)
PGgS Levels
High PGgS (+1 SD)
Average PGgS
Low PGgS (−1 SD)

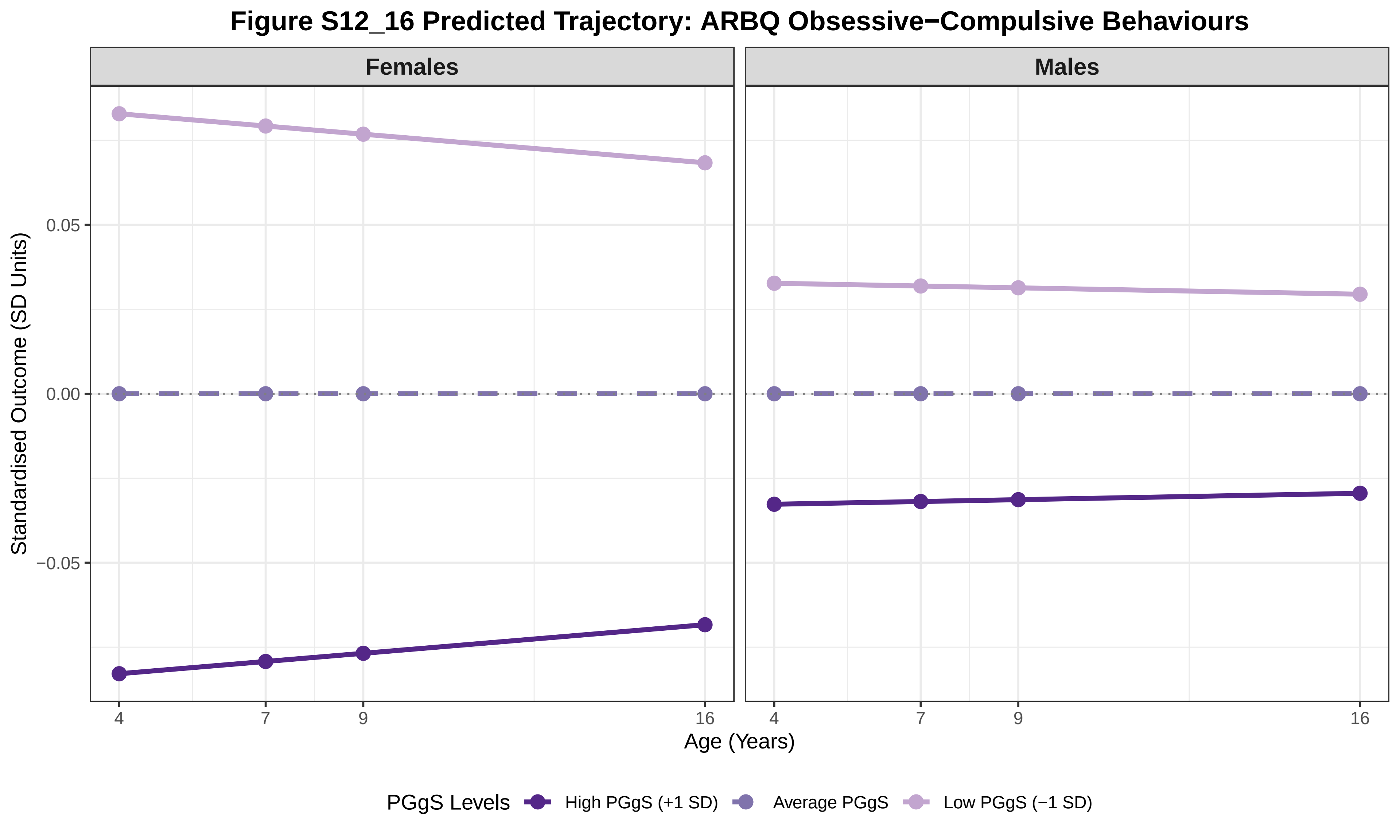

Figure S12_16 Predicted Trajectory: ARBQ Obsessive−Compulsive Behaviours
Females
Males
0.05
0.00
−0.05
Standardised Outcome (SD Units)
4
7
9
16
Age (Years)
PGgS Levels
High PGgS (+1 SD)
Average PGgS
Low PGgS (−1 SD)

### Figure S12_17 Predicted Trajectory: ARBQ Negative Affect

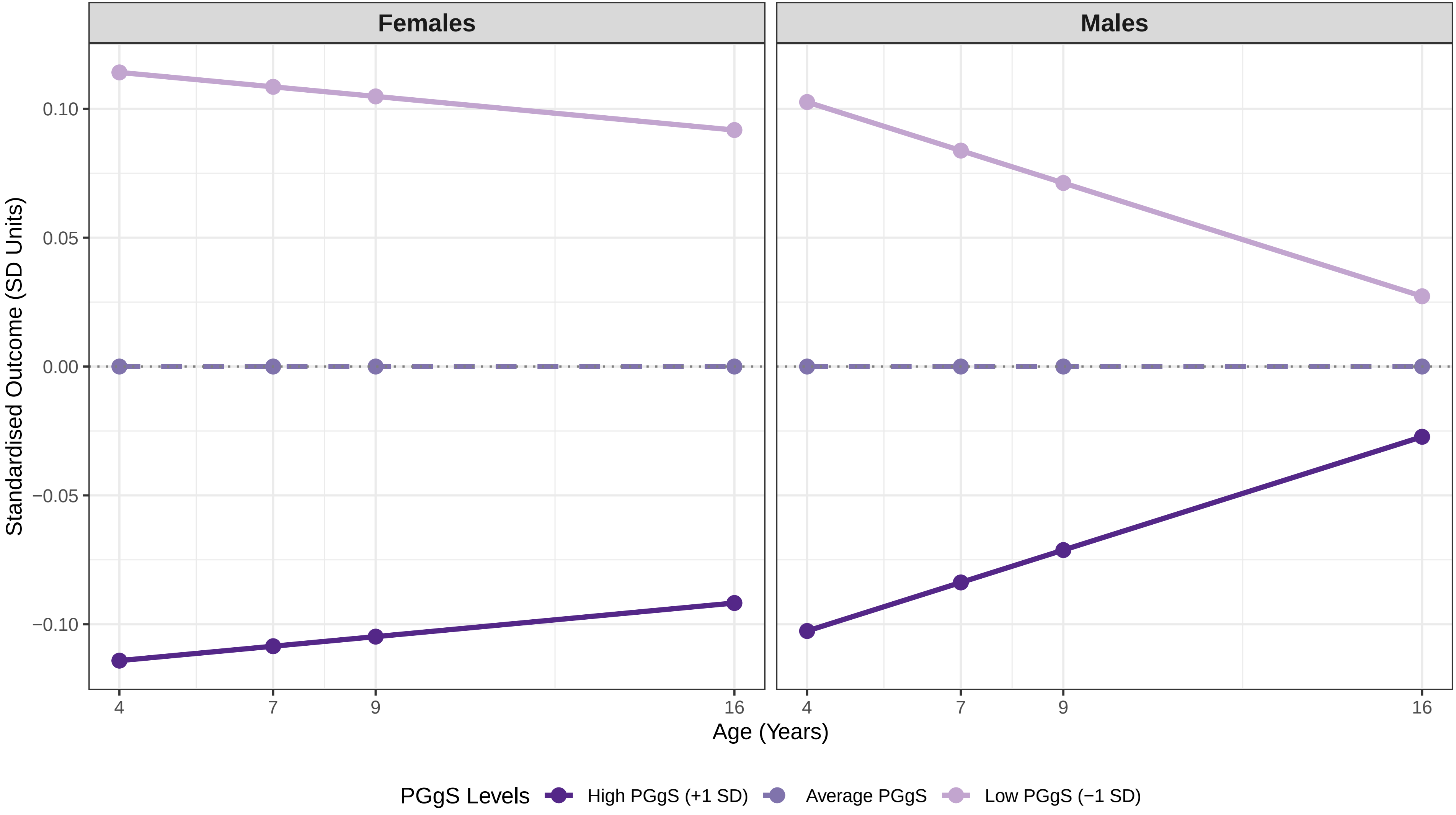

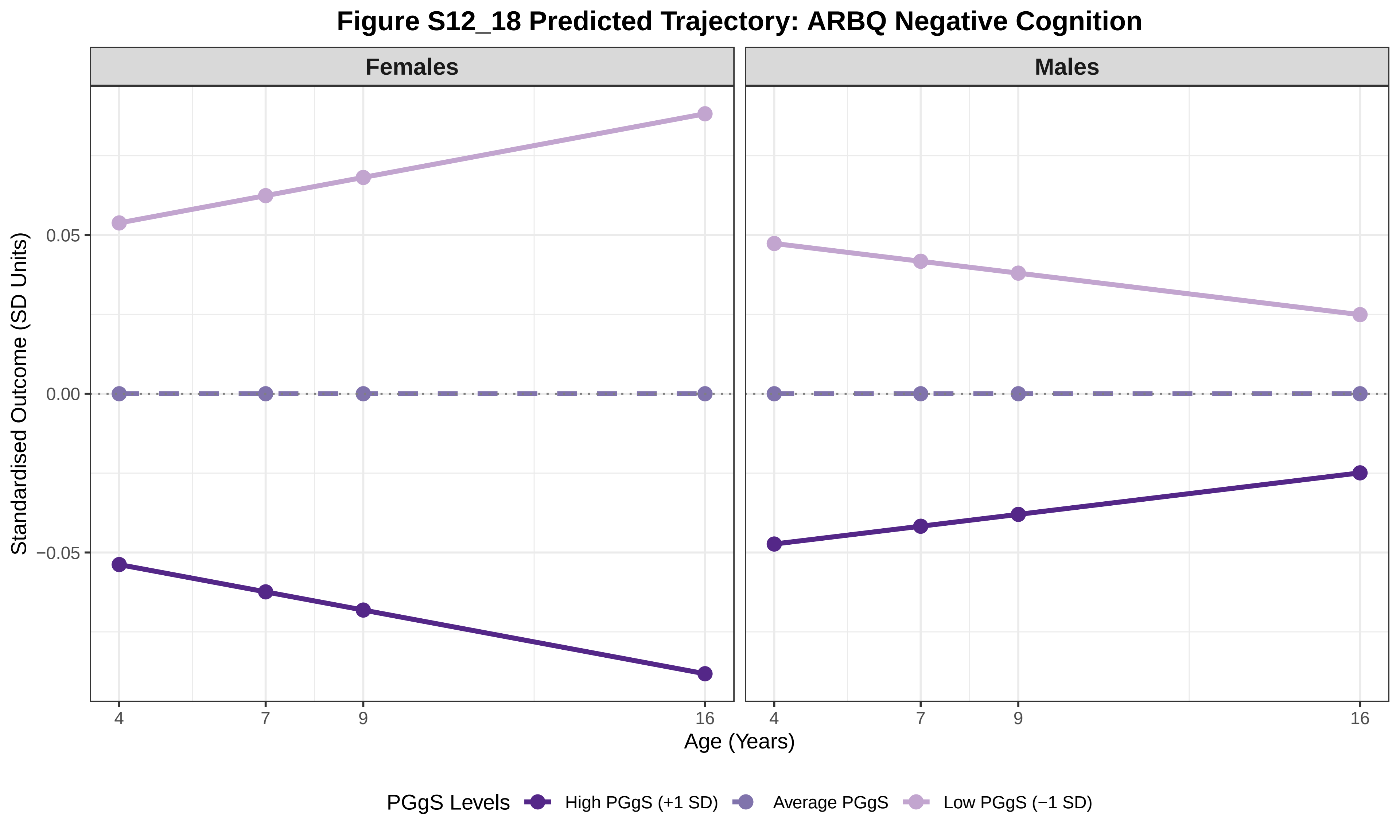

Figure S12_18 Predicted Trajectory: ARBQ Negative Cognition
Females
Males
Standardised Outcome (SD Units)
0.05
0.00
−0.05
4
7
9
16
Age (Years)
PGgS Levels
High PGgS (+1 SD)
Average PGgS
Low PGgS (−1 SD)

#### Figure S12_19 Predicted Trajectory: ARBQ Anxiety Total

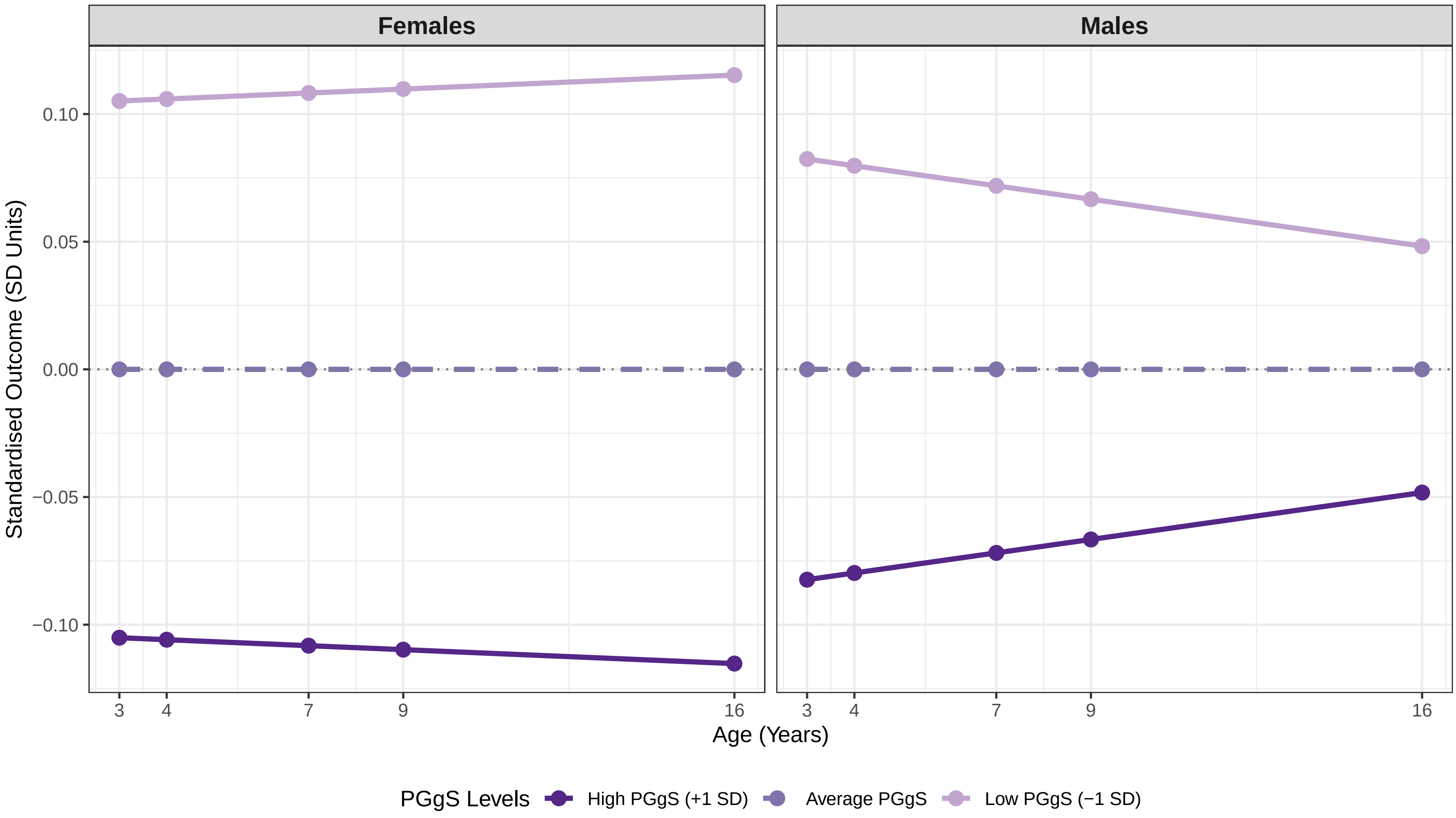

### Figure S12_20 Predicted Trajectory: Conners ADHD Inattention

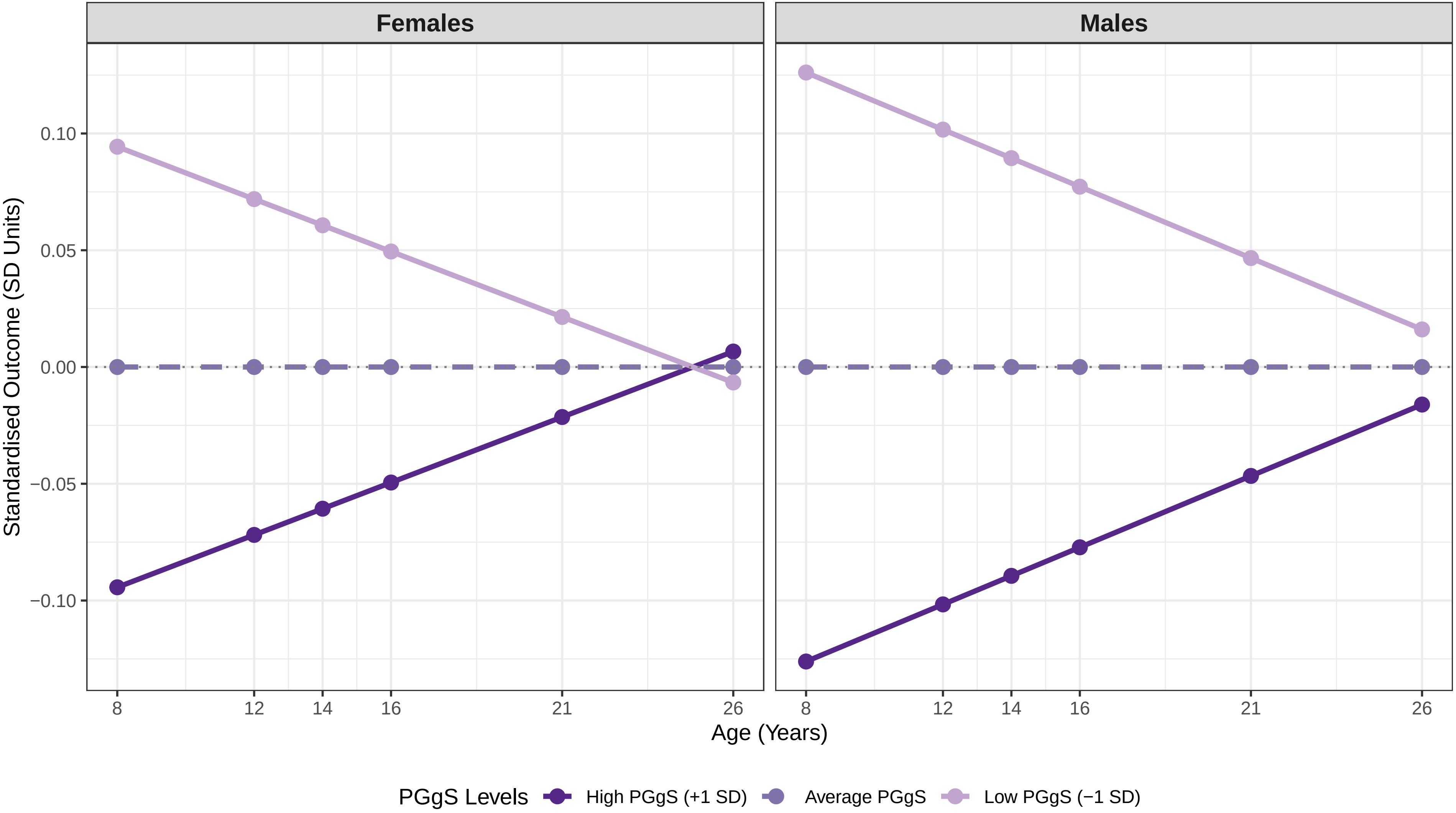

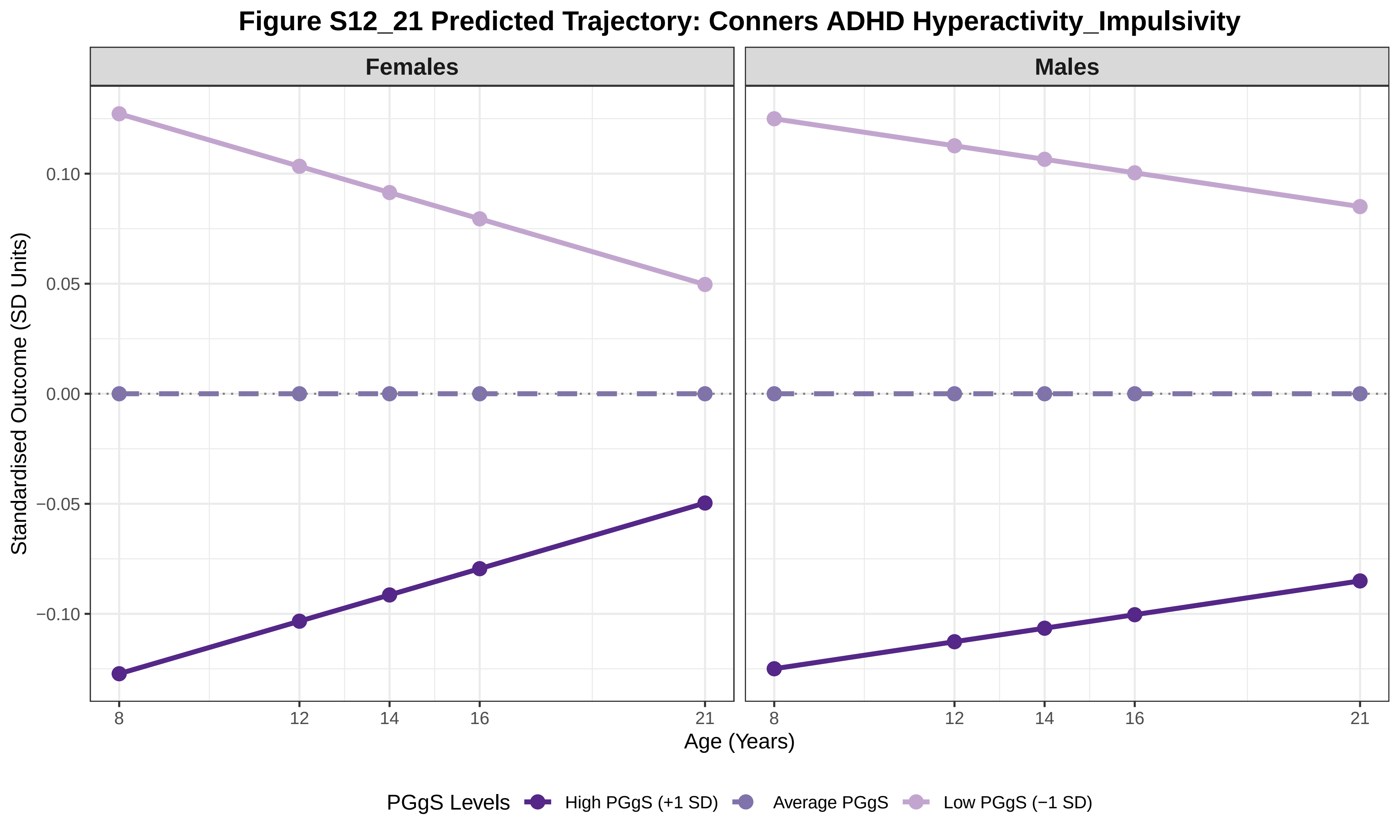

Figure S12_21 Predicted Trajectory: Conners ADHD Hyperactivity_Impulsivity
Females
Males
Standardised Outcome (SD Units)
0.10
0.05
0.00
−0.05
−0.10
8
12
14
16
21
Age (Years)
PGgS Levels
High PGgS (+1 SD)
Average PGgS
Low PGgS (−1 SD)

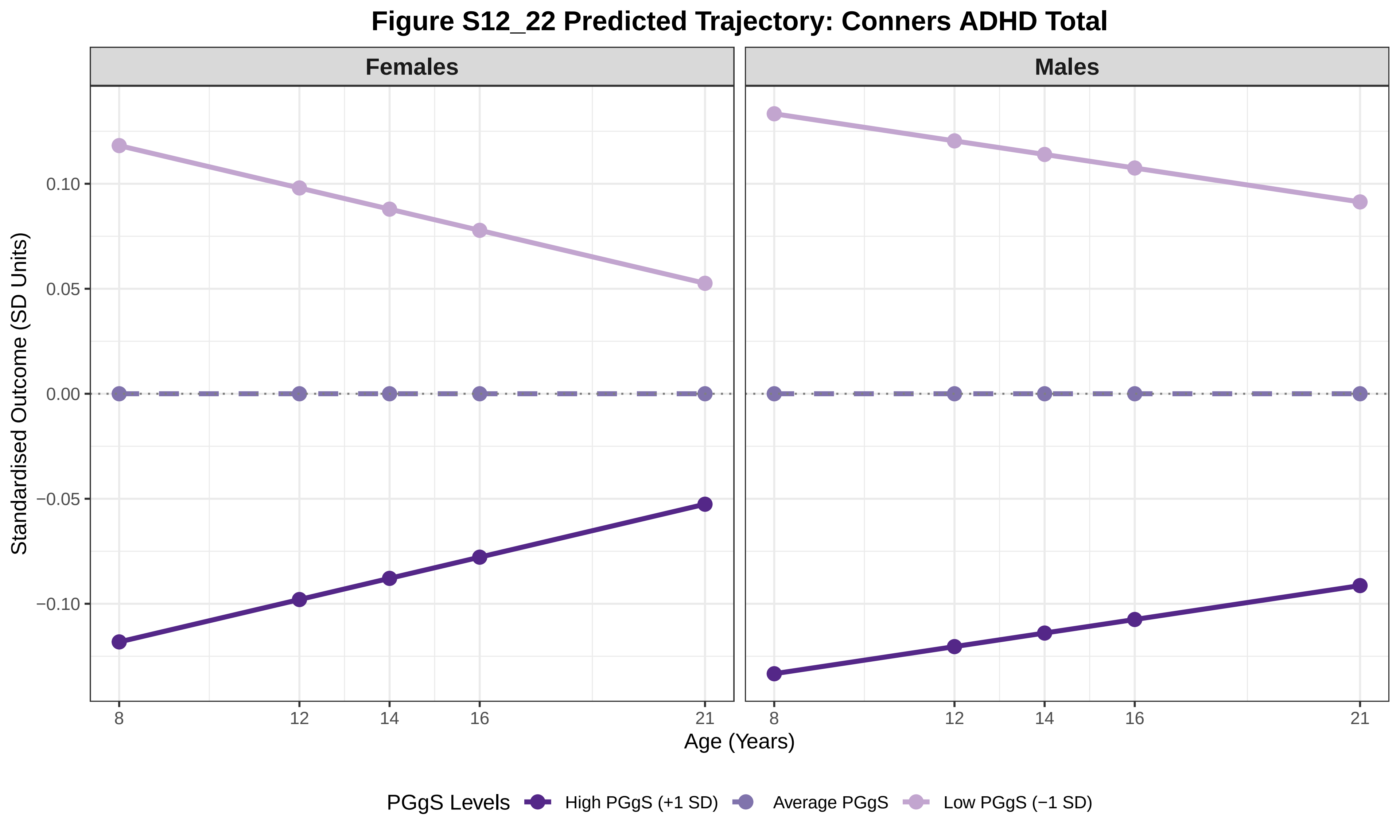

Figure S12_22 Predicted Trajectory: Conners ADHD Total
Females
Males
Standardised Outcome (SD Units)
0.10
0.05
0.00
−0.05
−0.10
8
12
14
16
21
Age (Years)
PGgS Levels
High PGgS (+1 SD)
Average PGgS
Low PGgS (−1 SD)

### Figure S12_23 Predicted Trajectory: Height

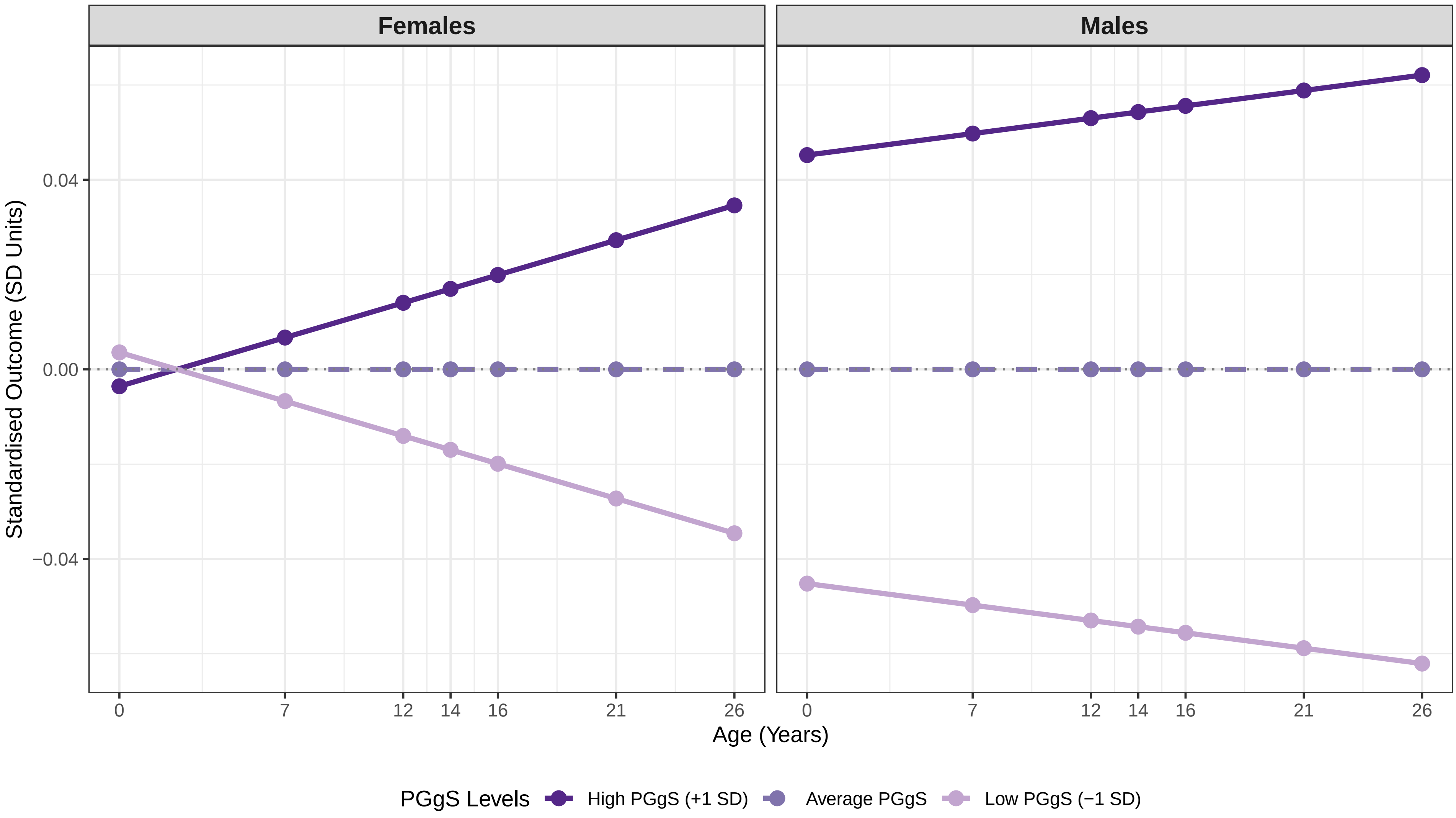

#### Figure S12_24 Predicted Trajectory: BMI (as weight at birth)

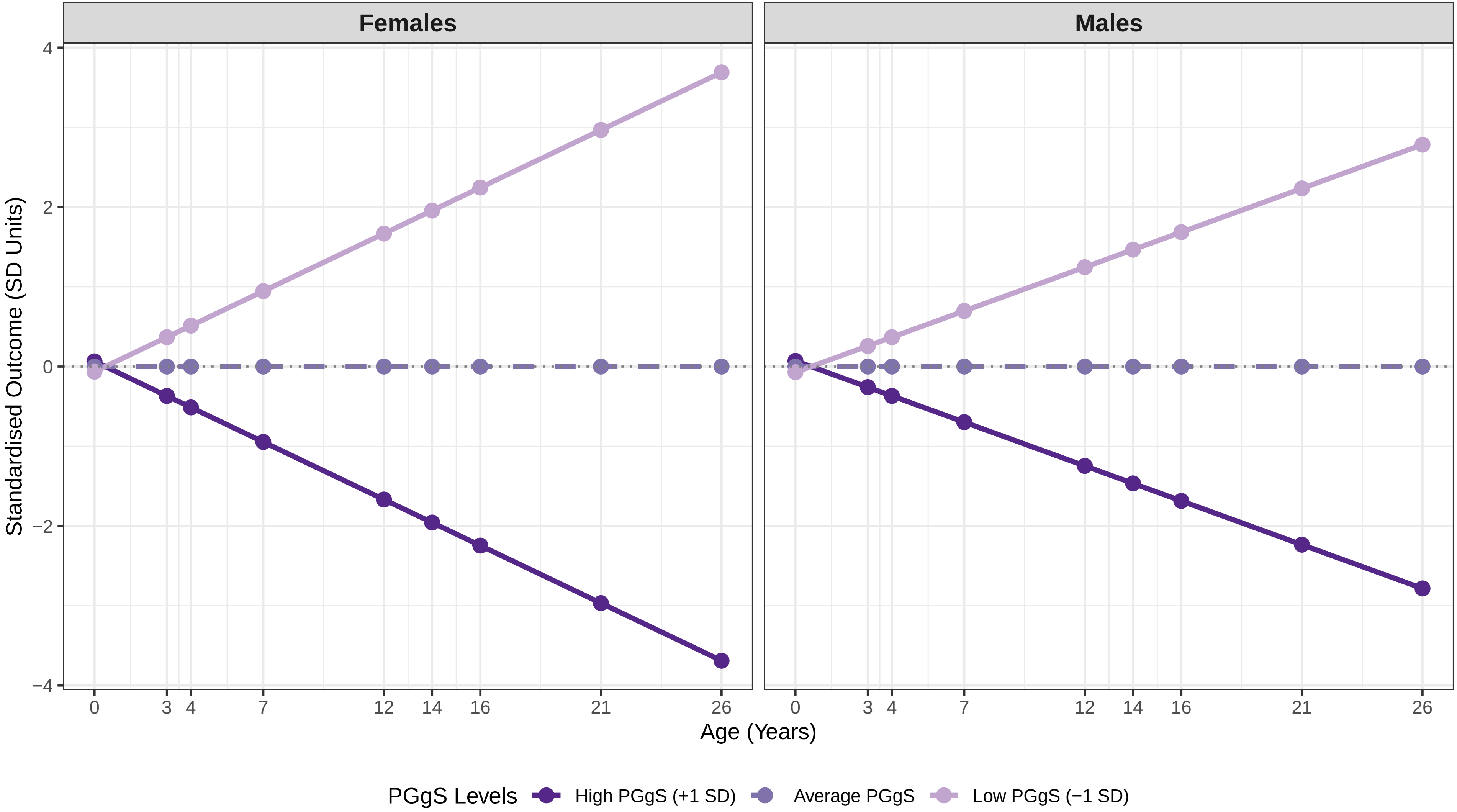

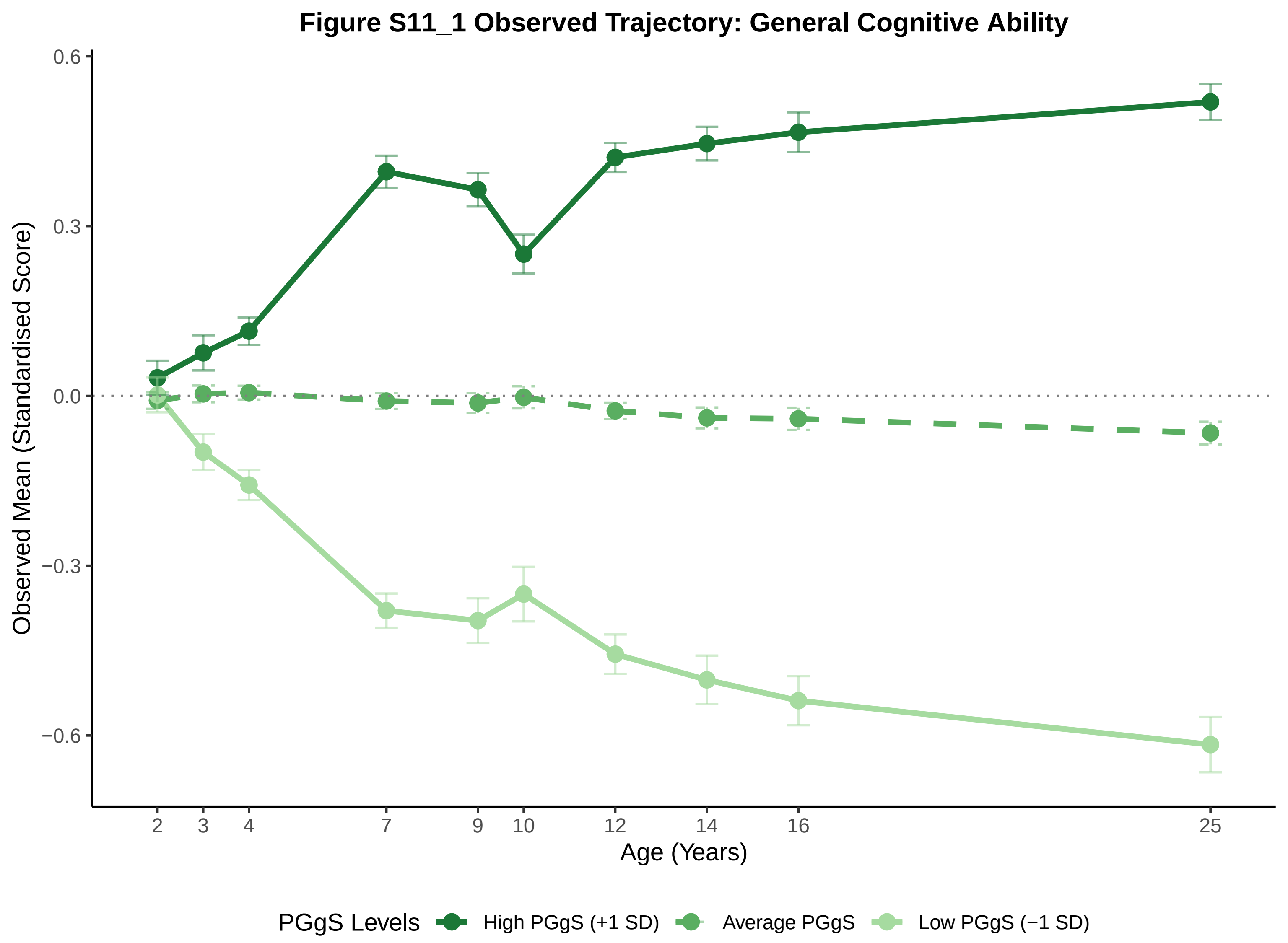
Figure S11_1 Observed Trajectory: General Cognitive Ability
Observed Mean (Standardised Score)
0.6
0.3
0.0
−0.3
−0.6
2
3
4
7
9
10
12
14
16
25
Age (Years)
PGgS Levels
High PGgS (+1 SD)
Average PGgS
Low PGgS (−1 SD)

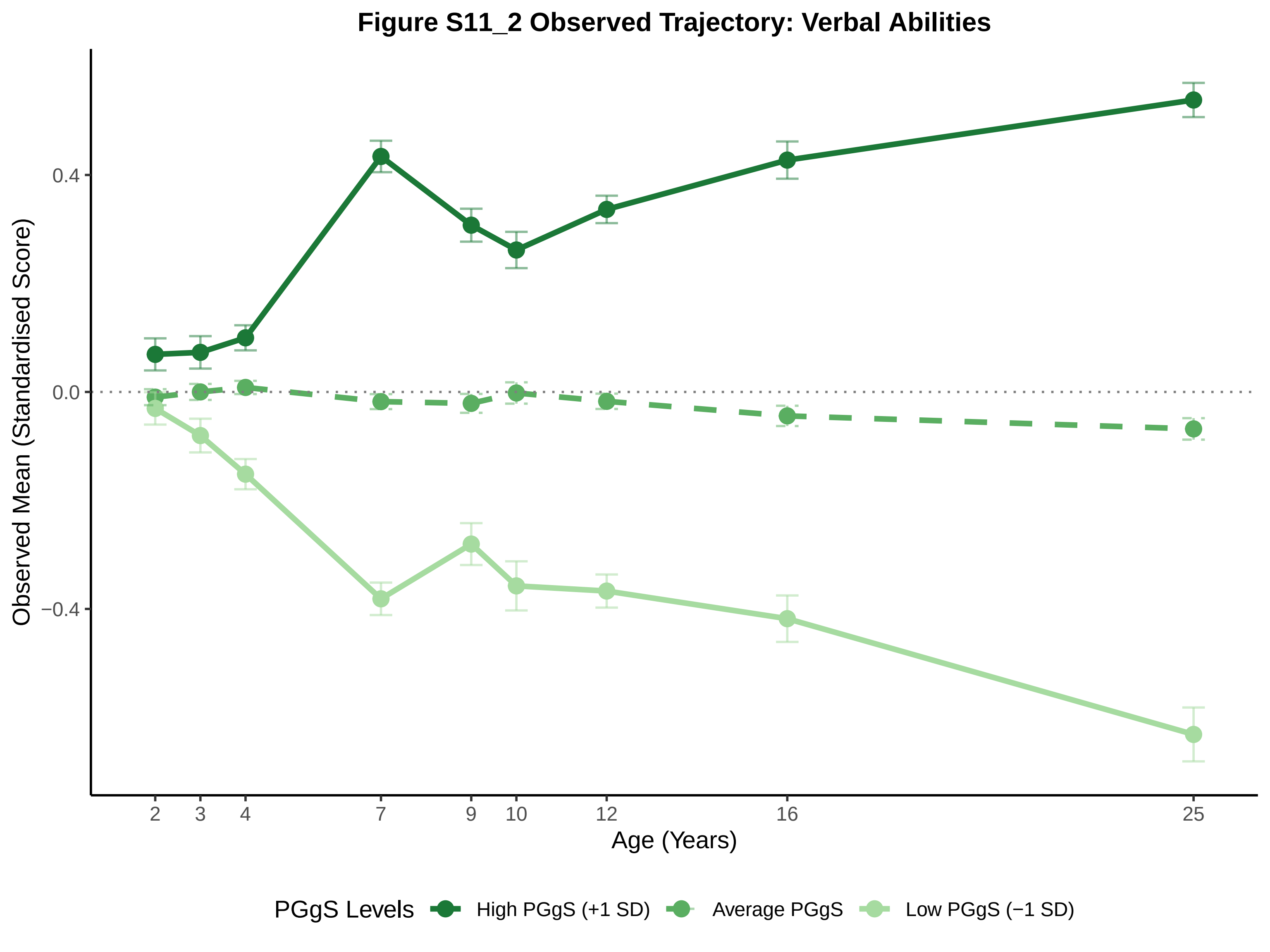
Figure S11_2 Observed Trajectory: Verbal Abilities
Observed Mean (Standardised Score)
0.4
0.0
−0.4
2
3
4
7
9
10
12
16
25
Age (Years)
PGgS Levels
High PGgS (+1 SD)
Average PGgS
Low PGgS (−1 SD)

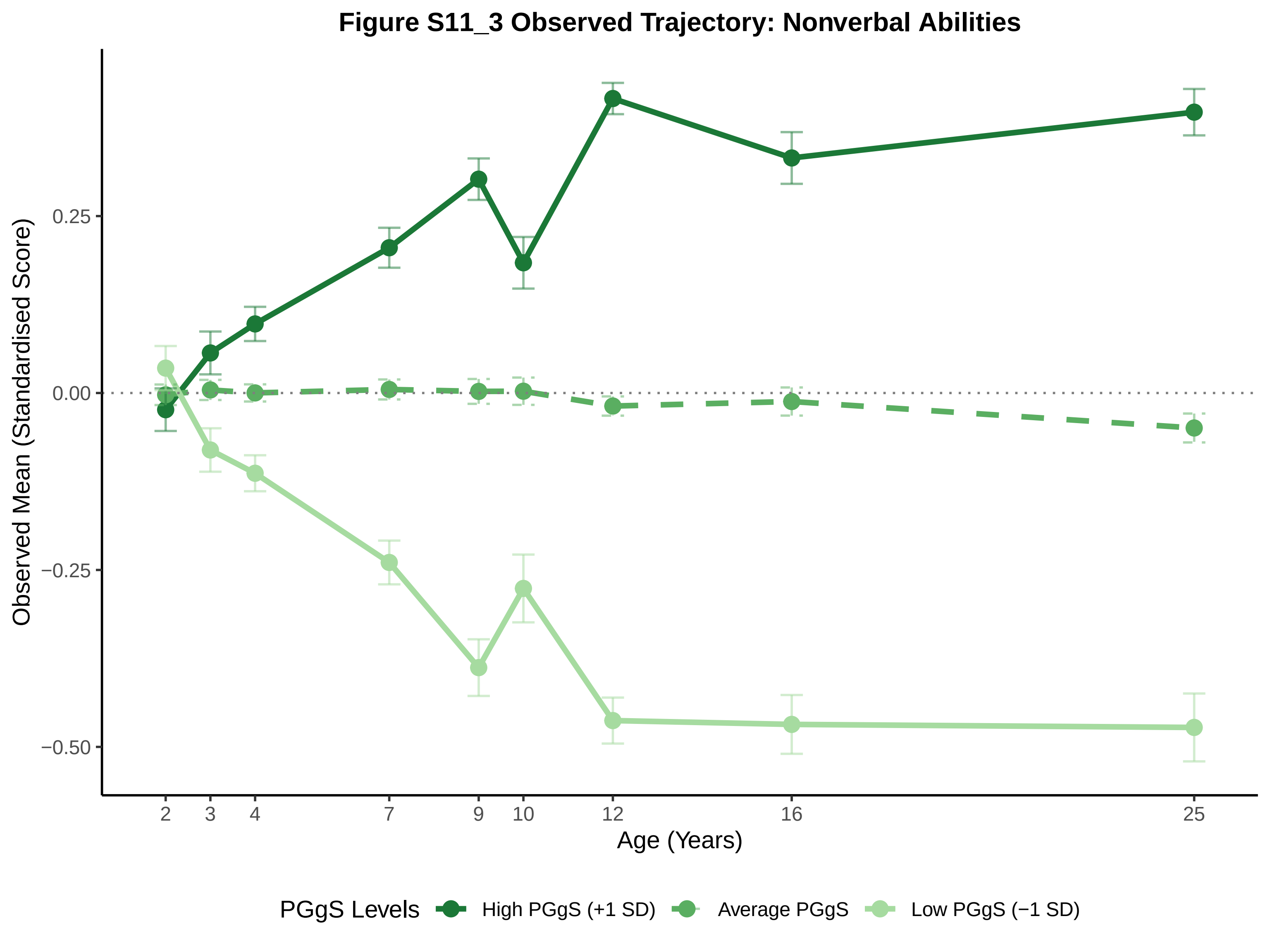
Figure S11_3 Observed Trajectory: Nonverbal Abilities
Observed Mean (Standardised Score)
0.25
0.00
−0.25
−0.50
2
3
4
7
9
10
12
16
25
Age (Years)
PGgS Levels
High PGgS (+1 SD)
Average PGgS
Low PGgS (−1 SD)

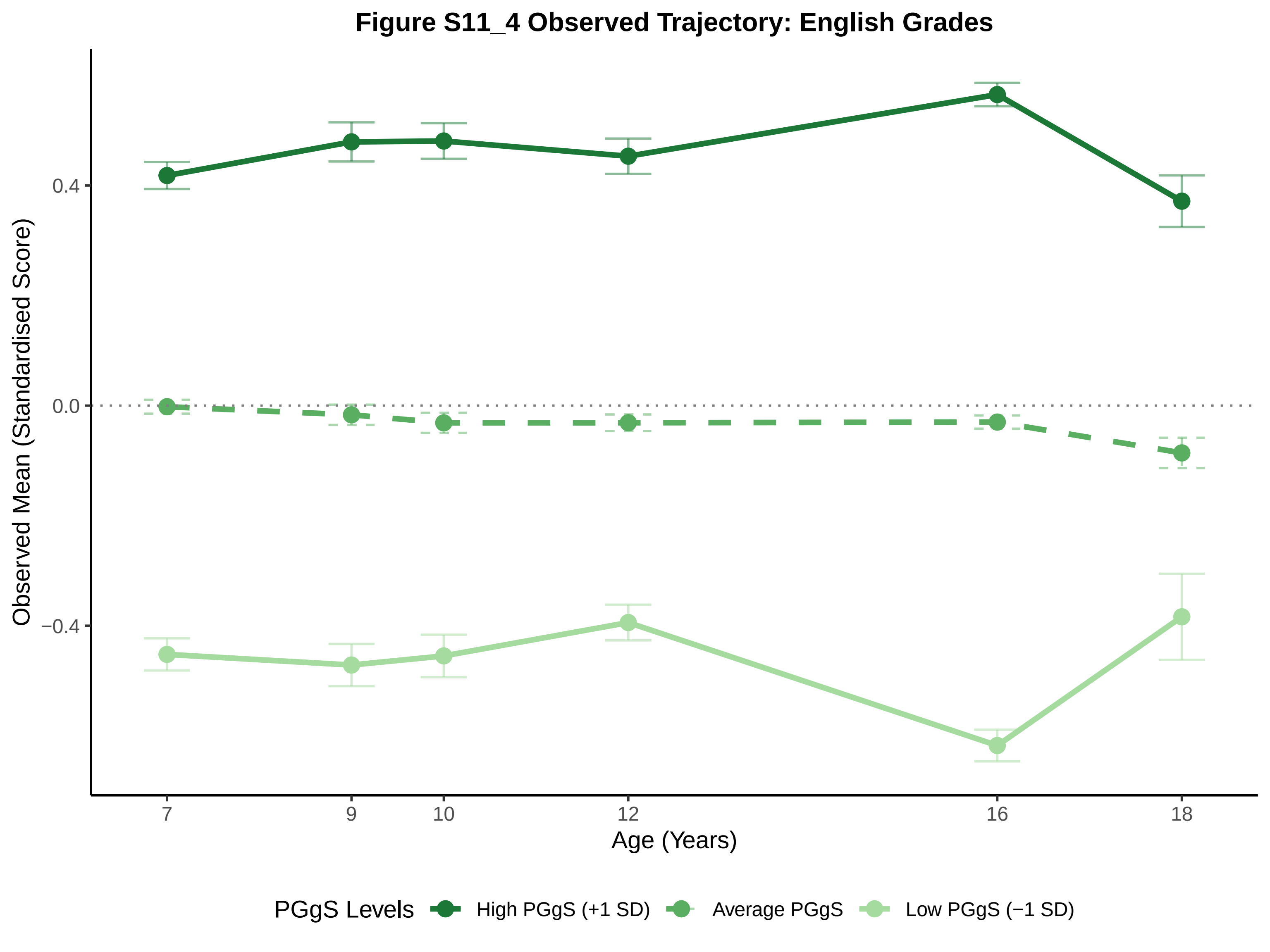
Figure S11_4 Observed Trajectory: English Grades
Observed Mean (Standardised Score)
0.4
0.0
−0.4
7
9
10
12
16
18
Age (Years)
PGgS Levels
High PGgS (+1 SD)
Average PGgS
Low PGgS (−1 SD)

### Figure S11_5 Observed Trajectory: Maths Grades

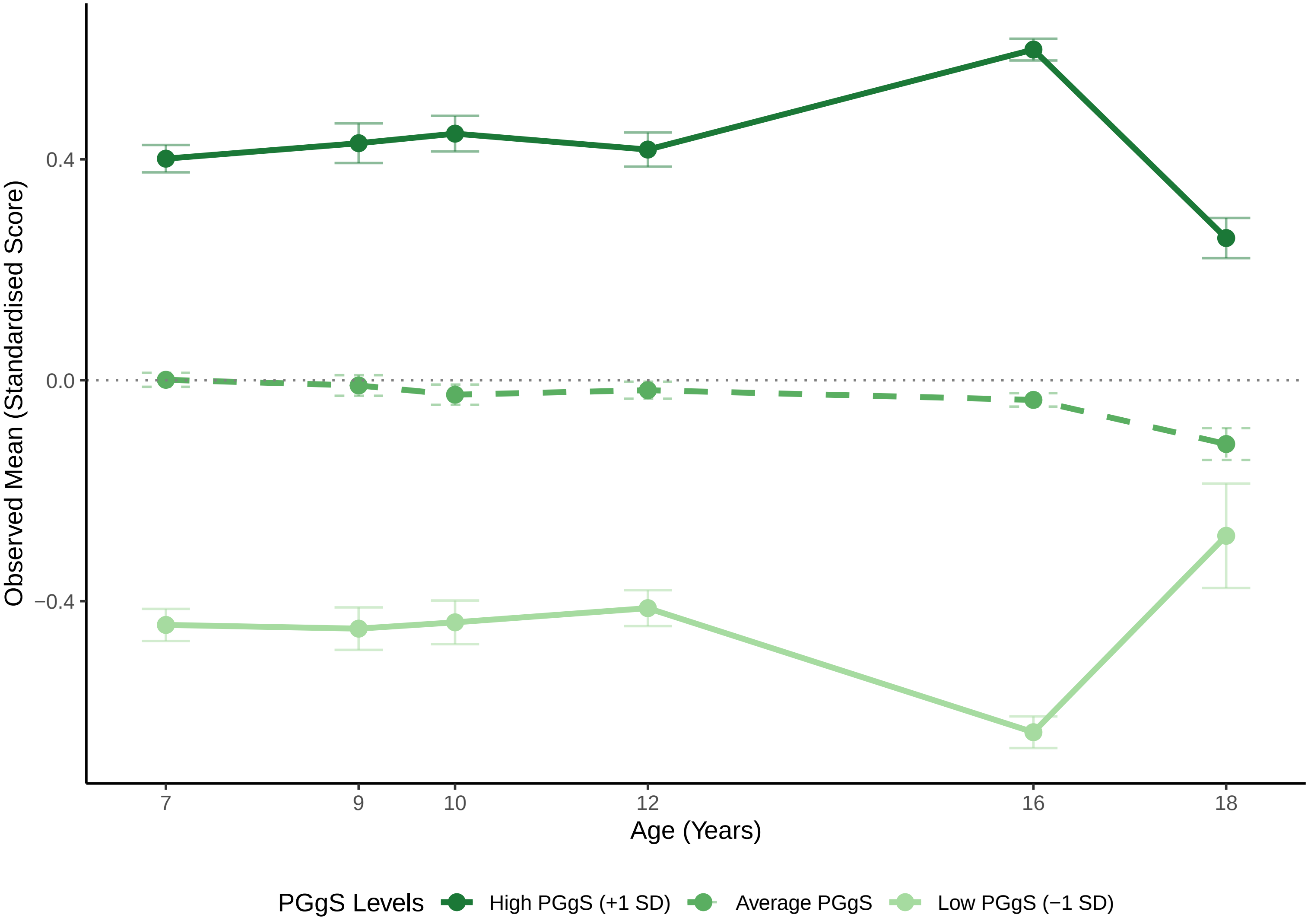

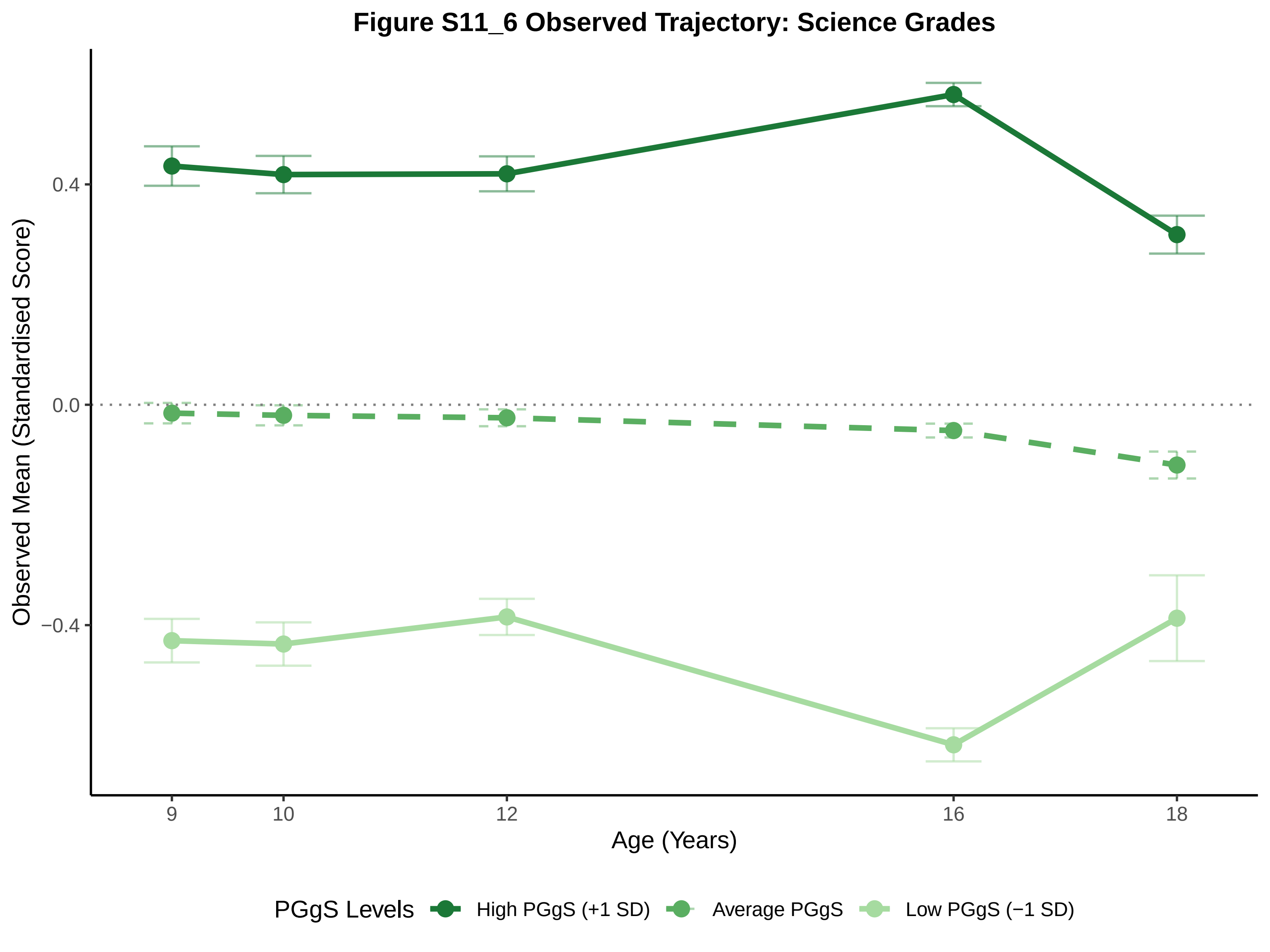
Figure S11_6 Observed Trajectory: Science Grades
Observed Mean (Standardised Score)
0.4
0.0
−0.4
9
10
12
16
18
Age (Years)
PGgS Levels
High PGgS (+1 SD)
Average PGgS
Low PGgS (−1 SD)

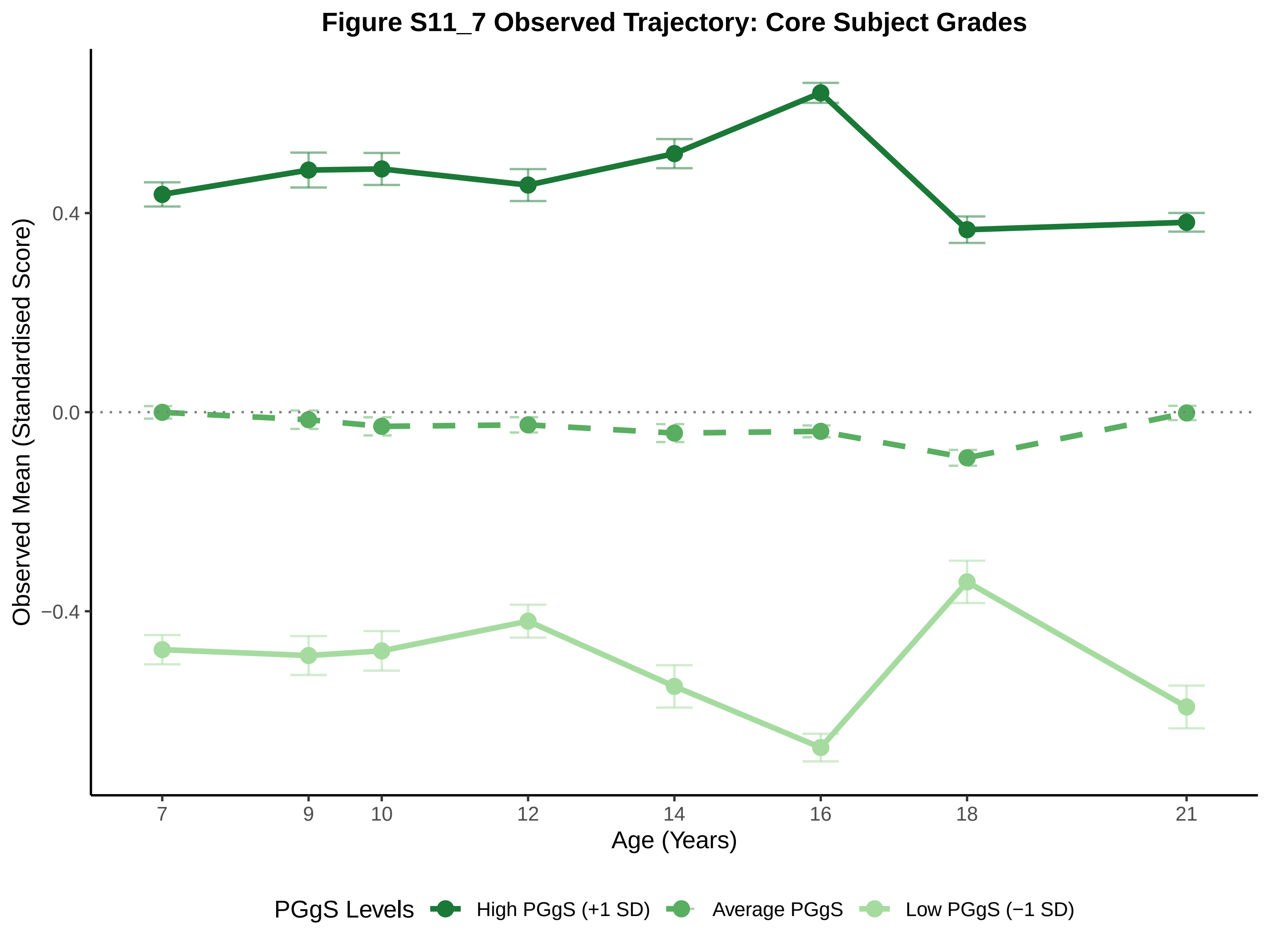
Figure S11_7 Observed Trajectory: Core Subject Grades
Observed Mean (Standardised Score)
0.4
0.0
−0.4
7
9
10
12
14
16
18
21
Age (Years)
PGgS Levels
High PGgS (+1 SD)
Average PGgS
Low PGgS (−1 SD)

#### Figure S11_8 Observed Trajectory: SDQ Conduct

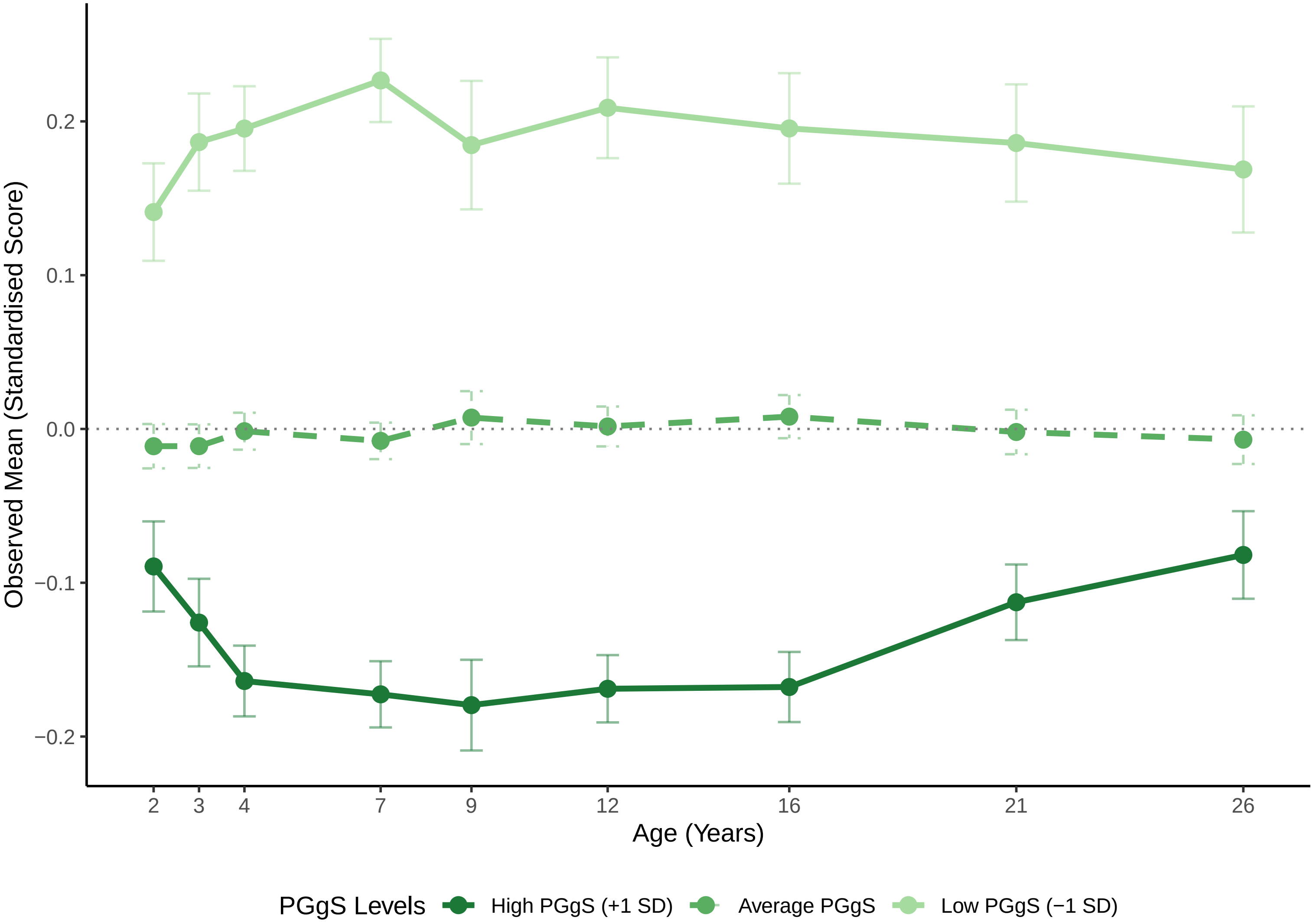

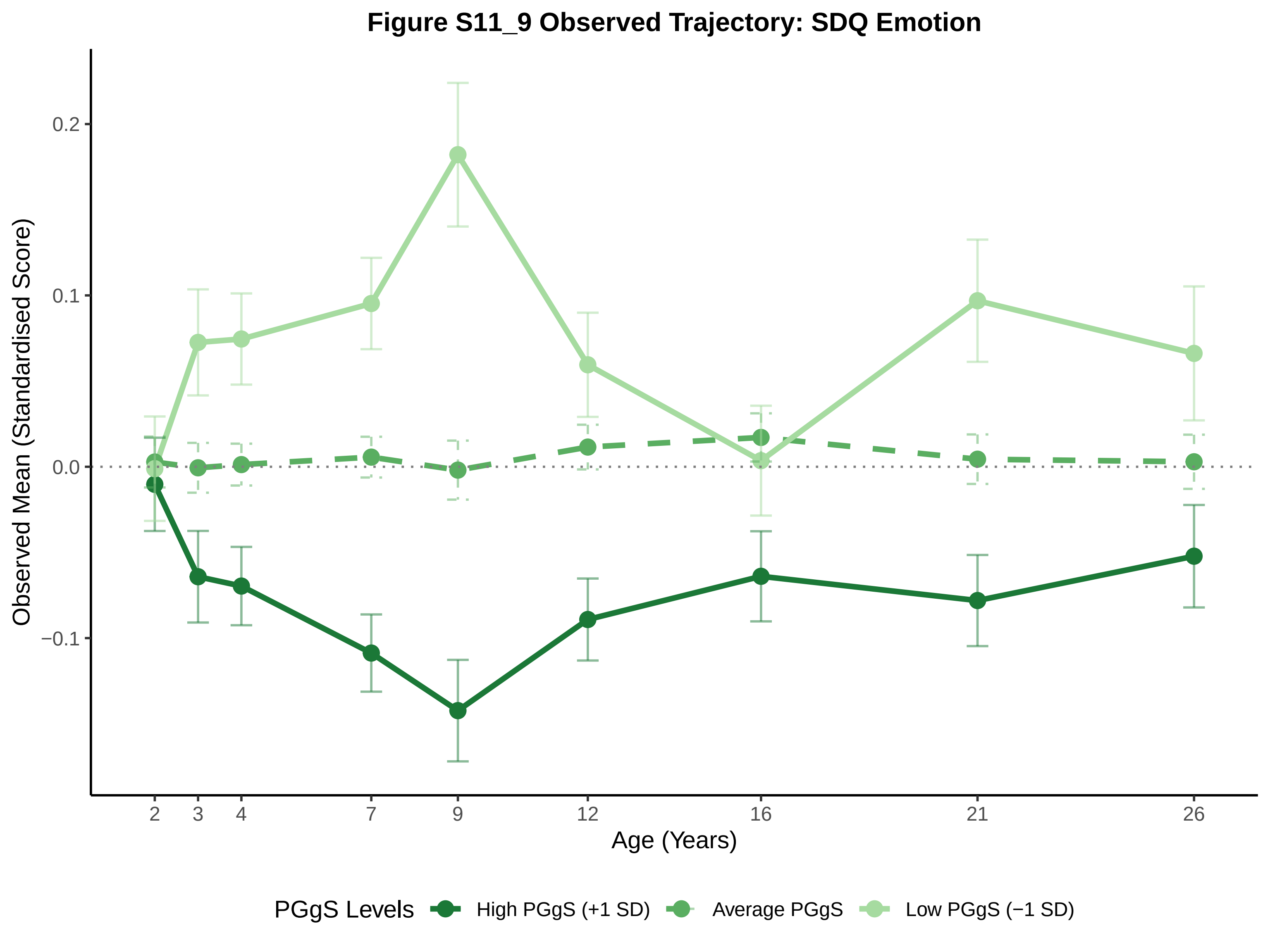

Figure S11_9 Observed Trajectory: SDQ Emotion
Observed Mean (Standardised Score)
0.2
0.1
0.0
−0.1
2
3
4
7
9
12
16
21
26
Age (Years)
PGgS Levels
High PGgS (+1 SD)
Average PGgS
Low PGgS (−1 SD)

**Figure S11_10 Observed Trajectory: SDQ Hyperactivity_SDQ**

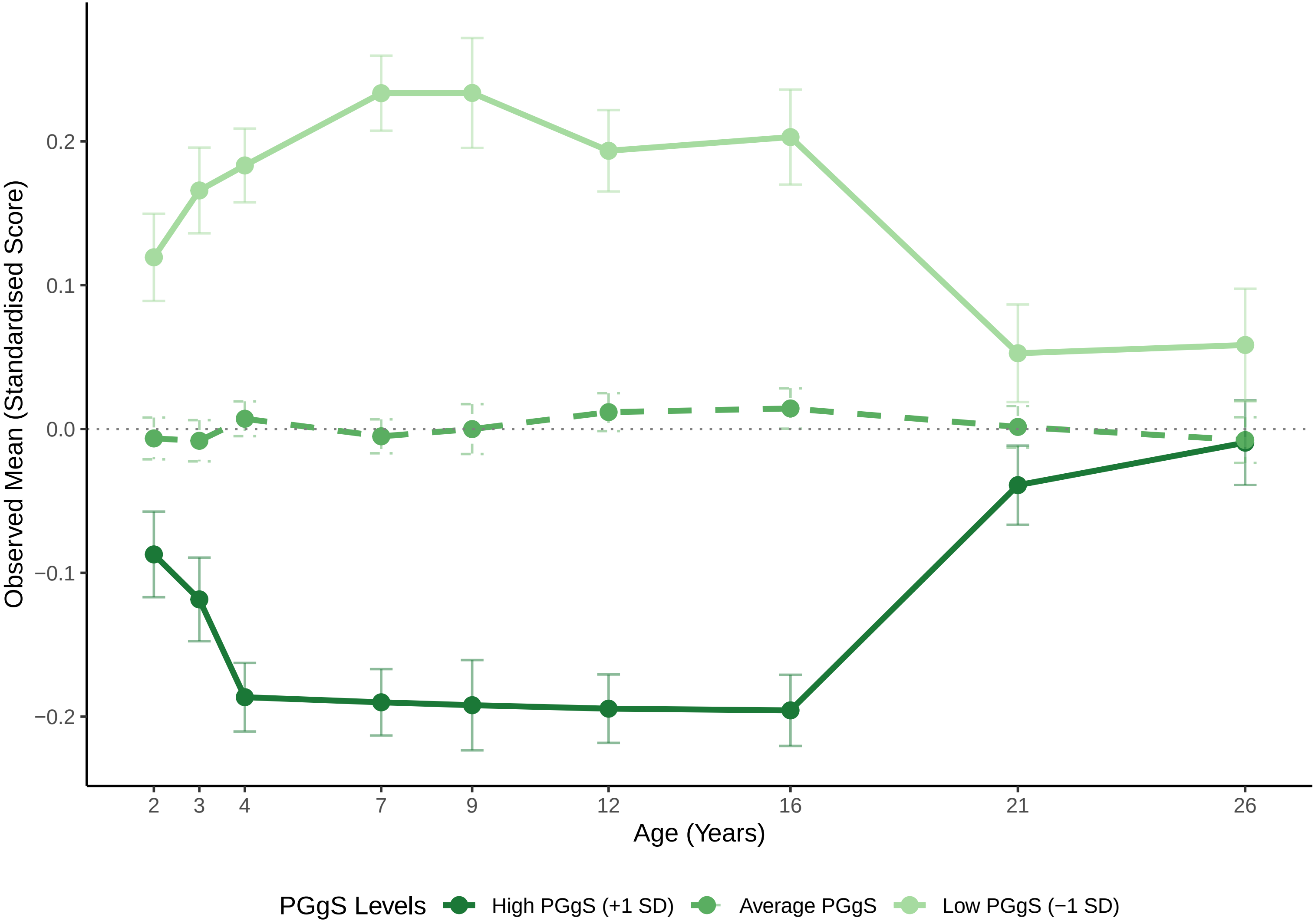

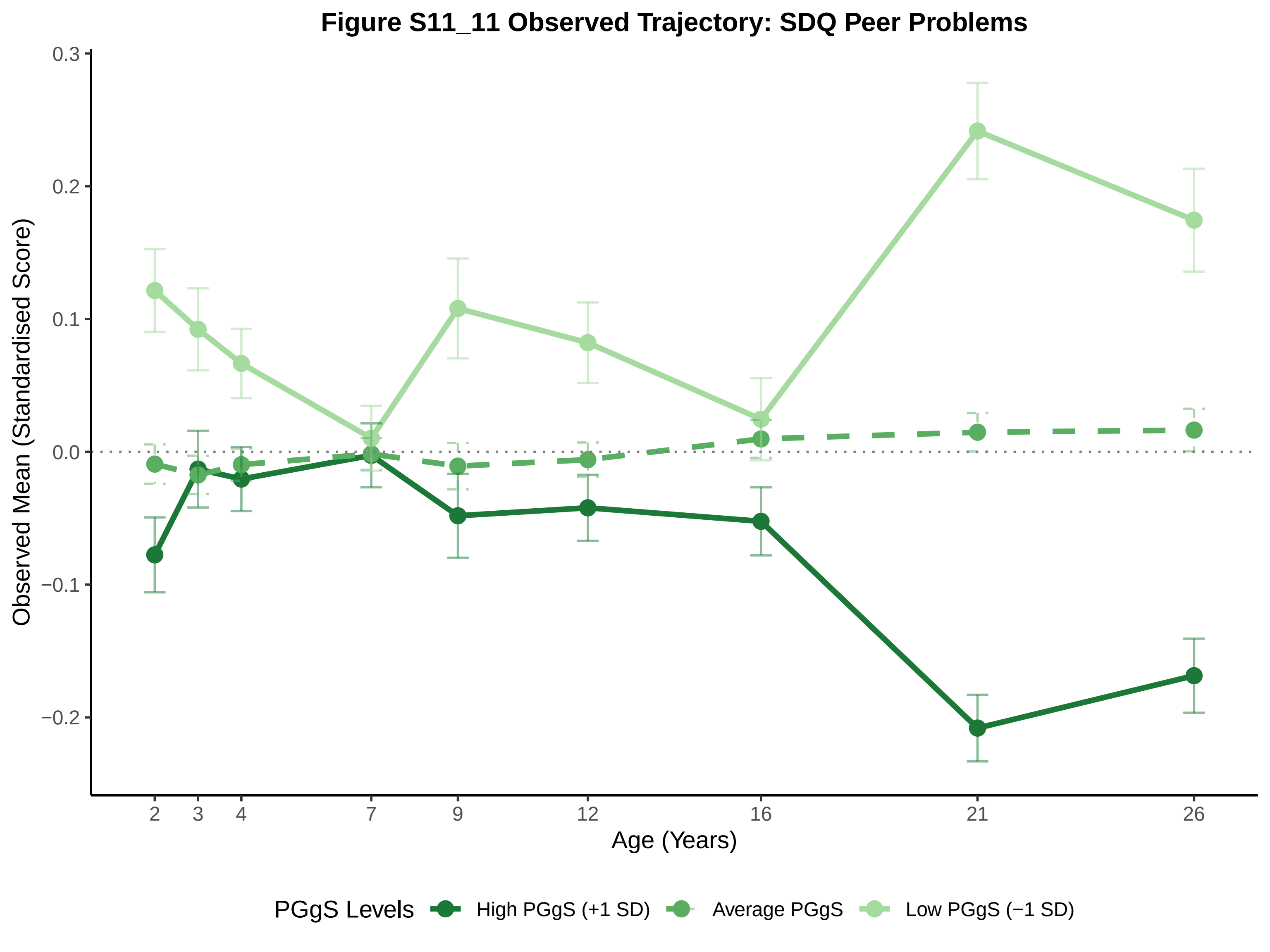

Figure S11_11 Observed Trajectory: SDQ Peer Problems
Observed Mean (Standardised Score)
0.3
0.2
0.1
0.0
−0.1
−0.2
2
3
4
7
9
12
16
21
26
Age (Years)
PGgS Levels
High PGgS (+1 SD)
Average PGgS
Low PGgS (−1 SD)

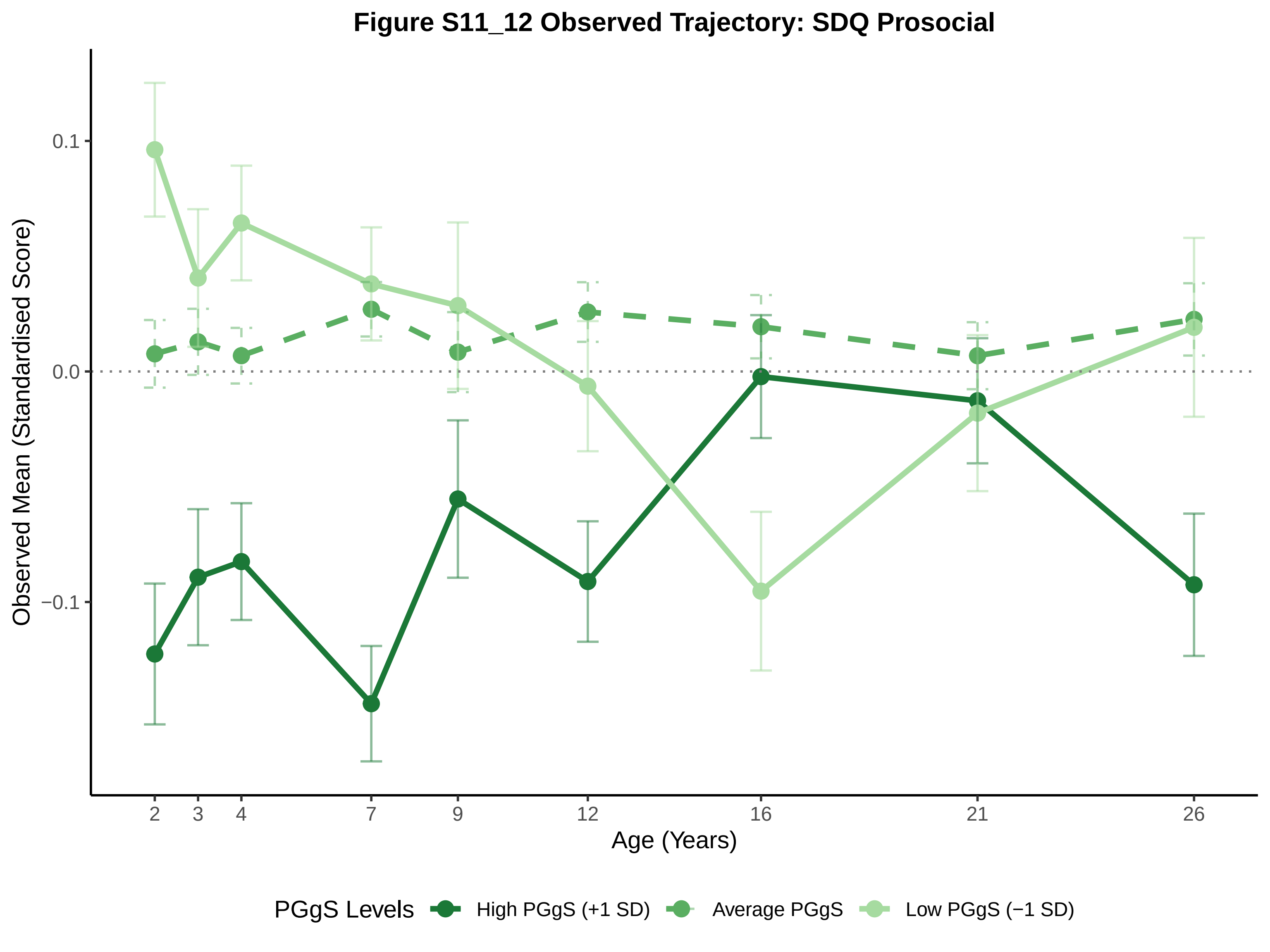

Figure S11_12 Observed Trajectory: SDQ Prosocial
Observed Mean (Standardised Score)
0.1
0.0
−0.1
2
3
4
7
9
12
16
21
26
Age (Years)
PGgS Levels
High PGgS (+1 SD)
Average PGgS
Low PGgS (−1 SD)

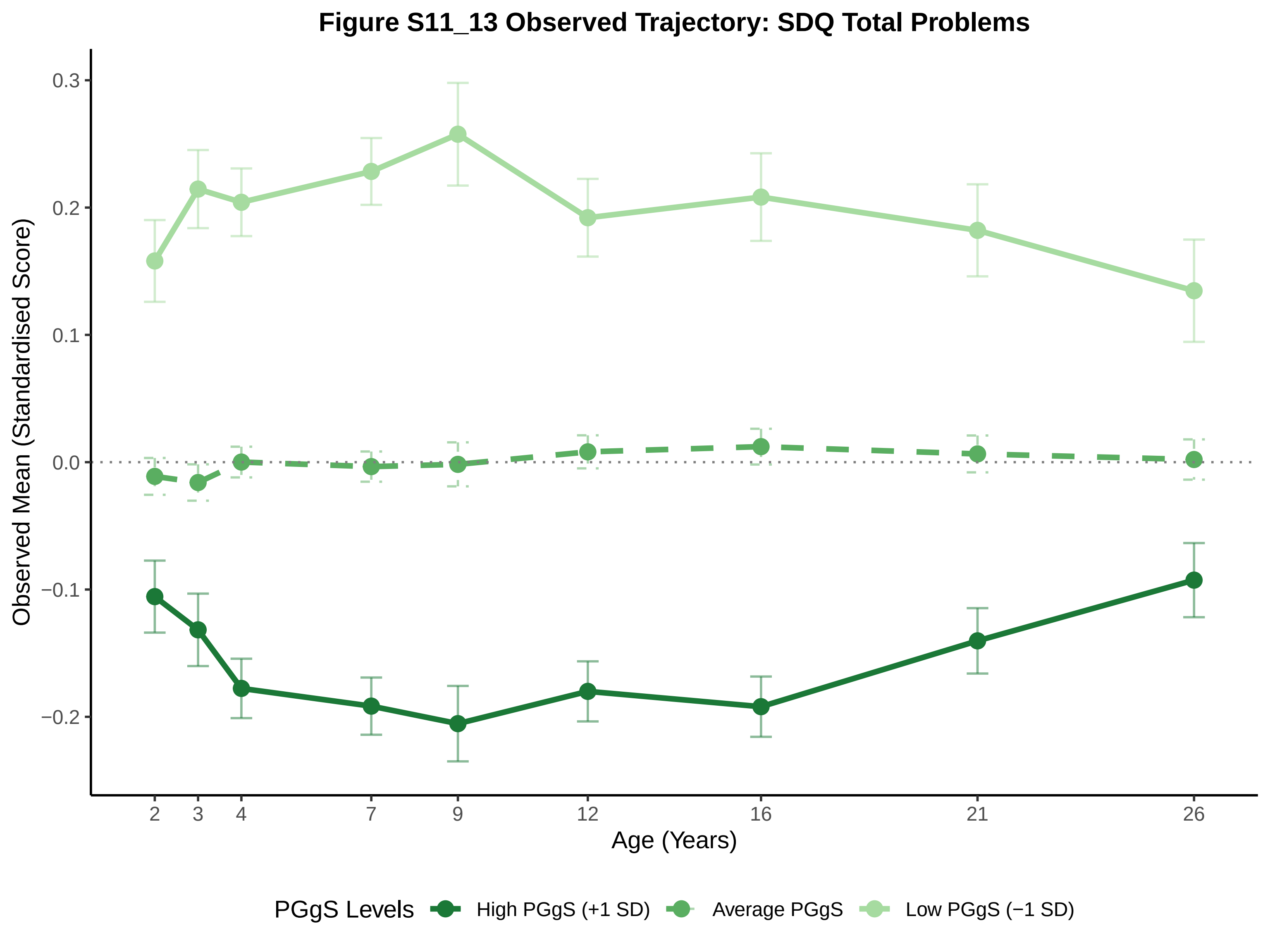
Figure S11_13 Observed Trajectory: SDQ Total Problems
Observed Mean (Standardised Score)
0.3
0.2
0.1
0.0
−0.1
−0.2
2
3
4
7
9
12
16
21
26
Age (Years)
PGgS Levels
High PGgS (+1 SD)
Average PGgS
Low PGgS (−1 SD)

### Figure S11_14 Observed Trajectory: ARBQ Shyness

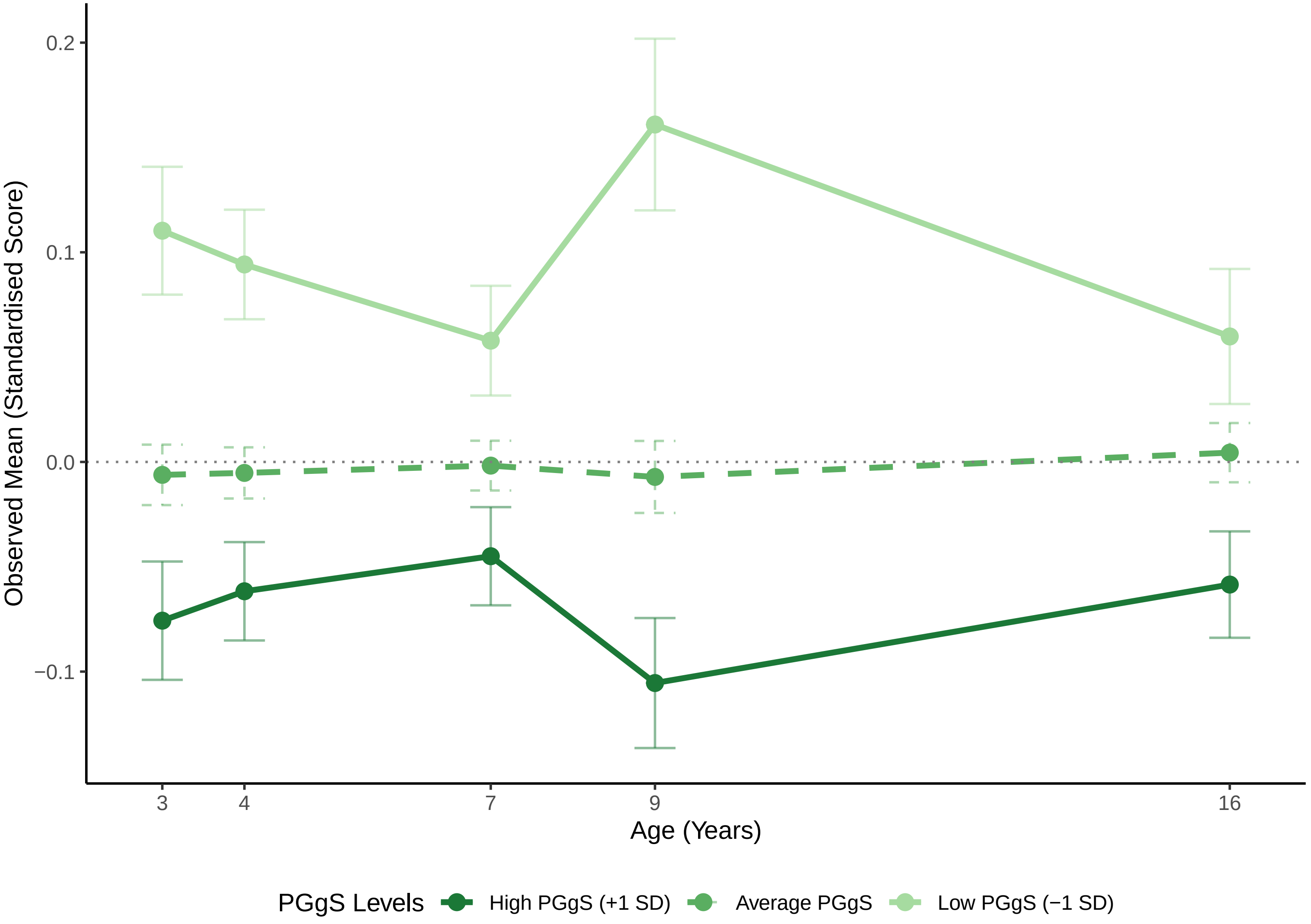

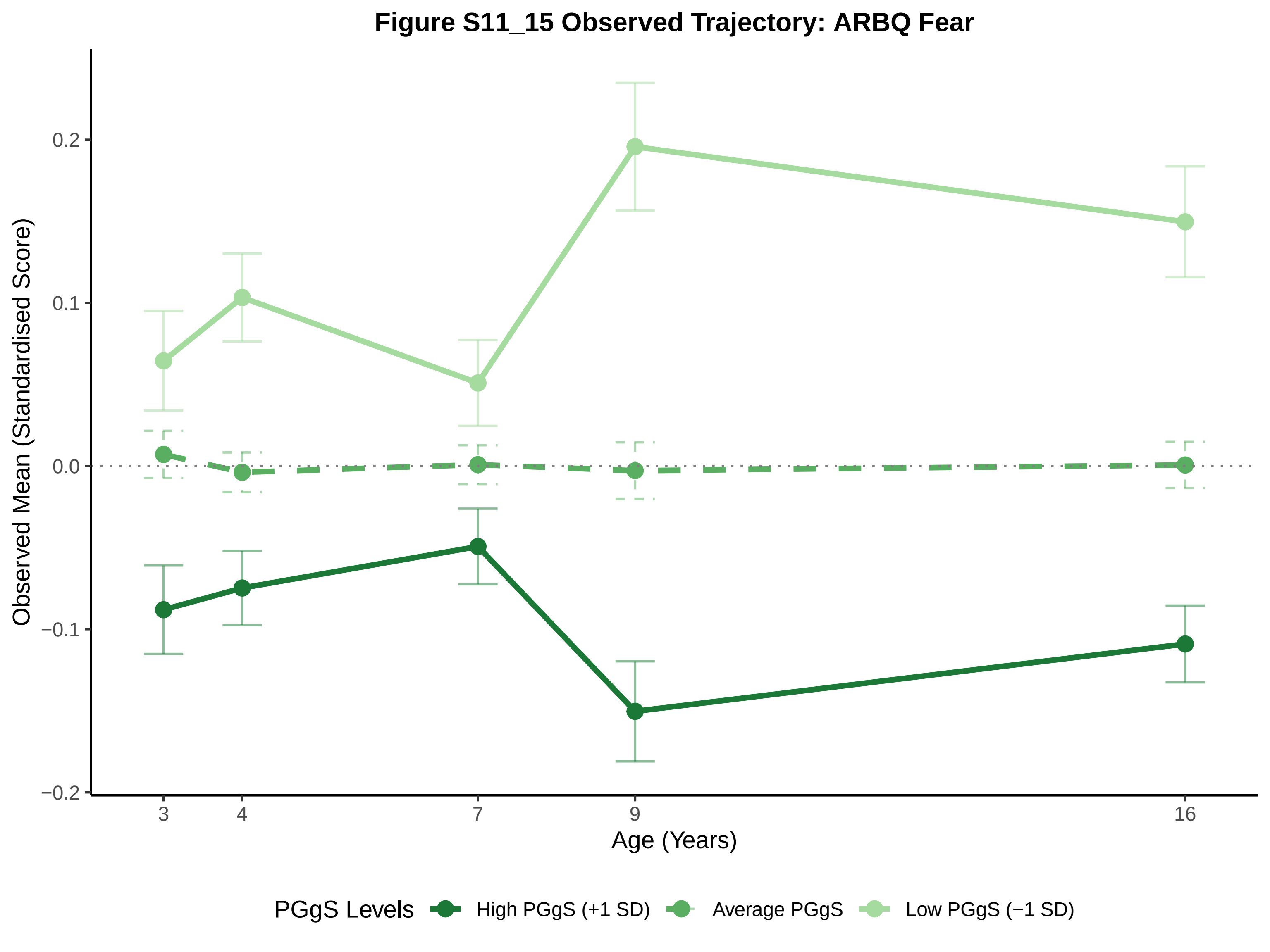
Figure S11_15 Observed Trajectory: ARBQ Fear
Observed Mean (Standardised Score)
0.2
0.1
0.0
−0.1
−0.2
3
4
7
9
16
Age (Years)
PGgS Levels
High PGgS (+1 SD)
Average PGgS
Low PGgS (−1 SD)

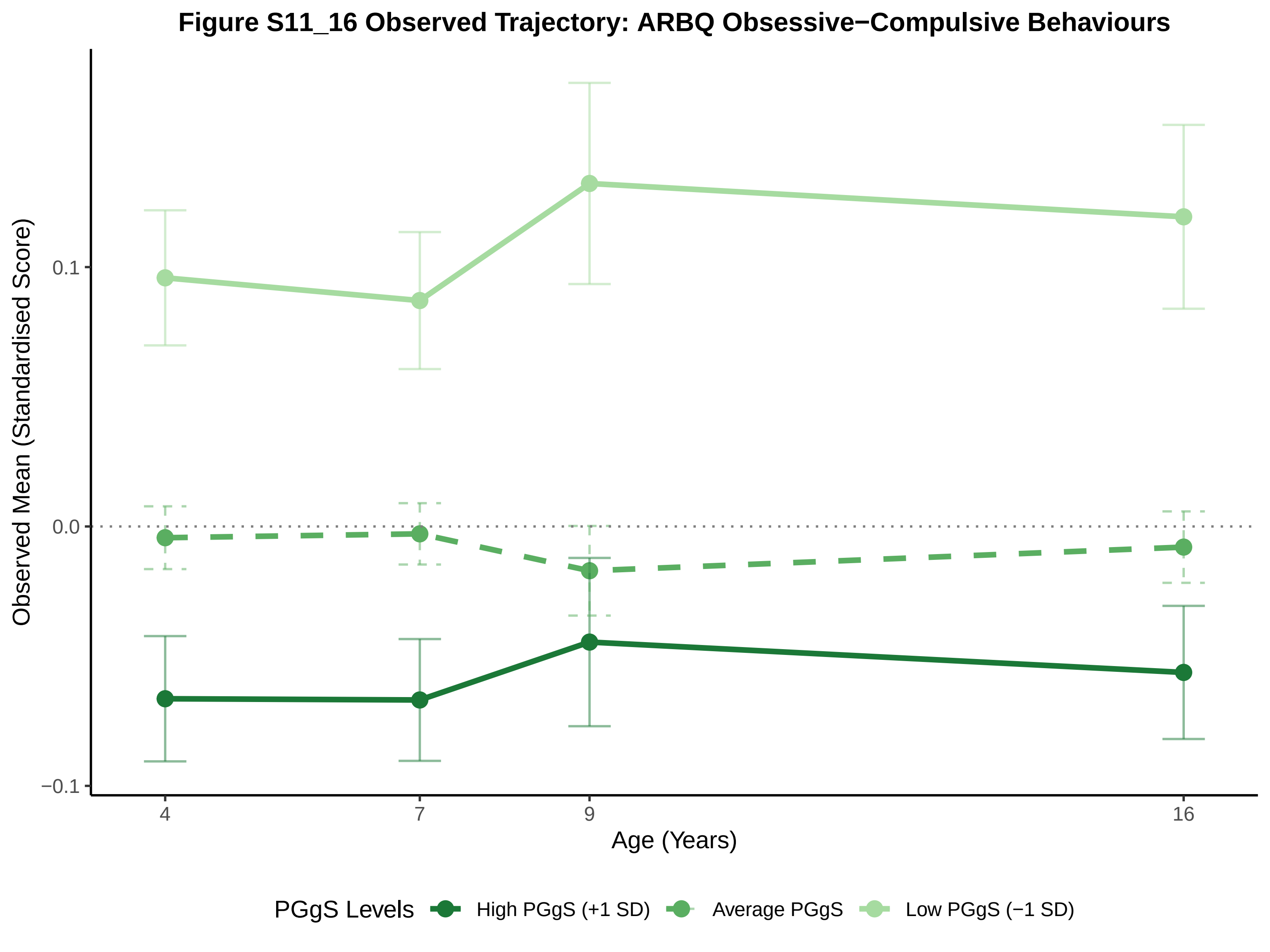
Figure S11_16 Observed Trajectory: ARBQ Obsessive−Compulsive Behaviours
Observed Mean (Standardised Score)
0.1
0.0
−0.1
4
7
9
16
Age (Years)
PGgS Levels
High PGgS (+1 SD)
Average PGgS
Low PGgS (−1 SD)

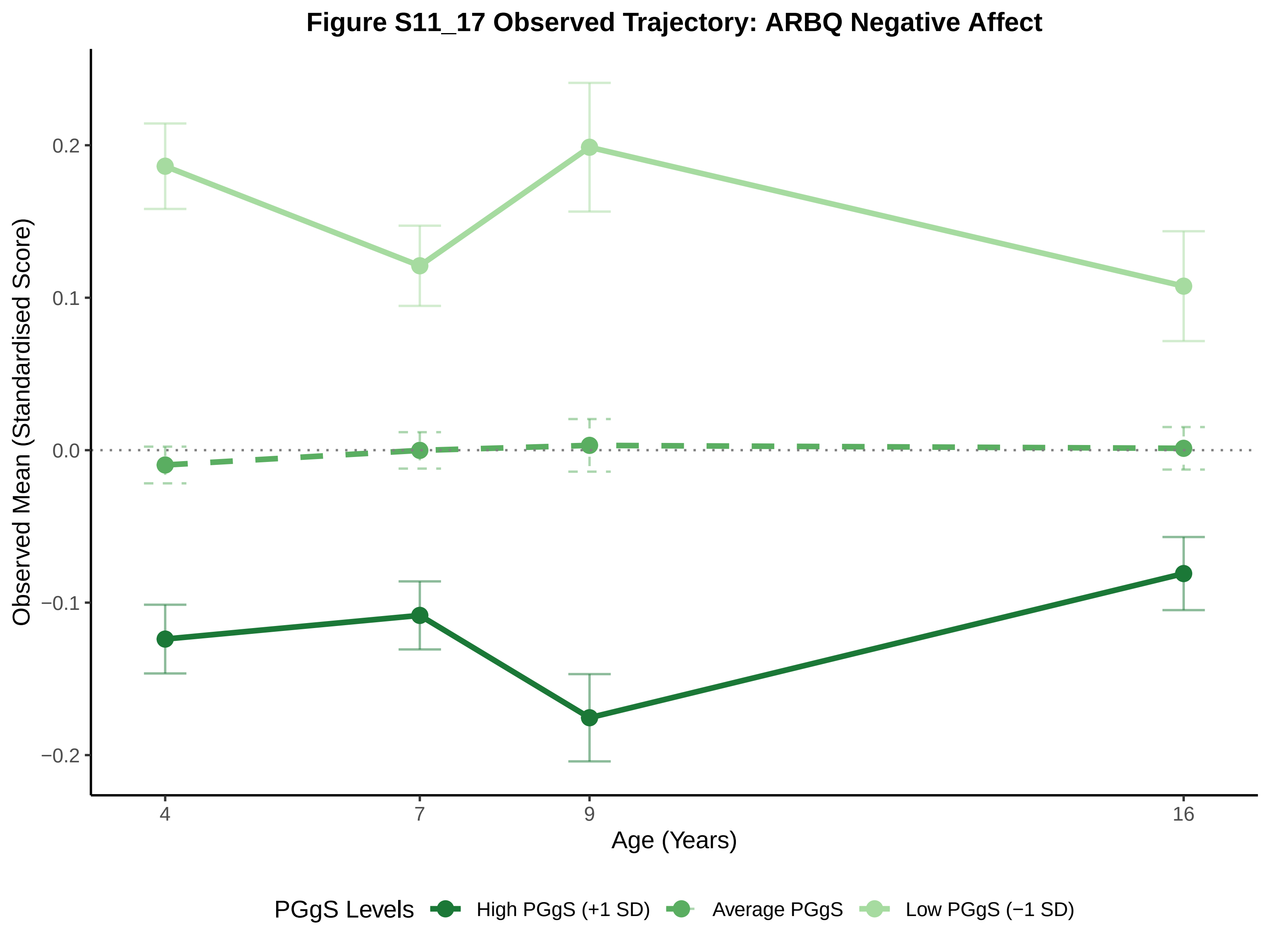

Figure S11_17 Observed Trajectory: ARBQ Negative Affect
Observed Mean (Standardised Score)
0.2
0.1
0.0
−0.1
−0.2
4
7
9
16
Age (Years)
PGgS Levels
High PGgS (+1 SD)
Average PGgS
Low PGgS (−1 SD)

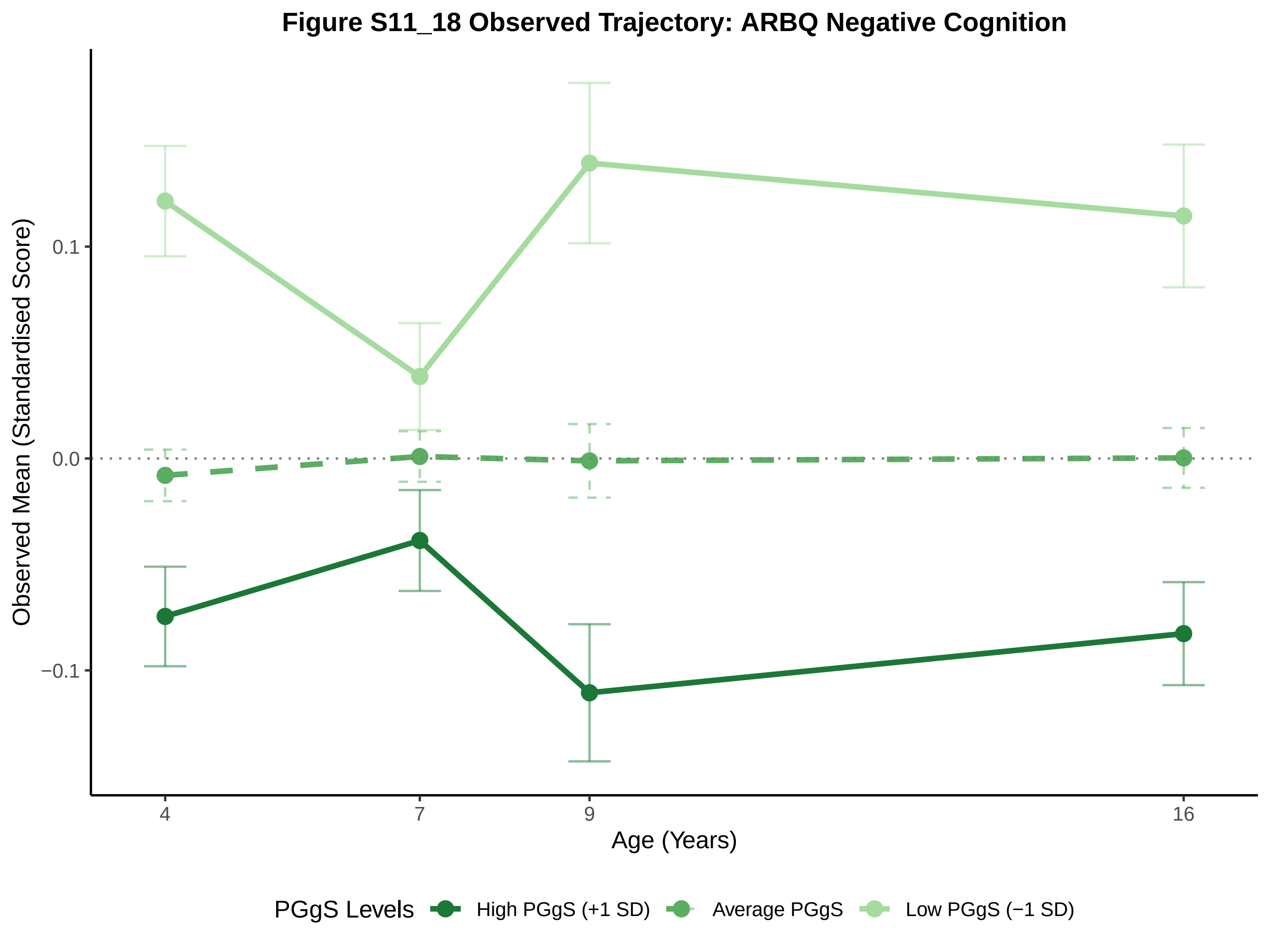

Figure S11_18 Observed Trajectory: ARBQ Negative Cognition
Observed Mean (Standardised Score)
0.1
0.0
−0.1
4
7
9
16
Age (Years)
PGgS Levels
High PGgS (+1 SD)
Average PGgS
Low PGgS (−1 SD)

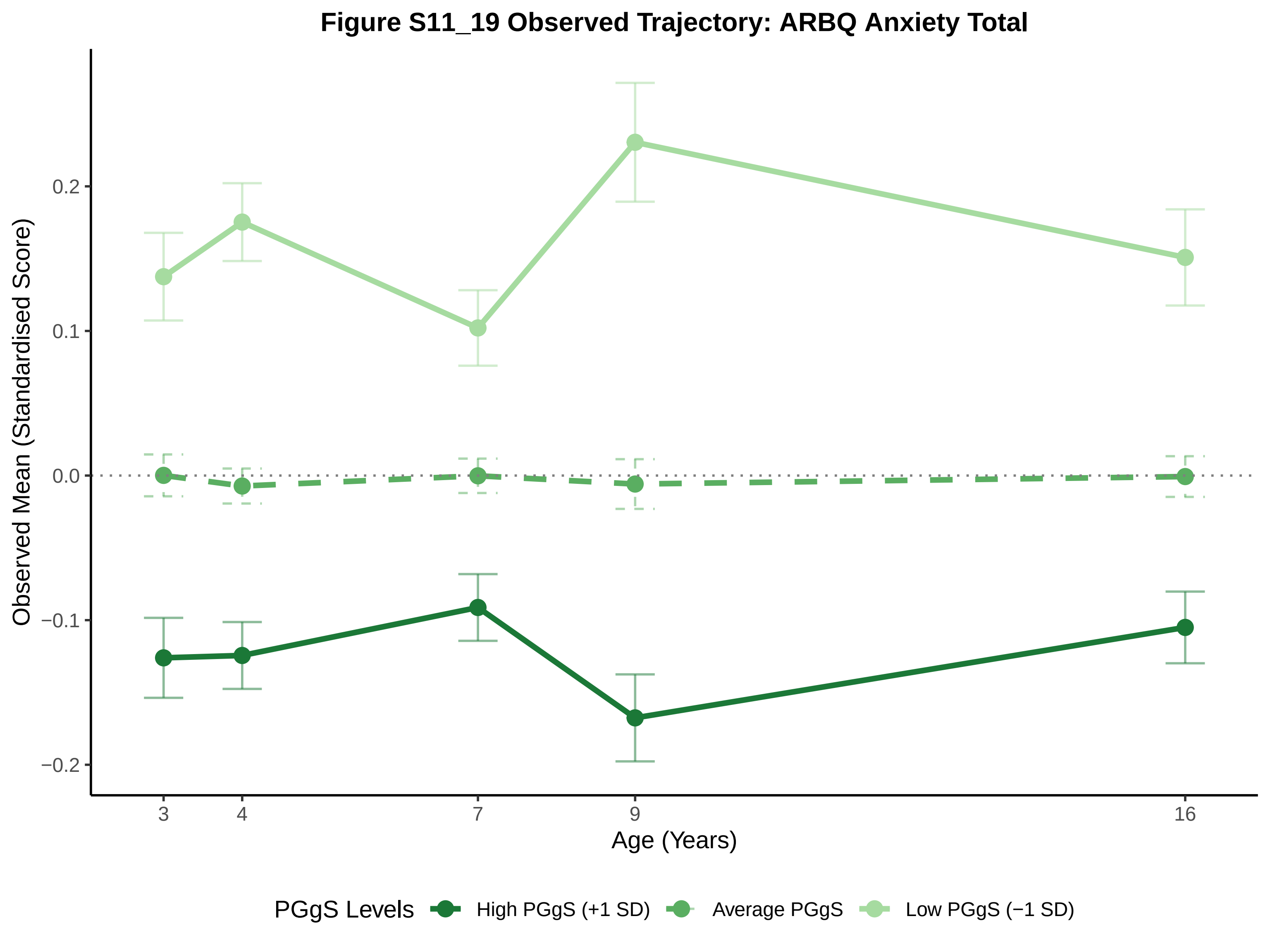
Figure S11_19 Observed Trajectory: ARBQ Anxiety Total
Observed Mean (Standardised Score)
0.2
0.1
0.0
−0.1
−0.2
3
4
7
9
16
Age (Years)
PGgS Levels
High PGgS (+1 SD)
Average PGgS
Low PGgS (−1 SD)

#### Figure S11_20 Observed Trajectory: Conners ADHD Inattention

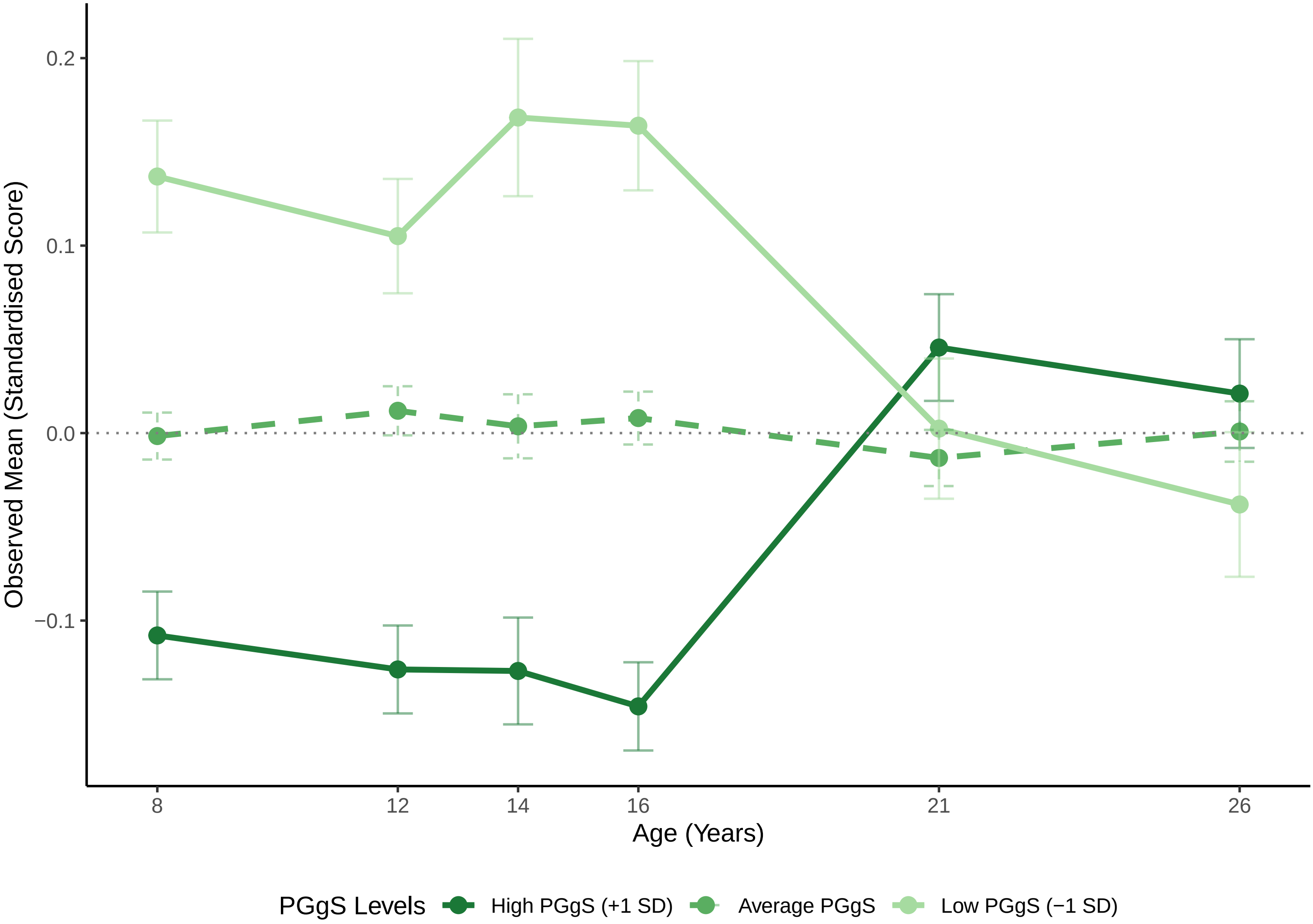

**Figure S11_21 Observed Trajectory: Conners ADHD Hyperactivity_Impulsivity**

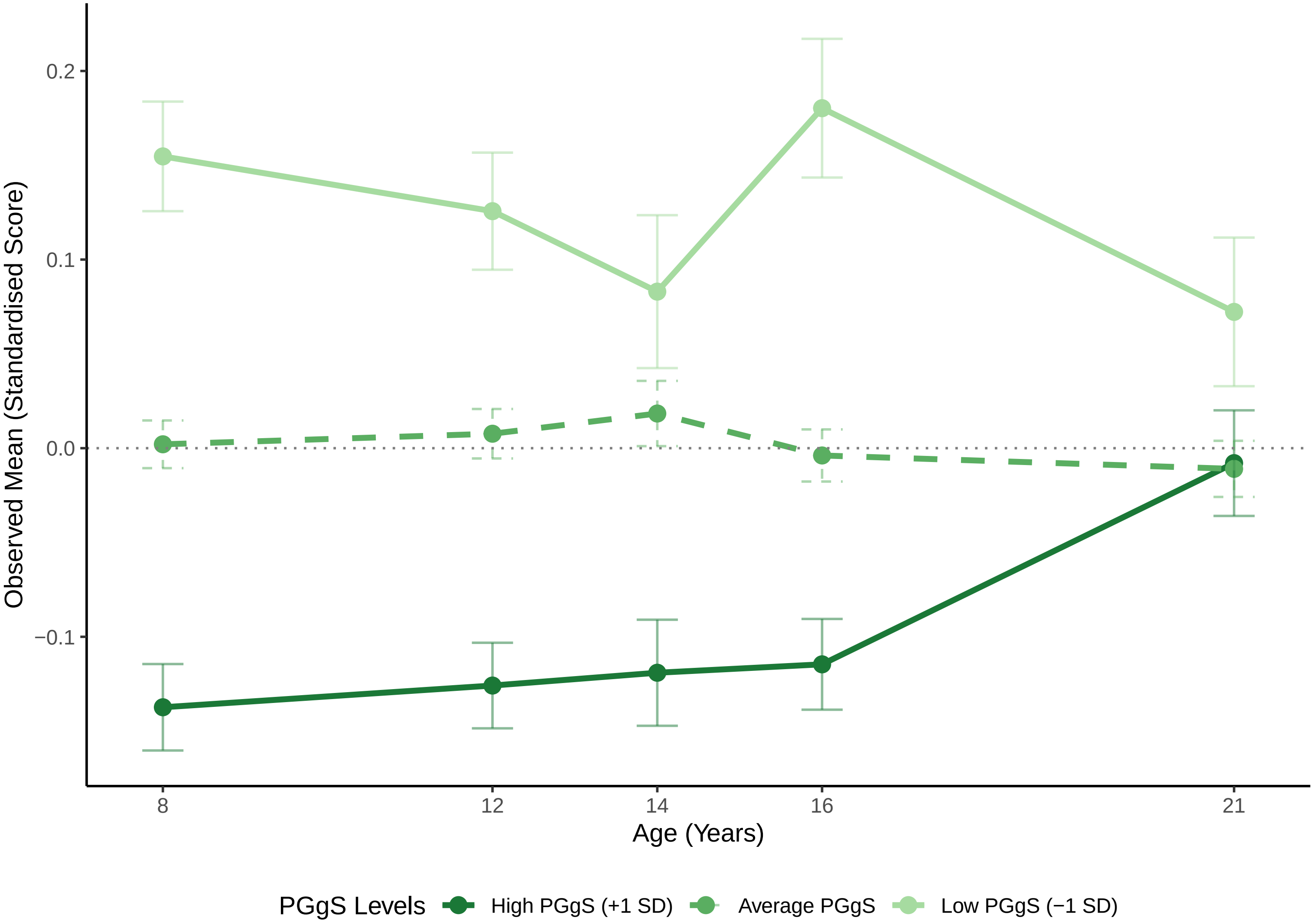

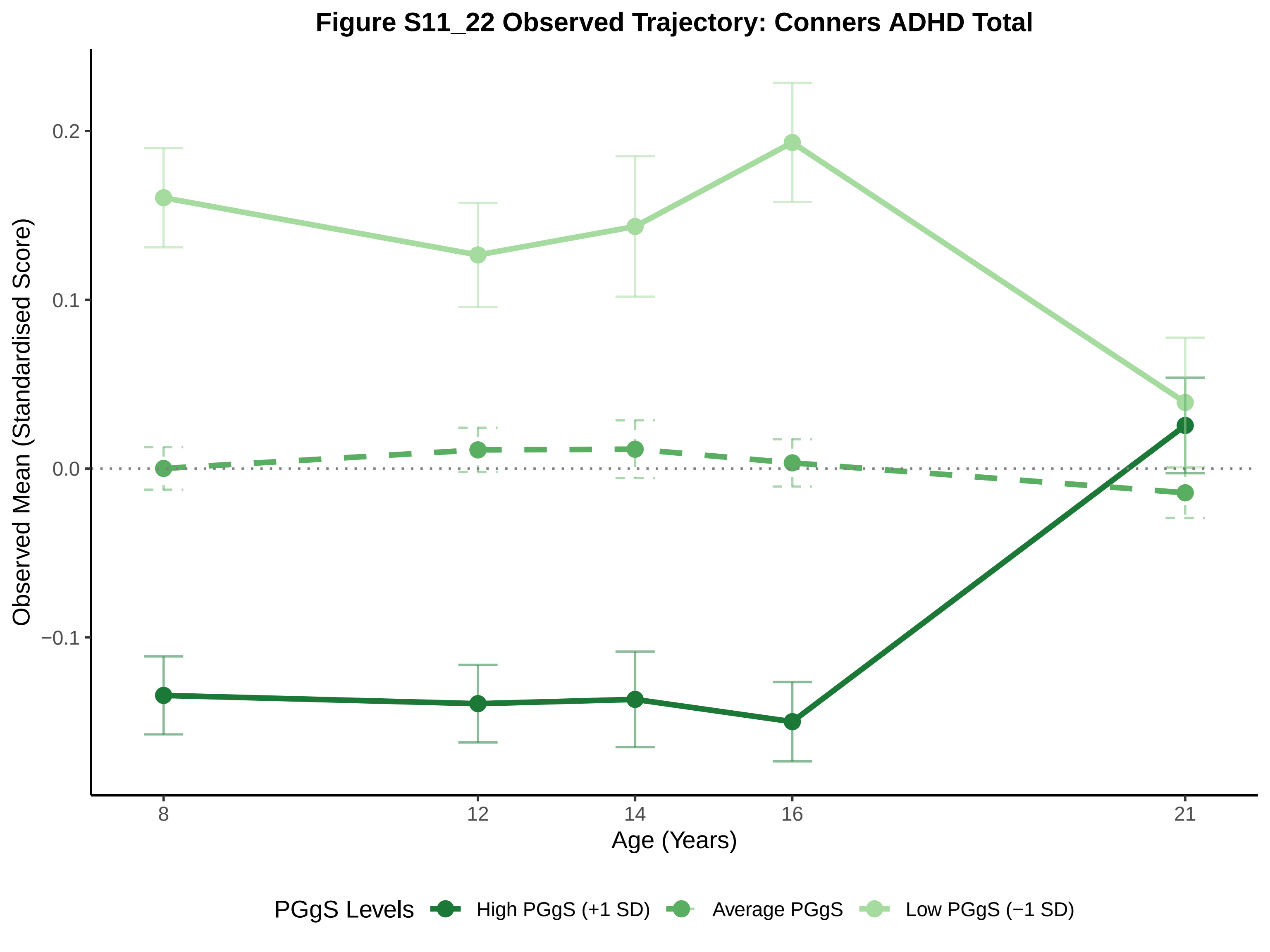
Figure S11_22 Observed Trajectory: Conners ADHD Total
Observed Mean (Standardised Score)
0.2
0.1
0.0
−0.1
8
12
14
16
21
Age (Years)
PGgS Levels
High PGgS (+1 SD)
Average PGgS
Low PGgS (−1 SD)

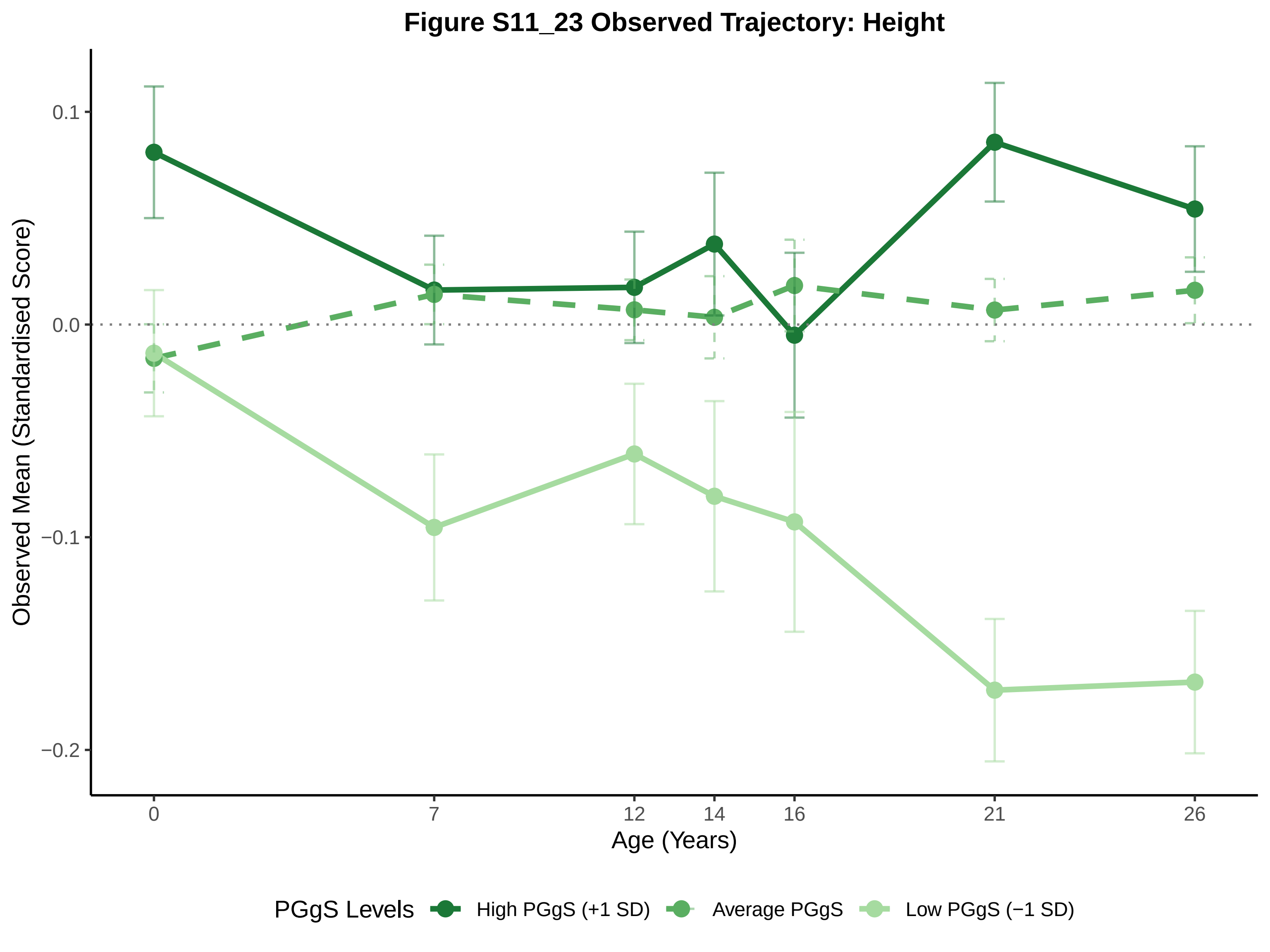

Figure S11_23 Observed Trajectory: Height
Observed Mean (Standardised Score)
0.1
0.0
−0.1
−0.2
0
7
12
14
16
21
26
Age (Years)
PGgS Levels
High PGgS (+1 SD)
Average PGgS
Low PGgS (−1 SD)

### Figure S11_24 Observed Trajectory: BMI (as weight at birth)

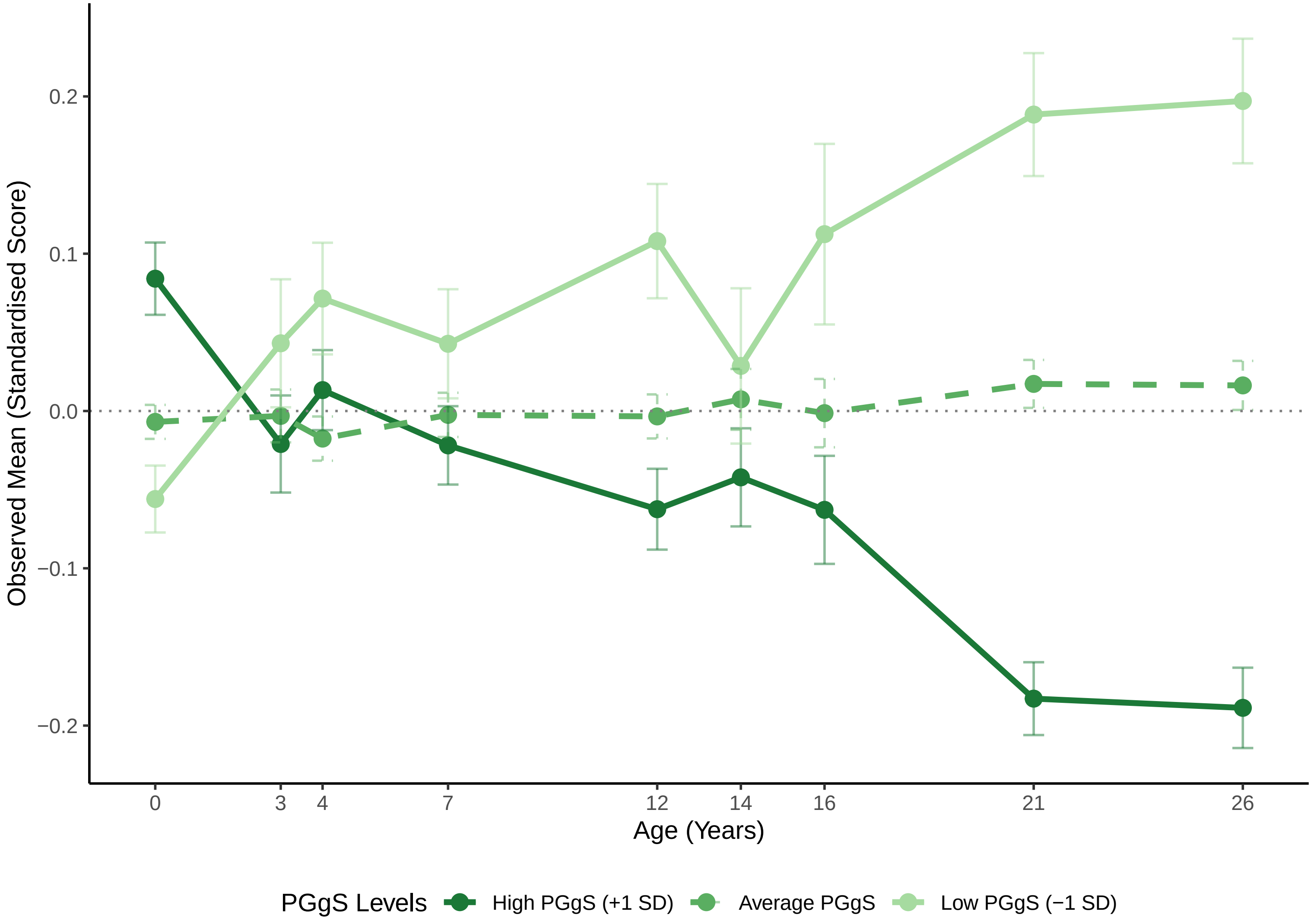

**Figure S13_1 Observed Trajectory: General Cognitive Ability**

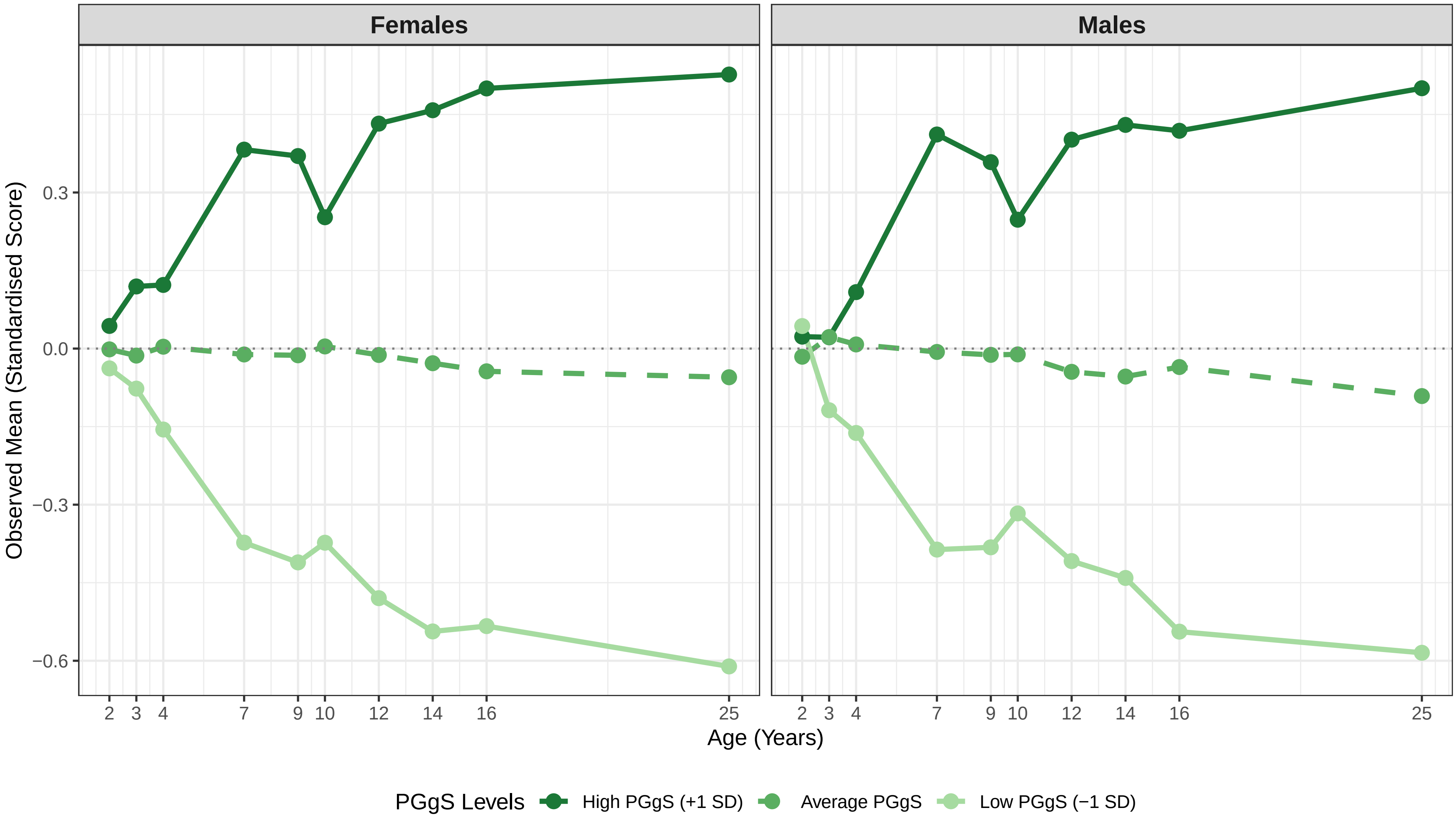

**Figure S13_2 Observed Trajectory: Verbal Abilities**

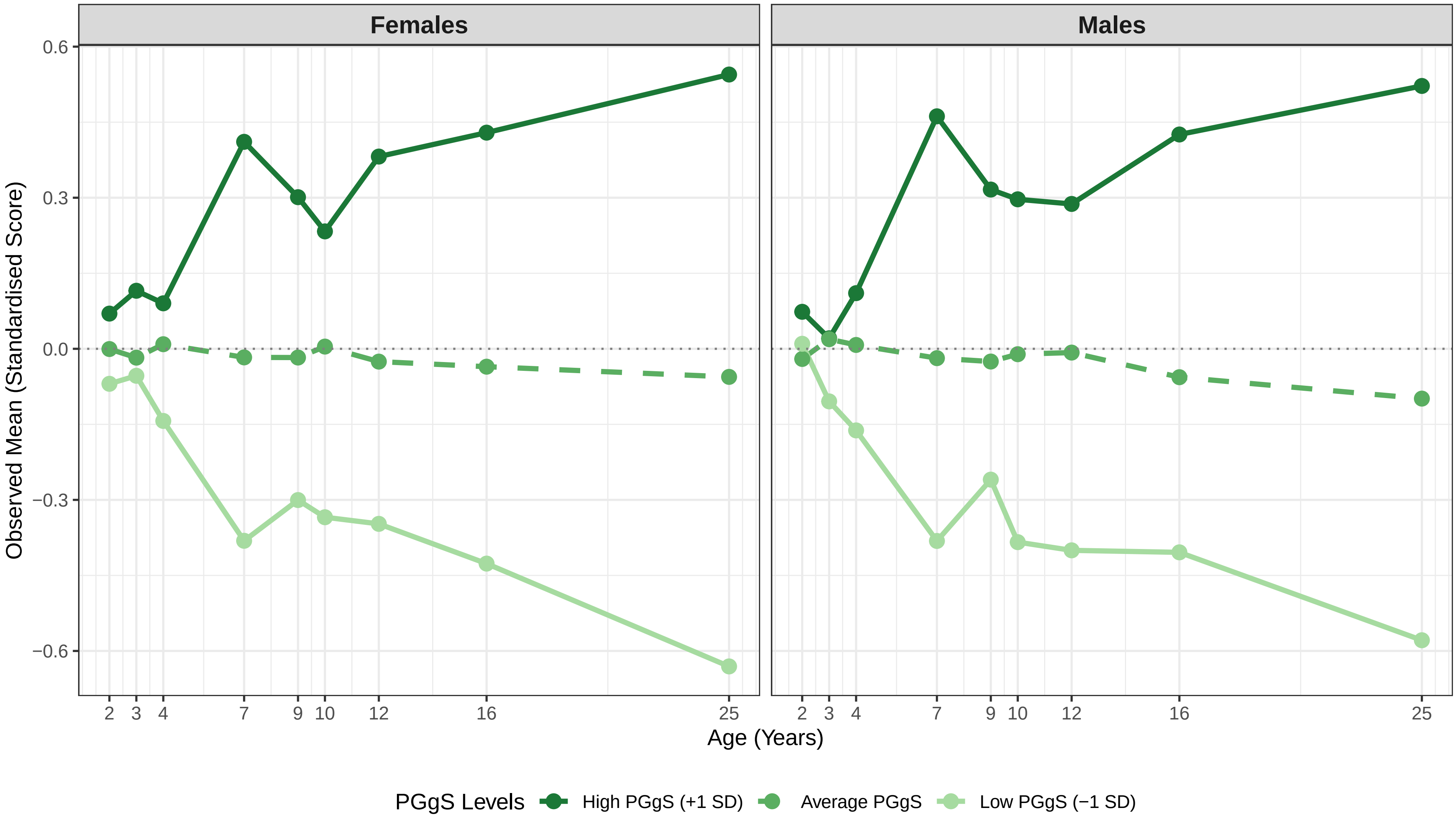

Figure S13_3 Observed Trajectory: Nonverbal Abilities

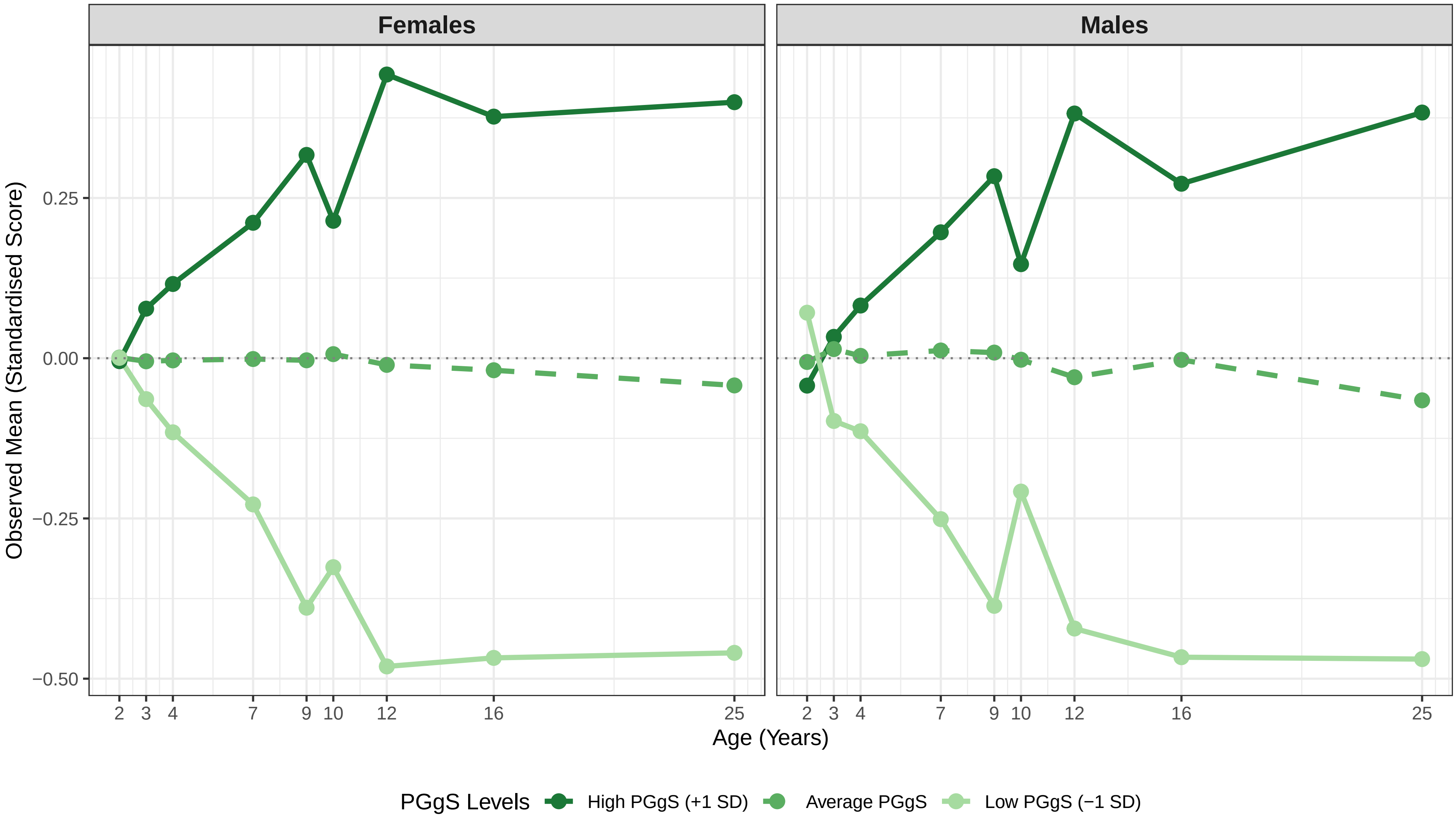

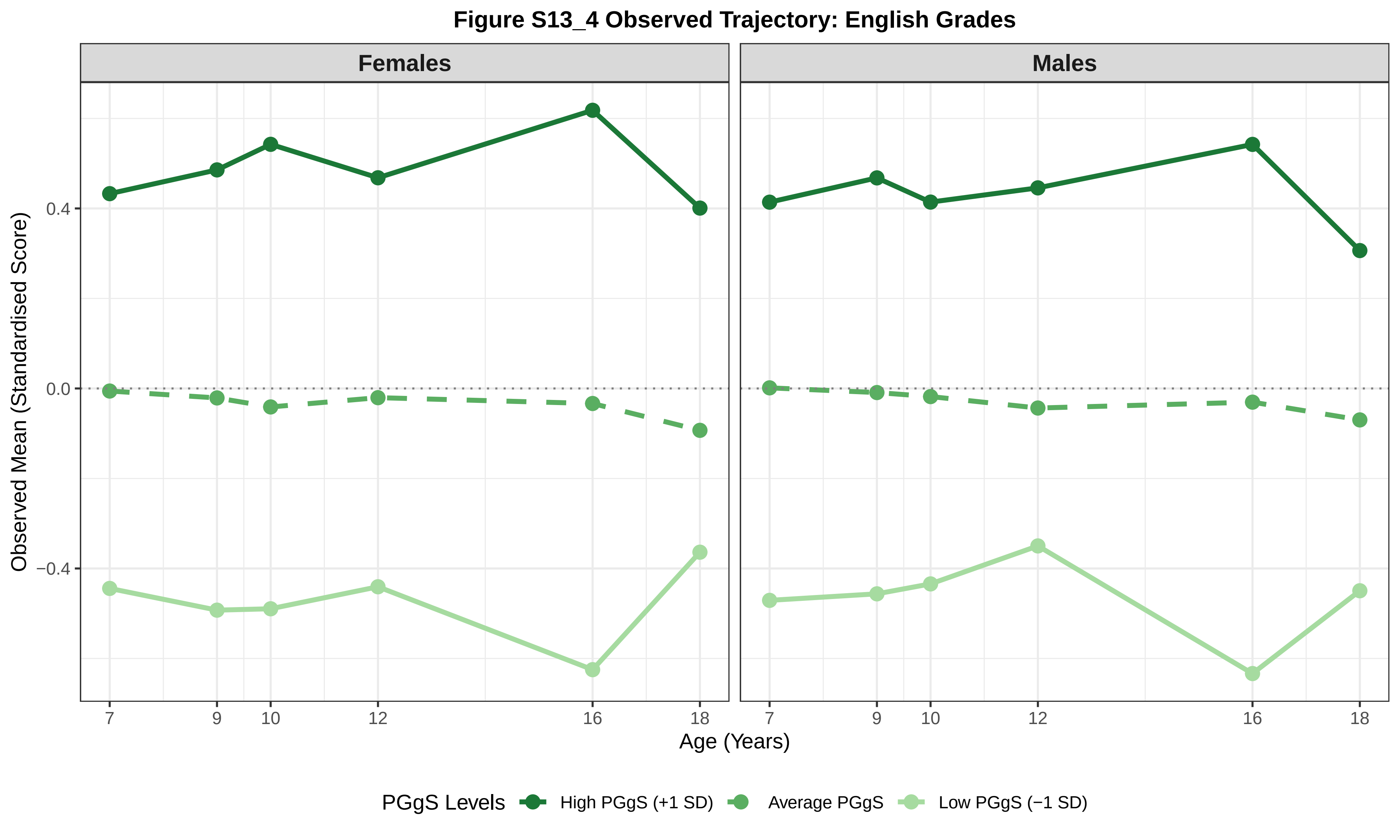

Figure S13_4 Observed Trajectory: English Grades
Females
Males
Observed Mean (Standardised Score)
0.4
0.0
−0.4
7
9
10
12
16
18
Age (Years)
PGgS Levels
High PGgS (+1 SD)
Average PGgS
Low PGgS (−1 SD)

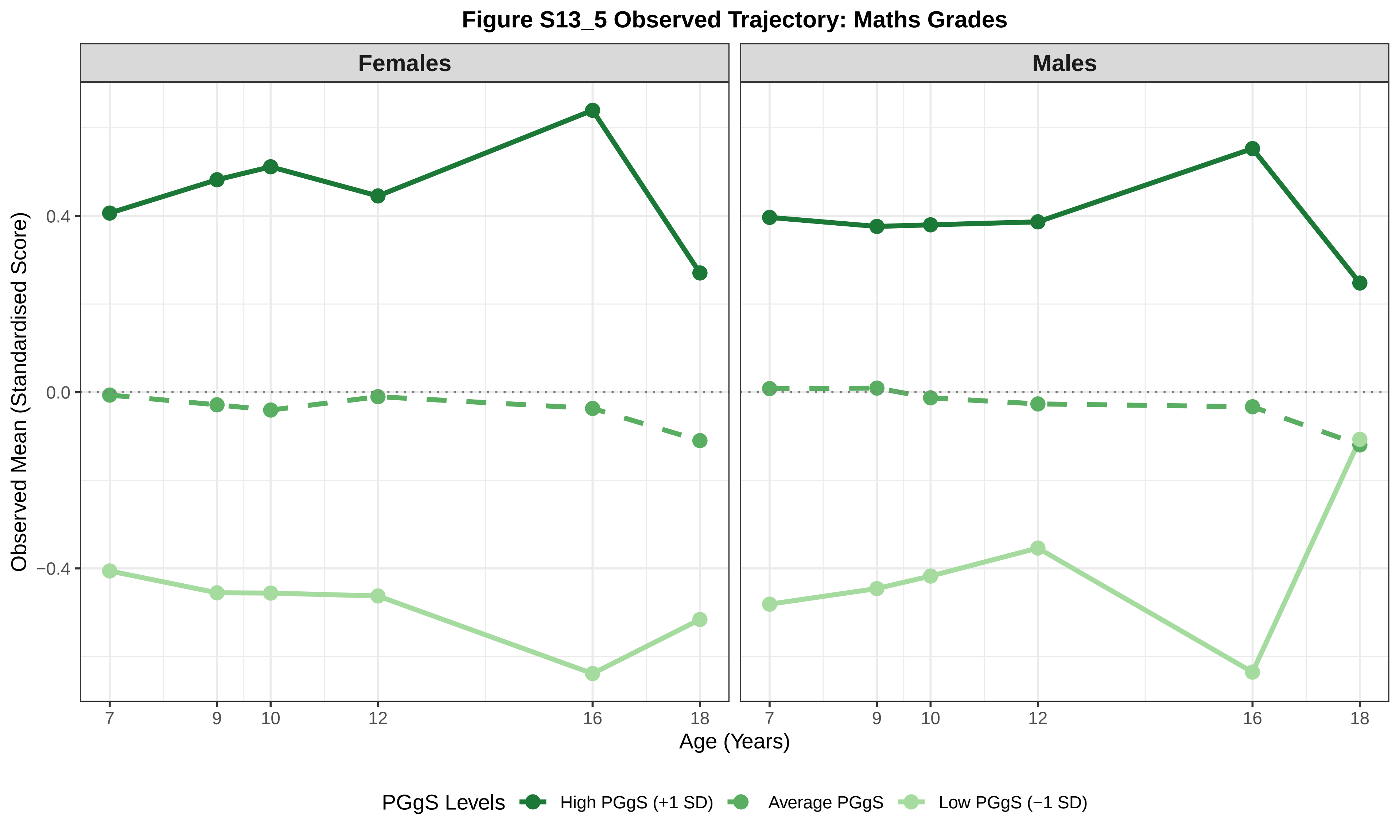

Figure S13_5 Observed Trajectory: Maths Grades
Females
Males
Observed Mean (Standardised Score)
0.4
0.0
−0.4
7
9
10
12
16
18
Age (Years)
PGgS Levels
High PGgS (+1 SD)
Average PGgS
Low PGgS (−1 SD)

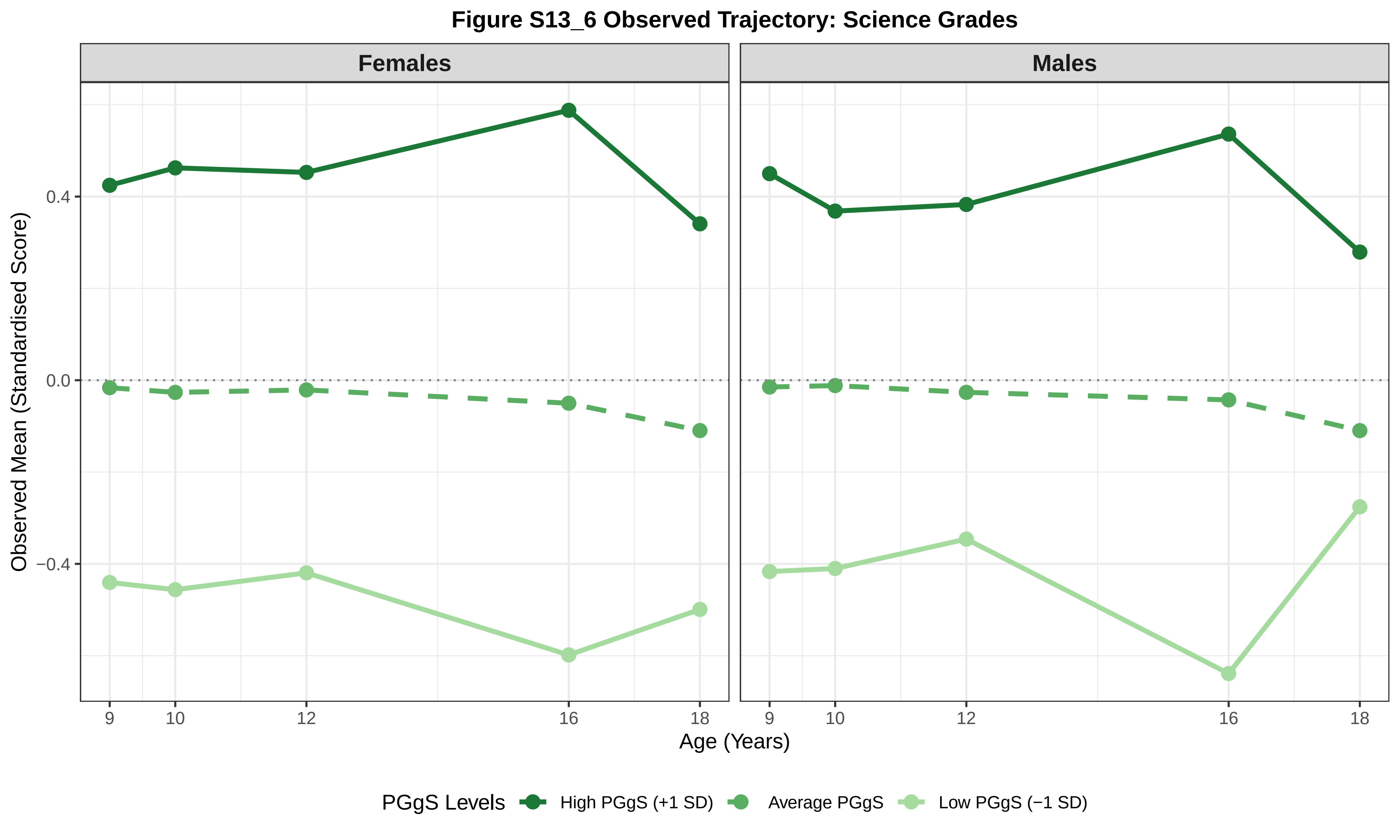
Figure S13_6 Observed Trajectory: Science Grades
Females
Males
Observed Mean (Standardised Score)
0.4
0.0
−0.4
9
10
12
16
18
Age (Years)
PGgS Levels
High PGgS (+1 SD)
Average PGgS
Low PGgS (−1 SD)

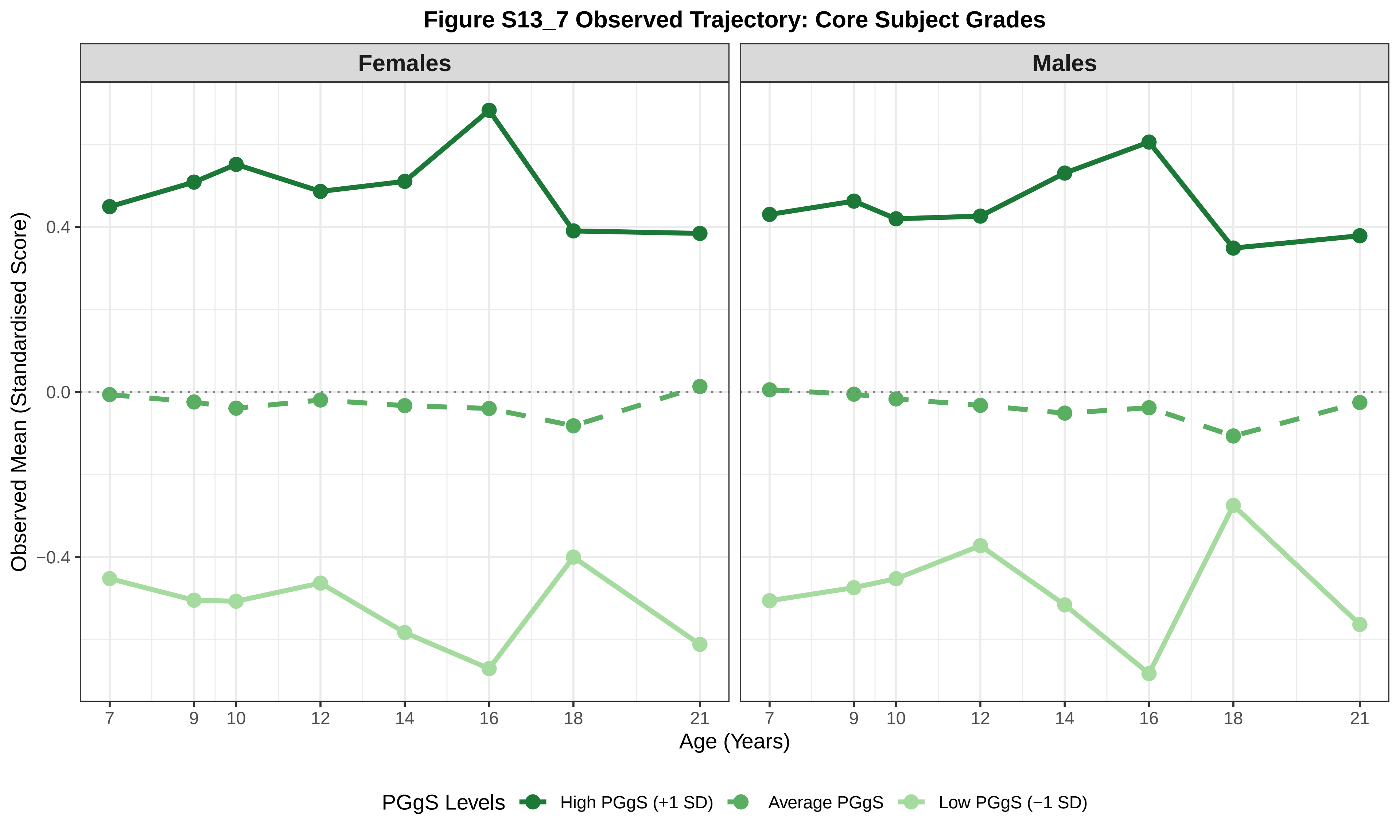
Figure S13_7 Observed Trajectory: Core Subject Grades
Females
Males
Observed Mean (Standardised Score)
0.4
0.0
−0.4
7
9
10
12
14
16
18
21
Age (Years)
PGgS Levels
High PGgS (+1 SD)
Average PGgS
Low PGgS (−1 SD)

**Figure S13_8 Observed Trajectory: SDQ Conduct**

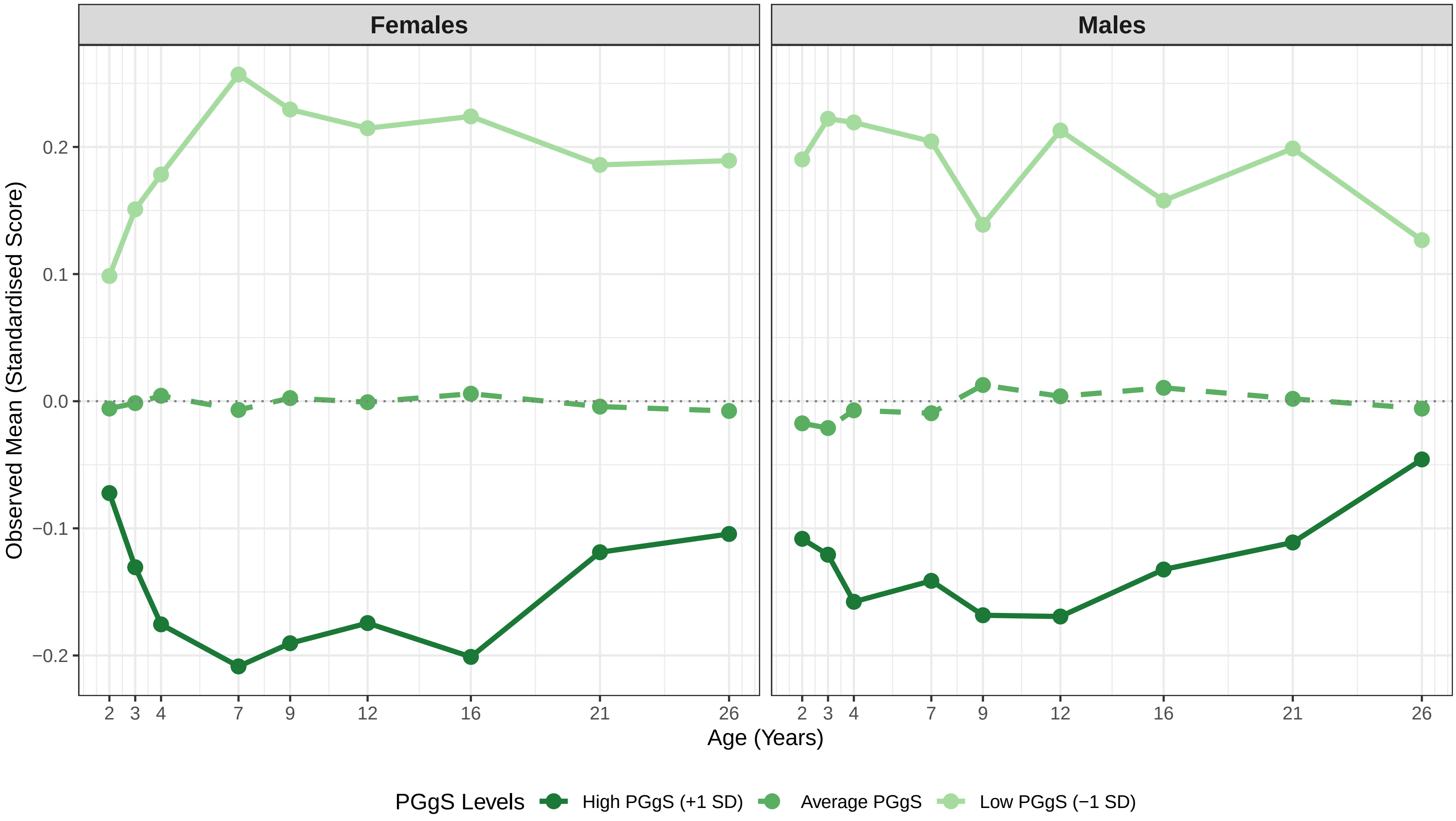

Figure S13_9 Observed Trajectory: SDQ Emotion

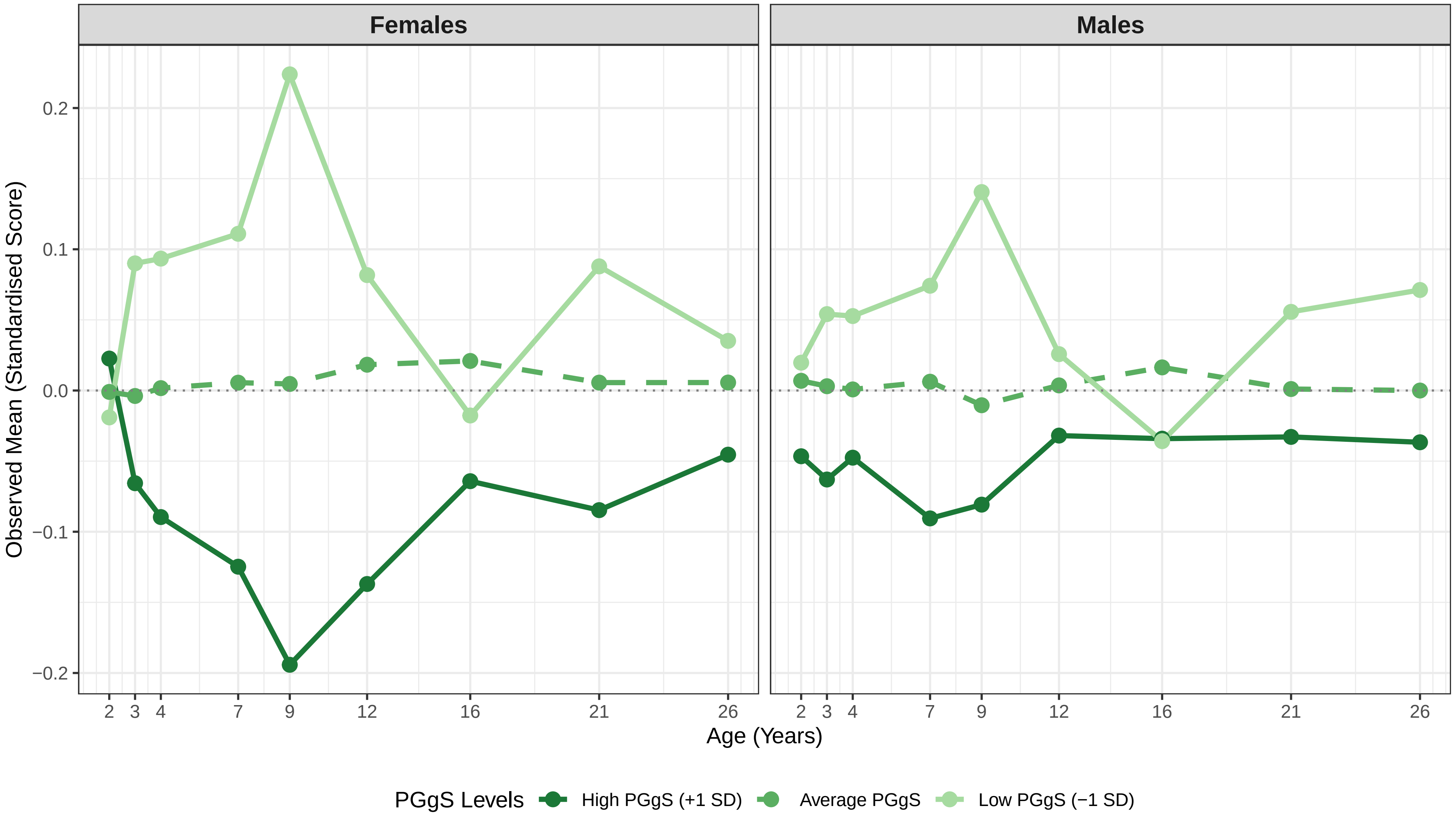

Figure S13_10 Observed Trajectory: SDQ Hyperactivity_SDQ

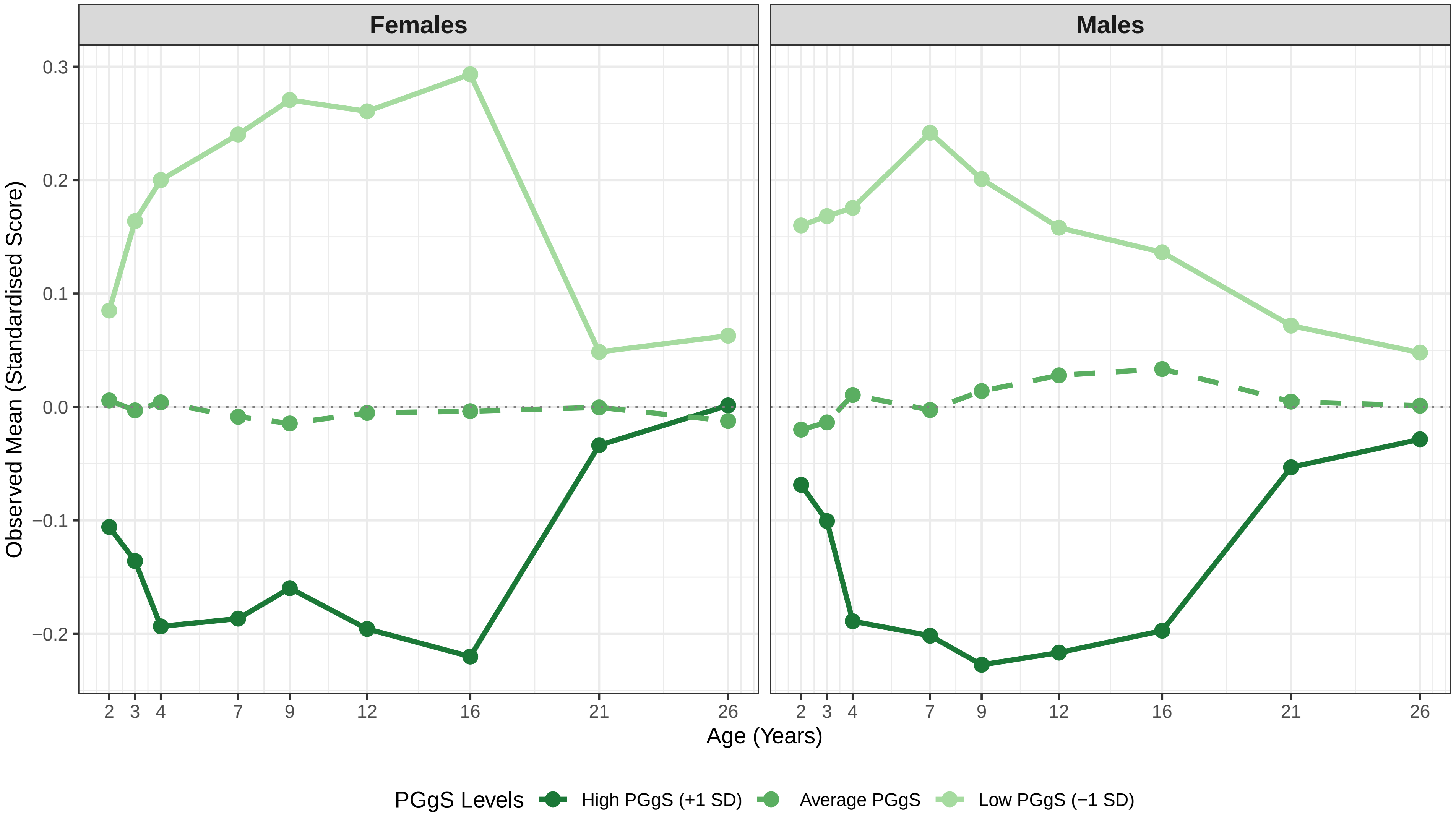

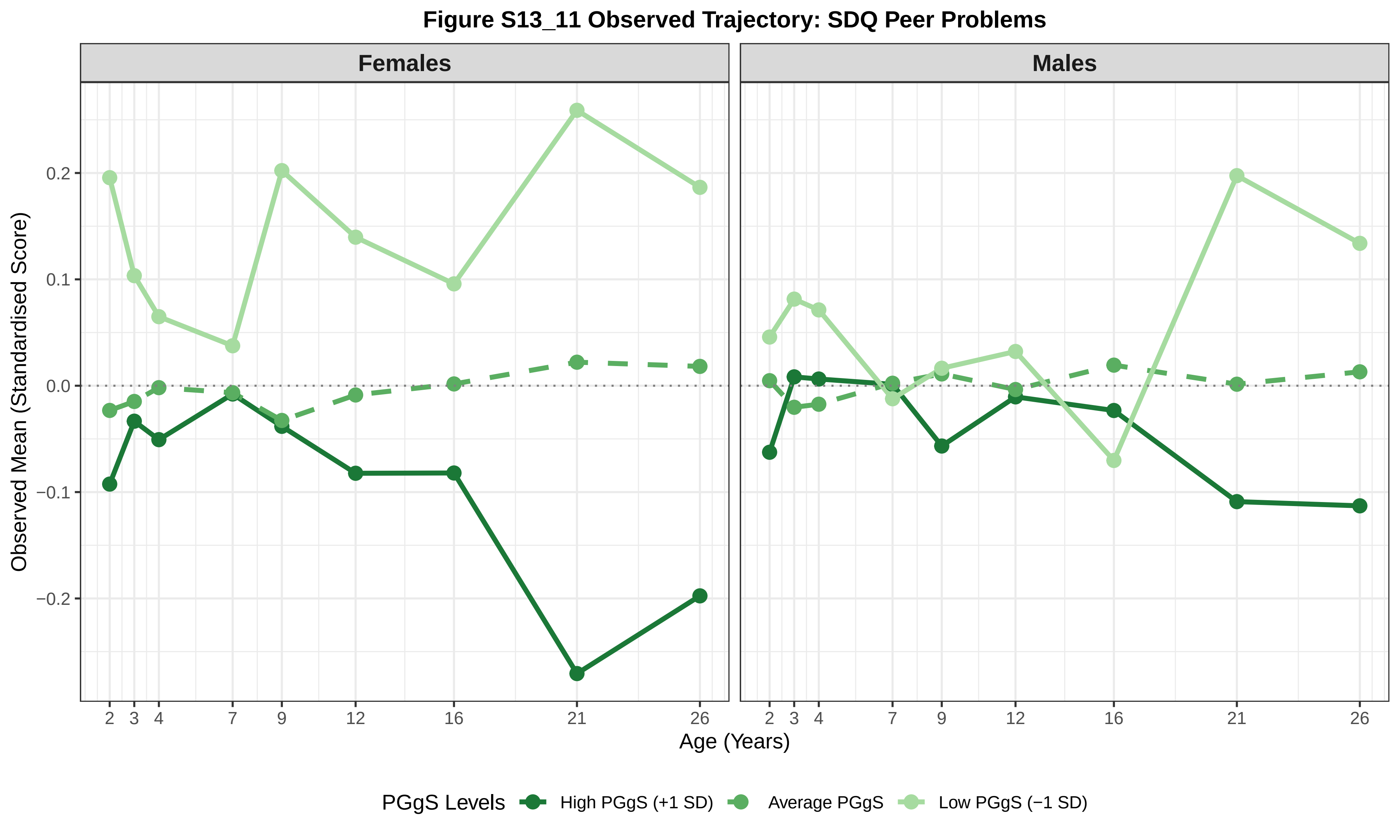
Figure S13_11 Observed Trajectory: SDQ Peer Problems
Females
Males
Observed Mean (Standardised Score)
0.2
0.1
0.0
−0.1
−0.2
2
3
4
7
9
12
16
21
26
Age (Years)
PGgS Levels
High PGgS (+1 SD)
Average PGgS
Low PGgS (−1 SD)

**Figure S13_12 Observed Trajectory: SDQ Prosocial**

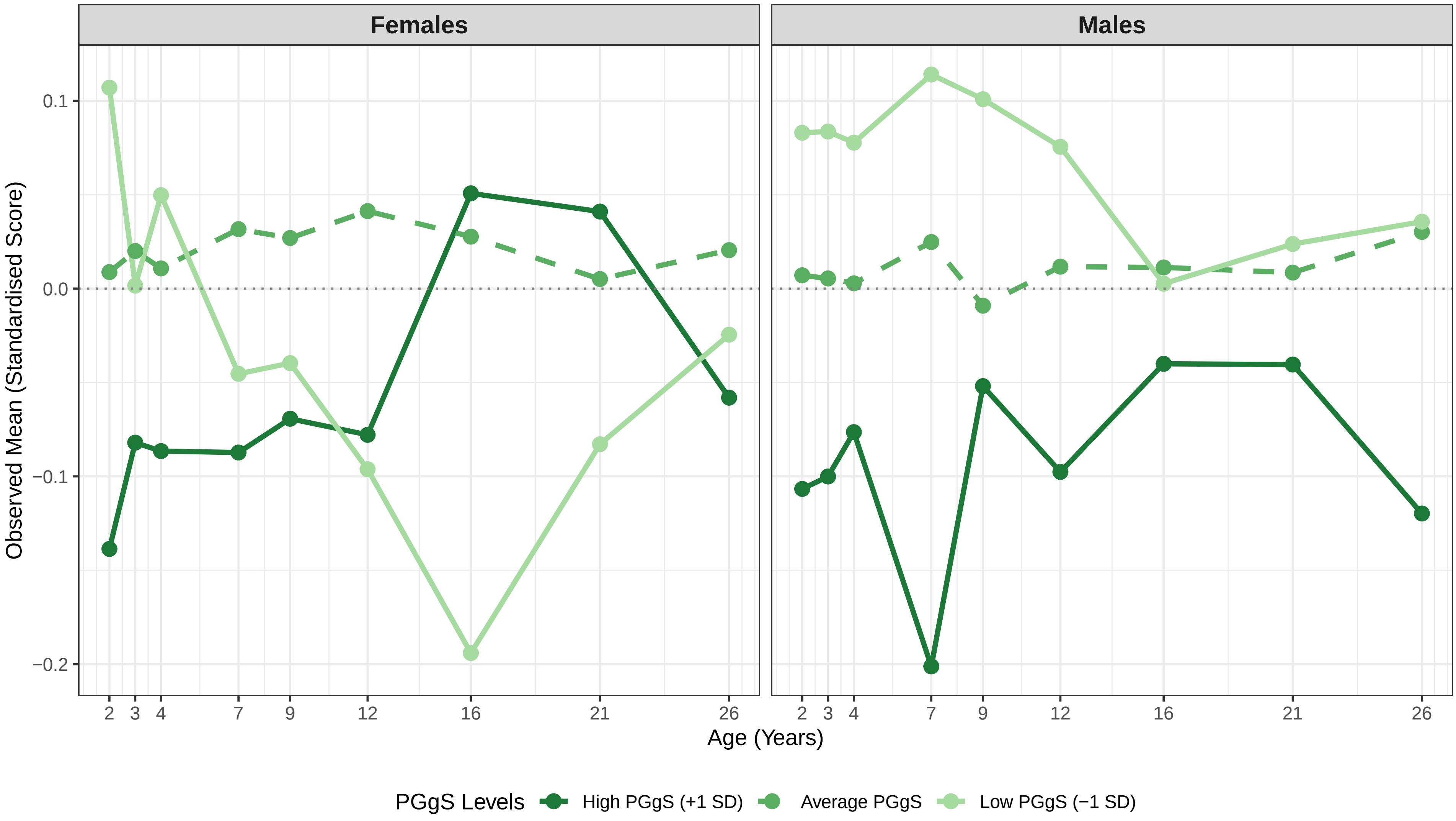

Figure S13_13 Observed Trajectory: SDQ Total Problems

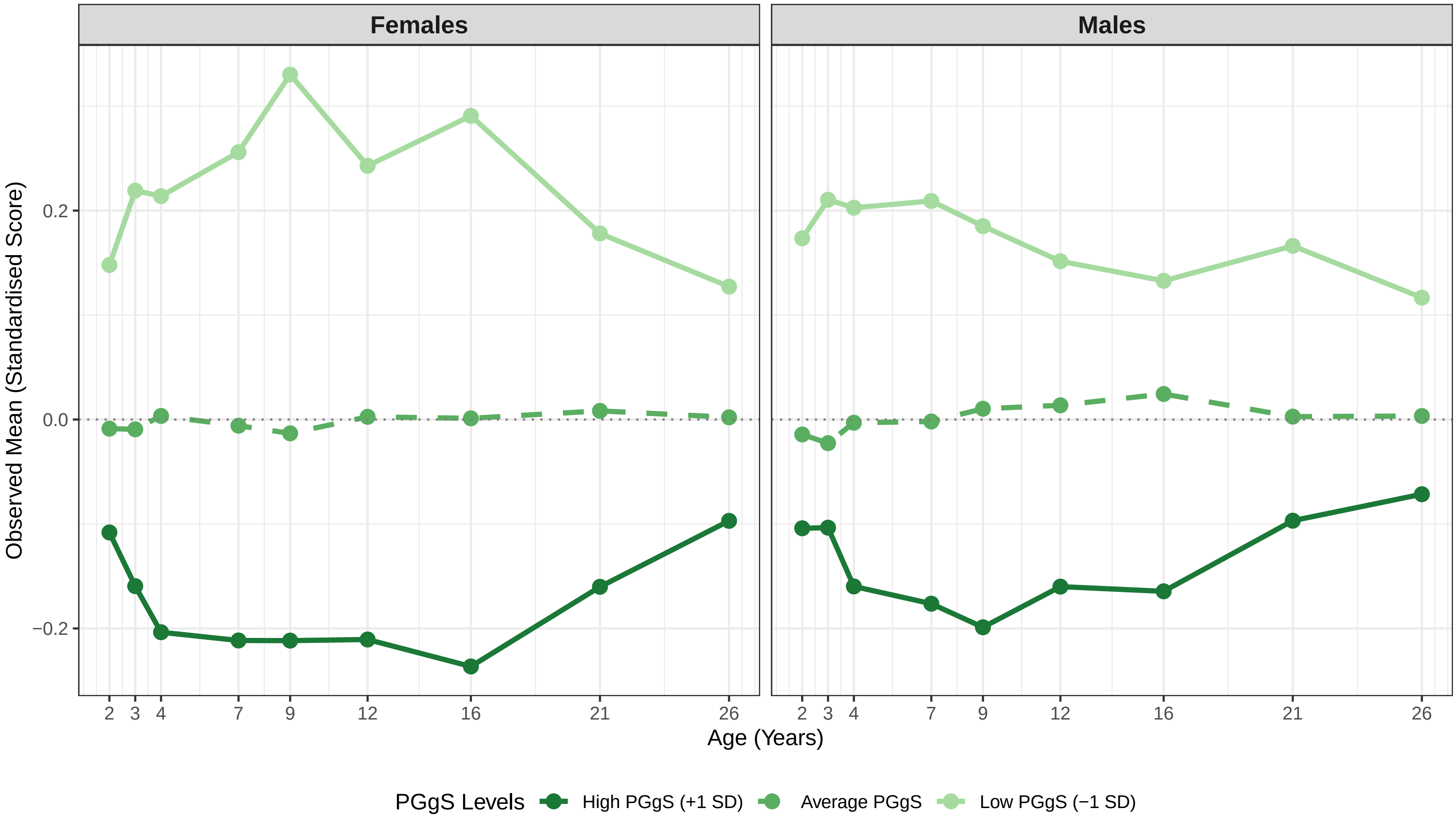

**Figure S13_14 Observed Trajectory: ARBQ Shyness**

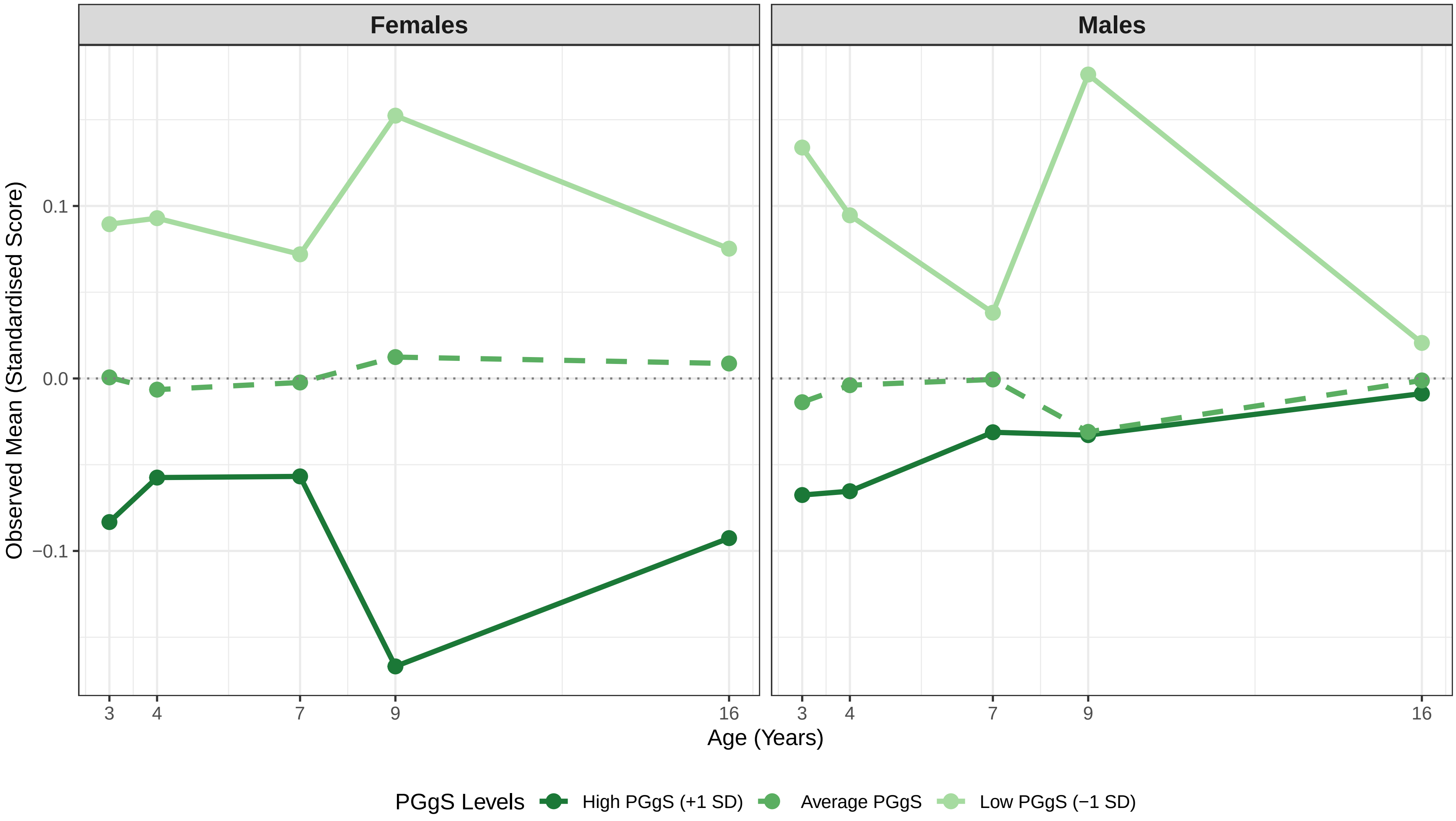

Figure S13_15 Observed Trajectory: ARBQ Fear

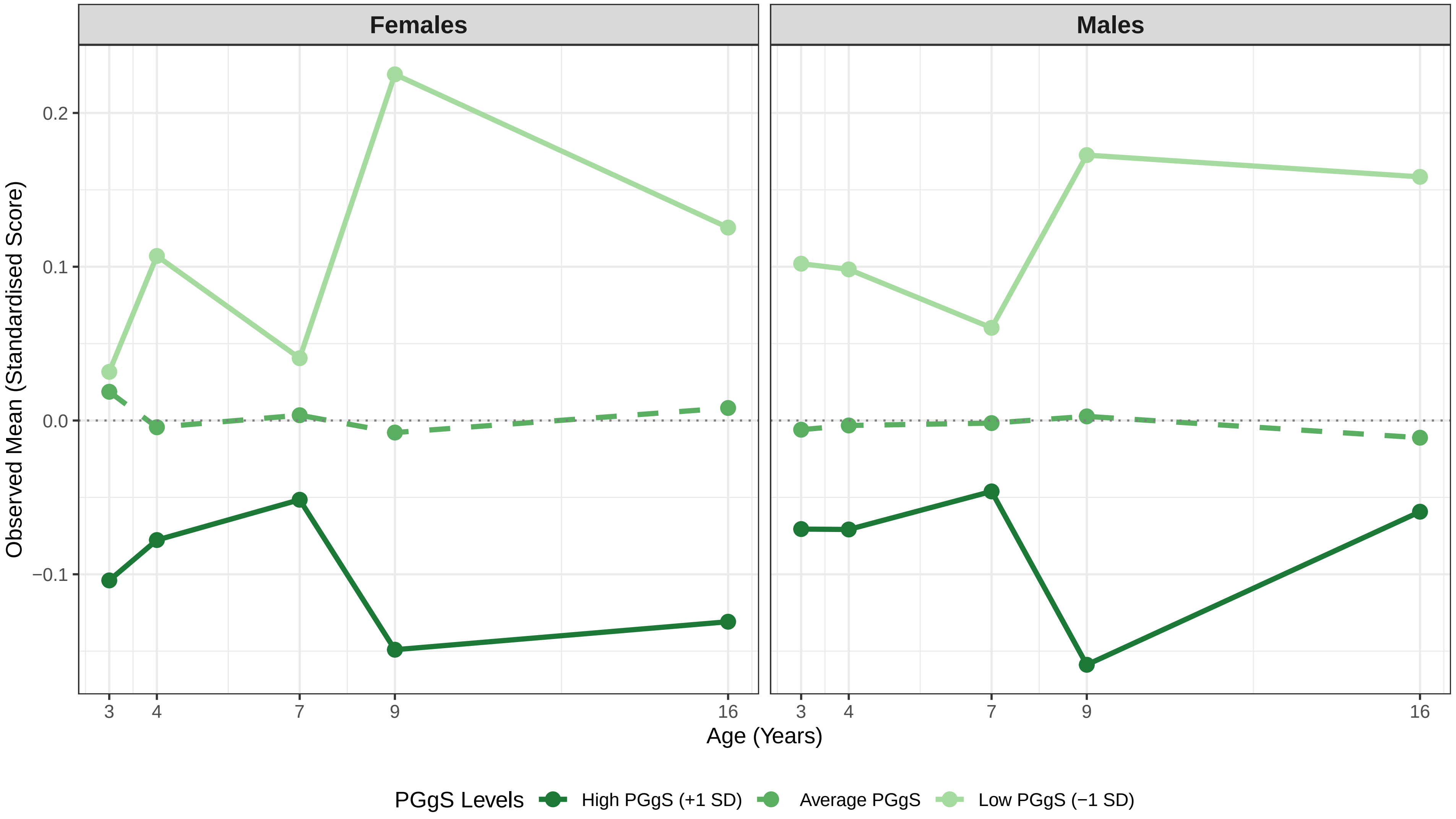

Figure S13_16 Observed Trajectory: ARBQ Obsessive−Compulsive Behaviours

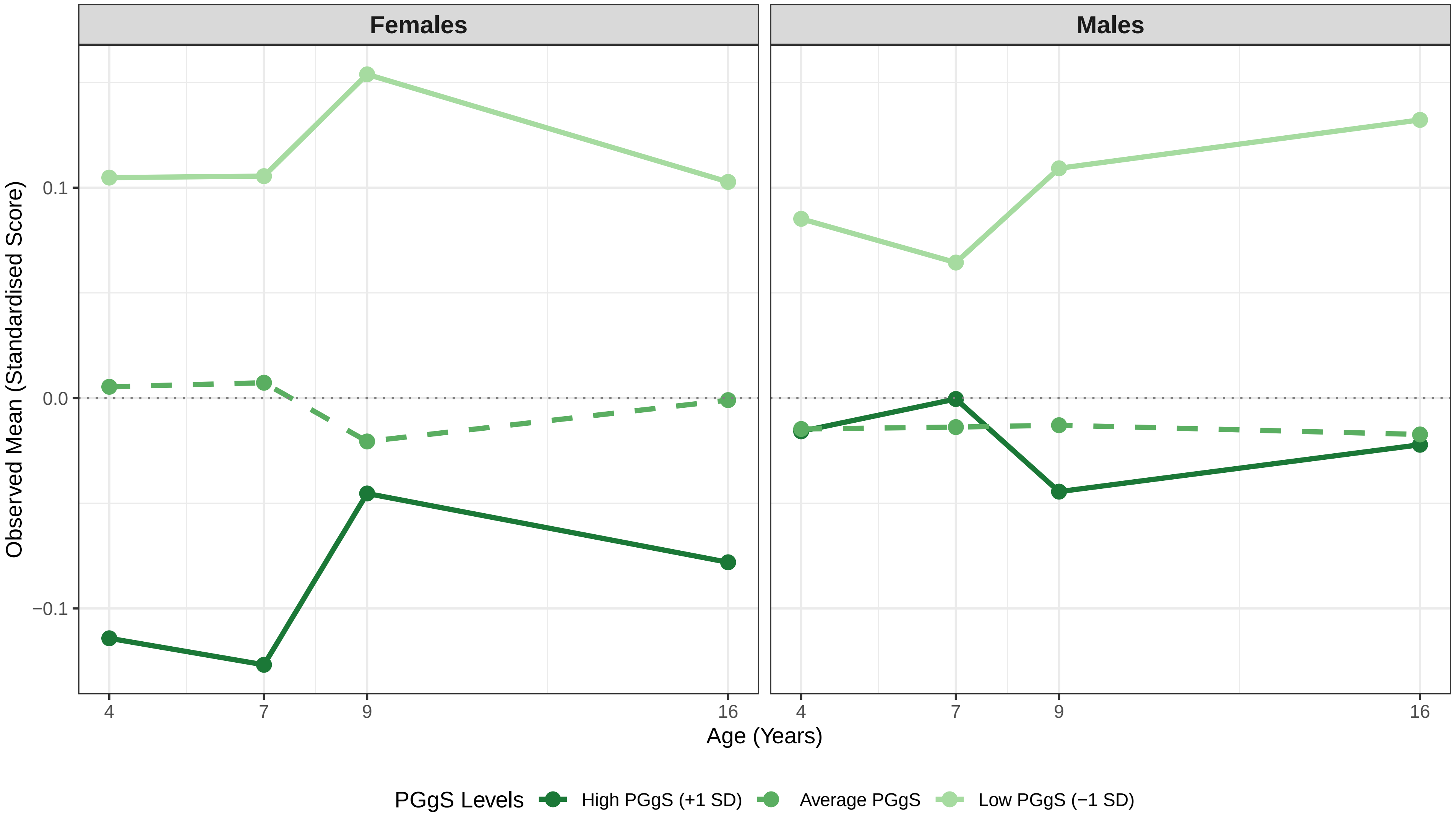

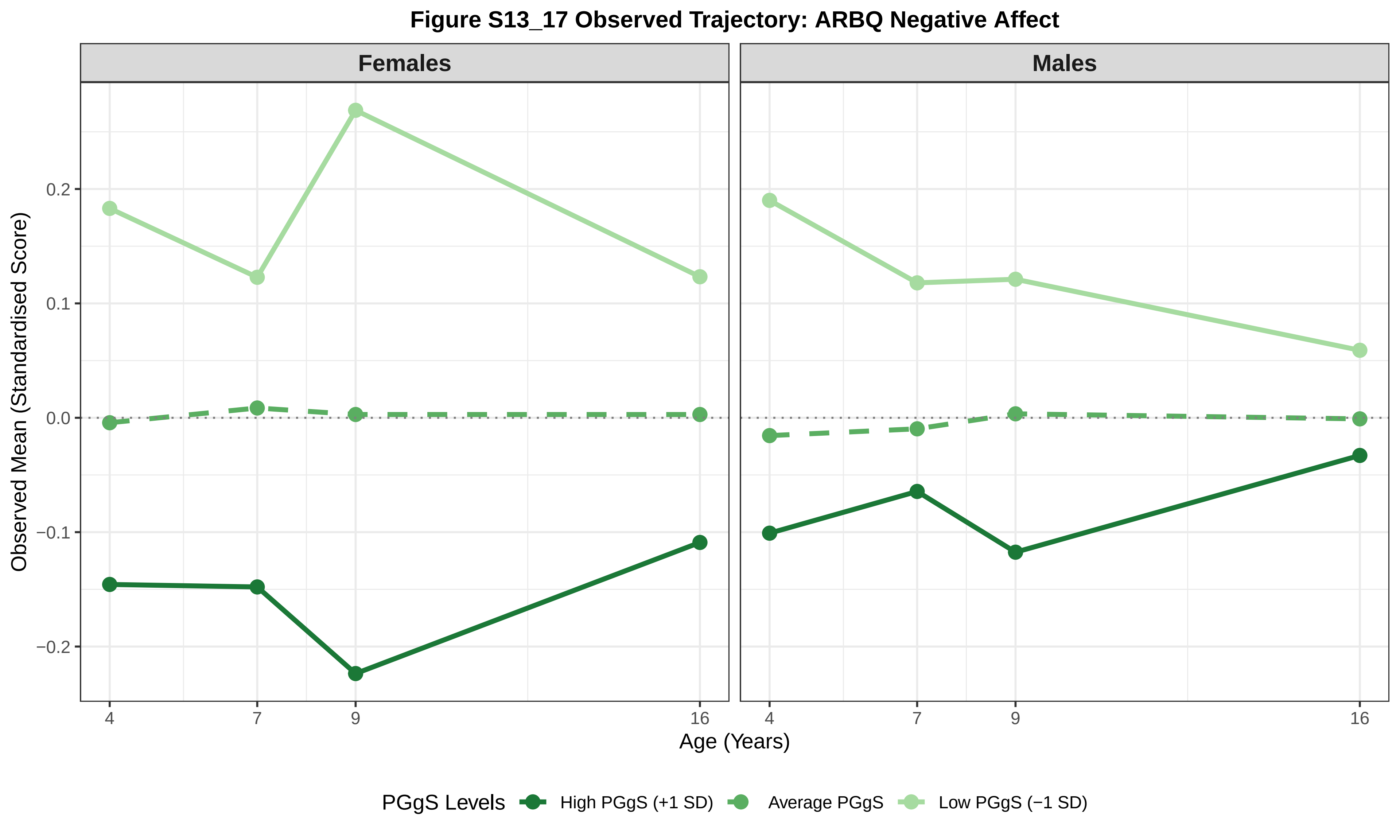
Figure S13_17 Observed Trajectory: ARBQ Negative Affect
Females
Males
Observed Mean (Standardised Score)
0.2
0.1
0.0
−0.1
−0.2
4
7
9
16
Age (Years)
PGgS Levels
High PGgS (+1 SD)
Average PGgS
Low PGgS (−1 SD)

### Figure S13_18 Observed Trajectory: ARBQ Negative Cognition

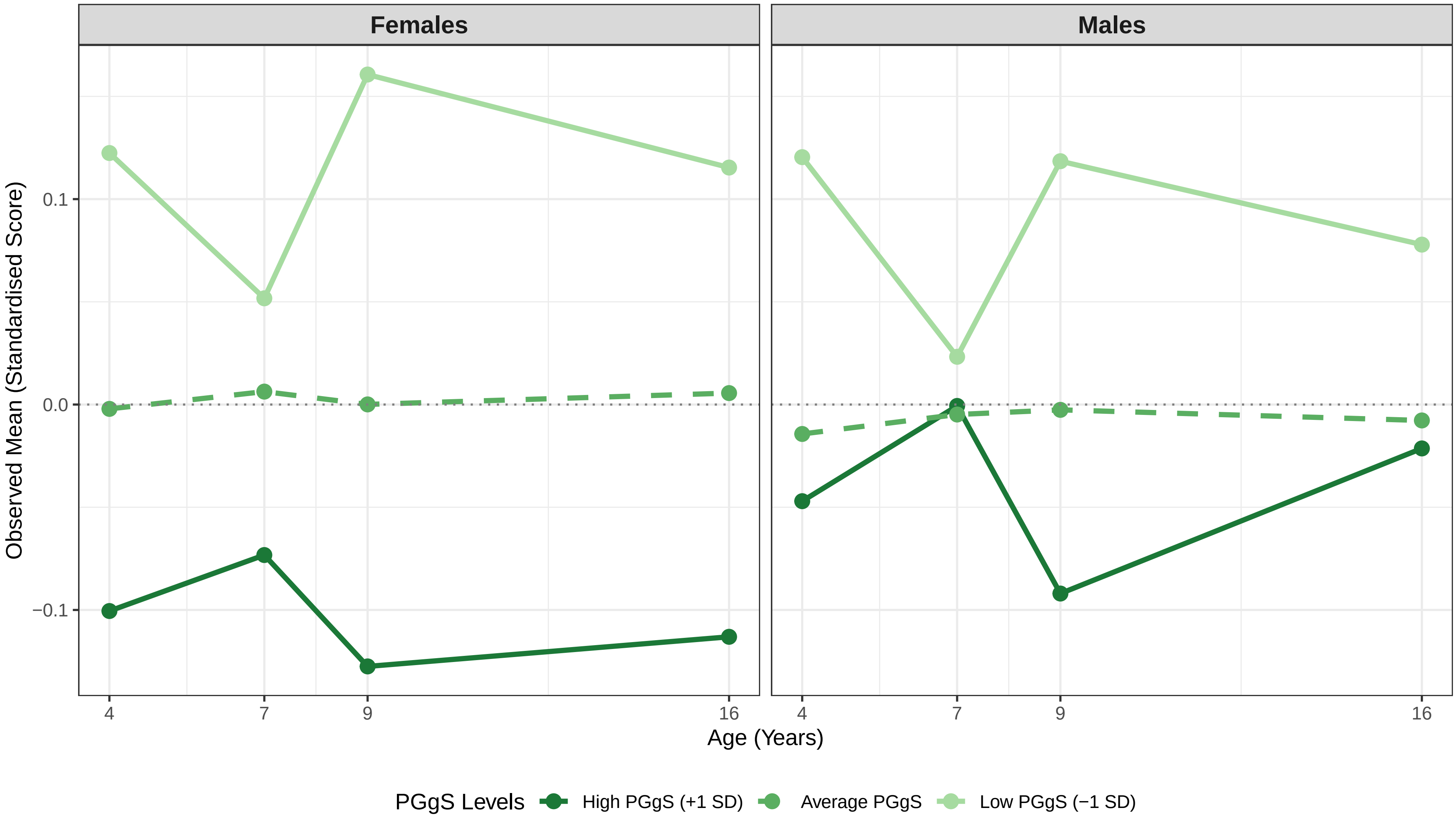

**Figure S13_19 Observed Trajectory: ARBQ Anxiety Total**

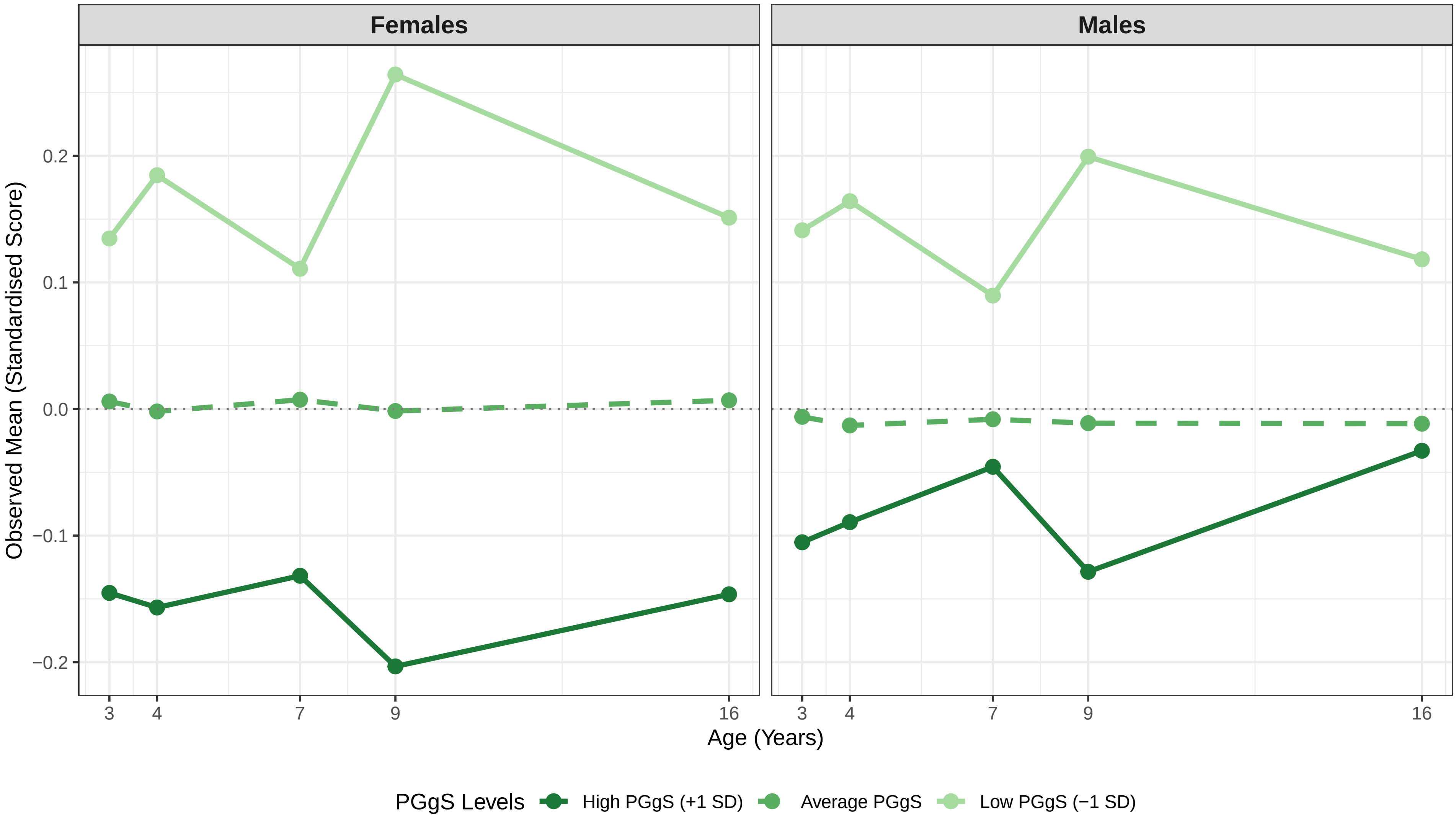

**Figure S13_20 Observed Trajectory: Conners ADHD Inattention**

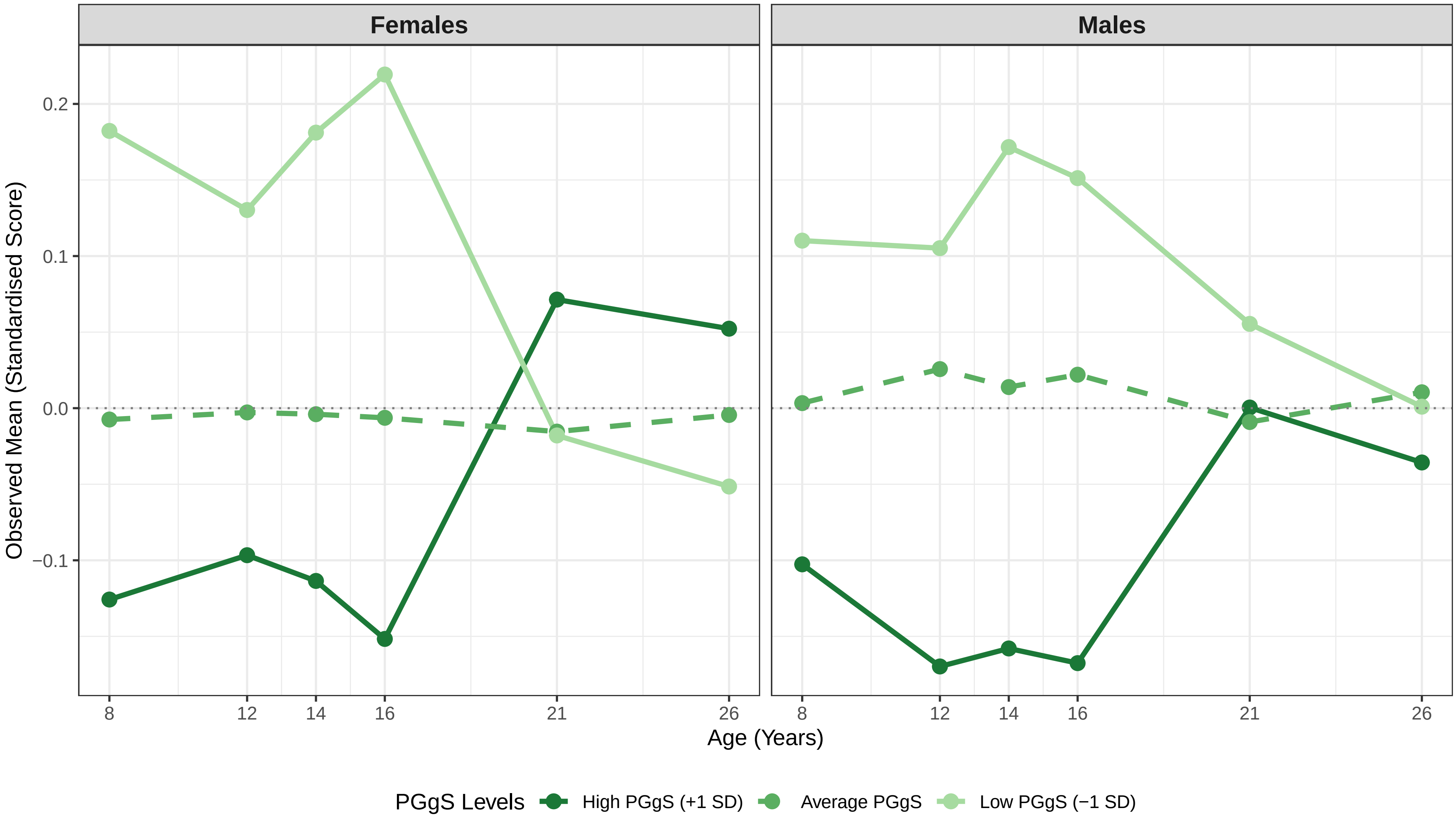

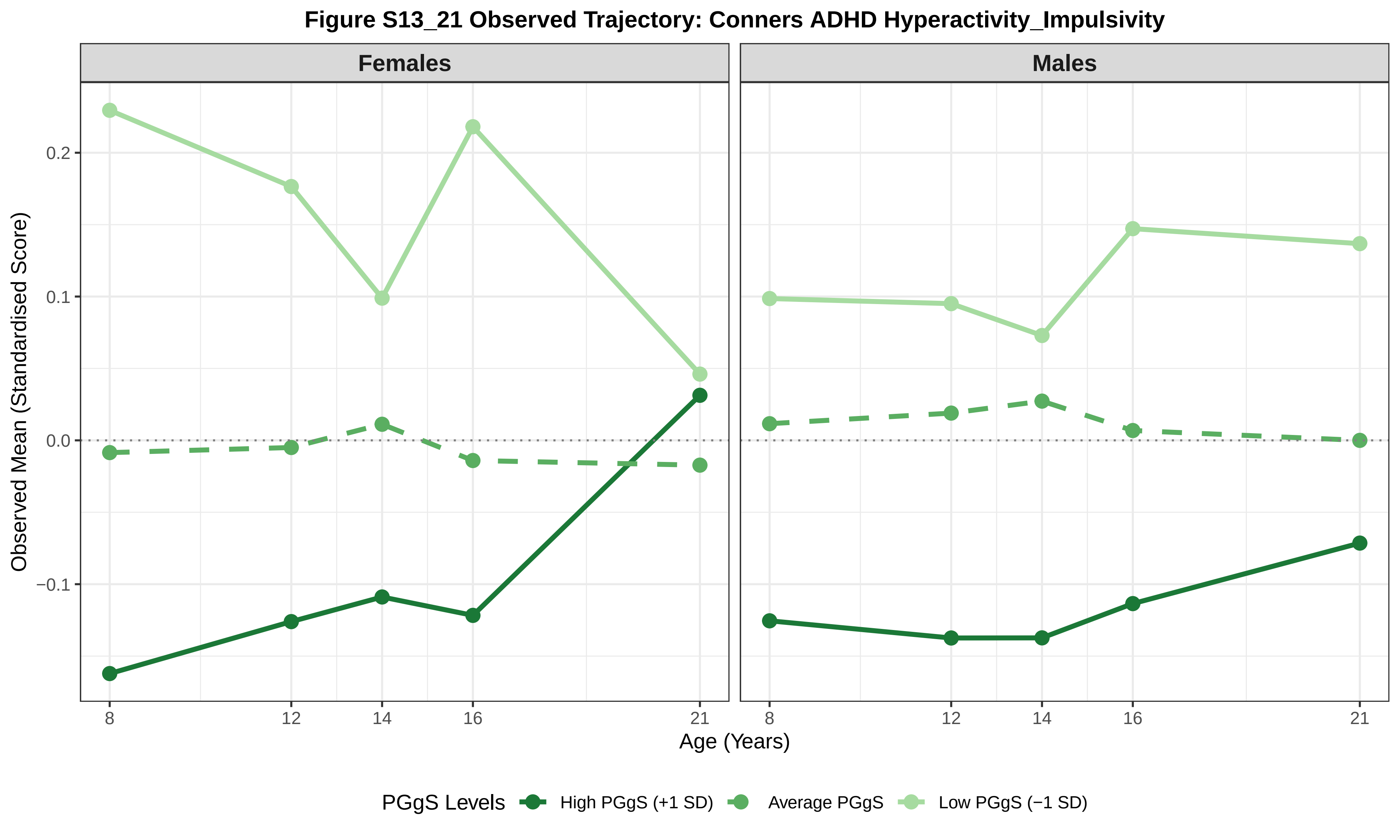
Figure S13_21 Observed Trajectory: Conners ADHD Hyperactivity_Impulsivity
Females
Males
Observed Mean (Standardised Score)
0.2
0.1
0.0
−0.1
8
12
14
16
21
Age (Years)
PGgS Levels
High PGgS (+1 SD)
Average PGgS
Low PGgS (−1 SD)

Figure S13_22 Observed Trajectory: Conners ADHD Total

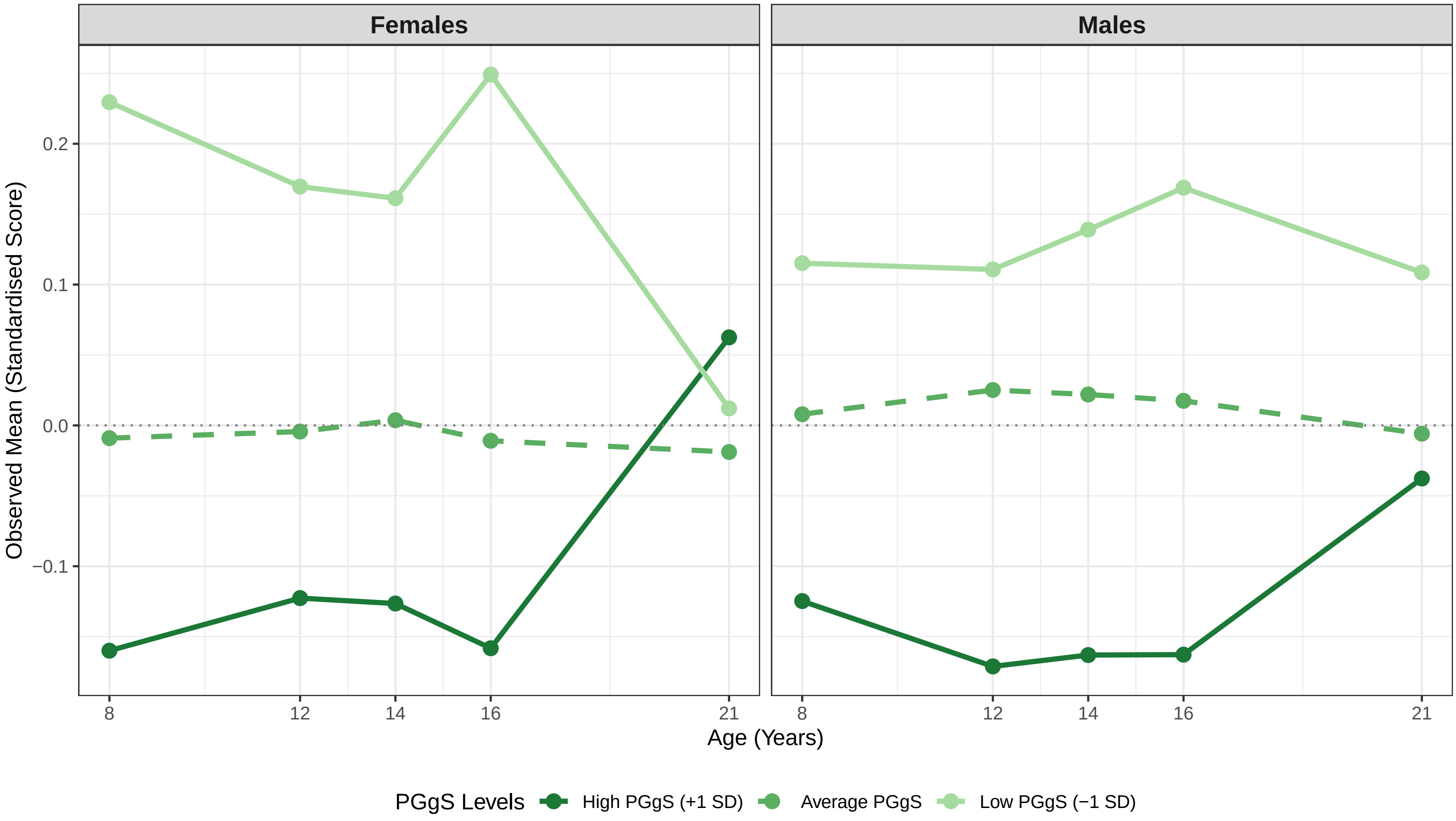

### Figure S13_23 Observed Trajectory: Height

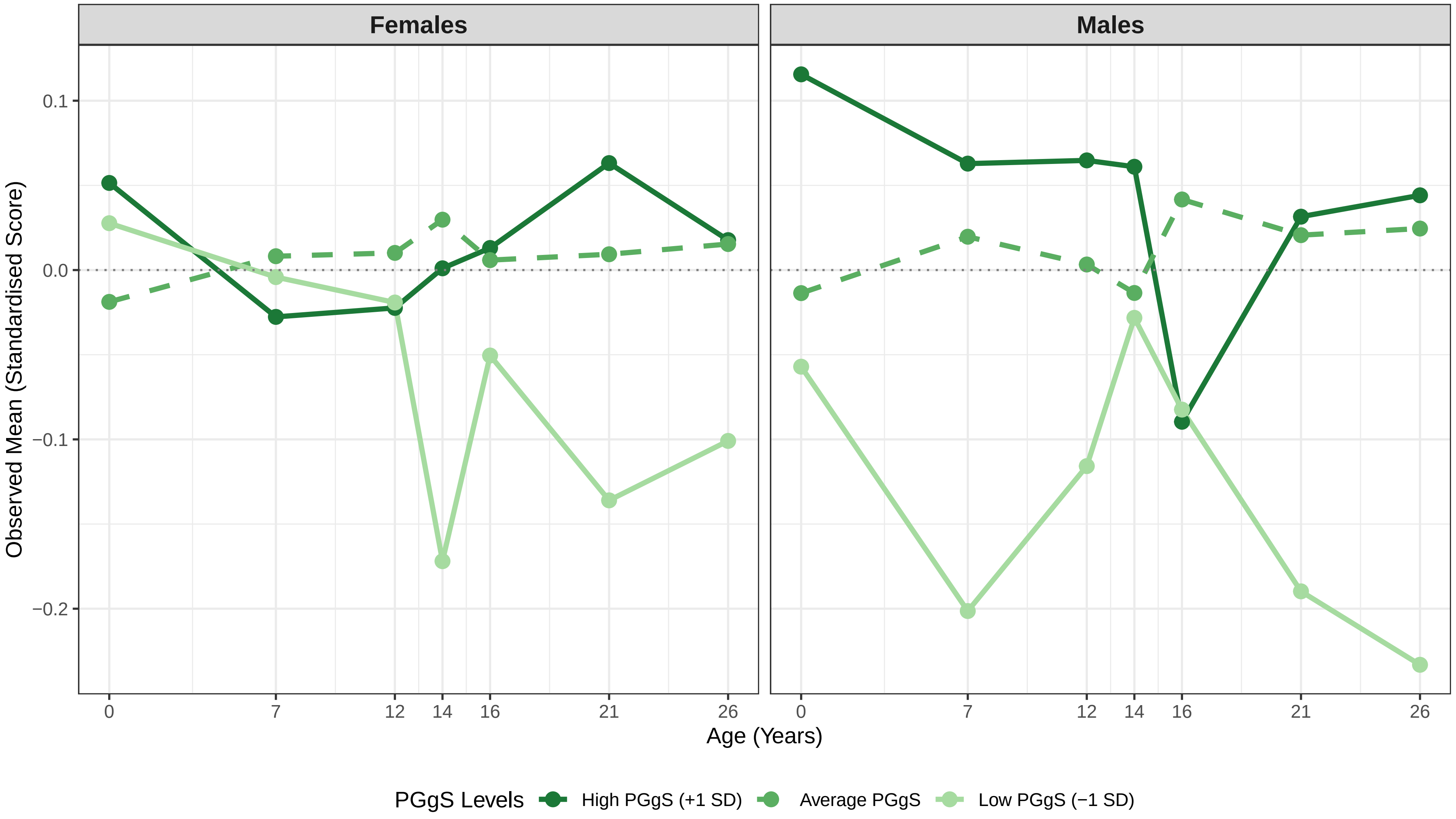

#### Figure S13_24 Observed Trajectory: BMI (as weight at birth)

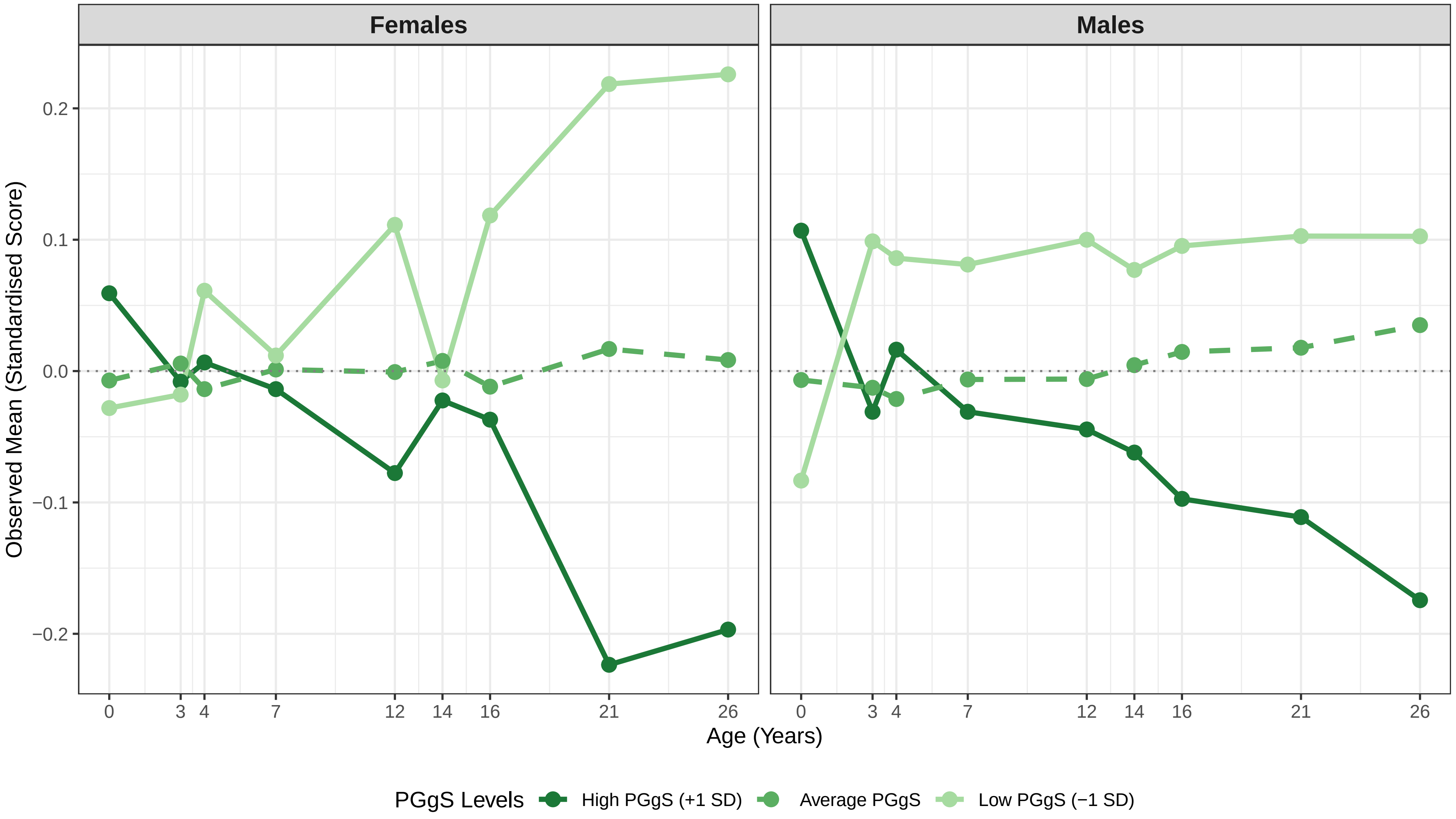
