## Supplementary Materials for "Charting the cognitive development of children using adult ‘polygenic g scores’"

Yujing Lin [1], Robert Plomin [1]
[1] Social, Genetic and Developmental Psychiatry Centre, King's College London

The Supplementary Materials are organised into three sections: 1) Supplementary Text, 2) Supplementary Tables, and 3) Supplementary Figures

### Table of Contents

***Supplementary Text: Supplementary Phenotypic Measures*** **... 5**
- **ADHD Symptoms** ... 5
- **Anxiety** ... 5
- **Height and BMI** ... 5

**Additional Measures ... 5**
- ***Reading and Mathematics Environment*** ... 5
- ***PISA Measures*** ... 5
- ***Spatial and Mathematical Anxiety*** ... 6

***Supplementary Tables (provided in a separate spreadsheet)*** **... 8**

All supplementary tables referenced in the main text are compiled in the accompanying Excel spreadsheet (SupTables.xlsx). The following list provides an overview of the tables included. ... 8

**Table S1. Sensitivity Analyses Examining Effects of Age, Sex, Zygosity, Twin Birth Order, Genotyping Chip Type, and the First Ten Principal Components on Phenotypes and Polygenic g Score ... 8**

**Table S2. Pearson's r Correlation Matrix Between all Variables and Polygenic Scores used in the Study ... 8**

**Table S3. Sample Characteristics for Phenotypic and Genotypic Outcomes Across the Whole Sample and Subsamples ... 8**

**Table S4. Regression Results of Polygenic g Score Predicting Cognitive, Educational, and Behavioural Outcomes for Whole Sample, Sex-Stratified, and Zygosity-Stratified Samples ... 8**

**Table S5. Standardised Factor Loadings for Cross-Age Latent Factors: Cross-Rater and/or Within-Rater Confirmatory Factor Analyses by Developmental Stage or by Measures of Cognitive, Educational, and Behavioural Outcomes ... 8**

**Table S6. Confirmatory Factor Analysis Model Fit Indices ... 8**

**Table S7. Confirmatory Factor Analysis Model Fit Indices and Polygenic g Score Prediction of Cross-Age Latent Factors ... 8**

**Table S8. Latent Growth Curve Model Fit Indices and Polygenic g Score Prediction of Latent Intercepts and Slopes ... 8**

**Table S9. Sex-Stratified Latent Growth Curve Model Fit Indices and Polygenic g Score Prediction of Latent Intercepts and Slopes ... 8**

**Table S10. Polynomial Regression Results Testing Nonlinear Associations Between Polygenic g Score and Cognitive, Educational, and Behavioural Outcomes ... 8**

### Supplementary Text: Supplementary Phenotypic Measures

#### ADHD Symptoms

Attention-Deficit/Hyperactivity Disorder (ADHD) symptoms were assessed using the Conners Rating Scale based on Diagnostic and Statistical Manual of Mental Disorders, Fourth Edition (DSM-IV) criteria (Conners, 2003, 2008). The standard 18-item scale comprises two 9-item subscales measuring hyperactivity and inattentive behaviours.

Parent ratings were obtained at ages 8, 12, 14, 16, and 21, teacher ratings at age 14, and self-ratings at ages 14 and 21, all using the full 18-item scale. At age 26, only the 9-item inattention subscale was administered via self-report (with 2 additional quality control items included in the questionnaire).

#### Anxiety

Anxiety was assessed using the Anxiety-Related Behaviours Questionnaire (ARBQ), a measure designed to capture anxiety symptoms and anxiety-related temperamental traits in the general population (Eley et al., 2003). The ARBQ comprises five subscales: negative cognition, negative affect, fear, social anxiety (shyness), and obsessive-compulsive behaviours (OCB). Psychometric properties and construct validity of the measure have been reported in previous TEDS publications (Hallett et al., 2009; Trzaskowski et al., 2012).

Parent ratings were collected at ages 3 (5 items), 4 (12 items), 7 (21 items), 9 (20 items), and 16 (19 items). Teacher ratings (17 items) were collected at age 7. The questionnaire's length and content were adapted for developmental appropriateness by the TEDS researchers. At age 3, only the fear and social anxiety subscales were assessed; the OCB, negative affect, and negative cognition subscales were added at age 4. Total anxiety scores were calculated as the standardised mean of the available subscales for each participant at each age.

#### Height and BMI

Parent-reported birth weight (grams) and length (centimetres) were collected retrospectively during the first contact at 18 months. Height was also collected in centimetres at ages 7, 12, 14, 16, 21, and 26. Body Mass Index (BMI) was calculated at ages 3, 4, 7, 12, 14, 16, 21, and 26.

#### Additional Measures

##### *Reading and Mathematics Environment*

Reading and mathematics environments were assessed at ages 10 and 12 using child-completed web-based questionnaires adapted from the National Assessment of Educational Progress (NAEP) Grade 4 student background questionnaires for mathematics (https://nces.ed.gov/nationsreportcard/pdf/05BQstudentG4math.pdf) and reading (https://nces.ed.gov/nationsreportcard/pdf/05BQstudentG4read.pdf). The adapted questionnaire included 15 items assessing children’s experiences with mathematics and reading environments at home and school.

##### *PISA Measures*

At age 16, several measures were adapted from the Programme for International Student Assessment (PISA; OECD: https://www.oecd.org/en/about/programmes/pisa.html), drawing on items from the 2000, 2003, and 2006 student questionnaires. The measures included homework behaviours (5 items), attitudes toward school (4 items), mathematics self-efficacy (8 items), mathematics interest (3 items), and time spent on mathematics (3 items).

***Spatial and Mathematical Anxiety***

At age 18, spatial and mathematical anxiety was measured using the Self-Perceived Ability and Anxiety (SPAA) questionnaire, developed by the TEDS team as part of the spatial ability testing battery based on existing scales (Hopko et al., 2003; Lawton, 1994). The questionnaire included subscales assessing spatial ability (8 items), spatial anxiety (10 items), and mathematics anxiety (9 items).

***Financial and Socioeconomic Measures***

At age 21, participants completed self-report measures assessing financial wellbeing, literacy, and attitudes. Financial wellbeing was measured using a 5-item shortened version of the Contentment with Life Assessment Scale (CLAS), assessing satisfaction with their financial situation (Lavallee et al., 2007). Financial literacy regarding products was measured using a 13-item adaptation of the OECD Financial Literacy instrument, shortened from 15 items with an additional quality control item (https://www.oecd.org/en/topics/financial-education.html). General money attitudes and behaviours were assessed using six relevant items from the OECD scale.

At age 26, socioeconomic outcomes were captured using derived indicators of economic vulnerability and a twin-specific SES composite. Economic vulnerability was quantified as an ordinal score (0–4), summing indicators of NEET status (not in education, employment, or training), low income, receipt of benefits, presence of children, and zero-hours employment. NEET status was coded as a binary variable, indicating whether the participant was not in education, employment, or training. The twin SES composite was a standardised continuous measure derived from educational attainment, income level, and economic vulnerability, calculated to reflect the individual twin's socioeconomic status. All variables were devised by TEDS researchers using survey responses and administrative coding according to the Standard Occupational Classification 2000 (https://www.ons.gov.uk/methodology/classificationsandstandards/standardoccupationalclassificationsoc/socarchive).

***Wellbeing and Noncognitive Traits***

At age 16, wellbeing and non-cognitive traits were assessed using a combination of established self-report scales. The measures included a 21-item life satisfaction scale, 6-item hopefulness scale, 6-item gratitude scale, 7-item curiosity scale with subscales for exploration and flow, 4-item subjective happiness scale, 13-item grit and ambition scale, 6-item optimism scale, and a 10-item academic self-concept scale.

Life satisfaction was measured using the Multidimensional Students' Life Satisfaction Scale, assessing satisfaction with family, friends, living environment, school, and self, as well as overall life satisfaction (Huebner, 1994). Hopefulness was assessed using the Children's Hope Scale, comprising agency and pathways subscales (Snyder et al., 1997). Additional wellbeing measures included Subjective Happiness (Lyubomirsky & Lepper, 1999), Gratitude (McCullough et al., 2002), and Curiosity (Kashdan et al., 2004), which captured both exploration and flow components.

Non-cognitive traits were measured using Ambition and Grit scales (Duckworth et al., 2007; Duckworth & Quinn, 2009), with Grit subdivided into consistency of interests and perseverance of effort subscales. Optimism was assessed using the Life Orientation Test–Revised (Scheier et al., 1994), and

Academic Self-Concept using a shortened version of Perceptions of Self as Learners and Problem Solvers scale (Burden, 1998).

Finally, composite indices of psychological wellbeing and subjective wellbeing were derived from relevant subsets of these wellbeing measures, including life satisfaction, hopefulness, gratitude, curiosity, and subjective happiness.

At age 21, several measures overlapped with those administered at age 16, alongside additional assessments of self-control, purpose in life, and depressive symptoms. Self-control was measured using a 6-item version of the Brief Self-Control Scale (Tangney et al., 2004), shortened from the original 13-item measure. The Consideration of Future Consequences Scale was abbreviated to 4 items, with the addition of a quality control item (Strathman et al., 1994). Life goals were assessed using a 9-item adaptation of the GOALS questionnaire, translated into English and focused on fulfilment- and relationship-oriented goals, excluding items related to social status or likelihood of success (Pöhlmann & Brunstein, 1997). Purpose in Life was measured using a 5-item short form of the Purpose in Life Test (Crumbaugh & Maholick, 1964). Ambition was assessed using the same 5-item Grit measure previously administered at age 16. Finally, depressive symptoms were measured using an 8-item version of the Short Mood and Feelings Questionnaire, adapted from the original 13-item version with one additional quality control item (Angold et al., 1995).

At age 26, wellbeing and mental health were re-evaluated. Subjective wellbeing (quality of life) was measured using a 3-item scale including two euthymic (positive emotion) items and one eudaimonic (meaning) item, adapted from the UK Biobank and Genetic Links to Anxiety and Depression (GLAD) Study mental health questionnaires. Depressive symptoms were assessed using the 13-item Mood and Feelings Questionnaire, a validated screening measure for current depressive symptoms, also including a quality control item (Angold et al., 1995).

### Supplementary Tables (provided in a separate spreadsheet)

All supplementary tables referenced in the main text are compiled in the accompanying Excel spreadsheet (SupTables.xlsx). The following list provides an overview of the tables included.

Table S1. Sensitivity Analyses Examining Effects of Age, Sex, Zygosity, Twin Birth Order, Genotyping Chip Type, and the First Ten Principal Components on Phenotypes and Polygenic g Score

Table S2. Pearson's r Correlation Matrix Between all Variables and Polygenic Scores used in the Study

Table S3. Sample Characteristics for Phenotypic and Genotypic Outcomes Across the Whole Sample and Subsamples

Table S4. Regression Results of Polygenic g Score Predicting Cognitive, Educational, and Behavioural Outcomes for Whole Sample, Sex-Stratified, and Zygosity-Stratified Samples

Table S5. Standardised Factor Loadings for Cross-Age Latent Factors: Cross-Rater and/or Within-Rater Confirmatory Factor Analyses by Developmental Stage or by Measures of Cognitive, Educational, and Behavioural Outcomes

Table S6. Confirmatory Factor Analysis Model Fit Indices

Table S7. Confirmatory Factor Analysis Model Fit Indices and Polygenic g Score Prediction of Cross-Age Latent Factors

Table S8. Latent Growth Curve Model Fit Indices and Polygenic g Score Prediction of Latent Intercepts and Slopes

Table S9. Sex-Stratified Latent Growth Curve Model Fit Indices and Polygenic g Score Prediction of Latent Intercepts and Slopes

Table S10. Polynomial Regression Results Testing Nonlinear Associations Between Polygenic g Score and Cognitive, Educational, and Behavioural Outcomes

Table S11. Mean Differences in Cognitive, Educational, and Behavioural Outcomes Between Top and Bottom Deciles of Polygenic g Score

Table S12. Phenotypic Differences Between High and Low Polygenic g Score Groups

### Supplementary Figures

All supplementary figures referenced in the main text are presented in this supplement and in an accompanying compiled PDF file: SupFigures.pdf.

**Figure S1. Heatmaps of Pearson correlations between the polygenic g score and observed phenotypic outcomes (provided in a separate PDF file)**

Nine correlation matrices compiled in the PDF file correspond to the correlation matrix presented in Supplementary Table S2. Each matrix presents a heatmap of Pearson's r correlation coefficients among repeated measures or composite variables within a trait domain. Above the diagonal, the shape and colour indicate the strength and direction of the correlation; below the diagonal, the corresponding correlation coefficients are displayed.

The heatmaps include correlations between the polygenic g score, Savage g PGS, Okbay EA PGS and:

1. Cognitive ability composites — general cognitive ability (g), verbal ability, and nonverbal ability at each age and across ages
2. General cognitive ability composites at each age and across ages
3. Individual verbal tests administered at each age and across ages
4. Individual nonverbal tests administered at each age and across ages
5. Educational achievement and attainment outcomes at each age and across ages
6. Strengths and Difficulties Questionnaire (SDQ) subscales and total problem score at each age and across ages
7. Anxiety-Related Behaviours Questionnaire (ARBQ) subscales and total anxiety score at each age and across ages
8. Conners Rating Scale subscales and total ADHD score at each age and across ages
9. Anthropometric traits — height and BMI at each age and weight at birth

**Figure S2 to S7. Polygenic g score prediction of phenotypic outcomes in the whole sample**

Figures S2 to S7 present the standardised beta coefficients and 95% bootstrapped confidence intervals, estimated using percentile bootstrapping with 1000 iterations. Analyses were conducted in the unrelated sample by randomly selecting one twin from each pair. Each figure corresponds to a specific trait domain. Raters are specified in the plot when multiple raters are available for the same measure. When only one rater is present across ages (typically the child), or when rater differences are not relevant (e.g., height and BMI), the rater is not specified. Please refer to the measure descriptions for abbreviations of the variable names.

The main results for general cognitive ability, verbal and nonverbal ability composites, educational outcomes, and behavioural outcomes (SDQ) are presented in the main text (Figure 2a to 2c). Numerical estimates, model variance explained, and statistical significance values for these analyses, as well as sex-stratified and zygosity-stratified results, are reported in Table S4. The following list provides an overview of Figures S2 to S7:

- Figure S2. Individual verbal test scores
- Figure S3. Individual nonverbal test scores
- Figure S4. Anxiety subscales and total score
- Figure S5. ADHD subscales and total score
- Figure S6. Height, birth weight, and BMI from age 3 onward
- Figure S7. Other outcomes (educational, environmental, behavioural, and wellbeing measures)

### Figure S2. Polygenic prediction of individual verbal test scores

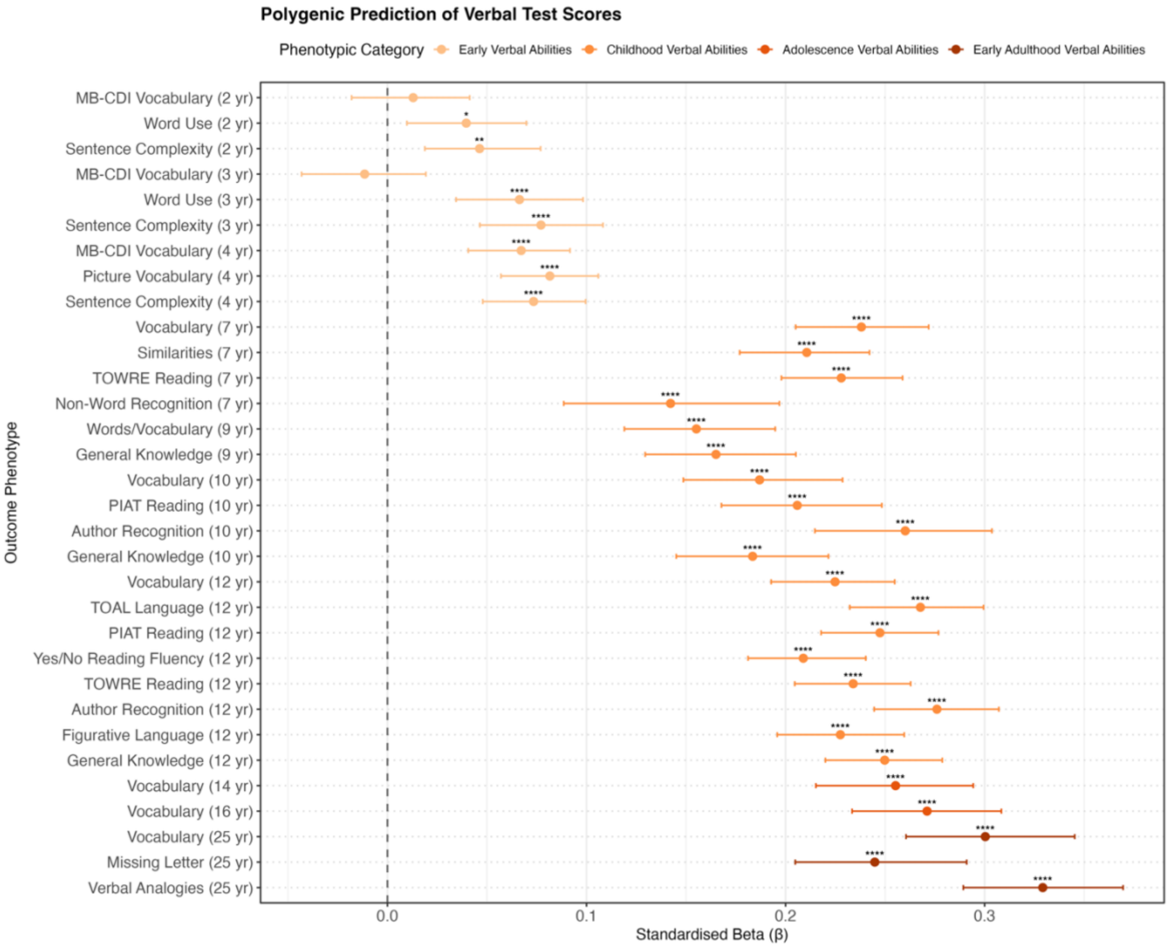

### Figure S3. Polygenic prediction of individual nonverbal test scores

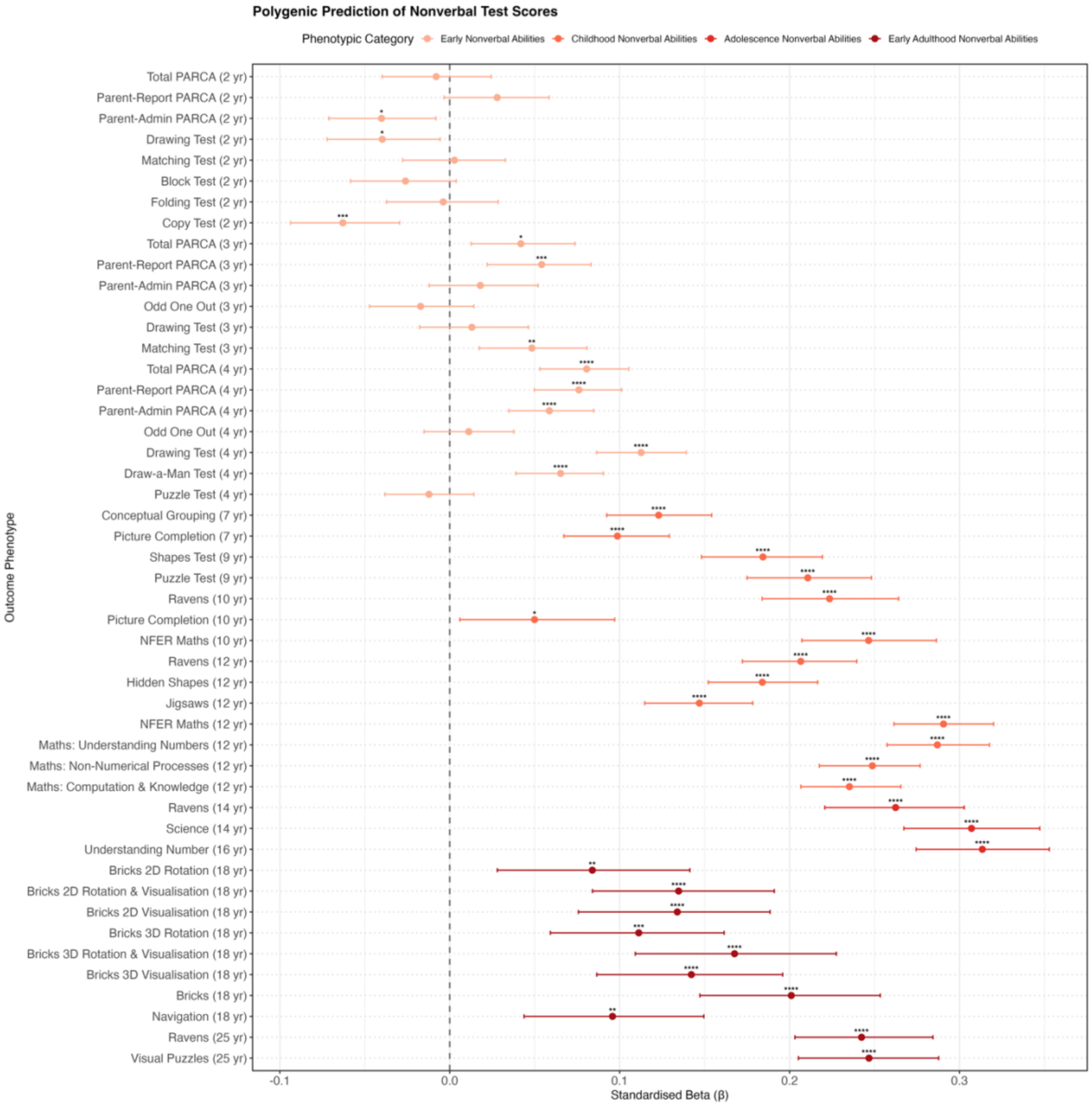

### Figure S4. Polygenic prediction of anxiety subscales and total score

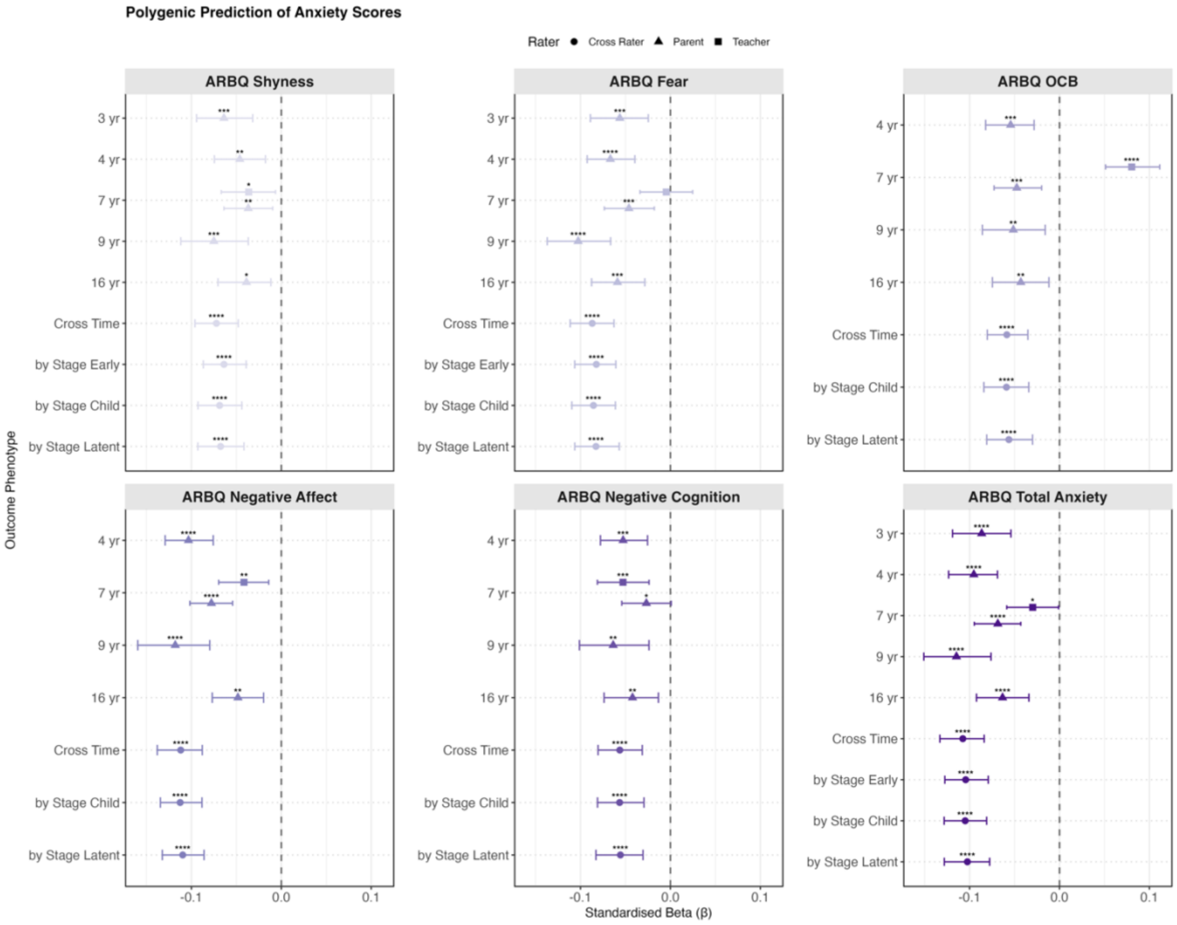

### Figure S5. Polygenic prediction of ADHD subscales and total score

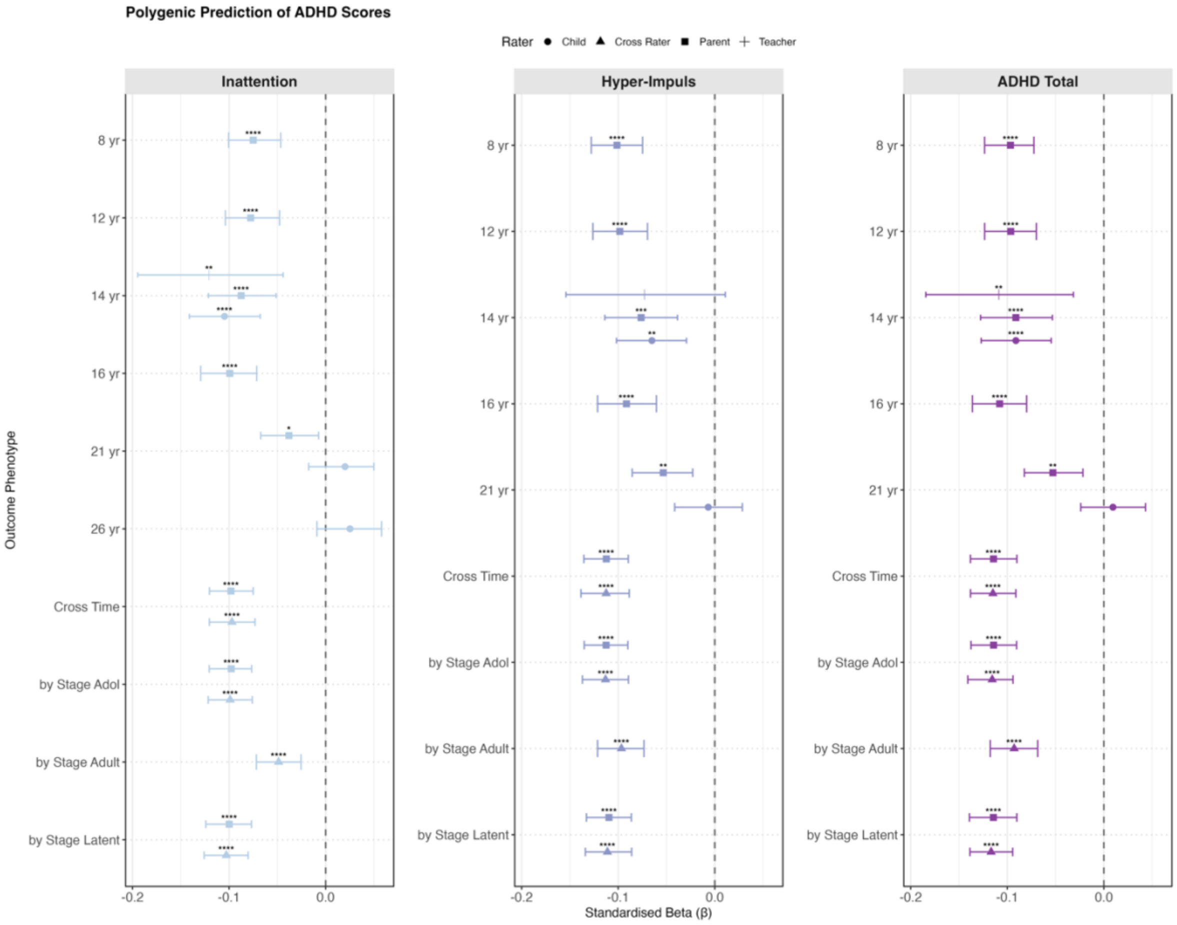

### Figure S6. Polygenic prediction of height and BMI

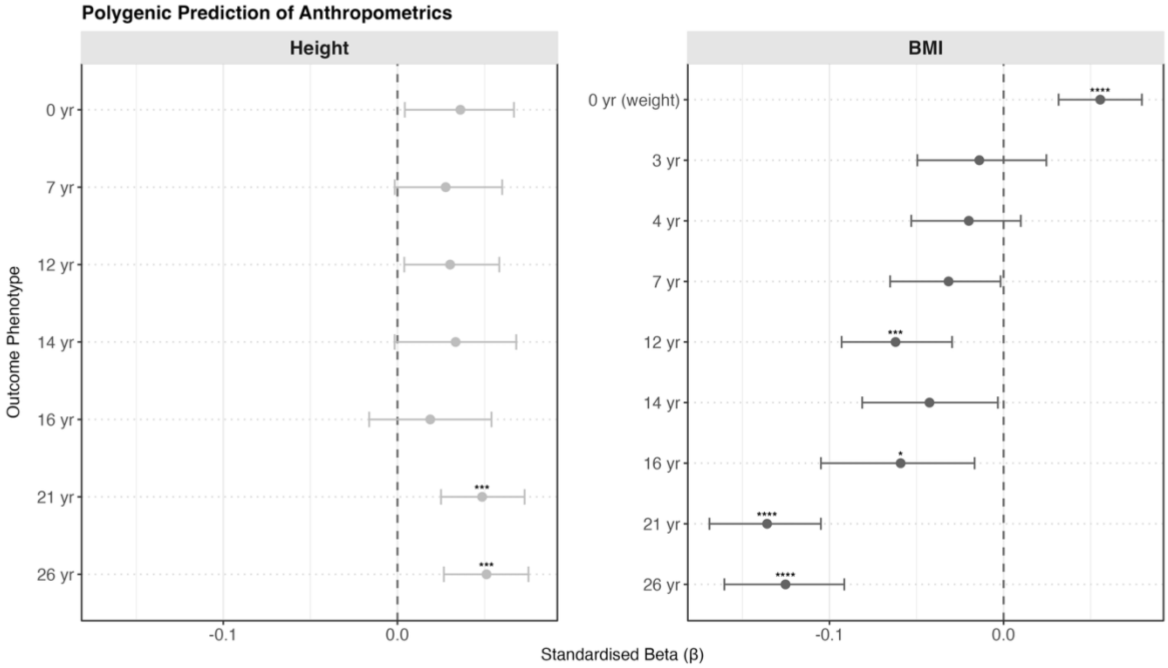

### Figure S7. Polygenic prediction of other outcomes (educational, environmental, behavioural, and wellbeing measures)

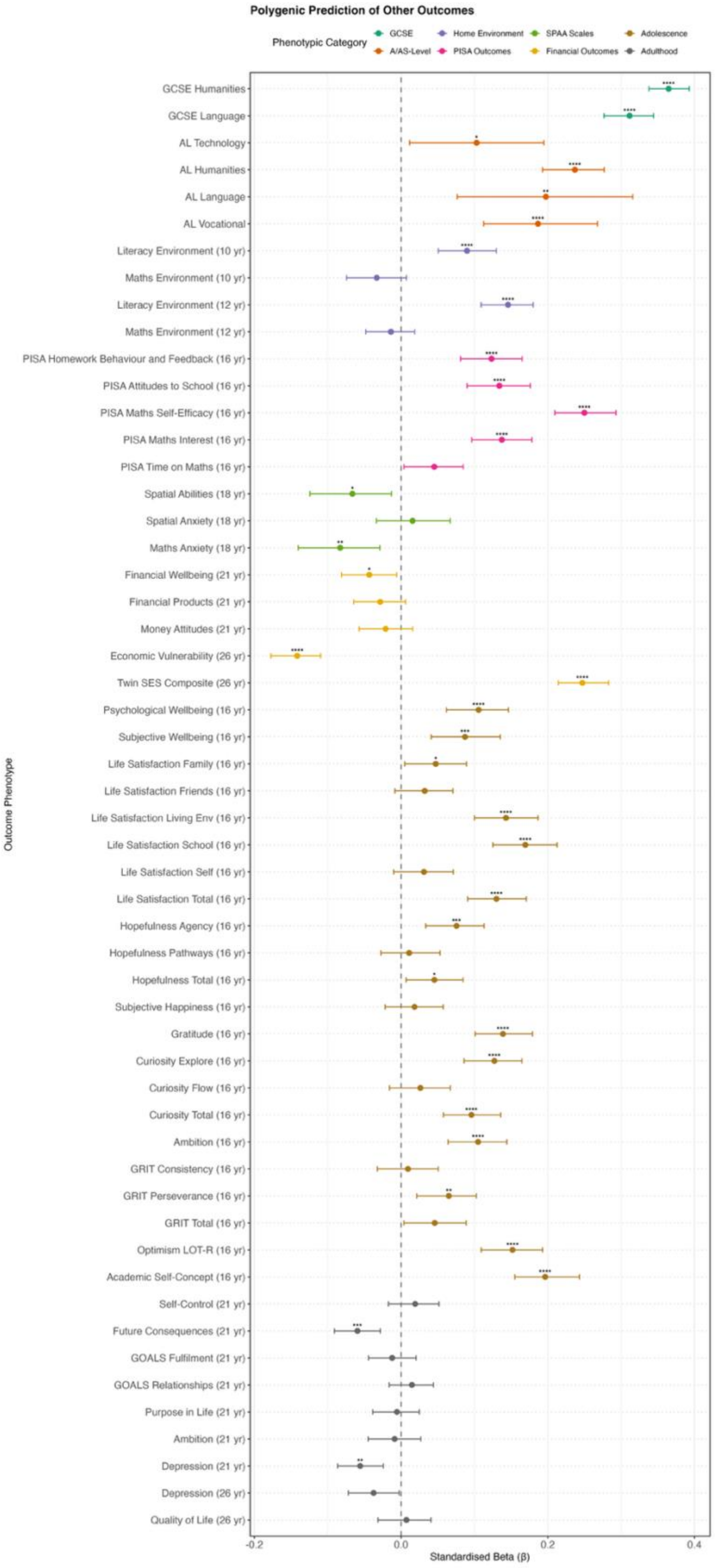

### Figure S8. Confirmatory factor analysis (CFA) path diagrams (provided in a separate PDF file)

All 73 CFAs are compiled in the PDF file ('SupFigures.pdf'), with models labelled Figure S8-1 through Figure S8-73. Path coefficients are standardised estimates. Models were fitted using confirmatory factor analysis in the lavaan package with full information maximum likelihood (FIML) estimation. The full twin sample was used with clustering by family specified to account for non-independence and increase statistical power. Complete numerical results are detailed in Tables S5 to S6.

All plots are constructed using the *semPath*s function from the *semPlot* package. Latent constructs are depicted as ovals, representing unobserved traits extracted from multiple observed indicators, which are shown as rectangles. These models are structured to extract a cross-time latent factor, or higher-order factors that account for specific developmental stages and measurement types. Observed variable labels specify the age of measurement and, for behavioural outcomes, the rater (e.g., yr7P for parent-rated phenotype at age 7, yr7T for teacher-rated). When multiple raters are used, the rater is specified in the label (P = Parent, T = Teacher, C = Child/Self); otherwise, a single rater (typically the child) is used across time, or non-overlapping raters are used developmentally (typically parent at younger ages, child at older ages).

All coefficients are standardised. Arrows from the latent factors (ovals) to observed variables (rectangles) represent factor loadings (also in Table S5), indicating the strength of association between each measure and the underlying latent construct. The dotted line on the path to one indicator is a scaling artefact (this variable serves as the reference indicator) and does not affect the interpretation of the standardised loading.

Residual variances are displayed for both latent and observed variables. In higher-order models, values on the arrows pointing to lower-order latent factors represent their standardised residual variances (the proportion of variance unexplained by the higher-order factor). Values on arrows pointing to observed variables at the bottom represent their standardised residual variances (the proportion of variance unexplained by the latent factor), calculated as $1 - (\text{loading})^2$.

Small triangles represent the constant (1) used in the model specification to estimate intercepts (means) of variables. These are standard CFA notation and indicate that means are being modelled.

Path colours denote the direction of associations, with green indicating positive associations and red indicating negative associations.

The following list provides an overview of Figures S8-1 to S8-73 for confirmatory factor analyses. CTCR = cross-time cross rater

Figure S8-1. General Cognitive Ability (g) - Overall
Figure S8-2. General Cognitive Ability (g) - Stage
Figure S8-3. General Cognitive Ability (g) - Method
Figure S8-4. Verbal Ability - Overall
Figure S8-5. Verbal Ability - Stage
Figure S8-6. Verbal Ability - Method
Figure S8-7. Nonverbal Ability - Overall
Figure S8-8. Nonverbal Ability - Stage
Figure S8-9. Nonverbal Ability - Method
Figure S8-10. English Achievement Latent (Teacher 7-12)
Figure S8-11. Maths Achievement Latent (Teacher 7-12)
Figure S8-12. Science Achievement Latent (Teacher 9-12)
Figure S8-13. Core-Subject Achievement Latent (Teacher 7-12)
Figure S8-14. ARBQ Shyness - CTCR Overall
Figure S8-15. ARBQ Shyness - CTCR Stage
Figure S8-16. ARBQ Fear - CTCR Overall
Figure S8-17. ARBQ Fear - CTCR Stage
Figure S8-18. ARBQ Obsessive-Compulsive - CTCR Overall
Figure S8-19. ARBQ Obsessive-Compulsive - CTCR Stage
Figure S8-20. ARBQ Negative Affect - CTCR Overall
Figure S8-21. ARBQ Negative Affect - CTCR Stage
Figure S8-22. ARBQ Negative Cognition - CTCR Overall
Figure S8-23. ARBQ Negative Cognition - CTCR Stage
Figure S8-24. ARBQ Anxiety Total - CTCR Overall
Figure S8-25. ARBQ Anxiety Total - CTCR Stage
Figure S8-26. Conners Inattention - CTCR Overall
Figure S8-27. Conners Inattention - CTCR Stage
Figure S8-28. Conners Inattention - Parent Overall
Figure S8-29. Conners Inattention - Parent Stage
Figure S8-30. Conners Hyperactivity-Impulsivity - CTCR Overall
Figure S8-31. Conners Hyperactivity-Impulsivity - CTCR Stage
Figure S8-32. Conners Hyperactivity-Impulsivity - Parent Overall

Figure S8-33. Conners Hyperactivity-Impulsivity - Parent Stage

Figure S8-34. Conners Total - CTCR Overall

Figure S8-35. Conners Total - CTCR Stage

Figure S8-36. Conners Total - Parent Overall

Figure S8-37. Conners Total - Parent Stage

Figure S8-38. SDQ Conduct - CTCR Overall

Figure S8-39. SDQ Conduct - CTCR Stage

Figure S8-40. SDQ Conduct - Parent Overall

Figure S8-41. SDQ Conduct - Parent Stage

Figure S8-42. SDQ Conduct - Teacher Overall

Figure S8-43. SDQ Conduct - Child Overall

Figure S8-44. SDQ Emotion - CTCR Overall

Figure S8-45. SDQ Emotion - CTCR Stage

Figure S8-46. SDQ Emotion - Parent Overall

Figure S8-47. SDQ Emotion - Parent Stage

Figure S8-48. SDQ Emotion - Teacher Overall

Figure S8-49. SDQ Emotion - Child Overall

Figure S8-50. SDQ Hyperactivity - CTCR Overall

Figure S8-51. SDQ Hyperactivity - CTCR Stage

Figure S8-52. SDQ Hyperactivity - Parent Overall

Figure S8-53. SDQ Hyperactivity - Parent Stage

Figure S8-54. SDQ Hyperactivity - Teacher Overall

Figure S8-55. SDQ Hyperactivity - Child Overall

Figure S8-56. SDQ Peer Problems - CTCR Overall

Figure S8-57. SDQ Peer Problems - CTCR Stage

Figure S8-58. SDQ Peer Problems - Parent Overall

Figure S8-59. SDQ Peer Problems - Parent Stage

Figure S8-60. SDQ Peer Problems - Teacher Overall

Figure S8-61. SDQ Peer Problems - Child Overall

Figure S8-62. SDQ Prosocial - CTCR Overall

Figure S8-63. SDQ Prosocial - CTCR Stage

Figure S8-64. SDQ Prosocial - Parent Overall

Figure S8-65. SDQ Prosocial - Parent Stage

Figure S8-66. SDQ Prosocial - Teacher Overall

Figure S8-67. SDQ Prosocial - Child Overall

Figure S8-68. SDQ Total Problems - CTCR Overall

Figure S8-69. SDQ Total Problems - CTCR Stage

Figure S8-70. SDQ Total Problems - Parent Overall

Figure S8-71. SDQ Total Problems - Parent Stage

Figure S8-72. SDQ Total Problems - Teacher Overall

Figure S8-73. SDQ Total Problems - Child Overall

**Figure S9. Confirmatory factor analysis (CFA) with polygenic g score prediction path diagrams (provided in a separate PDF file)**

All 73 CFAs are compiled in the PDF file ('SupFigures.pdf'), with models labelled Figure S9-1 through Figure S9-73. The path diagrams follow the exact same method and logic, except for the additional polygenic g score prediction path. All coefficients are standardised. Complete numerical results are detailed in Table S7.

The observed polygenic g score (PGgS) appears as a rectangle at the top, modelled as a predictor of the latent factor. The path from PGgS to the latent factor shows the standardised regression coefficient ($\beta$), quantifying the direct predictive association between the polygenic score and the latent trait (Table S7). This coefficient is estimated within the CFA framework, where the measurement model and structural paths are fitted simultaneously. This approach differs from a two-stage analysis where factor scores are first extracted and then used in a separate regression, as the CFA accounts for measurement uncertainty in the latent construct during estimation.

Residual variances are displayed for both latent and observed variables. The value on the arrow pointing to the latent factor represents its standardised residual variance (the proportion of variance unexplained by PGgS), calculated as $1 - \beta^2$. Values on arrows pointing to observed variables on the bottom represent their standardised residual variances (the proportion unexplained by the latent factor), calculated as $1 - (\text{loading})^2$.

**Figures S10 to S13. Latent Growth Curve (LGC) Observed and Predicted Trajectories (provided in a separate PDF file)**

Figure S10: Whole Sample Predicted Trajectories (S10-1 to S10-24)

Figure S11: Whole Sample Observed Trajectories (S11-1 to S11-24)

Figure S12: Sex-Stratified Observed Trajectories (S12-1 to S12-24)

Figure S13: Sex-Stratified Predicted Trajectories (S13-1 to S13-24)

All 4*24 LGC predicted and observed trajectory plots are compiled into a separate PDF (‘SupFigures.pdf’). Trajectories are presented to illustrate longitudinal rank-order change across cognitive, educational, behavioural, and anthropometric outcomes as a function of the polygenic g score (PGgS). To facilitate comparison, all outcome variables were standardised (mean = 0, SD = 1) at each age of measurement, ensuring the y-axis represents the outcome in standard deviation (SD) units.

Participants were stratified into three groups based on their PGgS continuous distribution: high PGgS (≥ +1 SD), low PGgS (≤ -1 SD), and average PGgS (within 1 SD of the mean). In all plots, solid lines represent the high and low PGgS groups, while dashed lines represent the average PGgS group. A dotted grey horizontal line denotes the sample mean baseline of 0 SD.

Predicted Trajectories (Figures S10 & S12): These plots depict the modelled trajectories derived directly from the LGC model results (intercepts and slopes).

Observed Trajectories (Figures S11 & S13): These plots depict the raw, empirical data. They are constructed by calculating the mean standardised score of each PGgS group at each available age point.

The following list provides an overview of the 24 plots contained within each of the four main figure files:

1. General Cognitive Ability
2. Verbal Abilities
3. Nonverbal Abilities
4. English Grades
5. Maths Grades
6. Science Grades
7. Core Subject Grades
8. SDQ Conduct
9. SDQ Emotion
10. SDQ Hyperactivity
11. SDQ Peer Problems
12. SDQ Prosocial
13. SDQ Total Problems
14. ARBQ Shyness
15. ARBQ Fear
16. ARBQ Obsessive-Compulsive Behaviours
17. ARBQ Negative Affect
18. ARBQ Negative Cognition
19. ARBQ Anxiety Total
20. Conners ADHD Inattention
21. Conners ADHD Hyperactivity-Impulsivity
22. Conners ADHD Total
23. Height
24. BMI (as weight at birth)

**Figure S14. Mean developmental trajectories for behavioural and anthropometric outcomes in individuals with high and low polygenic g scores**

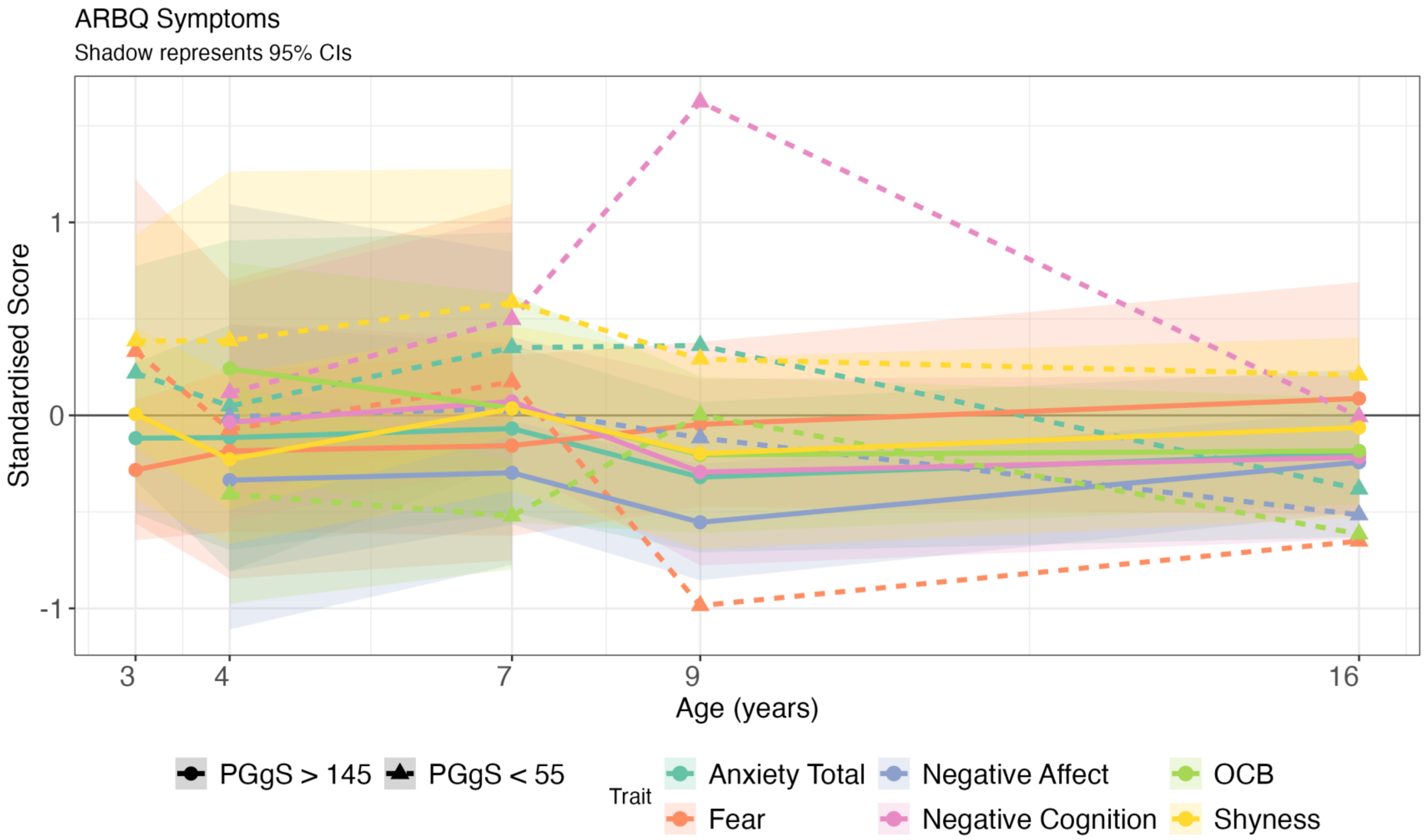

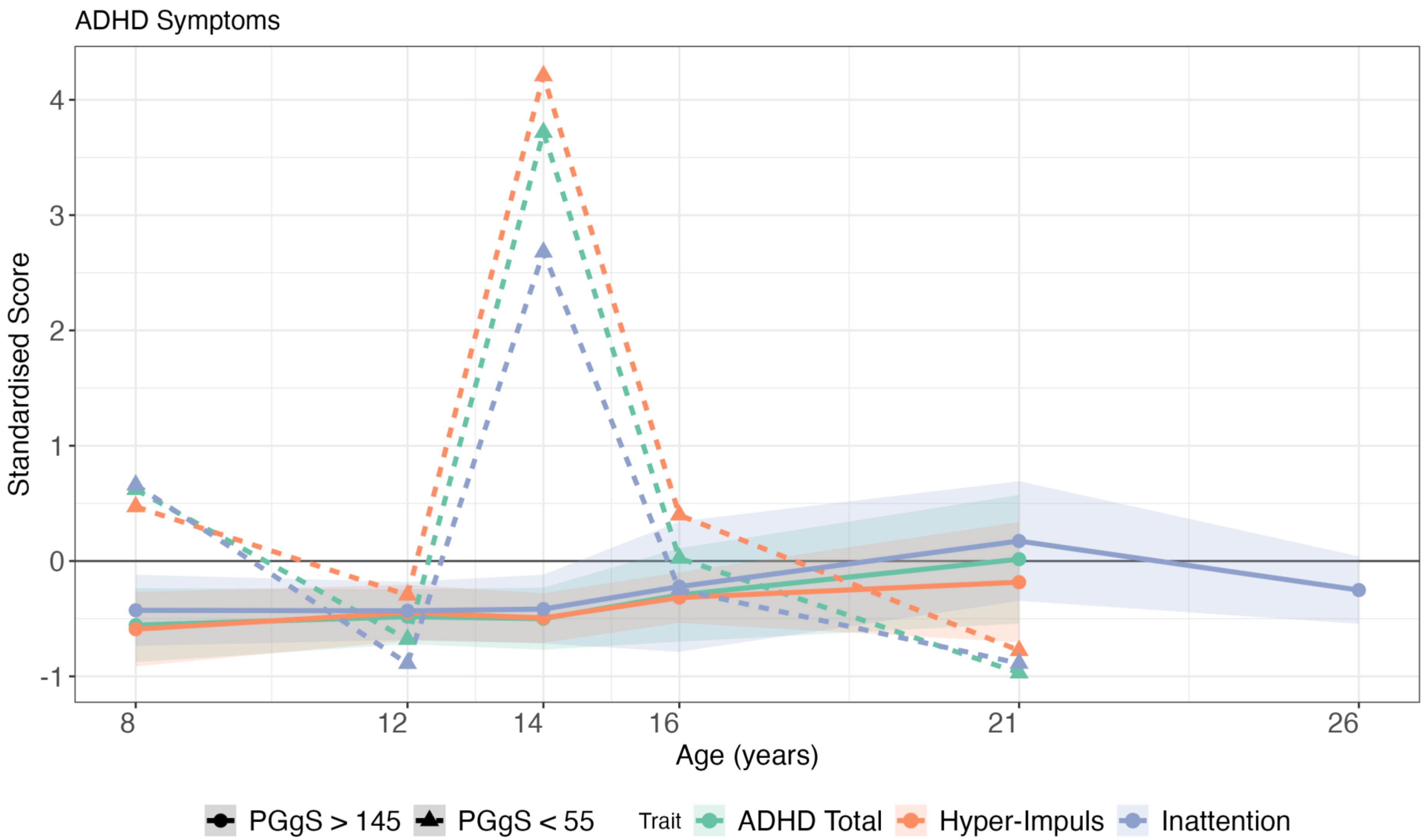
ADHD Symptoms
Standardised Score
4
3
2
1
0
-1
8
12
14
16
21
26
Age (years)
PGgS > 145
PGgS < 55
Trait
ADHD Total
Hyper-Impuls
Inattention

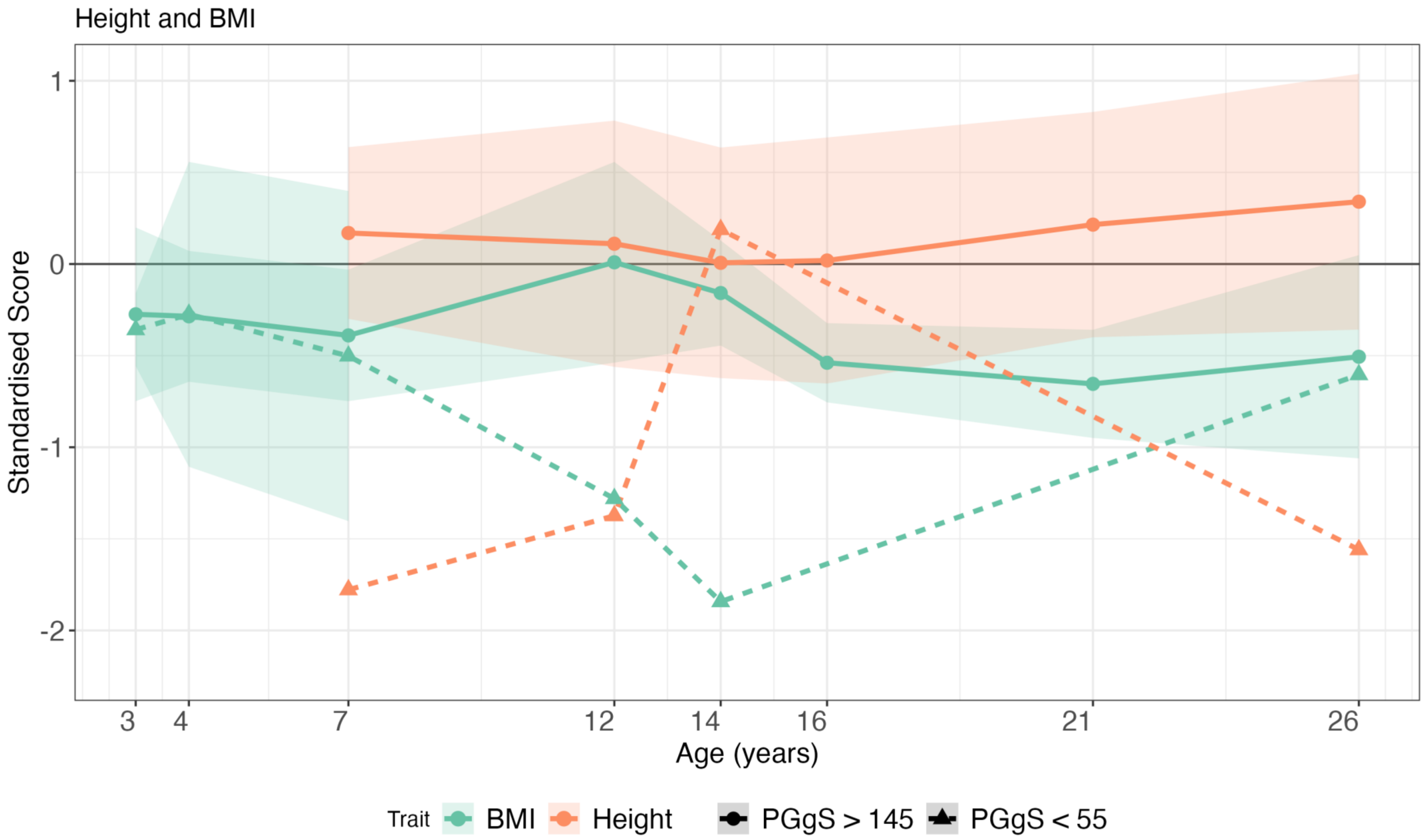

*Note*. Mean trajectories across development for individuals with polygenic g scores >145 (panels a-c) and <55 (panels d-f). Panel a/d: Anxiety-Related Behaviours Questionnaire (ARBQ) five subscales and total score. Panel b/e: Conners' Rating Scale ADHD two subscales (inattention and hyperactivity/impulsivity) and total score. Panel c/f: Height and body mass index (BMI). All measures are standardised to sample mean = 0, SD = 1. Shaded areas represent 95% confidence intervals; absence of confidence intervals indicates only one individual was measured at that age. For phenotypes with multiple raters, one rater per age is used: parent ratings before age 18 and child/self-ratings at age 18 and older. The black horizontal line at y = 0 represents the population average.
